# The molecular basis of socially-mediated phenotypic plasticity in a eusocial paper wasp

**DOI:** 10.1101/2020.07.15.203943

**Authors:** Benjamin A. Taylor, Alessandro Cini, Christopher D. R. Wyatt, Max Reuter, Seirian Sumner

## Abstract

Phenotypic plasticity, the ability to produce multiple phenotypes from a single genotype, represents an excellent model with which to examine the relationship between gene expression and phenotypes. Despite this, analyses of the molecular bases of plasticity have been limited by the challenges of linking individual phenotypes with individual-level gene expression profiles, especially in the case of complex social phenotypes. Here, we tackle this challenge by analysing the individual-level gene expression profiles of *Polistes dominula* paper wasps following the loss of a queen, a perturbation that induces some individuals to undergo a significant phenotypic shift and become replacement reproductives. Using a machine learning approach, we find a strong response of caste-associated gene expression to queen loss, wherein individuals’ expression profiles become intermediate between queen and worker states. Importantly, this change occurs even in individuals that appear phenotypically unaffected. Part of this response is explained by individual attributes, most prominently age. These results demonstrate that large changes in gene expression may occur in the absence of detectable phenotypic changes, resulting here in a socially mediated de-differentiation of individuals at the transcriptomic but not the phenotypic level. Our findings also highlight the complexity of the relationship between gene expression and phenotype, where transcriptomes are neither a direct reflection of the genotype nor a proxy for the molecular underpinnings of the external phenotype.

## Introduction

The relationship between gene expression and external phenotype is complex and unresolved. Much research in behavioural and evolutionary ecology is based on the implicit assumption that phenotypic traits can be modelled as though they directly reflect gene expression patterns, and that evolutionary trajectories can therefore be studied while remaining agnostic with regard to the underlying molecular mechanisms (Hadfield et al 2007; Rittschof & Robinson 2014; Rubin 2016). This ‘phenotypic gambit’ has proven a useful rule of thumb, permitting the establishment of a rich body of literature surrounding the evolution of complex traits, despite a lack of data relating to the genetic basis of these traits (e.g. Réale et al 2010; Chapman et al 2011; Fowler-Finn & Rodríguez 2012). In the past decade, however, advances in the affordability of ‘omic’ data and availability of powerful bioinformatic methods have greatly enhanced our ability to assess the assumptions made by the phenotypic gambit (Rittschof & Robinson 2014; Heyes 2016). The time is right to disentangle the molecular bases of complex phenotypic traits.

Phenotypic plasticity, the ability of an individual to effect phenotypic changes in response to external cues, is an ideal phenomenon with which to study the relationship between gene expression and phenotype because it involves the production of multiple phenotypes without gene sequence changes. Of particular value are species in which adult individuals can be experimentally induced to transition between distinct, measurable phenotypes. By comparing the gene expression profiles of groups of individuals that differ in their phenotypes as a result of plasticity, it is possible to isolate phenotypic effects of gene expression while controlling for genetic differences. Using this approach, significant progress has been made in unravelling the molecular underpinnings of sequential sex changes in hermaphroditic fish (Horiguchi et al 2013; Casas et al 2016; Todd et al 2019), the distinct gregarious social phenotype of desert locusts (Cullen et al 2017; Lo et al 2018), and the reproductive castes of social insects (Simola et al 2016; Libbrecht et al 2018; Rehan et al 2018; Shell & Rehan 2018). Such studies typically rely on comparisons between groups of individuals with well-differentiated phenotypes, however. As a result, little is known about more subtle effects during the transition from one morph to the other, and the relationship between expression patterns and phenotypic traits at the individual level.

The reproductive castes found in the colonies of social insects provide excellent model systems for determining the extent to which fine-scale changes in phenotype are reflected at the molecular level. With a few exceptions (reviewed in Schwander et al 2010), the distinct queen and worker phenotypes found in such colonies are plastically determined either during development or in adulthood. Workers in some species can be experimentally induced to transition to a reproductive role in response to the removal of a colony’s queen (e.g. Strassmann et al 2004; Tibbetts & Huang 2010) or as a result of exposure to varying levels of brood (e.g. Chandra et al 2018; Libbrecht et al 2018), allowing changes in the behavioural, physiological and molecular traits that define caste identity to be tracked. An additional benefit—and challenge—of studying social insect colonies is that they involve complex social structures. Such interactions can be hard to study, but offer the opportunity to assess the effects of social interactions upon phenotypes and transcriptomes.

In the present study, we explore the relationship between transcriptome and phenotype in the context of plastic caste expression in the European paper wasp *Polistes dominula* (Christ 1791), which is a model organism often used in studies of social insect behaviour (reviewed in Starks & Turillazzi 2006; Jandt et al 2014) and, more recently, for analyses of caste gene expression (e.g. Patalano et al 2015; Standage et al 2016; Geffre et al 2017; Manfredini et al 2018). In this species, removing the established queen from a single-foundress colony induces a queen succession process in which one (or very few) workers transition to a queen phenotype, with age playing a key role in predicting which individual will do so (Pardi 1948; Tsuji &Tsuji 2005): almost invariably, the new queen is one of the oldest individuals, and there is little conflict over succession (Strassmann et al 2004; Taylor et al 2020). In a recent paper (Taylor et al 2020) we followed responses to queen removal in *P. dominula* on a fine scale by measuring individual-level behavioural and physiological traits and generating a univariate measure of caste identity (‘queenness’) to describe individuals’ phenotypic profiles (**Box 1**). We found that phenotypic responses were limited to the subset of individuals that transitioned to the queen role, while other individuals exhibited little or no measurable change in measured traits. This system, in which individuals within a controlled environment vary strongly and predictably in their phenotypic response to a shared stimulus, affords an excellent opportunity to track the relationship between gene expression and phenotypic expression at the individual level.

To do so, here we complement the individual-level phenotypic data of Taylor et al. (2020) by assaying the brain gene expression profiles of 107 individuals for which fine-scale behavioural and ovarian data have been obtained, including queens and workers from stable colonies and individuals from colonies that had their queens experimentally removed. We then assess the relationship between the transcriptome and the phenotypic changes induced by queen removal. To match the univariate measure of caste identity that we applied to our phenotypic data (**Box 1**), we use a support vector classification approach to generate a univariate measure of caste-associated gene expression. Support vector classification is a powerful tool with which to transform complex patterns in multidimensional data into a continuous classification score, allowing the detection of subtle, widespread signals of differential expression between phenotypic states that are likely to be missed in conventional differential expression analyses. Support vector machines (SVMs) have become a key tool in the early identification of phenotypically indistinguishable cancer subtypes (Segal et al 2003; Abeel et al 2010; Huang et al 2018) and their value has recently been demonstrated in animal behaviour studies (e.g. Chakravarty et al 2019; Rast et al 2020). Given their sensitivity, SVMs should be ideally suited to quantify gene expression variation across the spectrum between differentiated worker and queen roles, but to our knowledge this is the first time SVMs have been used to interrogate the molecular basis of behavioural traits.

Applying an SVM approach along with standard differential expression and gene co-expression analyses, we show that brain gene expression responses to queen removal in *P. dominula* include a colony-wide response that does not match that observed at the phenotypic level. Our results indicate that gene expression in *P. dominula* colonies reflects both a generalised response to queen loss which is independent of phenotype, and a phenotype-specific response that tracks individuals’ expression of plastic phenotypic changes. To our knowledge, this study provides the most comprehensive analysis to date of the ways that plastic phenotypes are reflected at the transcriptomic level; our results expose the complexity of the relationship between individual-level gene expression and individual phenotypes and call into question the assumptions of the phenotypic gambit.

## Box 1.

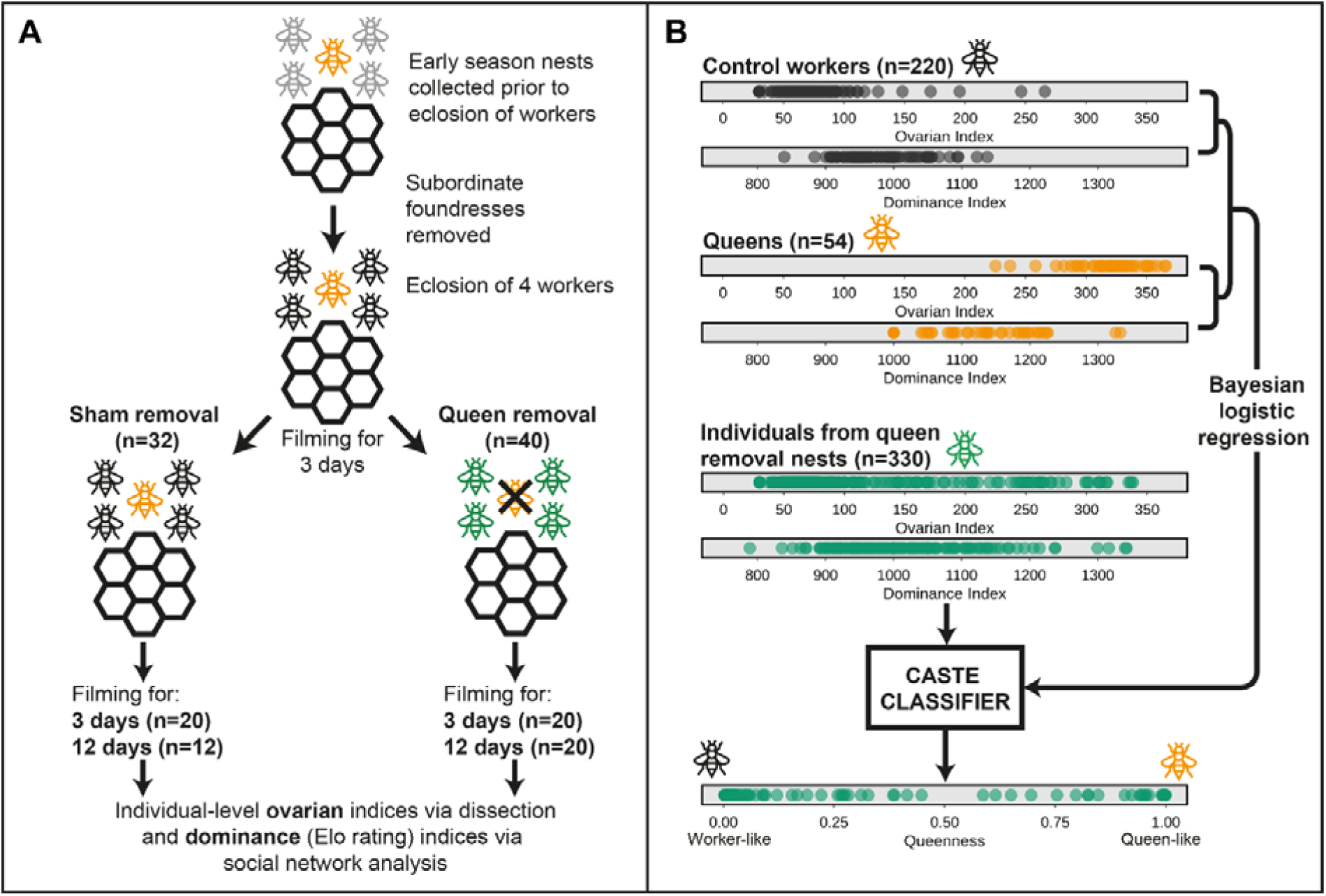

Summarized experimental design describing the generation of phenotypic data pertaining to individual-level responses to queen removal in *Polistes dominula* (Taylor et al 2020) used in this study. **(A)** Early-season nests were transferred to the lab prior to the eclosion of workers and subordinate foundresses were removed. After at least 4 workers had eclosed, nests were filmed for three days and then assigned to either control or queen removal treatments. Following treatment (sham or queen removal) nests were filmed for a further three or twelve days and then all individuals were dissected to generate ovarian development indices following Cini et al (Cini et al. 2013). Footage of nests was used to generate dominance indices in the form of Elo ratings (Elo 1978; Neumann et al. 2011). **(B)** Ovarian and dominance indices from queens and control workers were used to produce a logistic regression model for caste classification (0=worker, 1=queen). Data from individuals on queenless post-removal nests were then passed through this model to fit caste estimates and thereby identify individuals with high ‘queenness’, i.e. those that exhibited strongly queen-like phenotypes following queen removal and thus represented possible replacement queens. Of the individuals for which data are shown here, 27 queens, 12 workers from control nests, and 62 individuals from queen removal nests were subsequently sequenced to generate the data discussed in the present study.

## Results

### Support vector classification reveals consistent patterns of caste gene expression differentiation involving many genes

Our support vector approach allowed us to reduce complex variation in brain gene expression data down to a single dimension of caste identity, matching the Bayesian logistic regression model we used in Taylor et al. (2020) to condense ovarian and behavioural data into a unidimensional metric of phenotypic caste (‘queenness’; **Box 1**). We trained an SVM using gene expression data from 27 queens and 12 workers from stable, unmanipulated colonies. An initial model, based on all 10734 genes annotated in our experiment, achieved a root mean squared validation error of 0.065 in three-fold cross-validation, i.e. a model trained on a random subset of two thirds of workers and queens classified the remaining third within 25.5% of their true values (worker=0, queen=1). By applying a process of iterative feature selection we were able to remove genes that had low weights in the classification and contributed to overfitting (**Figure S1A; Table S1**). Doing so allowed us to identify an optimal model containing 1992 genes with a substantially reduced root mean squared classification error of 0.021 (**Table S2**). This model classified queens and workers very consistently, with strong separation of queens from workers (**Figure 1A**). Thirty-four gene ontology (GO) terms were significantly enriched among these 1992 genes, including a number of terms associated with translation such as *rRNA processing, tRNA aminoacylation*, and *ribosomal large subunit biogenesis* (**Figure S1B**).

**Figure 1.**
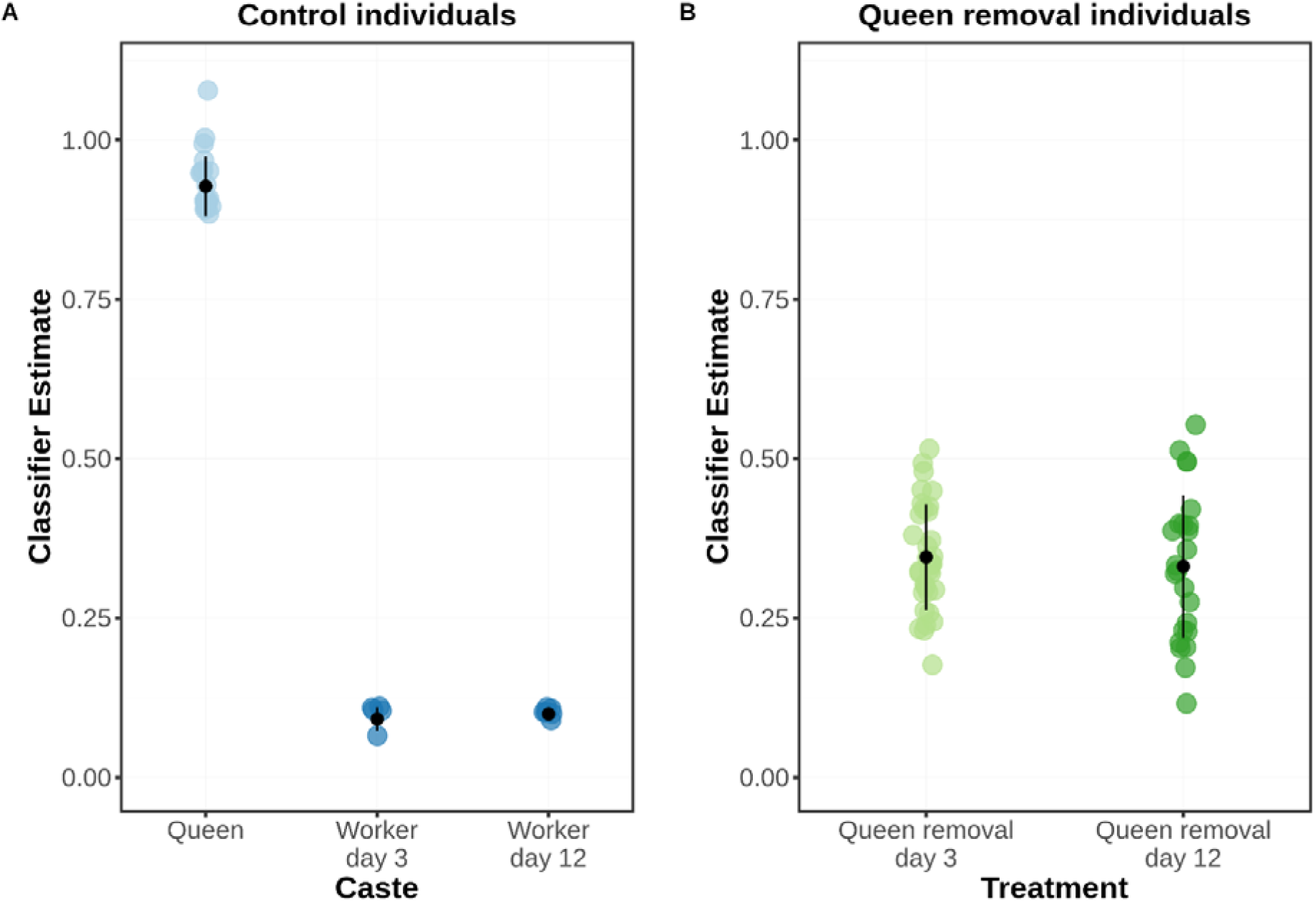
Classifier estimates generated by the lowest-error SVM using 1992 caste-informative genes for **(A)** queens and workers from control colonies and **(B)** individuals from experimental colonies following queen removal. Mean and standard deviation for each group are shown in black.

The 1992 genes in our optimised SVM represent a much larger set than that which is found to differentiate queens and workers based on a standard differential expression approach. When we applied a DESeq2 analysis with a 1.5 fold-change threshold to the same set of queens and workers used to train the SVM, we identified just 81 differentially expressed genes (with no associated GO terms), a small number that is typical for similar analyses in *Polistes* (e.g. Toth et al 2014; Patalano et al 2015; Geffre et al 2017). All but four of these genes were present in the larger SVM set (**Figure S1C**; **Table S3**), suggesting that the SVM captures almost all the information obtained from these standard methods. Conversely, an SVM trained using just these 81 differentially-expressed genes exhibited a cross-validation error rate of 0.065, no better than the original un-optimised model. Thus, the picture of caste differentiation provided by standard differential expression analysis appears to miss a great number of subtle differences in individuals’ gene expression profiles that contribute to caste differentiation.

### Colony-wide brain gene expression responses to queen removal

Following the loss of a queen from a *Polistes* colony, typically one or a few individuals undergo a phenotypic transition to become a replacement queen while the rest of the colony members remain workers (Miyano 1991; Dapporto et al 2005; Taylor et al 2020). If this response is reflected at the level of gene expression, then we would expect to observe significant changes in the transcriptional profiles of individuals that exhibit a phenotypic transition, but not in those that do not. Our individual-level expression data allowed us to track responses in 62 individuals’ gene expression profiles following queen removal across 19 nests. We analysed individuals’ gene expression profiles at three days after queen removal, when queen replacement is ongoing (n=24), and at twelve days after queen removal, when succession is largely settled at the phenotypic level (Strassmann et al 2004; Taylor et al 2020; n=38). Individuals for sequencing were selected to cover a wide range of phenotypes, including those that remained entirely worker-like, those that had transitioned to highly queen-like phenotypes, and those with intermediate phenotypes at the time of sampling (**Figure S1D-F**).

Analysing shifts in SVM estimates of individual wasps allowed us to assess the degree to which these concurred with the changes visible at the phenotypic level. Doing so, we found that the SVM estimates of individuals from post-removal nests were intermediate between those of queens and workers from control nests (**Figure 1B**), a finding which concurs with the placement of these individuals according to principal component analysis (**Figure S1G**). This result is surprising given that the majority of individuals on queen removal nests are phenotypically indistinguishable from workers on control nests in terms of ovarian development and behavioural dominance (Taylor et al 2020). Thus, the large majority of individuals on queen removal nests exhibited perturbation of their caste-associated gene expression, even though only a few of these individuals exhibited responses to queen loss at the level of physiology or behaviour and transitioned from a worker to a queen role.

Furthermore, we found no evidence that the degree of transcriptional perturbation declined over time following queen removal. SVM classification estimates of individuals from queen removal nests did not differ significantly between days three and twelve following queen removal (QR3 mean 0.331±0.111; QR12 mean 0.345±0.082; Wilcoxon W=369, *p*=0.55). The effects of queen removal on individuals’ gene expression profiles therefore appear to be both widespread and persistent, affecting all individuals in a nest and lasting beyond the point at which a new queen has already become phenotypically established.

Interestingly, the strong colony-wide perturbation following queen loss that this SVM approach identifies would have been entirely missed using a standard differential expression approach: DESeq2 identified just five genes as differentially expressed between control workers and individuals from manipulated nests, with no associated GO enrichment (**Table S4**).

### Age and queenness explain variation in individual-level molecular responses to queen loss

While SVM classification indicates that queen removal causes colony-wide perturbation to brain expression profiles, classifier estimates varied substantially between individuals following queen removal, spanning a much greater range of values (0.116-0.540) than those of queens (0.900-1.100) or workers (0.057-0.100) from queenright colonies (**Figure 1**). To better understand this variation, we examined whether the classifier estimates for individuals from manipulated colonies were predicted by those individuals’ phenotypic traits—specifically ovarian development, behavioural dominance, and age (**Box 1**; Taylor et al 2020).

Our phenotypic measure of caste identity (queenness) was a significant predictor of expression-based SVM classifier estimates when fitted using a linear model (slope±SE = 0.0414±0.0114, *p* = 6.0×10^−4^; **Figure 2A**). The individual components of queenness, ovarian development and dominance were also significant or near-significant predictors of SVM classification individually (ovarian development: slope±SE = 0.0426±0.0113, *p* = 4.0×10^−4^; dominance: slope±SE = 0.0231±0.0122, *p* = 0.06). Notably, however, because phenotypic queenness was strongly correlated with age among post-removal individuals (cor = 0.4832, p = 8.0×10^−5^; Taylor et al 2020), age was an equally strong predictor of caste estimates when fitted in a separate linear model (slope±SE = 0.0516±0.0106, *p* = 9.8×10^−6^; **Figure 2B**). Thus, the significance of queenness as a predictor of caste estimates might have been an artefact of the fact that both are correlated with age. To test whether queenness had an effect over and above that accounted for by age, we calculated the residuals of phenotypic queenness on age and fitted the caste estimates against these. These residuals were not significantly predictive of SVM classification (slope±SE = 0.0202±0.0123, *p* = 0.11; **Figure 2C**), nor were the residuals of ovarian development alone on age (slope±SE = 0.0223±0.0123, *p* = 0.07) or the residuals of dominance alone (slope±SE = 0.0114±0.0126, *p* = 0.37). Age is thus the strongest determinant both of individuals’ caste phenotypes and of their caste-associated gene expression.

**Figure 2.**
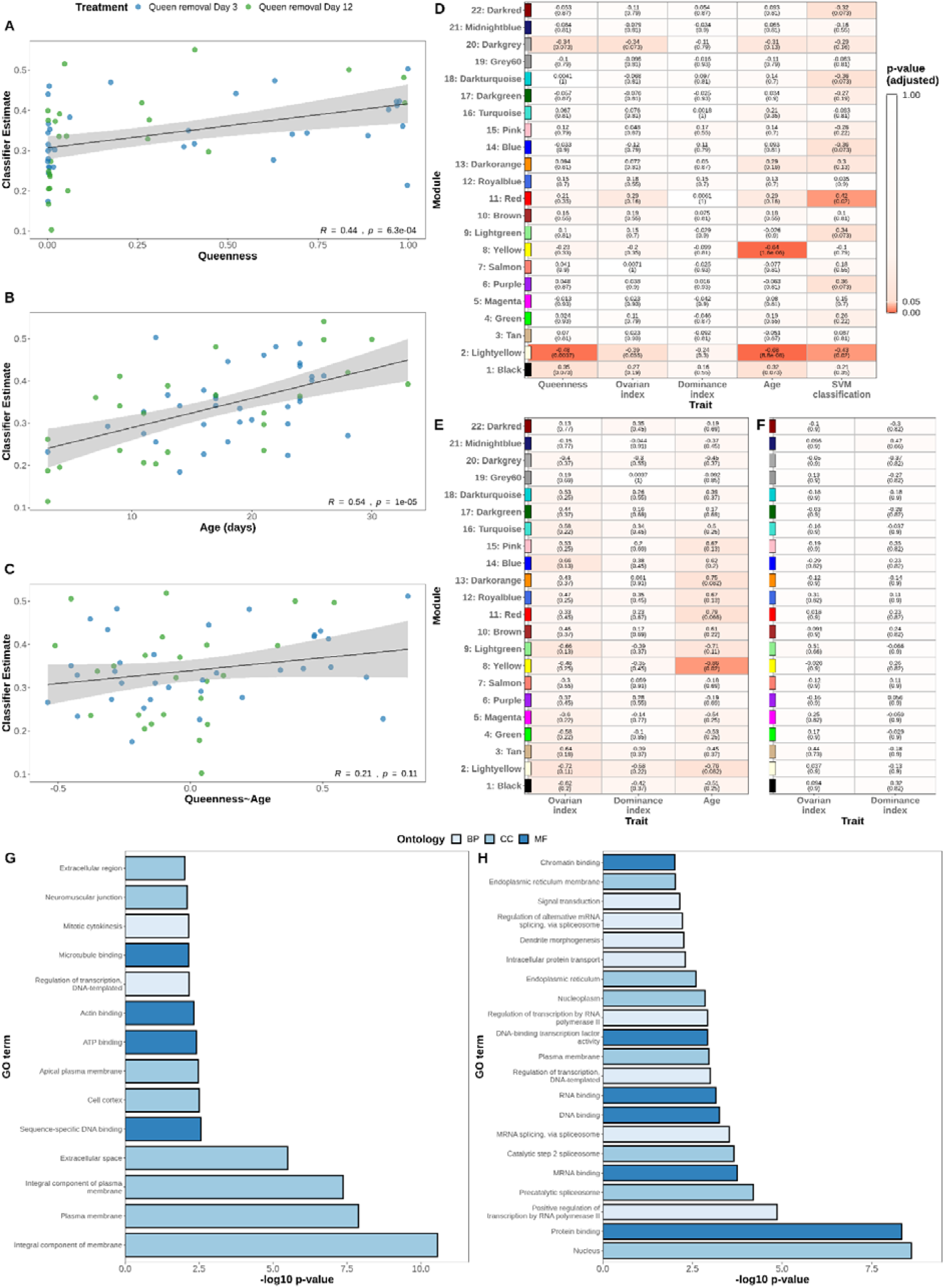
**(A-C)** Scatterplots of SVM classifier estimates for individuals from post-removal colonies plotted against **(A)** phenotypic queenness, **(B)** age, and **(C)** the residuals of queenness on age respectively. **(D-F)** Module-trait correlations for consensus modules identified by WGCNA in queen removal individuals, control workers and queens respectively. Within each cell, the Pearson correlation coefficient is given above and the FDR-adjusted p-value is given below in parentheses. **(G-H)** Enriched gene ontology terms at p<0.01 for gene module 11 and gene module 8.

In order to further discern the factors that shaped individuals’ gene expression profiles following queen removal, we performed weighted gene co-expression network analysis (WGCNA) using the full set of annotated genes. We generated 22 consensus modules across all samples, and then determined which traits significantly predicted the expression of a given module within each group (**Figure 2D-F**). In doing so, we identified three modules of particular interest.

Module 11 (*Red*) consists of 519 genes associated with 14 enriched GO terms, including a number of terms associated with molecular binding (**Figure 2G**). This module is notable in that its expression in individuals from queen removal colonies is positively associated with SVM classifications but not with phenotypic correlates of caste. A second module, Module 2 (*Lightyellow*), consists of 102 genes and appears to be associated with many caste-related traits in the queen removal condition, being negatively correlated with queenness, age, SVM classification and (more weakly) with ovarian development among queen removal individuals. We also found weak evidence that this module is negatively associated with age among workers from control nests. Unexpectedly, despite the seeming importance of Module 2 in predicting caste identity among workers, not a single GO term was enriched among the genes in the module at p<0.01. Finally, Module 8 (*Yellow*), consists of 614 genes and has a strongly negative correlation with age among both control workers and queen removal individuals. This module is associated with 21 GO terms, including a number of terms associated with molecular binding, such as protein binding, DNA binding, RNA binding and chromatin binding (**Figure 2H**). The latter term is particularly notable: regulation of chromatin accessibility is one of several epigenetic processes that have been implicated in the control of caste expression in social insects (Simola et al 2016; Wojciechowski et al 2018; Duncan et al 2020). Downregulation of this module with age might plausibly contribute to the increased phenotypic plasticity exhibited by older individuals in *P. dominula*.

## Discussion

This study provides one of the most comprehensive analyses to date regarding the transcriptomic signatures of plastic phenotypes, and the first to employ an SVM approach to interrogate the molecular basis of behaviour. We have applied this approach to identify caste-specific gene expression profiles in *P. dominula* and to explore the relationship between transcriptomic and plastic phenotypic changes following a major social perturbation in paper wasp colonies. We identify a set of nearly 2000 genes that optimally capture gene expression differences between established queens and workers. Using a caste classifier based on these genes, we find that queen removal leads to a colony-wide shift in expression, where the expression profiles of all individuals move towards a state intermediate between those of established queens and workers. Individual variation around this intermediate state is related to age and phenotypic attributes, with older (and therefore more queen-like) individuals showing expression profiles that are closer to that expected for established queens. Our results show that molecular responses to queen removal in *P. dominula* consist of both a general colony-wide response independent of phenotypic change and a response that reflects the plastic phenotypic transition.

Our study contributes to the enormous progress in our understanding of the relationship between molecular changes and changes in phenotypic expression that has been made in the past decade, facilitated by the increased availability of ‘omic’ data and complex bioinformatic analyses. Recent studies have started to challenge the view that there is a direct correspondence between transcriptomic states and external phenotypes. Libbrecht et al (2018), for example, show that gene expression responses associated with a reversible phenotypic change differ qualitatively based on the directionality of the change (from reproductive to non-reproductive or vice versa). Meanwhile, molecular manipulations have revealed a surprising degree of plasticity in canonically implastic traits such as mammalian sex (Matson et al 2011) or ant castes (Simola et al 2016). Our results go further, showing a shift in caste-specific brain gene expression profiles among individuals whose phenotypic caste expression remains otherwise apparently unchanged. This shows that the expectation of a close match between expression profiles and phenotypes is excessively simplistic, or at least that detecting such a match requires detailed knowledge of relevant genes and/or an exhaustive phenotypic characterisation. This, in turn, suggests that the use of expression data to infer the molecular basis of phenotypes is more challenging than hitherto appreciated.

A major advantage of this study is the use of individual-level gene expression data from a large number of subjects, including individuals reared in a shared social environment but exhibiting very different phenotypic responses to perturbation. By sequencing individuals rather than pools, we were able to match each gene expression profile to high-resolution phenotypic data that captures the scale of naturally-occurring variation in features such as age, ovarian development and dominance behavior. This resolution allows us to address questions that are otherwise inaccessible in gene expression analyses. For example, we have been able to show that caste identity, but not the residuals of caste identity on age, are significantly predictive of individuals’ change in transcriptomic caste identity following queen loss.

Our discovery of colony-wide responses to queen loss suggests that this social perturbation provokes a significant reaction even from individuals that have little hope of attaining the vacant reproductive role. This is a surprising finding given that *P. dominula* is thought to express a ‘conventional’ gerontocratic mechanism of dominance and queen succession that mitigates the need for costly intragroup conflicts over the identity of the replacement queen (Pardi 1948; Tsuji & Tsuji 2005; Monnin et al 2009), which should greatly reduce the need for young, low-ranking workers to respond to queen loss (Taylor et al 2020). The gene expression responses of lower-ranked workers to queen removal might plausibly represent a form of safeguard against queen loss: if queen loss sometimes occurs multiple times in quick succession or is frequently associated with a general decimation of the nest population (i.e. through predation), there might be kin-selected benefits of a colony-wide ‘de-differentiation’ of individuals that facilitates a quicker succession process.

Replacement queens in our colonies did not have access to males and therefore remained unmated even after queen succession. This may partially explain the fact that even individuals with fully developed ovaries and very high dominance ratings did not transition to a fully queen-like gene expression profile, as mating can induce significant gene expression changes in insects (e.g. Gomulski et al 2012; Zhou et al 2014). The lack of immediate mating opportunities for new queens in our experiment is not necessarily unrealistic, however: unmated Hymenopteran females can lay unfertilized eggs, which develop as males. Moreover, in naturally-occurring early *P. dominula* nests, replacement queens may be established a month or more before they are mated (Strassmann et al 2004), presumably due to a scarcity of early males. The unmated replacement queens analysed here are therefore representative of those that would be present on wild nests shortly after queen loss.

To our knowledge, this study represents the first application of a support vector classification approach to behavior-associated transcriptomic data. Using this approach we identified a large group of genes as differing meaningfully between *Polistes* castes—over 10% of annotated genes, a much larger set than those found using standard analytic approaches in this study and others (e.g. Ferreira et al 2013; Toth et al 2014; Patalano et al 2015; Geffre et al 2017). Standard approaches using packages such as edgeR (Robinson et al 2010), DESeq2 (Love et al 2014), or NOISeq (Tarazona et al 2011), which typically include information-sharing between genes and relatively strict fold change cutoffs, allow differential expression to be assessed with a high degree of confidence at the level of individual genes. However, because such approaches assess genes individually and do not reduce the dimensionality of the samples’ transcriptional profiles, they are not well-suited to the tracking of subtle but consistent broad-scale changes. This is especially true for gene expression data that are noisy and heterogenous, or those that are dominated by many genes of small effect (Huang et al 2018). The molecular bases of caste in simple social insect societies represent such a class of data.

Using a machine learning approach, here we have undertaken a detailed analysis of the relationship between gene expression and socially-mediated phenotypic plasticity, revealing broad-scale changes in caste-associated gene expression profiles following a major social disruption. Our results reveal a hitherto unrecognized capacity for large scale disruption to caste-biased gene expression profiles even in the absence of apparent changes in caste phenotype, a disconnect that undermines simplistic models of the relationship between transcriptome and phenotype. Future studies should continue to marry detailed phenotypic and gene expression data in order to assess the prevalence and provenance of such discontinuities.

## Supporting information

Supplementary Phenotypic Data

## Acknowledgements

We would like to thank F. Cappa, I. Pepiciello, L. Dapporto and others at the Dipartimento di Biologia, Università di Firenze for assistance with field work, and E. Favreau and M. Bentley for their assistance with analyses. Special thanks to R. Cervo, without whose generous support this study would not have been possible. This work was supported by the Natural Environment Research Council (grant code NE/L002485/1).

## Author contributions

BAT, SS and MR conceived of and designed the experiment. BAT and AC carried out the initial experiment. BAT performed behavioural analyses, dissections and RNA extractions, and wrote the initial draft of the manuscript. BAT conducted the analyses with advice from CDRW, MR and SS. BAT and CDRW designed the figures. All authors read and contributed to the manuscript.

## Data availability

Sequencing data associated with this paper have been deposited in the NCBI Gene Expression Omnibus (www.ncbi.nlm.nih.gov/geo) under accession number GSE153532 with reviewer access token *adctccsahjgjbqt*.

## Supplementary methods

### Gene expression quantification

Brain tissue was extracted from the heads of individual samples and RNA was extracted using the RNeasy Mini Kit (Qiagen) according to manufacturer’s instructions. Library preparation was performed by Novogene Co. followed by sequencing on an Illumina HiSeq 2000 platform with 150-base pair paired-end reads. Reads were filtered with SortMeRNA (Kopylova et al 2012) using default options to remove ribosomal sequences. Trimmomatic (Bolger et al 2014) was then used to perform quality trimming. First, we trimmed adapter sequences and leading and trailing bases with low phred scores (<3). We then used the MAXINFO option with target length 36 and strictness 0.7 to trim low-quality sequences from the remaining reads. Reads were next mapped to 11313 transcripts from the *P. dominula* genome annotation 1.0 (Standage et al. 2016) using STAR (Dobin et al 2013) with default options. All 101 samples produced >85% uniquely mapped reads. Reads were assembled into transcripts using StringTie2 (Kovaka et al 2019) before being passed on as raw counts to downstream analyses. Prior to downstream analysis, transcripts were filtered to remove any gene which did not have >20 counts across all samples in at least one of the experimental groups (queens, control workers, and day three and day twelve post-manipulation workers). Following this filtering, 10734/11313 (94.9%) of transcripts remained.

## Support vector machine classification

Support vector classification was performed in R using the e1071 package (Meyer et al 2017) using the gene expression profiles for all available queens (coded with a value of 1; n=26) and control workers (coded with a value of 0; n=12). Classifiers were assessed via their 3-fold cross-validation error rates; classifiers with lower classification error were considered to be superior. Initially, we tested classifiers using a variety of kernel functions (radial, linear, sigmoid and polynomial) combined with grid searches across a wide range of kernel parameters. A radial kernel with γ=10^−6^ and cost parameter C=2^5^ was found to produce the lowest error rate of all combinations, so all subsequent classifiers were fit using this kernel and a more focused grid search in a parameter space of 2^4^<C<2^6^ and 10^−5^<γ<10^−7^ in order to minimise the processing power necessary to perform feature selection. For feature selection, we took the classifier fitted with all genes, and iteratively performed the following process: (1) The 3-fold cross-validation error of the model was calculated twenty times using randomly-assigned bins, and the mean of the resulting errors was recorded as the true validation error of the classifier; (2) The feature weights of all genes were calculated by taking the matrix product of that classifier’s coefficients with its support vectors; (3) The gene with the smallest absolute weight in the model was dropped; (4) A new classifier was calculated using the remaining set of genes. This process was repeated until just 100 genes remained, and the optimal support vector classifier was then taken as that for which the cross-validation error reached its minimum.

## Differential expression analysis

Differential expression analyses were performed in R using the DESeq2 package (Love et al 2014). DESeq2 was run on all groups and contrasts were then calculated for each pair of groups. Differential expression was calculated relative to a baseline fold change of 1.5, i.e. *p*-values refer to the probability that absolute change between two groups was greater than 50%. Genes were considered differentially expressed between conditions if *p*<0.05 after false discovery rate correction according to the Benjamini-Hochberg procedure.

## Gene co-expression network analysis

Weighted gene co-expression network analysis was performed in R using the WGCNA package (Langfelder & Horvath 2008). As WGCNA is particularly sensitive to genes with low expression, data were first subjected to a second round of filtering in which genes that had <10 reads in >90% of samples were removed. Consensus gene modules across all samples were then constructed using a soft-threshold power of 9. Initially, 26 gene modules were identified. Modules whose eigengene correlation was >0.75 were subsequently merged, after which 22 consensus modules remained. Finally, the Pearson correlation of each module with each phenotypic trait within each group (queens, control workers and individuals from queen removal nests) was calculated and subjected to Benjamini-Hochberg FDR correction.

## Gene ontology (GO) enrichment analysis

In order to perform GO enrichment analysis, we first used OrthoFinder (Emms & Kelly 2019) to identify orthologues for each *P. dominula* gene in *D. melanogaster*, a model species for which GO annotations are much more complete. GO annotations for each *D. melanogaster* gene were acquired from BioMart (Smedley et al 2015) and each *P. dominula* gene was then assigned GO terms permissively, i.e. a given *P. dominula* gene was assigned a GO term if that term appeared as an annotation to any of its orthologues. GO enrichment analysis was then performed in R via the topGO package (Alexa & Rahnenfuhrer 2009) using TopGO’s weight01 algorithm and Fisher’s exact test to identify GO terms that were significantly overrepresented (*p*<0.01) in a focal set of genes against a background consisting of all genes that appeared in the relevant analysis.

## Notes

https://www.ncbi.nlm.nih.gov/geo/query/acc.cgi?acc=GSE153532

### Competing Interest Statement

The authors have declared no competing interest.

