## Supplementary Table S1 for "The molecular basis of socially-mediated phenotypic plasticity in a eusocial paper wasp"

**Table S1.** Genes removed following each feature selection iteration of the the support vector machine. **RSMCVerror:** The root mean squared 3-fold cross-validation error of the model that resulted following the removal of each given gene. **Weight:** The weight of each feature in the original, unoptimised SVM. Weights were calculated as the matrix product of the SVM's coefficients and its support vector values. **Optimal:** Whether the gene appears in the final, optimised caste-classifying SVM. **DE:** Whether the gene was identified by DESeq2 as differentially expressed between queens and workers.

| GeneID | ProteinID | RSMCVerror | Weight | Optimal | DE |
|---|---|---|---|---|---|
| LOC107064170 | E3 ubiquitin-protein ligase TRIM9 | 0.05328 | 0.00909 | FALSE | FALSE |
| LOC107068113 | prolyl 3-hydroxylase OGFOD1 | 0.05263 | 0.01269 | FALSE | FALSE |
| LOC107070651 | RING finger protein 157 | 0.05588 | 0.01977 | FALSE | FALSE |
| LOC107068744 | uncharacterized protein LOC107068744 | 0.05536 | 0.04059 | FALSE | FALSE |
| LOC107073484 | uncharacterized protein LOC107073484 | 0.05471 | 0.04085 | FALSE | FALSE |
| LOC107067867 | zinc finger protein-like 1 | 0.05314 | 0.04653 | FALSE | FALSE |
| LOC107066601 | uncharacterized protein LOC107066601 | 0.05592 | 0.04744 | FALSE | FALSE |
| LOC107065109 | neuroguidin | 0.05369 | 0.05077 | FALSE | FALSE |
| LOC107072370 | carbonyl reductase [NADPH | 0.0534 | 0.0538 | FALSE | FALSE |
| LOC107068889 | G-box-binding factor-like | 0.05452 | 0.05543 | FALSE | FALSE |
| LOC107073639 | uncharacterized protein LOC107073639 | 0.05418 | 0.06643 | FALSE | FALSE |
| LOC107068612 | protein AF-9 | 0.05599 | 0.07398 | FALSE | FALSE |
| LOC107065665 | dynactin subunit 5 | 0.05303 | 0.07866 | FALSE | FALSE |
| LOC107069087 | programmed cell death 6-interacting protei | 0.05491 | 0.08369 | FALSE | FALSE |
| LOC107069065 | uncharacterized protein LOC107069065 | 0.05379 | 0.08702 | FALSE | FALSE |
| LOC107067233 | DNA ligase 4-like | 0.05441 | 0.09185 | FALSE | FALSE |
| LOC107073014 | cytochrome c oxidase subunit 5B, mitochon | 0.05664 | 0.10013 | FALSE | FALSE |
| LOC107064642 | mini-chromosome maintenance complex-bi | 0.05502 | 0.10464 | FALSE | FALSE |
| LOC107066191 | TBC1 domain family member 10A | 0.05742 | 0.11131 | FALSE | FALSE |
| LOC107069765 | cystinosin homolog | 0.05441 | 0.12533 | FALSE | FALSE |
| LOC107063994 | solute carrier family 25 member 40 | 0.05389 | 0.12892 | FALSE | FALSE |
| LOC107071981 | solute carrier family 41 member 1-like | 0.05468 | 0.15352 | FALSE | FALSE |
| LOC107065428 | uncharacterized protein C1orf158 homolog | 0.05352 | 0.1592 | FALSE | FALSE |
| LOC107065448 | cysteine protease ATG4B | 0.05625 | 0.16857 | FALSE | FALSE |
| LOC107070840 | la-related protein 7 | 0.05563 | 0.17126 | FALSE | FALSE |
| LOC107069757 | uncharacterized protein LOC107069757 | 0.05506 | 0.17265 | FALSE | FALSE |
| LOC107067679 | DNA-directed RNA polymerase III subunit RI | 0.05412 | 0.17406 | FALSE | FALSE |
| LOC107064050 | dolichyl-diphosphooligosaccharide--protein | 0.05299 | 0.17886 | FALSE | FALSE |
| LOC107068153 | phosphatidylinositol 5-phosphate 4-kinase t | 0.05447 | 0.19433 | FALSE | FALSE |
| LOC107072601 | lysM and putative peptidoglycan-binding dc | 0.05606 | 0.2172 | FALSE | FALSE |
| LOC107072500 | U8-agatoxin-Ao1a-like | 0.05478 | 0.2403 | FALSE | FALSE |
| LOC107068455 | glucose dehydrogenase [FAD, quinone | 0.05607 | 0.24589 | FALSE | FALSE |
| LOC107071275 | meteorin-like protein | 0.05311 | 0.25714 | FALSE | FALSE |
| LOC107071954 | talin-1 | 0.05283 | 0.2797 | FALSE | FALSE |
| LOC107067720 | 28S ribosomal protein S10, mitochondrial | 0.05495 | 0.28303 | FALSE | FALSE |
| LOC107069269 | CB1 cannabinoid receptor-interacting prote | 0.05516 | 0.31056 | FALSE | FALSE |
| LOC107072742 | vascular endothelial growth factor receptor | 0.05237 | 0.33502 | FALSE | FALSE |
| LOC107070023 | calcium/calmodulin-dependent 3',5'-cyclic r | 0.0548 | 0.34494 | FALSE | FALSE |
| LOC107067371 | putative fatty acyl-CoA reductase CG5065 | 0.05519 | 0.36793 | FALSE | FALSE |
| LOC107068322 | cyclin-G-associated kinase | 0.05365 | 0.37357 | FALSE | FALSE |
| LOC107065786 | uncharacterized protein LOC107065786 | 0.05339 | 0.38537 | FALSE | FALSE |

| | | | | | |
|---|---|---|---|---|---|
| LOC107067791 | T-complex protein 1 subunit theta | 0.05584 | 0.38689 | FALSE | FALSE |
| LOC107064640 | uncharacterized protein LOC107064640 | 0.05498 | 0.40989 | FALSE | FALSE |
| LOC107066470 | REST corepressor 3 | 0.05349 | 0.41076 | FALSE | FALSE |
| LOC107074365 | DNA-directed RNA polymerase III subunit RI | 0.05174 | 0.41197 | FALSE | FALSE |
| LOC107067534 | leucine-rich PPR motif-containing protein, m | 0.05341 | 0.42848 | FALSE | FALSE |
| LOC107065062 | dipeptidase 1-like | 0.05383 | 0.43398 | FALSE | FALSE |
| LOC107063918 | transcription factor HIVEP3 | 0.0548 | 0.44153 | FALSE | FALSE |
| LOC107070921 | collagen alpha-1(I) chain-like | 0.05415 | 0.4446 | FALSE | FALSE |
| LOC107065168 | prisilkin-39-like | 0.05308 | 0.45603 | FALSE | FALSE |
| LOC107071233 | suppressor protein SRP40-like | 0.055 | 0.46949 | FALSE | FALSE |
| LOC107067011 | tachykinin-like peptides receptor 99D | 0.05494 | 0.47004 | FALSE | FALSE |
| LOC107069463 | iron/zinc purple acid phosphatase-like prote | 0.05341 | 0.47786 | FALSE | FALSE |
| LOC107073406 | uncharacterized protein C19orf52 homolog | 0.05593 | 0.48335 | FALSE | FALSE |
| LOC107067655 | roquin-1 | 0.05581 | 0.48658 | FALSE | FALSE |
| LOC107066461 | ATP-dependent RNA helicase DDX24 | 0.0542 | 0.49285 | FALSE | FALSE |
| LOC107066614 | metal transporter CNNM4 | 0.05335 | 0.50162 | FALSE | FALSE |
| LOC107072022 | uncharacterized protein LOC107072022 | 0.05351 | 0.50407 | FALSE | FALSE |
| LOC107063823 | cysteine and glycine-rich protein 2-like | 0.05405 | 0.52792 | FALSE | FALSE |
| LOC107073263 | uncharacterized protein LOC107073263 | 0.05481 | 0.53396 | FALSE | FALSE |
| LOC107066114 | transcription initiation factor TFIID subunit 5 | 0.0532 | 0.53506 | FALSE | FALSE |
| LOC107071259 | zinc finger protein 148-like | 0.05361 | 0.53796 | FALSE | FALSE |
| LOC107071944 | uncharacterized protein LOC107071944 | 0.05541 | 0.54092 | FALSE | FALSE |
| LOC107072396 | protein RRNAD1-like | 0.05515 | 0.54834 | FALSE | FALSE |
| LOC107067855 | neural-cadherin | 0.05336 | 0.54981 | FALSE | FALSE |
| LOC107068325 | kelch-like ECH-associated protein 1 | 0.05408 | 0.54997 | FALSE | FALSE |
| LOC107071579 | mitoferrin-1 | 0.05401 | 0.55062 | FALSE | FALSE |
| LOC107070683 | uncharacterized protein LOC107070683 | 0.05594 | 0.55252 | FALSE | FALSE |
| LOC107070264 | large proline-rich protein BAG6-like | 0.05463 | 0.57281 | FALSE | FALSE |
| LOC107064436 | RUN and FYVE domain-containing protein 2 | 0.05534 | 0.57407 | FALSE | FALSE |
| LOC107073016 | uncharacterized protein LOC107073016 | 0.05478 | 0.57491 | FALSE | FALSE |
| LOC107066243 | PH-interacting protein | 0.05482 | 0.58523 | FALSE | FALSE |
| LOC107071328 | TBC1 domain family member 24 | 0.05211 | 0.58853 | FALSE | FALSE |
| LOC107068692 | MAM and LDL-receptor class A domain-cont | 0.05276 | 0.5946 | FALSE | FALSE |
| LOC107073885 | uncharacterized protein LOC107073885 | 0.05332 | 0.6016 | FALSE | FALSE |
| LOC107068872 | uncharacterized protein LOC107068872 | 0.0532 | 0.60522 | FALSE | FALSE |
| LOC107068006 | uncharacterized protein LOC107068006 | 0.05429 | 0.60998 | FALSE | FALSE |
| LOC107069157 | very-long-chain 3-oxoacyl-CoA reductase-lik | 0.05562 | 0.63267 | FALSE | FALSE |
| LOC107073004 | protein PAT1 homolog 1-like | 0.0535 | 0.63673 | FALSE | FALSE |
| LOC107069999 | maestro heat-like repeat-containing protein | 0.055 | 0.65327 | FALSE | FALSE |
| LOC107065553 | ion transport peptide | 0.05236 | 0.66233 | FALSE | FALSE |
| LOC107063730 | probable sodium/potassium/calcium exchar | 0.05496 | 0.6624 | FALSE | FALSE |
| LOC107074519 | fez family zinc finger protein 1-like | 0.05437 | 0.66362 | FALSE | FALSE |
| LOC107064344 | facilitated trehalose transporter Tret1-2 hor | 0.05448 | 0.6784 | FALSE | FALSE |
| LOC107072907 | retinoid-inducible serine carboxypeptidase- | 0.05314 | 0.68104 | FALSE | FALSE |
| LOC107069348 | uncharacterized protein LOC107069348 | 0.05536 | 0.69358 | FALSE | FALSE |
| LOC107073130 | coiled-coil domain-containing protein 39-lik | 0.05468 | 0.6996 | FALSE | FALSE |
| LOC107071657 | uncharacterized protein LOC107071657 | 0.05464 | 0.70785 | FALSE | FALSE |
| LOC107064143 | 39S ribosomal protein L28, mitochondrial | 0.05458 | 0.71268 | FALSE | FALSE |
| LOC107069213 | uncharacterized protein LOC107069213 | 0.05517 | 0.71537 | FALSE | FALSE |
| LOC107066454 | uncharacterized protein LOC107066454 | 0.05469 | 0.72422 | FALSE | FALSE |

| | | | | | |
|---|---|---|---|---|---|
| LOC107065142 | tyrosine kinase receptor Cad96Ca | 0.05513 | 0.72495 | FALSE | FALSE |
| LOC107069522 | Bardet-Biedl syndrome 1 protein | 0.05401 | 0.72708 | FALSE | FALSE |
| LOC107070060 | protein LLP homolog | 0.05424 | 0.73323 | FALSE | FALSE |
| LOC107074377 | 26S proteasome non-ATPase regulatory sub | 0.05174 | 0.73479 | FALSE | FALSE |
| LOC107068727 | PR domain zinc finger protein 16-like | 0.05323 | 0.73695 | FALSE | FALSE |
| LOC107073802 | sorting nexin-6 | 0.05377 | 0.77426 | FALSE | FALSE |
| LOC107066502 | ABC transporter G family member 20-like | 0.0541 | 0.77577 | FALSE | FALSE |
| LOC107074554 | uncharacterized protein LOC107074554 | 0.05363 | 0.77711 | FALSE | FALSE |
| LOC107070313 | myosin-IA | 0.05238 | 0.77757 | FALSE | FALSE |
| LOC107074678 | protein lozenge-like | 0.05312 | 0.78217 | FALSE | FALSE |
| LOC107065467 | E3 ubiquitin-protein ligase TM129 | 0.05242 | 0.78351 | FALSE | FALSE |
| LOC107068492 | cytochrome P450 6A1-like | 0.05259 | 0.79158 | FALSE | FALSE |
| LOC107073829 | fas-associated death domain protein | 0.0536 | 0.80703 | FALSE | FALSE |
| LOC107074256 | extracellular serine/threonine protein CG31 | 0.0541 | 0.81335 | FALSE | FALSE |
| LOC107067012 | keratin-associated protein 19-2-like | 0.05347 | 0.81586 | FALSE | FALSE |
| LOC107065045 | amyloid beta A4 precursor protein-binding f | 0.05374 | 0.83165 | FALSE | FALSE |
| LOC107067874 | heat shock 70 kDa protein cognate 3 | 0.0546 | 0.83171 | FALSE | FALSE |
| LOC107073358 | mediator of RNA polymerase II transcriptior | 0.05426 | 0.8411 | FALSE | FALSE |
| LOC107069143 | ubiquitin carboxyl-terminal hydrolase 46 | 0.05314 | 0.84753 | FALSE | FALSE |
| LOC107069815 | uncharacterized protein LOC107069815 | 0.05522 | 0.86383 | FALSE | FALSE |
| LOC107065410 | fibroblast growth factor 17-like | 0.05456 | 0.86571 | FALSE | FALSE |
| LOC107066174 | COMM domain-containing protein 5-like | 0.05545 | 0.87392 | FALSE | FALSE |
| LOC107070663 | LOW QUALITY PROTEIN: neuralized-like pro | 0.05279 | 0.87698 | FALSE | FALSE |
| LOC107073120 | receptor-binding cancer antigen expressed ( | 0.05249 | 0.87785 | FALSE | FALSE |
| LOC107071562 | carnitine O-acetyltransferase-like | 0.05445 | 0.88514 | FALSE | FALSE |
| LOC107067386 | heterogeneous nuclear ribonucleoprotein K | 0.05326 | 0.88535 | FALSE | FALSE |
| LOC107065889 | bystin | 0.05559 | 0.89602 | FALSE | FALSE |
| LOC107074046 | small proline-rich protein 2H-like | 0.05523 | 0.89655 | FALSE | FALSE |
| LOC107069854 | myelin expression factor 2 | 0.05222 | 0.89685 | FALSE | FALSE |
| LOC107071909 | LOW QUALITY PROTEIN: armadillo repeat-co | 0.05493 | 0.90836 | FALSE | FALSE |
| LOC107066913 | SH3 domain-containing kinase-binding prote | 0.05265 | 0.92422 | FALSE | FALSE |
| LOC107068002 | uncharacterized protein LOC107068002 | 0.05386 | 0.92896 | FALSE | FALSE |
| LOC107071746 | uncharacterized protein LOC107071746 | 0.05333 | 0.9312 | FALSE | FALSE |
| LOC107068026 | COP9 signalosome complex subunit 4 | 0.05393 | 0.93594 | FALSE | FALSE |
| LOC107073238 | uncharacterized protein LOC107073238 | 0.05348 | 0.94041 | FALSE | FALSE |
| LOC107071955 | spermidine synthase | 0.05359 | 0.96716 | FALSE | FALSE |
| LOC107069326 | uncharacterized protein LOC107069326 | 0.05249 | 0.96781 | FALSE | FALSE |
| LOC107070118 | nucleoporin Nup43 | 0.05455 | 0.99835 | FALSE | FALSE |
| LOC107069126 | CD109 antigen-like | 0.05237 | 1.00459 | FALSE | FALSE |
| LOC107074000 | aristaless-related homeobox protein-like | 0.05293 | 1.00831 | FALSE | FALSE |
| LOC107069121 | vesicle transport protein SFT2C | 0.05385 | 1.01416 | FALSE | FALSE |
| LOC107065068 | cytochrome P450 4C1-like | 0.05315 | 1.01971 | FALSE | FALSE |
| LOC107073575 | uncharacterized protein LOC107073575 | 0.05338 | 1.02649 | FALSE | FALSE |
| LOC107072634 | mitochondrial amidoxime-reducing compon | 0.05784 | 1.04396 | FALSE | FALSE |
| LOC107070748 | ATP-binding cassette sub-family G member | 0.05477 | 1.05054 | FALSE | FALSE |
| LOC107074478 | exportin-6-A | 0.05352 | 1.05333 | FALSE | FALSE |
| LOC107066234 | UPF0505 protein C16orf62 homolog | 0.05328 | 1.05558 | FALSE | FALSE |
| LOC107068484 | calcium/calmodulin-dependent protein kina | 0.05431 | 1.06087 | FALSE | FALSE |
| LOC107072623 | cullin-4A-like | 0.05271 | 1.09452 | FALSE | FALSE |
| LOC107069356 | rho guanine nucleotide exchange factor 17- | 0.05478 | 1.0962 | FALSE | FALSE |

| | | | | | |
|---|---|---|---|---|---|
| LOC107072920 | guanosine-3',5'-bis(diphosphate) 3'-pyropho | 0.05409 | 1.09708 | FALSE | FALSE |
| LOC107068874 | uncharacterized protein LOC107068874 | 0.05227 | 1.1214 | FALSE | FALSE |
| LOC107069970 | GPI mannosyltransferase 3 | 0.05322 | 1.12341 | FALSE | FALSE |
| LOC107069825 | mucin-17-like | 0.05461 | 1.13126 | FALSE | FALSE |
| LOC107064273 | zinc finger matrin-type protein CG9776 | 0.05376 | 1.13675 | FALSE | FALSE |
| LOC107073041 | histone-lysine N-methyltransferase pr-set7 | 0.05309 | 1.14139 | FALSE | FALSE |
| LOC107070232 | cell division cycle protein 27 homolog | 0.05265 | 1.15359 | FALSE | FALSE |
| LOC107071155 | anoctamin-1 | 0.05353 | 1.15974 | FALSE | FALSE |
| LOC107074529 | uncharacterized protein LOC107074529 | 0.05473 | 1.16709 | FALSE | FALSE |
| LOC107063648 | nuclear pore complex protein Nup98-Nup96 | 0.05209 | 1.1995 | FALSE | FALSE |
| LOC107072450 | uncharacterized protein LOC107072450 | 0.05319 | 1.22083 | FALSE | FALSE |
| LOC107069724 | GTP-binding nuclear protein Ran | 0.05425 | 1.22564 | FALSE | FALSE |
| LOC107066009 | transmembrane protein 170A | 0.05199 | 1.22968 | FALSE | FALSE |
| LOC107072199 | mRNA export factor | 0.05384 | 1.23508 | FALSE | FALSE |
| LOC107064939 | 60S ribosomal protein L37a | 0.0537 | 1.24603 | FALSE | FALSE |
| LOC107064881 | zinc transporter ZIP11 | 0.05407 | 1.25659 | FALSE | FALSE |
| LOC107071376 | uncharacterized protein LOC107071376 | 0.05557 | 1.26647 | FALSE | FALSE |
| LOC107073293 | apyrase | 0.05278 | 1.27229 | FALSE | FALSE |
| LOC107074275 | hexokinase-1-like | 0.05187 | 1.27739 | FALSE | FALSE |
| LOC107064102 | serine/threonine-protein kinase SIK2-like | 0.05343 | 1.27922 | FALSE | FALSE |
| LOC107064282 | peptidyl-prolyl cis-trans isomerase E | 0.0541 | 1.28647 | FALSE | FALSE |
| LOC107068786 | uncharacterized protein C16orf52 homolog | 0.05387 | 1.29023 | FALSE | FALSE |
| LOC107069467 | exportin-5 | 0.05542 | 1.29126 | FALSE | FALSE |
| LOC107074590 | protein lethal(2)denticleless | 0.0525 | 1.30438 | FALSE | FALSE |
| LOC107072374 | tuftelin-interacting protein 11 | 0.05375 | 1.30475 | FALSE | FALSE |
| LOC107064366 | uncharacterized protein LOC107064366 | 0.05309 | 1.30892 | FALSE | FALSE |
| LOC107072053 | ribosomal RNA small subunit methyltransfe | 0.05638 | 1.31005 | FALSE | FALSE |
| LOC107071003 | beta-arrestin-1-like | 0.05177 | 1.31947 | FALSE | FALSE |
| LOC107070544 | RNA-binding protein 25 | 0.05351 | 1.32911 | FALSE | FALSE |
| LOC107070211 | kelch domain-containing protein 4 | 0.05311 | 1.33548 | FALSE | FALSE |
| LOC107073470 | probable serine/threonine-protein kinase n | 0.0566 | 1.34752 | FALSE | FALSE |
| LOC107064324 | protein daughterless | 0.05455 | 1.35771 | FALSE | FALSE |
| LOC107072895 | 39S ribosomal protein L38, mitochondrial | 0.05189 | 1.36581 | FALSE | FALSE |
| LOC107070283 | cyclin-dependent kinase 1 | 0.05114 | 1.37421 | FALSE | FALSE |
| LOC107070954 | BUB3-interacting and GLEBS motif-containir | 0.05397 | 1.39077 | FALSE | FALSE |
| LOC107068983 | uncharacterized protein C1orf131 homolog | 0.05441 | 1.39631 | FALSE | FALSE |
| LOC107066415 | axonemal dynein light intermediate polypep | 0.05738 | 1.40076 | FALSE | FALSE |
| LOC107067420 | uncharacterized protein LOC107067420 | 0.05364 | 1.40499 | FALSE | FALSE |
| LOC107066983 | sec1 family domain-containing protein 2-like | 0.05397 | 1.42513 | FALSE | FALSE |
| LOC107069548 | disco-interacting protein 2 homolog A | 0.05114 | 1.45251 | FALSE | FALSE |
| LOC107073688 | cytochrome P450 4V2-like | 0.05347 | 1.4555 | FALSE | FALSE |
| LOC107072017 | cat eye syndrome critical region protein 5-li | 0.05222 | 1.45898 | FALSE | FALSE |
| LOC107068650 | uncharacterized protein LOC107068650 | 0.05452 | 1.47279 | FALSE | FALSE |
| LOC107067400 | programmed cell death protein 6 | 0.05476 | 1.47539 | FALSE | FALSE |
| LOC107069313 | protein N-terminal glutamine amidohydrola | 0.05444 | 1.50399 | FALSE | FALSE |
| LOC107063936 | guanine nucleotide-binding protein G(i) sub | 0.05227 | 1.51196 | FALSE | FALSE |
| LOC107066483 | LOW QUALITY PROTEIN: uncharacterized pr | 0.05323 | 1.51377 | FALSE | FALSE |
| LOC107065546 | mitogen-activated protein kinase kinase kin | 0.05357 | 1.51779 | FALSE | FALSE |
| LOC107068778 | serine/threonine-protein kinase VRK1-like | 0.05261 | 1.51977 | FALSE | FALSE |
| LOC107072512 | acetylcholinesterase-like | 0.05396 | 1.54866 | FALSE | FALSE |

| | | | | | |
|---|---|---|---|---|---|
| LOC107072554 | uncharacterized protein LOC107072554 | 0.05391 | 1.5513 | FALSE | FALSE |
| LOC107070531 | myotubularin-related protein 13 | 0.05373 | 1.56079 | FALSE | FALSE |
| LOC107071603 | protein C10 | 0.05469 | 1.57221 | FALSE | FALSE |
| LOC107070307 | uncharacterized protein LOC107070307 | 0.05384 | 1.57419 | FALSE | FALSE |
| LOC107072940 | 28S ribosomal protein S17, mitochondrial | 0.05261 | 1.57633 | FALSE | FALSE |
| LOC107066198 | dehydrodolichyl diphosphate syntase compl | 0.05383 | 1.57789 | FALSE | FALSE |
| LOC107074467 | AP-3 complex subunit beta-2 | 0.05078 | 1.58059 | FALSE | FALSE |
| LOC107064243 | lisH domain-containing protein FOPNL-like | 0.05308 | 1.58312 | FALSE | FALSE |
| LOC107063960 | enhancer of split mbeta protein-like | 0.05225 | 1.59766 | FALSE | FALSE |
| LOC107063625 | pentatricopeptide repeat-containing protei | 0.05295 | 1.60456 | FALSE | FALSE |
| LOC107065802 | FGGY carbohydrate kinase domain-containi | 0.05308 | 1.61417 | FALSE | FALSE |
| LOC107072200 | WD repeat, SAM and U-box domain-contain | 0.05222 | 1.61904 | FALSE | FALSE |
| LOC107073058 | pyruvate kinase-like | 0.05371 | 1.62117 | FALSE | FALSE |
| LOC107072433 | uncharacterized protein LOC107072433 | 0.05596 | 1.62207 | FALSE | FALSE |
| LOC107074526 | acyl-protein thioesterase 1 | 0.05292 | 1.63386 | FALSE | FALSE |
| LOC107064848 | nectin-3-like | 0.05329 | 1.63947 | FALSE | FALSE |
| LOC107067051 | probable serine/threonine-protein kinase D | 0.05385 | 1.64929 | FALSE | FALSE |
| LOC107067268 | uncharacterized protein LOC107067268 | 0.05193 | 1.65566 | FALSE | FALSE |
| LOC107068510 | uncharacterized protein LOC107068510 | 0.0547 | 1.66053 | FALSE | FALSE |
| LOC107072642 | uncharacterized protein LOC107072642 | 0.05242 | 1.66251 | FALSE | FALSE |
| LOC107063806 | uncharacterized protein LOC107063806 | 0.05194 | 1.66707 | FALSE | FALSE |
| LOC107065899 | leucine-zipper-like transcriptional regulator | 0.05359 | 1.66818 | FALSE | FALSE |
| LOC107072606 | UBX domain-containing protein 7 | 0.0533 | 1.67655 | FALSE | FALSE |
| LOC107067097 | cAMP-dependent protein kinase type II regu | 0.05211 | 1.6807 | FALSE | FALSE |
| LOC107064510 | zinc finger protein 76-like | 0.05232 | 1.68689 | FALSE | FALSE |
| LOC107064938 | protein FAM98B | 0.05333 | 1.70298 | FALSE | FALSE |
| LOC107065140 | U6 snRNA-associated Sm-like protein LSm8 | 0.05245 | 1.71026 | FALSE | FALSE |
| LOC107064689 | uncharacterized protein LOC107064689 | 0.05588 | 1.7169 | FALSE | FALSE |
| LOC107066927 | homocysteine S-methyltransferase 1-like | 0.05454 | 1.73318 | FALSE | FALSE |
| LOC107066945 | ras-related and estrogen-regulated growth i | 0.05496 | 1.7419 | FALSE | FALSE |
| LOC107068725 | 39S ribosomal protein L27, mitochondrial | 0.05433 | 1.7469 | FALSE | FALSE |
| LOC107067933 | short/branched chain specific acyl-CoA dehy | 0.05341 | 1.75094 | FALSE | FALSE |
| LOC107065447 | protein phosphatase 1L | 0.05234 | 1.75887 | FALSE | FALSE |
| LOC107066103 | cysteine-rich venom protein 6-like | 0.05599 | 1.78009 | FALSE | FALSE |
| LOC107067562 | methyltransferase-like protein 22 | 0.05401 | 1.81197 | FALSE | FALSE |
| LOC107067478 | uncharacterized protein LOC107067478 | 0.05082 | 1.81489 | FALSE | FALSE |
| LOC107070030 | endoribonuclease Dicer | 0.05372 | 1.83029 | FALSE | FALSE |
| LOC107069811 | uncharacterized protein LOC107069811 | 0.05296 | 1.86117 | FALSE | FALSE |
| LOC107067686 | GRIP and coiled-coil domain-containing prot | 0.05638 | 1.86457 | FALSE | FALSE |
| LOC107070448 | coiled-coil domain-containing protein 186 | 0.05389 | 1.87146 | FALSE | FALSE |
| LOC107071813 | uncharacterized protein LOC107071813 | 0.05294 | 1.87685 | FALSE | FALSE |
| LOC107069272 | uncharacterized protein LOC107069272 | 0.05266 | 1.89956 | FALSE | FALSE |
| LOC107065690 | stress-associated endoplasmic reticulum pro | 0.05306 | 1.90486 | FALSE | FALSE |
| LOC107065936 | zinc finger HIT domain-containing protein 1 | 0.0526 | 1.91587 | FALSE | FALSE |
| LOC107069071 | cytochrome P450 4C1-like | 0.05329 | 1.91687 | FALSE | FALSE |
| LOC107070859 | translation initiation factor eIF-2B subunit d | 0.05325 | 1.91786 | FALSE | FALSE |
| LOC107064632 | T-box protein H15-like | 0.05346 | 1.91983 | FALSE | FALSE |
| LOC107074249 | adenylate cyclase type 10-like | 0.05328 | 1.93537 | FALSE | FALSE |
| LOC107067115 | heterochromatin protein 1-like | 0.05333 | 1.94184 | FALSE | FALSE |
| LOC107066865 | cytochrome P450 9e2-like | 0.05296 | 1.94217 | FALSE | FALSE |

| | | | | | |
|---|---|---|---|---|---|
| LOC107065245 | uncharacterized protein LOC107065245 | 0.05506 | 1.9439 | FALSE | FALSE |
| LOC107066881 | FMRFamide receptor | 0.0541 | 1.94883 | FALSE | FALSE |
| LOC107065947 | transmembrane channel-like protein 5 | 0.04977 | 1.95032 | FALSE | FALSE |
| LOC107070116 | TM2 domain-containing protein almondex | 0.05397 | 1.95713 | FALSE | FALSE |
| LOC107071338 | uncharacterized protein LOC107071338 | 0.05252 | 1.9626 | FALSE | FALSE |
| LOC107064680 | UDP-glucuronic acid decarboxylase 1 | 0.05294 | 1.9958 | FALSE | FALSE |
| LOC107068785 | uncharacterized protein LOC107068785 | 0.05383 | 2.0003 | FALSE | FALSE |
| LOC107072902 | uncharacterized protein LOC107072902 | 0.05316 | 2.01192 | FALSE | FALSE |
| LOC107064438 | protein disabled | 0.05397 | 2.02346 | FALSE | FALSE |
| LOC107072224 | uncharacterized protein LOC107072224 | 0.05268 | 2.03387 | FALSE | FALSE |
| LOC107066622 | laminin subunit alpha | 0.05161 | 2.07701 | FALSE | FALSE |
| LOC107074513 | uncharacterized protein LOC107074513 | 0.05496 | 2.09719 | FALSE | FALSE |
| LOC107066255 | trafficking protein particle complex subunit | 0.05623 | 2.12665 | FALSE | FALSE |
| LOC107067120 | flightin | 0.05387 | 2.12759 | FALSE | FALSE |
| LOC107074577 | rap guanine nucleotide exchange factor 6-lil | 0.05234 | 2.14076 | FALSE | FALSE |
| LOC107066287 | oxysterol-binding protein-related protein 11 | 0.05441 | 2.14253 | FALSE | FALSE |
| LOC107072202 | uncharacterized protein LOC107072202 | 0.05139 | 2.14564 | FALSE | FALSE |
| LOC107065488 | la-related protein CG11505 | 0.05184 | 2.14716 | FALSE | FALSE |
| LOC107067639 | uncharacterized protein LOC107067639 | 0.05492 | 2.17459 | FALSE | FALSE |
| LOC107068128 | synaptic vesicle 2-related protein | 0.05203 | 2.18547 | FALSE | FALSE |
| LOC107074283 | neurofilament medium polypeptide | 0.053 | 2.18564 | FALSE | FALSE |
| LOC107071149 | uncharacterized protein LOC107071149 | 0.05237 | 2.18856 | FALSE | FALSE |
| LOC107072107 | protein patched | 0.05353 | 2.19344 | FALSE | FALSE |
| LOC107064795 | amphoterin-induced protein 1-like | 0.05289 | 2.20427 | FALSE | FALSE |
| LOC107070998 | pyrimidodiazepine synthase-like | 0.05236 | 2.20474 | FALSE | FALSE |
| LOC107065738 | protein msta, isoform B-like | 0.05239 | 2.22136 | FALSE | FALSE |
| LOC107066514 | V-type proton ATPase catalytic subunit A | 0.05333 | 2.22604 | FALSE | FALSE |
| LOC107074107 | trichohyalin | 0.05348 | 2.22835 | FALSE | FALSE |
| LOC107065997 | LOW QUALITY PROTEIN: uncharacterized pr | 0.05045 | 2.24083 | FALSE | FALSE |
| LOC107066376 | V-type proton ATPase 116 kDa subunit a iso | 0.0541 | 2.25253 | FALSE | FALSE |
| LOC107067076 | vacuolar protein sorting-associated protein | 0.05465 | 2.25281 | FALSE | FALSE |
| LOC107065724 | uncharacterized protein LOC107065724 | 0.05315 | 2.2533 | FALSE | FALSE |
| LOC107066035 | methylcrotonoyl-CoA carboxylase subunit a | 0.05184 | 2.25362 | FALSE | FALSE |
| LOC107066405 | surfeit locus protein 4 homolog | 0.05545 | 2.26127 | FALSE | FALSE |
| LOC107070972 | poly(U)-binding-splicing factor half pint | 0.05269 | 2.26704 | FALSE | FALSE |
| LOC107072737 | scm-like with four MBT domains protein 1 | 0.05246 | 2.27891 | FALSE | FALSE |
| LOC107074376 | uncharacterized protein LOC107074376 | 0.05208 | 2.28756 | FALSE | FALSE |
| LOC107066325 | probable ATP-dependent RNA helicase kurz | 0.05555 | 2.2935 | FALSE | FALSE |
| LOC107071317 | uncharacterized protein LOC107071317 | 0.05201 | 2.29665 | FALSE | FALSE |
| LOC107071391 | exonuclease 1 | 0.05422 | 2.30467 | FALSE | FALSE |
| LOC107064163 | DNA-directed RNA polymerase III subunit RI | 0.05221 | 2.31451 | FALSE | FALSE |
| LOC107070344 | uncharacterized protein LOC107070344 | 0.05456 | 2.31894 | FALSE | FALSE |
| LOC107072389 | WASH complex subunit strumpellin | 0.05179 | 2.36042 | FALSE | FALSE |
| LOC107068203 | trifunctional purine biosynthetic protein ade | 0.05352 | 2.3684 | FALSE | FALSE |
| LOC107066224 | transcriptional repressor p66-beta | 0.05245 | 2.40212 | FALSE | FALSE |
| LOC107068277 | uncharacterized protein LOC107068277 | 0.05254 | 2.40607 | FALSE | FALSE |
| LOC107070356 | acyl-CoA synthetase family member 4 | 0.05216 | 2.41688 | FALSE | FALSE |
| LOC107070927 | anaphase-promoting complex subunit 4 | 0.05146 | 2.41954 | FALSE | FALSE |
| LOC107073365 | activator of basal transcription 1 | 0.05288 | 2.42865 | FALSE | FALSE |
| LOC107065400 | uncharacterized protein LOC107065400 | 0.05256 | 2.43045 | FALSE | FALSE |

| | | | | | |
|---|---|---|---|---|---|
| LOC107068761 | putative methyltransferase NSUN6 | 0.05339 | 2.43857 | FALSE | FALSE |
| LOC107072952 | autophagy-related protein 9A | 0.0518 | 2.44994 | FALSE | FALSE |
| LOC107065146 | multiple PDZ domain protein-like | 0.05295 | 2.45739 | FALSE | FALSE |
| LOC107071257 | protein gooseberry-like | 0.05363 | 2.46861 | FALSE | FALSE |
| LOC107067017 | uncharacterized protein LOC107067017 | 0.05245 | 2.47446 | FALSE | FALSE |
| LOC107070769 | probable glutamine--tRNA ligase | 0.05378 | 2.48687 | FALSE | FALSE |
| LOC107073767 | acidic endochitinase SP2-like | 0.05245 | 2.50212 | FALSE | FALSE |
| LOC107073799 | acidic endochitinase SP2-like | 0.05224 | 2.50212 | FALSE | FALSE |
| LOC107072004 | inositol oxygenase | 0.05391 | 2.50903 | FALSE | FALSE |
| LOC107071874 | ribosome biogenesis protein BOP1 homolog | 0.0531 | 2.51144 | FALSE | FALSE |
| LOC107065723 | homeobox protein Hox-B2a-like | 0.05399 | 2.52105 | FALSE | FALSE |
| LOC107066332 | beta-1,3-galactosyltransferase 1-like | 0.05208 | 2.53698 | FALSE | FALSE |
| LOC107069687 | uncharacterized protein LOC107069687 | 0.05311 | 2.56976 | FALSE | FALSE |
| LOC107069662 | matrix metalloproteinase-2 | 0.05277 | 2.57272 | FALSE | FALSE |
| LOC107070015 | DNA ligase 1 | 0.05531 | 2.5756 | FALSE | FALSE |
| LOC107066988 | guanine nucleotide-binding protein subunit | 0.05297 | 2.58995 | FALSE | FALSE |
| LOC107073976 | uncharacterized protein LOC107073976 | 0.05275 | 2.59265 | FALSE | FALSE |
| LOC107073312 | condensin complex subunit 2-like | 0.05454 | 2.5947 | FALSE | FALSE |
| LOC107071952 | eukaryotic translation initiation factor 3 sub | 0.05376 | 2.59808 | FALSE | FALSE |
| Trnah-gug | Trnah-gug | 0.05469 | 2.60338 | FALSE | FALSE |
| LOC107069706 | RWD domain-containing protein 4 | 0.05402 | 2.61794 | FALSE | FALSE |
| LOC107063980 | protein prune homolog | 0.05257 | 2.63672 | FALSE | FALSE |
| LOC107068264 | probable ATP-dependent RNA helicase DDX | 0.05237 | 2.63725 | FALSE | FALSE |
| LOC107065211 | protein real-time | 0.05369 | 2.6388 | FALSE | FALSE |
| LOC107066713 | protein KRTCAP2 homolog | 0.05253 | 2.64143 | FALSE | FALSE |
| LOC107072130 | KRAB-A domain-containing protein 2-like | 0.05382 | 2.64354 | FALSE | FALSE |
| LOC107067547 | uncharacterized protein LOC107067547 | 0.05282 | 2.66149 | FALSE | FALSE |
| LOC107067999 | probable tRNA (guanine(26)-N(2))-dimethyl | 0.05279 | 2.66505 | FALSE | FALSE |
| LOC107073253 | uncharacterized protein LOC107073253 | 0.05222 | 2.66682 | FALSE | FALSE |
| LOC107070876 | ATPase inhibitor mai-2, mitochondrial-like | 0.05375 | 2.67319 | FALSE | FALSE |
| LOC107072231 | myoneurin-like | 0.0514 | 2.68832 | FALSE | FALSE |
| LOC107073116 | protein CREG1 | 0.05177 | 2.70135 | FALSE | FALSE |
| LOC107073808 | probable protein BRICK1-B | 0.05257 | 2.70964 | FALSE | FALSE |
| LOC107074081 | membrane-associated guanylate kinase, W\ | 0.05159 | 2.71569 | FALSE | FALSE |
| LOC107074589 | protein ARV1 | 0.05294 | 2.73252 | FALSE | FALSE |
| LOC107073701 | uncharacterized protein LOC107073701 | 0.05278 | 2.7381 | FALSE | FALSE |
| LOC107072831 | uncharacterized protein LOC107072831 | 0.05456 | 2.74287 | FALSE | FALSE |
| LOC107074384 | PRKC apoptosis WT1 regulator protein-like | 0.05187 | 2.75284 | FALSE | FALSE |
| LOC107070926 | uncharacterized protein LOC107070926 | 0.05272 | 2.76112 | FALSE | FALSE |
| LOC107067118 | NADH dehydrogenase [ubiquinone | 0.05428 | 2.76333 | FALSE | FALSE |
| LOC107069787 | probable cytochrome P450 305a1 | 0.05339 | 2.764 | FALSE | FALSE |
| LOC107064409 | poly(U)-specific endoribonuclease homolog | 0.05316 | 2.77214 | FALSE | FALSE |
| LOC107066857 | probable serine/threonine-protein kinase D | 0.05402 | 2.80326 | FALSE | FALSE |
| LOC107072657 | methionine--tRNA ligase, mitochondrial | 0.05131 | 2.81657 | FALSE | FALSE |
| LOC107064285 | rhythmically expressed gene 2 protein-like | 0.05237 | 2.84559 | FALSE | FALSE |
| LOC107064629 | longitudinals lacking protein-like | 0.05218 | 2.85115 | FALSE | FALSE |
| LOC107063699 | monocarboxylate transporter 12-like | 0.05275 | 2.85425 | FALSE | FALSE |
| LOC107064497 | unconventional myosin-XVIIIa | 0.05429 | 2.86105 | FALSE | FALSE |
| LOC107069796 | uncharacterized protein LOC107069796 | 0.0536 | 2.86199 | FALSE | FALSE |
| LOC107071715 | uncharacterized protein LOC107071715 | 0.05396 | 2.87308 | FALSE | FALSE |

| | | | | | |
|---|---|---|---|---|---|
| LOC107074291 | uncharacterized protein LOC107074291 | 0.05246 | 2.88814 | FALSE | FALSE |
| LOC107069308 | LOW QUALITY PROTEIN: dihydrofolate redu | 0.05641 | 2.8908 | FALSE | FALSE |
| LOC107064449 | lachesin-like | 0.05389 | 2.89508 | FALSE | FALSE |
| LOC107064525 | stromal membrane-associated protein 1 | 0.05395 | 2.90409 | FALSE | FALSE |
| LOC107066749 | PRKR-interacting protein 1 homolog | 0.05227 | 2.91331 | FALSE | FALSE |
| LOC107065476 | DNA mismatch repair protein Mlh1 | 0.05281 | 2.92474 | FALSE | FALSE |
| LOC107065373 | transcription factor collier | 0.0519 | 2.92478 | FALSE | FALSE |
| LOC107066350 | protein-L-isoaspartate(D-aspartate) O-meth | 0.05297 | 2.9274 | FALSE | FALSE |
| LOC107068557 | odorant receptor 94a-like | 0.05249 | 2.93549 | FALSE | FALSE |
| LOC107065996 | kinesin-like protein unc-104 | 0.05235 | 2.93767 | FALSE | FALSE |
| LOC107071546 | constitutive coactivator of PPAR-gamma-lik | 0.05254 | 2.94663 | FALSE | FALSE |
| LOC107067651 | uncharacterized protein LOC107067651 | 0.05133 | 2.94992 | FALSE | FALSE |
| LOC107068436 | NEDD8-conjugating enzyme UBE2F-like | 0.05324 | 2.95821 | FALSE | FALSE |
| LOC107067328 | aspartate--tRNA ligase, cytoplasmic | 0.05233 | 2.96259 | FALSE | FALSE |
| LOC107069380 | uncharacterized protein LOC107069380 | 0.05431 | 2.96511 | FALSE | FALSE |
| LOC107065666 | uncharacterized protein LOC107065666 | 0.05303 | 2.96705 | FALSE | FALSE |
| LOC107072451 | 6-pyruvoyl tetrahydrobiopterin synthase | 0.05193 | 2.97079 | FALSE | FALSE |
| LOC107073023 | uncharacterized protein LOC107073023 | 0.05276 | 2.99272 | FALSE | FALSE |
| LOC107074297 | small integral membrane protein 14 | 0.0527 | 3.00139 | FALSE | FALSE |
| LOC107074568 | katanin p60 ATPase-containing subunit A-lik | 0.053 | 3.00501 | FALSE | FALSE |
| LOC107067167 | putative ATP-dependent RNA helicase TDRD | 0.05369 | 3.01045 | FALSE | FALSE |
| LOC107072984 | putative inositol monophosphatase 3 | 0.05215 | 3.0124 | FALSE | FALSE |
| LOC107070554 | E3 SUMO-protein ligase RanBP2 | 0.05196 | 3.01604 | FALSE | FALSE |
| LOC107070161 | guanine nucleotide exchange factor for Rab | 0.05172 | 3.01991 | FALSE | FALSE |
| LOC107070071 | pseudouridine-metabolizing bifunctional pro | 0.05176 | 3.02407 | FALSE | FALSE |
| LOC107070353 | lachesin-like | 0.05061 | 3.03834 | FALSE | FALSE |
| LOC107070909 | 50S ribosomal protein L1 | 0.05305 | 3.04423 | FALSE | FALSE |
| LOC107064306 | Golgi-specific brefeldin A-resistance guanine | 0.05238 | 3.04983 | FALSE | FALSE |
| LOC107067873 | uncharacterized protein LOC107067873 | 0.05316 | 3.05369 | FALSE | FALSE |
| LOC107073609 | uncharacterized protein LOC107073609 | 0.05155 | 3.06548 | FALSE | FALSE |
| LOC107070338 | putative uncharacterized protein DDB_G028 | 0.05282 | 3.08027 | FALSE | FALSE |
| LOC107073862 | D-2-hydroxyglutarate dehydrogenase, mitoc | 0.05229 | 3.09112 | FALSE | FALSE |
| LOC107073240 | exonuclease mut-7 homolog | 0.05212 | 3.10149 | FALSE | FALSE |
| LOC107072661 | serine-rich adhesin for platelets | 0.05458 | 3.10229 | FALSE | FALSE |
| LOC107073057 | probable cyclin-dependent serine/threonine | 0.05298 | 3.10712 | FALSE | FALSE |
| LOC107069944 | cysteine sulfinic acid decarboxylase | 0.05105 | 3.11243 | FALSE | FALSE |
| LOC107065105 | mRNA cap guanine-N7 methyltransferase | 0.05411 | 3.12651 | FALSE | FALSE |
| LOC107071802 | uncharacterized protein LOC107071802 | 0.05213 | 3.13259 | FALSE | FALSE |
| LOC107072647 | uncharacterized protein LOC107072647 | 0.05067 | 3.1695 | FALSE | FALSE |
| LOC107068596 | stromal interaction molecule homolog | 0.05367 | 3.1726 | FALSE | FALSE |
| LOC107071732 | regulator of microtubule dynamics protein 1 | 0.05426 | 3.18191 | FALSE | FALSE |
| LOC107073003 | uncharacterized protein LOC107073003 | 0.05156 | 3.18568 | FALSE | FALSE |
| LOC107067504 | uncharacterized protein LOC107067504 | 0.0545 | 3.21251 | FALSE | FALSE |
| LOC107069410 | tRNA (uracil-5-)-methyltransferase homolog | 0.05129 | 3.22849 | FALSE | FALSE |
| LOC107073999 | DNA (cytosine-5)-methyltransferase PliMCI- | 0.05369 | 3.22944 | FALSE | FALSE |
| LOC107068041 | nudix hydrolase 7 | 0.05235 | 3.23176 | FALSE | FALSE |
| LOC107066896 | translation initiation factor IF-2-like | 0.05494 | 3.2326 | FALSE | FALSE |
| LOC107066725 | diacylglycerol O-acyltransferase 1 | 0.05345 | 3.25694 | FALSE | FALSE |
| LOC107074321 | coiled-coil-helix-coiled-coil-helix domain-co | 0.05145 | 3.25712 | FALSE | FALSE |
| LOC107064410 | transmembrane emp24 domain-containing | 0.05202 | 3.25935 | FALSE | FALSE |

| | | | | | |
|---|---|---|---|---|---|
| LOC107064864 | histone H2A | 0.05597 | 3.2598 | FALSE | FALSE |
| LOC107066871 | uncharacterized protein CG3556 | 0.0523 | 3.26319 | FALSE | FALSE |
| LOC107067860 | mitochondrial chaperone BCS1 | 0.05317 | 3.27849 | FALSE | FALSE |
| LOC107068418 | kynurenine/alpha-aminoadipate aminotrans | 0.05198 | 3.27892 | FALSE | FALSE |
| LOC107064724 | ATP synthase subunit s, mitochondrial | 0.05317 | 3.28368 | FALSE | FALSE |
| LOC107067925 | NHL repeat-containing protein 2 | 0.05353 | 3.28676 | FALSE | FALSE |
| LOC107066839 | succinate--hydroxymethylglutarate CoA-tra | 0.05159 | 3.29063 | FALSE | FALSE |
| LOC107074704 | uncharacterized protein LOC107074704 | 0.05126 | 3.30038 | FALSE | FALSE |
| LOC107070214 | non-specific lipid-transfer protein | 0.05204 | 3.31257 | FALSE | FALSE |
| LOC107066873 | PAXIP1-associated glutamate-rich protein 1 | 0.05315 | 3.33036 | FALSE | FALSE |
| LOC107065915 | transcription initiation factor IIA subunit 2 | 0.05309 | 3.3322 | FALSE | FALSE |
| LOC107068344 | uncharacterized protein LOC107068344 | 0.05439 | 3.33702 | FALSE | FALSE |
| LOC107065394 | GPI ethanolamine phosphate transferase 2 | 0.05324 | 3.34192 | FALSE | FALSE |
| LOC107065949 | FAD-dependent oxidoreductase domain-cor | 0.05201 | 3.36595 | FALSE | FALSE |
| LOC107067058 | EGF domain-specific O-linked N-acetylgluco | 0.05177 | 3.37507 | FALSE | FALSE |
| LOC107064068 | spondin-1 | 0.05109 | 3.38788 | FALSE | FALSE |
| LOC107072156 | uncharacterized protein LOC107072156 | 0.0528 | 3.39939 | FALSE | FALSE |
| LOC107067798 | uncharacterized protein LOC107067798 | 0.05283 | 3.40296 | FALSE | FALSE |
| LOC107072317 | PAX3- and PAX7-binding protein 1 | 0.05105 | 3.40305 | FALSE | FALSE |
| LOC107065602 | cytochrome P450 4C1-like | 0.05407 | 3.41455 | FALSE | FALSE |
| LOC107067588 | 3-hydroxyisobutyryl-CoA hydrolase, mitoch | 0.05478 | 3.42503 | FALSE | FALSE |
| LOC107071423 | uncharacterized protein LOC107071423 | 0.05125 | 3.43517 | FALSE | FALSE |
| LOC107072253 | ruvB-like 1 | 0.05165 | 3.47431 | FALSE | FALSE |
| LOC107071761 | zinc finger CCCH domain-containing protein | 0.0514 | 3.48171 | FALSE | FALSE |
| LOC107074608 | uncharacterized protein LOC107074608 | 0.05195 | 3.48435 | FALSE | FALSE |
| LOC107071007 | uncharacterized protein LOC107071007 | 0.05138 | 3.49641 | FALSE | FALSE |
| LOC107067887 | solute carrier family 12 member 9 | 0.05144 | 3.49838 | FALSE | FALSE |
| LOC107073170 | endoplasmic reticulum-Golgi intermediate c | 0.05165 | 3.50199 | FALSE | FALSE |
| LOC107064094 | organic cation transporter 1-like | 0.05162 | 3.50835 | FALSE | FALSE |
| LOC107066159 | uncharacterized protein LOC107066159 | 0.0534 | 3.51577 | FALSE | FALSE |
| LOC107065627 | phosphatidate cytidylyltransferase, photore | 0.05091 | 3.51629 | FALSE | FALSE |
| LOC107073689 | cytochrome P450 4C1-like | 0.05319 | 3.53178 | FALSE | FALSE |
| LOC107066451 | 40S ribosomal protein S14 | 0.05395 | 3.53263 | FALSE | FALSE |
| LOC107069108 | rab3 GTPase-activating protein catalytic sub | 0.05245 | 3.53355 | FALSE | FALSE |
| LOC107073102 | uncharacterized protein LOC107073102 | 0.05237 | 3.53494 | FALSE | FALSE |
| LOC107066847 | probable RNA-binding protein 18 | 0.05218 | 3.54921 | FALSE | FALSE |
| LOC107068273 | uncharacterized protein LOC107068273 | 0.05155 | 3.57196 | FALSE | FALSE |
| LOC107066819 | putative Polycomb group protein ASXL3 | 0.0514 | 3.59237 | FALSE | FALSE |
| LOC107073441 | WD repeat-containing protein 20 | 0.05181 | 3.59418 | FALSE | FALSE |
| LOC107063943 | mitochondrial carrier homolog 2-like | 0.05182 | 3.59543 | FALSE | FALSE |
| LOC107069634 | integrator complex subunit 11 | 0.05216 | 3.60968 | FALSE | FALSE |
| LOC107065255 | zinc finger CCCH domain-containing protein | 0.05161 | 3.61041 | FALSE | FALSE |
| LOC107072847 | serine/threonine-protein kinase 16 | 0.05048 | 3.61698 | FALSE | FALSE |
| LOC107067081 | uncharacterized protein LOC107067081 | 0.05109 | 3.62543 | FALSE | FALSE |
| LOC107072523 | UPF0396 protein-like | 0.05209 | 3.63756 | FALSE | FALSE |
| LOC107070831 | odorant receptor 10a-like | 0.05128 | 3.65606 | FALSE | FALSE |
| LOC107063659 | ras-like protein 3 | 0.05297 | 3.66112 | FALSE | FALSE |
| LOC107074176 | uncharacterized protein LOC107074176 | 0.05131 | 3.66112 | FALSE | FALSE |
| LOC107066296 | sex-lethal homolog | 0.05148 | 3.66633 | FALSE | FALSE |
| LOC107070520 | SWI/SNF-related matrix-associated actin-de | 0.05177 | 3.66651 | FALSE | FALSE |

| | | | | | |
|---|---|---|---|---|---|
| LOC107072157 | branched-chain-amino-acid aminotransfera: | 0.05349 | 3.69059 | FALSE | FALSE |
| LOC107067563 | uncharacterized protein LOC107067563 | 0.05313 | 3.69609 | FALSE | FALSE |
| LOC107073257 | putative fatty acyl-CoA reductase CG5065 | 0.05173 | 3.72215 | FALSE | FALSE |
| LOC107064634 | transmembrane protein 11 homolog, mitoc | 0.05238 | 3.72715 | FALSE | FALSE |
| LOC107071927 | transcription initiation factor TFIID subunit 6 | 0.05138 | 3.7301 | FALSE | FALSE |
| LOC107066740 | uncharacterized protein LOC107066740 | 0.0541 | 3.73669 | FALSE | FALSE |
| LOC107072273 | disrupted in renal carcinoma protein 2 hom | 0.05462 | 3.74585 | FALSE | FALSE |
| LOC107067845 | beta-1,3-glucosyltransferase | 0.05258 | 3.75773 | FALSE | FALSE |
| LOC107074477 | transcription elongation factor S-II | 0.05307 | 3.77064 | FALSE | FALSE |
| LOC107073113 | intersectin-1 | 0.05399 | 3.77363 | FALSE | FALSE |
| LOC107072143 | uncharacterized protein LOC107072143 | 0.05223 | 3.77375 | FALSE | FALSE |
| LOC107068218 | potassium voltage-gated channel protein ea | 0.05284 | 3.78128 | FALSE | FALSE |
| LOC107066241 | protein GPR107 | 0.05119 | 3.80799 | FALSE | FALSE |
| LOC107068174 | potassium channel subfamily K member 13- | 0.05091 | 3.81552 | FALSE | FALSE |
| LOC107069091 | choline O-acetyltransferase | 0.05211 | 3.81867 | FALSE | FALSE |
| LOC107063945 | echinoderm microtubule-associated proteir | 0.0504 | 3.83 | FALSE | FALSE |
| LOC107064805 | serine proteinase stubble-like | 0.05091 | 3.8331 | FALSE | FALSE |
| LOC107064091 | pre-mRNA-splicing factor ISY1 homolog | 0.05244 | 3.84394 | FALSE | FALSE |
| LOC107069359 | spermine oxidase | 0.05314 | 3.84411 | FALSE | FALSE |
| LOC107067040 | uncharacterized protein LOC107067040 | 0.05257 | 3.84572 | FALSE | FALSE |
| LOC107067084 | cell division control protein 31-like | 0.05244 | 3.84631 | FALSE | FALSE |
| LOC107073619 | breast cancer type 2 susceptibility protein h | 0.05446 | 3.87967 | FALSE | FALSE |
| LOC107072526 | uncharacterized protein LOC107072526 | 0.05073 | 3.88073 | FALSE | FALSE |
| LOC107067283 | uncharacterized protein LOC107067283 | 0.05309 | 3.88978 | FALSE | FALSE |
| LOC107073580 | uncharacterized protein LOC107073580 | 0.05227 | 3.91482 | FALSE | FALSE |
| LOC107067757 | uncharacterized protein LOC107067757 | 0.05271 | 3.91575 | FALSE | FALSE |
| LOC107065971 | uncharacterized protein LOC107065971 | 0.05367 | 3.91891 | FALSE | FALSE |
| LOC107072070 | dnaJ homolog subfamily C member 24-like | 0.04951 | 3.92602 | FALSE | FALSE |
| LOC107066425 | nuclear pore complex protein DDB_G02749 | 0.05239 | 3.93734 | FALSE | FALSE |
| LOC107070903 | homeobox protein orthopedia B-like | 0.0517 | 3.9396 | FALSE | FALSE |
| LOC107068410 | exocyst complex component 1 | 0.05162 | 3.94418 | FALSE | FALSE |
| LOC107066446 | uncharacterized protein LOC107066446 | 0.05323 | 3.94797 | FALSE | FALSE |
| LOC107067445 | protein N-lysine methyltransferase METTL2( | 0.05134 | 3.95374 | FALSE | FALSE |
| LOC107073651 | uncharacterized protein LOC107073651 | 0.05121 | 3.95445 | FALSE | FALSE |
| LOC107073892 | U3 small nucleolar RNA-associated protein 1 | 0.05101 | 3.96037 | FALSE | FALSE |
| LOC107074285 | intraflagellar transport protein 140 homolo | 0.05265 | 3.97673 | FALSE | FALSE |
| LOC107064480 | uncharacterized protein LOC107064480 | 0.05218 | 3.98434 | FALSE | FALSE |
| LOC107065141 | protein slit | 0.05108 | 3.98543 | FALSE | FALSE |
| LOC107068148 | uncharacterized protein LOC107068148 | 0.05221 | 3.9865 | FALSE | FALSE |
| LOC107065236 | uncharacterized protein LOC107065236 | 0.05141 | 4.00466 | FALSE | FALSE |
| LOC107071682 | aminopeptidase N | 0.05345 | 4.00502 | FALSE | FALSE |
| LOC107066953 | cytochrome P450 4C1-like | 0.05195 | 4.02773 | FALSE | FALSE |
| LOC107070577 | uncharacterized protein LOC107070577 | 0.05233 | 4.03378 | FALSE | FALSE |
| LOC107067572 | uncharacterized protein LOC107067572 | 0.04952 | 4.05638 | FALSE | FALSE |
| LOC107066496 | uncharacterized protein LOC107066496 | 0.0533 | 4.06535 | FALSE | FALSE |
| LOC107073142 | uncharacterized protein C45G9.7 | 0.05127 | 4.0692 | FALSE | FALSE |
| LOC107072198 | vacuolar protein sorting-associated protein | 0.05066 | 4.07307 | FALSE | FALSE |
| LOC107073048 | protein AF-10-like | 0.05196 | 4.08477 | FALSE | FALSE |
| LOC107074194 | extended synaptotagmin-2-B | 0.05237 | 4.09112 | FALSE | FALSE |
| LOC107069115 | uncharacterized protein LOC107069115 | 0.05151 | 4.09646 | FALSE | FALSE |

| | | | | | |
|---|---|---|---|---|---|
| LOC107074395 | uncharacterized protein LOC107074395 | 0.05127 | 4.10552 | FALSE | FALSE |
| LOC107068527 | iduronate 2-sulfatase | 0.04996 | 4.10601 | FALSE | FALSE |
| LOC107072489 | dihydrolipoyllysine-residue succinyltransfera | 0.04924 | 4.10719 | FALSE | FALSE |
| LOC107072313 | spastin | 0.05299 | 4.12091 | FALSE | FALSE |
| LOC107072585 | N-acetylgalactosaminyltransferase 7 | 0.05389 | 4.12554 | FALSE | FALSE |
| LOC107068253 | nuclear inhibitor of protein phosphatase 1 | 0.05105 | 4.12615 | FALSE | FALSE |
| LOC107072933 | protein brunelleschi | 0.05149 | 4.13086 | FALSE | FALSE |
| LOC107068267 | uncharacterized protein LOC107068267 | 0.05333 | 4.13193 | FALSE | FALSE |
| LOC107066580 | myb-like protein AA | 0.05063 | 4.14555 | FALSE | FALSE |
| LOC107074643 | large proline-rich protein BAG6 | 0.05082 | 4.14878 | FALSE | FALSE |
| LOC107073108 | protein TAPT1 homolog | 0.05254 | 4.15437 | FALSE | FALSE |
| LOC107065180 | alpha-tocopherol transfer protein-like | 0.05092 | 4.15511 | FALSE | FALSE |
| LOC107069908 | CTD small phosphatase-like protein 2 | 0.05101 | 4.15871 | FALSE | FALSE |
| LOC107072992 | putative mediator of RNA polymerase II trar | 0.05102 | 4.16229 | FALSE | FALSE |
| LOC107072864 | MAM and LDL-receptor class A domain-cont | 0.05058 | 4.19122 | FALSE | FALSE |
| LOC107072288 | splicing factor 45 | 0.05192 | 4.19337 | FALSE | FALSE |
| LOC107068223 | protein broad-minded-like | 0.05163 | 4.19486 | FALSE | FALSE |
| LOC107073859 | peroxisome assembly factor 2 | 0.05038 | 4.19897 | FALSE | FALSE |
| LOC107067512 | zinc finger protein 775-like | 0.05163 | 4.19986 | FALSE | FALSE |
| LOC107067524 | zinc finger E-box-binding homeobox 1-like | 0.0518 | 4.20109 | FALSE | FALSE |
| LOC107074102 | sentrin-specific protease 1-like | 0.05117 | 4.20881 | FALSE | FALSE |
| LOC107068496 | aldehyde dehydrogenase, dimeric NADP-pre | 0.05312 | 4.21461 | FALSE | FALSE |
| LOC107063889 | actin-related protein 3 | 0.05212 | 4.22942 | FALSE | FALSE |
| LOC107071826 | putative fatty acyl-CoA reductase CG8306 | 0.05249 | 4.23059 | FALSE | FALSE |
| LOC107072284 | bifunctional methylenetetrahydrofolate deh | 0.05207 | 4.24627 | FALSE | FALSE |
| LOC107067946 | cytochrome P450 4c3-like | 0.0517 | 4.25221 | FALSE | FALSE |
| LOC107073251 | ubiquinone biosynthesis protein COQ9, mito | 0.04999 | 4.25306 | FALSE | FALSE |
| LOC107070203 | zinc finger protein 709-like | 0.05237 | 4.26232 | FALSE | FALSE |
| LOC107071477 | nuclear pore glycoprotein p62 | 0.05179 | 4.2788 | FALSE | FALSE |
| LOC107068051 | proclotting enzyme-like | 0.05142 | 4.29192 | FALSE | FALSE |
| LOC107068979 | uncharacterized protein LOC107068979 | 0.05075 | 4.29735 | FALSE | FALSE |
| LOC107068450 | zinc finger protein 341-like | 0.05037 | 4.30615 | FALSE | FALSE |
| LOC107070078 | uncharacterized protein LOC107070078 | 0.05223 | 4.30758 | FALSE | FALSE |
| LOC107065391 | uncharacterized protein LOC107065391 | 0.05239 | 4.30855 | FALSE | FALSE |
| LOC107070503 | uncharacterized protein LOC107070503 | 0.05114 | 4.31225 | FALSE | FALSE |
| LOC107066964 | conserved oligomeric Golgi complex subunit | 0.05137 | 4.32601 | FALSE | FALSE |
| LOC107069504 | probable serine incorporator | 0.05063 | 4.32793 | FALSE | FALSE |
| LOC107065048 | uncharacterized protein LOC107065048 | 0.05202 | 4.33787 | FALSE | FALSE |
| LOC107065803 | G-protein coupled receptor 143-like | 0.0511 | 4.33996 | FALSE | FALSE |
| LOC107068820 | 39S ribosomal protein L4, mitochondrial | 0.0516 | 4.34097 | FALSE | FALSE |
| LOC107069166 | uncharacterized protein LOC107069166 | 0.05102 | 4.35848 | FALSE | FALSE |
| LOC107069240 | uncharacterized protein LOC107069240 | 0.05403 | 4.36768 | FALSE | FALSE |
| LOC107072521 | rRNA 2'-O-methyltransferase fibrillarin | 0.05148 | 4.37467 | FALSE | FALSE |
| LOC107070508 | pyridine nucleotide-disulfide oxidoreductase | 0.0507 | 4.3765 | FALSE | FALSE |
| LOC107072809 | acetyl-CoA acetyltransferase, mitochondrial | 0.05109 | 4.4023 | FALSE | FALSE |
| LOC107072349 | sprT-like domain-containing protein Spartar | 0.05069 | 4.41606 | FALSE | FALSE |
| LOC107067483 | sequestosome-1 | 0.05166 | 4.42528 | FALSE | FALSE |
| LOC107067213 | peptidoglycan-recognition protein LC-like | 0.05202 | 4.42763 | FALSE | FALSE |
| LOC107068680 | zinc finger matrin-type protein 2 | 0.05169 | 4.42853 | FALSE | FALSE |
| LOC107070158 | uncharacterized protein LOC107070158 | 0.04989 | 4.4356 | FALSE | FALSE |

| | | | | | |
|---|---|---|---|---|---|
| LOC107066046 | uncharacterized protein LOC107066046 | 0.051 | 4.44846 | FALSE | FALSE |
| LOC107073897 | importin-11 | 0.05241 | 4.44877 | FALSE | FALSE |
| LOC107067680 | pyridoxal kinase | 0.05195 | 4.45166 | FALSE | FALSE |
| LOC107065080 | major royal jelly protein 1-like | 0.0499 | 4.45208 | FALSE | FALSE |
| LOC107070677 | uncharacterized protein LOC107070677 | 0.05177 | 4.45734 | FALSE | FALSE |
| LOC107068227 | 60S ribosomal protein L10a | 0.05131 | 4.46793 | FALSE | FALSE |
| LOC107073851 | uncharacterized protein LOC107073851 | 0.05054 | 4.46969 | FALSE | FALSE |
| LOC107065689 | restin homolog | 0.05424 | 4.48894 | FALSE | FALSE |
| LOC107072442 | uncharacterized protein LOC107072442 | 0.05167 | 4.4893 | FALSE | FALSE |
| LOC107072629 | gametocyte-specific factor 1-like | 0.05067 | 4.49766 | FALSE | FALSE |
| LOC107074185 | hippocampus abundant transcript 1 protein | 0.05141 | 4.50236 | FALSE | FALSE |
| LOC107074632 | apyrase-like | 0.05198 | 4.50262 | FALSE | FALSE |
| LOC107071084 | growth arrest-specific protein 1-like | 0.05235 | 4.50725 | FALSE | FALSE |
| LOC107070986 | transmembrane channel-like protein 2 | 0.05236 | 4.51248 | FALSE | FALSE |
| LOC107067532 | UDP-glucuronosyltransferase 2C1-like | 0.05147 | 4.51852 | FALSE | FALSE |
| LOC107070069 | uncharacterized protein LOC107070069 | 0.05168 | 4.51878 | FALSE | FALSE |
| LOC107072212 | protein PFC0760c-like | 0.05081 | 4.52451 | FALSE | FALSE |
| LOC107064518 | probable dolichol-phosphate mannosyltrans | 0.05107 | 4.52629 | FALSE | FALSE |
| LOC107068926 | uncharacterized protein LOC107068926 | 0.05056 | 4.52966 | FALSE | FALSE |
| LOC107068362 | tetratricopeptide repeat protein 7B | 0.05159 | 4.53666 | FALSE | FALSE |
| LOC107064797 | uncharacterized protein LOC107064797 | 0.05161 | 4.55338 | FALSE | FALSE |
| LOC107071585 | regucalcin-like | 0.05215 | 4.55479 | FALSE | FALSE |
| LOC107068443 | UPF0553 protein C9orf64 homolog | 0.05327 | 4.6213 | FALSE | FALSE |
| LOC107067542 | cyclin-dependent kinase 2-like | 0.05276 | 4.62906 | FALSE | FALSE |
| LOC107073444 | uncharacterized protein LOC107073444 | 0.05204 | 4.62948 | FALSE | FALSE |
| LOC107072646 | complement C1s subcomponent-like | 0.05178 | 4.63068 | FALSE | FALSE |
| LOC107070287 | nuclear transcription factor Y subunit alpha | 0.0519 | 4.63141 | FALSE | FALSE |
| LOC107074160 | rho guanine nucleotide exchange factor 3-lil | 0.05142 | 4.65357 | FALSE | FALSE |
| LOC107069279 | angio-associated migratory cell protein | 0.04969 | 4.66563 | FALSE | FALSE |
| LOC107067438 | E3 ubiquitin-protein ligase Bre1 | 0.05382 | 4.67659 | FALSE | FALSE |
| LOC107068788 | bromodomain-containing protein DDB_G02 | 0.05268 | 4.67919 | FALSE | FALSE |
| LOC107068397 | mpv17-like protein 2 | 0.05165 | 4.71012 | FALSE | FALSE |
| LOC107070480 | DNA primase small subunit-like | 0.05259 | 4.71565 | FALSE | FALSE |
| LOC107068493 | cytochrome P450 6A1-like | 0.05097 | 4.71687 | FALSE | FALSE |
| LOC107065866 | protein grindelwald | 0.05132 | 4.71791 | FALSE | FALSE |
| LOC107074580 | uncharacterized protein LOC107074580 | 0.05303 | 4.71817 | FALSE | FALSE |
| LOC107067656 | probable 28S ribosomal protein S25, mitoch | 0.05179 | 4.72882 | FALSE | FALSE |
| LOC107069098 | sodium channel protein Nach | 0.05208 | 4.73149 | FALSE | FALSE |
| LOC107071622 | dolichyl-diphosphooligosaccharide--protein | 0.05317 | 4.73927 | FALSE | FALSE |
| LOC107068408 | trafficking protein particle complex subunit | 0.05157 | 4.74007 | FALSE | FALSE |
| LOC107067526 | zinc finger protein 768-like | 0.05083 | 4.74245 | FALSE | FALSE |
| LOC107070042 | protein-methionine sulfoxide oxidase mical3 | 0.04925 | 4.75262 | FALSE | FALSE |
| LOC107065067 | extensin-1-like | 0.0502 | 4.77768 | FALSE | FALSE |
| LOC107064725 | uncharacterized protein LOC107064725 | 0.05699 | 4.77836 | FALSE | FALSE |
| LOC107069915 | protein CutA homolog | 0.0508 | 4.78685 | FALSE | FALSE |
| LOC107068125 | uncharacterized protein LOC107068125 | 0.05291 | 4.78924 | FALSE | FALSE |
| LOC107069215 | uncharacterized protein LOC107069215 | 0.05323 | 4.79765 | FALSE | FALSE |
| LOC107073409 | uncharacterized protein LOC107073409 | 0.05465 | 4.82544 | FALSE | FALSE |
| LOC107070498 | NF-kappa-B inhibitor-interacting Ras-like pro | 0.05142 | 4.82976 | FALSE | FALSE |
| LOC107073020 | isocitrate dehydrogenase [NAD | 0.05178 | 4.83741 | FALSE | FALSE |

| | | | | | |
|---|---|---|---|---|---|
| LOC107073954 | dnaJ homolog subfamily A member 4 | 0.05196 | 4.8391 | FALSE | FALSE |
| LOC107072551 | uncharacterized protein LOC107072551 | 0.05124 | 4.83954 | FALSE | FALSE |
| LOC107068741 | uncharacterized protein LOC107068741 | 0.05226 | 4.85795 | FALSE | FALSE |
| LOC107071195 | uncharacterized protein LOC107071195 | 0.05011 | 4.86056 | FALSE | FALSE |
| LOC107072217 | uncharacterized protein LOC107072217 | 0.05077 | 4.86827 | FALSE | FALSE |
| LOC107072191 | uncharacterized protein LOC107072191 | 0.05004 | 4.88381 | FALSE | FALSE |
| LOC107065033 | uncharacterized protein LOC107065033 | 0.05254 | 4.88491 | FALSE | FALSE |
| LOC107073604 | acyl-CoA Delta(11) desaturase-like | 0.04876 | 4.89915 | FALSE | FALSE |
| LOC107074196 | uncharacterized protein LOC107074196 | 0.05125 | 4.90912 | FALSE | FALSE |
| LOC107068092 | uncharacterized protein LOC107068092 | 0.05003 | 4.91363 | FALSE | FALSE |
| LOC107066579 | uncharacterized protein LOC107066579 | 0.05246 | 4.91694 | FALSE | FALSE |
| LOC107070527 | general transcription factor IIH subunit 1 | 0.05079 | 4.91834 | FALSE | FALSE |
| LOC107063639 | ribosome biogenesis protein BRX1 homolog | 0.05056 | 4.92111 | FALSE | FALSE |
| LOC107073154 | structural maintenance of chromosomes pr | 0.05187 | 4.93881 | FALSE | FALSE |
| LOC107071281 | uncharacterized protein LOC107071281 | 0.05177 | 4.94999 | FALSE | FALSE |
| LOC107069816 | putative uncharacterized protein DDB_G02 | 0.05235 | 4.95313 | FALSE | FALSE |
| LOC107066011 | metastasis-associated protein MTA3 | 0.05041 | 4.95768 | FALSE | FALSE |
| LOC107069402 | von Willebrand factor A domain-containing | 0.04967 | 4.96402 | FALSE | FALSE |
| LOC107066569 | fibroblast growth factor receptor substrate | 0.04984 | 4.96682 | FALSE | FALSE |
| LOC107065682 | protein SERAC1 | 0.05105 | 4.98437 | FALSE | FALSE |
| LOC107069868 | uncharacterized protein LOC107069868 | 0.05278 | 4.99155 | FALSE | FALSE |
| LOC107066971 | uncharacterized protein LOC107066971 | 0.05052 | 5.00188 | FALSE | FALSE |
| LOC107064524 | orexin receptor type 1-like | 0.05086 | 5.00282 | FALSE | FALSE |
| LOC107066000 | UDP-xylose and UDP-N-acetylglucosamine t | 0.05152 | 5.00444 | FALSE | FALSE |
| LOC107069628 | Hermansky-Pudlak syndrome 5 protein hom | 0.04996 | 5.0048 | FALSE | FALSE |
| LOC107067358 | RING finger protein 212B | 0.05235 | 5.00553 | FALSE | FALSE |
| LOC107066522 | protein giant-lens | 0.05216 | 5.0065 | FALSE | FALSE |
| LOC107072603 | neuroligin-4, X-linked-like | 0.04972 | 5.00991 | FALSE | FALSE |
| LOC107067520 | protein bric-a-brac 2 | 0.05223 | 5.01195 | FALSE | FALSE |
| LOC107067112 | COP9 signalosome complex subunit 8 | 0.05155 | 5.01386 | FALSE | FALSE |
| LOC107065829 | U2 snRNP-associated SURP motif-containing | 0.05074 | 5.01669 | FALSE | FALSE |
| LOC107067568 | probable phosphorylase b kinase regulatory | 0.05155 | 5.01954 | FALSE | FALSE |
| LOC107073968 | serine/threonine-protein kinase TBK1 | 0.05124 | 5.02228 | FALSE | FALSE |
| LOC107074199 | nucleolar protein of 40 kDa-like | 0.05093 | 5.02328 | FALSE | FALSE |
| LOC107073869 | CCR4-NOT transcription complex subunit 10 | 0.05043 | 5.04519 | FALSE | FALSE |
| LOC107064981 | CCR4-NOT transcription complex subunit 1 | 0.05062 | 5.04818 | FALSE | FALSE |
| LOC107068398 | FHA domain-containing protein FhaA-like | 0.05167 | 5.05494 | FALSE | FALSE |
| LOC107067689 | methylosome protein 50-like | 0.05226 | 5.05852 | FALSE | FALSE |
| LOC107065584 | inducible metalloproteinase inhibitor protei | 0.05189 | 5.07305 | FALSE | FALSE |
| LOC107073168 | uncharacterized protein LOC107073168 | 0.0513 | 5.07652 | FALSE | FALSE |
| LOC107073706 | piggyBac transposable element-derived pro | 0.04964 | 5.08486 | FALSE | FALSE |
| LOC107064503 | adenylate kinase isoenzyme 6 | 0.05084 | 5.09551 | FALSE | FALSE |
| LOC107070275 | uncharacterized protein LOC107070275 | 0.05116 | 5.09893 | FALSE | FALSE |
| LOC107071129 | uncharacterized protein LOC107071129 | 0.05029 | 5.10019 | FALSE | FALSE |
| LOC107068105 | CDGSH iron-sulfur domain-containing prote | 0.05161 | 5.10744 | FALSE | FALSE |
| LOC107067425 | uncharacterized protein LOC107067425 | 0.05212 | 5.11048 | FALSE | FALSE |
| LOC107073673 | fatty acyl-CoA reductase 1-like | 0.051 | 5.1126 | FALSE | FALSE |
| LOC107068630 | peroxisomal membrane protein PEX14 | 0.05119 | 5.12678 | FALSE | FALSE |
| LOC107065589 | acidic phospholipase A2 1 | 0.05149 | 5.12937 | FALSE | FALSE |
| LOC107067298 | tRNA:m(4)X modification enzyme TRM13 ho | 0.05113 | 5.14075 | FALSE | FALSE |

| | | | | | |
|---|---|---|---|---|---|
| LOC107073399 | vacuolar protein sorting-associated protein | 0.05035 | 5.15356 | FALSE | FALSE |
| LOC107067983 | protein lethal(2)essential for life-like | 0.04968 | 5.17273 | FALSE | FALSE |
| LOC107072028 | F-BAR and double SH3 domains protein 2 | 0.05158 | 5.19114 | FALSE | FALSE |
| LOC107073033 | trypsin-like | 0.05196 | 5.2013 | FALSE | FALSE |
| LOC107068159 | translin | 0.05064 | 5.21466 | FALSE | FALSE |
| LOC107069044 | innexin shaking-B-like | 0.05122 | 5.21555 | FALSE | FALSE |
| LOC107071643 | fatty-acid amide hydrolase 2-B-like | 0.05105 | 5.22068 | FALSE | FALSE |
| LOC107073727 | putative fatty acyl-CoA reductase CG5065 | 0.05223 | 5.22734 | FALSE | FALSE |
| LOC107069734 | dopamine N-acetyltransferase-like | 0.0535 | 5.2336 | FALSE | FALSE |
| LOC107068905 | uncharacterized protein LOC107068905 | 0.04879 | 5.23476 | FALSE | FALSE |
| LOC107069051 | uncharacterized protein C2orf42 homolog | 0.05193 | 5.23675 | FALSE | FALSE |
| LOC107069786 | methyl farnesoate epoxidase-like | 0.05023 | 5.2484 | FALSE | FALSE |
| LOC107065896 | short-chain dehydrogenase/reductase famil | 0.04956 | 5.25612 | FALSE | FALSE |
| LOC107066434 | uncharacterized protein LOC107066434 | 0.05366 | 5.27326 | FALSE | FALSE |
| LOC107069176 | putative uncharacterized protein DDB_G027 | 0.0508 | 5.30095 | FALSE | FALSE |
| LOC107065907 | laminin subunit gamma-1 | 0.05347 | 5.30192 | FALSE | FALSE |
| LOC107070697 | uncharacterized protein LOC107070697 | 0.05043 | 5.30456 | FALSE | FALSE |
| LOC107071457 | active breakpoint cluster region-related pro | 0.04923 | 5.31082 | FALSE | FALSE |
| LOC107070234 | G-protein coupled receptor moody-like | 0.05185 | 5.31183 | FALSE | FALSE |
| LOC107072287 | alpha-1,3/1,6-mannosyltransferase ALG2 | 0.05074 | 5.31783 | FALSE | FALSE |
| LOC107065094 | peritrophin-1-like | 0.04996 | 5.3186 | FALSE | FALSE |
| LOC107066815 | colorectal mutant cancer protein | 0.05087 | 5.32857 | FALSE | FALSE |
| LOC107070586 | sodium-independent sulfate anion transpor | 0.0513 | 5.32872 | FALSE | FALSE |
| LOC107068216 | arrestin homolog | 0.05097 | 5.33343 | FALSE | FALSE |
| LOC107074070 | zinc finger protein ZFP69-like | 0.05128 | 5.34189 | FALSE | FALSE |
| LOC107069642 | cleavage and polyadenylation specificity fac | 0.05105 | 5.34641 | FALSE | FALSE |
| LOC107071008 | uncharacterized protein LOC107071008 | 0.05193 | 5.36076 | FALSE | FALSE |
| LOC107071358 | uncharacterized protein LOC107071358 | 0.05086 | 5.36662 | FALSE | FALSE |
| LOC107067114 | uncharacterized protein LOC107067114 | 0.05086 | 5.37728 | FALSE | FALSE |
| LOC107066591 | uncharacterized protein LOC107066591 | 0.05072 | 5.37865 | FALSE | FALSE |
| LOC107068822 | sorbitol dehydrogenase-like | 0.05244 | 5.37987 | FALSE | FALSE |
| LOC107069980 | UPF0693 protein C10orf32 homolog | 0.04986 | 5.38145 | FALSE | FALSE |
| LOC107066767 | uncharacterized protein LOC107066767 | 0.04954 | 5.38765 | FALSE | FALSE |
| LOC107068427 | osteomodulin-like | 0.05189 | 5.39249 | FALSE | FALSE |
| LOC107067957 | uncharacterized protein LOC107067957 | 0.05134 | 5.39388 | FALSE | FALSE |
| LOC107068091 | retinal rod rhodopsin-sensitive cGMP 3',5'-c | 0.05152 | 5.39413 | FALSE | FALSE |
| LOC107074532 | hemicentin-1 | 0.04992 | 5.39761 | FALSE | FALSE |
| LOC107066301 | tetratricopeptide repeat protein 14 homolo | 0.05168 | 5.40429 | FALSE | FALSE |
| LOC107073555 | 2-oxoisovalerate dehydrogenase subunit be | 0.05131 | 5.41217 | FALSE | FALSE |
| LOC107066551 | serine/arginine repetitive matrix protein 1 | 0.05112 | 5.42454 | FALSE | FALSE |
| LOC107068925 | histone H2B | 0.04998 | 5.43576 | FALSE | FALSE |
| LOC107071163 | uncharacterized protein LOC107071163 | 0.05041 | 5.44417 | FALSE | FALSE |
| LOC107068652 | facilitated trehalose transporter Tret1-2 hor | 0.05271 | 5.45675 | FALSE | FALSE |
| LOC107071170 | uncharacterized protein LOC107071170 | 0.05082 | 5.46525 | FALSE | FALSE |
| LOC107067917 | uncharacterized protein LOC107067917 | 0.0494 | 5.46648 | FALSE | FALSE |
| LOC107070984 | papilin | 0.0499 | 5.47178 | FALSE | FALSE |
| LOC107074130 | CWF19-like protein 1 | 0.04807 | 5.48459 | FALSE | FALSE |
| LOC107065017 | uncharacterized protein LOC107065017 | 0.05042 | 5.48532 | FALSE | FALSE |
| LOC107066904 | probable proline--tRNA ligase, mitochondria | 0.05109 | 5.50109 | FALSE | FALSE |
| LOC107064166 | uncharacterized protein LOC107064166 | 0.05038 | 5.50184 | FALSE | FALSE |

| | | | | | |
|---|---|---|---|---|---|
| LOC107065672 | nuclear RNA export factor 1-like | 0.05142 | 5.52361 | FALSE | FALSE |
| LOC107074153 | N6-adenosine-methyltransferase subunit M | 0.05033 | 5.52947 | FALSE | FALSE |
| LOC107068854 | armadillo segment polarity protein | 0.05101 | 5.5549 | FALSE | FALSE |
| LOC107070217 | uncharacterized protein LOC107070217 | 0.0506 | 5.57438 | FALSE | FALSE |
| LOC107071408 | icarapin-like | 0.05011 | 5.59601 | FALSE | FALSE |
| LOC107065598 | orcokinin peptides type A-like | 0.05171 | 5.59715 | FALSE | FALSE |
| LOC107064025 | uncharacterized protein LOC107064025 | 0.05026 | 5.60803 | FALSE | FALSE |
| LOC107067046 | nuclear receptor subfamily 2 group E memb | 0.05218 | 5.61008 | FALSE | FALSE |
| LOC107069556 | ATP-dependent DNA helicase Q1-like | 0.05175 | 5.62875 | FALSE | FALSE |
| LOC107069861 | pro-resilin-like | 0.05145 | 5.63241 | FALSE | FALSE |
| LOC107065369 | GATA zinc finger domain-containing protein | 0.0507 | 5.63587 | FALSE | FALSE |
| LOC107063903 | protein NPC2 homolog | 0.05052 | 5.639 | FALSE | FALSE |
| LOC107070982 | two pore potassium channel protein sup-9 | 0.05234 | 5.64143 | FALSE | FALSE |
| LOC107065205 | uncharacterized protein LOC107065205 | 0.05015 | 5.66269 | FALSE | FALSE |
| LOC107063701 | uncharacterized protein LOC107063701 | 0.0502 | 5.67644 | FALSE | FALSE |
| LOC107072704 | nuclear factor of activated T-cells 5 | 0.04989 | 5.67881 | FALSE | FALSE |
| LOC107067779 | cuticle protein 79, isoform B-like | 0.05112 | 5.68252 | FALSE | FALSE |
| LOC107072707 | 40S ribosomal protein S28-like | 0.05217 | 5.6982 | FALSE | FALSE |
| LOC107070382 | neuronal membrane glycoprotein M6-a | 0.05131 | 5.70106 | FALSE | FALSE |
| LOC107071478 | RING finger and CHY zinc finger domain-con | 0.05137 | 5.70615 | FALSE | FALSE |
| LOC107068768 | NADP-dependent malic enzyme | 0.0508 | 5.71026 | FALSE | FALSE |
| LOC107070872 | lysosomal alpha-glucosidase-like | 0.05116 | 5.7123 | FALSE | FALSE |
| LOC107071103 | CWF19-like protein 2 | 0.05088 | 5.72331 | FALSE | FALSE |
| LOC107066452 | dynactin subunit 4 | 0.05011 | 5.73306 | FALSE | FALSE |
| LOC107070235 | phosphotriesterase-related protein | 0.05071 | 5.73924 | FALSE | FALSE |
| LOC107066066 | cilia- and flagella-associated protein 43-like | 0.05121 | 5.75227 | FALSE | FALSE |
| LOC107067266 | mediator of RNA polymerase II transcriptior | 0.04959 | 5.77052 | FALSE | FALSE |
| LOC107067792 | uncharacterized protein LOC107067792 | 0.0518 | 5.77839 | FALSE | FALSE |
| LOC107074574 | kelch-like protein 10 | 0.05092 | 5.77985 | FALSE | FALSE |
| LOC107071628 | alpha-endosulfine | 0.05284 | 5.78368 | FALSE | FALSE |
| LOC107071043 | uncharacterized protein LOC107071043 | 0.05124 | 5.78394 | FALSE | FALSE |
| LOC107070348 | unconventional myosin-Va | 0.05248 | 5.78561 | FALSE | FALSE |
| LOC107070027 | uncharacterized protein KIAA0513 | 0.05097 | 5.80931 | FALSE | FALSE |
| LOC107066516 | uncharacterized protein LOC107066516 | 0.05114 | 5.8241 | FALSE | FALSE |
| LOC107072999 | homeobox protein Hmx | 0.05257 | 5.83246 | FALSE | FALSE |
| LOC107065581 | WD40 repeat-containing protein SMU1 | 0.05022 | 5.84055 | FALSE | FALSE |
| LOC107063988 | uncharacterized protein LOC107063988 | 0.05142 | 5.8519 | FALSE | FALSE |
| LOC107074140 | uncharacterized protein LOC107074140 | 0.05074 | 5.87736 | FALSE | FALSE |
| LOC107071034 | secretin receptor-like | 0.04929 | 5.89308 | FALSE | FALSE |
| LOC107064112 | titin | 0.05259 | 5.89944 | FALSE | FALSE |
| LOC107063631 | glutamyl aminopeptidase-like | 0.05087 | 5.9052 | FALSE | FALSE |
| LOC107072777 | uncharacterized protein LOC107072777 | 0.0501 | 5.91495 | FALSE | FALSE |
| LOC107073208 | peroxisomal biogenesis factor 3 | 0.05047 | 5.92323 | FALSE | FALSE |
| LOC107067476 | uncharacterized protein LOC107067476 | 0.05055 | 5.92753 | FALSE | FALSE |
| LOC107071041 | transcription factor CP2-like protein 1 | 0.05051 | 5.93114 | FALSE | FALSE |
| LOC107066374 | uncharacterized protein LOC107066374 | 0.05129 | 5.93346 | FALSE | FALSE |
| LOC107068016 | uncharacterized protein LOC107068016 | 0.05319 | 5.93638 | FALSE | FALSE |
| LOC107067858 | uncharacterized protein LOC107067858 | 0.05157 | 5.9493 | FALSE | FALSE |
| LOC107070483 | tyrosine-protein phosphatase corkscrew | 0.05106 | 5.94997 | FALSE | FALSE |
| LOC107065384 | FK506-binding protein 59-like | 0.05029 | 5.95343 | FALSE | FALSE |

| | | | | | |
|---|---|---|---|---|---|
| LOC107073195 | GATA zinc finger domain-containing protein | 0.05173 | 5.96009 | FALSE | FALSE |
| LOC107072829 | solute carrier organic anion transporter fam | 0.05292 | 5.96209 | FALSE | FALSE |
| LOC107072123 | uncharacterized protein LOC107072123 | 0.05151 | 5.96672 | FALSE | FALSE |
| LOC107073616 | uncharacterized protein LOC107073616 | 0.04952 | 5.96834 | FALSE | FALSE |
| LOC107071714 | uncharacterized protein LOC107071714 | 0.05055 | 5.98211 | FALSE | FALSE |
| LOC107069112 | deleted in autism protein 1 homolog | 0.05084 | 5.98243 | FALSE | FALSE |
| LOC107066512 | serine hydroxymethyltransferase, cytosolic | 0.05062 | 5.98492 | FALSE | FALSE |
| LOC107066121 | sodium-dependent nutrient amino acid tran | 0.05007 | 5.98801 | FALSE | FALSE |
| LOC107073721 | ATP-dependent RNA helicase vasa, isoform | 0.05102 | 6.00592 | FALSE | FALSE |
| LOC107074106 | pleckstrin homology domain-containing fam | 0.04996 | 6.01621 | FALSE | FALSE |
| LOC107072751 | DNA-directed RNA polymerase I subunit RP/ | 0.05035 | 6.01817 | FALSE | FALSE |
| LOC107063789 | uncharacterized protein LOC107063789 | 0.04942 | 6.02709 | FALSE | FALSE |
| LOC107065084 | peroxisomal hydratase-dehydrogenase-epir | 0.05038 | 6.03168 | FALSE | FALSE |
| LOC107069512 | DNA fragmentation factor subunit alpha-like | 0.05021 | 6.06566 | FALSE | FALSE |
| LOC107074242 | A-kinase anchor protein 17A | 0.05138 | 6.08538 | FALSE | FALSE |
| LOC107073586 | NAD-dependent protein deacylase Sirt4 | 0.05137 | 6.08836 | FALSE | FALSE |
| LOC107069304 | brain-specific homeobox protein homolog | 0.05145 | 6.09293 | FALSE | FALSE |
| LOC107071455 | probable phospholipid-transporting ATPase | 0.04946 | 6.09645 | FALSE | FALSE |
| LOC107066636 | protein FAM73B | 0.05056 | 6.10822 | FALSE | FALSE |
| LOC107069388 | prohormone-1-like | 0.05073 | 6.11342 | FALSE | FALSE |
| LOC107069365 | paired box protein Pax-6-like | 0.05324 | 6.12178 | FALSE | FALSE |
| LOC107064342 | protein abrupt-like | 0.05213 | 6.14874 | FALSE | FALSE |
| LOC107072592 | pyruvate dehydrogenase E1 component sub | 0.05019 | 6.17412 | FALSE | FALSE |
| LOC107063728 | nucleoprotein TPR-like | 0.05329 | 6.17917 | FALSE | FALSE |
| LOC107072071 | thioredoxin, mitochondrial | 0.05189 | 6.18123 | FALSE | FALSE |
| LOC107067310 | uncharacterized protein LOC107067310 | 0.04927 | 6.20872 | FALSE | FALSE |
| LOC107067598 | uncharacterized protein LOC107067598 | 0.05159 | 6.20957 | FALSE | FALSE |
| LOC107069652 | ER membrane protein complex subunit 2-lik | 0.05014 | 6.21345 | FALSE | FALSE |
| LOC107066311 | inosine triphosphate pyrophosphatase | 0.05025 | 6.21952 | FALSE | FALSE |
| LOC107069005 | mediator of RNA polymerase II transcriptior | 0.05075 | 6.22715 | FALSE | FALSE |
| LOC107065203 | uncharacterized protein LOC107065203 | 0.0491 | 6.22735 | FALSE | FALSE |
| LOC107065036 | uncharacterized protein LOC107065036 | 0.04864 | 6.25053 | FALSE | FALSE |
| LOC107064403 | alpha,alpha-trehalose-phosphate synthase [ | 0.05004 | 6.27291 | FALSE | FALSE |
| LOC107066919 | alpha-ketoglutarate-dependent dioxygenase | 0.04992 | 6.28452 | FALSE | FALSE |
| LOC107066708 | angiotensin-converting enzyme-like | 0.0501 | 6.29006 | FALSE | FALSE |
| LOC107066620 | esterase E4-like | 0.05163 | 6.30256 | FALSE | FALSE |
| LOC107064636 | T-box transcription factor TBX20-like | 0.05091 | 6.30354 | FALSE | FALSE |
| LOC107067277 | lysine-specific demethylase 8 | 0.04968 | 6.32091 | FALSE | FALSE |
| LOC107073273 | cell division cycle 5-like protein | 0.0522 | 6.3258 | FALSE | FALSE |
| LOC107069004 | uncharacterized protein LOC107069004 | 0.04915 | 6.34627 | FALSE | FALSE |
| LOC107064793 | probable serine/threonine-protein kinase D | 0.04873 | 6.35668 | FALSE | FALSE |
| LOC107068005 | probable serine/threonine-protein kinase D | 0.05087 | 6.35756 | FALSE | FALSE |
| LOC107067416 | cell division cycle protein 23 homolog | 0.05171 | 6.35899 | FALSE | FALSE |
| LOC107074092 | uncharacterized protein LOC107074092 | 0.05029 | 6.36636 | FALSE | FALSE |
| LOC107070552 | E3 ubiquitin-protein ligase Su(dx) | 0.0503 | 6.39331 | FALSE | FALSE |
| LOC107069953 | proteasome subunit alpha type-3 | 0.04934 | 6.41034 | FALSE | FALSE |
| LOC107064542 | cytochrome c | 0.05063 | 6.4111 | FALSE | FALSE |
| LOC107073824 | poly(U)-specific endoribonuclease homolog | 0.04999 | 6.41535 | FALSE | FALSE |
| LOC107066694 | histone H2B.v2 | 0.05063 | 6.41603 | FALSE | FALSE |
| LOC107068927 | uncharacterized protein LOC107068927 | 0.05134 | 6.42197 | FALSE | FALSE |

| | | | | | |
|---|---|---|---|---|---|
| LOC107067077 | U3 small nucleolar RNA-associated protein 1 | 0.05076 | 6.42295 | FALSE | FALSE |
| LOC107064180 | malectin-A | 0.0497 | 6.46792 | FALSE | FALSE |
| LOC107070797 | DNA topoisomerase 3-alpha | 0.05217 | 6.50399 | FALSE | FALSE |
| LOC107066628 | A disintegrin and metalloproteinase with th | 0.05084 | 6.50442 | FALSE | FALSE |
| LOC107065053 | neprilysin-like | 0.05136 | 6.51308 | FALSE | FALSE |
| LOC107066285 | golgin subfamily A member 7 | 0.04852 | 6.51907 | FALSE | FALSE |
| LOC107071501 | multiple epidermal growth factor-like doma | 0.04942 | 6.52669 | FALSE | FALSE |
| LOC107065111 | uncharacterized protein LOC107065111 | 0.04941 | 6.52953 | FALSE | FALSE |
| LOC107065691 | myeloid differentiation primary response pr | 0.04998 | 6.53293 | FALSE | FALSE |
| LOC107069801 | uncharacterized protein LOC107069801 | 0.052 | 6.53582 | FALSE | FALSE |
| LOC107071647 | myosin-IB | 0.05217 | 6.53765 | FALSE | FALSE |
| LOC107074255 | uncharacterized protein LOC107074255 | 0.0495 | 6.56498 | FALSE | FALSE |
| LOC107073843 | ragulator complex protein LAMTOR4 homol | 0.05097 | 6.56504 | FALSE | FALSE |
| LOC107070418 | WD repeat-containing protein 19 | 0.04838 | 6.5762 | FALSE | FALSE |
| LOC107065387 | uncharacterized protein LOC107065387 | 0.05064 | 6.5777 | FALSE | FALSE |
| LOC107069146 | phospholipase B1, membrane-associated-lik | 0.05086 | 6.57808 | FALSE | FALSE |
| LOC107073827 | INO80 complex subunit B | 0.05089 | 6.58785 | FALSE | FALSE |
| LOC107072818 | CBP80/20-dependent translation initiation f | 0.05065 | 6.59513 | FALSE | FALSE |
| LOC107065983 | katanin p60 ATPase-containing subunit A-lik | 0.05072 | 6.59886 | FALSE | FALSE |
| LOC107071347 | mitochondrial tRNA-specific 2-thiouridylase | 0.04937 | 6.60011 | FALSE | FALSE |
| LOC107068699 | LOW QUALITY PROTEIN: endocuticle structu | 0.05081 | 6.6041 | FALSE | FALSE |
| LOC107072120 | integrin beta-nu-like | 0.05154 | 6.60683 | FALSE | FALSE |
| LOC107070862 | carbonic anhydrase 1-like | 0.04966 | 6.60903 | FALSE | FALSE |
| LOC107063780 | uncharacterized protein LOC107063780 | 0.05046 | 6.63471 | FALSE | FALSE |
| LOC107072068 | uncharacterized protein LOC107072068 | 0.0488 | 6.65052 | FALSE | FALSE |
| LOC107067056 | uncharacterized protein LOC107067056 | 0.05142 | 6.66692 | FALSE | FALSE |
| LOC107064070 | calmodulin-lysine N-methyltransferase | 0.04962 | 6.67821 | FALSE | FALSE |
| LOC107069053 | uncharacterized protein LOC107069053 | 0.05044 | 6.68921 | FALSE | FALSE |
| LOC107066535 | UNC93-like protein | 0.04916 | 6.69182 | FALSE | FALSE |
| LOC107063724 | RNA-binding protein Nova-1 | 0.0501 | 6.70122 | FALSE | FALSE |
| LOC107070846 | forkhead box protein J1-B-like | 0.0513 | 6.71391 | FALSE | FALSE |
| LOC107074113 | importin-5 | 0.05023 | 6.71746 | FALSE | FALSE |
| LOC107071940 | ethanolamine kinase 1 | 0.05044 | 6.77316 | FALSE | FALSE |
| LOC107073340 | putative nuclease HARBI1 | 0.05127 | 6.77439 | FALSE | FALSE |
| LOC107064018 | phosphoserine phosphatase | 0.05268 | 6.78937 | FALSE | FALSE |
| LOC107064877 | E3 ubiquitin-protein ligase MYLIP | 0.04992 | 6.79087 | FALSE | FALSE |
| LOC107073483 | histone deacetylase complex subunit SAP18 | 0.05088 | 6.79701 | FALSE | FALSE |
| LOC107064128 | gamma-aminobutyric acid receptor alpha-lil | 0.05147 | 6.80195 | FALSE | FALSE |
| LOC107066511 | endothelin-converting enzyme 1 | 0.05015 | 6.8149 | FALSE | FALSE |
| LOC107072586 | uncharacterized protein LOC107072586 | 0.04904 | 6.82096 | FALSE | FALSE |
| LOC107068657 | U3 small nucleolar ribonucleoprotein protei | 0.05002 | 6.82588 | FALSE | FALSE |
| LOC107066135 | protein asunder homolog | 0.05011 | 6.82831 | FALSE | FALSE |
| LOC107074672 | uncharacterized protein DDB_G0271670-lik | 0.05001 | 6.83116 | FALSE | FALSE |
| LOC107073200 | COP9 signalosome complex subunit 2 | 0.05122 | 6.83429 | FALSE | FALSE |
| LOC107063803 | uncharacterized protein LOC107063803 | 0.05039 | 6.83704 | FALSE | FALSE |
| LOC107063627 | uncharacterized protein LOC107063627 | 0.04993 | 6.86168 | FALSE | FALSE |
| LOC107065702 | uncharacterized protein LOC107065702 | 0.0495 | 6.87075 | FALSE | FALSE |
| LOC107073169 | transcription termination factor 3, mitochor | 0.05028 | 6.87503 | FALSE | FALSE |
| LOC107068916 | protein D2-like | 0.04948 | 6.87813 | FALSE | FALSE |
| LOC107068661 | aurora kinase B-like | 0.0511 | 6.87831 | FALSE | FALSE |

| | | | | | |
|---|---|---|---|---|---|
| LOC107070951 | THO complex subunit 3 | 0.05016 | 6.88466 | FALSE | FALSE |
| LOC107065161 | cytochrome c1, heme protein, mitochondria | 0.05079 | 6.89628 | FALSE | FALSE |
| LOC107064354 | death-associated protein kinase related-like | 0.05192 | 6.90412 | FALSE | FALSE |
| LOC107072038 | uncharacterized protein LOC107072038 | 0.05055 | 6.91865 | FALSE | FALSE |
| LOC107068454 | glucose dehydrogenase [FAD, quinone | 0.0502 | 6.95995 | FALSE | FALSE |
| LOC107066975 | putative uncharacterized protein DDB_G029 | 0.05072 | 6.96036 | FALSE | FALSE |
| LOC107070196 | cystathionine gamma-lyase | 0.05066 | 6.97976 | FALSE | FALSE |
| LOC107063964 | uncharacterized protein LOC107063964 | 0.04912 | 7.00537 | FALSE | FALSE |
| LOC107068307 | 60S ribosomal protein L23a-like | 0.04958 | 7.02225 | FALSE | FALSE |
| LOC107064032 | uncharacterized protein LOC107064032 | 0.04862 | 7.02354 | FALSE | FALSE |
| LOC107070424 | tectonin beta-propeller repeat-containing p | 0.05065 | 7.03961 | FALSE | FALSE |
| LOC107070346 | glutamate receptor 1 | 0.05075 | 7.0532 | FALSE | FALSE |
| LOC107073910 | elongation factor 1-beta' | 0.05191 | 7.0619 | FALSE | FALSE |
| LOC107069789 | synaptotagmin 1 | 0.04902 | 7.06554 | FALSE | FALSE |
| LOC107065363 | uncharacterized protein LOC107065363 | 0.04882 | 7.07494 | FALSE | FALSE |
| LOC107068547 | uncharacterized protein LOC107068547 | 0.04862 | 7.09717 | FALSE | FALSE |
| LOC107063965 | protein enabled homolog | 0.04878 | 7.10226 | FALSE | FALSE |
| LOC107073079 | nucleoporin p58/p45 | 0.04885 | 7.10536 | FALSE | FALSE |
| LOC107063673 | ubinuclein-1 | 0.05145 | 7.11324 | FALSE | FALSE |
| LOC107073502 | ADAMTS-like protein 1 | 0.04991 | 7.11745 | FALSE | FALSE |
| LOC107074624 | uncharacterized protein LOC107074624 | 0.04841 | 7.12444 | FALSE | FALSE |
| LOC107065309 | amidophosphoribosyltransferase-like | 0.04743 | 7.13253 | FALSE | FALSE |
| LOC107069344 | recQ-mediated genome instability protein 1 | 0.04976 | 7.13832 | FALSE | FALSE |
| LOC107074555 | protein unc-13 homolog B | 0.04869 | 7.14017 | FALSE | FALSE |
| LOC107068054 | serine-rich adhesin for platelets-like | 0.05023 | 7.14597 | FALSE | FALSE |
| LOC107063727 | uncharacterized protein LOC107063727 | 0.05067 | 7.1659 | FALSE | FALSE |
| LOC107072925 | host cell factor 2-like | 0.0521 | 7.18628 | FALSE | FALSE |
| LOC107070781 | uncharacterized protein LOC107070781 | 0.04996 | 7.18964 | FALSE | FALSE |
| LOC107067781 | uncharacterized protein LOC107067781 | 0.04923 | 7.19121 | FALSE | FALSE |
| LOC107065089 | flap endonuclease GEN | 0.04884 | 7.1961 | FALSE | FALSE |
| LOC107065325 | ATP synthase mitochondrial F1 complex ass | 0.05115 | 7.19763 | FALSE | FALSE |
| LOC107066707 | translocating chain-associated membrane p | 0.0504 | 7.20035 | FALSE | FALSE |
| LOC107064408 | xyloside xylosyltransferase 1 | 0.05186 | 7.20547 | FALSE | FALSE |
| LOC107073593 | cytochrome P450 4C1-like | 0.0497 | 7.20553 | FALSE | FALSE |
| LOC107071527 | D-tyrosyl-tRNA(Tyr) deacylase 1 | 0.05173 | 7.20776 | FALSE | FALSE |
| LOC107065773 | GATA zinc finger domain-containing protein | 0.05038 | 7.20974 | FALSE | FALSE |
| LOC107074040 | dynein heavy chain 2, axonemal | 0.04897 | 7.2136 | FALSE | FALSE |
| LOC107070583 | lachesin-like | 0.04952 | 7.21572 | FALSE | FALSE |
| LOC107064357 | potassium voltage-gated channel subfamily | 0.05044 | 7.21995 | FALSE | FALSE |
| LOC107063721 | glycogen debranching enzyme | 0.0501 | 7.22306 | FALSE | FALSE |
| LOC107069175 | bone morphogenetic protein 1 | 0.04967 | 7.22584 | FALSE | FALSE |
| LOC107065835 | otoferlin-like | 0.04886 | 7.2261 | FALSE | FALSE |
| LOC107071463 | uncharacterized protein LOC107071463 | 0.05273 | 7.26088 | FALSE | FALSE |
| LOC107074373 | serine/threonine-protein kinase polo-like | 0.04971 | 7.27639 | FALSE | FALSE |
| LOC107074273 | uncharacterized protein LOC107074273 | 0.04938 | 7.27857 | FALSE | FALSE |
| LOC107071329 | muscarinic acetylcholine receptor DM1 | 0.05142 | 7.29755 | FALSE | FALSE |
| LOC107068578 | probable phenylalanine--tRNA ligase, mitocl | 0.04922 | 7.29988 | FALSE | FALSE |
| LOC107064780 | elongator complex protein 4 | 0.05002 | 7.30656 | FALSE | FALSE |
| LOC107074244 | uncharacterized protein LOC107074244 | 0.04963 | 7.30938 | FALSE | FALSE |
| LOC107068717 | transmembrane protein 181 | 0.04793 | 7.33447 | FALSE | FALSE |

| | | | | | |
|---|---|---|---|---|---|
| LOC107073845 | 40S ribosomal protein S27-like | 0.04978 | 7.33605 | FALSE | FALSE |
| LOC107064751 | uncharacterized protein LOC107064751 | 0.05036 | 7.33644 | FALSE | FALSE |
| LOC107064298 | adenosine kinase 1 | 0.04882 | 7.34205 | FALSE | FALSE |
| LOC107067191 | geranylgeranyl transferase type-2 subunit b | 0.05036 | 7.34995 | FALSE | FALSE |
| LOC107070978 | U11/U12 small nuclear ribonucleoprotein 3! | 0.05113 | 7.36112 | FALSE | FALSE |
| LOC107072937 | golgin subfamily A member 4-like | 0.05076 | 7.36418 | FALSE | FALSE |
| LOC107074205 | uncharacterized protein LOC107074205 | 0.04874 | 7.36626 | FALSE | FALSE |
| LOC107072582 | tetratricopeptide repeat protein 19 homolo | 0.04968 | 7.37519 | FALSE | FALSE |
| LOC107071184 | serine/threonine-protein kinase pakA | 0.05084 | 7.38148 | FALSE | FALSE |
| LOC107064762 | uncharacterized protein LOC107064762 | 0.04993 | 7.38302 | FALSE | FALSE |
| LOC107074260 | KRR1 small subunit processome component | 0.05009 | 7.38512 | FALSE | FALSE |
| LOC107068724 | zinc finger HIT domain-containing protein 2 | 0.04975 | 7.3926 | FALSE | FALSE |
| LOC107070947 | zinc finger protein 616-like | 0.05205 | 7.40042 | FALSE | FALSE |
| LOC107074351 | bifunctional purine biosynthesis protein PUI | 0.04969 | 7.40231 | FALSE | FALSE |
| LOC107064059 | tyrosine-protein phosphatase non-receptor | 0.05042 | 7.40586 | FALSE | FALSE |
| LOC107072544 | UBA-like domain-containing protein 2 | 0.05136 | 7.40707 | FALSE | FALSE |
| LOC107067716 | uncharacterized protein LOC107067716 | 0.04999 | 7.40781 | FALSE | FALSE |
| LOC107063835 | IQ motif-containing protein H-like | 0.0489 | 7.42356 | FALSE | FALSE |
| LOC107069080 | serine/threonine-protein phosphatase PP1- | 0.04982 | 7.44583 | FALSE | FALSE |
| LOC107066146 | DNA polymerase alpha catalytic subunit-like | 0.04845 | 7.44676 | FALSE | FALSE |
| LOC107068228 | tyrosine-protein phosphatase 10D | 0.04897 | 7.44937 | FALSE | FALSE |
| LOC107064379 | 60S ribosomal protein L34-like | 0.04886 | 7.4524 | FALSE | FALSE |
| LOC107068323 | eukaryotic peptide chain release factor subu | 0.04999 | 7.45714 | FALSE | FALSE |
| LOC107064185 | tRNA dimethylallyltransferase, mitochondri | 0.0496 | 7.46631 | FALSE | FALSE |
| LOC107074396 | uncharacterized protein LOC107074396 | 0.04969 | 7.47418 | FALSE | FALSE |
| LOC107069819 | uncharacterized protein CG43867 | 0.05031 | 7.47753 | FALSE | FALSE |
| LOC107069171 | vesicle-fusing ATPase 1-like | 0.04982 | 7.48108 | FALSE | FALSE |
| LOC107065528 | mitogen-activated protein kinase 14B-like | 0.04817 | 7.50366 | FALSE | FALSE |
| LOC107067274 | UPF0668 protein C10orf76 homolog | 0.05083 | 7.50638 | FALSE | FALSE |
| LOC107065278 | inositol monophosphatase 1-like | 0.04884 | 7.52485 | FALSE | FALSE |
| LOC107065272 | uncharacterized protein LOC107065272 | 0.04938 | 7.53147 | FALSE | FALSE |
| LOC107071695 | laccase-like | 0.04945 | 7.53631 | FALSE | FALSE |
| LOC107065209 | helicase domino | 0.0495 | 7.54032 | FALSE | FALSE |
| LOC107064286 | protein FAM60A | 0.04842 | 7.54157 | FALSE | FALSE |
| LOC107065840 | 40S ribosomal protein S12 | 0.05015 | 7.543 | FALSE | FALSE |
| LOC107071893 | pre-mRNA-splicing factor 38A | 0.04844 | 7.55261 | FALSE | FALSE |
| LOC107066261 | probable WRKY transcription factor protein | 0.05069 | 7.55311 | FALSE | FALSE |
| LOC107072135 | tRNA (cytosine(38)-C(5))-methyltransferase | 0.04908 | 7.56154 | FALSE | FALSE |
| LOC107069337 | ubiquitin carboxyl-terminal hydrolase 8-like | 0.04951 | 7.56968 | FALSE | FALSE |
| LOC107072225 | dnaJ homolog subfamily C member 1 | 0.04889 | 7.57236 | FALSE | FALSE |
| LOC107070272 | uncharacterized protein LOC107070272 | 0.05006 | 7.57421 | FALSE | FALSE |
| LOC107067116 | rac GTPase-activating protein 1-like | 0.04952 | 7.57447 | FALSE | FALSE |
| LOC107073810 | voltage-dependent T-type calcium channel s | 0.0499 | 7.57899 | FALSE | FALSE |
| LOC107070201 | 6-phosphogluconate dehydrogenase, decarl | 0.05018 | 7.5808 | FALSE | FALSE |
| LOC107072608 | T-complex protein 1 subunit delta | 0.04958 | 7.58313 | FALSE | FALSE |
| LOC107074443 | serine/threonine-protein phosphatase PGAI | 0.05076 | 7.5997 | FALSE | FALSE |
| LOC107073363 | pre-mRNA-splicing regulator WTAP-like | 0.05065 | 7.61724 | FALSE | FALSE |
| LOC107071781 | rhomboid-related protein 4-like | 0.04865 | 7.6258 | FALSE | FALSE |
| LOC107074135 | COMM domain-containing protein 10-like | 0.05129 | 7.63213 | FALSE | FALSE |
| LOC107070904 | disintegrin and metalloproteinase domain-c | 0.05015 | 7.65285 | FALSE | FALSE |

| | | | | | |
|---|---|---|---|---|---|
| LOC107067677 | protein FAM13A | 0.04886 | 7.66763 | FALSE | FALSE |
| LOC107073889 | lipid storage droplets surface-binding protei | 0.05008 | 7.67047 | FALSE | FALSE |
| LOC107071844 | histone-lysine N-methyltransferase SETD1 | 0.04909 | 7.68453 | FALSE | FALSE |
| LOC107067315 | lysosomal alpha-mannosidase | 0.04858 | 7.69159 | FALSE | FALSE |
| LOC107071645 | AP-1 complex subunit mu-1-like | 0.0507 | 7.71099 | FALSE | FALSE |
| LOC107074162 | rho GTPase-activating protein 190 | 0.04916 | 7.71768 | FALSE | FALSE |
| LOC107068122 | pyruvate dehydrogenase phosphatase regul | 0.05017 | 7.76168 | FALSE | FALSE |
| LOC107069688 | translation initiation factor eIF-2B subunit a | 0.04931 | 7.76323 | FALSE | FALSE |
| LOC107068755 | piggyBac transposable element-derived pro | 0.05021 | 7.77723 | FALSE | FALSE |
| LOC107071780 | prostaglandin E synthase 2 | 0.05009 | 7.77973 | FALSE | FALSE |
| LOC107072304 | uncharacterized protein LOC107072304 | 0.05049 | 7.7894 | FALSE | FALSE |
| LOC107071636 | transcription factor Dp-1 | 0.04867 | 7.79372 | FALSE | FALSE |
| LOC107074343 | another transcription unit protein | 0.05054 | 7.79542 | FALSE | FALSE |
| LOC107068816 | bone morphogenetic protein 3-like | 0.05005 | 7.79575 | FALSE | FALSE |
| LOC107072484 | THO complex subunit 4 | 0.04949 | 7.80126 | FALSE | FALSE |
| LOC107064815 | zinc finger protein 431-like | 0.04861 | 7.81539 | FALSE | FALSE |
| LOC107063677 | serine protease easter-like | 0.0498 | 7.85461 | FALSE | FALSE |
| LOC107066438 | monocarboxylate transporter 12 | 0.05015 | 7.86137 | FALSE | FALSE |
| LOC107066056 | protein hairless-like | 0.0486 | 7.8775 | FALSE | FALSE |
| LOC107068707 | uncharacterized protein LOC107068707 | 0.04707 | 7.88185 | FALSE | FALSE |
| LOC107072269 | ATPase family AAA domain-containing prote | 0.04918 | 7.88463 | FALSE | FALSE |
| LOC107070421 | poly [ADP-ribose | 0.04976 | 7.89147 | FALSE | FALSE |
| LOC107073163 | uncharacterized protein LOC107073163 | 0.051 | 7.89374 | FALSE | FALSE |
| LOC107068100 | uncharacterized protein LOC107068100 | 0.04943 | 7.89509 | FALSE | FALSE |
| LOC107070476 | zinc finger MYND domain-containing protei | 0.04854 | 7.9139 | FALSE | FALSE |
| LOC107071480 | uncharacterized protein LOC107071480 | 0.0499 | 7.91418 | FALSE | FALSE |
| LOC107063860 | uncharacterized protein LOC107063860 | 0.05249 | 7.91818 | FALSE | FALSE |
| LOC107071049 | elongation of very long chain fatty acids pro | 0.05098 | 7.91988 | FALSE | FALSE |
| LOC107067000 | FK506-binding protein 5-like | 0.05103 | 7.93099 | FALSE | FALSE |
| LOC107070532 | calmodulin | 0.04802 | 7.93945 | FALSE | FALSE |
| LOC107073209 | uncharacterized protein LOC107073209 | 0.05026 | 7.94382 | FALSE | FALSE |
| LOC107070783 | ATP-binding cassette sub-family G member | 0.04904 | 7.94693 | FALSE | FALSE |
| LOC107074546 | rhotekin-2-like | 0.04907 | 7.95234 | FALSE | FALSE |
| LOC107068032 | 4-hydroxybutyrate coenzyme A transferase- | 0.04953 | 7.95806 | FALSE | FALSE |
| LOC107070945 | methyltransferase-like protein 25 | 0.05056 | 7.97751 | FALSE | FALSE |
| LOC107064955 | plasma membrane calcium-transporting ATI | 0.04856 | 7.98519 | FALSE | FALSE |
| LOC107071811 | cyclic AMP response element-binding prote | 0.04827 | 7.98535 | FALSE | FALSE |
| LOC107070585 | alpha-(1,3)-fucosyltransferase C-like | 0.04709 | 8.00982 | FALSE | FALSE |
| LOC107067270 | uncharacterized protein LOC107067270 | 0.04806 | 8.01111 | FALSE | FALSE |
| LOC107073262 | uncharacterized protein LOC107073262 | 0.04922 | 8.01448 | FALSE | FALSE |
| LOC107072435 | neuroendocrine convertase 2 | 0.04912 | 8.01871 | FALSE | FALSE |
| LOC107069929 | ATPase family AAA domain-containing prote | 0.04987 | 8.02277 | FALSE | FALSE |
| LOC107070044 | protein TolA-like | 0.04976 | 8.03138 | FALSE | FALSE |
| LOC107070182 | protein ecdysoneless | 0.05061 | 8.04474 | FALSE | FALSE |
| LOC107070349 | general transcription factor IIE subunit 2 | 0.04874 | 8.04656 | FALSE | FALSE |
| LOC107068714 | nose resistant to fluoxetine protein 6-like | 0.04884 | 8.04693 | FALSE | FALSE |
| LOC107069721 | ribonuclease 3 | 0.04948 | 8.049 | FALSE | FALSE |
| LOC107074313 | uncharacterized protein LOC107074313 | 0.0491 | 8.05002 | FALSE | FALSE |
| LOC107064256 | CCR4-NOT transcription complex subunit 2 | 0.04975 | 8.05227 | FALSE | FALSE |
| LOC107067407 | uncharacterized protein LOC107067407 | 0.04971 | 8.05262 | FALSE | FALSE |

| | | | | | |
|---|---|---|---|---|---|
| LOC107065883 | uncharacterized protein LOC107065883 | 0.04819 | 8.06454 | FALSE | FALSE |
| LOC107069597 | 26S proteasome non-ATPase regulatory sub | 0.05064 | 8.07583 | FALSE | FALSE |
| LOC107071625 | NADH-ubiquinone oxidoreductase 75 kDa su | 0.04873 | 8.076 | FALSE | FALSE |
| LOC107066709 | fumarylacetoacetase | 0.04979 | 8.08311 | FALSE | FALSE |
| LOC107065095 | JNK-interacting protein 1 | 0.04907 | 8.09222 | FALSE | FALSE |
| LOC107066545 | 60S ribosomal protein L6-like | 0.0506 | 8.1059 | FALSE | FALSE |
| LOC107072054 | myotubularin-related protein 6 | 0.05059 | 8.10662 | FALSE | FALSE |
| LOC107070126 | diuretic hormone class 2 | 0.04956 | 8.11901 | FALSE | FALSE |
| LOC107072397 | lachesin-like | 0.04983 | 8.13963 | FALSE | FALSE |
| LOC107068101 | uncharacterized protein LOC107068101 | 0.04932 | 8.14066 | FALSE | FALSE |
| LOC107073950 | NAD-dependent protein deacetylase sirtuin- | 0.05048 | 8.15343 | FALSE | FALSE |
| LOC107067801 | protein transport protein Sec61 subunit gan | 0.0501 | 8.16544 | FALSE | FALSE |
| LOC107068605 | apidaecins type 22-like | 0.0507 | 8.17433 | FALSE | FALSE |
| LOC107070171 | protein yellow-like | 0.0502 | 8.18212 | FALSE | FALSE |
| LOC107069178 | methionine aminopeptidase 1 | 0.04971 | 8.19493 | FALSE | FALSE |
| LOC107066340 | Y+L amino acid transporter 2 | 0.04775 | 8.22074 | FALSE | FALSE |
| LOC107068126 | kinesin-like protein KIF23 | 0.04882 | 8.22087 | FALSE | FALSE |
| LOC107074626 | protein transport protein Sec24A | 0.049 | 8.22409 | FALSE | FALSE |
| LOC107068946 | glypican-6 | 0.04964 | 8.23679 | FALSE | FALSE |
| LOC107068903 | odorant receptor 13a-like | 0.05154 | 8.24569 | FALSE | FALSE |
| LOC107065058 | uncharacterized protein LOC107065058 | 0.04823 | 8.24974 | FALSE | FALSE |
| LOC107073997 | aldehyde dehydrogenase, mitochondrial | 0.04998 | 8.25222 | FALSE | FALSE |
| LOC107074349 | tRNA:m(4)X modification enzyme TRM13 hc | 0.05033 | 8.25824 | FALSE | FALSE |
| LOC107066537 | probable glucarate transporter | 0.04869 | 8.27334 | FALSE | FALSE |
| LOC107066605 | dynamin-1-like protein | 0.04924 | 8.2771 | FALSE | FALSE |
| LOC107065939 | nucleolar pre-ribosomal-associated protein | 0.04881 | 8.27722 | FALSE | FALSE |
| LOC107074664 | sulfotransferase 1C4-like | 0.04964 | 8.2893 | FALSE | FALSE |
| LOC107068196 | monocarboxylate transporter 10-like | 0.04834 | 8.32036 | FALSE | FALSE |
| LOC107072660 | glycine receptor subunit alpha-3 | 0.04677 | 8.32116 | FALSE | FALSE |
| LOC107073909 | pre-mRNA-splicing factor 38B | 0.04873 | 8.33418 | FALSE | FALSE |
| LOC107065424 | general transcription factor IIH subunit 3 | 0.04947 | 8.34249 | FALSE | FALSE |
| LOC107069390 | uncharacterized protein LOC107069390 | 0.04869 | 8.35287 | FALSE | FALSE |
| LOC107072930 | transient receptor potential cation channel | 0.04835 | 8.35699 | FALSE | FALSE |
| LOC107067216 | uncharacterized protein LOC107067216 | 0.05097 | 8.36045 | FALSE | FALSE |
| LOC107065635 | ATPase family AAA domain-containing prote | 0.04855 | 8.36458 | FALSE | FALSE |
| LOC107072027 | probable 26S proteasome non-ATPase regul | 0.04928 | 8.37872 | FALSE | FALSE |
| LOC107064563 | 5-aminolevulinate synthase, erythroid-speci | 0.04936 | 8.38491 | FALSE | FALSE |
| LOC107074305 | uncharacterized protein LOC107074305 | 0.04991 | 8.39983 | FALSE | FALSE |
| LOC107064437 | uncharacterized protein LOC107064437 | 0.04906 | 8.43983 | FALSE | FALSE |
| LOC107073278 | uncharacterized protein LOC107073278 | 0.04988 | 8.4446 | FALSE | FALSE |
| LOC107068756 | intraflagellar transport protein 43 homolog | 0.0478 | 8.44601 | FALSE | FALSE |
| LOC107073499 | uncharacterized protein LOC107073499 | 0.04908 | 8.44852 | FALSE | FALSE |
| LOC107065365 | teneurin-m | 0.04764 | 8.45976 | FALSE | FALSE |
| LOC107074147 | tyrosine-protein phosphatase 99A | 0.04923 | 8.46145 | FALSE | FALSE |
| LOC107068830 | 26S proteasome non-ATPase regulatory sub | 0.049 | 8.47167 | FALSE | FALSE |
| LOC107070155 | uncharacterized protein LOC107070155 | 0.0479 | 8.47305 | FALSE | FALSE |
| LOC107065385 | WD repeat-containing protein 61 | 0.04914 | 8.47442 | FALSE | FALSE |
| LOC107068062 | cullin-2 | 0.04739 | 8.48751 | FALSE | FALSE |
| LOC107074460 | uncharacterized protein LOC107074460 | 0.04856 | 8.48763 | FALSE | FALSE |
| LOC107073494 | poly(A) polymerase type 3 | 0.04961 | 8.49432 | FALSE | FALSE |

| | | | | | |
|---|---|---|---|---|---|
| LOC107070780 | TBC1 domain family member 15 | 0.0481 | 8.49711 | FALSE | FALSE |
| LOC107070686 | organic cation transporter protein | 0.05206 | 8.50523 | FALSE | FALSE |
| LOC107068821 | DNA replication complex GINS protein PSF1 | 0.04912 | 8.5206 | FALSE | FALSE |
| LOC107074237 | homeobox protein HMX1-like | 0.05112 | 8.53004 | FALSE | FALSE |
| LOC107072717 | serine/threonine-protein phosphatase 2A 6 | 0.04977 | 8.5431 | FALSE | FALSE |
| LOC107073000 | nuclear hormone receptor HR96 | 0.05127 | 8.54352 | FALSE | FALSE |
| LOC107068814 | putative fatty acyl-CoA reductase CG5065 | 0.051 | 8.56497 | FALSE | FALSE |
| LOC107074656 | uncharacterized protein LOC107074656 | 0.04832 | 8.58088 | FALSE | FALSE |
| LOC107072817 | location of vulva defective 1-like | 0.04875 | 8.58123 | FALSE | FALSE |
| LOC107066822 | glycerol-3-phosphate dehydrogenase, mitoc | 0.04919 | 8.58952 | FALSE | FALSE |
| LOC107064852 | uncharacterized protein LOC107064852 | 0.04926 | 8.59917 | FALSE | FALSE |
| LOC107069903 | protein rhomboid-like | 0.04937 | 8.59919 | FALSE | FALSE |
| LOC107072011 | uncharacterized protein LOC107072011 | 0.04908 | 8.60622 | FALSE | FALSE |
| LOC107069972 | gamma-glutamylcyclotransferase-like | 0.04985 | 8.60815 | FALSE | FALSE |
| LOC107066124 | protein-serine O-palmitoleoyltransferase pc | 0.05042 | 8.61524 | FALSE | FALSE |
| LOC107070428 | uncharacterized protein C6orf106 homolog | 0.05033 | 8.61568 | FALSE | FALSE |
| LOC107070882 | lateral signaling target protein 2 homolog | 0.04803 | 8.61587 | FALSE | FALSE |
| LOC107071816 | probable serine/threonine-protein kinase d | 0.04796 | 8.61845 | FALSE | FALSE |
| LOC107065000 | clavesin-2-like | 0.04822 | 8.61942 | FALSE | FALSE |
| LOC107069679 | serine/threonine-protein kinase RIO1-like | 0.04884 | 8.62736 | FALSE | FALSE |
| LOC107072501 | trafficking protein particle complex subunit | 0.05042 | 8.62879 | FALSE | FALSE |
| LOC107063824 | ruvB-like 2 | 0.0496 | 8.63906 | FALSE | FALSE |
| LOC107072458 | ras-related protein Rab-30-like | 0.04774 | 8.64318 | FALSE | FALSE |
| LOC107071072 | serine/threonine-protein phosphatase 4 reg | 0.05057 | 8.68742 | FALSE | FALSE |
| LOC107069475 | dual oxidase 1-like | 0.04971 | 8.69141 | FALSE | FALSE |
| LOC107068732 | trafficking protein particle complex subunit | 0.04907 | 8.69569 | FALSE | FALSE |
| LOC107068734 | uncharacterized protein LOC107068734 | 0.04882 | 8.69717 | FALSE | FALSE |
| LOC107070387 | A disintegrin and metalloproteinase with th | 0.04976 | 8.69905 | FALSE | FALSE |
| LOC107070482 | 60S ribosome subunit biogenesis protein NI | 0.04844 | 8.70209 | FALSE | FALSE |
| LOC107072779 | cytochrome P450 4C1-like | 0.04838 | 8.71397 | FALSE | FALSE |
| LOC107069690 | uncharacterized protein LOC107069690 | 0.04837 | 8.72403 | FALSE | FALSE |
| LOC107066702 | uncharacterized protein LOC107066702 | 0.05038 | 8.72585 | FALSE | FALSE |
| LOC107066615 | uncharacterized protein LOC107066615 | 0.04789 | 8.73199 | FALSE | FALSE |
| LOC107066923 | peptidyl-prolyl cis-trans isomerase-like 2 | 0.04908 | 8.73358 | FALSE | FALSE |
| LOC107064887 | sodium-coupled monocarboxylate transport | 0.04994 | 8.73395 | FALSE | FALSE |
| LOC107066196 | 39S ribosomal protein L22, mitochondrial | 0.04953 | 8.73534 | FALSE | FALSE |
| LOC107069655 | N-acetyltransferase ESCO1 | 0.04922 | 8.73679 | FALSE | FALSE |
| LOC107073659 | uncharacterized protein LOC107073659 | 0.04813 | 8.754 | FALSE | FALSE |
| LOC107073112 | BTB/POZ domain-containing protein KCTD5 | 0.04771 | 8.76385 | FALSE | FALSE |
| LOC107071997 | putative polypeptide N-acetylgalactosaminy | 0.04832 | 8.76751 | FALSE | FALSE |
| LOC107066524 | ATP-dependent Clp protease ATP-binding su | 0.04975 | 8.77846 | FALSE | FALSE |
| LOC107066227 | mitochondrial inner membrane protease su | 0.0507 | 8.78344 | FALSE | FALSE |
| LOC107064752 | head-specific guanylate cyclase | 0.04832 | 8.78463 | FALSE | FALSE |
| LOC107073229 | uncharacterized protein LOC107073229 | 0.0479 | 8.78647 | FALSE | FALSE |
| LOC107068133 | hyaluronidase | 0.0491 | 8.78695 | FALSE | FALSE |
| LOC107064614 | nicalin | 0.04864 | 8.79617 | FALSE | FALSE |
| LOC107074085 | 3-hydroxyisobutyryl-CoA hydrolase, mitoch | 0.048 | 8.80978 | FALSE | FALSE |
| LOC107069198 | ras-related protein Rab-27A | 0.04828 | 8.83496 | FALSE | FALSE |
| LOC107063856 | vacuolar protein sorting-associated protein | 0.04859 | 8.84045 | FALSE | FALSE |
| LOC107071441 | cationic amino acid transporter 2 | 0.04898 | 8.84639 | FALSE | FALSE |

| | | | | | |
|---|---|---|---|---|---|
| LOC107070225 | uncharacterized protein LOC107070225 | 0.04863 | 8.86398 | FALSE | FALSE |
| LOC107066064 | serine/threonine-protein kinase ULK3 | 0.04813 | 8.86741 | FALSE | FALSE |
| LOC107073670 | uncharacterized protein LOC107073670 | 0.04855 | 8.89381 | FALSE | FALSE |
| LOC107066830 | barH-like 2 homeobox protein | 0.05002 | 8.89562 | FALSE | FALSE |
| LOC107067275 | phosphatidylinositol 4-kinase type 2-alpha | 0.04871 | 8.91342 | FALSE | FALSE |
| LOC107065246 | pancreatic lipase-related protein 2-like | 0.04987 | 8.91744 | FALSE | FALSE |
| LOC107064979 | proteasome subunit beta type-1 | 0.04871 | 8.91745 | FALSE | FALSE |
| LOC107071285 | RNA pseudouridylate synthase domain-cont | 0.0485 | 8.9219 | FALSE | FALSE |
| LOC107072265 | uncharacterized protein LOC107072265 | 0.04886 | 8.9239 | FALSE | FALSE |
| LOC107073162 | phospholipid scramblase 1-like | 0.04974 | 8.92398 | FALSE | FALSE |
| LOC107072681 | uncharacterized protein LOC107072681 | 0.0498 | 8.94113 | FALSE | FALSE |
| LOC107069292 | transmembrane protein 62-like | 0.04841 | 8.95447 | FALSE | FALSE |
| LOC107070330 | polycomb protein eed-B-like | 0.04761 | 8.96018 | FALSE | FALSE |
| LOC107067650 | SH3 domain-containing RING finger protein | 0.04935 | 8.9608 | FALSE | FALSE |
| LOC107070410 | CREB-binding protein | 0.04944 | 8.96242 | FALSE | FALSE |
| LOC107068783 | protein grainyhead | 0.04891 | 8.96691 | FALSE | FALSE |
| LOC107066150 | RNA-binding protein 28-like | 0.04899 | 8.98511 | FALSE | FALSE |
| LOC107071160 | potassium channel subfamily K member 18- | 0.04933 | 9.00324 | FALSE | FALSE |
| LOC107066494 | uncharacterized protein LOC107066494 | 0.04797 | 9.00675 | FALSE | FALSE |
| LOC107074448 | serine-rich adhesin for platelets | 0.0486 | 9.00755 | FALSE | FALSE |
| LOC107073025 | zinc finger MIZ domain-containing protein 1 | 0.04768 | 9.02202 | FALSE | FALSE |
| LOC107071449 | uncharacterized protein LOC107071449 | 0.04916 | 9.02302 | FALSE | FALSE |
| LOC107065417 | putative serine protease K12H4.7 | 0.04746 | 9.0444 | FALSE | FALSE |
| LOC107073378 | ubiquitin thioesterase OTU1 | 0.04925 | 9.05147 | FALSE | FALSE |
| LOC107065117 | uncharacterized protein LOC107065117 | 0.04863 | 9.05363 | FALSE | FALSE |
| LOC107068373 | sphingomyelin phosphodiesterase | 0.04621 | 9.05444 | FALSE | FALSE |
| LOC107064617 | arrestin domain-containing protein 2-like | 0.04824 | 9.06756 | FALSE | FALSE |
| LOC107065431 | mitochondrial import inner membrane trans | 0.04764 | 9.07803 | FALSE | FALSE |
| LOC107066617 | pre-mRNA 3' end processing protein WDR33 | 0.04887 | 9.07928 | FALSE | FALSE |
| LOC107065937 | uncharacterized protein LOC107065937 | 0.04696 | 9.09683 | FALSE | FALSE |
| LOC107071204 | uncharacterized protein LOC107071204 | 0.04961 | 9.09992 | FALSE | FALSE |
| LOC107066732 | FK506-binding protein 15 | 0.04756 | 9.1025 | FALSE | FALSE |
| LOC107068984 | cytochrome b5 | 0.04855 | 9.10788 | FALSE | FALSE |
| LOC107069274 | U6 snRNA-associated Sm-like protein LSm7 | 0.04866 | 9.11915 | FALSE | FALSE |
| LOC107064321 | aminomethyltransferase, mitochondrial | 0.0478 | 9.14572 | FALSE | FALSE |
| LOC107068022 | WD repeat-containing protein 91 | 0.04775 | 9.14666 | FALSE | FALSE |
| LOC107064808 | 4-coumarate--CoA ligase 1-like | 0.0479 | 9.15197 | FALSE | FALSE |
| LOC107071640 | long-chain-fatty-acid--CoA ligase ACSBG2 | 0.0488 | 9.15921 | FALSE | FALSE |
| LOC107065968 | uncharacterized protein LOC107065968 | 0.05038 | 9.15924 | FALSE | FALSE |
| LOC107071961 | vesicle transport through interaction with t- | 0.04848 | 9.15946 | FALSE | FALSE |
| LOC107070649 | ethanolamine-phosphate cytidylyltransferas | 0.04898 | 9.17309 | FALSE | FALSE |
| LOC107063981 | bifunctional lysine-specific demethylase and | 0.04746 | 9.17633 | FALSE | FALSE |
| LOC107073394 | DNA polymerase alpha catalytic subunit | 0.04763 | 9.18572 | FALSE | FALSE |
| LOC107066885 | palmitoleoyl-protein carboxylesterase NOTU | 0.04747 | 9.19256 | FALSE | FALSE |
| LOC107074312 | ubiquitin carboxyl-terminal hydrolase 22 | 0.04814 | 9.20733 | FALSE | FALSE |
| LOC107069809 | sodium channel protein Nach-like | 0.04799 | 9.22299 | FALSE | FALSE |
| LOC107063870 | AN1-type zinc finger protein 5 | 0.04903 | 9.2247 | FALSE | FALSE |
| LOC107064105 | probable protein phosphatase 2C T23F11.1 | 0.04838 | 9.23632 | FALSE | FALSE |
| LOC107070706 | protein amnionless-like | 0.04949 | 9.23914 | FALSE | FALSE |
| LOC107074364 | translocator protein | 0.04896 | 9.24229 | FALSE | FALSE |

| | | | | | |
|---|---|---|---|---|---|
| LOC107065376 | UPF0428 protein CXorf56 homolog | 0.04839 | 9.26391 | FALSE | FALSE |
| LOC107069991 | mediator of RNA polymerase II transcriptior | 0.04849 | 9.26962 | FALSE | FALSE |
| LOC107065857 | cleft lip and palate transmembrane protein | 0.04788 | 9.27876 | FALSE | FALSE |
| LOC107066931 | CD151 antigen-like | 0.0478 | 9.28464 | FALSE | FALSE |
| LOC107071316 | uncharacterized protein LOC107071316 | 0.04823 | 9.28645 | FALSE | FALSE |
| LOC107065910 | histone chaperone asf1 | 0.04792 | 9.29327 | FALSE | FALSE |
| LOC107065154 | protein DDI1 homolog 2 | 0.04779 | 9.302 | FALSE | FALSE |
| LOC107069069 | UPF0536 protein C12orf66 homolog | 0.04843 | 9.30282 | FALSE | FALSE |
| LOC107072590 | acidic fibroblast growth factor intracellular-l | 0.04868 | 9.30373 | FALSE | FALSE |
| LOC107074114 | uncharacterized protein LOC107074114 | 0.04842 | 9.31121 | FALSE | FALSE |
| LOC107064994 | F-box only protein 11 | 0.04838 | 9.31138 | FALSE | FALSE |
| LOC107066570 | lipoyl synthase, mitochondrial | 0.04769 | 9.31554 | FALSE | FALSE |
| LOC107065310 | multifunctional protein ADE2 | 0.04835 | 9.32006 | FALSE | FALSE |
| LOC107073218 | uncharacterized protein LOC107073218 | 0.04871 | 9.33359 | FALSE | FALSE |
| LOC107073546 | purine nucleoside phosphorylase | 0.04716 | 9.34424 | FALSE | FALSE |
| LOC107064347 | E3 ubiquitin-protein ligase RBBP6 | 0.05021 | 9.37078 | FALSE | FALSE |
| LOC107071929 | serine/threonine-protein kinase polo | 0.05124 | 9.39933 | FALSE | FALSE |
| LOC107064978 | 39S ribosomal protein L46, mitochondrial | 0.04826 | 9.40513 | FALSE | FALSE |
| LOC107065612 | 60 kDa heat shock protein, mitochondrial-lil | 0.04635 | 9.407 | FALSE | FALSE |
| LOC107065793 | DNA-directed RNA polymerase II subunit RP | 0.04983 | 9.41926 | FALSE | FALSE |
| LOC107067595 | shootin-1 | 0.04649 | 9.42525 | FALSE | FALSE |
| LOC107064271 | chloride intracellular channel exc-4 | 0.04874 | 9.42726 | FALSE | FALSE |
| LOC107064373 | structural maintenance of chromosomes pr | 0.04643 | 9.43135 | FALSE | FALSE |
| LOC107074673 | endothelial lipase | 0.04824 | 9.43374 | FALSE | FALSE |
| LOC107063991 | ephrin-B2 | 0.05036 | 9.43918 | FALSE | FALSE |
| LOC107072209 | homeotic protein empty spiracles-like | 0.04877 | 9.44155 | FALSE | FALSE |
| LOC107070867 | uncharacterized protein LOC107070867 | 0.04799 | 9.4465 | FALSE | FALSE |
| LOC107072420 | uncharacterized protein LOC107072420 | 0.04814 | 9.44993 | FALSE | FALSE |
| LOC107064846 | trace amine-associated receptor 9 | 0.04928 | 9.45391 | FALSE | FALSE |
| LOC107067997 | pickpocket protein 28-like | 0.04817 | 9.46354 | FALSE | FALSE |
| LOC107067835 | uncharacterized protein LOC107067835 | 0.04746 | 9.47827 | FALSE | FALSE |
| LOC107071319 | putative glutamate synthase [NADPH | 0.04913 | 9.47866 | FALSE | FALSE |
| LOC107071772 | lipopolysaccharide-induced tumor necrosis | 0.04691 | 9.48391 | FALSE | FALSE |
| LOC107074404 | ras-like protein 2 | 0.04819 | 9.48735 | FALSE | FALSE |
| LOC107074202 | hemicentin-1 | 0.04809 | 9.49031 | FALSE | FALSE |
| LOC107074236 | glutamate receptor ionotropic, delta-1-like | 0.04851 | 9.50298 | FALSE | FALSE |
| LOC107064854 | acidic leucine-rich nuclear phosphoprotein 3 | 0.04748 | 9.51172 | FALSE | FALSE |
| LOC107073562 | uncharacterized protein LOC107073562 | 0.04993 | 9.51702 | FALSE | FALSE |
| LOC107065975 | UDP-glucuronosyltransferase 2C1-like | 0.04708 | 9.52973 | FALSE | FALSE |
| LOC107065013 | membrane metallo-endopeptidase-like 1 | 0.04964 | 9.5299 | FALSE | FALSE |
| LOC107072724 | voltage-dependent calcium channel subunit | 0.04835 | 9.53372 | FALSE | FALSE |
| LOC107065237 | uncharacterized protein LOC107065237 | 0.04745 | 9.53856 | FALSE | FALSE |
| LOC107066077 | xaa-Pro aminopeptidase 1-like | 0.04642 | 9.54 | FALSE | FALSE |
| Trnak-cuu | Trnak-cuu | 0.04944 | 9.54293 | FALSE | FALSE |
| LOC107071942 | DTW domain-containing protein 2 | 0.05016 | 9.54957 | FALSE | FALSE |
| LOC107065754 | ragulator complex protein LAMTOR3-A | 0.04822 | 9.55499 | FALSE | FALSE |
| LOC107064600 | anaphase-promoting complex subunit 1 | 0.04953 | 9.55576 | FALSE | FALSE |
| LOC107066328 | abhydrolase domain-containing protein 2 | 0.04795 | 9.55666 | FALSE | FALSE |
| LOC107071832 | la-related protein 1-like | 0.04845 | 9.57304 | FALSE | FALSE |
| LOC107066106 | H(+)/Cl(-) exchange transporter 7 | 0.04834 | 9.57468 | FALSE | FALSE |

| | | | | | |
|---|---|---|---|---|---|
| LOC107070053 | zinc transporter ZIP9-A | 0.04757 | 9.58196 | FALSE | FALSE |
| LOC107067073 | tyrosine-protein phosphatase non-receptor | 0.04831 | 9.60082 | FALSE | FALSE |
| LOC107074367 | uncharacterized protein LOC107074367 | 0.04779 | 9.60401 | FALSE | FALSE |
| LOC107067762 | uncharacterized protein LOC107067762 | 0.04972 | 9.60612 | FALSE | FALSE |
| LOC107071349 | H(+)/Cl(-) exchange transporter 3 | 0.04774 | 9.61079 | FALSE | FALSE |
| LOC107071070 | uncharacterized protein LOC107071070 | 0.04831 | 9.62993 | FALSE | FALSE |
| LOC107070659 | uncharacterized protein LOC107070659 | 0.04775 | 9.6336 | FALSE | FALSE |
| LOC107074681 | post-GPI attachment to proteins factor 2-lik | 0.04875 | 9.63569 | FALSE | FALSE |
| LOC107068696 | palmitoyl-protein thioesterase 1 | 0.04825 | 9.64248 | FALSE | FALSE |
| LOC107074669 | catenin alpha | 0.04899 | 9.64594 | FALSE | FALSE |
| LOC107066764 | uncharacterized protein LOC107066764 | 0.04925 | 9.64773 | FALSE | FALSE |
| LOC107072679 | protein quiver | 0.04719 | 9.64879 | FALSE | FALSE |
| LOC107069627 | meiotic nuclear division protein 1 homolog | 0.04824 | 9.6789 | FALSE | FALSE |
| LOC107064514 | protein lingerer | 0.04774 | 9.68594 | FALSE | FALSE |
| LOC107069684 | uncharacterized protein LOC107069684 | 0.0488 | 9.71589 | FALSE | FALSE |
| LOC107064307 | cyclin-dependent kinase-like 1 | 0.04684 | 9.71659 | FALSE | FALSE |
| LOC107073064 | myb-like protein Q | 0.04792 | 9.73808 | FALSE | FALSE |
| LOC107069691 | acetylcholine receptor subunit alpha-like 1 | 0.0488 | 9.75226 | FALSE | FALSE |
| LOC107068584 | 32 kDa beta-galactoside-binding lectin lec-3 | 0.04958 | 9.75408 | FALSE | FALSE |
| LOC107071725 | uncharacterized protein DDB_G0283697-lik | 0.0477 | 9.76904 | FALSE | FALSE |
| LOC107067947 | U4/U6.U5 small nuclear ribonucleoprotein 2 | 0.04767 | 9.77387 | FALSE | FALSE |
| LOC107065631 | carbonic anhydrase-related protein 10 | 0.04894 | 9.78073 | FALSE | FALSE |
| LOC107071048 | DNA-binding protein inhibitor ID-1 | 0.04901 | 9.79061 | FALSE | FALSE |
| LOC107068678 | NAD(P)H-hydrate epimerase | 0.0474 | 9.79828 | FALSE | FALSE |
| LOC107066646 | derlin-1 | 0.04929 | 9.79919 | FALSE | FALSE |
| LOC107064646 | acetyl-CoA acetyltransferase, cytosolic | 0.04706 | 9.80727 | FALSE | FALSE |
| LOC107065459 | uncharacterized protein LOC107065459 | 0.04925 | 9.81017 | FALSE | FALSE |
| LOC107069245 | uncharacterized protein LOC107069245 | 0.05014 | 9.81928 | FALSE | FALSE |
| LOC107064506 | 14-3-3 protein epsilon | 0.04825 | 9.83006 | FALSE | FALSE |
| LOC107066208 | WASH complex subunit 7 | 0.04667 | 9.84648 | FALSE | FALSE |
| LOC107071261 | uncharacterized protein LOC107071261 | 0.04745 | 9.84978 | FALSE | FALSE |
| LOC107066416 | apoptosis-inducing factor 3 | 0.04692 | 9.85514 | FALSE | FALSE |
| LOC107067672 | uncharacterized protein LOC107067672 | 0.04711 | 9.85602 | FALSE | FALSE |
| LOC107071006 | phosphatidylinositide phosphatase SAC2 | 0.04791 | 9.85846 | FALSE | FALSE |
| LOC107069265 | cytochrome P450 9e2-like | 0.04729 | 9.86788 | FALSE | FALSE |
| LOC107066144 | uncharacterized protein LOC107066144 | 0.04708 | 9.87078 | FALSE | FALSE |
| LOC107068009 | probable fatty acid-binding protein | 0.05032 | 9.8735 | FALSE | FALSE |
| LOC107067565 | synaptotagmin-6 | 0.04819 | 9.87367 | FALSE | FALSE |
| LOC107074180 | elongation factor G, mitochondrial | 0.04766 | 9.87547 | FALSE | FALSE |
| LOC107068112 | diacylglycerol kinase epsilon-like | 0.04753 | 9.89306 | FALSE | FALSE |
| LOC107064767 | putative aminopeptidase W07G4.4 | 0.04962 | 9.89781 | FALSE | FALSE |
| LOC107073707 | uncharacterized protein LOC107073707 | 0.04847 | 9.90394 | FALSE | FALSE |
| LOC107064760 | protein mab-21 | 0.04839 | 9.94345 | FALSE | FALSE |
| LOC107065684 | receptor-type tyrosine-protein phosphatase | 0.04655 | 9.94778 | FALSE | FALSE |
| LOC107072137 | uncharacterized protein LOC107072137 | 0.04728 | 9.95395 | FALSE | FALSE |
| LOC107068448 | tyrosine-protein kinase Btk29A-like | 0.04898 | 9.9643 | FALSE | FALSE |
| LOC107070318 | odorant receptor 30a-like | 0.0462 | 9.96899 | FALSE | FALSE |
| LOC107073107 | protein numb | 0.04809 | 9.97463 | FALSE | FALSE |
| LOC107068757 | uncharacterized protein LOC107068757 | 0.04807 | 9.98603 | FALSE | FALSE |
| LOC107064552 | major centromere autoantigen B-like | 0.04822 | 9.99951 | FALSE | FALSE |

| | | | | | |
|---|---|---|---|---|---|
| LOC107064300 | uncharacterized protein LOC107064300 | 0.04778 | 10.01741 | FALSE | FALSE |
| LOC107072459 | homeobox protein orthopedia-like | 0.04771 | 10.01747 | FALSE | FALSE |
| LOC107067825 | uncharacterized protein LOC107067825 | 0.04798 | 10.02357 | FALSE | FALSE |
| LOC107065077 | uncharacterized protein LOC107065077 | 0.04814 | 10.02373 | FALSE | FALSE |
| LOC107073517 | uncharacterized protein LOC107073517 | 0.05039 | 10.02443 | FALSE | FALSE |
| LOC107069563 | uncharacterized protein LOC107069563 | 0.04763 | 10.03421 | FALSE | FALSE |
| LOC107071514 | uncharacterized protein LOC107071514 | 0.04919 | 10.04026 | FALSE | FALSE |
| LOC107065730 | nose resistant to fluoxetine protein 6 | 0.04928 | 10.06007 | FALSE | FALSE |
| LOC107064231 | uncharacterized protein LOC107064231 | 0.04906 | 10.07389 | FALSE | FALSE |
| LOC107067208 | facilitated trehalose transporter Tret1-like | 0.04898 | 10.07422 | FALSE | FALSE |
| LOC107067170 | leucine-rich repeat flightless-interacting pro | 0.04761 | 10.07558 | FALSE | FALSE |
| LOC107068340 | glutathione S-transferase C-terminal domaiı | 0.04668 | 10.08932 | FALSE | FALSE |
| LOC107068901 | ADP-ribosylation factor GTPase-activating p | 0.04833 | 10.0924 | FALSE | FALSE |
| LOC107066676 | protein twist | 0.04909 | 10.09867 | FALSE | FALSE |
| LOC107068145 | histone-lysine N-methyltransferase set1 | 0.04984 | 10.11992 | FALSE | FALSE |
| LOC107071677 | MKI67 FHA domain-interacting nucleolar ph | 0.04825 | 10.12983 | FALSE | FALSE |
| LOC107065090 | uncharacterized protein LOC107065090 | 0.04808 | 10.13039 | FALSE | FALSE |
| LOC107065734 | sodium/hydrogen exchanger 3 | 0.04791 | 10.13071 | FALSE | FALSE |
| LOC107070220 | vam6/Vps39-like protein | 0.0484 | 10.14705 | FALSE | FALSE |
| LOC107070062 | uncharacterized protein LOC107070062 | 0.0474 | 10.14726 | FALSE | FALSE |
| LOC107066743 | serine-rich adhesin for platelets | 0.04679 | 10.15587 | FALSE | FALSE |
| LOC107068568 | tryptophan--tRNA ligase, mitochondrial | 0.04651 | 10.15702 | FALSE | FALSE |
| LOC107072262 | uncharacterized protein LOC107072262 | 0.04854 | 10.16927 | FALSE | FALSE |
| LOC107067988 | glycerol-3-phosphate acyltransferase 1, mit | 0.04824 | 10.17601 | FALSE | FALSE |
| LOC107069755 | RNA-binding protein Musashi homolog 2 | 0.04755 | 10.18792 | FALSE | FALSE |
| LOC107069371 | protein vav | 0.04794 | 10.19268 | FALSE | FALSE |
| LOC107074423 | rab GTPase-binding effector protein 1 | 0.04778 | 10.19422 | FALSE | FALSE |
| LOC107071503 | alpha-mannosidase 2 | 0.04686 | 10.20261 | FALSE | FALSE |
| LOC107073983 | nuclear valosin-containing protein-like | 0.0476 | 10.20535 | FALSE | FALSE |
| LOC107070887 | ras-related protein Rap-2a | 0.04938 | 10.2184 | FALSE | FALSE |
| LOC107065301 | alpha-1,3-mannosyl-glycoprotein 2-beta-N-a | 0.04743 | 10.22203 | FALSE | FALSE |
| LOC107072600 | ADP-ribosylation factor-binding protein GG/ | 0.04655 | 10.24847 | FALSE | FALSE |
| LOC107073002 | hydroxypyruvate reductase | 0.04992 | 10.25587 | FALSE | FALSE |
| LOC107068551 | leucine-rich repeat-containing protein let-4- | 0.04687 | 10.32068 | FALSE | FALSE |
| LOC107064172 | post-GPI attachment to proteins factor 2-lik | 0.04883 | 10.32487 | FALSE | FALSE |
| LOC107071738 | uncharacterized protein LOC107071738 | 0.04839 | 10.3367 | FALSE | FALSE |
| LOC107072834 | nose resistant to fluoxetine protein 6-like | 0.04763 | 10.33925 | FALSE | FALSE |
| LOC107069040 | ribonuclease ZC3H12A-like | 0.04846 | 10.35026 | FALSE | FALSE |
| LOC107068288 | PQ-loop repeat-containing protein 1 | 0.04907 | 10.37076 | FALSE | FALSE |
| LOC107068967 | protein rtoA | 0.04903 | 10.37406 | FALSE | FALSE |
| LOC107070919 | cell division cycle-associated protein 7-like | 0.04892 | 10.3895 | FALSE | FALSE |
| LOC107063745 | immunoglobulin domain-containing protein | 0.04692 | 10.3936 | FALSE | FALSE |
| LOC107071530 | prosaposin | 0.04847 | 10.39814 | FALSE | FALSE |
| LOC107065120 | gamma-glutamyltransferase 7-like | 0.04645 | 10.40047 | FALSE | FALSE |
| LOC107072926 | acetyl-coenzyme A transporter 1 | 0.04772 | 10.4102 | FALSE | FALSE |
| LOC107074596 | methionine--tRNA ligase, cytoplasmic | 0.04909 | 10.41154 | FALSE | FALSE |
| LOC107067355 | sialin-like | 0.04839 | 10.42056 | FALSE | FALSE |
| LOC107070529 | gamma-interferon-inducible lysosomal thiol | 0.04682 | 10.43098 | FALSE | FALSE |
| LOC107066508 | uncharacterized protein LOC107066508 | 0.048 | 10.43236 | FALSE | FALSE |
| LOC107064787 | partner of bursicon | 0.04939 | 10.44281 | FALSE | FALSE |

| | | | | | |
|---|---|---|---|---|---|
| LOC107070293 | F-box/LRR-repeat protein 7 | 0.04742 | 10.44316 | FALSE | FALSE |
| LOC107066970 | uncharacterized protein LOC107066970 | 0.04755 | 10.45053 | FALSE | FALSE |
| LOC107072326 | 28S ribosomal protein S9, mitochondrial-like | 0.0472 | 10.45965 | FALSE | FALSE |
| LOC107067316 | uncharacterized protein LOC107067316 | 0.04652 | 10.45976 | FALSE | FALSE |
| LOC107071011 | ras-related protein Rab6 | 0.04823 | 10.46608 | FALSE | FALSE |
| LOC107066391 | F-box only protein 39-like | 0.04711 | 10.4662 | FALSE | FALSE |
| LOC107072892 | uncharacterized protein LOC107072892 | 0.04671 | 10.48857 | FALSE | FALSE |
| LOC107073463 | ras GTPase-activating-like protein IQGAP1 | 0.0481 | 10.48932 | FALSE | FALSE |
| LOC107063740 | transcription initiation factor IIA subunit 1 | 0.04835 | 10.49467 | FALSE | FALSE |
| LOC107072192 | uncharacterized protein LOC107072192 | 0.04778 | 10.50646 | FALSE | FALSE |
| LOC107072591 | protein arginine N-methyltransferase 1 | 0.04534 | 10.50982 | FALSE | FALSE |
| LOC107063656 | ADP-ribose pyrophosphatase, mitochondria | 0.04549 | 10.51402 | FALSE | FALSE |
| LOC107072126 | protein regulator of cytokinesis 1-like | 0.04817 | 10.5215 | FALSE | FALSE |
| LOC107064772 | uncharacterized protein LOC107064772 | 0.04883 | 10.52903 | FALSE | FALSE |
| LOC107063800 | uncharacterized protein LOC107063800 | 0.04804 | 10.5312 | FALSE | FALSE |
| LOC107065778 | fatty acyl-CoA reductase 1-like | 0.04689 | 10.53144 | FALSE | FALSE |
| LOC107067309 | adipocyte plasma membrane-associated pro | 0.04566 | 10.54896 | FALSE | FALSE |
| LOC107072061 | ran-specific GTPase-activating protein-like | 0.04831 | 10.5548 | FALSE | FALSE |
| LOC107070491 | centrosomal protein of 89 kDa-like | 0.04838 | 10.55871 | FALSE | FALSE |
| LOC107068116 | tubulin glycylase 3A-like | 0.04825 | 10.5702 | FALSE | FALSE |
| LOC107063992 | probable serine/threonine-protein kinase pa | 0.04893 | 10.5745 | FALSE | FALSE |
| LOC107069636 | protein PTCD3 homolog, mitochondrial | 0.04827 | 10.59708 | FALSE | FALSE |
| LOC107073761 | uncharacterized protein LOC107073761 | 0.04856 | 10.6016 | FALSE | FALSE |
| LOC107070067 | solute carrier organic anion transporter fam | 0.04887 | 10.60207 | FALSE | FALSE |
| LOC107071357 | PERQ amino acid-rich with GYF domain-con | 0.04899 | 10.61889 | FALSE | FALSE |
| LOC107066096 | mitochondrial dimethyladenosine transfera | 0.04918 | 10.64638 | FALSE | FALSE |
| LOC107066149 | valine--tRNA ligase, mitochondrial-like | 0.04775 | 10.64878 | FALSE | FALSE |
| LOC107071843 | nischarin-like | 0.04818 | 10.6492 | FALSE | FALSE |
| LOC107069575 | uncharacterized protein LOC107069575 | 0.04819 | 10.66667 | FALSE | FALSE |
| LOC107069441 | uncharacterized protein LOC107069441 | 0.04685 | 10.68091 | FALSE | FALSE |
| LOC107067546 | uncharacterized protein LOC107067546 | 0.04862 | 10.70403 | FALSE | FALSE |
| LOC107072153 | cytochrome P450 4C1-like | 0.04605 | 10.71267 | FALSE | FALSE |
| LOC107069212 | ribosomal RNA processing protein 1 homolo | 0.04777 | 10.7152 | FALSE | FALSE |
| LOC107069644 | putative FERM domain-containing protein F | 0.04713 | 10.72422 | FALSE | FALSE |
| LOC107068799 | septin-7 | 0.04633 | 10.72731 | FALSE | FALSE |
| LOC107070744 | G protein-coupled receptor kinase 1 | 0.04724 | 10.73447 | FALSE | FALSE |
| LOC107068207 | putative inorganic phosphate cotransporter | 0.04788 | 10.74542 | FALSE | FALSE |
| LOC107071763 | uncharacterized protein LOC107071763 | 0.04696 | 10.75216 | FALSE | FALSE |
| LOC107064458 | uncharacterized protein LOC107064458 | 0.04717 | 10.762 | FALSE | FALSE |
| LOC107065346 | protein arginine N-methyltransferase 5 | 0.04839 | 10.76884 | FALSE | FALSE |
| LOC107074560 | WD repeat-containing protein 37 | 0.04658 | 10.77411 | FALSE | FALSE |
| LOC107074232 | uncharacterized protein LOC107074232 | 0.04856 | 10.78276 | FALSE | FALSE |
| LOC107069225 | RNA-binding protein 8A | 0.04641 | 10.79988 | FALSE | FALSE |
| LOC107073316 | mediator of RNA polymerase II transcriptior | 0.04805 | 10.80086 | FALSE | FALSE |
| LOC107064491 | MLX-interacting protein | 0.04964 | 10.80916 | FALSE | FALSE |
| LOC107065091 | mediator of RNA polymerase II transcriptior | 0.04858 | 10.82162 | FALSE | FALSE |
| LOC107070385 | antizyme inhibitor 2-like | 0.04754 | 10.82518 | FALSE | FALSE |
| LOC107072476 | uncharacterized protein LOC107072476 | 0.04865 | 10.83307 | FALSE | FALSE |
| LOC107069194 | myophilin | 0.04847 | 10.84449 | FALSE | FALSE |
| LOC107064811 | glycosyltransferase-like protein LARGE2 | 0.04825 | 10.84774 | FALSE | FALSE |

| | | | | | |
|---|---|---|---|---|---|
| LOC107064283 | dehydrogenase/reductase SDR family meml | 0.04668 | 10.84973 | FALSE | FALSE |
| LOC107065551 | serine/threonine-protein phosphatase 2A a | 0.04822 | 10.86062 | FALSE | FALSE |
| LOC107067203 | epithelial discoidin domain-containing recep | 0.0487 | 10.87312 | FALSE | FALSE |
| LOC107070824 | uncharacterized protein LOC107070824 | 0.04773 | 10.88494 | FALSE | FALSE |
| LOC107067808 | uncharacterized protein LOC107067808 | 0.04701 | 10.8881 | FALSE | FALSE |
| LOC107071560 | microtubule-associated protein Jupiter | 0.04653 | 10.8908 | FALSE | FALSE |
| LOC107063926 | hybrid signal transduction histidine kinase G | 0.04676 | 10.89172 | FALSE | FALSE |
| LOC107074047 | LOW QUALITY PROTEIN: adenylate cyclase t | 0.04534 | 10.89244 | FALSE | FALSE |
| LOC107066653 | coiled-coil domain-containing protein 12 | 0.04781 | 10.90253 | FALSE | FALSE |
| LOC107066145 | sodium-dependent nutrient amino acid tran | 0.04698 | 10.90692 | FALSE | FALSE |
| LOC107065060 | uncharacterized protein LOC107065060 | 0.04679 | 10.93679 | FALSE | FALSE |
| LOC107064201 | arylsulfatase B-like | 0.04668 | 10.94242 | FALSE | FALSE |
| LOC107073287 | protein toll-like | 0.04623 | 10.94855 | FALSE | FALSE |
| LOC107064562 | potassium channel subfamily K member 1-li | 0.04793 | 10.95017 | FALSE | FALSE |
| LOC107069837 | WD repeat and HMG-box DNA-binding prot | 0.04672 | 10.9608 | FALSE | FALSE |
| LOC107070364 | OTU domain-containing protein 7B-like | 0.04944 | 10.9656 | FALSE | FALSE |
| LOC107071077 | serine/threonine-protein kinase SIK3-like | 0.0478 | 10.96826 | FALSE | FALSE |
| LOC107068067 | uncharacterized protein LOC107068067 | 0.04723 | 10.97537 | FALSE | FALSE |
| LOC107072881 | retinoid-inducible serine carboxypeptidase- | 0.04662 | 10.98741 | FALSE | FALSE |
| LOC107070602 | synaptosomal-associated protein 29 | 0.04725 | 11.00378 | FALSE | FALSE |
| LOC107068102 | lipoma HMGIC fusion partner-like 2 protein | 0.04741 | 11.006 | FALSE | FALSE |
| LOC107071481 | uncharacterized protein LOC107071481 | 0.0478 | 11.01042 | FALSE | FALSE |
| LOC107064178 | heme oxygenase 1 | 0.04731 | 11.0148 | FALSE | FALSE |
| LOC107069172 | putative gustatory receptor 28b | 0.04816 | 11.04728 | FALSE | FALSE |
| LOC107066765 | major facilitator superfamily domain-contai | 0.04668 | 11.05159 | FALSE | FALSE |
| LOC107073044 | pancreas transcription factor 1 subunit alph | 0.04684 | 11.06316 | FALSE | FALSE |
| LOC107074096 | paired box pox-meso protein | 0.04693 | 11.06959 | FALSE | FALSE |
| LOC107064547 | carbohydrate sulfotransferase 11-like | 0.04689 | 11.07975 | FALSE | FALSE |
| LOC107067421 | death-associated inhibitor of apoptosis 2-lik | 0.04793 | 11.08329 | FALSE | FALSE |
| LOC107068986 | serine-rich adhesin for platelets | 0.04916 | 11.10287 | FALSE | FALSE |
| LOC107066600 | uncharacterized protein LOC107066600 | 0.04875 | 11.11387 | FALSE | FALSE |
| LOC107069605 | uncharacterized protein LOC107069605 | 0.04602 | 11.11709 | FALSE | FALSE |
| LOC107065816 | ral GTPase-activating protein subunit beta | 0.04812 | 11.12054 | FALSE | FALSE |
| LOC107069535 | hairy/enhancer-of-split related with YRPW r | 0.04638 | 11.12988 | FALSE | FALSE |
| LOC107071996 | serine-enriched protein | 0.0473 | 11.13582 | FALSE | FALSE |
| LOC107064893 | mucin-5AC | 0.04687 | 11.13667 | FALSE | FALSE |
| LOC107067194 | transmembrane 9 superfamily member 3 | 0.04717 | 11.15283 | FALSE | FALSE |
| LOC107067329 | tubulin polyglutamylase complex subunit 2 | 0.04831 | 11.1781 | FALSE | FALSE |
| LOC107065347 | protein odr-4 homolog | 0.04861 | 11.20924 | FALSE | FALSE |
| LOC107067763 | uncharacterized protein LOC107067763 | 0.0483 | 11.22189 | FALSE | FALSE |
| LOC107066590 | zinc finger protein 239-like | 0.04787 | 11.23247 | FALSE | FALSE |
| LOC107070248 | Williams-Beuren syndrome chromosomal re | 0.04906 | 11.2345 | FALSE | FALSE |
| LOC107069822 | uncharacterized protein LOC107069822 | 0.04703 | 11.24885 | FALSE | FALSE |
| LOC107068190 | delta(14)-sterol reductase | 0.04578 | 11.26104 | FALSE | FALSE |
| LOC107065206 | long-chain-fatty-acid--CoA ligase 4 | 0.04685 | 11.29286 | FALSE | FALSE |
| LOC107067863 | uncharacterized protein LOC107067863 | 0.04701 | 11.29435 | FALSE | FALSE |
| LOC107069152 | homeobox protein engrailed-1a-like | 0.04704 | 11.30394 | FALSE | FALSE |
| LOC107065681 | uncharacterized protein LOC107065681 | 0.04708 | 11.31192 | FALSE | FALSE |
| LOC107072301 | negative elongation factor D | 0.04744 | 11.33079 | FALSE | FALSE |
| LOC107067678 | pre-mRNA-processing factor 40 homolog A | 0.04737 | 11.33097 | FALSE | FALSE |

| | | | | | |
|---|---|---|---|---|---|
| LOC107066263 | late secretory pathway protein AVL9 homol | 0.04744 | 11.3391 | FALSE | FALSE |
| LOC107066283 | transmembrane protease serine 9-like | 0.04711 | 11.34032 | FALSE | FALSE |
| LOC107064765 | sodium-coupled monocarboxylate transport | 0.04694 | 11.35219 | FALSE | FALSE |
| LOC107068318 | homeobox protein 13-like | 0.04912 | 11.36295 | FALSE | FALSE |
| LOC107069400 | chromodomain-helicase-DNA-binding prote | 0.04781 | 11.37408 | FALSE | FALSE |
| LOC107067090 | 28S ribosomal protein S14, mitochondrial | 0.04753 | 11.3768 | FALSE | FALSE |
| LOC107073792 | uncharacterized protein LOC107073792 | 0.04781 | 11.37706 | FALSE | FALSE |
| LOC107071269 | ankyrin repeat domain-containing protein 2 | 0.04658 | 11.38254 | FALSE | FALSE |
| LOC107064336 | uncharacterized protein LOC107064336 | 0.04726 | 11.3866 | FALSE | FALSE |
| LOC107066609 | ubiquitin-like modifier-activating enzyme at | 0.04814 | 11.39323 | FALSE | FALSE |
| LOC107070988 | tyrosine-protein phosphatase non-receptor | 0.04776 | 11.40585 | FALSE | FALSE |
| LOC107071762 | beta-1,3-glucan-binding protein 1-like | 0.04684 | 11.41864 | FALSE | FALSE |
| LOC107070746 | WD repeat and FYVE domain-containing pro | 0.04651 | 11.41984 | FALSE | FALSE |
| LOC107069006 | protein PFC0760c-like | 0.04696 | 11.42816 | FALSE | FALSE |
| LOC107066959 | elongation of very long chain fatty acids pro | 0.04728 | 11.45285 | FALSE | FALSE |
| LOC107065625 | vesicle transport through interaction with t- | 0.0471 | 11.45506 | FALSE | FALSE |
| LOC107066321 | chromaffin granule amine transporter | 0.04762 | 11.45509 | FALSE | FALSE |
| LOC107069174 | synaptic vesicle glycoprotein 2B-like | 0.04672 | 11.45926 | FALSE | FALSE |
| LOC107072569 | cysteine-rich protein 2-binding protein | 0.04749 | 11.46701 | FALSE | FALSE |
| LOC107064056 | WD repeat-containing protein WRAP73-like | 0.04846 | 11.46819 | FALSE | FALSE |
| LOC107071379 | RNA-binding protein pno1-like | 0.04606 | 11.47474 | FALSE | FALSE |
| LOC107073178 | piggyBac transposable element-derived pro | 0.04549 | 11.48488 | FALSE | FALSE |
| LOC107071548 | integrator complex subunit 6 | 0.04757 | 11.4944 | FALSE | FALSE |
| LOC107074281 | anaphase-promoting complex subunit 7 | 0.04622 | 11.50028 | FALSE | FALSE |
| LOC107073078 | protein FAM114A2 | 0.04787 | 11.50731 | FALSE | FALSE |
| LOC107069777 | hydroxysteroid dehydrogenase-like protein | 0.04724 | 11.50986 | FALSE | FALSE |
| LOC107073369 | uncharacterized protein LOC107073369 | 0.04693 | 11.51112 | FALSE | FALSE |
| LOC107069227 | 39S ribosomal protein L21, mitochondrial | 0.04907 | 11.51639 | FALSE | FALSE |
| LOC107066059 | mitochondrial import inner membrane tran: | 0.04755 | 11.5273 | FALSE | FALSE |
| LOC107065348 | dual specificity protein kinase splA-like | 0.04832 | 11.54458 | FALSE | FALSE |
| LOC107068588 | glycogen phosphorylase | 0.04609 | 11.549 | FALSE | FALSE |
| LOC107069275 | ropporin-1-like protein | 0.04651 | 11.5495 | FALSE | FALSE |
| LOC107064261 | zinc finger with UFM1-specific peptidase do | 0.0472 | 11.56423 | FALSE | FALSE |
| LOC107065354 | RB1-inducible coiled-coil protein 1 | 0.04888 | 11.56545 | FALSE | FALSE |
| LOC107071143 | paramyosin, short form-like | 0.04846 | 11.57687 | FALSE | FALSE |
| LOC107066898 | myb-like protein I | 0.04799 | 11.59543 | FALSE | FALSE |
| LOC107070821 | uncharacterized protein LOC107070821 | 0.04618 | 11.59873 | FALSE | FALSE |
| LOC107070206 | uncharacterized protein LOC107070206 | 0.04484 | 11.61524 | FALSE | FALSE |
| LOC107068844 | uncharacterized protein LOC107068844 | 0.04761 | 11.62256 | FALSE | FALSE |
| LOC107071154 | uncharacterized protein LOC107071154 | 0.04672 | 11.63361 | FALSE | FALSE |
| LOC107066911 | trafficking kinesin-binding protein milt | 0.0469 | 11.63411 | FALSE | FALSE |
| LOC107074214 | methylmalonic aciduria and homocystinuria | 0.04668 | 11.64485 | FALSE | FALSE |
| LOC107069490 | TATA-binding protein-associated factor 172 | 0.04586 | 11.66976 | FALSE | FALSE |
| LOC107068780 | serine/arginine-rich splicing factor 2 | 0.04722 | 11.67433 | FALSE | FALSE |
| LOC107068123 | charged multivesicular body protein 4b | 0.04687 | 11.67615 | FALSE | FALSE |
| LOC107064200 | protein yellow-like | 0.0494 | 11.68128 | FALSE | FALSE |
| LOC107065732 | probable serine/threonine-protein kinase D | 0.04723 | 11.69843 | FALSE | FALSE |
| LOC107072747 | cytochrome b reductase 1 | 0.047 | 11.70537 | FALSE | FALSE |
| LOC107070370 | protein sly1 homolog | 0.04843 | 11.70626 | FALSE | FALSE |
| LOC107065326 | transmembrane protein 216-like | 0.04657 | 11.71169 | FALSE | FALSE |

| | | | | | |
|---|---|---|---|---|---|
| LOC107069386 | uncharacterized protein LOC107069386 | 0.04883 | 11.72091 | FALSE | FALSE |
| LOC107071407 | uncharacterized protein LOC107071407 | 0.04722 | 11.73956 | FALSE | FALSE |
| LOC107069657 | dimethyladenosine transferase 2, mitochon | 0.04723 | 11.74333 | FALSE | FALSE |
| LOC107073655 | endoplasmic reticulum aminopeptidase 1 | 0.0498 | 11.75176 | FALSE | FALSE |
| LOC107064165 | tumor susceptibility gene 101 protein | 0.04759 | 11.75589 | FALSE | FALSE |
| LOC107068314 | cytochrome P450 302a1, mitochondrial-like | 0.04724 | 11.75593 | FALSE | FALSE |
| LOC107066031 | WD repeat-containing protein 60 | 0.04806 | 11.77225 | FALSE | FALSE |
| LOC107073805 | farnesyl pyrophosphate synthase | 0.04681 | 11.78287 | FALSE | FALSE |
| LOC107069454 | vacuolar protein sorting-associated protein | 0.04772 | 11.78545 | FALSE | FALSE |
| LOC107068590 | UBX domain-containing protein 6 | 0.04634 | 11.79108 | FALSE | FALSE |
| LOC107064704 | extracellular sulfatase SULF-1 homolog | 0.04659 | 11.81121 | FALSE | FALSE |
| LOC107069555 | TBC1 domain family member 16 | 0.04723 | 11.83283 | FALSE | FALSE |
| LOC107070327 | uncharacterized protein LOC107070327 | 0.04681 | 11.83509 | FALSE | FALSE |
| LOC107068097 | DET1- and DDB1-associated protein 1 | 0.04751 | 11.83905 | FALSE | FALSE |
| LOC107069343 | glutaminyl-peptide cyclotransferase | 0.04679 | 11.85148 | FALSE | FALSE |
| LOC107073643 | 60S ribosomal protein L7a | 0.0467 | 11.85651 | FALSE | FALSE |
| LOC107064228 | spermatogenesis-defective protein 39 hom( | 0.04704 | 11.8705 | FALSE | FALSE |
| LOC107072340 | supervillin-like | 0.04875 | 11.87602 | FALSE | FALSE |
| LOC107071782 | mannosyl-oligosaccharide glucosidase GCS1 | 0.04797 | 11.91445 | FALSE | FALSE |
| LOC107064714 | 7-methylguanosine phosphate-specific 5'-nu | 0.04655 | 11.94015 | FALSE | FALSE |
| LOC107070517 | cysteine-rich hydrophobic domain-containir | 0.04757 | 11.97115 | FALSE | FALSE |
| LOC107073303 | SUN domain-containing protein 1-like | 0.04882 | 11.9827 | FALSE | FALSE |
| LOC107069239 | uncharacterized protein LOC107069239 | 0.04826 | 11.98484 | FALSE | FALSE |
| LOC107066070 | zinc finger protein 112-like | 0.04874 | 11.98729 | FALSE | FALSE |
| LOC107068817 | hydroxymethylglutaryl-CoA synthase 1 | 0.04597 | 12.02328 | FALSE | FALSE |
| LOC107067334 | heparan sulfate 2-O-sulfotransferase pipe | 0.04655 | 12.03314 | FALSE | FALSE |
| LOC107065071 | cytochrome P450 4C1-like | 0.04894 | 12.03799 | FALSE | FALSE |
| LOC107071433 | ATP synthase subunit s-like protein | 0.04565 | 12.03839 | FALSE | FALSE |
| LOC107072886 | uncharacterized protein LOC107072886 | 0.04707 | 12.04022 | FALSE | FALSE |
| LOC107071896 | protein lin-52 homolog | 0.04811 | 12.04423 | FALSE | FALSE |
| LOC107066574 | polymerase delta-interacting protein 2 | 0.04585 | 12.05902 | FALSE | FALSE |
| LOC107069182 | vacuolar protein sorting-associated protein | 0.04584 | 12.06572 | FALSE | FALSE |
| LOC107071612 | 3-oxoacyl-[acyl-carrier-protein | 0.04719 | 12.0691 | FALSE | FALSE |
| LOC107072872 | F-BAR domain only protein 2 | 0.04737 | 12.06966 | FALSE | FALSE |
| LOC107073842 | axin-1 | 0.04707 | 12.07195 | FALSE | FALSE |
| LOC107074544 | integrator complex subunit 12-like | 0.04689 | 12.08333 | FALSE | FALSE |
| LOC107071747 | zinc finger protein 385B-like | 0.04972 | 12.08376 | FALSE | FALSE |
| LOC107064019 | MATH and LRR domain-containing protein F | 0.04612 | 12.08756 | FALSE | FALSE |
| LOC107070756 | uncharacterized protein LOC107070756 | 0.04881 | 12.08803 | FALSE | FALSE |
| LOC107065290 | zinc finger protein 615-like | 0.04574 | 12.10577 | FALSE | FALSE |
| LOC107068167 | uncharacterized protein LOC107068167 | 0.04669 | 12.11416 | FALSE | FALSE |
| LOC107067360 | putative uncharacterized protein DDB_G027 | 0.04745 | 12.12006 | FALSE | FALSE |
| LOC107064414 | hexamerin-like | 0.04738 | 12.12267 | FALSE | FALSE |
| LOC107068049 | protein Wnt-5b-like | 0.04701 | 12.13687 | FALSE | FALSE |
| LOC107067473 | uncharacterized protein LOC107067473 | 0.04788 | 12.13891 | FALSE | FALSE |
| LOC107072362 | forkhead box protein O-like | 0.04742 | 12.13924 | FALSE | FALSE |
| LOC107067614 | YEATS domain-containing protein 4 | 0.04577 | 12.15665 | FALSE | FALSE |
| LOC107064688 | transmembrane protein 147 | 0.04843 | 12.1575 | FALSE | FALSE |
| LOC107073899 | histone-lysine N-methyltransferase SETMAF | 0.04646 | 12.17409 | FALSE | FALSE |
| LOC107074149 | uncharacterized protein LOC107074149 | 0.04705 | 12.18077 | FALSE | FALSE |

| | | | | | |
|---|---|---|---|---|---|
| LOC107068919 | alpha-1,6-mannosyl-glycoprotein 2-beta-N-a | 0.04912 | 12.1816 | FALSE | FALSE |
| LOC107065215 | choline/ethanolamine kinase | 0.04629 | 12.19958 | FALSE | FALSE |
| LOC107070698 | gamma-soluble NSF attachment protein-like | 0.04711 | 12.20261 | FALSE | FALSE |
| LOC107065972 | transmembrane protein 19 | 0.0461 | 12.21501 | FALSE | FALSE |
| LOC107074309 | locomotion-related protein Hikaru genki | 0.04611 | 12.21626 | FALSE | FALSE |
| LOC107065800 | zinc finger protein 260-like | 0.04602 | 12.22002 | FALSE | FALSE |
| LOC107068415 | PCTP-like protein | 0.04534 | 12.2356 | FALSE | FALSE |
| LOC107074118 | uncharacterized protein LOC107074118 | 0.04565 | 12.23978 | FALSE | FALSE |
| LOC107074125 | 40S ribosomal protein S3 | 0.0474 | 12.24575 | FALSE | FALSE |
| LOC107074032 | dynactin subunit 1 | 0.04765 | 12.24713 | FALSE | FALSE |
| LOC107068271 | centromere-associated protein E-like | 0.04763 | 12.24949 | FALSE | FALSE |
| LOC107067458 | protein lethal(2)essential for life-like | 0.05018 | 12.24958 | FALSE | FALSE |
| LOC107065092 | uncharacterized protein LOC107065092 | 0.04655 | 12.25369 | FALSE | FALSE |
| LOC107068516 | protein spaetzle | 0.04633 | 12.25492 | FALSE | FALSE |
| LOC107068188 | uncharacterized protein LOC107068188 | 0.04733 | 12.26049 | FALSE | FALSE |
| LOC107069805 | 39S ribosomal protein L47, mitochondrial | 0.04787 | 12.27306 | FALSE | FALSE |
| LOC107066777 | cytochrome P450 9e2-like | 0.04775 | 12.27365 | FALSE | FALSE |
| LOC107064275 | uncharacterized protein LOC107064275 | 0.04772 | 12.28763 | FALSE | FALSE |
| LOC107072174 | CMP-sialic acid transporter 1 | 0.04708 | 12.28794 | FALSE | FALSE |
| LOC107071089 | F-actin-capping protein subunit beta | 0.04692 | 12.32258 | FALSE | FALSE |
| LOC107074103 | nitrogen permease regulator 3-like protein | 0.04758 | 12.32337 | FALSE | FALSE |
| LOC107072583 | E3 ubiquitin-protein ligase RNF180-like | 0.04912 | 12.33667 | FALSE | FALSE |
| LOC107065333 | NADH dehydrogenase [ubiquinone | 0.04728 | 12.35462 | FALSE | FALSE |
| LOC107066473 | pro-resilin | 0.04729 | 12.35494 | FALSE | FALSE |
| LOC107067649 | probable ATP-dependent RNA helicase DDX | 0.04705 | 12.35914 | FALSE | FALSE |
| LOC107070620 | endoglucanase E-4-like | 0.04691 | 12.36076 | FALSE | FALSE |
| LOC107070699 | uncharacterized protein LOC107070699 | 0.04641 | 12.37761 | FALSE | FALSE |
| LOC107067259 | E3 ubiquitin-protein ligase TRIP12 | 0.04739 | 12.40069 | FALSE | FALSE |
| LOC107065295 | probable RNA-binding protein EIF1AD | 0.04708 | 12.42357 | FALSE | FALSE |
| LOC107066110 | protein enhancer of sevenless 2B | 0.04575 | 12.42637 | FALSE | FALSE |
| LOC107071456 | cytochrome P450 315a1, mitochondrial | 0.04569 | 12.44069 | FALSE | FALSE |
| LOC107070342 | probable G-protein coupled receptor CG317 | 0.04801 | 12.44756 | FALSE | FALSE |
| LOC107068471 | chitinase-3-like protein 2 | 0.04505 | 12.45339 | FALSE | FALSE |
| LOC107069457 | signal transducing adapter molecule 1 | 0.04801 | 12.47466 | FALSE | FALSE |
| LOC107067351 | uncharacterized protein LOC107067351 | 0.04694 | 12.47548 | FALSE | FALSE |
| LOC107072706 | neurofilament heavy polypeptide-like | 0.04707 | 12.47922 | FALSE | FALSE |
| LOC107065039 | uncharacterized protein LOC107065039 | 0.0458 | 12.49046 | FALSE | FALSE |
| LOC107072125 | SET and MYND domain-containing protein 4 | 0.04529 | 12.49369 | FALSE | FALSE |
| LOC107068039 | aquaporin AQPAe.a-like | 0.04737 | 12.49566 | FALSE | FALSE |
| LOC107068386 | globin CTT-E/E' | 0.04665 | 12.49917 | FALSE | FALSE |
| LOC107073986 | uncharacterized protein LOC107073986 | 0.04662 | 12.5007 | FALSE | FALSE |
| LOC107068047 | IQ and AAA domain-containing protein 1-lik | 0.04725 | 12.50105 | FALSE | FALSE |
| LOC107069206 | signal peptidase complex catalytic subunit S | 0.04774 | 12.50562 | FALSE | FALSE |
| LOC107069355 | uncharacterized protein LOC107069355 | 0.0474 | 12.51194 | FALSE | FALSE |
| LOC107072687 | uncharacterized protein LOC107072687 | 0.04739 | 12.51494 | FALSE | FALSE |
| LOC107068413 | succinyl-CoA:3-ketoacid coenzyme A transfe | 0.04639 | 12.55304 | FALSE | FALSE |
| LOC107073861 | polyhomeotic-proximal chromatin protein-l | 0.04721 | 12.55619 | FALSE | FALSE |
| LOC107071769 | polyprenol reductase | 0.04642 | 12.56516 | FALSE | FALSE |
| LOC107073495 | lipase 3-like | 0.04594 | 12.57894 | FALSE | FALSE |
| LOC107066131 | DNA-directed RNA polymerases I, II, and III s | 0.04847 | 12.58244 | FALSE | FALSE |

| | | | | | |
|---|---|---|---|---|---|
| LOC107067809 | serine/threonine-protein kinase MPS1 | 0.04518 | 12.58422 | FALSE | FALSE |
| LOC107067263 | mRNA-capping enzyme | 0.04659 | 12.58667 | FALSE | FALSE |
| LOC107068515 | dnaJ homolog subfamily C member 7 | 0.04659 | 12.60621 | FALSE | FALSE |
| LOC107071633 | vigilin | 0.04558 | 12.61062 | FALSE | FALSE |
| LOC107073329 | uncharacterized protein LOC107073329 | 0.04626 | 12.62239 | FALSE | FALSE |
| LOC107074183 | trimethylguanosine synthase | 0.04671 | 12.62636 | FALSE | FALSE |
| LOC107070684 | aminopeptidase N-like | 0.04588 | 12.63653 | FALSE | FALSE |
| LOC107071661 | protein germ cell-less | 0.04788 | 12.64439 | FALSE | FALSE |
| LOC107067747 | mitochondrial-processing peptidase subunit | 0.04528 | 12.64628 | FALSE | FALSE |
| LOC107065086 | serine/threonine-protein kinase RIO3-like | 0.04706 | 12.64762 | FALSE | FALSE |
| LOC107065965 | rhodopsin, long-wavelength-like | 0.04561 | 12.64816 | FALSE | FALSE |
| LOC107073217 | beta-catenin-like protein 1 | 0.04629 | 12.66157 | FALSE | FALSE |
| LOC107064753 | guanylate cyclase soluble subunit beta-1 | 0.04576 | 12.66157 | FALSE | FALSE |
| LOC107072350 | uncharacterized protein LOC107072350 | 0.04613 | 12.67209 | FALSE | FALSE |
| LOC107072800 | baculoviral IAP repeat-containing protein 3 | 0.04733 | 12.67296 | FALSE | FALSE |
| LOC107073296 | replication factor C subunit 5 | 0.0469 | 12.68325 | FALSE | FALSE |
| LOC107068885 | homeobox protein OTX1-like | 0.04435 | 12.69557 | FALSE | FALSE |
| LOC107070034 | uncharacterized protein LOC107070034 | 0.04747 | 12.70615 | FALSE | FALSE |
| LOC107067784 | major facilitator superfamily domain-contai | 0.04599 | 12.73808 | FALSE | FALSE |
| LOC107067774 | DNA oxidative demethylase ALKBH1 | 0.04717 | 12.74113 | FALSE | FALSE |
| LOC107067452 | vanin-like protein 2 | 0.04686 | 12.74827 | FALSE | FALSE |
| LOC107071487 | putative fatty acyl-CoA reductase CG5065 | 0.04632 | 12.75208 | FALSE | FALSE |
| LOC107074289 | uncharacterized protein LOC107074289 | 0.04679 | 12.75795 | FALSE | FALSE |
| LOC107065544 | retinol dehydrogenase 14 | 0.04691 | 12.76129 | FALSE | FALSE |
| LOC107070180 | LIM and senescent cell antigen-like-containi | 0.04633 | 12.76464 | FALSE | FALSE |
| LOC107071046 | puromycin-sensitive aminopeptidase | 0.04587 | 12.77872 | FALSE | FALSE |
| LOC107065072 | thymidylate synthase | 0.04767 | 12.78171 | FALSE | FALSE |
| LOC107068200 | ras-related protein RabX-like | 0.04593 | 12.78967 | FALSE | FALSE |
| LOC107071479 | checkpoint protein HUS1 | 0.04971 | 12.7899 | FALSE | FALSE |
| LOC107066207 | MICAL-like protein 1 | 0.04537 | 12.79989 | FALSE | FALSE |
| LOC107072315 | protein wings apart-like | 0.04968 | 12.8023 | FALSE | FALSE |
| LOC107070688 | uncharacterized protein LOC107070688 | 0.04832 | 12.82547 | FALSE | FALSE |
| LOC107070874 | urea transporter 1-like | 0.04874 | 12.83741 | FALSE | FALSE |
| LOC107066318 | protein EFR3 homolog cmp44E | 0.04682 | 12.85591 | FALSE | FALSE |
| LOC107069823 | uncharacterized protein LOC107069823 | 0.04677 | 12.85791 | FALSE | FALSE |
| LOC107068150 | translation machinery-associated protein 7 | 0.0458 | 12.85972 | FALSE | FALSE |
| LOC107071058 | breast cancer anti-estrogen resistance prote | 0.04576 | 12.86475 | FALSE | FALSE |
| LOC107072378 | thyrotroph embryonic factor-like | 0.04595 | 12.86877 | FALSE | FALSE |
| LOC107066381 | nuclear pore complex protein Nup85 | 0.04557 | 12.86902 | FALSE | FALSE |
| LOC107068134 | cyclin-dependent kinase 8-like | 0.04532 | 12.87218 | FALSE | FALSE |
| LOC107072108 | 26S protease regulatory subunit 6A-B | 0.04752 | 12.87273 | FALSE | FALSE |
| LOC107066194 | nitrilase and fragile histidine triad fusion pro | 0.048 | 12.87328 | FALSE | FALSE |
| LOC107071553 | cilia- and flagella-associated protein 20 | 0.0463 | 12.87549 | FALSE | FALSE |
| LOC107070291 | transmembrane protein C5orf28 | 0.04689 | 12.87756 | FALSE | FALSE |
| LOC107067098 | uncharacterized protein LOC107067098 | 0.04603 | 12.88314 | FALSE | FALSE |
| LOC107073762 | uncharacterized protein LOC107073762 | 0.04621 | 12.88435 | FALSE | FALSE |
| LOC107065834 | kinesin-like protein KIF13A | 0.04609 | 12.89904 | FALSE | FALSE |
| LOC107065343 | probable DNA replication complex GINS pro | 0.04616 | 12.90058 | FALSE | FALSE |
| LOC107068871 | odorant receptor 22c-like | 0.04505 | 12.90183 | FALSE | FALSE |
| LOC107066176 | E3 ubiquitin-protein ligase Nedd-4 | 0.04691 | 12.91178 | FALSE | FALSE |

| | | | | | |
|---|---|---|---|---|---|
| LOC107070892 | protein scarlet-like | 0.04611 | 12.9239 | FALSE | FALSE |
| LOC107070798 | 26S protease regulatory subunit 6B | 0.04598 | 12.9286 | FALSE | FALSE |
| LOC107068055 | uncharacterized protein LOC107068055 | 0.04701 | 12.93023 | FALSE | FALSE |
| LOC107063979 | discoidin domain-containing receptor 2-like | 0.04626 | 12.94926 | FALSE | FALSE |
| LOC107068591 | basigin | 0.04669 | 12.95617 | FALSE | FALSE |
| LOC107067164 | putative fatty acyl-CoA reductase CG5065 | 0.04627 | 12.95894 | FALSE | FALSE |
| LOC107071051 | ADP-ribosylation factor-like protein 8B-A | 0.048 | 12.96713 | FALSE | FALSE |
| LOC107071793 | lactosylceramide 4-alpha-galactosyltransfer | 0.0463 | 12.9732 | FALSE | FALSE |
| LOC107069725 | 14 kDa phosphohistidine phosphatase-like | 0.04595 | 12.98359 | FALSE | FALSE |
| LOC107070562 | lachesin-like | 0.04666 | 12.98452 | FALSE | FALSE |
| LOC107065755 | uncharacterized protein LOC107065755 | 0.04815 | 12.98584 | FALSE | FALSE |
| LOC107069794 | TATA-box-binding protein-like | 0.0458 | 12.98592 | FALSE | FALSE |
| LOC107073250 | mucin-5AC | 0.04524 | 12.98976 | FALSE | FALSE |
| LOC107068019 | uncharacterized protein LOC107068019 | 0.04721 | 12.99283 | FALSE | FALSE |
| LOC107066118 | zinc finger protein GLI2-like | 0.04601 | 13.00077 | FALSE | FALSE |
| LOC107065018 | uncharacterized protein LOC107065018 | 0.04603 | 13.03366 | FALSE | FALSE |
| LOC107070730 | cuticle protein 19-like | 0.04987 | 13.039 | FALSE | FALSE |
| LOC107064952 | ataxin-7-like protein 1 | 0.04716 | 13.04711 | FALSE | FALSE |
| LOC107072658 | galactosylgalactosylxylosylprotein 3-beta-gl | 0.04563 | 13.05288 | FALSE | FALSE |
| LOC107066360 | uncharacterized protein LOC107066360 | 0.04661 | 13.07762 | FALSE | FALSE |
| LOC107064652 | regulating synaptic membrane exocytosis pr | 0.0468 | 13.09115 | FALSE | FALSE |
| LOC107070339 | dual 3',5'-cyclic-AMP and -GMP phosphodie | 0.04713 | 13.09137 | FALSE | FALSE |
| LOC107067290 | tubulin polyglutamylase TTLL6 | 0.04613 | 13.09557 | FALSE | FALSE |
| LOC107069460 | uncharacterized protein LOC107069460 | 0.04695 | 13.1234 | FALSE | FALSE |
| LOC107068697 | myosin heavy chain, skeletal muscle-like | 0.04769 | 13.12527 | FALSE | FALSE |
| LOC107064718 | juvenile hormone acid O-methyltransferase | 0.0459 | 13.12698 | FALSE | FALSE |
| LOC107068941 | N-terminal kinase-like protein | 0.04621 | 13.13336 | FALSE | FALSE |
| LOC107064576 | uncharacterized protein LOC107064576 | 0.04738 | 13.14115 | FALSE | FALSE |
| LOC107070627 | pre-mRNA-splicing factor 18 | 0.04657 | 13.14543 | FALSE | FALSE |
| LOC107074666 | ceramide-1-phosphate transfer protein | 0.0455 | 13.16642 | FALSE | FALSE |
| LOC107068077 | nitrogen permease regulator 2-like protein | 0.04668 | 13.1797 | FALSE | FALSE |
| LOC107064337 | GMP synthase [glutamine-hydrolyzing | 0.04624 | 13.19059 | FALSE | FALSE |
| LOC107070041 | E3 ubiquitin-protein ligase SIAH1-like | 0.04578 | 13.19301 | FALSE | FALSE |
| LOC107064553 | 60S ribosomal protein L44 | 0.04654 | 13.19593 | FALSE | FALSE |
| LOC107069983 | ADAM 17-like protease | 0.04584 | 13.1973 | FALSE | FALSE |
| LOC107067430 | small heat shock protein hspG3-like | 0.04782 | 13.20044 | FALSE | FALSE |
| LOC107064194 | probable dynactin subunit 2 | 0.04642 | 13.20077 | FALSE | FALSE |
| LOC107066921 | putative uncharacterized protein DDB_G028 | 0.04768 | 13.20454 | FALSE | FALSE |
| LOC107066382 | ATP synthase lipid-binding protein, mitocho | 0.04455 | 13.22004 | FALSE | FALSE |
| LOC107068895 | uncharacterized protein LOC107068895 | 0.04649 | 13.22351 | FALSE | FALSE |
| LOC107065993 | histone acetyltransferase type B catalytic su | 0.04592 | 13.22782 | FALSE | FALSE |
| LOC107071617 | nitric oxide synthase, salivary gland | 0.0466 | 13.23643 | FALSE | FALSE |
| LOC107072618 | uncharacterized protein LOC107072618 | 0.04516 | 13.25903 | FALSE | FALSE |
| LOC107067264 | uncharacterized protein LOC107067264 | 0.04585 | 13.25908 | FALSE | FALSE |
| LOC107070469 | protocadherin-like wing polarity protein sta | 0.0468 | 13.2618 | FALSE | FALSE |
| LOC107066087 | sodium/potassium-transporting ATPase sub | 0.04684 | 13.26397 | FALSE | FALSE |
| LOC107065101 | prohormone-2-like | 0.04553 | 13.27222 | FALSE | FALSE |
| LOC107071355 | uncharacterized protein LOC107071355 | 0.04635 | 13.27409 | FALSE | FALSE |
| LOC107072796 | 60S ribosomal protein L18a | 0.04567 | 13.28418 | FALSE | FALSE |
| LOC107069155 | putative uncharacterized protein DDB_G028 | 0.04621 | 13.28687 | FALSE | FALSE |

| | | | | | |
|---|---|---|---|---|---|
| LOC107063869 | beta-ureidopropionase | 0.04737 | 13.30035 | FALSE | FALSE |
| LOC107070463 | coatomer subunit gamma | 0.046 | 13.30362 | FALSE | FALSE |
| LOC107071922 | annexin B9-like | 0.04551 | 13.30548 | FALSE | FALSE |
| LOC107070890 | transmembrane protein 205 | 0.04649 | 13.3143 | FALSE | FALSE |
| LOC107064992 | mediator of RNA polymerase II transcriptior | 0.04471 | 13.32517 | FALSE | FALSE |
| LOC107065923 | zinc finger protein 184 | 0.04494 | 13.33092 | FALSE | FALSE |
| LOC107069927 | ras-related protein Rab-40C | 0.04583 | 13.35038 | FALSE | FALSE |
| LOC107065330 | uncharacterized protein LOC107065330 | 0.04562 | 13.35469 | FALSE | FALSE |
| LOC107071629 | uncharacterized protein LOC107071629 | 0.04545 | 13.36847 | FALSE | FALSE |
| LOC107074261 | uncharacterized protein LOC107074261 | 0.04558 | 13.379 | FALSE | FALSE |
| LOC107065586 | uncharacterized protein LOC107065586 | 0.04442 | 13.38039 | FALSE | FALSE |
| LOC107071754 | tubulin-specific chaperone D | 0.04533 | 13.38688 | FALSE | FALSE |
| LOC107067730 | alpha/beta hydrolase domain-containing pr | 0.04586 | 13.38707 | FALSE | FALSE |
| LOC107068198 | ATPase family AAA domain-containing prote | 0.0472 | 13.39381 | FALSE | FALSE |
| LOC107069745 | odorant receptor 85f-like | 0.04642 | 13.39428 | FALSE | FALSE |
| LOC107068809 | serine--pyruvate aminotransferase, mitocho | 0.04654 | 13.40933 | FALSE | FALSE |
| LOC107071906 | nucleoporin seh1-A | 0.0454 | 13.41086 | FALSE | FALSE |
| LOC107071840 | alpha-crystallin B chain | 0.04513 | 13.41901 | FALSE | FALSE |
| LOC107065620 | uncharacterized protein LOC107065620 | 0.0465 | 13.42097 | FALSE | FALSE |
| LOC107064310 | FACT complex subunit SSRP1-like | 0.04685 | 13.43181 | FALSE | FALSE |
| LOC107071415 | dolichol kinase | 0.04442 | 13.43642 | FALSE | FALSE |
| LOC107073098 | putative inhibitor of apoptosis | 0.04727 | 13.44574 | FALSE | FALSE |
| LOC107063864 | uncharacterized protein LOC107063864 | 0.04621 | 13.44597 | FALSE | FALSE |
| LOC107063614 | uncharacterized protein LOC107063614 | 0.04635 | 13.45516 | FALSE | FALSE |
| LOC107073039 | chorion peroxidase | 0.04654 | 13.46369 | FALSE | FALSE |
| LOC107070505 | 40S ribosomal protein S19-like | 0.04557 | 13.46958 | FALSE | FALSE |
| LOC107069168 | multiple epidermal growth factor-like doma | 0.04747 | 13.47132 | FALSE | FALSE |
| LOC107063956 | arf-GAP with dual PH domain-containing pro | 0.04633 | 13.47219 | FALSE | FALSE |
| LOC107065527 | homeobox protein abdominal-B-like | 0.04716 | 13.47681 | FALSE | FALSE |
| LOC107071550 | TBC1 domain family member 9 | 0.04459 | 13.48933 | FALSE | FALSE |
| Trnaa-ggc | Trnaa-ggc | 0.04532 | 13.49908 | FALSE | FALSE |
| LOC107072400 | uncharacterized protein LOC107072400 | 0.04701 | 13.51274 | FALSE | FALSE |
| LOC107071923 | poly(A) RNA polymerase gld-2 homolog A-lil | 0.04783 | 13.52828 | FALSE | FALSE |
| LOC107071828 | tumor necrosis factor alpha-induced proteir | 0.04499 | 13.5415 | FALSE | FALSE |
| LOC107068036 | signal recognition particle subunit SRP72 | 0.0469 | 13.5457 | FALSE | FALSE |
| LOC107070262 | DENN domain-containing protein 1A | 0.04591 | 13.55431 | FALSE | FALSE |
| LOC107068958 | dynein heavy chain 7, axonemal-like | 0.04688 | 13.56114 | FALSE | FALSE |
| LOC107067559 | la protein homolog | 0.04586 | 13.56259 | FALSE | FALSE |
| LOC107067918 | uncharacterized protein LOC107067918 | 0.04435 | 13.56532 | FALSE | FALSE |
| LOC107068291 | natterin-3-like | 0.04497 | 13.58142 | FALSE | FALSE |
| LOC107065611 | DEAD-box helicase Dbp80 | 0.04448 | 13.58214 | FALSE | FALSE |
| LOC107072091 | factor VIII intron 22 protein-like | 0.0453 | 13.58606 | FALSE | FALSE |
| LOC107068668 | chromatin assembly factor 1 subunit B | 0.04643 | 13.59098 | FALSE | FALSE |
| LOC107066557 | gamma-interferon-inducible-lysosomal thio | 0.04638 | 13.60322 | FALSE | FALSE |
| LOC107064997 | protein smoothened | 0.04581 | 13.60436 | FALSE | FALSE |
| LOC107066576 | zinc finger protein 330 homolog | 0.04603 | 13.60994 | FALSE | FALSE |
| LOC107069288 | protein trachealess | 0.04417 | 13.61011 | FALSE | FALSE |
| LOC107072929 | gamma-tubulin complex component 2-like | 0.04535 | 13.61201 | FALSE | FALSE |
| LOC107067829 | ADP-ribosylation factor-like protein 3 | 0.04542 | 13.61328 | FALSE | FALSE |
| LOC107071739 | xaa-Pro dipeptidase | 0.04612 | 13.62146 | FALSE | FALSE |

| | | | | | |
|---|---|---|---|---|---|
| LOC107074450 | potassium voltage-gated channel protein Sh | 0.04659 | 13.64117 | FALSE | FALSE |
| LOC107068634 | mRNA-decapping enzyme 1A | 0.045 | 13.65396 | FALSE | FALSE |
| LOC107063905 | nuclear pore complex protein Nup205 | 0.04637 | 13.65878 | FALSE | FALSE |
| LOC107067064 | uncharacterized protein LOC107067064 | 0.04652 | 13.67481 | FALSE | FALSE |
| LOC107068394 | octopamine receptor beta-1R-like | 0.04448 | 13.68582 | FALSE | FALSE |
| LOC107073204 | proteasome subunit beta type-6 | 0.04521 | 13.69295 | FALSE | FALSE |
| LOC107070445 | suppressor of fused homolog | 0.04427 | 13.70249 | FALSE | FALSE |
| LOC107067611 | putative glycine-rich cell wall structural prot | 0.04561 | 13.71784 | FALSE | FALSE |
| LOC107073089 | GTP-binding protein Di-Ras2 | 0.04698 | 13.71943 | FALSE | FALSE |
| LOC107072848 | dexamethasone-induced Ras-related protei | 0.04599 | 13.7196 | FALSE | FALSE |
| LOC107074487 | serine-rich adhesin for platelets-like | 0.04599 | 13.76905 | FALSE | FALSE |
| LOC107069592 | uncharacterized protein LOC107069592 | 0.04534 | 13.77608 | FALSE | FALSE |
| LOC107070847 | uncharacterized protein LOC107070847 | 0.04564 | 13.77975 | FALSE | FALSE |
| LOC107073269 | conserved oligomeric Golgi complex subuni | 0.04664 | 13.78683 | FALSE | FALSE |
| LOC107067417 | protein bric-a-brac 2-like | 0.04413 | 13.79269 | FALSE | FALSE |
| LOC107070672 | protein kintoun | 0.04533 | 13.79794 | FALSE | FALSE |
| LOC107072964 | F-box only protein 28 | 0.04726 | 13.80029 | FALSE | FALSE |
| LOC107072319 | trafficking protein particle complex subunit | 0.0464 | 13.80541 | FALSE | FALSE |
| LOC107074109 | ecdysone-induced protein 74EF-like | 0.04583 | 13.81333 | FALSE | FALSE |
| LOC107071708 | uncharacterized protein LOC107071708 | 0.04648 | 13.82016 | FALSE | FALSE |
| LOC107074537 | uncharacterized protein LOC107074537 | 0.04346 | 13.8247 | FALSE | FALSE |
| LOC107068658 | fatty acyl-CoA reductase 1-like | 0.04611 | 13.82646 | FALSE | FALSE |
| LOC107067163 | 60S ribosomal protein L38 | 0.04635 | 13.83308 | FALSE | FALSE |
| LOC107069398 | uncharacterized protein LOC107069398 | 0.04386 | 13.84087 | FALSE | FALSE |
| LOC107069753 | ZZ-type zinc finger-containing protein 3 | 0.04578 | 13.84501 | FALSE | FALSE |
| LOC107071068 | myotrophin-like | 0.0468 | 13.85125 | FALSE | FALSE |
| LOC107066602 | uncharacterized protein LOC107066602 | 0.04638 | 13.85432 | FALSE | FALSE |
| LOC107073729 | cytochrome P450 4V2-like | 0.0452 | 13.86714 | FALSE | FALSE |
| LOC107070612 | ADP-ribosylation factor-like protein 6-intera | 0.04545 | 13.86919 | FALSE | FALSE |
| LOC107068061 | leucine-rich repeat-containing protein 15 | 0.0451 | 13.87007 | FALSE | FALSE |
| LOC107065159 | FAS-associated factor 2 | 0.04391 | 13.89301 | FALSE | FALSE |
| LOC107073264 | tetratricopeptide repeat protein 39C-like | 0.04694 | 13.89635 | FALSE | FALSE |
| LOC107066651 | dopamine N-acetyltransferase-like | 0.04573 | 13.89802 | FALSE | FALSE |
| LOC107067177 | UPF0568 protein C14orf166 homolog | 0.04538 | 13.90319 | FALSE | FALSE |
| LOC107067704 | signal peptide peptidase-like 3 | 0.04645 | 13.90555 | FALSE | FALSE |
| LOC107072024 | frequenin-1 | 0.04657 | 13.91674 | FALSE | FALSE |
| LOC107074403 | uncharacterized protein LOC107074403 | 0.04449 | 13.9447 | FALSE | FALSE |
| LOC107065660 | glutamate receptor ionotropic, kainate 5-lik | 0.04566 | 13.94721 | FALSE | FALSE |
| LOC107064139 | uncharacterized protein LOC107064139 | 0.04684 | 13.95507 | FALSE | FALSE |
| LOC107074012 | kynurenine/alpha-aminoadipate aminotrans | 0.04565 | 13.97195 | FALSE | FALSE |
| LOC107070963 | presenilin-1 | 0.04748 | 13.97384 | FALSE | FALSE |
| LOC107071268 | syntaxin-binding protein 5 | 0.04309 | 13.97451 | FALSE | FALSE |
| LOC107067281 | translation initiation factor IF-2, mitochondı | 0.04652 | 14.01567 | FALSE | FALSE |
| LOC107073029 | doublesex- and mab-3-related transcription | 0.04544 | 14.01663 | FALSE | FALSE |
| LOC107064435 | anamorsin homolog | 0.0453 | 14.03241 | FALSE | FALSE |
| LOC107063932 | coatomer subunit beta | 0.04487 | 14.03364 | FALSE | FALSE |
| LOC107073927 | ubiquinol-cytochrome-c reductase complex | 0.04501 | 14.04359 | FALSE | FALSE |
| LOC107066441 | pre-mRNA-splicing factor CWC22 homolog | 0.04566 | 14.0445 | FALSE | FALSE |
| LOC107063715 | glycogen [starch | 0.04596 | 14.04799 | FALSE | FALSE |
| LOC107071299 | uncharacterized protein LOC107071299 | 0.04454 | 14.05365 | FALSE | FALSE |

| | | | | | |
|---|---|---|---|---|---|
| LOC107069866 | cytoplasmic phosphatidylinositol transfer pr | 0.04398 | 14.08423 | FALSE | FALSE |
| LOC107071990 | uncharacterized protein LOC107071990 | 0.04525 | 14.08765 | FALSE | FALSE |
| LOC107068147 | anaphase-promoting complex subunit 11 | 0.04576 | 14.08873 | FALSE | FALSE |
| LOC107066854 | protein phosphatase 1 regulatory subunit 1 | 0.04508 | 14.10041 | FALSE | FALSE |
| LOC107070740 | uncharacterized protein LOC107070740 | 0.0448 | 14.10372 | FALSE | FALSE |
| LOC107071535 | rRNA methyltransferase 3, mitochondrial | 0.04663 | 14.10735 | FALSE | FALSE |
| LOC107070770 | parkin coregulated gene protein homolog | 0.04723 | 14.11815 | FALSE | FALSE |
| LOC107071323 | putative uncharacterized transmembrane p | 0.04546 | 14.12806 | FALSE | FALSE |
| LOC107065235 | probable small nuclear ribonucleoprotein E | 0.04591 | 14.13914 | FALSE | FALSE |
| LOC107069992 | maternal embryonic leucine zipper kinase-li | 0.04543 | 14.13994 | FALSE | FALSE |
| LOC107072515 | trans-1,2-dihydrobenzene-1,2-diol dehydro | 0.04621 | 14.14234 | FALSE | FALSE |
| LOC107064478 | sodium-dependent phosphate transporter 2 | 0.04487 | 14.16426 | FALSE | FALSE |
| LOC107065721 | WD repeat-containing protein 78-like | 0.0464 | 14.16702 | FALSE | FALSE |
| LOC107067635 | TBC1 domain family member 14-like | 0.04464 | 14.17646 | FALSE | FALSE |
| LOC107072260 | nucleophosmin-like | 0.04512 | 14.19602 | FALSE | FALSE |
| LOC107071865 | uncharacterized protein LOC107071865 | 0.04634 | 14.20536 | FALSE | FALSE |
| LOC107071346 | putative fatty acyl-CoA reductase CG5065 | 0.04708 | 14.20866 | FALSE | FALSE |
| LOC107073972 | NAD kinase 2, mitochondrial-like | 0.04632 | 14.21196 | FALSE | FALSE |
| LOC107074327 | structural maintenance of chromosomes pr | 0.04638 | 14.22123 | FALSE | FALSE |
| LOC107066990 | uncharacterized protein LOC107066990 | 0.04655 | 14.22196 | FALSE | FALSE |
| LOC107065217 | transferrin-like | 0.04477 | 14.22269 | FALSE | FALSE |
| LOC107066540 | protein BOLA2-like | 0.04681 | 14.22838 | FALSE | FALSE |
| LOC107068335 | eukaryotic translation initiation factor 4 gan | 0.04581 | 14.23755 | FALSE | FALSE |
| LOC107067934 | limulus clotting factor C-like | 0.04543 | 14.25478 | FALSE | FALSE |
| LOC107063704 | indole-3-acetaldehyde oxidase-like | 0.04596 | 14.26112 | FALSE | FALSE |
| LOC107064196 | uncharacterized protein LOC107064196 | 0.0472 | 14.2874 | FALSE | FALSE |
| LOC107066845 | uncharacterized protein LOC107066845 | 0.04567 | 14.29295 | FALSE | FALSE |
| LOC107073091 | protein farnesyltransferase/geranylgeranylt | 0.0465 | 14.30592 | FALSE | FALSE |
| LOC107070823 | probable enoyl-CoA hydratase, mitochondri | 0.04611 | 14.30838 | FALSE | FALSE |
| LOC107072466 | adenylate cyclase type 8-like | 0.04608 | 14.31024 | FALSE | FALSE |
| LOC107067186 | iron-sulfur cluster assembly 1 homolog, mit | 0.04615 | 14.31348 | FALSE | FALSE |
| LOC107067321 | serine protease 44-like | 0.04583 | 14.32449 | FALSE | FALSE |
| LOC107072133 | hemicentin-1-like | 0.04578 | 14.32777 | FALSE | FALSE |
| LOC107066850 | protein phosphatase PP2A 55 kDa regulator | 0.04558 | 14.33298 | FALSE | FALSE |
| LOC107074614 | 26S proteasome non-ATPase regulatory sub | 0.04482 | 14.33782 | FALSE | FALSE |
| LOC107068250 | MICOS complex subunit Mic10-like | 0.04668 | 14.33851 | FALSE | FALSE |
| LOC107064001 | cohesin subunit SA-2 | 0.04601 | 14.33931 | FALSE | FALSE |
| LOC107073728 | uncharacterized protein LOC107073728 | 0.04433 | 14.34342 | FALSE | FALSE |
| LOC107070188 | uncharacterized protein LOC107070188 | 0.04511 | 14.35685 | FALSE | FALSE |
| LOC107066314 | transmembrane protein 68 | 0.04454 | 14.35952 | FALSE | FALSE |
| LOC107070121 | uncharacterized protein LOC107070121 | 0.04633 | 14.36118 | FALSE | FALSE |
| LOC107073795 | uncharacterized protein LOC107073795 | 0.04328 | 14.3622 | FALSE | FALSE |
| LOC107069580 | calcium uniporter protein, mitochondrial | 0.04623 | 14.36965 | FALSE | FALSE |
| LOC107074581 | myb-like protein X | 0.04627 | 14.37754 | FALSE | FALSE |
| LOC107067536 | protein unc-80 homolog | 0.04602 | 14.39187 | FALSE | FALSE |
| LOC107069710 | general odorant-binding protein 56a | 0.04464 | 14.39374 | FALSE | FALSE |
| LOC107071326 | uncharacterized protein LOC107071326 | 0.04522 | 14.39971 | FALSE | FALSE |
| LOC107074495 | spidroin-1-like | 0.04607 | 14.40505 | FALSE | FALSE |
| LOC107069824 | uncharacterized protein LOC107069824 | 0.04418 | 14.40951 | FALSE | FALSE |
| LOC107067898 | uncharacterized protein LOC107067898 | 0.04745 | 14.40963 | FALSE | FALSE |

| | | | | | |
|---|---|---|---|---|---|
| LOC107067187 | DNA-directed RNA polymerase II subunit RP | 0.04637 | 14.41027 | FALSE | FALSE |
| LOC107065296 | ubiquitin-like-conjugating enzyme ATG3 | 0.0461 | 14.41266 | FALSE | FALSE |
| LOC107066063 | cyclin-dependent kinase 2-associated protei | 0.04465 | 14.41687 | FALSE | FALSE |
| LOC107065913 | transmembrane protein 39A | 0.04605 | 14.42046 | FALSE | FALSE |
| LOC107067048 | uncharacterized protein LOC107067048 | 0.04575 | 14.42377 | FALSE | FALSE |
| LOC107065914 | 8-oxo-dGDP phosphatase NUDT18 | 0.04496 | 14.44299 | FALSE | FALSE |
| LOC107066233 | homeobox protein OTX2-B-like | 0.04549 | 14.45521 | FALSE | FALSE |
| LOC107069856 | BRO1 domain-containing protein BROX-like | 0.04554 | 14.47096 | FALSE | FALSE |
| LOC107069035 | uncharacterized protein LOC107069035 | 0.04654 | 14.47728 | FALSE | FALSE |
| LOC107067620 | uncharacterized protein LOC107067620 | 0.04417 | 14.48168 | FALSE | FALSE |
| LOC107073676 | uncharacterized protein LOC107073676 | 0.04621 | 14.48958 | FALSE | FALSE |
| LOC107063749 | rho GTPase-activating protein gacF-like | 0.04603 | 14.51032 | FALSE | FALSE |
| LOC107068751 | paired mesoderm homeobox protein 2B-like | 0.04499 | 14.52414 | FALSE | FALSE |
| LOC107073579 | uncharacterized protein LOC107073579 | 0.04431 | 14.53597 | FALSE | FALSE |
| LOC107072256 | G1/S-specific cyclin-D2-like | 0.04601 | 14.53658 | FALSE | FALSE |
| LOC107064919 | gamma-aminobutyric acid type B receptor s | 0.04564 | 14.53723 | FALSE | FALSE |
| LOC107072464 | uncharacterized protein LOC107072464 | 0.04526 | 14.55372 | FALSE | FALSE |
| LOC107073314 | chitooligosaccharidolytic beta-N-acetylgluco | 0.04713 | 14.55508 | FALSE | FALSE |
| LOC107070390 | transmembrane protein 14C | 0.04434 | 14.55623 | FALSE | FALSE |
| LOC107071097 | uncharacterized protein LOC107071097 | 0.04497 | 14.56081 | FALSE | FALSE |
| LOC107074378 | COX assembly mitochondrial protein homol | 0.04556 | 14.56135 | FALSE | FALSE |
| LOC107064686 | protein cornichon homolog 4 | 0.04653 | 14.57702 | FALSE | FALSE |
| LOC107069362 | protein Shroom-like | 0.04567 | 14.58603 | FALSE | FALSE |
| LOC107065497 | osmotic avoidance abnormal protein 3-like | 0.04504 | 14.61807 | FALSE | FALSE |
| LOC107071081 | craniofacial development protein 1 | 0.04592 | 14.61953 | FALSE | FALSE |
| LOC107069966 | SUN domain-containing ossification factor | 0.04514 | 14.62193 | FALSE | FALSE |
| LOC107065119 | U4/U6 small nuclear ribonucleoprotein Prp3 | 0.04582 | 14.63167 | FALSE | FALSE |
| LOC107073328 | uncharacterized protein LOC107073328 | 0.04443 | 14.63522 | FALSE | FALSE |
| LOC107070900 | 60S ribosomal protein L19 | 0.04533 | 14.64001 | FALSE | FALSE |
| LOC107073377 | uncharacterized protein LOC107073377 | 0.04402 | 14.6825 | FALSE | FALSE |
| LOC107066642 | tyrosine-protein kinase transmembrane rec | 0.04371 | 14.6848 | FALSE | FALSE |
| LOC107074320 | ethanolaminephosphotransferase 1-like | 0.04629 | 14.6947 | FALSE | FALSE |
| LOC107071181 | protein FAM177A1 | 0.0458 | 14.70576 | FALSE | FALSE |
| LOC107063664 | uncharacterized protein LOC107063664 | 0.04542 | 14.70748 | FALSE | FALSE |
| LOC107071224 | uncharacterized protein LOC107071224 | 0.04471 | 14.72519 | FALSE | FALSE |
| LOC107073490 | PBAN-type neuropeptides-like | 0.04491 | 14.74238 | FALSE | FALSE |
| LOC107071173 | dynamin | 0.04489 | 14.75197 | FALSE | FALSE |
| LOC107064691 | probable 28S ribosomal protein S6, mitocho | 0.04689 | 14.76331 | FALSE | FALSE |
| LOC107069325 | uncharacterized protein LOC107069325 | 0.04517 | 14.76418 | FALSE | FALSE |
| LOC107069599 | lipase member H-like | 0.04547 | 14.7732 | FALSE | FALSE |
| LOC107072641 | uncharacterized protein LOC107072641 | 0.04512 | 14.77927 | FALSE | FALSE |
| LOC107065426 | uncharacterized protein C9orf117 homolog | 0.04514 | 14.78542 | FALSE | FALSE |
| LOC107064950 | uncharacterized protein LOC107064950 | 0.04407 | 14.79227 | FALSE | FALSE |
| LOC107072294 | 1-phosphatidylinositol 3-phosphate 5-kinase | 0.04487 | 14.79575 | FALSE | FALSE |
| LOC107065302 | sterol regulatory element-binding protein 1 | 0.04652 | 14.81123 | FALSE | FALSE |
| LOC107074269 | pleckstrin homology-like domain family B m | 0.04427 | 14.83346 | FALSE | FALSE |
| LOC107069615 | ribonuclease P protein subunit p29 | 0.04503 | 14.85499 | FALSE | FALSE |
| LOC107066380 | E3 ubiquitin-protein ligase HERC2 | 0.04459 | 14.8677 | FALSE | FALSE |
| LOC107069965 | uncharacterized protein LOC107069965 | 0.04622 | 14.87355 | FALSE | FALSE |
| LOC107072359 | uncharacterized protein LOC107072359 | 0.04531 | 14.87659 | FALSE | FALSE |

| | | | | | |
|---|---|---|---|---|---|
| LOC107074573 | mitochondrial basic amino acids transporter | 0.04532 | 14.88341 | FALSE | FALSE |
| LOC107066583 | fumarate hydratase, mitochondrial-like | 0.04495 | 14.88533 | FALSE | FALSE |
| LOC107074579 | alcohol dehydrogenase [NADP(+) | 0.04538 | 14.89046 | FALSE | FALSE |
| LOC107064975 | LOW QUALITY PROTEIN: histone acetyltrans | 0.04705 | 14.89454 | FALSE | FALSE |
| LOC107074457 | G2/mitotic-specific cyclin-B3 | 0.04584 | 14.90757 | FALSE | FALSE |
| LOC107070289 | ragulator complex protein LAMTOR1 | 0.04596 | 14.90852 | FALSE | FALSE |
| LOC107074668 | CAAX prenyl protease 1 homolog | 0.04451 | 14.90864 | FALSE | FALSE |
| LOC107064619 | proteasome subunit beta type-5 | 0.04464 | 14.9095 | FALSE | FALSE |
| LOC107066929 | uncharacterized protein LOC107066929 | 0.04386 | 14.91244 | FALSE | FALSE |
| LOC107072473 | U4/U6.U5 tri-snRNP-associated protein 1 | 0.04615 | 14.93055 | FALSE | FALSE |
| LOC107063801 | synaptotagmin-2-like | 0.04473 | 14.94134 | FALSE | FALSE |
| LOC107069195 | uncharacterized protein LOC107069195 | 0.04463 | 14.94206 | FALSE | FALSE |
| LOC107068810 | dorsal-ventral patterning protein Sog | 0.04569 | 14.94793 | FALSE | FALSE |
| LOC107074510 | mediator of RNA polymerase II transcription | 0.04593 | 14.97322 | FALSE | FALSE |
| LOC107067676 | myb-like protein V | 0.04468 | 14.97569 | FALSE | FALSE |
| LOC107074247 | H/ACA ribonucleoprotein complex subunit 4 | 0.04331 | 14.98475 | FALSE | FALSE |
| LOC107068280 | uncharacterized protein MAL8P1.12-like | 0.04577 | 15.003 | FALSE | FALSE |
| LOC107074401 | uncharacterized protein LOC107074401 | 0.04515 | 15.00861 | FALSE | FALSE |
| LOC107068878 | uncharacterized protein LOC107068878 | 0.04478 | 15.01585 | FALSE | FALSE |
| LOC107068095 | synaptobrevin-1-like | 0.04548 | 15.02287 | FALSE | FALSE |
| LOC107064613 | diuretic hormone receptor-like | 0.04594 | 15.02534 | FALSE | FALSE |
| LOC107069852 | importin subunit alpha-7-like | 0.04548 | 15.0254 | FALSE | FALSE |
| LOC107065753 | probable serine/threonine-protein kinase m | 0.04566 | 15.02626 | FALSE | FALSE |
| LOC107066479 | N-alpha-acetyltransferase 15, NatA auxiliary | 0.04585 | 15.03508 | FALSE | FALSE |
| LOC107072965 | thioredoxin-related transmembrane protein | 0.0448 | 15.0368 | FALSE | FALSE |
| LOC107064459 | innexin inx2-like | 0.04557 | 15.06983 | FALSE | FALSE |
| LOC107065943 | peptidyl-prolyl cis-trans isomerase cyp11 | 0.04492 | 15.07381 | FALSE | FALSE |
| LOC107066448 | DNA-directed RNA polymerase I subunit RPA | 0.04479 | 15.07895 | FALSE | FALSE |
| LOC107066775 | uncharacterized protein LOC107066775 | 0.0463 | 15.09359 | FALSE | FALSE |
| LOC107069405 | ATP-binding cassette sub-family F member 3 | 0.04467 | 15.09508 | FALSE | FALSE |
| LOC107069506 | unconventional myosin-If-like | 0.04456 | 15.09906 | FALSE | FALSE |
| LOC107064429 | solute carrier family 35 member F6 | 0.04584 | 15.12528 | FALSE | FALSE |
| LOC107064012 | 60S ribosomal protein L35a | 0.04532 | 15.1336 | FALSE | FALSE |
| LOC107074271 | uncharacterized protein LOC107074271 | 0.04562 | 15.14872 | FALSE | FALSE |
| LOC107070139 | uncharacterized protein LOC107070139 | 0.04444 | 15.14896 | FALSE | FALSE |
| LOC107069742 | pescadillo homolog | 0.04469 | 15.1492 | FALSE | FALSE |
| LOC107066888 | uncharacterized protein LOC107066888 | 0.04379 | 15.15071 | FALSE | FALSE |
| LOC107067198 | transcription elongation factor B polypeptid | 0.04648 | 15.17355 | FALSE | FALSE |
| LOC107074662 | hemicentin-1-like | 0.04513 | 15.18342 | FALSE | FALSE |
| LOC107073692 | uncharacterized protein LOC107073692 | 0.04809 | 15.18759 | FALSE | FALSE |
| LOC107073451 | phosphatidylinositol-binding clathrin assem | 0.04509 | 15.19029 | FALSE | FALSE |
| LOC107070127 | hepatocyte nuclear factor 4-gamma | 0.04433 | 15.23501 | FALSE | FALSE |
| LOC107064849 | uncharacterized protein LOC107064849 | 0.04352 | 15.2412 | FALSE | FALSE |
| LOC107071272 | serine/threonine-protein phosphatase 4 reg | 0.044 | 15.24353 | FALSE | FALSE |
| LOC107073259 | histone-lysine N-methyltransferase SETMAR | 0.04579 | 15.24535 | FALSE | FALSE |
| LOC107071145 | 70 kDa peptidyl-prolyl isomerase-like | 0.04502 | 15.25038 | FALSE | FALSE |
| LOC107070290 | dentin sialophosphoprotein | 0.04481 | 15.25213 | FALSE | FALSE |
| LOC107071448 | uncharacterized protein LOC107071448 | 0.04548 | 15.26877 | FALSE | FALSE |
| LOC107070413 | general transcription factor IIH subunit 2 | 0.04385 | 15.26967 | FALSE | FALSE |
| LOC107066423 | uncharacterized protein LOC107066423 | 0.04437 | 15.28744 | FALSE | FALSE |

| | | | | | |
|---|---|---|---|---|---|
| LOC107064477 | EF-hand domain-containing protein 1-like | 0.04345 | 15.30349 | FALSE | FALSE |
| LOC107065132 | uncharacterized protein LOC107065132 | 0.04303 | 15.31079 | FALSE | FALSE |
| LOC107066670 | G patch domain-containing protein 2 | 0.04458 | 15.3162 | FALSE | FALSE |
| LOC107066648 | F-box only protein 28 | 0.04574 | 15.31971 | FALSE | FALSE |
| LOC107071917 | glycine-rich protein DOT1-like | 0.04493 | 15.3271 | FALSE | FALSE |
| LOC107069020 | tetraspanin-9 | 0.04532 | 15.33468 | FALSE | FALSE |
| LOC107069310 | myrosinase 1-like | 0.04453 | 15.33524 | FALSE | FALSE |
| LOC107073856 | uncharacterized protein LOC107073856 | 0.04525 | 15.34328 | FALSE | FALSE |
| LOC107074699 | myrosinase 1-like | 0.04549 | 15.35088 | FALSE | FALSE |
| LOC107066655 | inositol-tetrakisphosphate 1-kinase-like | 0.04743 | 15.35451 | FALSE | FALSE |
| LOC107066189 | uncharacterized protein LOC107066189 | 0.04505 | 15.38379 | FALSE | FALSE |
| LOC107067648 | mitochondrial ornithine transporter 1 | 0.04338 | 15.39394 | FALSE | FALSE |
| LOC107070509 | differentially expressed in FDCP 8 homolog | 0.04649 | 15.3947 | FALSE | FALSE |
| LOC107067782 | nuclear receptor coactivator 2 | 0.04679 | 15.39678 | FALSE | FALSE |
| LOC107066618 | esterase E4-like | 0.04555 | 15.39758 | FALSE | FALSE |
| LOC107064826 | transcription factor SOX-9-like | 0.04507 | 15.39854 | FALSE | FALSE |
| LOC107065897 | uncharacterized protein LOC107065897 | 0.04359 | 15.40391 | FALSE | FALSE |
| LOC107067790 | uncharacterized protein LOC107067790 | 0.04401 | 15.41101 | FALSE | FALSE |
| LOC107072176 | cell wall protein IFF6 | 0.04431 | 15.41694 | FALSE | FALSE |
| LOC107072367 | uncharacterized protein LOC107072367 | 0.0445 | 15.44274 | FALSE | FALSE |
| LOC107064824 | transcription factor Sox-8-like | 0.04559 | 15.45003 | FALSE | FALSE |
| LOC107065280 | inositol-pentakisphosphate 2-kinase | 0.0445 | 15.45322 | FALSE | FALSE |
| LOC107073297 | eukaryotic translation initiation factor 2A | 0.04384 | 15.46272 | FALSE | FALSE |
| LOC107071424 | serine/arginine-rich splicing factor 11-like | 0.04468 | 15.46541 | FALSE | FALSE |
| LOC107071262 | kinectin-like | 0.04663 | 15.47062 | FALSE | FALSE |
| LOC107067394 | uncharacterized protein LOC107067394 | 0.04727 | 15.47275 | FALSE | FALSE |
| LOC107068631 | BMP and activin membrane-bound inhibitor | 0.04458 | 15.47674 | FALSE | FALSE |
| LOC107066700 | voltage-dependent L-type calcium channel s | 0.0447 | 15.48908 | FALSE | FALSE |
| LOC107073920 | uncharacterized protein LOC107073920 | 0.0445 | 15.49286 | FALSE | FALSE |
| LOC107072204 | uncharacterized protein LOC107072204 | 0.0453 | 15.50282 | FALSE | FALSE |
| LOC107070151 | flavin-containing monooxygenase FMO GS-( | 0.04548 | 15.51221 | FALSE | FALSE |
| LOC107066585 | uncharacterized protein LOC107066585 | 0.04513 | 15.51804 | FALSE | FALSE |
| LOC107069184 | charged multivesicular body protein 3 | 0.04482 | 15.53383 | FALSE | FALSE |
| LOC107070409 | uncharacterized protein LOC107070409 | 0.04677 | 15.53468 | FALSE | FALSE |
| LOC107072749 | protein EMSY | 0.04403 | 15.54089 | FALSE | FALSE |
| LOC107065047 | bestrophin homolog 15 | 0.04667 | 15.54479 | FALSE | FALSE |
| LOC107071789 | ubiquitin carboxyl-terminal hydrolase 47 | 0.04502 | 15.56187 | FALSE | FALSE |
| LOC107072527 | equilibrative nucleoside transporter 4 | 0.04568 | 15.56948 | FALSE | FALSE |
| LOC107073860 | epidermal growth factor receptor substrate | 0.04421 | 15.58654 | FALSE | FALSE |
| LOC107072810 | facilitated trehalose transporter Tret1-like | 0.04577 | 15.59314 | FALSE | FALSE |
| LOC107068321 | uncharacterized protein LOC107068321 | 0.04602 | 15.60428 | FALSE | FALSE |
| LOC107064726 | maltase 1-like | 0.04409 | 15.61001 | FALSE | FALSE |
| LOC107065836 | anoctamin-8-like | 0.04387 | 15.61436 | FALSE | FALSE |
| LOC107068231 | protein PF14_0175-like | 0.04507 | 15.61643 | FALSE | FALSE |
| LOC107066994 | serine-rich adhesin for platelets-like | 0.04536 | 15.61788 | FALSE | FALSE |
| LOC107073021 | pro-resilin | 0.04415 | 15.62002 | FALSE | FALSE |
| LOC107073632 | uncharacterized protein LOC107073632 | 0.04297 | 15.62816 | FALSE | FALSE |
| LOC107070871 | zinc finger protein 37-like | 0.04394 | 15.63182 | FALSE | FALSE |
| LOC107063822 | transmembrane protein 115 | 0.04374 | 15.63805 | FALSE | FALSE |
| LOC107071488 | putative U5 small nuclear ribonucleoprotein | 0.04432 | 15.67134 | FALSE | FALSE |

| | | | | | |
|---|---|---|---|---|---|
| LOC107072702 | peptidyl-prolyl cis-trans isomerase-like 4 | 0.04535 | 15.68051 | FALSE | FALSE |
| LOC107068839 | ribosomal RNA-processing protein 7 homolo | 0.04418 | 15.68529 | FALSE | FALSE |
| LOC107069376 | plancitoxin-1 | 0.04454 | 15.68838 | FALSE | FALSE |
| LOC107071869 | putative mediator of RNA polymerase II trar | 0.04482 | 15.69311 | FALSE | FALSE |
| LOC107068117 | testis-expressed sequence 10 protein homo | 0.04391 | 15.6974 | FALSE | FALSE |
| LOC107072280 | DNA ligase 1 | 0.04263 | 15.70224 | FALSE | FALSE |
| LOC107064527 | 40S ribosomal protein S28 | 0.04323 | 15.7162 | FALSE | FALSE |
| LOC107067938 | uncharacterized protein LOC107067938 | 0.04364 | 15.72145 | FALSE | FALSE |
| LOC107074701 | regulator of telomere elongation helicase 1 | 0.04313 | 15.72444 | FALSE | FALSE |
| LOC107070135 | prohormone-3 | 0.04443 | 15.7315 | FALSE | FALSE |
| LOC107067967 | (E3-independent) E2 ubiquitin-conjugating e | 0.04474 | 15.7335 | FALSE | FALSE |
| LOC107066699 | hemocytin | 0.04413 | 15.75561 | FALSE | FALSE |
| LOC107069912 | ras-responsive element-binding protein 1 | 0.04385 | 15.75982 | FALSE | FALSE |
| LOC107065624 | uncharacterized protein LOC107065624 | 0.04537 | 15.76948 | FALSE | FALSE |
| LOC107067992 | heparanase-like | 0.04361 | 15.77123 | FALSE | FALSE |
| LOC107069485 | protein O-mannosyl-transferase 2-like | 0.04435 | 15.77133 | FALSE | FALSE |
| LOC107074146 | mitochondrial import inner membrane trans | 0.04539 | 15.78107 | FALSE | FALSE |
| LOC107063678 | serine protease easter-like | 0.04495 | 15.7869 | FALSE | FALSE |
| LOC107067269 | vitamin K epoxide reductase complex subur | 0.04564 | 15.79074 | FALSE | FALSE |
| LOC107072394 | vesicular inhibitory amino acid transporter | 0.04416 | 15.79482 | FALSE | FALSE |
| LOC107067622 | uncharacterized protein LOC107067622 | 0.04474 | 15.7965 | FALSE | FALSE |
| LOC107074024 | fork head domain-containing protein FD4 | 0.04517 | 15.80362 | FALSE | FALSE |
| LOC107069776 | coiled-coil and C2 domain-containing protei | 0.04575 | 15.81282 | FALSE | FALSE |
| LOC107068329 | trafficking protein particle complex subunit | 0.04632 | 15.81422 | FALSE | FALSE |
| LOC107065200 | SAP domain-containing ribonucleoprotein | 0.04392 | 15.83406 | FALSE | FALSE |
| LOC107073265 | uncharacterized protein LOC107073265 | 0.04505 | 15.83969 | FALSE | FALSE |
| LOC107067826 | spermine oxidase-like | 0.0444 | 15.84885 | FALSE | FALSE |
| LOC107070333 | cyclin-Y-like protein 1 | 0.04444 | 15.8566 | FALSE | FALSE |
| LOC107065561 | uncharacterized protein LOC107065561 | 0.04605 | 15.8623 | FALSE | FALSE |
| LOC107065409 | myb-like protein AA | 0.04347 | 15.87049 | FALSE | FALSE |
| LOC107074116 | cathepsin O-like | 0.04913 | 15.87351 | FALSE | FALSE |
| LOC107064794 | selenocysteine insertion sequence-binding p | 0.04497 | 15.87824 | FALSE | FALSE |
| LOC107073940 | slit homolog 2 protein-like | 0.04526 | 15.90423 | FALSE | FALSE |
| LOC107073923 | sorting nexin-30-like | 0.0441 | 15.91726 | FALSE | FALSE |
| LOC107066171 | dnaJ homolog subfamily C member 22 | 0.04538 | 15.92276 | FALSE | FALSE |
| LOC107072572 | facilitated trehalose transporter Tret1-2 hor | 0.04394 | 15.93261 | FALSE | FALSE |
| LOC107065116 | solute carrier family 2, facilitated glucose tra | 0.04339 | 15.94115 | FALSE | FALSE |
| LOC107071621 | glycerol kinase | 0.04471 | 15.95031 | FALSE | FALSE |
| LOC107068286 | 26S proteasome non-ATPase regulatory sub | 0.04542 | 15.95114 | FALSE | FALSE |
| LOC107071608 | uncharacterized protein LOC107071608 | 0.0446 | 15.9515 | FALSE | FALSE |
| LOC107066078 | intraflagellar transport protein 46 homolog | 0.04409 | 15.95217 | FALSE | FALSE |
| LOC107073969 | ceramide synthase 5-like | 0.04635 | 15.96244 | FALSE | FALSE |
| LOC107064496 | uncharacterized protein LOC107064496 | 0.04424 | 15.98569 | FALSE | FALSE |
| LOC107074551 | nuclear anchorage protein 1-like | 0.04439 | 15.98814 | FALSE | FALSE |
| LOC107070791 | uncharacterized protein LOC107070791 | 0.04448 | 15.99714 | FALSE | FALSE |
| LOC107066050 | transformer-2 protein homolog beta | 0.04512 | 15.9975 | FALSE | FALSE |
| LOC107067727 | superkiller viralicidic activity 2-like 2 | 0.04423 | 16.00331 | FALSE | FALSE |
| LOC107066346 | protein PFC0760c-like | 0.04381 | 16.00631 | FALSE | FALSE |
| LOC107074412 | uncharacterized protein CG7065-like | 0.04342 | 16.01674 | FALSE | FALSE |
| LOC107068951 | protein-associating with the carboxyl-termir | 0.04567 | 16.02137 | FALSE | FALSE |

| | | | | | |
|---|---|---|---|---|---|
| LOC107068917 | odorant receptor 13a-like | 0.04524 | 16.03327 | FALSE | FALSE |
| LOC107070198 | sin3 histone deacetylase corepressor compl | 0.04376 | 16.04346 | FALSE | FALSE |
| LOC107073346 | putative nuclease HARBI1 | 0.04522 | 16.04531 | FALSE | FALSE |
| LOC107069317 | protein furry | 0.04452 | 16.05406 | FALSE | FALSE |
| LOC107074464 | uncharacterized protein C19orf47 homolog | 0.04512 | 16.06269 | FALSE | FALSE |
| LOC107066217 | D-aspartate oxidase | 0.04434 | 16.08704 | FALSE | FALSE |
| LOC107073746 | putative fatty acyl-CoA reductase CG5065 | 0.04474 | 16.08719 | FALSE | FALSE |
| LOC107064027 | uncharacterized protein LOC107064027 | 0.04493 | 16.08996 | FALSE | FALSE |
| LOC107070369 | probable cytochrome P450 6a13 | 0.04381 | 16.1065 | FALSE | FALSE |
| LOC107070571 | neurochondrin homolog | 0.04244 | 16.1163 | FALSE | FALSE |
| LOC107063775 | probable maleylacetoacetate isomerase 2 | 0.04447 | 16.12652 | FALSE | FALSE |
| LOC107066647 | 60S ribosomal protein L12 | 0.04646 | 16.12873 | FALSE | FALSE |
| LOC107071318 | uncharacterized protein LOC107071318 | 0.04541 | 16.13274 | FALSE | FALSE |
| LOC107073693 | uncharacterized protein LOC107073693 | 0.04401 | 16.13435 | FALSE | FALSE |
| LOC107066256 | leucine-rich repeat protein soc-2 homolog | 0.04403 | 16.15493 | FALSE | FALSE |
| LOC107072441 | putative ATP-dependent RNA helicase DHX3 | 0.04395 | 16.16343 | FALSE | FALSE |
| LOC107069414 | uncharacterized protein LOC107069414 | 0.04501 | 16.16965 | FALSE | FALSE |
| LOC107065207 | DTW domain-containing protein 1 | 0.04464 | 16.17099 | FALSE | FALSE |
| LOC107069600 | myrosinase 1 | 0.0446 | 16.18235 | FALSE | FALSE |
| LOC107065253 | D-glucuronyl C5-epimerase | 0.04532 | 16.18257 | FALSE | FALSE |
| LOC107066132 | uncharacterized protein LOC107066132 | 0.04245 | 16.19693 | FALSE | FALSE |
| LOC107072285 | guanine nucleotide-binding protein subunit | 0.04402 | 16.20869 | FALSE | FALSE |
| LOC107064069 | WW domain-binding protein 4 | 0.04507 | 16.24791 | FALSE | FALSE |
| LOC107071807 | glyoxylate reductase/hydroxypyruvate redu | 0.04398 | 16.25112 | FALSE | FALSE |
| LOC107066071 | 60S ribosomal protein L17 | 0.04486 | 16.25412 | FALSE | FALSE |
| LOC107065163 | neurogenic locus protein delta-like | 0.04392 | 16.26012 | FALSE | FALSE |
| LOC107069473 | U3 small nucleolar RNA-interacting protein | 0.04387 | 16.26509 | FALSE | FALSE |
| LOC107065928 | nibrin | 0.04397 | 16.28321 | FALSE | FALSE |
| LOC107064311 | exocyst complex component 7 | 0.04452 | 16.29684 | FALSE | FALSE |
| LOC107064912 | ADP-ribosylation factor-like protein 6 | 0.04503 | 16.31672 | FALSE | FALSE |
| LOC107067621 | uncharacterized protein LOC107067621 | 0.04283 | 16.31714 | FALSE | FALSE |
| LOC107067950 | uncharacterized protein LOC107067950 | 0.04406 | 16.34082 | FALSE | FALSE |
| LOC107064365 | uncharacterized protein LOC107064365 | 0.04344 | 16.36616 | FALSE | FALSE |
| LOC107073855 | E3 ubiquitin-protein ligase CBL-B | 0.0443 | 16.36971 | FALSE | FALSE |
| LOC107066004 | phytanoyl-CoA dioxygenase, peroxisomal-lik | 0.04458 | 16.37932 | FALSE | FALSE |
| LOC107068215 | erythroid differentiation-related factor 1 | 0.04331 | 16.39217 | FALSE | FALSE |
| LOC107069179 | uncharacterized protein LOC107069179 | 0.04381 | 16.39526 | FALSE | FALSE |
| LOC107074498 | uncharacterized protein LOC107074498 | 0.04381 | 16.41222 | FALSE | FALSE |
| LOC107064823 | CD82 antigen-like | 0.04402 | 16.41862 | FALSE | FALSE |
| LOC107065898 | PMS1 protein homolog 1-like | 0.04552 | 16.42865 | FALSE | FALSE |
| LOC107070306 | uncharacterized protein LOC107070306 | 0.04465 | 16.43095 | FALSE | FALSE |
| LOC107064207 | uncharacterized protein LOC107064207 | 0.04324 | 16.43619 | FALSE | FALSE |
| LOC107063795 | uncharacterized protein LOC107063795 | 0.04378 | 16.43986 | FALSE | FALSE |
| LOC107070468 | dihydrolipoyl dehydrogenase, mitochondria | 0.04482 | 16.44519 | FALSE | FALSE |
| LOC107068932 | splicing factor 3B subunit 1 | 0.04436 | 16.4481 | FALSE | FALSE |
| LOC107066422 | probable ATP-dependent RNA helicase DDX | 0.04307 | 16.44976 | FALSE | FALSE |
| LOC107067585 | protein disulfide-isomerase TMX3 | 0.04428 | 16.45276 | FALSE | FALSE |
| LOC107065195 | LOW QUALITY PROTEIN: protein SHQ1 hom | 0.04447 | 16.45354 | FALSE | FALSE |
| LOC107072870 | uncharacterized protein C20orf24 homolog | 0.04461 | 16.45675 | FALSE | FALSE |
| LOC107070811 | actin-related protein 8 | 0.04595 | 16.47085 | FALSE | FALSE |

| | | | | | |
|---|---|---|---|---|---|
| LOC107067444 | protein-L-isoaspartate O-methyltransferase | 0.04329 | 16.47155 | FALSE | FALSE |
| LOC107067402 | afadin- and alpha-actinin-binding protein-lik | 0.04387 | 16.48138 | FALSE | FALSE |
| LOC107068719 | uncharacterized protein LOC107068719 | 0.04529 | 16.48221 | FALSE | FALSE |
| LOC107070961 | peroxiredoxin-5, mitochondrial | 0.04291 | 16.49244 | FALSE | FALSE |
| LOC107072405 | eukaryotic translation initiation factor 2 sub | 0.04337 | 16.49861 | FALSE | FALSE |
| LOC107067715 | bumetanide-sensitive sodium-(potassium)-c | 0.04277 | 16.49884 | FALSE | FALSE |
| LOC107068940 | probable glucosamine 6-phosphate N-acety | 0.04424 | 16.50048 | FALSE | FALSE |
| LOC107068255 | uncharacterized protein LOC107068255 | 0.04493 | 16.50901 | FALSE | FALSE |
| LOC107064037 | microtubule-associated protein futsch | 0.04288 | 16.52473 | FALSE | FALSE |
| LOC107074480 | LOW QUALITY PROTEIN: methyltransferase- | 0.04423 | 16.53323 | FALSE | FALSE |
| LOC107064539 | rap1 GTPase-activating protein 1-like | 0.04482 | 16.54264 | FALSE | FALSE |
| LOC107074431 | inositol monophosphatase 1-like | 0.04503 | 16.54992 | FALSE | FALSE |
| LOC107069449 | importin subunit alpha-4 | 0.04363 | 16.55469 | FALSE | FALSE |
| LOC107071490 | janus kinase and microtubule-interacting pr | 0.04469 | 16.55517 | FALSE | FALSE |
| LOC107069820 | uncharacterized protein LOC107069820 | 0.04379 | 16.57032 | FALSE | FALSE |
| LOC107067291 | single-strand selective monofunctional urac | 0.04454 | 16.58028 | FALSE | FALSE |
| LOC107070459 | syntaxin-12 | 0.0435 | 16.59995 | FALSE | FALSE |
| LOC107066008 | uridine diphosphate glucose pyrophosphata | 0.04417 | 16.6222 | FALSE | FALSE |
| LOC107064014 | dymeclin | 0.04407 | 16.63001 | FALSE | FALSE |
| LOC107067510 | cryptochrome-1-like | 0.04382 | 16.63771 | FALSE | FALSE |
| LOC107067911 | odorant receptor 4-like | 0.04423 | 16.64142 | FALSE | FALSE |
| LOC107071120 | thyroid receptor-interacting protein 11-like | 0.04419 | 16.64601 | FALSE | FALSE |
| LOC107065824 | tudor and KH domain-containing protein | 0.04498 | 16.65958 | FALSE | FALSE |
| LOC107073472 | cytochrome P450 4V2-like | 0.04577 | 16.67594 | FALSE | FALSE |
| LOC107074683 | uncharacterized protein LOC107074683 | 0.04452 | 16.68096 | FALSE | FALSE |
| LOC107071672 | transmembrane protein 230 | 0.04527 | 16.68854 | FALSE | FALSE |
| LOC107064114 | menin | 0.04445 | 16.70675 | FALSE | FALSE |
| LOC107074660 | phosphatidate phosphatase PPAPDC1B | 0.04464 | 16.70806 | FALSE | FALSE |
| LOC107069494 | cadherin-87A | 0.04292 | 16.70815 | FALSE | FALSE |
| LOC107073343 | 28 kDa heat- and acid-stable phosphoprotei | 0.04508 | 16.7175 | FALSE | FALSE |
| LOC107067787 | tubulin alpha chain-like | 0.04442 | 16.73794 | FALSE | FALSE |
| LOC107072517 | uncharacterized protein LOC107072517 | 0.04316 | 16.73942 | FALSE | FALSE |
| LOC107070025 | alpha-glucosidase-like | 0.04487 | 16.74902 | FALSE | FALSE |
| LOC107064584 | zinc finger protein 271-like | 0.0442 | 16.77357 | FALSE | FALSE |
| LOC107067143 | uncharacterized protein LOC107067143 | 0.04396 | 16.79893 | FALSE | FALSE |
| LOC107069223 | CD9 antigen | 0.04443 | 16.8191 | FALSE | FALSE |
| LOC107071283 | macoilin-1-like | 0.04228 | 16.82724 | FALSE | FALSE |
| LOC107067573 | uncharacterized protein LOC107067573 | 0.04421 | 16.83493 | FALSE | FALSE |
| LOC107069368 | uncharacterized protein LOC107069368 | 0.04332 | 16.85539 | FALSE | FALSE |
| LOC107071529 | regulator of nonsense transcripts 2 | 0.04369 | 16.87942 | FALSE | FALSE |
| LOC107071101 | endoribonuclease Dcr-1 | 0.04427 | 16.88251 | FALSE | FALSE |
| LOC107067159 | fizzy-related protein homolog | 0.0442 | 16.89826 | FALSE | FALSE |
| LOC107064208 | probable ATP-dependent RNA helicase DDX | 0.04301 | 16.90962 | FALSE | FALSE |
| LOC107072508 | non-canonical poly(A) RNA polymerase PAP | 0.04283 | 16.92034 | FALSE | FALSE |
| LOC107074444 | transcriptional protein SWT1-like | 0.04379 | 16.92192 | FALSE | FALSE |
| LOC107073368 | snRNA-activating protein complex subunit 1 | 0.04287 | 16.93162 | FALSE | FALSE |
| LOC107064922 | pupal cuticle protein C1B-like | 0.04454 | 16.93167 | FALSE | FALSE |
| LOC107070743 | uncharacterized protein LOC107070743 | 0.04483 | 16.93168 | FALSE | FALSE |
| LOC107069113 | protein decapentaplegic-like | 0.04474 | 16.95384 | FALSE | FALSE |
| LOC107063931 | protein fem-1 homolog B | 0.0434 | 16.95784 | FALSE | FALSE |

| | | | | | |
|---|---|---|---|---|---|
| LOC107064930 | ATP-dependent RNA helicase DBP2-like | 0.04382 | 16.97428 | FALSE | FALSE |
| LOC107066814 | 60S ribosomal protein L4 | 0.04211 | 16.97783 | FALSE | FALSE |
| LOC107071236 | potassium voltage-gated channel subfamily | 0.04473 | 16.98002 | FALSE | FALSE |
| LOC107071572 | acyl carrier protein, mitochondrial | 0.04486 | 16.98399 | FALSE | FALSE |
| LOC107070097 | N-acetylated-alpha-linked acidic dipeptidase | 0.04388 | 16.98683 | FALSE | FALSE |
| LOC107074301 | ubiquitin-conjugating enzyme E2-22 kDa | 0.04547 | 16.98693 | FALSE | FALSE |
| LOC107070298 | complement component 1 Q subcomponen | 0.04497 | 16.99767 | FALSE | FALSE |
| LOC107065701 | uncharacterized protein LOC107065701 | 0.04211 | 17.00022 | FALSE | FALSE |
| LOC107067503 | uncharacterized protein LOC107067503 | 0.04255 | 17.0246 | FALSE | FALSE |
| LOC107068610 | translation initiation factor IF-2-like | 0.04346 | 17.0358 | FALSE | FALSE |
| LOC107064380 | type-1 angiotensin II receptor-associated pr | 0.04367 | 17.03709 | FALSE | FALSE |
| LOC107068507 | uncharacterized protein LOC107068507 | 0.04173 | 17.04671 | FALSE | FALSE |
| LOC107074336 | facilitated trehalose transporter Tret1 | 0.0436 | 17.0558 | FALSE | FALSE |
| LOC107066107 | uncharacterized protein LOC107066107 | 0.04499 | 17.06484 | FALSE | FALSE |
| LOC107072295 | Kv channel-interacting protein 1-like | 0.04457 | 17.0766 | FALSE | FALSE |
| LOC107064546 | uncharacterized protein LOC107064546 | 0.04263 | 17.09173 | FALSE | FALSE |
| LOC107073069 | frataxin homolog, mitochondrial | 0.0441 | 17.09846 | FALSE | FALSE |
| LOC107072844 | ATP-citrate synthase | 0.04365 | 17.09866 | FALSE | FALSE |
| LOC107065850 | uncharacterized protein LOC107065850 | 0.04375 | 17.11431 | FALSE | FALSE |
| LOC107072344 | acylamino-acid-releasing enzyme-like | 0.04412 | 17.11477 | FALSE | FALSE |
| LOC107072012 | protein capicua homolog | 0.04455 | 17.11616 | FALSE | FALSE |
| LOC107066645 | translocon-associated protein subunit alpha | 0.04297 | 17.12329 | FALSE | FALSE |
| LOC107065811 | uncharacterized protein LOC107065811 | 0.04335 | 17.12509 | FALSE | FALSE |
| LOC107070197 | solute carrier family 35 member E2-like | 0.04391 | 17.14448 | FALSE | FALSE |
| LOC107063967 | dipeptidyl peptidase 3 | 0.04395 | 17.15473 | FALSE | FALSE |
| LOC107072579 | inosine-5'-monophosphate dehydrogenase | 0.04478 | 17.16268 | FALSE | FALSE |
| LOC107064644 | trichoplein keratin filament-binding protein | 0.04435 | 17.16436 | FALSE | FALSE |
| LOC107067080 | deformed epidermal autoregulatory factor | 0.04442 | 17.16633 | FALSE | FALSE |
| LOC107064866 | protein croquemort-like | 0.04393 | 17.17124 | FALSE | FALSE |
| LOC107072023 | hemK methyltransferase family member 2 | 0.04522 | 17.18242 | FALSE | FALSE |
| LOC107071027 | protein FAM8A1 | 0.04465 | 17.19745 | FALSE | FALSE |
| LOC107072542 | putative uncharacterized protein DDB_G028 | 0.04365 | 17.20428 | FALSE | FALSE |
| LOC107065241 | 15-hydroxyprostaglandin dehydrogenase [N | 0.04479 | 17.20434 | FALSE | FALSE |
| LOC107069459 | bromodomain-containing protein DDB_G02 | 0.04402 | 17.2259 | FALSE | FALSE |
| LOC107071443 | uncharacterized protein LOC107071443 | 0.04396 | 17.23441 | FALSE | FALSE |
| LOC107072249 | glycine dehydrogenase (decarboxylating), m | 0.04449 | 17.23735 | FALSE | FALSE |
| LOC107069235 | nicastrin | 0.04382 | 17.2402 | FALSE | FALSE |
| LOC107068811 | uncharacterized protein LOC107068811 | 0.04378 | 17.24124 | FALSE | FALSE |
| LOC107067593 | translocation protein SEC62 | 0.04514 | 17.26876 | FALSE | FALSE |
| LOC107070896 | uncharacterized protein LOC107070896 | 0.04421 | 17.26893 | FALSE | FALSE |
| LOC107068094 | fatty acid-binding protein 12-like | 0.04417 | 17.27151 | FALSE | FALSE |
| LOC107068833 | serine/threonine-protein kinase fused | 0.04506 | 17.29846 | FALSE | FALSE |
| LOC107073183 | rho-associated protein kinase 2 | 0.04362 | 17.29957 | FALSE | FALSE |
| LOC107064970 | snRNA-activating protein complex subunit 4 | 0.04439 | 17.30842 | FALSE | FALSE |
| LOC107071716 | serine/threonine-protein phosphatase 2B ca | 0.04473 | 17.32337 | FALSE | FALSE |
| LOC107070555 | delta-1-pyrroline-5-carboxylate synthase | 0.04338 | 17.32633 | FALSE | FALSE |
| LOC107068902 | uncharacterized protein LOC107068902 | 0.04486 | 17.33 | FALSE | FALSE |
| LOC107066746 | probable 39S ribosomal protein L24, mitoch | 0.04247 | 17.34202 | FALSE | FALSE |
| LOC107072659 | laminin subunit beta-1 | 0.04292 | 17.34578 | FALSE | FALSE |
| LOC107074013 | major royal jelly protein 1-like | 0.04458 | 17.35888 | FALSE | FALSE |

| | | | | | |
|---|---|---|---|---|---|
| LOC107068703 | endocuticle structural glycoprotein SgAbd-2 | 0.04347 | 17.38609 | FALSE | FALSE |
| LOC107065485 | uncharacterized protein LOC107065485 | 0.04359 | 17.39404 | FALSE | FALSE |
| LOC107065716 | transcription factor AP-1-like | 0.0431 | 17.3971 | FALSE | FALSE |
| LOC107070321 | putative uncharacterized protein DDB_G028 | 0.04389 | 17.40012 | FALSE | FALSE |
| LOC107070528 | DNA-directed RNA polymerases I and III sub | 0.04514 | 17.40082 | FALSE | FALSE |
| LOC107066036 | uncharacterized protein LOC107066036 | 0.04461 | 17.40189 | FALSE | FALSE |
| LOC107073182 | ribosomal protein VAR1, mitochondrial-like | 0.04337 | 17.40943 | FALSE | FALSE |
| LOC107068435 | excitatory amino acid transporter 3 | 0.04237 | 17.41091 | FALSE | FALSE |
| LOC107069648 | clathrin heavy chain | 0.04285 | 17.41316 | FALSE | FALSE |
| LOC107069735 | BMP-binding endothelial regulator protein | 0.04292 | 17.41691 | FALSE | FALSE |
| LOC107074588 | homeobox protein goosecoid-like | 0.04394 | 17.43938 | FALSE | FALSE |
| LOC107072357 | hemicentin-1-like | 0.04282 | 17.44601 | FALSE | FALSE |
| LOC107073881 | PAB-dependent poly(A)-specific ribonucleas | 0.04322 | 17.44702 | FALSE | FALSE |
| LOC107070038 | uncharacterized protein LOC107070038 | 0.04442 | 17.48025 | FALSE | FALSE |
| LOC107069855 | probable isocitrate dehydrogenase [NAD | 0.04376 | 17.48423 | FALSE | FALSE |
| LOC107065797 | nuclear receptor subfamily 2 group F memb | 0.04436 | 17.48688 | FALSE | FALSE |
| LOC107074514 | SEC14-like protein 3 | 0.04328 | 17.49964 | FALSE | FALSE |
| LOC107068332 | uncharacterized protein LOC107068332 | 0.0448 | 17.50631 | FALSE | FALSE |
| LOC107071017 | hamartin | 0.04368 | 17.52759 | FALSE | FALSE |
| LOC107071339 | activator of 90 kDa heat shock protein ATPa | 0.04468 | 17.53483 | FALSE | FALSE |
| LOC107063755 | nuclear envelope integral membrane protei | 0.04221 | 17.56281 | FALSE | FALSE |
| LOC107064829 | IQ domain-containing protein K-like | 0.04338 | 17.57039 | FALSE | FALSE |
| LOC107065608 | probable tRNA N6-adenosine threonylcarba | 0.04602 | 17.57778 | FALSE | FALSE |
| LOC107068299 | U6 snRNA-associated Sm-like protein LSm4 | 0.04413 | 17.57922 | FALSE | FALSE |
| LOC107068561 | Niemann-Pick C1 protein-like | 0.04404 | 17.58591 | FALSE | FALSE |
| LOC107072272 | dynein beta chain, ciliary-like | 0.04452 | 17.60516 | FALSE | FALSE |
| LOC107063952 | neuropeptide-like 1 | 0.04345 | 17.60856 | FALSE | FALSE |
| LOC107069932 | asparagine synthetase [glutamine-hydrolyzi | 0.04257 | 17.61072 | FALSE | FALSE |
| LOC107066375 | activin receptor type-2B | 0.04248 | 17.62295 | FALSE | FALSE |
| LOC107066499 | uncharacterized protein LOC107066499 | 0.04253 | 17.62496 | FALSE | FALSE |
| LOC107072079 | BTB/POZ domain-containing protein KCTD9 | 0.04296 | 17.63319 | FALSE | FALSE |
| LOC107071085 | nicotinamidase-like | 0.04292 | 17.63663 | FALSE | FALSE |
| LOC107064776 | mitochondrial protein import protein ZIM17 | 0.04245 | 17.65981 | FALSE | FALSE |
| LOC107066532 | uncharacterized protein LOC107066532 | 0.04443 | 17.67296 | FALSE | FALSE |
| LOC107068305 | two pore potassium channel protein sup-9 | 0.0436 | 17.68237 | FALSE | FALSE |
| LOC107065320 | solute carrier family 23 member 2 | 0.04319 | 17.68367 | FALSE | FALSE |
| LOC107067190 | WD repeat-containing protein 43-like | 0.04322 | 17.70333 | FALSE | FALSE |
| LOC107070860 | ras guanine nucleotide exchange factor R-lik | 0.04253 | 17.71688 | FALSE | FALSE |
| LOC107065558 | uncharacterized protein LOC107065558 | 0.0431 | 17.73383 | FALSE | FALSE |
| LOC107063718 | uncharacterized protein LOC107063718 | 0.04284 | 17.73441 | FALSE | FALSE |
| LOC107073121 | diamine acetyltransferase 2 | 0.0425 | 17.73907 | FALSE | FALSE |
| LOC107065466 | oxygen-dependent coproporphyrinogen-III c | 0.04333 | 17.74032 | FALSE | FALSE |
| LOC107064798 | nuclear pore complex protein Nup153-like | 0.04385 | 17.74394 | FALSE | FALSE |
| LOC107073242 | U11/U12 small nuclear ribonucleoprotein 48 | 0.04435 | 17.75437 | FALSE | FALSE |
| LOC107071216 | aquaporin-like | 0.04365 | 17.76065 | FALSE | FALSE |
| LOC107064171 | L-2-hydroxyglutarate dehydrogenase, mitoc | 0.04319 | 17.78119 | FALSE | FALSE |
| LOC107064097 | afadin | 0.04322 | 17.78948 | FALSE | FALSE |
| LOC107070626 | maternal protein exuperantia | 0.04424 | 17.8 | FALSE | FALSE |
| LOC107069000 | S-formylglutathione hydrolase | 0.04385 | 17.80161 | FALSE | FALSE |
| LOC107070507 | kinesin-like protein KIF14 | 0.04439 | 17.80938 | FALSE | FALSE |

| | | | | | |
|---|---|---|---|---|---|
| LOC107065328 | sphingomyelin phosphodiesterase 4 | 0.04208 | 17.81374 | FALSE | FALSE |
| LOC107065676 | neural/ectodermal development factor IMP | 0.04344 | 17.82038 | FALSE | FALSE |
| LOC107073669 | putative fatty acyl-CoA reductase CG5065 | 0.04161 | 17.85688 | FALSE | FALSE |
| LOC107070331 | uncharacterized protein LOC107070331 | 0.04507 | 17.87077 | FALSE | FALSE |
| LOC107070105 | protein phosphatase inhibitor 2-like | 0.04446 | 17.88324 | FALSE | FALSE |
| LOC107064606 | serine-rich adhesin for platelets | 0.04392 | 17.89492 | FALSE | FALSE |
| LOC107064727 | probable 28S rRNA (cytosine-C(5))-methyltr | 0.04281 | 17.90903 | FALSE | FALSE |
| LOC107064338 | DNA repair protein REV1 | 0.04342 | 17.91511 | FALSE | FALSE |
| LOC107070210 | putative uncharacterized protein DDB_G029 | 0.04451 | 17.93159 | FALSE | FALSE |
| LOC107074357 | MIP18 family protein CG7949 | 0.04349 | 17.93808 | FALSE | FALSE |
| LOC107065954 | uncharacterized protein LOC107065954 | 0.04224 | 17.94068 | FALSE | FALSE |
| LOC107070277 | uncharacterized protein LOC107070277 | 0.04276 | 17.94982 | FALSE | FALSE |
| LOC107072757 | uncharacterized protein LOC107072757 | 0.04223 | 17.95658 | FALSE | FALSE |
| LOC107069153 | neuropeptide Y receptor | 0.04349 | 17.9706 | FALSE | FALSE |
| LOC107069260 | WD repeat-containing protein on Y chromos | 0.04325 | 17.97336 | FALSE | FALSE |
| LOC107072076 | UPF0047 protein YjbQ-like | 0.04247 | 17.98554 | FALSE | FALSE |
| LOC107068894 | snurportin-1 | 0.04298 | 17.98823 | FALSE | FALSE |
| LOC107071770 | UDP-N-acetylglucosamine--peptide N-acety | 0.04567 | 17.9907 | FALSE | FALSE |
| LOC107070414 | mediator of RNA polymerase II transcription | 0.04506 | 18.00362 | FALSE | FALSE |
| LOC107068293 | testis-specific serine/threonine-protein kina | 0.0437 | 18.01485 | FALSE | FALSE |
| LOC107070085 | dynein assembly factor with WDR repeat do | 0.0432 | 18.0301 | FALSE | FALSE |
| LOC107074208 | cytochrome b-c1 complex subunit 9 | 0.04333 | 18.03135 | FALSE | FALSE |
| LOC107070179 | golgin-45 | 0.04218 | 18.03881 | FALSE | FALSE |
| LOC107069705 | RUS1 family protein C16orf58 homolog | 0.04411 | 18.04408 | FALSE | FALSE |
| LOC107063712 | uncharacterized protein LOC107063712 | 0.04338 | 18.05301 | FALSE | FALSE |
| LOC107071908 | serine-rich adhesin for platelets-like | 0.04377 | 18.07023 | FALSE | FALSE |
| LOC107073967 | codanin-1 | 0.04346 | 18.07525 | FALSE | FALSE |
| LOC107072766 | negative elongation factor B-like | 0.04363 | 18.07931 | FALSE | FALSE |
| LOC107069246 | protein NRDE2 homolog | 0.04397 | 18.0798 | FALSE | FALSE |
| LOC107066552 | nuclear factor NF-kappa-B p100 subunit | 0.04339 | 18.07996 | FALSE | FALSE |
| LOC107066657 | peptidyl-prolyl cis-trans isomerase-like 1 | 0.04467 | 18.08537 | FALSE | FALSE |
| LOC107073584 | uncharacterized protein LOC107073584 | 0.04304 | 18.09561 | FALSE | FALSE |
| LOC107067662 | PDZ and LIM domain protein 3 | 0.04265 | 18.10166 | FALSE | FALSE |
| LOC107065430 | uncharacterized protein LOC107065430 | 0.04232 | 18.10771 | FALSE | FALSE |
| LOC107064728 | uncharacterized protein LOC107064728 | 0.04389 | 18.10987 | FALSE | FALSE |
| LOC107066177 | tektin-4 | 0.0418 | 18.11287 | FALSE | FALSE |
| LOC107070134 | E3 ubiquitin-protein ligase AMFR-like | 0.04343 | 18.11952 | FALSE | FALSE |
| LOC107066908 | teneurin-a | 0.04145 | 18.12506 | FALSE | FALSE |
| LOC107074521 | DDB1- and CUL4-associated factor 11 | 0.04328 | 18.12786 | FALSE | FALSE |
| LOC107067998 | putative uncharacterized protein DDB_G028 | 0.0427 | 18.13051 | FALSE | FALSE |
| LOC107072557 | GTP-binding protein Di-Ras2 | 0.04275 | 18.13274 | FALSE | FALSE |
| LOC107071491 | uncharacterized protein LOC107071491 | 0.04312 | 18.13863 | FALSE | FALSE |
| LOC107067133 | FACT complex subunit spt16 | 0.04393 | 18.13887 | FALSE | FALSE |
| LOC107065366 | protein takeout | 0.04304 | 18.15121 | FALSE | FALSE |
| LOC107071691 | putative carbonic anhydrase 3 | 0.04208 | 18.15385 | FALSE | FALSE |
| LOC107074161 | multiple C2 and transmembrane domain-co | 0.04451 | 18.15627 | FALSE | FALSE |
| LOC107070050 | uncharacterized protein LOC107070050 | 0.04496 | 18.17329 | FALSE | FALSE |
| LOC107071483 | THAP domain-containing protein 5-like | 0.04212 | 18.19667 | FALSE | FALSE |
| LOC107070727 | trehalase-like | 0.04414 | 18.21828 | FALSE | FALSE |
| LOC107069518 | putative glucose-6-phosphate 1-epimerase | 0.04236 | 18.2188 | FALSE | FALSE |

| | | | | | |
|---|---|---|---|---|---|
| LOC107064989 | coatomer subunit alpha | 0.04251 | 18.22808 | FALSE | FALSE |
| LOC107072728 | transportin-1 | 0.04335 | 18.23015 | FALSE | FALSE |
| LOC107066290 | carboxypeptidase B-like | 0.04324 | 18.25196 | FALSE | FALSE |
| LOC107071863 | paired box protein Pax-6 | 0.04399 | 18.25336 | FALSE | FALSE |
| LOC107066309 | vitellogenin-6-like | 0.04265 | 18.26644 | FALSE | FALSE |
| LOC107069653 | tRNA (guanine(10)-N2)-methyltransferase h | 0.04268 | 18.27185 | FALSE | FALSE |
| LOC107067005 | protein lin-37 homolog | 0.04269 | 18.29132 | FALSE | FALSE |
| LOC107072425 | peptidyl-tRNA hydrolase 2, mitochondrial-lil | 0.04232 | 18.29289 | FALSE | FALSE |
| LOC107071018 | uncharacterized protein LOC107071018 | 0.04472 | 18.31476 | FALSE | FALSE |
| LOC107071676 | uncharacterized protein LOC107071676 | 0.04407 | 18.31588 | FALSE | FALSE |
| LOC107070518 | peptidyl-tRNA hydrolase 2, mitochondrial-lil | 0.04302 | 18.32346 | FALSE | FALSE |
| Trnag-gcc | Trnag-gcc | 0.04298 | 18.3258 | FALSE | FALSE |
| LOC107074294 | zinc finger CCHC domain-containing protein | 0.04423 | 18.34625 | FALSE | FALSE |
| LOC107070434 | uncharacterized protein LOC107070434 | 0.04189 | 18.34784 | FALSE | FALSE |
| LOC107067945 | cuticle protein 67, isoform B-like | 0.04465 | 18.34814 | FALSE | FALSE |
| LOC107069956 | sulfide:quinone oxidoreductase, mitochond | 0.04265 | 18.36016 | FALSE | FALSE |
| LOC107071002 | uncharacterized protein LOC107071002 | 0.04348 | 18.37372 | FALSE | FALSE |
| LOC107071815 | DNA polymerase delta small subunit-like | 0.04342 | 18.38224 | FALSE | FALSE |
| LOC107074496 | glycine-rich cell wall structural protein-like | 0.04297 | 18.4065 | FALSE | FALSE |
| LOC107067463 | uncharacterized protein LOC107067463 | 0.04331 | 18.42408 | FALSE | FALSE |
| LOC107063949 | rho guanine nucleotide exchange factor 18 | 0.04322 | 18.44227 | FALSE | FALSE |
| LOC107071854 | uncharacterized protein LOC107071854 | 0.04337 | 18.45988 | FALSE | FALSE |
| LOC107064505 | malate dehydrogenase, mitochondrial | 0.04368 | 18.49281 | FALSE | FALSE |
| LOC107063729 | protein virilizer | 0.04285 | 18.49523 | FALSE | FALSE |
| LOC107067583 | pre-mRNA-splicing factor CWC25 homolog | 0.04136 | 18.50807 | FALSE | FALSE |
| LOC107064592 | uncharacterized protein LOC107064592 | 0.04408 | 18.51342 | FALSE | FALSE |
| LOC107065450 | lachesin | 0.04211 | 18.52567 | FALSE | FALSE |
| LOC107066297 | mannose-P-dolichol utilization defect 1 prot | 0.04382 | 18.52798 | FALSE | FALSE |
| LOC107070066 | eukaryotic translation initiation factor 2-alp | 0.04334 | 18.55313 | FALSE | FALSE |
| LOC107068531 | uncharacterized protein LOC107068531 | 0.04466 | 18.57278 | FALSE | FALSE |
| LOC107070125 | dolichyl-diphosphooligosaccharide--protein | 0.0436 | 18.57822 | FALSE | FALSE |
| LOC107067859 | UDP-glucose:glycoprotein glucosyltransfera | 0.04333 | 18.58127 | FALSE | FALSE |
| LOC107074048 | myosin-4-like | 0.04174 | 18.58729 | FALSE | FALSE |
| LOC107072532 | IQ and AAA domain-containing protein 1 | 0.04213 | 18.60247 | FALSE | FALSE |
| LOC107068550 | phosphorylated adapter RNA export proteir | 0.04267 | 18.6075 | FALSE | FALSE |
| LOC107069285 | uncharacterized protein LOC107069285 | 0.04306 | 18.60947 | FALSE | FALSE |
| LOC107072555 | translocation protein SEC63 homolog | 0.04237 | 18.63665 | FALSE | FALSE |
| LOC107073690 | uncharacterized protein LOC107073690 | 0.04326 | 18.64111 | FALSE | FALSE |
| LOC107064648 | ryanodine receptor | 0.04475 | 18.64268 | FALSE | FALSE |
| LOC107073284 | probable ubiquitin carboxyl-terminal hydrol | 0.04203 | 18.64627 | FALSE | FALSE |
| LOC107068931 | bromodomain adjacent to zinc finger domai | 0.04328 | 18.64886 | FALSE | FALSE |
| LOC107070728 | transient-receptor-potential-like protein | 0.04447 | 18.64973 | FALSE | FALSE |
| LOC107074307 | nucleolar and coiled-body phosphoprotein 1 | 0.04197 | 18.65333 | FALSE | FALSE |
| LOC107067465 | venom protease-like | 0.04449 | 18.65364 | FALSE | FALSE |
| LOC107066154 | UDP-N-acetylglucosamine transferase subu | 0.0417 | 18.66335 | FALSE | FALSE |
| LOC107070806 | suppressor of hairless protein | 0.0433 | 18.66992 | FALSE | FALSE |
| LOC107074507 | chitinase domain-containing protein 1 | 0.04309 | 18.67075 | FALSE | FALSE |
| LOC107066895 | helicase SRCAP-like | 0.04284 | 18.67307 | FALSE | FALSE |
| LOC107071417 | enhancer of rudimentary homolog | 0.04294 | 18.69105 | FALSE | FALSE |
| LOC107063645 | uncharacterized protein LOC107063645 | 0.04177 | 18.70739 | FALSE | FALSE |

| | | | | | |
|---|---|---|---|---|---|
| LOC107074497 | dirigent protein 10-like | 0.04541 | 18.71804 | FALSE | FALSE |
| LOC107066890 | TATA box-binding protein-associated factor | 0.04249 | 18.72056 | FALSE | FALSE |
| LOC107066669 | zinc-responsive transcriptional regulator ZA | 0.04429 | 18.72403 | FALSE | FALSE |
| LOC107073635 | uncharacterized protein LOC107073635 | 0.04338 | 18.72406 | FALSE | FALSE |
| LOC107067556 | uncharacterized protein LOC107067556 | 0.0447 | 18.74015 | FALSE | FALSE |
| LOC107064079 | lipoyltransferase 1, mitochondrial | 0.04435 | 18.74556 | FALSE | FALSE |
| LOC107064463 | uncharacterized protein LOC107064463 | 0.04251 | 18.74732 | FALSE | FALSE |
| LOC107066582 | deoxycytidylate deaminase | 0.04239 | 18.76432 | FALSE | FALSE |
| LOC107074618 | zinc finger CCCH domain-containing protein | 0.04344 | 18.76833 | FALSE | FALSE |
| LOC107071312 | protein-cysteine N-palmitoyltransferase HH | 0.04369 | 18.78159 | FALSE | FALSE |
| LOC107073344 | tetraspanin-1 | 0.04433 | 18.78853 | FALSE | FALSE |
| LOC107073351 | DNA-binding protein RFX7 | 0.04288 | 18.79608 | FALSE | FALSE |
| LOC107072630 | telomerase reverse transcriptase-like | 0.04264 | 18.79925 | FALSE | FALSE |
| LOC107068350 | U4/U6 small nuclear ribonucleoprotein Prp3 | 0.04379 | 18.82412 | FALSE | FALSE |
| LOC107071720 | ras-related protein Rab-35 | 0.04299 | 18.82606 | FALSE | FALSE |
| LOC107069109 | vasorin | 0.04262 | 18.86823 | FALSE | FALSE |
| LOC107069382 | uncharacterized protein LOC107069382 | 0.04428 | 18.88661 | FALSE | FALSE |
| LOC107065687 | eukaryotic translation initiation factor 5 | 0.04205 | 18.89292 | FALSE | FALSE |
| LOC107072281 | protein SDA1 homolog | 0.04319 | 18.91199 | FALSE | FALSE |
| LOC107066543 | nitric oxide synthase-interacting protein hor | 0.04148 | 18.91755 | FALSE | FALSE |
| LOC107071599 | chymotrypsin-2 | 0.042 | 18.91882 | FALSE | FALSE |
| LOC107071729 | patatin-like phospholipase domain-containi | 0.04435 | 18.91951 | FALSE | FALSE |
| LOC107073929 | ribonucleases P/MRP protein subunit POP1 | 0.04262 | 18.92989 | FALSE | FALSE |
| LOC107069395 | transmembrane protein 8A-like | 0.04259 | 18.94242 | FALSE | FALSE |
| LOC107073135 | KIF1-binding protein homolog | 0.044 | 18.95003 | FALSE | FALSE |
| LOC107073814 | centrosome-associated zinc finger protein C | 0.0432 | 18.95194 | FALSE | FALSE |
| LOC107072533 | COP9 signalosome complex subunit 3 | 0.04264 | 18.97296 | FALSE | FALSE |
| LOC107067201 | pre-mRNA-splicing factor RBM22 | 0.04287 | 18.97391 | FALSE | FALSE |
| LOC107068646 | protein ST7 homolog | 0.04194 | 18.97767 | FALSE | FALSE |
| LOC107067471 | insulin-like growth factor-binding protein cc | 0.04284 | 18.98793 | FALSE | FALSE |
| LOC107064412 | acyl-CoA-binding protein homolog | 0.04237 | 18.99148 | FALSE | FALSE |
| LOC107064564 | TIP41-like protein | 0.04385 | 19.00065 | FALSE | FALSE |
| LOC107072979 | PAB-dependent poly(A)-specific ribonucleas | 0.04284 | 19.00567 | FALSE | FALSE |
| LOC107064482 | probable Ufm1-specific protease 1 | 0.04341 | 19.01718 | FALSE | FALSE |
| LOC107069914 | E3 ubiquitin-protein ligase UHRF1-like | 0.0419 | 19.02228 | FALSE | FALSE |
| LOC107073354 | thioredoxin-like protein 1 | 0.04274 | 19.02508 | FALSE | FALSE |
| LOC107073718 | putative fatty acyl-CoA reductase CG5065 | 0.04417 | 19.03791 | FALSE | FALSE |
| LOC107073280 | uncharacterized protein LOC107073280 | 0.04287 | 19.04337 | FALSE | FALSE |
| LOC107069357 | uncharacterized protein LOC107069357 | 0.04394 | 19.04826 | FALSE | FALSE |
| LOC107074009 | endocuticle structural glycoprotein SgAbd-1 | 0.04241 | 19.05474 | FALSE | FALSE |
| LOC107070848 | uncharacterized protein LOC107070848 | 0.0428 | 19.06914 | FALSE | FALSE |
| LOC107065239 | probable medium-chain specific acyl-CoA de | 0.04495 | 19.09186 | FALSE | FALSE |
| LOC107071563 | uncharacterized protein LOC107071563 | 0.04267 | 19.106 | FALSE | FALSE |
| LOC107064508 | open rectifier potassium channel protein 1 | 0.04217 | 19.12085 | FALSE | FALSE |
| LOC107073268 | uncharacterized protein LOC107073268 | 0.04308 | 19.12613 | FALSE | FALSE |
| LOC107070296 | zinc transporter ZIP13 homolog | 0.04334 | 19.12842 | FALSE | FALSE |
| LOC107065809 | uncharacterized protein LOC107065809 | 0.04323 | 19.12927 | FALSE | FALSE |
| LOC107070681 | prefoldin subunit 5 | 0.04193 | 19.1305 | FALSE | FALSE |
| LOC107071698 | LIM and SH3 domain protein Lasp | 0.04199 | 19.13511 | FALSE | FALSE |
| LOC107066265 | uncharacterized protein LOC107066265 | 0.0419 | 19.14107 | FALSE | FALSE |

| | | | | | |
|---|---|---|---|---|---|
| LOC107065243 | uncharacterized protein LOC107065243 | 0.04553 | 19.14329 | FALSE | FALSE |
| LOC107071394 | WD repeat-containing protein 55 homolog | 0.04413 | 19.15949 | FALSE | FALSE |
| LOC107071744 | transportin-3 | 0.04353 | 19.1637 | FALSE | FALSE |
| LOC107072563 | kinesin-like protein KIF21A | 0.04288 | 19.16694 | FALSE | FALSE |
| LOC107071593 | putative ferric-chelate reductase 1 homolog | 0.04319 | 19.17904 | FALSE | FALSE |
| LOC107069938 | uncharacterized protein LOC107069938 | 0.0419 | 19.17991 | FALSE | FALSE |
| LOC107068763 | uncharacterized protein LOC107068763 | 0.04371 | 19.1803 | FALSE | FALSE |
| LOC107065511 | DNA excision repair protein ERCC-6 | 0.04207 | 19.20843 | FALSE | FALSE |
| LOC107063986 | calsyntenin-1 | 0.04559 | 19.21716 | FALSE | FALSE |
| LOC107064371 | ester hydrolase C11orf54 homolog | 0.04201 | 19.24023 | FALSE | FALSE |
| LOC107066055 | PHD finger protein 14 | 0.04253 | 19.2434 | FALSE | FALSE |
| LOC107064771 | protein Skeletor, isoforms B/C | 0.04385 | 19.24668 | FALSE | FALSE |
| LOC107067954 | protein TANC2 | 0.04344 | 19.25218 | FALSE | FALSE |
| LOC107069732 | V-type proton ATPase subunit D | 0.04313 | 19.25561 | FALSE | FALSE |
| LOC107066681 | protein DDB_G0268328 | 0.04288 | 19.26828 | FALSE | FALSE |
| LOC107064388 | transmembrane protease serine 9-like | 0.04349 | 19.26865 | FALSE | FALSE |
| LOC107063807 | aspartate aminotransferase, mitochondrial | 0.04472 | 19.27436 | FALSE | FALSE |
| LOC107071650 | uncharacterized protein LOC107071650 | 0.04354 | 19.2799 | FALSE | FALSE |
| LOC107069955 | dolichyldiphosphatase 1-like | 0.04225 | 19.29006 | FALSE | FALSE |
| LOC107068261 | cyclin-K | 0.042 | 19.29769 | FALSE | FALSE |
| LOC107069863 | uncharacterized protein DDB_G0283357 | 0.04219 | 19.31694 | FALSE | FALSE |
| LOC107065711 | uncharacterized protein LOC107065711 | 0.04238 | 19.34097 | FALSE | FALSE |
| LOC107065867 | uncharacterized protein LOC107065867 | 0.04315 | 19.34879 | FALSE | FALSE |
| LOC107066348 | vascular endothelial growth factor receptor | 0.04174 | 19.35997 | FALSE | FALSE |
| LOC107067612 | uncharacterized protein LOC107067612 | 0.04348 | 19.36858 | FALSE | FALSE |
| LOC107072968 | fatty acyl-CoA reductase 1-like | 0.04176 | 19.37479 | FALSE | FALSE |
| LOC107073813 | protein FRG1 homolog | 0.04302 | 19.39451 | FALSE | FALSE |
| LOC107065507 | protein glass-like | 0.04122 | 19.40284 | FALSE | FALSE |
| LOC107065402 | multiple epidermal growth factor-like doma | 0.04379 | 19.40374 | FALSE | FALSE |
| LOC107073481 | protein toll-like | 0.04266 | 19.41642 | FALSE | FALSE |
| LOC107066855 | uncharacterized protein LOC107066855 | 0.04157 | 19.41927 | FALSE | FALSE |
| LOC107073371 | synaptobrevin homolog YKT6 | 0.04283 | 19.43421 | FALSE | FALSE |
| LOC107068558 | uncharacterized protein LOC107068558 | 0.04322 | 19.44542 | FALSE | FALSE |
| LOC107066215 | DNA replication licensing factor Mcm2 | 0.04293 | 19.45887 | FALSE | FALSE |
| LOC107072045 | uncharacterized protein LOC107072045 | 0.04349 | 19.45973 | FALSE | FALSE |
| LOC107072401 | uncharacterized protein LOC107072401 | 0.04171 | 19.46044 | FALSE | FALSE |
| LOC107074299 | uncharacterized protein LOC107074299 | 0.04325 | 19.46314 | FALSE | FALSE |
| LOC107069200 | disintegrin and metalloproteinase domain-c | 0.04277 | 19.46321 | FALSE | FALSE |
| LOC107065308 | 5-hydroxytryptamine receptor 1D-like | 0.04313 | 19.465 | FALSE | FALSE |
| LOC107064013 | protein phosphatase 1H | 0.04244 | 19.47218 | FALSE | FALSE |
| LOC107073050 | uncharacterized protein LOC107073050 | 0.04204 | 19.4811 | FALSE | FALSE |
| LOC107072649 | collagen alpha-1(I) chain-like | 0.04323 | 19.48481 | FALSE | FALSE |
| LOC107066531 | uncharacterized protein LOC107066531 | 0.04174 | 19.48744 | FALSE | FALSE |
| LOC107068178 | CCAAT/enhancer-binding protein gamma | 0.04366 | 19.488 | FALSE | FALSE |
| LOC107070971 | cytoplasmic dynein 2 heavy chain 1 | 0.0423 | 19.49404 | FALSE | FALSE |
| LOC107071978 | uncharacterized protein LOC107071978 | 0.04313 | 19.49626 | FALSE | FALSE |
| LOC107068534 | forkhead box protein N3 | 0.04177 | 19.53337 | FALSE | FALSE |
| LOC107065845 | sodium-coupled monocarboxylate transport | 0.04299 | 19.53476 | FALSE | FALSE |
| LOC107070059 | uncharacterized protein LOC107070059 | 0.04316 | 19.5427 | FALSE | FALSE |
| LOC107063733 | Down syndrome cell adhesion molecule-like | 0.04109 | 19.54679 | FALSE | FALSE |

| | | | | | |
|---|---|---|---|---|---|
| LOC107063761 | GDP-Man:Man(3)GlcNAc(2)-PP-Dol alpha-1, | 0.04304 | 19.55491 | FALSE | FALSE |
| LOC107068258 | T-complex protein 11-like protein 1 | 0.04393 | 19.5603 | FALSE | FALSE |
| LOC107069817 | centrosomal protein of 131 kDa | 0.04191 | 19.56792 | FALSE | FALSE |
| LOC107072700 | methionine-R-sulfoxide reductase B1 | 0.04569 | 19.575 | FALSE | FALSE |
| LOC107072638 | mitochondrial amidoxime-reducing compon | 0.04419 | 19.57835 | FALSE | FALSE |
| LOC107068363 | unc-112-related protein | 0.04351 | 19.57913 | FALSE | FALSE |
| LOC107071445 | cytochrome c oxidase assembly protein CO) | 0.04271 | 19.58414 | FALSE | FALSE |
| LOC107071263 | early endosome antigen 1-like | 0.04172 | 19.58847 | FALSE | FALSE |
| LOC107063868 | peregrin | 0.04265 | 19.59823 | FALSE | FALSE |
| LOC107063623 | uncharacterized protein LOC107063623 | 0.0424 | 19.6083 | FALSE | FALSE |
| LOC107071792 | uncharacterized protein LOC107071792 | 0.04304 | 19.60895 | FALSE | FALSE |
| LOC107064137 | NADH dehydrogenase [ubiquinone | 0.04291 | 19.62988 | FALSE | FALSE |
| LOC107066370 | peroxisomal membrane protein 11C | 0.04131 | 19.63104 | FALSE | FALSE |
| LOC107065455 | uncharacterized protein LOC107065455 | 0.04248 | 19.63313 | FALSE | FALSE |
| LOC107067617 | uncharacterized protein LOC107067617 | 0.04179 | 19.65473 | FALSE | FALSE |
| LOC107071356 | N-acetylglucosaminyl-phosphatidylinositol b | 0.04172 | 19.66057 | FALSE | FALSE |
| LOC107074136 | centrosomal protein of 120 kDa | 0.04286 | 19.66591 | FALSE | FALSE |
| LOC107071820 | DNA repair endonuclease XPF | 0.04186 | 19.67013 | FALSE | FALSE |
| LOC107074120 | uncharacterized protein LOC107074120 | 0.04182 | 19.67021 | FALSE | FALSE |
| LOC107065515 | peroxisomal membrane protein 2 | 0.04287 | 19.68689 | FALSE | FALSE |
| LOC107070884 | ribonucleoside-diphosphate reductase subu | 0.04432 | 19.69176 | FALSE | FALSE |
| LOC107071040 | transcription factor 21 | 0.04212 | 19.69806 | FALSE | FALSE |
| LOC107065506 | macrophage erythroblast attacher | 0.04249 | 19.70164 | FALSE | FALSE |
| LOC107067558 | protein disulfide-isomerase | 0.04266 | 19.72021 | FALSE | FALSE |
| LOC107066339 | uncharacterized protein LOC107066339 | 0.04343 | 19.72134 | FALSE | FALSE |
| LOC107070866 | carbonic anhydrase 1-like | 0.04228 | 19.72503 | FALSE | FALSE |
| LOC107071342 | metabotropic glutamate receptor 2-like | 0.04143 | 19.73466 | FALSE | FALSE |
| LOC107071197 | alpha-N-acetylglucosaminidase | 0.0426 | 19.73713 | FALSE | FALSE |
| LOC107072973 | integrin alpha-PS2-like | 0.04277 | 19.75227 | FALSE | FALSE |
| LOC107073820 | uncharacterized protein LOC107073820 | 0.04177 | 19.76242 | FALSE | FALSE |
| LOC107064247 | probable serine/threonine-protein kinase M | 0.04259 | 19.76547 | FALSE | FALSE |
| LOC107065992 | uncharacterized protein LOC107065992 | 0.04263 | 19.78278 | FALSE | FALSE |
| LOC107064566 | glucose-fructose oxidoreductase domain-co | 0.04242 | 19.80315 | FALSE | FALSE |
| LOC107067671 | uncharacterized protein LOC107067671 | 0.04052 | 19.80771 | FALSE | FALSE |
| LOC107068317 | transcription factor Sox-10-like | 0.04198 | 19.82784 | FALSE | FALSE |
| LOC107071350 | triokinase/FMN cyclase-like | 0.04398 | 19.83261 | FALSE | FALSE |
| LOC107065453 | uncharacterized protein LOC107065453 | 0.04338 | 19.85601 | FALSE | FALSE |
| LOC107065707 | estradiol 17-beta-dehydrogenase 8-like | 0.04315 | 19.86696 | FALSE | FALSE |
| LOC107073902 | DNA-binding protein RFX2 | 0.04258 | 19.88061 | FALSE | FALSE |
| LOC107065486 | uncharacterized protein LOC107065486 | 0.04292 | 19.88934 | FALSE | FALSE |
| LOC107067840 | sodium/potassium-transporting ATPase sub | 0.04222 | 19.90241 | FALSE | FALSE |
| LOC107074173 | probable methyltransferase-like protein 15 | 0.04282 | 19.92829 | FALSE | FALSE |
| LOC107066863 | E3 ubiquitin-protein ligase Topors-like | 0.0423 | 19.93912 | FALSE | FALSE |
| LOC107067008 | piggyBac transposable element-derived pro | 0.04298 | 19.93985 | FALSE | FALSE |
| LOC107069207 | protein Skeletor, isoforms B/C | 0.04268 | 19.94387 | FALSE | FALSE |
| LOC107063893 | dynamin-binding protein-like | 0.0415 | 19.96404 | FALSE | FALSE |
| LOC107069532 | uncharacterized protein KIAA2013 homolog | 0.04236 | 19.97645 | FALSE | FALSE |
| LOC107067939 | uncharacterized protein LOC107067939 | 0.04299 | 19.98259 | FALSE | FALSE |
| LOC107074344 | glycerol-3-phosphate acyltransferase 3 | 0.04218 | 20.00398 | FALSE | FALSE |
| LOC107068687 | telomerase Cajal body protein 1 | 0.04111 | 20.00852 | FALSE | FALSE |

| | | | | | |
|---|---|---|---|---|---|
| LOC107072794 | protein Star | 0.0415 | 20.01048 | FALSE | FALSE |
| LOC107072492 | uncharacterized protein LOC107072492 | 0.04332 | 20.05672 | FALSE | FALSE |
| LOC107074337 | ubiquitin-related modifier 1 | 0.04126 | 20.05709 | FALSE | FALSE |
| LOC107068048 | uncharacterized protein LOC107068048 | 0.04282 | 20.07094 | FALSE | FALSE |
| LOC107069409 | myb-like protein AA | 0.04343 | 20.08307 | FALSE | FALSE |
| LOC107068666 | sarcalumenin | 0.04243 | 20.11161 | FALSE | FALSE |
| LOC107070113 | von Willebrand factor A domain-containing | 0.04352 | 20.11591 | FALSE | FALSE |
| LOC107072711 | carboxypeptidase N subunit 2-like | 0.04349 | 20.13045 | FALSE | FALSE |
| LOC107070629 | uncharacterized protein LOC107070629 | 0.04149 | 20.13053 | FALSE | FALSE |
| LOC107069853 | homeobox protein SIX3 | 0.04297 | 20.13113 | FALSE | FALSE |
| LOC107070647 | sugar phosphate exchanger 2 | 0.04244 | 20.13217 | FALSE | FALSE |
| LOC107073594 | cytochrome P450 4c3-like | 0.04186 | 20.14672 | FALSE | FALSE |
| LOC107074177 | uncharacterized protein LOC107074177 | 0.04251 | 20.14703 | FALSE | FALSE |
| LOC107067910 | odorant receptor 13a-like | 0.04114 | 20.16033 | FALSE | FALSE |
| LOC107065337 | selT-like protein | 0.04352 | 20.17171 | FALSE | FALSE |
| LOC107070614 | fructose-bisphosphate aldolase-like | 0.04201 | 20.17735 | FALSE | FALSE |
| LOC107065651 | lateral signaling target protein 2 homolog | 0.04116 | 20.19144 | FALSE | FALSE |
| LOC107068462 | calponin homology domain-containing prot | 0.04212 | 20.19824 | FALSE | FALSE |
| LOC107066599 | putative ATP-dependent RNA helicase TDRD | 0.04234 | 20.20216 | FALSE | FALSE |
| LOC107067948 | 60S ribosomal protein L26 | 0.043 | 20.21072 | FALSE | FALSE |
| LOC107064184 | tctex1 domain-containing protein 1-like | 0.04334 | 20.21861 | FALSE | FALSE |
| LOC107065958 | putative uncharacterized protein DDB_G02 | 0.04237 | 20.2212 | FALSE | FALSE |
| LOC107067518 | zinc finger protein 2 homolog | 0.04247 | 20.23066 | FALSE | FALSE |
| LOC107066029 | uncharacterized protein LOC107066029 | 0.04261 | 20.23816 | FALSE | FALSE |
| LOC107064274 | zinc finger protein 585A-like | 0.04188 | 20.25758 | FALSE | FALSE |
| LOC107069539 | diuretic hormone 44 | 0.0422 | 20.27259 | FALSE | FALSE |
| LOC107070128 | DNA replication complex GINS protein PSF3 | 0.04392 | 20.28468 | FALSE | FALSE |
| LOC107071565 | uncharacterized protein LOC107071565 | 0.0424 | 20.28716 | FALSE | FALSE |
| LOC107066045 | uncharacterized protein LOC107066045 | 0.04125 | 20.29509 | FALSE | FALSE |
| LOC107070227 | ribonuclease H2 subunit A | 0.04186 | 20.3003 | FALSE | FALSE |
| LOC107073215 | 40S ribosomal protein S8 | 0.04181 | 20.30862 | FALSE | FALSE |
| LOC107074250 | small nuclear ribonucleoprotein F | 0.04214 | 20.31579 | FALSE | FALSE |
| LOC107069483 | ATP synthase mitochondrial F1 complex ass | 0.04297 | 20.31637 | FALSE | FALSE |
| LOC107069797 | queuine tRNA-ribosyltransferase subunit Q | 0.04283 | 20.32294 | FALSE | FALSE |
| LOC107067566 | uncharacterized protein LOC107067566 | 0.04151 | 20.3429 | FALSE | FALSE |
| LOC107066141 | sodium-dependent nutrient amino acid tran | 0.04227 | 20.34505 | FALSE | FALSE |
| LOC107069593 | coagulation factor VII | 0.04287 | 20.36625 | FALSE | FALSE |
| LOC107069596 | uncharacterized protein LOC107069596 | 0.04197 | 20.36892 | FALSE | FALSE |
| LOC107073684 | putative fatty acyl-CoA reductase CG5065 | 0.04221 | 20.36932 | FALSE | FALSE |
| LOC107071913 | neurogenic protein big brain | 0.04162 | 20.36998 | FALSE | FALSE |
| LOC107064862 | UBX domain-containing protein 1 | 0.04279 | 20.37664 | FALSE | FALSE |
| LOC107071374 | piggyBac transposable element-derived pro | 0.04265 | 20.40234 | FALSE | FALSE |
| LOC107074278 | intraflagellar transport protein 88 homolog | 0.04085 | 20.40472 | FALSE | FALSE |
| LOC107063894 | quinone oxidoreductase | 0.03968 | 20.41092 | FALSE | FALSE |
| LOC107067405 | KDEL motif-containing protein 1-like | 0.04335 | 20.41624 | FALSE | FALSE |
| LOC107067936 | cytochrome P450 4C1-like | 0.04285 | 20.42014 | FALSE | FALSE |
| LOC107064011 | uncharacterized protein LOC107064011 | 0.04325 | 20.43388 | FALSE | FALSE |
| LOC107074058 | serine/threonine-protein kinase pakD-like | 0.04262 | 20.44333 | FALSE | FALSE |
| LOC107069038 | uncharacterized protein LOC107069038 | 0.04154 | 20.4453 | FALSE | FALSE |
| LOC107073849 | sodium-independent sulfate anion transpor | 0.04099 | 20.44928 | FALSE | FALSE |

| | | | | | |
|---|---|---|---|---|---|
| LOC107066147 | STE20-like serine/threonine-protein kinase | 0.04234 | 20.52138 | FALSE | FALSE |
| LOC107070108 | J domain-containing protein CG6693 | 0.0409 | 20.52493 | FALSE | FALSE |
| LOC107071705 | 3-hydroxyisobutyrate dehydrogenase, mito | 0.04331 | 20.52927 | FALSE | FALSE |
| LOC107069503 | extended synaptotagmin-2 | 0.04183 | 20.53268 | FALSE | FALSE |
| LOC107064415 | heterogeneous nuclear ribonucleoprotein L | 0.04148 | 20.55028 | FALSE | FALSE |
| LOC107074207 | leukocyte receptor cluster member 1 | 0.04206 | 20.55378 | FALSE | FALSE |
| LOC107073539 | target of rapamycin complex subunit lst8 | 0.04179 | 20.57205 | FALSE | FALSE |
| LOC107070801 | protein chibby homolog 1-like | 0.04327 | 20.61185 | FALSE | FALSE |
| LOC107064747 | 39S ribosomal protein L2, mitochondrial | 0.0423 | 20.61968 | FALSE | FALSE |
| LOC107068896 | acidic amino acid decarboxylase GADL1 | 0.04209 | 20.62504 | FALSE | FALSE |
| LOC107069631 | vacuolar protein sorting-associated protein | 0.04191 | 20.63142 | FALSE | FALSE |
| LOC107068464 | uncharacterized protein LOC107068464 | 0.04383 | 20.63454 | FALSE | FALSE |
| LOC107067010 | protein FAM117B-like | 0.0418 | 20.64381 | FALSE | FALSE |
| LOC107066686 | zinc finger protein 271-like | 0.04062 | 20.65544 | FALSE | FALSE |
| LOC107071586 | pyruvate dehydrogenase protein X compon | 0.04187 | 20.66671 | FALSE | FALSE |
| LOC107064405 | zinc finger protein 836-like | 0.04188 | 20.67376 | FALSE | FALSE |
| LOC107068934 | 2-hydroxyacylsphingosine 1-beta-galactosyl | 0.04059 | 20.68362 | FALSE | FALSE |
| LOC107064532 | ubiquitin carboxyl-terminal hydrolase 7 | 0.04236 | 20.68455 | FALSE | FALSE |
| LOC107070543 | protein Smaug homolog 1 | 0.04371 | 20.69906 | FALSE | FALSE |
| LOC107064842 | mitochondrial cardiolipin hydrolase | 0.0421 | 20.70628 | FALSE | FALSE |
| LOC107065339 | uncharacterized protein LOC107065339 | 0.04029 | 20.70951 | FALSE | FALSE |
| LOC107065397 | uncharacterized protein LOC107065397 | 0.04057 | 20.7104 | FALSE | FALSE |
| LOC107064453 | protein amalgam-like | 0.04107 | 20.71658 | FALSE | FALSE |
| LOC107070726 | probable aconitate hydratase, mitochondria | 0.04259 | 20.71725 | FALSE | FALSE |
| LOC107070112 | centaurin-gamma-1A | 0.0424 | 20.73646 | FALSE | FALSE |
| LOC107064980 | endothelial differentiation-related factor 1 | 0.04359 | 20.73684 | FALSE | FALSE |
| LOC107071075 | 1-acyl-sn-glycerol-3-phosphate acyltransfera | 0.04278 | 20.74884 | FALSE | FALSE |
| LOC107063819 | L-asparaginase 1 | 0.043 | 20.75645 | FALSE | FALSE |
| LOC107072090 | ejaculatory bulb-specific protein 3-like | 0.04225 | 20.76408 | FALSE | FALSE |
| LOC107064448 | tetraspanin-7-like | 0.04113 | 20.76753 | FALSE | FALSE |
| LOC107072908 | LIM domain only protein 3-like | 0.04184 | 20.7698 | FALSE | FALSE |
| LOC107074264 | mitotic spindle assembly checkpoint protein | 0.04052 | 20.77225 | FALSE | FALSE |
| LOC107072945 | autophagy protein 5 | 0.04249 | 20.77525 | FALSE | FALSE |
| LOC107074224 | cyclin-C | 0.04278 | 20.77875 | FALSE | FALSE |
| LOC107074564 | uncharacterized protein LOC107074564 | 0.04194 | 20.79242 | FALSE | FALSE |
| LOC107073915 | adapter molecule Crk | 0.0414 | 20.80178 | FALSE | FALSE |
| LOC107074319 | GDNF-inducible zinc finger protein 1-like | 0.04103 | 20.80843 | FALSE | FALSE |
| LOC107072218 | uncharacterized protein LOC107072218 | 0.04255 | 20.84451 | FALSE | FALSE |
| LOC107064047 | uncharacterized protein LOC107064047 | 0.04254 | 20.84818 | FALSE | FALSE |
| LOC107064749 | uncharacterized protein LOC107064749 | 0.04296 | 20.85314 | FALSE | FALSE |
| LOC107069254 | probable RNA polymerase II nuclear localiza | 0.04086 | 20.85394 | FALSE | FALSE |
| LOC107068058 | chloride channel protein 2 | 0.04356 | 20.87859 | FALSE | FALSE |
| LOC107064683 | uncharacterized protein LOC107064683 | 0.04177 | 20.88129 | FALSE | FALSE |
| LOC107069875 | COP9 signalosome complex subunit 5 | 0.04119 | 20.88302 | FALSE | FALSE |
| LOC107072480 | heme transporter hrg1-A-like | 0.04206 | 20.89147 | FALSE | FALSE |
| LOC107063629 | glutamate receptor ionotropic, kainate 2-lik | 0.04307 | 20.89461 | FALSE | FALSE |
| LOC107066853 | ATP-dependent RNA helicase abstrakt | 0.04308 | 20.90429 | FALSE | FALSE |
| LOC107070082 | uncharacterized protein LOC107070082 | 0.04146 | 20.92874 | FALSE | FALSE |
| LOC107072676 | dynactin subunit 6 | 0.04243 | 20.93823 | FALSE | FALSE |
| LOC107069150 | uncharacterized protein LOC107069150 | 0.04242 | 20.94853 | FALSE | FALSE |

| | | | | | |
|---|---|---|---|---|---|
| LOC107064111 | uncharacterized protein LOC107064111 | 0.04055 | 20.94931 | FALSE | FALSE |
| LOC107072619 | peptidyl-prolyl cis-trans isomerase NIMA-int | 0.04257 | 20.9667 | FALSE | FALSE |
| LOC107073974 | Bardet-Biedl syndrome 5 protein homolog | 0.04312 | 20.96685 | FALSE | FALSE |
| LOC107066126 | GDNF-inducible zinc finger protein 1 | 0.04157 | 20.97735 | FALSE | FALSE |
| LOC107073891 | myelin regulatory factor-like protein | 0.04074 | 20.98071 | FALSE | FALSE |
| LOC107066720 | cyclin-dependent kinase 12 | 0.04303 | 21.0189 | FALSE | FALSE |
| LOC107074383 | heparin sulfate O-sulfotransferase | 0.04369 | 21.03832 | FALSE | FALSE |
| LOC107072141 | transcription factor MafG | 0.0419 | 21.03886 | FALSE | FALSE |
| LOC107067675 | myb-like protein Q | 0.04243 | 21.04398 | FALSE | FALSE |
| LOC107071130 | centrosomal protein of 152 kDa-like | 0.04281 | 21.04497 | FALSE | FALSE |
| LOC107073926 | 60S ribosomal export protein NMD3 | 0.04213 | 21.06825 | FALSE | FALSE |
| LOC107064709 | RNA-binding protein 5-like | 0.0424 | 21.06902 | FALSE | FALSE |
| LOC107066726 | prohormone-4 | 0.04306 | 21.0823 | FALSE | FALSE |
| LOC107068483 | actin-related protein 2/3 complex subunit 3 | 0.04158 | 21.0881 | FALSE | FALSE |
| LOC107065619 | 10 kDa heat shock protein, mitochondrial | 0.04033 | 21.09432 | FALSE | FALSE |
| LOC107069862 | GTP-binding protein 128up | 0.04076 | 21.10002 | FALSE | FALSE |
| LOC107071060 | U3 small nucleolar RNA-associated protein 1 | 0.04169 | 21.12191 | FALSE | FALSE |
| LOC107067078 | uncharacterized protein LOC107067078 | 0.04246 | 21.12327 | FALSE | FALSE |
| LOC107067624 | probable tubulin polyglutamylase TTLL2 | 0.04225 | 21.12719 | FALSE | FALSE |
| LOC107069450 | phosphatidylinositol 4-phosphate 5-kinase t | 0.03999 | 21.15029 | FALSE | FALSE |
| LOC107074695 | transmembrane protein 26 | 0.04137 | 21.15119 | FALSE | FALSE |
| LOC107072255 | uncharacterized protein LOC107072255 | 0.04129 | 21.15564 | FALSE | FALSE |
| LOC107071387 | uncharacterized protein LOC107071387 | 0.0414 | 21.16521 | FALSE | FALSE |
| LOC107065367 | MFS-type transporter SLC18B1-like | 0.0409 | 21.18101 | FALSE | FALSE |
| LOC107072124 | protein hu-li tai shao | 0.04108 | 21.18253 | FALSE | FALSE |
| LOC107065408 | uncharacterized protein LOC107065408 | 0.04252 | 21.18362 | FALSE | FALSE |
| LOC107073244 | 1-phosphatidylinositol 4,5-bisphosphate pho | 0.04204 | 21.19248 | FALSE | FALSE |
| LOC107069493 | rotatin | 0.04188 | 21.19668 | FALSE | FALSE |
| LOC107074424 | activating signal cointegrator 1 complex sub | 0.04173 | 21.20309 | FALSE | FALSE |
| LOC107070395 | proton-coupled amino acid transporter 1 | 0.04318 | 21.20466 | FALSE | FALSE |
| LOC107068356 | probable serine hydrolase | 0.04158 | 21.20511 | FALSE | FALSE |
| LOC107070524 | copper transport protein ATOX1 | 0.04206 | 21.215 | FALSE | FALSE |
| LOC107071191 | hormone receptor 4-like | 0.04209 | 21.21703 | FALSE | FALSE |
| LOC107072379 | uncharacterized protein LOC107072379 | 0.04278 | 21.2206 | FALSE | FALSE |
| LOC107069889 | RING finger protein 207-like | 0.04178 | 21.22677 | FALSE | FALSE |
| LOC107068171 | methionine aminopeptidase 1D, mitochond | 0.0404 | 21.23165 | FALSE | FALSE |
| LOC107070431 | mitochondrial intermembrane space import | 0.04175 | 21.23731 | FALSE | FALSE |
| LOC107064215 | zinc finger protein 883-like | 0.0426 | 21.25762 | FALSE | FALSE |
| LOC107072404 | ubiquitin carboxyl-terminal hydrolase 33 | 0.04141 | 21.2807 | FALSE | FALSE |
| LOC107071571 | trichohyalin-like | 0.04251 | 21.30237 | FALSE | FALSE |
| LOC107073826 | glyoxalase domain-containing protein 4 | 0.04144 | 21.31304 | FALSE | FALSE |
| LOC107064647 | 5'-nucleotidase domain-containing protein 1 | 0.04176 | 21.31326 | FALSE | FALSE |
| LOC107068853 | atrial natriuretic peptide-converting enzyme | 0.04088 | 21.32049 | FALSE | FALSE |
| LOC107070102 | uncharacterized protein LOC107070102 | 0.04114 | 21.33236 | FALSE | FALSE |
| LOC107064433 | zinc finger Ran-binding domain-containing p | 0.042 | 21.35918 | FALSE | FALSE |
| LOC107072514 | pre-mRNA-splicing factor CWC22 homolog | 0.04131 | 21.36355 | FALSE | FALSE |
| LOC107066298 | uncharacterized protein LOC107066298 | 0.04057 | 21.36378 | FALSE | FALSE |
| LOC107074277 | uncharacterized protein LOC107074277 | 0.0411 | 21.36403 | FALSE | FALSE |
| LOC107074559 | uncharacterized protein LOC107074559 | 0.04108 | 21.39673 | FALSE | FALSE |
| LOC107067063 | E3 ubiquitin-protein ligase RAD18-like | 0.04335 | 21.40448 | FALSE | FALSE |

| | | | | | |
|---|---|---|---|---|---|
| LOC107070432 | ADP-ribosylation factor-like protein 2 | 0.04304 | 21.4081 | FALSE | FALSE |
| LOC107065881 | THO complex subunit 2 | 0.04154 | 21.41117 | FALSE | FALSE |
| LOC107071091 | serine/threonine-protein kinase STK11 | 0.04166 | 21.4167 | FALSE | FALSE |
| LOC107073988 | protein RRP5 homolog | 0.04114 | 21.44449 | FALSE | FALSE |
| LOC107072352 | L-threonine 3-dehydrogenase, mitochondria | 0.04137 | 21.44818 | FALSE | FALSE |
| LOC107065446 | sialin | 0.04072 | 21.45342 | FALSE | FALSE |
| LOC107072346 | uncharacterized protein LOC107072346 | 0.04118 | 21.46914 | FALSE | FALSE |
| LOC107073921 | zinc finger protein 800 | 0.04108 | 21.47649 | FALSE | FALSE |
| LOC107063762 | apoptosis regulatory protein Siva-like | 0.04214 | 21.47972 | FALSE | FALSE |
| LOC107066770 | peptidoglycan-recognition protein SC2-like | 0.0424 | 21.4881 | FALSE | FALSE |
| LOC107067467 | uncharacterized protein LOC107067467 | 0.04181 | 21.50146 | FALSE | FALSE |
| LOC107068205 | structural maintenance of chromosomes pr | 0.04193 | 21.50161 | FALSE | FALSE |
| LOC107064375 | UDP-glucose 4-epimerase-like | 0.04208 | 21.50704 | FALSE | FALSE |
| LOC107072740 | threonine--tRNA ligase, cytoplasmic | 0.04093 | 21.50869 | FALSE | FALSE |
| LOC107069813 | homeobox protein araucan | 0.04013 | 21.51075 | FALSE | FALSE |
| LOC107069609 | probable ATP-dependent RNA helicase DDX | 0.04194 | 21.52363 | FALSE | FALSE |
| LOC107068354 | gamma-1-syntrophin | 0.04191 | 21.52964 | FALSE | FALSE |
| LOC107066950 | cytochrome P450 4C1-like | 0.04043 | 21.53114 | FALSE | FALSE |
| LOC107070731 | N-alpha-acetyltransferase 60 | 0.04194 | 21.54287 | FALSE | FALSE |
| LOC107073741 | LOW QUALITY PROTEIN: uncharacterized pr | 0.04083 | 21.54703 | FALSE | FALSE |
| LOC107064974 | beclin 1-associated autophagy-related key r | 0.04259 | 21.54948 | FALSE | FALSE |
| LOC107070441 | uncharacterized protein LOC107070441 | 0.0412 | 21.55755 | FALSE | FALSE |
| LOC107068759 | uncharacterized protein LOC107068759 | 0.0413 | 21.56608 | FALSE | FALSE |
| LOC107066759 | uncharacterized protein LOC107066759 | 0.04137 | 21.57098 | FALSE | FALSE |
| LOC107068517 | uncharacterized protein LOC107068517 | 0.04163 | 21.57122 | FALSE | FALSE |
| LOC107064879 | UPF0547 protein C16orf87 homolog | 0.04171 | 21.57123 | FALSE | FALSE |
| LOC107065526 | WD repeat-containing and planar cell polari | 0.04085 | 21.57143 | FALSE | FALSE |
| LOC107073677 | uncharacterized protein LOC107073677 | 0.04315 | 21.5718 | FALSE | FALSE |
| LOC107071839 | AP-2 complex subunit alpha | 0.04247 | 21.59703 | FALSE | FALSE |
| LOC107068915 | zinc finger CCCH domain-containing protein | 0.04084 | 21.60674 | FALSE | FALSE |
| LOC107068965 | GTPase-activating Rap/Ran-GAP domain-like | 0.04159 | 21.60725 | FALSE | FALSE |
| LOC107063818 | protein-glutamate O-methyltransferase-like | 0.04231 | 21.62429 | FALSE | FALSE |
| LOC107066025 | probable palmitoyltransferase ZDHHC24 | 0.04056 | 21.63971 | FALSE | FALSE |
| LOC107072177 | uncharacterized protein LOC107072177 | 0.04092 | 21.64572 | FALSE | FALSE |
| LOC107070381 | serine/threonine-protein kinase PLK4 | 0.04051 | 21.64586 | FALSE | FALSE |
| LOC107072066 | F-box/WD repeat-containing protein 7 | 0.0413 | 21.64621 | FALSE | FALSE |
| LOC107064608 | protein RER1 | 0.04051 | 21.64711 | FALSE | FALSE |
| LOC107067681 | uncharacterized protein LOC107067681 | 0.04244 | 21.6538 | FALSE | FALSE |
| LOC107073952 | hemocyte protein-glutamine gamma-glutan | 0.04034 | 21.65424 | FALSE | FALSE |
| LOC107072503 | uncharacterized protein LOC107072503 | 0.04077 | 21.664 | FALSE | FALSE |
| LOC107067884 | protein FAM49B | 0.04144 | 21.66574 | FALSE | FALSE |
| LOC107072640 | uncharacterized protein LOC107072640 | 0.04183 | 21.68082 | FALSE | FALSE |
| LOC107067326 | anaphase-promoting complex subunit 5 | 0.04264 | 21.68085 | FALSE | FALSE |
| LOC107071818 | cytochrome b-c1 complex subunit Rieske, m | 0.04261 | 21.69126 | FALSE | FALSE |
| LOC107074167 | putative deoxyribonuclease TATDN1 | 0.04224 | 21.69185 | FALSE | FALSE |
| LOC107064580 | non-homologous end-joining factor 1-like | 0.04253 | 21.70648 | FALSE | FALSE |
| LOC107069277 | protein big brother-like | 0.04111 | 21.70933 | FALSE | FALSE |
| LOC107069514 | uncharacterized protein LOC107069514 | 0.04097 | 21.71174 | FALSE | FALSE |
| LOC107072467 | tudor domain-containing protein 7B-like | 0.041 | 21.71869 | FALSE | FALSE |
| LOC107074610 | brefeldin A-inhibited guanine nucleotide-ex | 0.0418 | 21.72176 | FALSE | FALSE |

| | | | | | |
|---|---|---|---|---|---|
| LOC107073072 | short neuropeptide F-like | 0.04235 | 21.74537 | FALSE | FALSE |
| LOC107073840 | sodium-independent sulfate anion transpor | 0.04129 | 21.75024 | FALSE | FALSE |
| LOC107065313 | targeting protein for Xklp2 homolog | 0.04128 | 21.75305 | FALSE | FALSE |
| LOC107074059 | homeobox protein HMX3-B-like | 0.04258 | 21.75802 | FALSE | FALSE |
| LOC107069865 | porphobilinogen deaminase | 0.04035 | 21.78919 | FALSE | FALSE |
| LOC107064061 | glycerol-3-phosphate dehydrogenase [NAD( | 0.04047 | 21.80579 | FALSE | FALSE |
| LOC107067067 | uncharacterized protein LOC107067067 | 0.0409 | 21.82591 | FALSE | FALSE |
| LOC107064861 | uncharacterized protein LOC107064861 | 0.04078 | 21.83641 | FALSE | FALSE |
| LOC107069827 | trafficking protein particle complex subunit | 0.04025 | 21.84923 | FALSE | FALSE |
| LOC107072136 | uncharacterized protein LOC107072136 | 0.04233 | 21.85199 | FALSE | FALSE |
| LOC107067018 | solute carrier organic anion transporter fam | 0.04057 | 21.8741 | FALSE | FALSE |
| LOC107073996 | probable small nuclear ribonucleoprotein Sr | 0.03897 | 21.87436 | FALSE | FALSE |
| LOC107064043 | amine oxidase [flavin-containing | 0.04347 | 21.8796 | FALSE | FALSE |
| LOC107063852 | putative uncharacterized protein DDB_G02 | 0.0411 | 21.89139 | FALSE | FALSE |
| LOC107072430 | uncharacterized protein LOC107072430 | 0.04278 | 21.89829 | FALSE | FALSE |
| LOC107063890 | UDP-galactose translocator | 0.04068 | 21.91594 | FALSE | FALSE |
| LOC107071132 | brain tumor protein-like | 0.03994 | 21.91779 | FALSE | FALSE |
| LOC107073194 | ATPase WRNIP1-like | 0.04111 | 21.92175 | FALSE | FALSE |
| LOC107069151 | troponin T, skeletal muscle-like | 0.04155 | 21.93108 | FALSE | FALSE |
| LOC107065126 | serine-arginine protein 55 | 0.04131 | 21.94283 | FALSE | FALSE |
| LOC107068457 | glucose dehydrogenase [FAD, quinone | 0.04249 | 21.94313 | FALSE | FALSE |
| LOC107065421 | required for meiotic nuclear division proteir | 0.04125 | 21.95718 | FALSE | FALSE |
| LOC107073342 | putative nuclease HARBI1 | 0.03877 | 21.96232 | FALSE | FALSE |
| LOC107071656 | lymphocyte antigen 6D | 0.04188 | 21.97037 | FALSE | FALSE |
| LOC107065274 | probable ATP-dependent RNA helicase DDX | 0.04299 | 21.97368 | FALSE | FALSE |
| LOC107066192 | uncharacterized protein LOC107066192 | 0.04129 | 21.99455 | FALSE | FALSE |
| LOC107070016 | uncharacterized protein LOC107070016 | 0.04085 | 22.00476 | FALSE | FALSE |
| LOC107067916 | odorant receptor 22b-like | 0.04062 | 22.00795 | FALSE | FALSE |
| LOC107073863 | uncharacterized protein LOC107073863 | 0.04126 | 22.01595 | FALSE | FALSE |
| LOC107073327 | surfeit locus protein 6 homolog | 0.04167 | 22.01714 | FALSE | FALSE |
| LOC107065165 | poly(A)-specific ribonuclease PARN-like dor | 0.0413 | 22.0199 | FALSE | FALSE |
| LOC107070765 | polyphosphoinositide phosphatase | 0.04217 | 22.02741 | FALSE | FALSE |
| LOC107064330 | transmembrane and TPR repeat-containing | 0.04097 | 22.03476 | FALSE | FALSE |
| LOC107071302 | uncharacterized protein LOC107071302 | 0.04425 | 22.03564 | FALSE | FALSE |
| LOC107073868 | MTSS1-like protein | 0.04142 | 22.04923 | FALSE | FALSE |
| LOC107072561 | uncharacterized protein LOC107072561 | 0.04212 | 22.06246 | FALSE | FALSE |
| LOC107074090 | uncharacterized protein LOC107074090 | 0.04271 | 22.08657 | FALSE | FALSE |
| LOC107071960 | protein ERGIC-53 | 0.04112 | 22.09174 | FALSE | FALSE |
| LOC107069974 | zinc transporter foi-like | 0.0421 | 22.10025 | FALSE | FALSE |
| LOC107074006 | sorting nexin lst-4 | 0.04091 | 22.10568 | FALSE | FALSE |
| LOC107065878 | high mobility group protein DSP1-like | 0.0415 | 22.1086 | FALSE | FALSE |
| LOC107065481 | uncharacterized protein LOC107065481 | 0.04132 | 22.11939 | FALSE | FALSE |
| LOC107067868 | 28S ribosomal protein S18a, mitochondrial | 0.04012 | 22.1232 | FALSE | FALSE |
| LOC107068831 | proteasome activator complex subunit 3 | 0.04286 | 22.14684 | FALSE | FALSE |
| LOC107072345 | helicase sen1 | 0.04057 | 22.1697 | FALSE | FALSE |
| LOC107070513 | dihydropyrimidine dehydrogenase [NADP(+ | 0.04216 | 22.17004 | FALSE | FALSE |
| LOC107066249 | probable tubulin polyglutamylase TTLL1 | 0.04231 | 22.17083 | FALSE | FALSE |
| LOC107065214 | eukaryotic translation initiation factor 4H | 0.04018 | 22.19352 | FALSE | FALSE |
| LOC107064610 | oxidative stress-induced growth inhibitor 2- | 0.0411 | 22.20751 | FALSE | FALSE |
| LOC107069614 | 60S acidic ribosomal protein P2 | 0.04173 | 22.21837 | FALSE | FALSE |

| | | | | | |
|---|---|---|---|---|---|
| LOC107066069 | uncharacterized protein LOC107066069 | 0.0406 | 22.21972 | FALSE | FALSE |
| LOC107067726 | heat shock protein 83-like | 0.04215 | 22.22054 | FALSE | FALSE |
| LOC107065686 | uncharacterized protein LOC107065686 | 0.04211 | 22.22643 | FALSE | FALSE |
| LOC107067805 | wolframin | 0.04044 | 22.25251 | FALSE | FALSE |
| LOC107073149 | sodium/potassium-transporting ATPase sub | 0.04085 | 22.25747 | FALSE | FALSE |
| LOC107067414 | protein SON-like | 0.04129 | 22.25981 | FALSE | FALSE |
| LOC107068326 | uncharacterized protein LOC107068326 | 0.04282 | 22.26306 | FALSE | FALSE |
| LOC107073439 | phospholipase D C-like | 0.04193 | 22.27034 | FALSE | FALSE |
| LOC107063847 | guanine nucleotide exchange factor subunit | 0.04082 | 22.2825 | FALSE | FALSE |
| LOC107071680 | TWiK family of potassium channels protein | 0.0426 | 22.29934 | FALSE | FALSE |
| LOC107064758 | neurobeachin | 0.04169 | 22.30007 | FALSE | FALSE |
| LOC107067240 | putative GPI-anchor transamidase | 0.0421 | 22.30898 | FALSE | FALSE |
| LOC107066424 | uncharacterized protein LOC107066424 | 0.04122 | 22.31003 | FALSE | FALSE |
| LOC107073888 | bromodomain adjacent to zinc finger domai | 0.04141 | 22.31191 | FALSE | FALSE |
| LOC107065813 | endocuticle structural protein SgAbd-6-like | 0.04082 | 22.32203 | FALSE | FALSE |
| LOC107066588 | superoxide dismutase [Mn | 0.04256 | 22.32245 | FALSE | FALSE |
| LOC107068429 | neprilysin-2-like | 0.04107 | 22.33813 | FALSE | FALSE |
| LOC107071578 | homeobox protein 5-like | 0.04104 | 22.33861 | FALSE | FALSE |
| LOC107074565 | activated RNA polymerase II transcriptional | 0.04059 | 22.33915 | FALSE | FALSE |
| LOC107065577 | uncharacterized protein LOC107065577 | 0.04177 | 22.34009 | FALSE | FALSE |
| LOC107065222 | WD repeat-containing protein 26 | 0.04173 | 22.34324 | FALSE | FALSE |
| LOC107071749 | bolA-like protein 3 | 0.04108 | 22.35078 | FALSE | FALSE |
| LOC107069495 | patronin | 0.04106 | 22.36019 | FALSE | FALSE |
| LOC107069945 | low-density lipoprotein receptor-related pr | 0.04064 | 22.36498 | FALSE | FALSE |
| LOC107069216 | probable cytochrome P450 301a1, mitocho | 0.04226 | 22.38045 | FALSE | FALSE |
| LOC107064313 | E3 ubiquitin-protein ligase HECTD1 | 0.04156 | 22.38073 | FALSE | FALSE |
| LOC107065259 | protein msta, isoform B-like | 0.04156 | 22.38209 | FALSE | FALSE |
| LOC107065636 | bromodomain-containing protein DDB_G02 | 0.0408 | 22.4007 | FALSE | FALSE |
| LOC107067442 | uncharacterized protein LOC107067442 | 0.04233 | 22.43813 | FALSE | FALSE |
| LOC107068513 | electron transfer flavoprotein subunit beta | 0.04135 | 22.44632 | FALSE | FALSE |
| LOC107070596 | DNA methyltransferase 1-associated protei | 0.04252 | 22.4493 | FALSE | FALSE |
| LOC107064272 | PTB domain-containing engulfment adapter | 0.04046 | 22.45063 | FALSE | FALSE |
| LOC107065025 | cysteine-rich with EGF-like domain protein | 0.04071 | 22.50505 | FALSE | FALSE |
| LOC107069826 | G kinase-anchoring protein 1-like | 0.04122 | 22.51364 | FALSE | FALSE |
| LOC107068428 | adenylosuccinate lyase | 0.04118 | 22.51416 | FALSE | FALSE |
| LOC107072873 | RWD domain-containing protein 2A | 0.04176 | 22.51663 | FALSE | FALSE |
| LOC107070845 | dual specificity mitogen-activated protein ki | 0.04029 | 22.53244 | FALSE | FALSE |
| LOC107065862 | uncharacterized protein LOC107065862 | 0.04184 | 22.54097 | FALSE | FALSE |
| LOC107066279 | uncharacterized protein LOC107066279 | 0.04247 | 22.54842 | FALSE | FALSE |
| LOC107071991 | zinc finger protein 423 | 0.04194 | 22.55855 | FALSE | FALSE |
| LOC107074172 | C-1-tetrahydrofolate synthase, cytoplasmic | 0.04095 | 22.566 | FALSE | FALSE |
| LOC107072018 | uncharacterized protein LOC107072018 | 0.04147 | 22.59243 | FALSE | FALSE |
| LOC107067890 | zinc finger protein MSN2-like | 0.0416 | 22.59275 | FALSE | FALSE |
| LOC107071073 | IST1 homolog | 0.04238 | 22.59939 | FALSE | FALSE |
| LOC107064133 | signal-induced proliferation-associated 1-lik | 0.0418 | 22.60704 | FALSE | FALSE |
| LOC107067864 | 26S proteasome complex subunit DSS1 | 0.04071 | 22.62537 | FALSE | FALSE |
| LOC107071385 | RNA-binding protein pno1-like | 0.04007 | 22.62553 | FALSE | FALSE |
| LOC107073015 | cyclin-dependent kinases regulatory subuni | 0.03946 | 22.62864 | FALSE | FALSE |
| LOC107073823 | protein lethal(2)essential for life-like | 0.04129 | 22.63737 | FALSE | FALSE |
| LOC107070374 | glutamate-gated chloride channel | 0.0412 | 22.64208 | FALSE | FALSE |

| | | | | | |
|---|---|---|---|---|---|
| LOC107066129 | H/ACA ribonucleoprotein complex subunit 2 | 0.04087 | 22.64371 | FALSE | FALSE |
| LOC107072013 | histone-lysine N-methyltransferase trithora: | 0.0421 | 22.65588 | FALSE | FALSE |
| LOC107074084 | adenosylhomocysteinase 2-like | 0.04033 | 22.66025 | FALSE | FALSE |
| LOC107064104 | cysteine and histidine-rich protein 1 homolc | 0.04066 | 22.66172 | FALSE | FALSE |
| LOC107071310 | small proline-rich protein 2B-like | 0.04089 | 22.67054 | FALSE | FALSE |
| LOC107065967 | DNA replication complex GINS protein SLD5 | 0.04036 | 22.67527 | FALSE | FALSE |
| LOC107064081 | phospholipase ABHD3 | 0.0409 | 22.68062 | FALSE | FALSE |
| LOC107069931 | eukaryotic translation initiation factor 3 sub | 0.04049 | 22.68676 | FALSE | FALSE |
| LOC107071230 | fork head domain transcription factor slp1-l | 0.04068 | 22.69523 | FALSE | FALSE |
| LOC107067295 | ATP-binding cassette sub-family A member | 0.04168 | 22.69838 | FALSE | FALSE |
| LOC107065202 | ATP synthase subunit g, mitochondrial | 0.03993 | 22.72651 | FALSE | FALSE |
| LOC107069864 | dr1-associated corepressor homolog | 0.04145 | 22.72719 | FALSE | FALSE |
| LOC107071499 | uncharacterized protein LOC107071499 | 0.04249 | 22.7276 | FALSE | FALSE |
| LOC107065350 | basic proline-rich protein-like | 0.04128 | 22.73508 | FALSE | FALSE |
| LOC107074209 | uncharacterized protein LOC107074209 | 0.04138 | 22.73702 | FALSE | FALSE |
| LOC107065247 | serine/threonine-protein kinase Warts | 0.03992 | 22.76261 | FALSE | FALSE |
| LOC107074585 | uncharacterized protein LOC107074585 | 0.04234 | 22.77973 | FALSE | FALSE |
| LOC107065772 | 26S proteasome non-ATPase regulatory sub | 0.04165 | 22.78939 | FALSE | FALSE |
| LOC107072718 | uncharacterized protein LOC107072718 | 0.04105 | 22.79031 | FALSE | FALSE |
| LOC107070964 | conserved oligomeric Golgi complex subunit | 0.04001 | 22.80406 | FALSE | FALSE |
| LOC107069629 | serine/threonine-protein kinase TAO1 | 0.04065 | 22.82317 | FALSE | FALSE |
| LOC107065751 | uncharacterized protein LOC107065751 | 0.04177 | 22.82566 | FALSE | FALSE |
| LOC107066366 | uncharacterized protein LOC107066366 | 0.04043 | 22.82617 | FALSE | FALSE |
| LOC107068999 | target of rapamycin complex 2 subunit MAF | 0.03983 | 22.82661 | FALSE | FALSE |
| LOC107070478 | protein LZIC-like | 0.03993 | 22.83638 | FALSE | FALSE |
| LOC107070019 | uncharacterized protein LOC107070019 | 0.04099 | 22.83663 | FALSE | FALSE |
| LOC107066355 | protein HIRA homolog | 0.04216 | 22.85197 | FALSE | FALSE |
| LOC107064863 | ribonuclease Oy | 0.04078 | 22.85457 | FALSE | FALSE |
| LOC107071229 | inhibitor of growth protein 1 | 0.04107 | 22.85714 | FALSE | FALSE |
| LOC107064361 | cone-rod homeobox protein-like | 0.04109 | 22.87828 | FALSE | FALSE |
| LOC107069878 | ADP-ribosylation factor-related protein 1 | 0.04023 | 22.88729 | FALSE | FALSE |
| LOC107073222 | uncharacterized protein LOC107073222 | 0.04063 | 22.89226 | FALSE | FALSE |
| LOC107067391 | soma ferritin-like | 0.04223 | 22.90319 | FALSE | FALSE |
| LOC107066021 | uncharacterized protein LOC107066021 | 0.0413 | 22.90597 | FALSE | FALSE |
| LOC107072858 | microtubule-associated serine/threonine-pr | 0.04163 | 22.91761 | FALSE | FALSE |
| LOC107072522 | nuclear receptor subfamily 2 group E memb | 0.04234 | 22.92665 | FALSE | FALSE |
| LOC107071574 | stromal cell-derived factor 2-like | 0.04061 | 22.92796 | FALSE | FALSE |
| LOC107066693 | LIM domain-containing protein jub | 0.04023 | 22.94633 | FALSE | FALSE |
| LOC107067647 | uncharacterized protein LOC107067647 | 0.04083 | 22.95232 | FALSE | FALSE |
| LOC107070837 | homeobox protein 2-like | 0.04078 | 22.96616 | FALSE | FALSE |
| LOC107074164 | transmembrane protein 214-A | 0.04038 | 22.97236 | FALSE | FALSE |
| LOC107064249 | uncharacterized protein LOC107064249 | 0.04 | 22.97238 | FALSE | FALSE |
| LOC107064713 | protein toll | 0.04186 | 22.97701 | FALSE | FALSE |
| LOC107072431 | uncharacterized protein LOC107072431 | 0.03906 | 22.98306 | FALSE | FALSE |
| LOC107067021 | eukaryotic translation initiation factor 2D | 0.04244 | 22.98493 | FALSE | FALSE |
| LOC107071591 | putative uncharacterized protein DDB_G028 | 0.04028 | 23.00141 | FALSE | FALSE |
| LOC107069581 | uncharacterized protein LOC107069581 | 0.03873 | 23.01006 | FALSE | FALSE |
| LOC107070928 | fork head domain transcription factor slp2-l | 0.0405 | 23.02269 | FALSE | FALSE |
| LOC107070455 | transcription elongation regulator 1-like | 0.04011 | 23.04502 | FALSE | FALSE |
| LOC107072556 | inhibitor of nuclear factor kappa-B kinase su | 0.04053 | 23.06133 | FALSE | FALSE |

| | | | | | |
|---|---|---|---|---|---|
| LOC107069425 | gamma-aminobutyric acid receptor alpha-lil | 0.04241 | 23.0635 | FALSE | FALSE |
| LOC107069873 | multiple inositol polyphosphate phosphatas | 0.04154 | 23.06355 | FALSE | FALSE |
| LOC107069814 | uncharacterized protein LOC107069814 | 0.04125 | 23.06364 | FALSE | FALSE |
| LOC107066656 | endoplasmic reticulum mannosyl-oligosacch | 0.04028 | 23.07829 | FALSE | FALSE |
| LOC107071196 | 4-nitrophenylphosphatase-like | 0.03998 | 23.08831 | FALSE | FALSE |
| LOC107072190 | uncharacterized protein LOC107072190 | 0.04131 | 23.08977 | FALSE | FALSE |
| LOC107064891 | uncharacterized Golgi apparatus membrane | 0.03925 | 23.09312 | FALSE | FALSE |
| LOC107071852 | rabankyrin-5 | 0.04047 | 23.09326 | FALSE | FALSE |
| LOC107068076 | uncharacterized oxidoreductase YjmC | 0.03975 | 23.10189 | FALSE | FALSE |
| LOC107071102 | uncharacterized protein LOC107071102 | 0.04089 | 23.1169 | FALSE | FALSE |
| LOC107071542 | interferon-related developmental regulator | 0.04122 | 23.13947 | FALSE | FALSE |
| LOC107071250 | dihydroxyacetone phosphate acyltransferas | 0.0417 | 23.14734 | FALSE | FALSE |
| LOC107071767 | uncharacterized protein LOC107071767 | 0.04043 | 23.14943 | FALSE | FALSE |
| LOC107068968 | uncharacterized protein LOC107068968 | 0.0417 | 23.15218 | FALSE | FALSE |
| LOC107063920 | uncharacterized protein LOC107063920 | 0.04118 | 23.15721 | FALSE | FALSE |
| LOC107070895 | RNA exonuclease 4 | 0.04167 | 23.16565 | FALSE | FALSE |
| LOC107070310 | LIM domain kinase 1 | 0.04189 | 23.16945 | FALSE | FALSE |
| LOC107072890 | dual oxidase maturation factor 1 | 0.04133 | 23.19639 | FALSE | FALSE |
| LOC107070156 | uncharacterized protein LOC107070156 | 0.04173 | 23.23719 | FALSE | FALSE |
| LOC107065358 | uncharacterized protein LOC107065358 | 0.04012 | 23.23761 | FALSE | FALSE |
| LOC107068302 | E3 ubiquitin-protein ligase MARCH8 | 0.042 | 23.24798 | FALSE | FALSE |
| LOC107071885 | leucine-rich repeat-containing protein 57-lik | 0.04193 | 23.26844 | FALSE | FALSE |
| LOC107068238 | scavenger receptor class B member 1-like | 0.04034 | 23.29292 | FALSE | FALSE |
| LOC107071510 | sterol regulatory element-binding protein cl | 0.04342 | 23.30327 | FALSE | FALSE |
| LOC107067497 | uncharacterized protein LOC107067497 | 0.04019 | 23.30633 | FALSE | FALSE |
| LOC107073006 | ribonuclease Z, mitochondrial | 0.04143 | 23.32779 | FALSE | FALSE |
| LOC107066289 | G patch domain-containing protein 4 | 0.04064 | 23.32832 | FALSE | FALSE |
| LOC107068581 | inositol hexakisphosphate and diphosphoin | 0.03987 | 23.33832 | FALSE | FALSE |
| LOC107073073 | FNIP repeat-containing protein DDB_G0290 | 0.04169 | 23.34446 | FALSE | FALSE |
| LOC107066658 | ATP synthase subunit beta, mitochondrial | 0.03982 | 23.37328 | FALSE | FALSE |
| LOC107069220 | uncharacterized protein LOC107069220 | 0.03995 | 23.38873 | FALSE | FALSE |
| LOC107067245 | myosuppressin | 0.04076 | 23.39437 | FALSE | FALSE |
| LOC107070940 | uncharacterized protein LOC107070940 | 0.0406 | 23.40046 | FALSE | FALSE |
| LOC107068003 | G-protein coupled receptor Mth2-like | 0.04056 | 23.41068 | FALSE | FALSE |
| LOC107072868 | uncharacterized protein LOC107072868 | 0.04109 | 23.41211 | FALSE | FALSE |
| LOC107073139 | 2-aminoethanethiol dioxygenase | 0.04187 | 23.42409 | FALSE | FALSE |
| LOC107065440 | uncharacterized protein LOC107065440 | 0.0419 | 23.42933 | FALSE | FALSE |
| LOC107064424 | uncharacterized protein LOC107064424 | 0.04126 | 23.44082 | FALSE | FALSE |
| LOC107070056 | dnaJ homolog subfamily C member 2 | 0.04245 | 23.44495 | FALSE | FALSE |
| LOC107065930 | centromere protein I-like | 0.04084 | 23.44574 | FALSE | FALSE |
| LOC107063827 | small G protein signaling modulator 3 homo | 0.0419 | 23.44992 | FALSE | FALSE |
| LOC107065524 | protein SCAF8 | 0.03977 | 23.45204 | FALSE | FALSE |
| LOC107070822 | histone lysine demethylase PHF8-like | 0.04001 | 23.46083 | FALSE | FALSE |
| LOC107064266 | uncharacterized protein LOC107064266 | 0.04108 | 23.46811 | FALSE | FALSE |
| LOC107066388 | mitochondrial import receptor subunit TOM | 0.04011 | 23.46972 | FALSE | FALSE |
| LOC107066867 | TOX high mobility group box family member | 0.04095 | 23.4709 | FALSE | FALSE |
| LOC107073199 | rapamycin-insensitive companion of mTOR | 0.03885 | 23.47532 | FALSE | FALSE |
| LOC107073141 | uncharacterized protein LOC107073141 | 0.04159 | 23.4859 | FALSE | FALSE |
| LOC107066604 | SET and MYND domain-containing protein 4 | 0.03862 | 23.50304 | FALSE | FALSE |
| LOC107066488 | Hermansky-Pudlak syndrome 1 protein hom | 0.04156 | 23.50308 | FALSE | FALSE |

| | | | | | |
|---|---|---|---|---|---|
| LOC107065383 | MATH and LRR domain-containing protein P | 0.04004 | 23.51548 | FALSE | FALSE |
| LOC107073900 | SKI family transcriptional corepressor 2 | 0.04119 | 23.52232 | FALSE | FALSE |
| LOC107064785 | G patch domain-containing protein 11 | 0.04125 | 23.52677 | FALSE | FALSE |
| LOC107069424 | CLK4-associating serine/arginine rich protei | 0.04065 | 23.5489 | FALSE | FALSE |
| LOC107068308 | twinfilin | 0.04188 | 23.56098 | FALSE | FALSE |
| LOC107070077 | proteasome inhibitor PI31 subunit | 0.04059 | 23.56132 | FALSE | FALSE |
| LOC107070083 | uncharacterized protein LOC107070083 | 0.04007 | 23.56708 | FALSE | FALSE |
| LOC107065650 | 60S ribosomal protein L37 | 0.04251 | 23.56935 | FALSE | FALSE |
| LOC107074486 | serine-rich adhesin for platelets-like | 0.04057 | 23.57262 | FALSE | FALSE |
| LOC107070905 | syndecan | 0.03898 | 23.57488 | FALSE | FALSE |
| LOC107068884 | 60S ribosomal protein L5-like | 0.04063 | 23.57615 | FALSE | FALSE |
| LOC107064903 | protein sel-1 homolog 1 | 0.04064 | 23.57733 | FALSE | FALSE |
| LOC107066387 | 39S ribosomal protein L40, mitochondrial | 0.04119 | 23.58795 | FALSE | FALSE |
| LOC107072655 | putative tyrosine-protein kinase Wsck | 0.03918 | 23.59195 | FALSE | FALSE |
| LOC107071157 | 2-aminoadipate transaminase-like | 0.04145 | 23.59482 | FALSE | FALSE |
| LOC107064281 | succinate-semialdehyde dehydrogenase [NA | 0.04054 | 23.59585 | FALSE | FALSE |
| LOC107067020 | protein dimmed-like | 0.04058 | 23.60369 | FALSE | FALSE |
| LOC107064368 | zinc finger protein 658B | 0.03917 | 23.60986 | FALSE | FALSE |
| LOC107067153 | ER membrane protein complex subunit 10 | 0.03967 | 23.62238 | FALSE | FALSE |
| LOC107069135 | ras GTPase-activating protein-binding prote | 0.0404 | 23.62633 | FALSE | FALSE |
| LOC107065817 | multidrug resistance-associated protein 1 | 0.04143 | 23.63436 | FALSE | FALSE |
| LOC107067949 | cytochrome P450 4C1-like | 0.04156 | 23.6361 | FALSE | FALSE |
| LOC107069352 | casein kinase I isoform epsilon-like | 0.0396 | 23.63919 | FALSE | FALSE |
| LOC107066322 | uncharacterized protein LOC107066322 | 0.04156 | 23.64371 | FALSE | FALSE |
| LOC107070157 | monocarboxylate transporter 12 | 0.0412 | 23.65138 | FALSE | FALSE |
| LOC107065275 | pancreatic lipase-related protein 2-like | 0.03971 | 23.65753 | FALSE | FALSE |
| LOC107064113 | 39S ribosomal protein L41, mitochondrial | 0.04122 | 23.69405 | FALSE | FALSE |
| LOC107066222 | ras-related protein Rab-7a | 0.03889 | 23.69582 | FALSE | FALSE |
| LOC107067540 | protein unc-80 homolog | 0.04031 | 23.71373 | FALSE | FALSE |
| LOC107064390 | serine/threonine-protein kinase PRP4 homo | 0.04066 | 23.71987 | FALSE | FALSE |
| LOC107068845 | uncharacterized protein LOC107068845 | 0.03987 | 23.72482 | FALSE | FALSE |
| LOC107070216 | putative uncharacterized protein DDB_G028 | 0.03978 | 23.74161 | FALSE | FALSE |
| LOC107065113 | U5 small nuclear ribonucleoprotein 40 kDa p | 0.03984 | 23.74987 | FALSE | FALSE |
| LOC107072924 | zinc finger CCHC-type and RNA-binding mot | 0.03999 | 23.75265 | FALSE | FALSE |
| LOC107065731 | myotubularin-related protein 2 | 0.03972 | 23.75505 | FALSE | FALSE |
| LOC107072664 | protein trapped in endoderm-1 | 0.04029 | 23.7632 | FALSE | FALSE |
| LOC107065776 | selenoprotein M-like | 0.04058 | 23.78085 | FALSE | FALSE |
| LOC107072626 | uncharacterized protein LOC107072626 | 0.03989 | 23.78116 | FALSE | FALSE |
| LOC107070457 | ras suppressor protein 1 | 0.0416 | 23.79124 | FALSE | FALSE |
| LOC107071662 | zinc finger protein 528-like | 0.04043 | 23.8058 | FALSE | FALSE |
| LOC107063763 | transmembrane protein 59-like | 0.04019 | 23.80874 | FALSE | FALSE |
| LOC107066795 | protein I'm not dead yet-like | 0.03963 | 23.82781 | FALSE | FALSE |
| LOC107064206 | somatostatin receptor type 2-like | 0.03914 | 23.83015 | FALSE | FALSE |
| LOC107066083 | sodium-dependent nutrient amino acid tran | 0.04125 | 23.83043 | FALSE | FALSE |
| LOC107073315 | epimerase family protein SDR39U1 | 0.0407 | 23.84795 | FALSE | FALSE |
| LOC107071172 | uncharacterized protein LOC107071172 | 0.04108 | 23.85006 | FALSE | FALSE |
| LOC107072931 | cationic amino acid transporter 4 | 0.04076 | 23.8541 | FALSE | FALSE |
| LOC107074063 | uncharacterized protein LOC107074063 | 0.04032 | 23.85988 | FALSE | FALSE |
| LOC107064987 | epidermal growth factor receptor kinase sul | 0.04162 | 23.8633 | FALSE | FALSE |
| LOC107067754 | uncharacterized protein LOC107067754 | 0.03927 | 23.86988 | FALSE | FALSE |

| | | | | | |
|---|---|---|---|---|---|
| LOC107067875 | adenomatous polyposis coli protein | 0.03942 | 23.88174 | FALSE | FALSE |
| LOC107071015 | male-specific lethal 3 homolog | 0.04037 | 23.88666 | FALSE | FALSE |
| LOC107069156 | uncharacterized protein LOC107069156 | 0.03975 | 23.89078 | FALSE | FALSE |
| LOC107074222 | mitochondrial import inner membrane trans | 0.03902 | 23.90848 | FALSE | FALSE |
| LOC107073111 | ataxin-2 homolog | 0.03923 | 23.91496 | FALSE | FALSE |
| LOC107073903 | ETS-like protein pointed, isoform P2/D | 0.0398 | 23.91906 | FALSE | FALSE |
| LOC107071963 | irregular chiasm C-roughest protein-like | 0.03909 | 23.92231 | FALSE | FALSE |
| LOC107069033 | solute carrier family 35 member B1 | 0.04073 | 23.92723 | FALSE | FALSE |
| LOC107065435 | glutamate receptor ionotropic, kainate 3 | 0.03916 | 23.94278 | FALSE | FALSE |
| LOC107064331 | zinc finger CCCH domain-containing protein | 0.03969 | 23.9465 | FALSE | FALSE |
| LOC107073975 | piggyBac transposable element-derived pro | 0.04006 | 23.95838 | FALSE | FALSE |
| LOC107071876 | nef-associated protein 1 | 0.03962 | 23.96746 | FALSE | FALSE |
| LOC107069511 | D-beta-hydroxybutyrate dehydrogenase, mi | 0.03929 | 23.98814 | FALSE | FALSE |
| LOC107066117 | chromatin assembly factor 1 subunit A-like | 0.04013 | 23.99082 | FALSE | FALSE |
| LOC107068079 | protein will die slowly | 0.0403 | 23.99717 | FALSE | FALSE |
| LOC107072758 | uncharacterized protein LOC107072758 | 0.03986 | 23.99844 | FALSE | FALSE |
| LOC107069554 | tetraspanin-5 | 0.04035 | 24.00827 | FALSE | FALSE |
| LOC107074697 | uncharacterized protein LOC107074697 | 0.03994 | 24.01358 | FALSE | FALSE |
| LOC107063784 | 5-methylcytosine rRNA methyltransferase N | 0.04096 | 24.02895 | FALSE | FALSE |
| LOC107067667 | uncharacterized protein LOC107067667 | 0.04029 | 24.03038 | FALSE | FALSE |
| LOC107064089 | uncharacterized protein LOC107064089 | 0.0408 | 24.04421 | FALSE | FALSE |
| LOC107070584 | probable multidrug resistance-associated pr | 0.03978 | 24.06656 | FALSE | FALSE |
| LOC107069117 | uncharacterized protein LOC107069117 | 0.03899 | 24.07312 | FALSE | FALSE |
| LOC107065029 | probable G-protein coupled receptor Mth-li | 0.03913 | 24.07929 | FALSE | FALSE |
| LOC107066930 | uncharacterized protein LOC107066930 | 0.04059 | 24.09644 | FALSE | FALSE |
| LOC107071211 | uncharacterized protein LOC107071211 | 0.04121 | 24.10392 | FALSE | FALSE |
| LOC107065709 | nose resistant to fluoxetine protein 6-like | 0.04157 | 24.12373 | FALSE | FALSE |
| LOC107067535 | barrier-to-autointegration factor B | 0.04125 | 24.13157 | FALSE | FALSE |
| LOC107067519 | histone-lysine N-methyltransferase PRDM9- | 0.03987 | 24.13204 | FALSE | FALSE |
| LOC107069696 | uncharacterized protein LOC107069696 | 0.04042 | 24.14604 | FALSE | FALSE |
| LOC107066404 | histone-lysine N-methyltransferase SMYD3 | 0.04174 | 24.15288 | FALSE | FALSE |
| LOC107071134 | uncharacterized protein LOC107071134 | 0.0405 | 24.18868 | FALSE | FALSE |
| LOC107068359 | coiled-coil domain-containing protein 94 | 0.0403 | 24.19205 | FALSE | FALSE |
| LOC107064078 | leucine-rich repeat-containing protein 15-lik | 0.03903 | 24.19753 | FALSE | FALSE |
| LOC107065468 | DNA-directed RNA polymerase II subunit RP | 0.03974 | 24.20281 | FALSE | FALSE |
| LOC107072447 | uncharacterized protein LOC107072447 | 0.03895 | 24.22769 | FALSE | FALSE |
| LOC107071014 | ubiquitin-like domain-containing CTD phosp | 0.0389 | 24.22901 | FALSE | FALSE |
| LOC107066169 | U2 small nuclear ribonucleoprotein auxiliary | 0.04187 | 24.23549 | FALSE | FALSE |
| LOC107068688 | WD repeat domain phosphoinositide-intera | 0.04023 | 24.2454 | FALSE | FALSE |
| LOC107064355 | ARF GTPase-activating protein GIT1 | 0.04136 | 24.25542 | FALSE | FALSE |
| LOC107065061 | DNA topoisomerase 3-beta-1 | 0.04002 | 24.27934 | FALSE | FALSE |
| LOC107065088 | major royal jelly protein 1-like | 0.04188 | 24.28188 | FALSE | FALSE |
| LOC107069907 | mitochondrial ribonuclease P protein 1 hom | 0.03992 | 24.29961 | FALSE | FALSE |
| LOC107067533 | uncharacterized protein LOC107067533 | 0.04066 | 24.31027 | FALSE | FALSE |
| LOC107070285 | acidic phospholipase A2 PA4-like | 0.0396 | 24.31409 | FALSE | FALSE |
| LOC107064175 | lutropin-choriogonadotropic hormone recep | 0.04212 | 24.31497 | FALSE | FALSE |
| LOC107067760 | uncharacterized protein LOC107067760 | 0.04008 | 24.31559 | FALSE | FALSE |
| LOC107071146 | atlastin | 0.03962 | 24.3255 | FALSE | FALSE |
| LOC107074356 | mitochondrial inner membrane protein OXA | 0.04078 | 24.32913 | FALSE | FALSE |
| LOC107071884 | venom protease-like | 0.0405 | 24.34266 | FALSE | FALSE |

| | | | | | |
|---|---|---|---|---|---|
| LOC107064911 | myophilin | 0.03888 | 24.34857 | FALSE | FALSE |
| LOC107063616 | tetra-peptide repeat homeobox protein 1-li | 0.03994 | 24.35202 | FALSE | FALSE |
| LOC107070288 | dynein assembly factor 3, axonemal homolo | 0.03977 | 24.36767 | FALSE | FALSE |
| LOC107064253 | 60S ribosomal protein L3 | 0.0391 | 24.3781 | FALSE | FALSE |
| LOC107071176 | cytochrome P450 4C1-like | 0.03877 | 24.37925 | FALSE | FALSE |
| LOC107066209 | protein-lysine N-methyltransferase EEF2KM | 0.03951 | 24.39151 | FALSE | FALSE |
| LOC107073045 | adenylate cyclase type 6 | 0.03946 | 24.40494 | FALSE | FALSE |
| LOC107072237 | uncharacterized protein LOC107072237 | 0.03965 | 24.4227 | FALSE | FALSE |
| LOC107071809 | lethal(2)neighbour of Tid protein | 0.04005 | 24.42687 | FALSE | FALSE |
| LOC107072010 | uncharacterized protein LOC107072010 | 0.03923 | 24.42765 | FALSE | FALSE |
| LOC107064507 | uncharacterized protein LOC107064507 | 0.03988 | 24.44156 | FALSE | FALSE |
| LOC107067721 | diphthamide biosynthesis protein 2-like | 0.04047 | 24.4528 | FALSE | FALSE |
| LOC107070576 | cytosolic carboxypeptidase 4-like | 0.04001 | 24.455 | FALSE | FALSE |
| LOC107071910 | uncharacterized protein LOC107071910 | 0.03903 | 24.47556 | FALSE | FALSE |
| LOC107072104 | mitotic spindle assembly checkpoint proteir | 0.0406 | 24.48861 | FALSE | FALSE |
| LOC107064031 | uncharacterized protein LOC107064031 | 0.03967 | 24.50354 | FALSE | FALSE |
| LOC107069443 | putative thiosulfate sulfurtransferase, mitoc | 0.03973 | 24.51803 | FALSE | FALSE |
| LOC107064543 | methyltransferase-like protein 13 | 0.0396 | 24.53092 | FALSE | FALSE |
| LOC107073361 | dual specificity mitogen-activated protein ki | 0.03864 | 24.53843 | FALSE | FALSE |
| LOC107069188 | putative methyltransferase C9orf114 homol | 0.03984 | 24.54256 | FALSE | FALSE |
| LOC107065541 | uncharacterized protein LOC107065541 | 0.03912 | 24.54316 | FALSE | FALSE |
| LOC107064511 | uncharacterized protein LOC107064511 | 0.03956 | 24.56538 | FALSE | FALSE |
| LOC107066832 | uncharacterized protein LOC107066832 | 0.04251 | 24.57433 | FALSE | FALSE |
| LOC107068802 | serine/threonine-protein kinase RIO2 | 0.04109 | 24.58868 | FALSE | FALSE |
| LOC107064301 | 60S ribosomal protein L18 | 0.03976 | 24.59931 | FALSE | FALSE |
| LOC107070511 | 28S ribosomal protein S18b, mitochondrial | 0.03944 | 24.61475 | FALSE | FALSE |
| LOC107065530 | putative leucine-rich repeat-containing prot | 0.03888 | 24.61712 | FALSE | FALSE |
| LOC107072383 | signal recognition particle receptor subunit | 0.04013 | 24.62889 | FALSE | FALSE |
| LOC107067429 | serine/threonine-protein kinase MARK2-like | 0.0401 | 24.63109 | FALSE | FALSE |
| LOC107074189 | histone H4 | 0.03956 | 24.63309 | FALSE | FALSE |
| LOC107074578 | homeobox protein MSH-A | 0.04037 | 24.6403 | FALSE | FALSE |
| LOC107074263 | transcription factor Sox-17-beta.1-like | 0.04164 | 24.64075 | FALSE | FALSE |
| LOC107065562 | uncharacterized protein LOC107065562 | 0.03966 | 24.65173 | FALSE | FALSE |
| LOC107073981 | F-box only protein 32 | 0.04285 | 24.65467 | FALSE | FALSE |
| LOC107066877 | MATH and LRR domain-containing protein P | 0.04033 | 24.66406 | FALSE | FALSE |
| LOC107074689 | hyaluronan mediated motility receptor-like | 0.04094 | 24.67055 | FALSE | FALSE |
| LOC107074479 | alpha-aminoadipic semialdehyde dehydroge | 0.04021 | 24.67611 | FALSE | FALSE |
| LOC107073652 | uncharacterized protein LOC107073652 | 0.04055 | 24.68654 | FALSE | FALSE |
| LOC107069917 | LIM homeobox transcription factor 1-beta-li | 0.0399 | 24.70186 | FALSE | FALSE |
| LOC107066242 | uncharacterized protein LOC107066242 | 0.03806 | 24.70314 | FALSE | FALSE |
| LOC107073998 | uncharacterized protein LOC107073998 | 0.03867 | 24.72122 | FALSE | FALSE |
| LOC107071786 | uncharacterized protein LOC107071786 | 0.03924 | 24.72327 | FALSE | FALSE |
| LOC107068800 | rab GTPase-activating protein 1-like | 0.03998 | 24.73623 | FALSE | FALSE |
| LOC107074001 | retinal homeobox protein Rx1-like | 0.03885 | 24.74352 | FALSE | FALSE |
| LOC107066292 | transmembrane protein 234 homolog | 0.04057 | 24.74561 | FALSE | FALSE |
| LOC107065287 | TP53-regulated inhibitor of apoptosis 1-like | 0.04031 | 24.75526 | FALSE | FALSE |
| LOC107067711 | insulin-like growth factor-binding protein co | 0.03866 | 24.76447 | FALSE | FALSE |
| LOC107065480 | grpE protein homolog 1, mitochondrial | 0.04025 | 24.76791 | FALSE | FALSE |
| LOC107068604 | odorant receptor Or1-like | 0.0406 | 24.76791 | FALSE | FALSE |
| LOC107073417 | uncharacterized protein LOC107073417 | 0.03923 | 24.77767 | FALSE | FALSE |

| | | | | | |
|---|---|---|---|---|---|
| LOC107073732 | gonadotropin-releasing hormone receptor-l | 0.03898 | 24.78852 | FALSE | FALSE |
| LOC107065979 | uncharacterized protein LOC107065979 | 0.0396 | 24.79154 | FALSE | FALSE |
| LOC107073151 | ubiquitin carboxyl-terminal hydrolase calyp | 0.039 | 24.81098 | FALSE | FALSE |
| LOC107064226 | RNA polymerase II degradation factor 1-like | 0.04197 | 24.82933 | FALSE | FALSE |
| LOC107063902 | glucose dehydrogenase [FAD, quinone | 0.03942 | 24.84026 | FALSE | FALSE |
| LOC107068549 | apoptosis inhibitor 5 | 0.04032 | 24.8573 | FALSE | FALSE |
| LOC107064856 | uncharacterized protein LOC107064856 | 0.03936 | 24.86198 | FALSE | FALSE |
| LOC107069041 | uncharacterized protein LOC107069041 | 0.03981 | 24.87348 | FALSE | FALSE |
| LOC107066082 | uncharacterized protein LOC107066082 | 0.03846 | 24.8914 | FALSE | FALSE |
| LOC107063767 | uncharacterized protein LOC107063767 | 0.03981 | 24.89172 | FALSE | FALSE |
| LOC107070819 | uncharacterized protein LOC107070819 | 0.03926 | 24.9005 | FALSE | FALSE |
| LOC107067527 | zinc finger protein squeeze-like | 0.04122 | 24.92028 | FALSE | FALSE |
| LOC107074154 | signal recognition particle 54 kDa protein | 0.03944 | 24.9445 | FALSE | FALSE |
| LOC107074137 | uncharacterized family 31 glucosidase KIAA | 0.04012 | 24.94452 | FALSE | FALSE |
| LOC107064193 | glyoxal reductase | 0.03903 | 24.94776 | FALSE | FALSE |
| LOC107071436 | uncharacterized protein LOC107071436 | 0.0392 | 24.95404 | FALSE | FALSE |
| LOC107063877 | probable ATP-dependent RNA helicase DDX | 0.0401 | 24.9588 | FALSE | FALSE |
| LOC107072727 | protein rolling stone-like | 0.04037 | 24.95897 | FALSE | FALSE |
| LOC107071564 | protein MANBAL | 0.03902 | 24.96072 | FALSE | FALSE |
| LOC107063830 | uncharacterized protein LOC107063830 | 0.03793 | 24.9661 | FALSE | FALSE |
| LOC107068542 | zinc finger protein 423-like | 0.03887 | 24.98852 | FALSE | FALSE |
| LOC107071409 | cell division cycle 5-like protein | 0.03858 | 25.00939 | FALSE | FALSE |
| LOC107069877 | AP-1 complex subunit gamma-1 | 0.03979 | 25.01332 | FALSE | FALSE |
| LOC107064901 | zinc finger protein 239-like | 0.03904 | 25.01431 | FALSE | FALSE |
| LOC107068729 | uncharacterized protein LOC107068729 | 0.04084 | 25.02007 | FALSE | FALSE |
| LOC107066204 | protein phosphatase 1 regulatory subunit 1 | 0.0392 | 25.02688 | FALSE | FALSE |
| LOC107071360 | uncharacterized protein LOC107071360 | 0.03988 | 25.03089 | FALSE | FALSE |
| LOC107066861 | replication factor C subunit 3 | 0.03922 | 25.03232 | FALSE | FALSE |
| LOC107072375 | uncharacterized protein LOC107072375 | 0.03939 | 25.03655 | FALSE | FALSE |
| LOC107070145 | peritrophin-44-like | 0.04046 | 25.0426 | FALSE | FALSE |
| LOC107070851 | aprataxin | 0.03929 | 25.0516 | FALSE | FALSE |
| LOC107073717 | uncharacterized protein LOC107073717 | 0.0395 | 25.05332 | FALSE | FALSE |
| LOC107068396 | CAS1 domain-containing protein 1 | 0.04131 | 25.05363 | FALSE | FALSE |
| LOC107071982 | uncharacterized protein LOC107071982 | 0.03914 | 25.0684 | FALSE | FALSE |
| LOC107071949 | protein krueppel | 0.04064 | 25.06979 | FALSE | FALSE |
| LOC107066109 | uncharacterized protein LOC107066109 | 0.03854 | 25.07059 | FALSE | FALSE |
| LOC107064545 | glutamic acid-rich protein-like | 0.03951 | 25.07124 | FALSE | FALSE |
| LOC107070983 | potassium/sodium hyperpolarization-activa | 0.03861 | 25.08481 | FALSE | FALSE |
| LOC107071413 | synaptotagmin-4-like | 0.04014 | 25.1046 | FALSE | FALSE |
| LOC107069181 | uncharacterized protein LOC107069181 | 0.03986 | 25.14077 | FALSE | FALSE |
| LOC107073756 | uncharacterized protein LOC107073756 | 0.03988 | 25.15392 | FALSE | FALSE |
| LOC107069747 | uncharacterized protein LOC107069747 | 0.04053 | 25.15582 | FALSE | FALSE |
| LOC107072452 | Down syndrome cell adhesion molecule-like | 0.04177 | 25.15789 | FALSE | FALSE |
| LOC107072750 | small nuclear ribonucleoprotein-associated | 0.03883 | 25.16988 | FALSE | FALSE |
| LOC107063634 | tubulin epsilon chain-like | 0.03941 | 25.17044 | FALSE | FALSE |
| LOC107065783 | uncharacterized protein LOC107065783 | 0.03912 | 25.17109 | FALSE | FALSE |
| LOC107068089 | vesicle-associated membrane protein 2-like | 0.03871 | 25.17189 | FALSE | FALSE |
| LOC107072957 | cdc42 homolog | 0.03959 | 25.17527 | FALSE | FALSE |
| LOC107071516 | U6 snRNA-associated Sm-like protein LSm1 | 0.03858 | 25.17892 | FALSE | FALSE |
| LOC107065427 | protein FRA10AC1 | 0.04155 | 25.18532 | FALSE | FALSE |

| | | | | | |
|---|---|---|---|---|---|
| LOC107070110 | probable cytochrome P450 6a14 | 0.03945 | 25.19291 | FALSE | FALSE |
| LOC107069480 | probable ATP-dependent RNA helicase DHX | 0.03944 | 25.20021 | FALSE | FALSE |
| LOC107072671 | peroxiredoxin 1-like | 0.03926 | 25.21204 | FALSE | FALSE |
| Trnar-ccu | Trnar-ccu | 0.04099 | 25.21411 | FALSE | FALSE |
| LOC107063911 | acetylcholine receptor subunit alpha-type a | 0.03941 | 25.23135 | FALSE | FALSE |
| LOC107072371 | periodic tryptophan protein 2 homolog | 0.04078 | 25.23412 | FALSE | FALSE |
| LOC107069119 | uncharacterized protein LOC107069119 | 0.03926 | 25.23806 | FALSE | FALSE |
| LOC107066453 | sarcosine dehydrogenase, mitochondrial | 0.03908 | 25.24796 | FALSE | FALSE |
| LOC107072093 | uncharacterized protein LOC107072093 | 0.04 | 25.26213 | FALSE | FALSE |
| LOC107068782 | uncharacterized protein LOC107068782 | 0.0393 | 25.27564 | FALSE | FALSE |
| LOC107066736 | uncharacterized protein LOC107066736 | 0.03892 | 25.2786 | FALSE | FALSE |
| LOC107064499 | protein kibra | 0.03896 | 25.2824 | FALSE | FALSE |
| LOC107071998 | uncharacterized protein LOC107071998 | 0.0415 | 25.28639 | FALSE | FALSE |
| LOC107068197 | cytochrome c oxidase assembly factor 5 | 0.0397 | 25.2906 | FALSE | FALSE |
| LOC107066892 | uncharacterized protein LOC107066892 | 0.03991 | 25.29243 | FALSE | FALSE |
| LOC107066396 | inositol-3-phosphate synthase 1-A | 0.04002 | 25.31075 | FALSE | FALSE |
| LOC107070159 | glutamate-rich WD repeat-containing protei | 0.03894 | 25.33156 | FALSE | FALSE |
| LOC107072029 | mitogen-activated protein kinase kinase kin | 0.0407 | 25.33223 | FALSE | FALSE |
| LOC107070268 | ribonuclease P protein subunit p30 | 0.04075 | 25.33705 | FALSE | FALSE |
| LOC107071914 | cholinesterase | 0.0391 | 25.34216 | FALSE | FALSE |
| LOC107066837 | cytochrome P450 9e2-like | 0.04029 | 25.34452 | FALSE | FALSE |
| LOC107069276 | uncharacterized protein LOC107069276 | 0.03935 | 25.35401 | FALSE | FALSE |
| LOC107071965 | contactin-4-like | 0.0392 | 25.37298 | FALSE | FALSE |
| LOC107065771 | mucin-3A | 0.03856 | 25.37452 | FALSE | FALSE |
| LOC107073678 | protein tramtrack, beta isoform-like | 0.0396 | 25.39975 | FALSE | FALSE |
| LOC107068383 | GPN-loop GTPase 3 | 0.03963 | 25.42848 | FALSE | FALSE |
| LOC107067550 | calcium-dependent secretion activator | 0.03906 | 25.43605 | FALSE | FALSE |
| LOC107066842 | golgin subfamily A member 6-like protein 1 | 0.04024 | 25.44256 | FALSE | FALSE |
| LOC107071557 | uncharacterized protein LOC107071557 | 0.0392 | 25.45017 | FALSE | FALSE |
| LOC107072402 | connectin | 0.03963 | 25.46471 | FALSE | FALSE |
| LOC107069874 | protein MEMO1 | 0.04053 | 25.49694 | FALSE | FALSE |
| LOC107070713 | B9 domain-containing protein 1 | 0.04019 | 25.50136 | FALSE | FALSE |
| LOC107072043 | uncharacterized protein LOC107072043 | 0.03913 | 25.51358 | FALSE | FALSE |
| LOC107064959 | NADH dehydrogenase [ubiquinone | 0.0388 | 25.51479 | FALSE | FALSE |
| LOC107069479 | dynein-1-beta heavy chain, flagellar inner ar | 0.03924 | 25.52128 | FALSE | FALSE |
| LOC107069608 | probable ATP-dependent RNA helicase DDX | 0.03901 | 25.53073 | FALSE | FALSE |
| LOC107066556 | trimeric intracellular cation channel type B | 0.03851 | 25.54525 | FALSE | FALSE |
| LOC107072455 | solute carrier family 12 member 8 | 0.03906 | 25.54722 | FALSE | FALSE |
| LOC107063871 | uncharacterized protein LOC107063871 | 0.03961 | 25.55337 | FALSE | FALSE |
| LOC107067079 | sperm-associated antigen 6 | 0.03956 | 25.56222 | FALSE | FALSE |
| LOC107073339 | uncharacterized protein LOC107073339 | 0.0392 | 25.56627 | FALSE | FALSE |
| LOC107070817 | gastrula zinc finger protein XlCGF26.1-like | 0.03865 | 25.56859 | FALSE | FALSE |
| LOC107073857 | probable phospholipid hydroperoxide glutat | 0.03999 | 25.56996 | FALSE | FALSE |
| LOC107065533 | beta-1,4-glucuronyltransferase 1 | 0.03919 | 25.57725 | FALSE | FALSE |
| LOC107067854 | uncharacterized protein LOC107067854 | 0.03978 | 25.58199 | FALSE | FALSE |
| LOC107064395 | ATP-binding cassette sub-family B member | 0.03901 | 25.59347 | FALSE | FALSE |
| LOC107071964 | GPI mannosyltransferase 1 | 0.03906 | 25.6041 | FALSE | FALSE |
| LOC107074616 | ankyrin repeat domain-containing protein 1 | 0.03823 | 25.61352 | FALSE | FALSE |
| LOC107072558 | exocyst complex component 3 | 0.04029 | 25.61498 | FALSE | FALSE |
| LOC107065364 | uncharacterized protein LOC107065364 | 0.0403 | 25.62282 | FALSE | FALSE |

| | | | | | |
|---|---|---|---|---|---|
| LOC107069096 | uncharacterized protein LOC107069096 | 0.03904 | 25.62878 | FALSE | FALSE |
| LOC107071382 | uncharacterized protein LOC107071382 | 0.03889 | 25.63218 | FALSE | FALSE |
| LOC107070955 | zinc finger protein 567-like | 0.03992 | 25.63474 | FALSE | FALSE |
| LOC107069282 | far upstream element-binding protein 3 | 0.03909 | 25.63619 | FALSE | FALSE |
| LOC107069689 | protein PTHB1 | 0.03894 | 25.63822 | FALSE | FALSE |
| LOC107072180 | E3 ubiquitin-protein ligase RNF14-like | 0.03898 | 25.64012 | FALSE | FALSE |
| LOC107070350 | ras-related and estrogen-regulated growth i | 0.03962 | 25.64253 | FALSE | FALSE |
| LOC107065728 | uncharacterized protein LOC107065728 | 0.03767 | 25.64462 | FALSE | FALSE |
| LOC107070254 | integrin alpha-PS4-like | 0.03873 | 25.66591 | FALSE | FALSE |
| LOC107063891 | neuroligin-4, Y-linked-like | 0.0401 | 25.6802 | FALSE | FALSE |
| LOC107072005 | probable RNA methyltransferase CG11342 | 0.0397 | 25.68841 | FALSE | FALSE |
| LOC107067138 | CD63 antigen | 0.03993 | 25.68878 | FALSE | FALSE |
| LOC107071522 | GDP-fucose protein O-fucosyltransferase 1 | 0.03929 | 25.68887 | FALSE | FALSE |
| LOC107073907 | lachesin-like | 0.03877 | 25.69619 | FALSE | FALSE |
| LOC107074155 | uncharacterized protein LOC107074155 | 0.03909 | 25.70867 | FALSE | FALSE |
| LOC107073080 | calpain-D-like | 0.03969 | 25.71201 | FALSE | FALSE |
| LOC107070628 | 40S ribosomal protein S5 | 0.04008 | 25.71343 | FALSE | FALSE |
| LOC107074311 | kalirin | 0.03821 | 25.71459 | FALSE | FALSE |
| LOC107066678 | stomatin-like protein 2, mitochondrial | 0.03854 | 25.71492 | FALSE | FALSE |
| LOC107068670 | XK-related protein 6-like | 0.03993 | 25.72293 | FALSE | FALSE |
| LOC107069785 | maltase A2-like | 0.03978 | 25.74331 | FALSE | FALSE |
| LOC107071248 | sorting nexin-2 | 0.03934 | 25.7498 | FALSE | FALSE |
| LOC107067725 | major facilitator superfamily domain-contai | 0.03956 | 25.76989 | FALSE | FALSE |
| LOC107070944 | ornithine aminotransferase, mitochondrial-l | 0.04006 | 25.78248 | FALSE | FALSE |
| LOC107064806 | uncharacterized protein LOC107064806 | 0.03896 | 25.78428 | FALSE | FALSE |
| LOC107068795 | germinal-center associated nuclear protein | 0.03898 | 25.79249 | FALSE | FALSE |
| LOC107071980 | solute carrier family 41 member 1-like | 0.03927 | 25.81706 | FALSE | FALSE |
| LOC107073241 | caspase-2-like | 0.03893 | 25.82372 | FALSE | FALSE |
| LOC107073887 | PCI domain-containing protein 2 | 0.03913 | 25.8418 | FALSE | FALSE |
| LOC107066894 | DNA-directed RNA polymerase II subunit RP | 0.04082 | 25.84675 | FALSE | FALSE |
| LOC107071785 | band 3 anion transport protein | 0.03809 | 25.8517 | FALSE | FALSE |
| LOC107063681 | protein FAM161A | 0.03866 | 25.85922 | FALSE | FALSE |
| LOC107067273 | signal transducer and activator of transcript | 0.03843 | 25.86156 | FALSE | FALSE |
| LOC107065884 | filamin-A | 0.04012 | 25.87558 | FALSE | FALSE |
| LOC107064498 | iron-sulfur cluster co-chaperone protein Hs | 0.03906 | 25.87817 | FALSE | FALSE |
| LOC107073664 | uncharacterized protein LOC107073664 | 0.03938 | 25.89266 | FALSE | FALSE |
| LOC107071567 | putative uncharacterized protein DDB_G02 | 0.03933 | 25.90196 | FALSE | FALSE |
| LOC107063622 | transmembrane protease serine 9-like | 0.04047 | 25.90445 | FALSE | FALSE |
| LOC107074665 | solute carrier family 28 member 3-like | 0.03909 | 25.92186 | FALSE | FALSE |
| LOC107070777 | trehalase | 0.03913 | 25.93435 | FALSE | FALSE |
| LOC107064668 | glycine--tRNA ligase | 0.03954 | 25.95234 | FALSE | FALSE |
| LOC107074490 | LIM/homeobox protein Lhx9-like | 0.03942 | 25.95526 | FALSE | FALSE |
| LOC107066889 | uncharacterized protein LOC107066889 | 0.03908 | 25.97065 | FALSE | FALSE |
| LOC107064457 | putative fatty acyl-CoA reductase CG5065 | 0.03902 | 25.98086 | FALSE | FALSE |
| LOC107064475 | ubiquitin-40S ribosomal protein S27a-like | 0.03923 | 25.99346 | FALSE | FALSE |
| LOC107065961 | eukaryotic translation initiation factor 6 | 0.0412 | 25.99933 | FALSE | FALSE |
| LOC107071266 | kinesin heavy chain-like | 0.03818 | 26.0023 | FALSE | FALSE |
| LOC107071206 | serine protease nudel | 0.03887 | 26.00726 | FALSE | FALSE |
| LOC107069798 | 40S ribosomal protein S10-like | 0.03791 | 26.0257 | FALSE | FALSE |
| LOC107065894 | KH domain-containing, RNA-binding, signal t | 0.04022 | 26.03387 | FALSE | FALSE |

| | | | | | |
|---|---|---|---|---|---|
| LOC107066238 | uncharacterized protein LOC107066238 | 0.04015 | 26.03718 | FALSE | FALSE |
| LOC107072211 | alpha-aminoadipic semialdehyde synthase, | 0.0384 | 26.03803 | FALSE | FALSE |
| LOC107068469 | cytohesin-1 | 0.03939 | 26.04293 | FALSE | FALSE |
| LOC107069333 | putative gustatory receptor 28b | 0.03962 | 26.04693 | FALSE | FALSE |
| LOC107072365 | uncharacterized protein LOC107072365 | 0.03936 | 26.05508 | FALSE | FALSE |
| LOC107073197 | DNA mismatch repair protein Msh2 | 0.03992 | 26.05659 | FALSE | FALSE |
| LOC107065441 | glycine-rich cell wall structural protein | 0.03764 | 26.05982 | FALSE | FALSE |
| LOC107070022 | rab11 family-interacting protein 1 | 0.03862 | 26.07049 | FALSE | FALSE |
| LOC107065178 | oxidative stress-responsive serine-rich prote | 0.0389 | 26.07209 | FALSE | FALSE |
| LOC107065900 | ketohexokinase-like | 0.0381 | 26.08704 | FALSE | FALSE |
| LOC107072032 | patched domain-containing protein 3-like | 0.03831 | 26.092 | FALSE | FALSE |
| LOC107067584 | uncharacterized protein LOC107067584 | 0.0386 | 26.09337 | FALSE | FALSE |
| LOC107068401 | 60S ribosomal protein L23a-like | 0.03892 | 26.10255 | FALSE | FALSE |
| LOC107072991 | glutaredoxin-3 | 0.03931 | 26.10766 | FALSE | FALSE |
| LOC107074527 | vacuolar protein sorting-associated protein | 0.03896 | 26.10769 | FALSE | FALSE |
| LOC107070411 | uncharacterized protein LOC107070411 | 0.04028 | 26.116 | FALSE | FALSE |
| LOC107069458 | uncharacterized protein LOC107069458 | 0.03912 | 26.11675 | FALSE | FALSE |
| LOC107072144 | uncharacterized protein LOC107072144 | 0.03926 | 26.13861 | FALSE | FALSE |
| LOC107073828 | uncharacterized protein LOC107073828 | 0.03952 | 26.13963 | FALSE | FALSE |
| LOC107070477 | pancreatic triacylglycerol lipase-like | 0.03974 | 26.15551 | FALSE | FALSE |
| LOC107073583 | uncharacterized protein LOC107073583 | 0.04009 | 26.17781 | FALSE | FALSE |
| LOC107064063 | complex III assembly factor LYRM7 | 0.03857 | 26.1835 | FALSE | FALSE |
| LOC107074485 | probable basic-leucine zipper transcription f | 0.03884 | 26.18699 | FALSE | FALSE |
| LOC107063646 | proteasome subunit alpha type-2 | 0.0383 | 26.19267 | FALSE | FALSE |
| LOC107068096 | gamma-aminobutyric acid receptor-associat | 0.03837 | 26.21171 | FALSE | FALSE |
| LOC107065059 | pyruvate dehydrogenase phosphatase regul | 0.03854 | 26.24402 | FALSE | FALSE |
| LOC107067299 | protein FAM69C | 0.03985 | 26.25297 | FALSE | FALSE |
| LOC107065609 | uncharacterized protein LOC107065609 | 0.0389 | 26.27115 | FALSE | FALSE |
| LOC107063953 | telomere-associated protein RIF1-like | 0.03949 | 26.29804 | FALSE | FALSE |
| LOC107064268 | uncharacterized protein LOC107064268 | 0.0383 | 26.30782 | FALSE | FALSE |
| LOC107071009 | DNA-binding protein D-ETS-3 | 0.04135 | 26.30968 | FALSE | FALSE |
| LOC107071249 | general transcriptional corepressor trfA-like | 0.03895 | 26.32576 | FALSE | FALSE |
| LOC107072889 | GTP-binding protein GEM | 0.03984 | 26.34403 | FALSE | FALSE |
| LOC107069498 | H2.0-like homeobox protein | 0.03904 | 26.3474 | FALSE | FALSE |
| LOC107072440 | probable RNA-binding protein CG14230 | 0.03988 | 26.34877 | FALSE | FALSE |
| LOC107071722 | alpha-(1,3)-fucosyltransferase 10 | 0.03864 | 26.35373 | FALSE | FALSE |
| LOC107069211 | dynein heavy chain 1, axonemal-like | 0.03821 | 26.35473 | FALSE | FALSE |
| LOC107067409 | PC4 and SFRS1-interacting protein | 0.03884 | 26.35946 | FALSE | FALSE |
| LOC107071689 | myosin regulatory light chain sqh | 0.03873 | 26.3606 | FALSE | FALSE |
| LOC107072118 | A disintegrin and metalloproteinase with th | 0.03924 | 26.36409 | FALSE | FALSE |
| LOC107065948 | UNC93-like protein MFSD11 | 0.03987 | 26.3659 | FALSE | FALSE |
| LOC107069268 | rho-related GTP-binding protein RhoU-like | 0.03807 | 26.37054 | FALSE | FALSE |
| LOC107069746 | uncharacterized protein LOC107069746 | 0.03811 | 26.37371 | FALSE | FALSE |
| LOC107073266 | ATP-dependent DNA helicase pif1-like | 0.03936 | 26.3862 | FALSE | FALSE |
| LOC107070230 | cell adhesion molecule 2-like | 0.03935 | 26.39041 | FALSE | FALSE |
| LOC107067865 | ribosome-releasing factor 2, mitochondrial | 0.04113 | 26.39048 | FALSE | FALSE |
| LOC107067288 | transmembrane protein 256-like | 0.03724 | 26.40493 | FALSE | FALSE |
| LOC107070565 | uncharacterized protein LOC107070565 | 0.03999 | 26.43544 | FALSE | FALSE |
| LOC107064967 | lipase maturation factor 2-like | 0.03974 | 26.46105 | FALSE | FALSE |
| LOC107065489 | probable basic-leucine zipper transcription f | 0.03888 | 26.46814 | FALSE | FALSE |

| | | | | | |
|---|---|---|---|---|---|
| LOC107065864 | guanine nucleotide-binding protein-like 1 | 0.03771 | 26.48075 | FALSE | FALSE |
| LOC107064445 | dolichol-phosphate mannosyltransferase su | 0.03836 | 26.4838 | FALSE | FALSE |
| LOC107068973 | uncharacterized protein LOC107068973 | 0.03931 | 26.49281 | FALSE | FALSE |
| LOC107063641 | neurabin-1 | 0.03904 | 26.49498 | FALSE | FALSE |
| LOC107065147 | alpha-tocopherol transfer protein-like | 0.03808 | 26.51342 | FALSE | FALSE |
| LOC107069841 | uncharacterized protein LOC107069841 | 0.03823 | 26.5163 | FALSE | FALSE |
| LOC107068617 | tropomyosin-like | 0.03936 | 26.5273 | FALSE | FALSE |
| LOC107074674 | circumsporozoite protein-like | 0.03805 | 26.52864 | FALSE | FALSE |
| LOC107071214 | uncharacterized protein LOC107071214 | 0.04026 | 26.53646 | FALSE | FALSE |
| LOC107073791 | uncharacterized protein LOC107073791 | 0.03591 | 26.54607 | FALSE | FALSE |
| LOC107071119 | threonine aspartase 1 | 0.03823 | 26.54771 | FALSE | FALSE |
| LOC107074629 | SET and MYND domain-containing protein 4 | 0.03968 | 26.55547 | FALSE | FALSE |
| LOC107064663 | PDZ domain-containing protein 4-like | 0.03816 | 26.57416 | FALSE | FALSE |
| LOC107072802 | dynein heavy chain, cytoplasmic | 0.04018 | 26.58579 | FALSE | FALSE |
| LOC107069560 | actin-5C | 0.0387 | 26.58793 | FALSE | FALSE |
| LOC107064616 | gamma-glutamyl hydrolase-like | 0.03898 | 26.59961 | FALSE | FALSE |
| LOC107070035 | uncharacterized protein LOC107070035 | 0.03914 | 26.60921 | FALSE | FALSE |
| LOC107069838 | protein DENND6A | 0.03862 | 26.61364 | FALSE | FALSE |
| LOC107071930 | UPF0489 protein C5orf22 homolog | 0.03799 | 26.61702 | FALSE | FALSE |
| LOC107071670 | uncharacterized protein LOC107071670 | 0.03909 | 26.61771 | FALSE | FALSE |
| LOC107072764 | geranylgeranyl pyrophosphate synthase | 0.03844 | 26.61936 | FALSE | FALSE |
| LOC107069018 | V-type proton ATPase subunit H | 0.03935 | 26.64244 | FALSE | FALSE |
| LOC107073035 | heat shock factor protein 1 | 0.04115 | 26.65227 | FALSE | FALSE |
| LOC107066401 | protein PRRC2C | 0.03872 | 26.65367 | FALSE | FALSE |
| LOC107068093 | 60S ribosomal protein L22-like | 0.03877 | 26.65765 | FALSE | FALSE |
| LOC107072867 | rab11 family-interacting protein 4A | 0.03908 | 26.67743 | FALSE | FALSE |
| LOC107071253 | uncharacterized protein LOC107071253 | 0.03864 | 26.68201 | FALSE | FALSE |
| LOC107064090 | synaptotagmin-14 | 0.03991 | 26.68632 | FALSE | FALSE |
| LOC107072330 | leucine-rich repeat and guanylate kinase do | 0.0387 | 26.69417 | FALSE | FALSE |
| LOC107065656 | solute carrier family 35 member C2 | 0.03918 | 26.69685 | FALSE | FALSE |
| LOC107065244 | ras-related protein Rab-18-B-like | 0.03953 | 26.72669 | FALSE | FALSE |
| LOC107070942 | troponin T, skeletal muscle | 0.03875 | 26.73515 | FALSE | FALSE |
| LOC107070051 | probable G-protein coupled receptor 52 | 0.03933 | 26.74598 | FALSE | FALSE |
| LOC107068246 | uncharacterized protein LOC107068246 | 0.0378 | 26.74667 | FALSE | FALSE |
| LOC107068472 | probable aspartate aminotransferase, cytop | 0.0385 | 26.74784 | FALSE | FALSE |
| LOC107064923 | protein vestigial | 0.03822 | 26.75009 | FALSE | FALSE |
| LOC107074359 | clustered mitochondria protein homolog | 0.03851 | 26.75369 | FALSE | FALSE |
| LOC107067127 | protein KIAA0100 | 0.03704 | 26.75974 | FALSE | FALSE |
| LOC107069561 | uncharacterized protein LOC107069561 | 0.03795 | 26.76115 | FALSE | FALSE |
| LOC107067313 | LOW QUALITY PROTEIN: putative uncharact | 0.0385 | 26.79133 | FALSE | FALSE |
| LOC107072131 | aquaporin AQPAn.G | 0.03906 | 26.80222 | FALSE | FALSE |
| LOC107069720 | ras-related protein Rab-9A | 0.03794 | 26.80854 | FALSE | FALSE |
| LOC107072962 | uncharacterized protein LOC107072962 | 0.03898 | 26.81495 | FALSE | FALSE |
| LOC107068158 | brefeldin A-inhibited guanine nucleotide-ex | 0.03981 | 26.81864 | FALSE | FALSE |
| LOC107063689 | ferritin subunit | 0.03827 | 26.83322 | FALSE | FALSE |
| LOC107064033 | facilitated trehalose transporter Tret1-like | 0.0395 | 26.83603 | FALSE | FALSE |
| LOC107068447 | arginine/serine-rich protein PNISR | 0.03817 | 26.85207 | FALSE | FALSE |
| LOC107063895 | WD repeat-containing protein 89 | 0.03884 | 26.86242 | FALSE | FALSE |
| LOC107074617 | uncharacterized protein LOC107074617 | 0.03844 | 26.86629 | FALSE | FALSE |
| LOC107069833 | derlin-2 | 0.03894 | 26.87245 | FALSE | FALSE |

| | | | | | |
|---|---|---|---|---|---|
| LOC107071611 | gametogenetin-binding protein 2-like | 0.0413 | 26.87518 | FALSE | FALSE |
| LOC107072567 | dnaJ protein homolog 1 | 0.03971 | 26.88112 | FALSE | FALSE |
| LOC107072016 | ALK tyrosine kinase receptor-like | 0.03964 | 26.89415 | FALSE | FALSE |
| LOC107071156 | ribosome-recycling factor, mitochondrial | 0.03912 | 26.90113 | FALSE | FALSE |
| LOC107072971 | glutaryl-CoA dehydrogenase, mitochondrial | 0.03787 | 26.91477 | FALSE | FALSE |
| LOC107069768 | 39S ribosomal protein L51, mitochondrial | 0.04071 | 26.91681 | FALSE | FALSE |
| LOC107071053 | elongation of very long chain fatty acids pro | 0.03822 | 26.9187 | FALSE | FALSE |
| LOC107064850 | amphiphysin | 0.03821 | 26.93008 | FALSE | FALSE |
| LOC107066518 | ectonucleotide pyrophosphatase/phosphod | 0.03907 | 26.93173 | FALSE | FALSE |
| LOC107068351 | NADH dehydrogenase [ubiquinone | 0.03839 | 26.95132 | FALSE | FALSE |
| LOC107067996 | uncharacterized protein LOC107067996 | 0.03939 | 26.95716 | FALSE | FALSE |
| LOC107073433 | translin-associated protein X | 0.03823 | 26.9786 | FALSE | FALSE |
| LOC107065129 | uncharacterized protein LOC107065129 | 0.0389 | 26.97936 | FALSE | FALSE |
| LOC107073145 | ELMO domain-containing protein 2 | 0.03803 | 26.98009 | FALSE | FALSE |
| LOC107065599 | uncharacterized protein LOC107065599 | 0.03902 | 26.98521 | FALSE | FALSE |
| LOC107071812 | zinc finger protein 277 | 0.04 | 26.98688 | FALSE | FALSE |
| LOC107071583 | exocyst complex component 6 | 0.03829 | 26.98903 | FALSE | FALSE |
| LOC107074547 | nucleolar complex protein 3 homolog | 0.03827 | 26.99949 | FALSE | FALSE |
| LOC107066891 | tyrosine--tRNA ligase, mitochondrial | 0.03788 | 27.04062 | FALSE | FALSE |
| LOC107064156 | DNA-directed primase/polymerase protein- | 0.03854 | 27.06645 | FALSE | FALSE |
| LOC107068647 | AF4/FMR2 family member 4-like | 0.03759 | 27.06824 | FALSE | FALSE |
| LOC107064920 | cuticle protein 65-like | 0.039 | 27.0698 | FALSE | FALSE |
| LOC107074133 | ankyrin repeat domain-containing protein 1 | 0.03878 | 27.07138 | FALSE | FALSE |
| LOC107070932 | uncharacterized protein LOC107070932 | 0.03898 | 27.07857 | FALSE | FALSE |
| LOC107070064 | very-long-chain (3R)-3-hydroxyacyl-CoA deh | 0.03764 | 27.0876 | FALSE | FALSE |
| LOC107063771 | uncharacterized protein LOC107063771 | 0.03759 | 27.09123 | FALSE | FALSE |
| LOC107073283 | uncharacterized protein LOC107073283 | 0.03884 | 27.09301 | FALSE | FALSE |
| LOC107064579 | synaptobrevin homolog 2-like | 0.03666 | 27.10001 | FALSE | FALSE |
| LOC107072048 | uncharacterized protein LOC107072048 | 0.03867 | 27.10362 | FALSE | FALSE |
| LOC107070124 | three-prime repair exonuclease 1 | 0.03826 | 27.1065 | FALSE | FALSE |
| LOC107072682 | phosphoinositide 3-kinase regulatory subun | 0.0392 | 27.11347 | FALSE | FALSE |
| LOC107070965 | uncharacterized protein LOC107070965 | 0.03877 | 27.1165 | FALSE | FALSE |
| LOC107067733 | nuclear receptor 2C2-associated protein | 0.03859 | 27.1223 | FALSE | FALSE |
| LOC107069111 | Y+L amino acid transporter 2 | 0.03873 | 27.13105 | FALSE | FALSE |
| LOC107072997 | negative elongation factor A | 0.04083 | 27.14195 | FALSE | FALSE |
| LOC107071768 | TNF receptor-associated factor 6 | 0.03906 | 27.14773 | FALSE | FALSE |
| LOC107074694 | nucleolar GTP-binding protein 2 | 0.03812 | 27.15002 | FALSE | FALSE |
| LOC107069492 | histone H2A.V | 0.03886 | 27.17928 | FALSE | FALSE |
| LOC107069930 | isoleucine--tRNA ligase, mitochondrial | 0.03709 | 27.18039 | FALSE | FALSE |
| LOC107065306 | low density lipoprotein receptor adapter pr | 0.03844 | 27.18168 | FALSE | FALSE |
| LOC107063955 | ecto-NOX disulfide-thiol exchanger 2 | 0.03794 | 27.19652 | FALSE | FALSE |
| LOC107063839 | uncharacterized protein LOC107063839 | 0.03911 | 27.21179 | FALSE | FALSE |
| LOC107069465 | Usher syndrome type-1G protein homolog | 0.03794 | 27.21715 | FALSE | FALSE |
| LOC107066661 | tRNA-dihydrouridine(20a/20b) synthase [N/ | 0.03882 | 27.23889 | FALSE | FALSE |
| LOC107072849 | uncharacterized protein LOC107072849 | 0.03855 | 27.24128 | FALSE | FALSE |
| LOC107073794 | sodium-coupled monocarboxylate transpor | 0.03846 | 27.24645 | FALSE | FALSE |
| LOC107074119 | integrin beta-PS-like | 0.03829 | 27.2554 | FALSE | FALSE |
| LOC107070002 | uncharacterized protein LOC107070002 | 0.0377 | 27.25641 | FALSE | FALSE |
| LOC107064773 | omega-amidase NIT2 | 0.03942 | 27.25773 | FALSE | FALSE |
| LOC107067851 | BTB/POZ domain-containing protein 6-B | 0.03893 | 27.26274 | FALSE | FALSE |

| | | | | | |
|---|---|---|---|---|---|
| LOC107070980 | exosome complex component RRP46 | 0.03842 | 27.26724 | FALSE | FALSE |
| LOC107067136 | probable small nuclear ribonucleoprotein Sr | 0.03901 | 27.27127 | FALSE | FALSE |
| LOC107068156 | kinesin-like protein KIF23 | 0.03802 | 27.27606 | FALSE | FALSE |
| LOC107074462 | regulatory-associated protein of mTOR | 0.03929 | 27.29229 | FALSE | FALSE |
| LOC107067441 | uncharacterized protein LOC107067441 | 0.03932 | 27.29351 | FALSE | FALSE |
| LOC107064831 | dentin sialophosphoprotein-like | 0.03836 | 27.30372 | FALSE | FALSE |
| LOC107067710 | dolichyl-diphosphooligosaccharide--protein | 0.03829 | 27.30607 | FALSE | FALSE |
| LOC107063635 | uncharacterized protein LOC107063635 | 0.03808 | 27.31314 | FALSE | FALSE |
| LOC107067029 | slowpoke-binding protein | 0.03826 | 27.31925 | FALSE | FALSE |
| LOC107066554 | FAD synthase-like | 0.03697 | 27.33465 | FALSE | FALSE |
| LOC107074099 | DNA polymerase epsilon catalytic subunit A | 0.03904 | 27.33801 | FALSE | FALSE |
| LOC107070427 | transmembrane protein 185A | 0.03794 | 27.33905 | FALSE | FALSE |
| LOC107072242 | protein SZT2-like | 0.03919 | 27.3422 | FALSE | FALSE |
| LOC107071879 | putative ATP-dependent RNA helicase me31 | 0.03849 | 27.34294 | FALSE | FALSE |
| LOC107072616 | transcriptional regulator ATRX-like | 0.03838 | 27.34508 | FALSE | FALSE |
| LOC107070719 | cyclic AMP response element-binding prote | 0.03835 | 27.35458 | FALSE | FALSE |
| LOC107067697 | N(G),N(G)-dimethylarginine dimethylaminol | 0.03841 | 27.3636 | FALSE | FALSE |
| LOC107069319 | thyrotropin-releasing hormone receptor | 0.03933 | 27.37351 | FALSE | FALSE |
| LOC107069077 | alkylated DNA repair protein alkB homolog | 0.0371 | 27.40155 | FALSE | FALSE |
| LOC107064255 | malonyl-CoA decarboxylase, mitochondrial- | 0.0385 | 27.41832 | FALSE | FALSE |
| LOC107066984 | DNA repair protein XRCC2 | 0.03839 | 27.43876 | FALSE | FALSE |
| LOC107067184 | coiled-coil domain-containing protein 40 | 0.03704 | 27.43894 | FALSE | FALSE |
| LOC107065082 | transmembrane 9 superfamily member 4 | 0.03772 | 27.44311 | FALSE | FALSE |
| LOC107067038 | 40S ribosomal protein S26 | 0.03898 | 27.44402 | FALSE | FALSE |
| LOC107064087 | complex I intermediate-associated protein 3 | 0.03868 | 27.45423 | FALSE | FALSE |
| LOC107065830 | glutamic acid-rich protein | 0.03762 | 27.46784 | FALSE | FALSE |
| LOC107070613 | uncharacterized protein LOC107070613 | 0.03949 | 27.46985 | FALSE | FALSE |
| LOC107070949 | protein king tubby | 0.03829 | 27.47129 | FALSE | FALSE |
| LOC107069986 | RNA-binding protein fusilli | 0.04012 | 27.48407 | FALSE | FALSE |
| LOC107070502 | wiskott-Aldrich syndrome protein family me | 0.03833 | 27.48424 | FALSE | FALSE |
| LOC107066017 | probable palmitoyltransferase ZDHHC16 | 0.03844 | 27.48445 | FALSE | FALSE |
| LOC107071427 | uncharacterized protein LOC107071427 | 0.03854 | 27.48769 | FALSE | FALSE |
| LOC107070440 | transcription factor SOX-5 | 0.03951 | 27.49907 | FALSE | FALSE |
| LOC107071926 | coatomer subunit epsilon | 0.03966 | 27.50013 | FALSE | FALSE |
| LOC107073184 | tyrosine-protein phosphatase non-receptor | 0.04001 | 27.52035 | FALSE | FALSE |
| LOC107073313 | uncharacterized protein LOC107073313 | 0.03665 | 27.5231 | FALSE | FALSE |
| LOC107074534 | N-acetylgalactosamine kinase-like | 0.03718 | 27.56066 | FALSE | FALSE |
| LOC107069604 | LOW QUALITY PROTEIN: methyltransferase- | 0.03821 | 27.58766 | FALSE | FALSE |
| LOC107068169 | vacuolar protein sorting-associated protein | 0.03826 | 27.58953 | FALSE | FALSE |
| LOC107065016 | uncharacterized protein LOC107065016 | 0.03728 | 27.5907 | FALSE | FALSE |
| LOC107067200 | protein retinal degeneration B | 0.03864 | 27.59555 | FALSE | FALSE |
| LOC107065155 | tyrosine-protein kinase Src64B | 0.03847 | 27.59591 | FALSE | FALSE |
| LOC107065746 | ubiquitin-protein ligase E3B | 0.03857 | 27.60855 | FALSE | FALSE |
| LOC107074195 | S-adenosylmethionine decarboxylase proen | 0.0374 | 27.61591 | FALSE | FALSE |
| LOC107066882 | uncharacterized protein LOC107066882 | 0.03902 | 27.62195 | FALSE | FALSE |
| LOC107067736 | leucine-rich repeat-containing protein C10o | 0.0381 | 27.63142 | FALSE | FALSE |
| LOC107071309 | peptidylprolyl isomerase domain and WD re | 0.03959 | 27.64024 | FALSE | FALSE |
| LOC107069250 | uncharacterized protein LOC107069250 | 0.03794 | 27.64091 | FALSE | FALSE |
| LOC107063747 | serine/threonine-protein phosphatase 4 reg | 0.03861 | 27.65159 | FALSE | FALSE |
| LOC107063989 | steroid hormone receptor ERR1 | 0.03812 | 27.65588 | FALSE | FALSE |

| | | | | | |
|---|---|---|---|---|---|
| LOC107066433 | uncharacterized protein LOC107066433 | 0.03848 | 27.66077 | FALSE | FALSE |
| LOC107069766 | prion-like-(Q/N-rich) domain-bearing protei | 0.03953 | 27.68965 | FALSE | FALSE |
| LOC107067892 | zinc finger protein 423 homolog | 0.03811 | 27.70254 | FALSE | FALSE |
| LOC107063817 | patched domain-containing protein 3 | 0.03838 | 27.70447 | FALSE | FALSE |
| LOC107071398 | neuropathy target esterase sws | 0.0387 | 27.70595 | FALSE | FALSE |
| LOC107064599 | uncharacterized protein LOC107064599 | 0.03964 | 27.72133 | FALSE | FALSE |
| LOC107066347 | histone-lysine N-methyltransferase SETMAR | 0.03818 | 27.72162 | FALSE | FALSE |
| LOC107067960 | alpha-(1,3)-fucosyltransferase 7-like | 0.03932 | 27.72882 | FALSE | FALSE |
| LOC107066534 | 60S ribosomal protein L6-like | 0.03966 | 27.73294 | FALSE | FALSE |
| LOC107065715 | uncharacterized protein LOC107065715 | 0.04049 | 27.73396 | FALSE | FALSE |
| LOC107071019 | 60S ribosomal protein L32 | 0.03782 | 27.75417 | FALSE | FALSE |
| LOC107064062 | peroxiredoxin-6-like | 0.03814 | 27.75654 | FALSE | FALSE |
| LOC107069906 | F-box/LRR-repeat protein 4 | 0.03852 | 27.76796 | FALSE | FALSE |
| LOC107070389 | ornithine decarboxylase 1-like | 0.03905 | 27.76802 | FALSE | FALSE |
| LOC107074221 | uncharacterized protein LOC107074221 | 0.0378 | 27.77917 | FALSE | FALSE |
| LOC107070492 | trypsin 5G1-like | 0.03755 | 27.78022 | FALSE | FALSE |
| LOC107073067 | pyruvate kinase-like | 0.03885 | 27.78506 | FALSE | FALSE |
| LOC107074170 | latrophilin Cirl-like | 0.03905 | 27.8065 | FALSE | FALSE |
| LOC107069470 | ribonuclease P protein subunit p21 | 0.03854 | 27.82124 | FALSE | FALSE |
| LOC107065613 | something about silencing protein 10 | 0.03714 | 27.84038 | FALSE | FALSE |
| LOC107073737 | glucose transporter type 1 | 0.03807 | 27.84864 | FALSE | FALSE |
| LOC107070247 | nuclear pore complex protein Nup133 | 0.0388 | 27.85868 | FALSE | FALSE |
| LOC107067246 | probable ATP-dependent RNA helicase DDX | 0.03806 | 27.86967 | FALSE | FALSE |
| LOC107074413 | kinesin-like protein KIF3B | 0.03784 | 27.87153 | FALSE | FALSE |
| LOC107065571 | ATP-binding cassette sub-family D member | 0.03814 | 27.88068 | FALSE | FALSE |
| LOC107071451 | mitochondrial amidoxime reducing compon | 0.03728 | 27.88146 | FALSE | FALSE |
| LOC107064898 | probable protein phosphatase 2C 58 | 0.03819 | 27.88197 | FALSE | FALSE |
| LOC107067481 | protein Wnt-1 | 0.0381 | 27.88981 | FALSE | FALSE |
| LOC107072860 | metalloendopeptidase OMA1, mitochondria | 0.03761 | 27.89494 | FALSE | FALSE |
| LOC107067866 | D-beta-hydroxybutyrate dehydrogenase, mi | 0.03856 | 27.91021 | FALSE | FALSE |
| LOC107065566 | cAMP-dependent protein kinase catalytic su | 0.03795 | 27.91696 | FALSE | FALSE |
| LOC107069606 | uncharacterized protein LOC107069606 | 0.03866 | 27.95374 | FALSE | FALSE |
| LOC107070858 | torso-like protein | 0.03826 | 27.95959 | FALSE | FALSE |
| LOC107066172 | uncharacterized protein LOC107066172 | 0.03756 | 27.96285 | FALSE | FALSE |
| LOC107071855 | nose resistant to fluoxetine protein 6 | 0.03823 | 27.96319 | FALSE | FALSE |
| LOC107064118 | kxDL motif-containing protein CG10681 | 0.03694 | 27.98074 | FALSE | FALSE |
| LOC107063950 | UV radiation resistance associated protein | 0.03899 | 27.99229 | FALSE | FALSE |
| LOC107064318 | probable ATP-dependent RNA helicase DDX | 0.03891 | 27.99262 | FALSE | FALSE |
| LOC107069748 | uncharacterized protein LOC107069748 | 0.03927 | 27.99647 | FALSE | FALSE |
| LOC107063752 | uncharacterized protein LOC107063752 | 0.03787 | 28.00316 | FALSE | FALSE |
| LOC107072602 | alanine and glycine-rich protein-like | 0.03786 | 28.00319 | FALSE | FALSE |
| LOC107068709 | slit homolog 1 protein-like | 0.03793 | 28.00518 | FALSE | FALSE |
| LOC107071164 | uncharacterized protein LOC107071164 | 0.03755 | 28.01504 | FALSE | FALSE |
| LOC107067930 | tubulin alpha-1 chain-like | 0.03774 | 28.01665 | FALSE | FALSE |
| LOC107066981 | 39S ribosomal protein L42, mitochondrial | 0.03775 | 28.02928 | FALSE | FALSE |
| LOC107070991 | SET and MYND domain-containing protein 5 | 0.03887 | 28.03506 | FALSE | FALSE |
| LOC107072385 | uncharacterized protein LOC107072385 | 0.03872 | 28.04596 | FALSE | FALSE |
| LOC107074075 | zinc finger protein ZPR1 | 0.03667 | 28.07532 | FALSE | FALSE |
| LOC107064289 | myb-like protein P | 0.03757 | 28.08378 | FALSE | FALSE |
| LOC107066500 | uncharacterized protein LOC107066500 | 0.03858 | 28.08879 | FALSE | FALSE |

| | | | | | |
|---|---|---|---|---|---|
| LOC107071210 | uncharacterized protein LOC107071210 | 0.03759 | 28.10012 | FALSE | FALSE |
| LOC107066937 | ejaculatory bulb-specific protein 3-like | 0.03762 | 28.11022 | FALSE | FALSE |
| LOC107066907 | CDGSH iron-sulfur domain-containing prote | 0.03704 | 28.13019 | FALSE | FALSE |
| LOC107071827 | putative fatty acyl-CoA reductase CG5065 | 0.03777 | 28.14572 | FALSE | FALSE |
| LOC107068071 | activating signal cointegrator 1 | 0.03908 | 28.1562 | FALSE | FALSE |
| LOC107066718 | transmembrane protein 98-like | 0.0366 | 28.16714 | FALSE | FALSE |
| LOC107070849 | probable H/ACA ribonucleoprotein complex | 0.03745 | 28.17057 | FALSE | FALSE |
| LOC107063944 | uncharacterized protein LOC107063944 | 0.03843 | 28.17167 | FALSE | FALSE |
| LOC107064073 | NADH dehydrogenase [ubiquinone | 0.03811 | 28.17455 | FALSE | FALSE |
| LOC107066166 | cytoplasmic dynein 1 light intermediate cha | 0.03773 | 28.1818 | FALSE | FALSE |
| LOC107072078 | alpha-protein kinase 1 | 0.03764 | 28.18223 | FALSE | FALSE |
| LOC107070305 | zinc finger protein 551-like | 0.03915 | 28.19006 | FALSE | FALSE |
| LOC107064048 | elongator complex protein 2 | 0.03912 | 28.19515 | FALSE | FALSE |
| LOC107068842 | syntaxin-7-like | 0.03902 | 28.19783 | FALSE | FALSE |
| LOC107065575 | NADH dehydrogenase [ubiquinone | 0.03781 | 28.19791 | FALSE | FALSE |
| LOC107067876 | band 4.1-like protein 5 | 0.03617 | 28.21212 | FALSE | FALSE |
| LOC107073801 | protein disulfide-isomerase A3 | 0.03915 | 28.22878 | FALSE | FALSE |
| LOC107073557 | uncharacterized protein LOC107073557 | 0.03805 | 28.24677 | FALSE | FALSE |
| LOC107074017 | hsp90 co-chaperone Cdc37 | 0.03775 | 28.254 | FALSE | FALSE |
| LOC107063907 | separin | 0.03643 | 28.25507 | FALSE | FALSE |
| LOC107071988 | RNA exonuclease 1 homolog | 0.03724 | 28.26813 | FALSE | FALSE |
| LOC107068536 | uncharacterized protein LOC107068536 | 0.03843 | 28.26983 | FALSE | FALSE |
| LOC107072580 | eukaryotic translation elongation factor 1 e| | 0.03779 | 28.2881 | FALSE | FALSE |
| LOC107064840 | uncharacterized protein LOC107064840 | 0.03753 | 28.28874 | FALSE | FALSE |
| LOC107066874 | histone-lysine N-methyltransferase EHMT2 | 0.0397 | 28.33226 | FALSE | FALSE |
| LOC107067888 | RCC1 domain-containing protein 1 | 0.03792 | 28.36049 | FALSE | FALSE |
| LOC107067927 | uncharacterized protein LOC107067927 | 0.03705 | 28.36931 | FALSE | FALSE |
| LOC107067477 | uncharacterized protein LOC107067477 | 0.03759 | 28.37131 | FALSE | FALSE |
| LOC107071465 | probable cytochrome P450 301a1, mitochoı | 0.03757 | 28.37412 | FALSE | FALSE |
| LOC107066139 | diphthine methyltransferase | 0.03845 | 28.38189 | FALSE | FALSE |
| LOC107070403 | alpha-tubulin N-acetyltransferase-like | 0.03757 | 28.38718 | FALSE | FALSE |
| LOC107070558 | 3-ketoacyl-CoA thiolase, mitochondrial | 0.03919 | 28.39027 | FALSE | FALSE |
| LOC107067982 | protein Cep78 homolog | 0.03842 | 28.39271 | FALSE | FALSE |
| LOC107065316 | pupal cuticle protein-like | 0.03779 | 28.39581 | FALSE | FALSE |
| LOC107066836 | cytochrome P450 9e2-like | 0.03644 | 28.39615 | FALSE | FALSE |
| LOC107074355 | T-cell activation inhibitor, mitochondrial-like | 0.03717 | 28.40656 | FALSE | FALSE |
| LOC107070873 | facilitated trehalose transporter Tret1-like | 0.03924 | 28.4386 | FALSE | FALSE |
| LOC107069058 | uncharacterized protein LOC107069058 | 0.03791 | 28.45099 | FALSE | FALSE |
| LOC107071359 | tuberin | 0.03786 | 28.45559 | FALSE | FALSE |
| LOC107071836 | phytochrome B | 0.03701 | 28.46284 | FALSE | FALSE |
| LOC107069898 | ankyrin repeat, SAM and basic leucine zippe | 0.03791 | 28.46606 | FALSE | FALSE |
| LOC107073427 | uncharacterized protein LOC107073427 | 0.03873 | 28.4666 | FALSE | FALSE |
| LOC107065123 | adiponectin receptor protein | 0.03749 | 28.47418 | FALSE | FALSE |
| LOC107069771 | uncharacterized protein LOC107069771 | 0.03811 | 28.48551 | FALSE | FALSE |
| LOC107070496 | acyl-CoA synthetase family member 2, mito | 0.03717 | 28.49333 | FALSE | FALSE |
| LOC107063698 | uncharacterized aarF domain-containing prc | 0.03762 | 28.50238 | FALSE | FALSE |
| LOC107065733 | Y+L amino acid transporter 2 | 0.0389 | 28.50242 | FALSE | FALSE |
| LOC107069185 | complexin | 0.03712 | 28.50487 | FALSE | FALSE |
| LOC107068840 | pleckstrin homology domain-containing fan | 0.03959 | 28.50842 | FALSE | FALSE |
| LOC107063874 | myb-like protein P | 0.03868 | 28.50935 | FALSE | FALSE |

| | | | | | |
|---|---|---|---|---|---|
| LOC107065331 | nuclear receptor subfamily 1 group D memb | 0.03838 | 28.51092 | FALSE | FALSE |
| LOC107063643 | YTH domain-containing protein 1 | 0.03868 | 28.51309 | FALSE | FALSE |
| LOC107066235 | 4-aminobutyrate aminotransferase, mitocho | 0.03783 | 28.52388 | FALSE | FALSE |
| LOC107064246 | nuclear nucleic acid-binding protein C1D-like | 0.03637 | 28.52521 | FALSE | FALSE |
| LOC107068737 | tubby-related protein 4 | 0.03605 | 28.53431 | FALSE | FALSE |
| LOC107069403 | dnaJ homolog subfamily B member 6-like | 0.03738 | 28.54118 | FALSE | FALSE |
| LOC107067551 | uncharacterized protein LOC107067551 | 0.03852 | 28.54735 | FALSE | FALSE |
| LOC107067390 | putative leucine-rich repeat-containing prot | 0.03854 | 28.55033 | FALSE | FALSE |
| LOC107068626 | succinate dehydrogenase assembly factor 3, | 0.03921 | 28.55524 | FALSE | FALSE |
| LOC107069978 | zinc finger protein 271-like | 0.03612 | 28.55895 | FALSE | FALSE |
| LOC107063983 | breast cancer metastasis-suppressor 1-like p | 0.03798 | 28.56042 | FALSE | FALSE |
| LOC107068969 | IQ domain-containing protein H-like | 0.03677 | 28.56262 | FALSE | FALSE |
| LOC107069064 | putative mediator of RNA polymerase II tran | 0.03761 | 28.56768 | FALSE | FALSE |
| LOC107064391 | glycerate kinase-like | 0.03897 | 28.59284 | FALSE | FALSE |
| LOC107065505 | LOW QUALITY PROTEIN: NADPH-dependent | 0.03838 | 28.59639 | FALSE | FALSE |
| LOC107069859 | RNA exonuclease 1 | 0.0376 | 28.60359 | FALSE | FALSE |
| LOC107070643 | uncharacterized protein LOC107070643 | 0.03855 | 28.61846 | FALSE | FALSE |
| LOC107071511 | ankyrin repeat and LEM domain-containing | 0.03834 | 28.61949 | FALSE | FALSE |
| LOC107071504 | uncharacterized protein LOC107071504 | 0.0379 | 28.63891 | FALSE | FALSE |
| LOC107067771 | uncharacterized protein LOC107067771 | 0.03855 | 28.64833 | FALSE | FALSE |
| LOC107070916 | nucleoporin NDC1 | 0.03868 | 28.6623 | FALSE | FALSE |
| LOC107067564 | jouberin-like | 0.03646 | 28.67071 | FALSE | FALSE |
| LOC107068659 | putative fatty acyl-CoA reductase CG5065 | 0.03682 | 28.69155 | FALSE | FALSE |
| LOC107068098 | uncharacterized protein LOC107068098 | 0.03771 | 28.69562 | FALSE | FALSE |
| LOC107065747 | pre-mRNA-processing factor 19 | 0.03783 | 28.70322 | FALSE | FALSE |
| LOC107068908 | uncharacterized protein LOC107068908 | 0.03688 | 28.71208 | FALSE | FALSE |
| LOC107067907 | uncharacterized protein LOC107067907 | 0.04004 | 28.73862 | FALSE | FALSE |
| LOC107068211 | uncharacterized protein LOC107068211 | 0.03773 | 28.76011 | FALSE | FALSE |
| LOC107067361 | ATP-binding cassette sub-family G member | 0.03802 | 28.76506 | FALSE | FALSE |
| LOC107066639 | sorting nexin-29 | 0.03752 | 28.77048 | FALSE | FALSE |
| LOC107067151 | BAG family molecular chaperone regulator 2 | 0.03815 | 28.7783 | FALSE | FALSE |
| LOC107074243 | LOW QUALITY PROTEIN: otoferlin-like | 0.038 | 28.78348 | FALSE | FALSE |
| LOC107066721 | syndetin | 0.03684 | 28.78649 | FALSE | FALSE |
| LOC107068403 | ras GTPase-activating protein 1 | 0.03833 | 28.78974 | FALSE | FALSE |
| LOC107074388 | centromere protein J | 0.03725 | 28.81215 | FALSE | FALSE |
| LOC107069045 | innexin inx1 | 0.03797 | 28.82022 | FALSE | FALSE |
| LOC107067398 | integrin-linked protein kinase homolog pat-4 | 0.03948 | 28.83161 | FALSE | FALSE |
| LOC107069622 | BTB/POZ domain-containing protein 17 | 0.0382 | 28.83451 | FALSE | FALSE |
| LOC107068118 | liprin-alpha-1 | 0.039 | 28.83709 | FALSE | FALSE |
| LOC107065289 | DNA polymerase subunit gamma-1, mitocho | 0.03736 | 28.84284 | FALSE | FALSE |
| LOC107071822 | peptidyl-prolyl cis-trans isomerase-like 6 | 0.03781 | 28.85214 | FALSE | FALSE |
| LOC107072386 | ammonium transporter Rh type B | 0.03777 | 28.85934 | FALSE | FALSE |
| LOC107066771 | receptor-type tyrosine-protein phosphatase | 0.03788 | 28.86877 | FALSE | FALSE |
| LOC107068622 | nucleoside diphosphate-linked moiety X mo | 0.03665 | 28.86901 | FALSE | FALSE |
| LOC107072803 | isocitrate dehydrogenase [NAD | 0.03823 | 28.88091 | FALSE | FALSE |
| LOC107066868 | acyl-CoA dehydrogenase family member 9, | 0.0375 | 28.88354 | FALSE | FALSE |
| LOC107066113 | E3 ubiquitin-protein ligase arih1 | 0.03792 | 28.88891 | FALSE | FALSE |
| LOC107066779 | ETS translocation variant 1 | 0.037 | 28.89202 | FALSE | FALSE |
| LOC107063680 | uncharacterized protein LOC107063680 | 0.03913 | 28.89727 | FALSE | FALSE |
| LOC107066816 | pre-mRNA-splicing factor SPF27 | 0.03859 | 28.89873 | FALSE | FALSE |

| | | | | | |
|---|---|---|---|---|---|
| LOC107065219 | aldose 1-epimerase | 0.03679 | 28.91741 | FALSE | FALSE |
| LOC107067549 | uncharacterized protein LOC107067549 | 0.03651 | 28.92634 | FALSE | FALSE |
| LOC107066886 | F-box only protein 33 | 0.03818 | 28.94258 | FALSE | FALSE |
| LOC107067492 | Sjoegren syndrome nuclear autoantigen 1 h | 0.03732 | 28.94394 | FALSE | FALSE |
| LOC107064441 | uncharacterized protein LOC107064441 | 0.03894 | 28.94841 | FALSE | FALSE |
| LOC107069893 | proton-coupled amino acid transporter 1-lik | 0.03789 | 28.96011 | FALSE | FALSE |
| LOC107073133 | ecotropic viral integration site 5 ortholog | 0.03718 | 28.96425 | FALSE | FALSE |
| LOC107063785 | scavenger receptor class B member 1 | 0.03935 | 28.96638 | FALSE | FALSE |
| LOC107064212 | serine proteinase stubble | 0.03873 | 28.96775 | FALSE | FALSE |
| LOC107073379 | uncharacterized protein LOC107073379 | 0.0385 | 28.98215 | FALSE | FALSE |
| LOC107070433 | nucleolar protein 16 | 0.03765 | 28.9965 | FALSE | FALSE |
| LOC107073411 | protein tipE | 0.0361 | 29.00221 | FALSE | FALSE |
| LOC107069059 | prostaglandin E2 receptor EP3 subtype | 0.03715 | 29.00893 | FALSE | FALSE |
| LOC107074219 | nucleoplasmin-like protein | 0.03872 | 29.00904 | FALSE | FALSE |
| LOC107073030 | protein strawberry notch-like | 0.03598 | 29.01207 | FALSE | FALSE |
| LOC107070429 | tubulin alpha-1C chain-like | 0.03783 | 29.0163 | FALSE | FALSE |
| LOC107068672 | protein fuzzy homolog | 0.03663 | 29.01782 | FALSE | FALSE |
| LOC107069964 | N-alpha-acetyltransferase 20 | 0.03749 | 29.02631 | FALSE | FALSE |
| LOC107068399 | dual specificity protein phosphatase Mpk3 | 0.03651 | 29.02814 | FALSE | FALSE |
| LOC107066410 | probable cysteine--tRNA ligase, mitochondr | 0.03736 | 29.03133 | FALSE | FALSE |
| LOC107065801 | A-kinase anchor protein 10, mitochondrial | 0.0392 | 29.03225 | FALSE | FALSE |
| LOC107067857 | ribosomal RNA processing protein 36 homo | 0.0372 | 29.03278 | FALSE | FALSE |
| LOC107065675 | cell growth-regulating nucleolar protein | 0.03732 | 29.04015 | FALSE | FALSE |
| LOC107072916 | putative N-acetylglucosamine-6-phosphate | 0.03815 | 29.05281 | FALSE | FALSE |
| LOC107071384 | protein TSSC1 | 0.03701 | 29.05697 | FALSE | FALSE |
| LOC107070523 | POC1 centriolar protein homolog A-like | 0.03775 | 29.0624 | FALSE | FALSE |
| LOC107066976 | SET and MYND domain-containing protein 4 | 0.03695 | 29.07444 | FALSE | FALSE |
| LOC107063862 | EF-hand calcium-binding domain-containing | 0.03689 | 29.07726 | FALSE | FALSE |
| LOC107067165 | putative leucine-rich repeat-containing prot | 0.03762 | 29.10729 | FALSE | FALSE |
| LOC107065616 | 39S ribosomal protein L20, mitochondrial | 0.03698 | 29.1089 | FALSE | FALSE |
| LOC107071105 | uncharacterized protein LOC107071105 | 0.03692 | 29.11444 | FALSE | FALSE |
| LOC107066143 | venom allergen 5 | 0.0383 | 29.11782 | FALSE | FALSE |
| LOC107071231 | uncharacterized protein LOC107071231 | 0.03677 | 29.12902 | FALSE | FALSE |
| LOC107070679 | aminopeptidase N-like | 0.03679 | 29.1324 | FALSE | FALSE |
| LOC107072422 | transmembrane protein 145 | 0.0353 | 29.13765 | FALSE | FALSE |
| LOC107068388 | probable 28S ribosomal protein S23, mitoch | 0.03758 | 29.14003 | FALSE | FALSE |
| LOC107066057 | sulfhydryl oxidase 2-like | 0.03962 | 29.14773 | FALSE | FALSE |
| LOC107063861 | uncharacterized protein LOC107063861 | 0.03881 | 29.15895 | FALSE | FALSE |
| LOC107070208 | tetra-peptide repeat homeobox protein 1-li | 0.03862 | 29.16469 | FALSE | FALSE |
| LOC107064367 | uncharacterized protein LOC107064367 | 0.0367 | 29.21391 | FALSE | FALSE |
| LOC107072173 | chitin-binding lectin 1-like | 0.03803 | 29.24602 | FALSE | FALSE |
| LOC107069756 | GDP-mannose 4,6 dehydratase | 0.03846 | 29.25494 | FALSE | FALSE |
| LOC107069876 | insulin-like peptide receptor | 0.03746 | 29.25872 | FALSE | FALSE |
| LOC107071013 | ATP-dependent 6-phosphofructokinase | 0.03854 | 29.26748 | FALSE | FALSE |
| LOC107065580 | F-box only protein 21-like | 0.03683 | 29.26878 | FALSE | FALSE |
| LOC107068834 | zinc metalloproteinase nas-13 | 0.03671 | 29.27102 | FALSE | FALSE |
| LOC107066133 | protein arginine N-methyltransferase 6 | 0.03817 | 29.28735 | FALSE | FALSE |
| LOC107074175 | sphingosine kinase 2-like | 0.0375 | 29.28868 | FALSE | FALSE |
| LOC107074015 | translocon-associated protein subunit beta | 0.03806 | 29.29028 | FALSE | FALSE |
| LOC107063723 | protein penguin | 0.03793 | 29.29646 | FALSE | FALSE |

| | | | | | |
|---|---|---|---|---|---|
| LOC107065282 | uncharacterized protein LOC107065282 | 0.03543 | 29.30865 | FALSE | FALSE |
| LOC107072905 | uncharacterized protein LOC107072905 | 0.03701 | 29.30888 | FALSE | FALSE |
| LOC107072856 | serine/threonine-protein kinase Nek5-like | 0.03615 | 29.31327 | FALSE | FALSE |
| LOC107072221 | uncharacterized protein LOC107072221 | 0.03787 | 29.3273 | FALSE | FALSE |
| LOC107074345 | uncharacterized protein LOC107074345 | 0.03653 | 29.33284 | FALSE | FALSE |
| LOC107071371 | uncharacterized protein LOC107071371 | 0.03832 | 29.34405 | FALSE | FALSE |
| LOC107066026 | fasciclin-1 | 0.03833 | 29.34673 | FALSE | FALSE |
| LOC107067179 | 28S ribosomal protein S7, mitochondrial | 0.03727 | 29.34704 | FALSE | FALSE |
| LOC107071697 | metalloproteinase inhibitor 1 | 0.03695 | 29.35661 | FALSE | FALSE |
| LOC107071588 | integrator complex subunit 5 | 0.03682 | 29.3692 | FALSE | FALSE |
| LOC107074218 | protein tilB-like | 0.03837 | 29.36932 | FALSE | FALSE |
| LOC107071137 | tRNA (guanine-N(7)-)-methyltransferase no | 0.03745 | 29.37688 | FALSE | FALSE |
| LOC107074282 | protein unc-79 homolog | 0.03686 | 29.39027 | FALSE | FALSE |
| LOC107070906 | tetratricopeptide repeat protein 5-like | 0.03752 | 29.39717 | FALSE | FALSE |
| LOC107072792 | uncharacterized protein LOC107072792 | 0.03705 | 29.40943 | FALSE | FALSE |
| LOC107072721 | protein shifted-like | 0.0367 | 29.42672 | FALSE | FALSE |
| LOC107067035 | AMMECR1-like protein | 0.03682 | 29.44039 | FALSE | FALSE |
| LOC107069527 | EH domain-binding protein 1 | 0.03709 | 29.44372 | FALSE | FALSE |
| LOC107068010 | uncharacterized protein LOC107068010 | 0.03772 | 29.44825 | FALSE | FALSE |
| LOC107068529 | BRCA1-A complex subunit BRE-like | 0.03671 | 29.44979 | FALSE | FALSE |
| LOC107066090 | dual specificity mitogen-activated protein ki | 0.0369 | 29.47471 | FALSE | FALSE |
| LOC107064465 | cyclin-dependent kinase 2-interacting prote | 0.03673 | 29.48161 | FALSE | FALSE |
| LOC107073289 | mitochondrial import inner membrane tran | 0.0377 | 29.48723 | FALSE | FALSE |
| LOC107067843 | nucleolar complex protein 4 homolog B | 0.03808 | 29.49135 | FALSE | FALSE |
| LOC107070865 | carbonic anhydrase 1-like | 0.03835 | 29.5052 | FALSE | FALSE |
| LOC107065567 | merlin | 0.03785 | 29.50801 | FALSE | FALSE |
| LOC107064452 | forkhead box protein D3-like | 0.03723 | 29.50999 | FALSE | FALSE |
| LOC107066801 | protein OPI10 homolog | 0.03693 | 29.51448 | FALSE | FALSE |
| LOC107066286 | probable deoxyhypusine synthase | 0.03931 | 29.52566 | FALSE | FALSE |
| LOC107064080 | uncharacterized protein LOC107064080 | 0.0364 | 29.52571 | FALSE | FALSE |
| LOC107064990 | nephrin-like | 0.03689 | 29.5564 | FALSE | FALSE |
| LOC107074178 | Fanconi anemia group M protein | 0.03764 | 29.56145 | FALSE | FALSE |
| LOC107070380 | mitochondrial fission process protein 1 | 0.03767 | 29.56596 | FALSE | FALSE |
| LOC107074421 | calcium release-activated calcium channel p | 0.03607 | 29.57414 | FALSE | FALSE |
| LOC107068815 | ubiquitin carboxyl-terminal hydrolase 10 | 0.03635 | 29.58312 | FALSE | FALSE |
| LOC107070889 | DDB1- and CUL4-associated factor 7 | 0.03817 | 29.58803 | FALSE | FALSE |
| LOC107067660 | 2-oxoglutarate dehydrogenase, mitochondr | 0.03689 | 29.58984 | FALSE | FALSE |
| LOC107065564 | protein spitz-like | 0.03654 | 29.59116 | FALSE | FALSE |
| LOC107065826 | broad-complex core protein isoforms 1/2/3 | 0.0387 | 29.60177 | FALSE | FALSE |
| LOC107063644 | carbohydrate sulfotransferase 11-like | 0.03694 | 29.60269 | FALSE | FALSE |
| LOC107067340 | transmembrane protein 245 | 0.03827 | 29.60454 | FALSE | FALSE |
| LOC107066337 | mitochondrial 10-formyltetrahydrofolate de | 0.03659 | 29.60482 | FALSE | FALSE |
| LOC107066349 | protein canopy 4 | 0.03817 | 29.61074 | FALSE | FALSE |
| LOC107069432 | guanine nucleotide-binding protein subunit | 0.03806 | 29.61945 | FALSE | FALSE |
| LOC107066013 | uncharacterized protein LOC107066013 | 0.03745 | 29.62013 | FALSE | FALSE |
| LOC107074540 | potassium voltage-gated channel protein Sh | 0.0386 | 29.62817 | FALSE | FALSE |
| LOC107073109 | peptidyl-prolyl cis-trans isomerase D | 0.03947 | 29.63048 | FALSE | FALSE |
| LOC107074053 | uncharacterized protein LOC107074053 | 0.03692 | 29.632 | FALSE | FALSE |
| LOC107066429 | uncharacterized protein LOC107066429 | 0.03829 | 29.68129 | FALSE | FALSE |
| LOC107071846 | transcriptional repressor CTCFL | 0.0361 | 29.68665 | FALSE | FALSE |

| | | | | | |
|---|---|---|---|---|---|
| LOC107072899 | uncharacterized protein LOC107072899 | 0.03648 | 29.68817 | FALSE | FALSE |
| LOC107068721 | octopamine receptor beta-2R | 0.03681 | 29.70518 | FALSE | FALSE |
| LOC107066353 | E3 ubiquitin-protein ligase TRIM37-like | 0.03627 | 29.72589 | FALSE | FALSE |
| LOC107064906 | nucleobindin-2-like | 0.03862 | 29.73088 | FALSE | FALSE |
| LOC107071581 | uncharacterized protein LOC107071581 | 0.0371 | 29.73314 | FALSE | FALSE |
| LOC107072094 | kynurenine--oxoglutarate transaminase 3 | 0.03796 | 29.74048 | FALSE | FALSE |
| LOC107071590 | protein transport protein Sec61 subunit alpl | 0.03975 | 29.74275 | FALSE | FALSE |
| LOC107072347 | extracellular matrix-binding protein ebh | 0.03561 | 29.75379 | FALSE | FALSE |
| LOC107065779 | uncharacterized protein LOC107065779 | 0.03774 | 29.76565 | FALSE | FALSE |
| LOC107072861 | protein MON2 homolog | 0.03666 | 29.76644 | FALSE | FALSE |
| LOC107071736 | glucose dehydrogenase [FAD, quinone | 0.03696 | 29.76732 | FALSE | FALSE |
| LOC107069063 | P17/29C-like protein DDB_G0287399 | 0.03753 | 29.81977 | FALSE | FALSE |
| LOC107064295 | histone acetyltransferase KAT6B | 0.03618 | 29.82546 | FALSE | FALSE |
| LOC107071290 | ras-related protein Rab-5B | 0.03607 | 29.83563 | FALSE | FALSE |
| LOC107065065 | probable cleavage and polyadenylation spe | 0.03758 | 29.83976 | FALSE | FALSE |
| LOC107074025 | transmembrane protein 203 | 0.03901 | 29.84337 | FALSE | FALSE |
| LOC107072244 | DENN domain-containing protein 5B | 0.03517 | 29.8478 | FALSE | FALSE |
| LOC107068247 | uncharacterized protein LOC107068247 | 0.03657 | 29.85751 | FALSE | FALSE |
| LOC107065871 | dynein heavy chain 3, axonemal | 0.03695 | 29.8583 | FALSE | FALSE |
| LOC107066924 | leucine--tRNA ligase, cytoplasmic | 0.03859 | 29.8588 | FALSE | FALSE |
| LOC107070639 | kinesin-like protein KIF2A | 0.03627 | 29.87085 | FALSE | FALSE |
| LOC107066813 | uncharacterized protein LOC107066813 | 0.03681 | 29.88969 | FALSE | FALSE |
| LOC107066407 | rab-like protein 6 | 0.03713 | 29.90203 | FALSE | FALSE |
| LOC107067472 | uncharacterized protein LOC107067472 | 0.03766 | 29.91278 | FALSE | FALSE |
| LOC107069808 | plexin domain-containing protein 2 | 0.03812 | 29.93472 | FALSE | FALSE |
| LOC107064999 | insulin-like growth factor-binding protein-re | 0.03655 | 29.94616 | FALSE | FALSE |
| LOC107067937 | MORN repeat-containing protein 5-like | 0.03643 | 29.9575 | FALSE | FALSE |
| LOC107066136 | mono [ADP-ribose | 0.03617 | 29.95949 | FALSE | FALSE |
| LOC107071947 | uncharacterized protein LOC107071947 | 0.03772 | 29.96203 | FALSE | FALSE |
| LOC107065054 | p21-activated protein kinase-interacting prc | 0.03671 | 29.97256 | FALSE | FALSE |
| LOC107068850 | uncharacterized protein LOC107068850 | 0.03837 | 29.98107 | FALSE | FALSE |
| LOC107067372 | kinase D-interacting substrate of 220 kDa-lil | 0.03751 | 29.98607 | FALSE | FALSE |
| LOC107071404 | uncharacterized protein LOC107071404 | 0.03649 | 29.99092 | FALSE | FALSE |
| LOC107073922 | uncharacterized protein LOC107073922 | 0.03776 | 29.99188 | FALSE | FALSE |
| LOC107070048 | RISC-loading complex subunit TARBP2-like | 0.03839 | 29.99384 | FALSE | FALSE |
| LOC107070454 | transmembrane protein 161B | 0.03789 | 30.03076 | FALSE | FALSE |
| LOC107065588 | LOW QUALITY PROTEIN: DNA polymerase b | 0.03838 | 30.03404 | FALSE | FALSE |
| Trnar-ccu | Trnar-ccu | 0.03717 | 30.03833 | FALSE | FALSE |
| LOC107073608 | protein hairy-like | 0.03637 | 30.04237 | FALSE | FALSE |
| LOC107069987 | uncharacterized protein LOC107069987 | 0.0374 | 30.0436 | FALSE | FALSE |
| LOC107064855 | histone-lysine N-methyltransferase ash1 | 0.03684 | 30.05461 | FALSE | FALSE |
| LOC107067824 | aromatic-L-amino-acid decarboxylase | 0.03637 | 30.05633 | FALSE | FALSE |
| LOC107072804 | uncharacterized protein LOC107072804 | 0.03639 | 30.05831 | FALSE | FALSE |
| LOC107064934 | uncharacterized protein LOC107064934 | 0.03796 | 30.06316 | FALSE | FALSE |
| LOC107066846 | FAD synthase-like | 0.03789 | 30.07806 | FALSE | FALSE |
| LOC107073890 | uncharacterized protein LOC107073890 | 0.03764 | 30.07819 | FALSE | FALSE |
| LOC107070106 | uncharacterized protein LOC107070106 | 0.03657 | 30.07991 | FALSE | FALSE |
| LOC107071254 | galanin receptor type 1-like | 0.03715 | 30.10913 | FALSE | FALSE |
| LOC107064988 | prominin-1-A | 0.03629 | 30.13502 | FALSE | FALSE |
| LOC107073648 | uncharacterized protein LOC107073648 | 0.03533 | 30.14943 | FALSE | FALSE |

| | | | | | |
|---|---|---|---|---|---|
| LOC107070654 | uncharacterized protein LOC107070654 | 0.0377 | 30.15728 | FALSE | FALSE |
| LOC107071594 | aminoacylase-1-like | 0.03681 | 30.17453 | FALSE | FALSE |
| LOC107071065 | histone-lysine N-methyltransferase eggless | 0.03692 | 30.18377 | FALSE | FALSE |
| LOC107073115 | NEDD8-activating enzyme E1 regulatory sub | 0.03891 | 30.1904 | FALSE | FALSE |
| LOC107074005 | ras-like protein 1 | 0.03721 | 30.20088 | FALSE | FALSE |
| LOC107069252 | hornerin-like | 0.03986 | 30.20847 | FALSE | FALSE |
| LOC107074505 | phytanoyl-CoA dioxygenase domain-contair | 0.03758 | 30.21123 | FALSE | FALSE |
| LOC107074612 | uncharacterized protein LOC107074612 | 0.03671 | 30.21151 | FALSE | FALSE |
| LOC107066996 | intraflagellar transport protein 172 homolo | 0.03664 | 30.21675 | FALSE | FALSE |
| LOC107070850 | uncharacterized protein LOC107070850 | 0.03699 | 30.219 | FALSE | FALSE |
| LOC107068898 | probable protein phosphatase 2C T23F11.1 | 0.03697 | 30.21996 | FALSE | FALSE |
| LOC107064456 | DBF4-type zinc finger-containing protein 2 h | 0.03716 | 30.22351 | FALSE | FALSE |
| LOC107073430 | uncharacterized protein LOC107073430 | 0.03723 | 30.23332 | FALSE | FALSE |
| LOC107071442 | protein kinase C, brain isozyme | 0.03597 | 30.2436 | FALSE | FALSE |
| LOC107070734 | sorting nexin-4-like | 0.03598 | 30.24913 | FALSE | FALSE |
| LOC107068582 | ATP-dependent RNA helicase p62-like | 0.03674 | 30.25027 | FALSE | FALSE |
| LOC107072160 | calcium-binding mitochondrial carrier prote | 0.03695 | 30.26612 | FALSE | FALSE |
| LOC107069090 | mitogen-activated protein kinase kinase kin | 0.03756 | 30.27185 | FALSE | FALSE |
| LOC107067036 | histone H3.3-like | 0.03709 | 30.27579 | FALSE | FALSE |
| LOC107064411 | troponin C, isoform 1 | 0.03735 | 30.28001 | FALSE | FALSE |
| LOC107064801 | WW domain-containing adapter protein wit | 0.03699 | 30.29224 | FALSE | FALSE |
| LOC107066806 | dynamin-like 120 kDa protein, mitochondri | 0.03692 | 30.31163 | FALSE | FALSE |
| LOC107070573 | uncharacterized protein LOC107070573 | 0.03827 | 30.32061 | FALSE | FALSE |
| LOC107070458 | myelin transcription factor 1-like | 0.036 | 30.32363 | FALSE | FALSE |
| LOC107072339 | serine-rich adhesin for platelets-like | 0.03611 | 30.3321 | FALSE | FALSE |
| LOC107066555 | opsin, ultraviolet-sensitive | 0.03705 | 30.34549 | FALSE | FALSE |
| LOC107069658 | glutamate-gated chloride channel | 0.03662 | 30.34985 | FALSE | FALSE |
| LOC107070712 | protein THEM6-like | 0.03787 | 30.35308 | FALSE | FALSE |
| LOC107065989 | protein RFT1 homolog | 0.03749 | 30.4018 | FALSE | FALSE |
| LOC107073787 | cytochrome P450 4C1-like | 0.03671 | 30.43361 | FALSE | FALSE |
| LOC107065843 | uncharacterized protein LOC107065843 | 0.03652 | 30.44032 | FALSE | FALSE |
| LOC107073913 | 5'-AMP-activated protein kinase catalytic su | 0.03768 | 30.44936 | FALSE | FALSE |
| LOC107065705 | raf homolog serine/threonine-protein kinas | 0.03679 | 30.46817 | FALSE | FALSE |
| LOC107072251 | eukaryotic translation initiation factor 3 sub | 0.03639 | 30.469 | FALSE | FALSE |
| LOC107074406 | uncharacterized protein LOC107074406 | 0.03659 | 30.47398 | FALSE | FALSE |
| LOC107070617 | mediator of RNA polymerase II transcriptior | 0.03565 | 30.48815 | FALSE | FALSE |
| LOC107063855 | 28S ribosomal protein S35, mitochondrial | 0.03637 | 30.49743 | FALSE | FALSE |
| LOC107073822 | poly(U)-specific endoribonuclease homolog | 0.03744 | 30.49815 | FALSE | FALSE |
| LOC107065568 | LOW QUALITY PROTEIN: zinc finger protein | 0.03616 | 30.49998 | FALSE | FALSE |
| LOC107067121 | uncharacterized protein LOC107067121 | 0.03679 | 30.52033 | FALSE | FALSE |
| LOC107071898 | DNA-directed RNA polymerases I, II, and III s | 0.0365 | 30.5361 | FALSE | FALSE |
| LOC107071403 | cell division cycle 5-like protein | 0.03733 | 30.55542 | FALSE | FALSE |
| LOC107067615 | ATP synthase subunit delta, mitochondrial | 0.0365 | 30.56209 | FALSE | FALSE |
| LOC107074121 | protein IWS1 homolog | 0.03573 | 30.56316 | FALSE | FALSE |
| LOC107064055 | V-type proton ATPase 116 kDa subunit a iso | 0.03724 | 30.56717 | FALSE | FALSE |
| LOC107067631 | ancient ubiquitous protein 1-like | 0.03571 | 30.57743 | FALSE | FALSE |
| LOC107069521 | transmembrane protein 8A-like | 0.03726 | 30.58402 | FALSE | FALSE |
| LOC107072513 | ski oncogene | 0.03716 | 30.59748 | FALSE | FALSE |
| LOC107067071 | sialin-like | 0.03684 | 30.60377 | FALSE | FALSE |
| LOC107064972 | mitochondrial import inner membrane tran | 0.03606 | 30.61042 | FALSE | FALSE |

| | | | | | |
|---|---|---|---|---|---|
| LOC107069695 | lipopolysaccharide-induced tumor necrosis | 0.03716 | 30.62461 | FALSE | FALSE |
| LOC107065622 | WD repeat-containing protein 92 | 0.03741 | 30.6294 | FALSE | FALSE |
| LOC107073982 | aurora kinase C-like | 0.03661 | 30.63564 | FALSE | FALSE |
| LOC107072213 | ralA-binding protein 1 | 0.03701 | 30.64693 | FALSE | FALSE |
| LOC107070542 | protein yippee-like 5 | 0.03633 | 30.66613 | FALSE | FALSE |
| LOC107067587 | programmed cell death protein 5 | 0.03754 | 30.67211 | FALSE | FALSE |
| LOC107074703 | uncharacterized protein LOC107074703 | 0.03634 | 30.71136 | FALSE | FALSE |
| LOC107069477 | low molecular weight phosphotyrosine prot | 0.03564 | 30.71189 | FALSE | FALSE |
| LOC107063858 | translocon-associated protein subunit delta | 0.03638 | 30.71456 | FALSE | FALSE |
| LOC107064520 | protein bcn92 | 0.03651 | 30.71986 | FALSE | FALSE |
| LOC107070297 | uncharacterized protein LOC107070297 | 0.03718 | 30.72869 | FALSE | FALSE |
| LOC107069353 | L-xylulose reductase-like | 0.03865 | 30.73676 | FALSE | FALSE |
| LOC107070487 | hemicentin-1 | 0.03659 | 30.73849 | FALSE | FALSE |
| LOC107069384 | uncharacterized protein LOC107069384 | 0.036 | 30.74201 | FALSE | FALSE |
| LOC107065413 | uncharacterized protein LOC107065413 | 0.03663 | 30.74732 | FALSE | FALSE |
| LOC107064904 | monocarboxylate transporter 12 | 0.03716 | 30.75605 | FALSE | FALSE |
| LOC107072008 | uncharacterized protein LOC107072008 | 0.03628 | 30.76229 | FALSE | FALSE |
| LOC107067769 | LOW QUALITY PROTEIN: uncharacterized pr | 0.03772 | 30.76385 | FALSE | FALSE |
| LOC107065504 | ubiquitin thioesterase trabid | 0.03764 | 30.76514 | FALSE | FALSE |
| LOC107064421 | plexin-B | 0.03601 | 30.77117 | FALSE | FALSE |
| LOC107069440 | uncharacterized protein LOC107069440 | 0.03587 | 30.78194 | FALSE | FALSE |
| LOC107068441 | MATH and LRR domain-containing protein P | 0.03787 | 30.80136 | FALSE | FALSE |
| LOC107073904 | uncharacterized protein LOC107073904 | 0.03681 | 30.80578 | FALSE | FALSE |
| LOC107064187 | maltase A3-like | 0.03542 | 30.8108 | FALSE | FALSE |
| LOC107069939 | facilitated trehalose transporter Tret1-2 hor | 0.03646 | 30.82584 | FALSE | FALSE |
| LOC107072683 | S phase cyclin A-associated protein in the er | 0.03671 | 30.82909 | FALSE | FALSE |
| LOC107063947 | MD-2-related lipid-recognition protein-like | 0.03738 | 30.85601 | FALSE | FALSE |
| LOC107065512 | zinc finger protein 26 | 0.03749 | 30.86678 | FALSE | FALSE |
| LOC107066665 | GATA zinc finger domain-containing protein | 0.03665 | 30.87224 | FALSE | FALSE |
| LOC107065596 | peroxisome biogenesis factor 2 | 0.03749 | 30.88095 | FALSE | FALSE |
| LOC107072185 | TBC1 domain family member 23 | 0.03575 | 30.88404 | FALSE | FALSE |
| LOC107067406 | transcription factor SUM-1 | 0.036 | 30.89027 | FALSE | FALSE |
| LOC107073873 | E3 ubiquitin-protein ligase CHIP | 0.03796 | 30.90077 | FALSE | FALSE |
| LOC107067722 | DNA polymerase delta subunit 3-like | 0.03621 | 30.90563 | FALSE | FALSE |
| LOC107073094 | kinesin-like protein KIF20A | 0.03621 | 30.9195 | FALSE | FALSE |
| LOC107069219 | uncharacterized protein LOC107069219 | 0.0374 | 30.92067 | FALSE | FALSE |
| LOC107064483 | uncharacterized protein LOC107064483 | 0.03747 | 30.9236 | FALSE | FALSE |
| LOC107066403 | integrator complex subunit 10 | 0.03624 | 30.93569 | FALSE | FALSE |
| LOC107069926 | developmentally-regulated GTP-binding pro | 0.03741 | 30.94476 | FALSE | FALSE |
| LOC107064578 | long-chain fatty acid transport protein 4-like | 0.03734 | 30.94825 | FALSE | FALSE |
| LOC107063996 | N-alpha-acetyltransferase 35, NatC auxiliary | 0.03584 | 30.94839 | FALSE | FALSE |
| LOC107064188 | uncharacterized protein LOC107064188 | 0.03593 | 30.95291 | FALSE | FALSE |
| LOC107073300 | RNA-directed DNA polymerase from mobile | 0.03653 | 30.95472 | FALSE | FALSE |
| LOC107066623 | PQ-loop repeat-containing protein 3 | 0.03495 | 30.96425 | FALSE | FALSE |
| LOC107070399 | DNA-directed RNA polymerases I, II, and III s | 0.0358 | 30.98468 | FALSE | FALSE |
| LOC107071107 | odorant receptor 4-like | 0.03549 | 30.99336 | FALSE | FALSE |
| LOC107069461 | WD repeat domain phosphoinositide-intera | 0.03629 | 31.00603 | FALSE | FALSE |
| LOC107065794 | NADH dehydrogenase [ubiquinone | 0.03829 | 31.00804 | FALSE | FALSE |
| LOC107064635 | protein THEM6-like | 0.0377 | 31.00907 | FALSE | FALSE |
| LOC107065814 | 40S ribosomal protein S17 | 0.03691 | 31.01024 | FALSE | FALSE |

| | | | | | |
|---|---|---|---|---|---|
| LOC107064722 | uncharacterized protein LOC107064722 | 0.03742 | 31.02153 | FALSE | FALSE |
| LOC107071080 | elongation of very long chain fatty acids pro | 0.03715 | 31.02274 | FALSE | FALSE |
| LOC107073687 | protein cycle-like | 0.03676 | 31.04432 | FALSE | FALSE |
| LOC107065386 | clathrin light chain | 0.03571 | 31.04682 | FALSE | FALSE |
| LOC107065490 | nuclear cap-binding protein subunit 2 | 0.03574 | 31.0529 | FALSE | FALSE |
| LOC107069872 | activating signal cointegrator 1 complex sub | 0.03684 | 31.06446 | FALSE | FALSE |
| LOC107067517 | zinc finger protein 468-like | 0.03791 | 31.06453 | FALSE | FALSE |
| LOC107069847 | uncharacterized protein LOC107069847 | 0.03627 | 31.08109 | FALSE | FALSE |
| LOC107067696 | fukutin-related protein | 0.0372 | 31.0854 | FALSE | FALSE |
| LOC107066211 | G patch domain and ankyrin repeat-contain | 0.03654 | 31.08927 | FALSE | FALSE |
| LOC107066214 | nucleolar protein 11 | 0.0353 | 31.09489 | FALSE | FALSE |
| LOC107066158 | aldehyde dehydrogenase X, mitochondrial | 0.03651 | 31.09558 | FALSE | FALSE |
| LOC107072611 | pantothenate kinase 1 | 0.03706 | 31.10153 | FALSE | FALSE |
| LOC107064657 | uncharacterized protein LOC107064657 | 0.03654 | 31.11388 | FALSE | FALSE |
| LOC107070662 | obg-like ATPase 1 | 0.03652 | 31.12456 | FALSE | FALSE |
| LOC107067990 | uncharacterized protein LOC107067990 | 0.03606 | 31.12753 | FALSE | FALSE |
| LOC107064995 | BTB/POZ domain-containing protein KCTD3 | 0.03833 | 31.12939 | FALSE | FALSE |
| LOC107073191 | condensin complex subunit 2-like | 0.03623 | 31.13317 | FALSE | FALSE |
| LOC107070933 | uncharacterized protein LOC107070933 | 0.03675 | 31.13332 | FALSE | FALSE |
| LOC107066872 | T-complex protein 1 subunit eta | 0.03608 | 31.16715 | FALSE | FALSE |
| LOC107069711 | general odorant-binding protein 56d-like | 0.03555 | 31.17186 | FALSE | FALSE |
| LOC107073255 | uncharacterized protein LOC107073255 | 0.03584 | 31.18004 | FALSE | FALSE |
| LOC107071113 | electron transfer flavoprotein-ubiquinone o | 0.03595 | 31.20594 | FALSE | FALSE |
| LOC107063808 | uncharacterized protein LOC107063808 | 0.03736 | 31.2074 | FALSE | FALSE |
| LOC107064517 | syntaxin-16 | 0.03591 | 31.22426 | FALSE | FALSE |
| LOC107068881 | EF-hand domain-containing family member | 0.0357 | 31.22548 | FALSE | FALSE |
| LOC107065322 | galactosylgalactosylxylosylprotein 3-beta-gl | 0.03645 | 31.23083 | FALSE | FALSE |
| LOC107065451 | probable methyltransferase BTM2 homolog | 0.03569 | 31.23191 | FALSE | FALSE |
| Trnak-uuu | Trnak-uuu | 0.03667 | 31.23778 | FALSE | FALSE |
| LOC107065044 | monocarboxylate transporter 10-like | 0.03686 | 31.2496 | FALSE | FALSE |
| LOC107073934 | rho GTPase-activating protein 44-like | 0.0363 | 31.2673 | FALSE | FALSE |
| LOC107069374 | uncharacterized protein LOC107069374 | 0.03655 | 31.29079 | FALSE | FALSE |
| LOC107074680 | methyl-CpG-binding domain protein 3 | 0.03702 | 31.29122 | FALSE | FALSE |
| LOC107070953 | uncharacterized protein LOC107070953 | 0.03642 | 31.31656 | FALSE | FALSE |
| LOC107063833 | uncharacterized protein LOC107063833 | 0.03467 | 31.32251 | FALSE | FALSE |
| LOC107073782 | receptor-type tyrosine-protein phosphatase | 0.03651 | 31.33167 | FALSE | FALSE |
| LOC107066041 | probable ATP-dependent RNA helicase Dbp | 0.03589 | 31.33932 | FALSE | FALSE |
| LOC107071304 | mitochondrial-processing peptidase subunit | 0.03659 | 31.34049 | FALSE | FALSE |
| LOC107065335 | nuclear cap-binding protein subunit 3-like | 0.0369 | 31.34378 | FALSE | FALSE |
| LOC107073201 | adenylyltransferase and sulfurtransferase M | 0.03613 | 31.34903 | FALSE | FALSE |
| LOC107069377 | coiled-coil domain-containing protein 6 | 0.03783 | 31.37911 | FALSE | FALSE |
| LOC107071428 | RNA polymerase II-associated protein 3-like | 0.03666 | 31.38642 | FALSE | FALSE |
| LOC107067643 | LDLR chaperone boca | 0.03541 | 31.40482 | FALSE | FALSE |
| LOC107066829 | zinc finger CCCH-type with G patch domain- | 0.03411 | 31.40614 | FALSE | FALSE |
| LOC107070636 | GTP-binding protein 1 | 0.03528 | 31.40663 | FALSE | FALSE |
| LOC107068572 | hormone-sensitive lipase | 0.03638 | 31.41412 | FALSE | FALSE |
| LOC107069692 | acetylcholine receptor subunit alpha-L1 | 0.03615 | 31.41823 | FALSE | FALSE |
| LOC107067397 | esterase E4-like | 0.03623 | 31.42153 | FALSE | FALSE |
| LOC107066271 | uncharacterized protein LOC107066271 | 0.03652 | 31.42791 | FALSE | FALSE |
| LOC107070073 | deoxyribodipyrimidine photo-lyase | 0.03635 | 31.42849 | FALSE | FALSE |

| | | | | | |
|---|---|---|---|---|---|
| LOC107064557 | beta-1,4-N-acetylgalactosaminyltransferase | 0.0369 | 31.451 | FALSE | FALSE |
| LOC107069116 | protein FAN-like | 0.03827 | 31.45383 | FALSE | FALSE |
| LOC107071878 | PI-PLC X domain-containing protein 1-like | 0.03755 | 31.48393 | FALSE | FALSE |
| LOC107066586 | fasciculation and elongation protein zeta-2 | 0.03765 | 31.48624 | FALSE | FALSE |
| LOC107065323 | alcohol dehydrogenase class-3 | 0.03752 | 31.48629 | FALSE | FALSE |
| LOC107072566 | A disintegrin and metalloproteinase with th | 0.03683 | 31.49623 | FALSE | FALSE |
| LOC107065041 | uncharacterized protein LOC107065041 | 0.03691 | 31.49929 | FALSE | FALSE |
| LOC107070302 | glutathione S-transferase-like | 0.03561 | 31.50588 | FALSE | FALSE |
| LOC107071867 | uncharacterized protein LOC107071867 | 0.03694 | 31.50624 | FALSE | FALSE |
| LOC107068046 | UPF0545 protein C22orf39 homolog | 0.03596 | 31.51177 | FALSE | FALSE |
| LOC107066228 | 28S ribosomal protein S36, mitochondrial | 0.03619 | 31.51789 | FALSE | FALSE |
| LOC107067926 | cytochrome P450 4C1-like | 0.03439 | 31.53384 | FALSE | FALSE |
| LOC107065573 | protein Mpv17 | 0.03556 | 31.54255 | FALSE | FALSE |
| LOC107070107 | zinc finger FYVE domain-containing protein | 0.03609 | 31.54627 | FALSE | FALSE |
| LOC107065457 | ATP-dependent (S)-NAD(P)H-hydrate dehyd | 0.0362 | 31.54648 | FALSE | FALSE |
| LOC107064591 | uncharacterized protein LOC107064591 | 0.03769 | 31.55242 | FALSE | FALSE |
| LOC107069624 | uncharacterized protein LOC107069624 | 0.03589 | 31.56539 | FALSE | FALSE |
| LOC107070913 | DNA replication licensing factor Mcm6 | 0.03647 | 31.56832 | FALSE | FALSE |
| LOC107069982 | nuclear pore complex protein DDB_G02749 | 0.03614 | 31.58596 | FALSE | FALSE |
| LOC107071422 | integral membrane protein 2C | 0.03659 | 31.6223 | FALSE | FALSE |
| LOC107065321 | uncharacterized protein LOC107065321 | 0.03727 | 31.62792 | FALSE | FALSE |
| LOC107073267 | centromere-associated protein E-like | 0.03584 | 31.64548 | FALSE | FALSE |
| LOC107067070 | uncharacterized protein LOC107067070 | 0.03604 | 31.64943 | FALSE | FALSE |
| LOC107069032 | sulfatase-modifying factor 1 | 0.03604 | 31.65324 | FALSE | FALSE |
| LOC107064880 | uncharacterized protein LOC107064880 | 0.03702 | 31.65578 | FALSE | FALSE |
| LOC107066774 | Golgi integral membrane protein 4-like | 0.03603 | 31.65699 | FALSE | FALSE |
| LOC107066408 | proto-oncogene tyrosine-protein kinase RO: | 0.03586 | 31.66364 | FALSE | FALSE |
| LOC107067796 | uncharacterized protein LOC107067796 | 0.0366 | 31.66471 | FALSE | FALSE |
| LOC107074463 | glutamate receptor ionotropic, kainate 2 | 0.03789 | 31.67209 | FALSE | FALSE |
| LOC107069707 | protein sprint | 0.03652 | 31.68038 | FALSE | FALSE |
| LOC107073971 | stAR-related lipid transfer protein 3 | 0.03652 | 31.68699 | FALSE | FALSE |
| LOC107067092 | uncharacterized protein LOC107067092 | 0.03531 | 31.69453 | FALSE | FALSE |
| LOC107073292 | protein SEC13 homolog | 0.03643 | 31.7027 | FALSE | FALSE |
| LOC107073522 | uncharacterized protein LOC107073522 | 0.03636 | 31.71034 | FALSE | FALSE |
| LOC107074409 | Fanconi anemia group D2 protein homolog | 0.03571 | 31.72848 | FALSE | FALSE |
| LOC107072419 | inositol 1,4,5-trisphosphate receptor | 0.03571 | 31.73573 | FALSE | FALSE |
| LOC107071979 | thyroid adenoma-associated protein | 0.03643 | 31.75471 | FALSE | FALSE |
| LOC107064327 | ephrin type-B receptor 1-B | 0.03592 | 31.75672 | FALSE | FALSE |
| LOC107067454 | cytochrome b-c1 complex subunit Rieske, m | 0.0371 | 31.75712 | FALSE | FALSE |
| LOC107074110 | potassium voltage-gated channel protein Sh | 0.03797 | 31.76676 | FALSE | FALSE |
| LOC107072776 | bifunctional coenzyme A synthase | 0.03723 | 31.76774 | FALSE | FALSE |
| LOC107066022 | kazrin | 0.03593 | 31.77582 | FALSE | FALSE |
| LOC107069525 | myb-like protein X | 0.03803 | 31.80379 | FALSE | FALSE |
| LOC107064659 | protein lethal(2)essential for life-like | 0.03593 | 31.82849 | FALSE | FALSE |
| LOC107066866 | F-box/WD repeat-containing protein 5 | 0.03567 | 31.85709 | FALSE | FALSE |
| LOC107064279 | ADP-dependent glucokinase | 0.03571 | 31.858 | FALSE | FALSE |
| LOC107072132 | short-chain dehydrogenase/reductase famil | 0.0357 | 31.86604 | FALSE | FALSE |
| LOC107072390 | reticulon-4-interacting protein 1 homolog, r | 0.03639 | 31.90437 | FALSE | FALSE |
| LOC107067575 | coronin-7 | 0.0361 | 31.90457 | FALSE | FALSE |
| LOC107068357 | solute carrier family 25 member 51 | 0.03656 | 31.92352 | FALSE | FALSE |

| | | | | | |
|---|---|---|---|---|---|
| LOC107067485 | villin-1 | 0.03579 | 31.92652 | FALSE | FALSE |
| LOC107072622 | cysteine-rich PDZ-binding protein | 0.03589 | 31.92809 | FALSE | FALSE |
| LOC107063999 | acyl-CoA Delta(11) desaturase-like | 0.03578 | 31.93444 | FALSE | FALSE |
| LOC107074087 | sn1-specific diacylglycerol lipase beta-like | 0.03697 | 31.93631 | FALSE | FALSE |
| LOC107073286 | leucine-rich repeat-containing protein 15-lik | 0.03592 | 31.95173 | FALSE | FALSE |
| LOC107069678 | microprocessor complex subunit DGCR8 | 0.0341 | 31.95395 | FALSE | FALSE |
| LOC107065179 | sialin | 0.03461 | 31.96624 | FALSE | FALSE |
| LOC107071012 | tyrosine-protein kinase Abl | 0.03608 | 31.968 | FALSE | FALSE |
| LOC107069502 | uncharacterized protein LOC107069502 | 0.03656 | 31.97602 | FALSE | FALSE |
| LOC107063711 | glutamyl aminopeptidase-like | 0.03624 | 31.98087 | FALSE | FALSE |
| LOC107073966 | protein VAC14 homolog | 0.03566 | 31.98836 | FALSE | FALSE |
| LOC107066747 | uncharacterized protein LOC107066747 | 0.0351 | 31.99967 | FALSE | FALSE |
| LOC107067418 | protein Wnt-10b | 0.03736 | 32.00599 | FALSE | FALSE |
| LOC107072113 | uncharacterized protein LOC107072113 | 0.0359 | 32.01692 | FALSE | FALSE |
| LOC107064700 | uncharacterized protein LOC107064700 | 0.03585 | 32.0303 | FALSE | FALSE |
| LOC107074054 | glutamate receptor ionotropic, delta-1-like | 0.03589 | 32.05644 | FALSE | FALSE |
| LOC107069920 | uncharacterized protein LOC107069920 | 0.03651 | 32.06764 | FALSE | FALSE |
| LOC107070725 | nucleoporin Nup37 | 0.03687 | 32.06888 | FALSE | FALSE |
| LOC107063726 | molybdopterin synthase catalytic subunit | 0.03681 | 32.07817 | FALSE | FALSE |
| LOC107071903 | myb-like protein A | 0.03618 | 32.08745 | FALSE | FALSE |
| LOC107063753 | uncharacterized protein LOC107063753 | 0.03708 | 32.09315 | FALSE | FALSE |
| LOC107073349 | uncharacterized protein LOC107073349 | 0.03667 | 32.1043 | FALSE | FALSE |
| LOC107072842 | uncharacterized protein LOC107072842 | 0.03627 | 32.11487 | FALSE | FALSE |
| LOC107067882 | general odorant-binding protein 69a-like | 0.03602 | 32.12377 | FALSE | FALSE |
| LOC107067199 | discoidin domain-containing receptor 2-like | 0.03604 | 32.12574 | FALSE | FALSE |
| LOC107067627 | deoxyribose-phosphate aldolase | 0.03623 | 32.13136 | FALSE | FALSE |
| LOC107074649 | 3-ketodihydrosphingosine reductase | 0.03642 | 32.13225 | FALSE | FALSE |
| LOC107064588 | vacuolar protein sorting-associated protein | 0.03557 | 32.13388 | FALSE | FALSE |
| LOC107073233 | protein crumbs | 0.03748 | 32.13485 | FALSE | FALSE |
| LOC107073095 | apoptotic protease-activating factor 1 | 0.03665 | 32.13542 | FALSE | FALSE |
| LOC107071076 | uncharacterized protein LOC107071076 | 0.03582 | 32.1742 | FALSE | FALSE |
| LOC107063706 | BUD13 homolog | 0.03571 | 32.19289 | FALSE | FALSE |
| LOC107063667 | uncharacterized protein LOC107063667 | 0.03619 | 32.20161 | FALSE | FALSE |
| LOC107072709 | uncharacterized protein LOC107072709 | 0.0366 | 32.207 | FALSE | FALSE |
| LOC107064126 | angiogenic factor with G patch and FHA don | 0.03565 | 32.22819 | FALSE | FALSE |
| LOC107067044 | myb-like protein I | 0.03606 | 32.23092 | FALSE | FALSE |
| LOC107072814 | uncharacterized protein LOC107072814 | 0.0358 | 32.24366 | FALSE | FALSE |
| LOC107071244 | uncharacterized protein LOC107071244 | 0.0355 | 32.25246 | FALSE | FALSE |
| LOC107064723 | peroxisome assembly protein 12 | 0.03655 | 32.28273 | FALSE | FALSE |
| LOC107064679 | exostosin-3 | 0.03571 | 32.2873 | FALSE | FALSE |
| LOC107071632 | protein quiver-like | 0.03582 | 32.29458 | FALSE | FALSE |
| LOC107068491 | uncharacterized protein LOC107068491 | 0.03459 | 32.31481 | FALSE | FALSE |
| LOC107064270 | GATA zinc finger domain-containing protein | 0.03678 | 32.32827 | FALSE | FALSE |
| LOC107067354 | RDS/peripherin-like protein xRDS36 | 0.03567 | 32.33352 | FALSE | FALSE |
| LOC107070861 | uncharacterized protein LOC107070861 | 0.03505 | 32.3361 | FALSE | FALSE |
| LOC107073511 | histone-lysine N-methyltransferase SETMAF | 0.03498 | 32.36839 | FALSE | FALSE |
| LOC107071834 | uncharacterized protein LOC107071834 | 0.03718 | 32.37685 | FALSE | FALSE |
| LOC107063781 | COMM domain-containing protein 4 | 0.0353 | 32.4026 | FALSE | FALSE |
| LOC107064471 | homeobox protein 5-like | 0.03564 | 32.40358 | FALSE | FALSE |
| LOC107065121 | acetyl-coenzyme A synthetase | 0.03584 | 32.43304 | FALSE | FALSE |

| | | | | | |
|---|---|---|---|---|---|
| LOC107065024 | transmembrane protein 47 | 0.03514 | 32.43598 | FALSE | FALSE |
| LOC107066281 | adenosine deaminase CECR1-like | 0.03571 | 32.44202 | FALSE | FALSE |
| LOC107073833 | NADH dehydrogenase (ubiquinone) complex | 0.03606 | 32.44218 | FALSE | FALSE |
| LOC107072550 | kinesin-like protein costa | 0.03586 | 32.45201 | FALSE | FALSE |
| LOC107067103 | 23 kDa integral membrane protein-like | 0.03648 | 32.46897 | FALSE | FALSE |
| LOC107074019 | ribonuclease P protein subunit p20 | 0.03551 | 32.4788 | FALSE | FALSE |
| LOC107068567 | reticulon-4-interacting protein 1, mitochonc | 0.03589 | 32.48566 | FALSE | FALSE |
| LOC107074226 | probable U3 small nucleolar RNA-associatec | 0.03491 | 32.5206 | FALSE | FALSE |
| LOC107065034 | plastin-2 | 0.03696 | 32.5442 | FALSE | FALSE |
| LOC107072112 | cytochrome P450 4C1-like | 0.03547 | 32.55347 | FALSE | FALSE |
| LOC107069602 | pancreatic triacylglycerol lipase-like | 0.03582 | 32.57156 | FALSE | FALSE |
| LOC107074387 | mediator of RNA polymerase II transcriptior | 0.03652 | 32.5788 | FALSE | FALSE |
| LOC107064874 | uncharacterized protein LOC107064874 | 0.03608 | 32.58002 | FALSE | FALSE |
| LOC107070704 | uncharacterized protein LOC107070704 | 0.03651 | 32.58611 | FALSE | FALSE |
| LOC107071406 | rho GTPase-activating protein 26 | 0.03679 | 32.61314 | FALSE | FALSE |
| LOC107064404 | dystrobrevin beta | 0.03562 | 32.61592 | FALSE | FALSE |
| LOC107071438 | LOW QUALITY PROTEIN: uncharacterized pr | 0.0353 | 32.62841 | FALSE | FALSE |
| LOC107070723 | hornerin-like | 0.03609 | 32.65395 | FALSE | FALSE |
| LOC107064486 | palmitoyltransferase ZDHHC3 | 0.03702 | 32.6729 | FALSE | FALSE |
| LOC107066295 | histone deacetylase 3 | 0.03551 | 32.68611 | FALSE | FALSE |
| LOC107067007 | ankyrin repeat domain-containing protein 1 | 0.0351 | 32.70546 | FALSE | FALSE |
| LOC107066536 | uncharacterized protein LOC107066536 | 0.03582 | 32.7116 | FALSE | FALSE |
| LOC107068391 | EF-hand domain-containing protein D2 hom | 0.03491 | 32.71897 | FALSE | FALSE |
| LOC107072333 | ankyrin repeat family A protein 2-like | 0.03667 | 32.72416 | FALSE | FALSE |
| LOC107064186 | neutral and basic amino acid transport prot | 0.03618 | 32.75057 | FALSE | FALSE |
| LOC107068127 | 60S ribosomal protein L13 | 0.03701 | 32.75785 | FALSE | FALSE |
| LOC107072328 | tectonic-3-like | 0.0363 | 32.77197 | FALSE | FALSE |
| LOC107064400 | MATH and LRR domain-containing protein P | 0.03612 | 32.77292 | FALSE | FALSE |
| LOC107071958 | microsomal triglyceride transfer protein larg | 0.03573 | 32.78165 | FALSE | FALSE |
| LOC107073711 | uncharacterized protein LOC107073711 | 0.03402 | 32.79293 | FALSE | FALSE |
| LOC107071692 | IQ domain-containing protein D-like | 0.03525 | 32.79359 | FALSE | FALSE |
| LOC107070423 | probable phospholipid-transporting ATPase | 0.03536 | 32.79747 | FALSE | FALSE |
| LOC107064413 | chromobox protein homolog 1-like | 0.03525 | 32.80151 | FALSE | FALSE |
| LOC107072009 | uncharacterized protein LOC107072009 | 0.03554 | 32.80664 | FALSE | FALSE |
| LOC107069888 | histone-lysine N-methyltransferase, H3 lysir | 0.03774 | 32.82037 | FALSE | FALSE |
| LOC107065371 | cytochrome P450 6B5-like | 0.03521 | 32.82875 | FALSE | FALSE |
| LOC107072236 | probable uridine nucleosidase 2 | 0.03585 | 32.83057 | FALSE | FALSE |
| LOC107070591 | neutral alpha-glucosidase AB | 0.0349 | 32.83171 | FALSE | FALSE |
| LOC107072990 | RRP12-like protein | 0.03597 | 32.83405 | FALSE | FALSE |
| LOC107072993 | parafibromin | 0.03571 | 32.90204 | FALSE | FALSE |
| LOC107063779 | uncharacterized protein LOC107063779 | 0.03578 | 32.92759 | FALSE | FALSE |
| LOC107066319 | protein bride of sevenless | 0.03466 | 32.94344 | FALSE | FALSE |
| LOC107068235 | uncharacterized protein LOC107068235 | 0.03682 | 32.94733 | FALSE | FALSE |
| LOC107065815 | lysosome membrane protein 2-like | 0.03635 | 32.95514 | FALSE | FALSE |
| LOC107064155 | phosphomevalonate kinase | 0.0358 | 32.95525 | FALSE | FALSE |
| LOC107069675 | synembryn | 0.03547 | 32.96665 | FALSE | FALSE |
| LOC107069708 | general odorant-binding protein 72-like | 0.03669 | 32.9701 | FALSE | FALSE |
| LOC107065590 | protein FAM45B-like | 0.035 | 32.99946 | FALSE | FALSE |
| LOC107069296 | RNA polymerase II-associated protein 1 | 0.03463 | 33.01595 | FALSE | FALSE |
| LOC107070703 | serine protease nudel-like | 0.03547 | 33.01713 | FALSE | FALSE |

| | | | | | |
|---|---|---|---|---|---|
| LOC107069595 | TSC22 domain family protein 1 | 0.03495 | 33.03268 | FALSE | FALSE |
| LOC107072316 | sodium- and chloride-dependent GABA tran | 0.03571 | 33.04149 | FALSE | FALSE |
| LOC107066679 | ranBP-type and C3HC4-type zinc finger-cont | 0.03467 | 33.05298 | FALSE | FALSE |
| LOC107071533 | armadillo repeat-containing protein 6 homo | 0.03427 | 33.05737 | FALSE | FALSE |
| LOC107072047 | protein kinase DC2 | 0.0341 | 33.05953 | FALSE | FALSE |
| LOC107074646 | ankycorbin | 0.03552 | 33.06007 | FALSE | FALSE |
| LOC107065991 | ras-related protein Rab-24-like | 0.03487 | 33.08046 | FALSE | FALSE |
| LOC107071057 | NADH dehydrogenase [ubiquinone | 0.03663 | 33.08388 | FALSE | FALSE |
| LOC107070281 | coiled-coil domain-containing protein 86 | 0.03618 | 33.08609 | FALSE | FALSE |
| LOC107068635 | pre-mRNA-splicing factor SYF1 | 0.03538 | 33.08647 | FALSE | FALSE |
| LOC107065251 | uncharacterized protein LOC107065251 | 0.03513 | 33.08905 | FALSE | FALSE |
| LOC107071375 | uncharacterized protein LOC107071375 | 0.0363 | 33.09949 | FALSE | FALSE |
| LOC107071260 | pickpocket protein 28-like | 0.03538 | 33.12222 | FALSE | FALSE |
| LOC107072520 | hornerin-like | 0.03588 | 33.12394 | FALSE | FALSE |
| LOC107063798 | eukaryotic translation initiation factor 2-alp | 0.03588 | 33.12651 | FALSE | FALSE |
| LOC107070760 | gamma-aminobutyric acid type B receptor s | 0.03573 | 33.13086 | FALSE | FALSE |
| LOC107073942 | DDB1- and CUL4-associated factor 13 | 0.03602 | 33.1397 | FALSE | FALSE |
| LOC107065293 | protein Mo25 | 0.0353 | 33.14526 | FALSE | FALSE |
| LOC107064495 | AKT-interacting protein-like | 0.03573 | 33.14911 | FALSE | FALSE |
| LOC107072536 | protein ECT2 | 0.03631 | 33.15317 | FALSE | FALSE |
| LOC107068523 | ninjurin-1-like | 0.03791 | 33.15804 | FALSE | FALSE |
| LOC107065606 | solute carrier family 2, facilitated glucose tra | 0.03497 | 33.17194 | FALSE | FALSE |
| LOC107068090 | adenine phosphoribosyltransferase | 0.03491 | 33.18137 | FALSE | FALSE |
| LOC107073962 | uncharacterized protein LOC107073962 | 0.03476 | 33.1854 | FALSE | FALSE |
| LOC107063985 | proteasome assembly chaperone 2 | 0.03614 | 33.18584 | FALSE | FALSE |
| LOC107073817 | myosin-11 | 0.03551 | 33.18903 | FALSE | FALSE |
| LOC107074212 | tubulin-folding cofactor B | 0.03464 | 33.19246 | FALSE | FALSE |
| LOC107066787 | odorant receptor 13a-like | 0.03515 | 33.21573 | FALSE | FALSE |
| LOC107071848 | prominin-like protein | 0.03671 | 33.22577 | FALSE | FALSE |
| LOC107073751 | uncharacterized protein LOC107073751 | 0.03516 | 33.23166 | FALSE | FALSE |
| LOC107066778 | ras-related protein Rab-23-like | 0.03503 | 33.23511 | FALSE | FALSE |
| LOC107071521 | semaphorin-1A-like | 0.0353 | 33.24348 | FALSE | FALSE |
| LOC107065540 | putative glutathione-specific gamma-glutam | 0.03665 | 33.24579 | FALSE | FALSE |
| LOC107063632 | uncharacterized protein LOC107063632 | 0.03579 | 33.25342 | FALSE | FALSE |
| LOC107074042 | LOW QUALITY PROTEIN: neuronal acetylcho | 0.03652 | 33.26901 | FALSE | FALSE |
| LOC107065073 | nucleolin-like | 0.03519 | 33.2696 | FALSE | FALSE |
| LOC107070829 | alpha-1,2-mannosyltransferase ALG9 | 0.03573 | 33.2737 | FALSE | FALSE |
| LOC107073402 | DNA polymerase delta subunit 2-like | 0.03613 | 33.28923 | FALSE | FALSE |
| LOC107065642 | uncharacterized protein LOC107065642 | 0.03524 | 33.3138 | FALSE | FALSE |
| LOC107066267 | DNA polymerase alpha catalytic subunit-like | 0.03408 | 33.33174 | FALSE | FALSE |
| LOC107066948 | zinc transporter 1 | 0.03809 | 33.33507 | FALSE | FALSE |
| LOC107069640 | DNA polymerase theta | 0.0357 | 33.34031 | FALSE | FALSE |
| LOC107066481 | uncharacterized protein C18orf63-like | 0.03472 | 33.34625 | FALSE | FALSE |
| LOC107074339 | uncharacterized protein LOC107074339 | 0.0351 | 33.36248 | FALSE | FALSE |
| LOC107072719 | neuroglian | 0.03482 | 33.37141 | FALSE | FALSE |
| LOC107065097 | protein FAM136A-like | 0.03529 | 33.38108 | FALSE | FALSE |
| LOC107065744 | erlin-1-like | 0.03504 | 33.39423 | FALSE | FALSE |
| LOC107072631 | uncharacterized protein LOC107072631 | 0.03547 | 33.40937 | FALSE | FALSE |
| LOC107065022 | CCAAT/enhancer-binding protein zeta | 0.03665 | 33.41483 | FALSE | FALSE |
| LOC107068361 | uncharacterized protein LOC107068361 | 0.03489 | 33.42646 | FALSE | FALSE |

| | | | | | |
|---|---|---|---|---|---|
| LOC107073587 | uncharacterized protein LOC107073587 | 0.0351 | 33.43095 | FALSE | FALSE |
| LOC107071723 | C-Maf-inducing protein-like | 0.03539 | 33.43636 | FALSE | FALSE |
| LOC107065999 | V-type proton ATPase subunit C | 0.03505 | 33.44565 | FALSE | FALSE |
| LOC107067332 | DNA replication ATP-dependent helicase/nu | 0.03659 | 33.44802 | FALSE | FALSE |
| LOC107074293 | inward rectifier potassium channel 2-like | 0.03503 | 33.46416 | FALSE | FALSE |
| LOC107072134 | uncharacterized protein LOC107072134 | 0.0365 | 33.48046 | FALSE | FALSE |
| LOC107071618 | SWI/SNF-related matrix-associated actin-de | 0.03516 | 33.4867 | FALSE | FALSE |
| LOC107067941 | UPF0562 protein C7orf55-like | 0.03568 | 33.49704 | FALSE | FALSE |
| LOC107065269 | iron-sulfur protein NUBPL | 0.03645 | 33.51921 | FALSE | FALSE |
| LOC107071437 | uncharacterized protein LOC107071437 | 0.03694 | 33.51994 | FALSE | FALSE |
| LOC107067953 | uncharacterized protein LOC107067953 | 0.03425 | 33.52287 | FALSE | FALSE |
| LOC107063938 | synaptojanin-1 | 0.03484 | 33.52358 | FALSE | FALSE |
| LOC107068463 | nuclear factor of activated T-cells 5-like | 0.03529 | 33.54225 | FALSE | FALSE |
| LOC107070193 | protein-S-isoprenylcysteine O-methyltransfe | 0.03545 | 33.56829 | FALSE | FALSE |
| LOC107068918 | protein D2-like | 0.0347 | 33.58001 | FALSE | FALSE |
| LOC107066838 | tyrosine-protein kinase Src42A | 0.03403 | 33.58291 | FALSE | FALSE |
| LOC107073245 | glycoprotein 3-alpha-L-fucosyltransferase A | 0.03533 | 33.58348 | FALSE | FALSE |
| LOC107068807 | cytosolic non-specific dipeptidase | 0.03643 | 33.61665 | FALSE | FALSE |
| LOC107073925 | sphingomyelin phosphodiesterase 1-like | 0.03487 | 33.6187 | FALSE | FALSE |
| LOC107069257 | proton-coupled folate transporter-like | 0.03652 | 33.62308 | FALSE | FALSE |
| LOC107071791 | protein TIPIN homolog | 0.03597 | 33.62881 | FALSE | FALSE |
| LOC107066869 | dehydrogenase/reductase SDR family meml | 0.03497 | 33.63132 | FALSE | FALSE |
| LOC107067443 | uncharacterized protein LOC107067443 | 0.03529 | 33.64088 | FALSE | FALSE |
| LOC107072116 | calumenin | 0.03442 | 33.64089 | FALSE | FALSE |
| LOC107064332 | uncharacterized protein LOC107064332 | 0.03681 | 33.64487 | FALSE | FALSE |
| LOC107067935 | cytochrome P450 4C1-like | 0.03348 | 33.65348 | FALSE | FALSE |
| LOC107066418 | prefoldin subunit 3 | 0.03665 | 33.66453 | FALSE | FALSE |
| LOC107064708 | uncharacterized protein C17orf59 homolog | 0.03584 | 33.68176 | FALSE | FALSE |
| LOC107072039 | bone morphogenetic protein 4 | 0.03515 | 33.68899 | FALSE | FALSE |
| LOC107067042 | uncharacterized protein LOC107067042 | 0.0361 | 33.70445 | FALSE | FALSE |
| LOC107070068 | adenylyl cyclase-associated protein 1 | 0.03505 | 33.70527 | FALSE | FALSE |
| LOC107066477 | zinc finger CCHC domain-containing protein | 0.03497 | 33.71228 | FALSE | FALSE |
| LOC107073212 | protein kinase C-binding protein 1 | 0.03511 | 33.73772 | FALSE | FALSE |
| LOC107067238 | probable phospholipid-transporting ATPase | 0.0345 | 33.73812 | FALSE | FALSE |
| LOC107066343 | uncharacterized protein LOC107066343 | 0.03519 | 33.73874 | FALSE | FALSE |
| LOC107065563 | uncharacterized protein LOC107065563 | 0.03604 | 33.74072 | FALSE | FALSE |
| LOC107067837 | solute carrier family 25 member 44 | 0.03654 | 33.74563 | FALSE | FALSE |
| LOC107067539 | chromatin accessibility complex protein 1 | 0.0351 | 33.74585 | FALSE | FALSE |
| LOC107068380 | uncharacterized protein LOC107068380 | 0.03454 | 33.75749 | FALSE | FALSE |
| LOC107070076 | nucleoside diphosphate kinase 7 | 0.0357 | 33.76664 | FALSE | FALSE |
| LOC107074228 | uncharacterized protein LOC107074228 | 0.03537 | 33.77375 | FALSE | FALSE |
| LOC107074576 | WD repeat-containing protein 63-like | 0.03593 | 33.78156 | FALSE | FALSE |
| LOC107070632 | probable cytosolic Fe-S cluster assembly fac | 0.03457 | 33.79268 | FALSE | FALSE |
| LOC107070917 | uncharacterized protein C15orf61 homolog | 0.03573 | 33.80176 | FALSE | FALSE |
| LOC107071237 | neuronal acetylcholine receptor subunit alp | 0.03554 | 33.80629 | FALSE | FALSE |
| LOC107074216 | clavesin-1-like | 0.03443 | 33.80944 | FALSE | FALSE |
| LOC107068520 | uncharacterized protein LOC107068520 | 0.03611 | 33.82099 | FALSE | FALSE |
| LOC107067117 | DEP domain-containing protein 5 | 0.03618 | 33.82737 | FALSE | FALSE |
| LOC107068431 | uncharacterized protein LOC107068431 | 0.03567 | 33.82955 | FALSE | FALSE |
| LOC107067490 | uncharacterized protein LOC107067490 | 0.03622 | 33.84415 | FALSE | FALSE |

| | | | | | |
|---|---|---|---|---|---|
| LOC107070588 | peptidyl-tRNA hydrolase ICT1, mitochondria | 0.03525 | 33.85006 | FALSE | FALSE |
| LOC107065273 | CCR4-NOT transcription complex subunit 3 | 0.03528 | 33.85161 | FALSE | FALSE |
| LOC107067922 | mucin-17 | 0.03517 | 33.85808 | FALSE | FALSE |
| LOC107066745 | uncharacterized protein LOC107066745 | 0.03604 | 33.88094 | FALSE | FALSE |
| LOC107074468 | 28S ribosomal protein S30, mitochondrial | 0.0348 | 33.88107 | FALSE | FALSE |
| LOC107066028 | tropomodulin | 0.03589 | 33.88133 | FALSE | FALSE |
| LOC107067618 | protein sister of odd and bowel | 0.03603 | 33.90519 | FALSE | FALSE |
| LOC107067142 | lanC-like protein 2 | 0.03603 | 33.90656 | FALSE | FALSE |
| LOC107068570 | bifunctional 3'-phosphoadenosine 5'-phospl | 0.03626 | 33.9125 | FALSE | FALSE |
| LOC107067581 | protein yellow-like | 0.03432 | 33.91593 | FALSE | FALSE |
| LOC107072206 | serine proteinase stubble-like | 0.03544 | 33.91914 | FALSE | FALSE |
| LOC107066533 | methyltransferase-like protein 9 | 0.03404 | 33.92517 | FALSE | FALSE |
| LOC107064570 | protein outspread | 0.03682 | 33.94047 | FALSE | FALSE |
| LOC107074602 | flavin reductase (NADPH) | 0.03484 | 33.94456 | FALSE | FALSE |
| LOC107068825 | ras-related protein Rab-43 | 0.03584 | 33.95989 | FALSE | FALSE |
| LOC107071728 | protein split ends | 0.03656 | 33.97166 | FALSE | FALSE |
| LOC107068303 | E3 ubiquitin-protein ligase RNF4-like | 0.03507 | 33.97506 | FALSE | FALSE |
| LOC107066005 | uncharacterized protein LOC107066005 | 0.03513 | 33.98347 | FALSE | FALSE |
| LOC107065136 | cysteine-rich motor neuron 1 protein-like | 0.03519 | 34.01242 | FALSE | FALSE |
| LOC107070184 | serine/threonine-protein kinase OSR1-like | 0.0339 | 34.02142 | FALSE | FALSE |
| LOC107069445 | sorting nexin-8-like | 0.03623 | 34.02872 | FALSE | FALSE |
| LOC107071175 | cap-specific mRNA (nucleoside-2'-O-)-methy | 0.03482 | 34.04552 | FALSE | FALSE |
| LOC107072429 | uncharacterized protein LOC107072429 | 0.03426 | 34.04819 | FALSE | FALSE |
| LOC107074093 | elongation of very long chain fatty acids pro | 0.03506 | 34.05564 | FALSE | FALSE |
| LOC107071939 | Bloom syndrome protein homolog | 0.03536 | 34.05614 | FALSE | FALSE |
| LOC107065098 | inositol-trisphosphate 3-kinase A | 0.03447 | 34.05765 | FALSE | FALSE |
| LOC107064982 | serine/threonine-protein kinase Genghis Kh | 0.03521 | 34.06033 | FALSE | FALSE |
| LOC107067001 | thioredoxin reductase 2, mitochondrial | 0.03522 | 34.07393 | FALSE | FALSE |
| LOC107063682 | protein tramtrack, alpha isoform | 0.03458 | 34.07942 | FALSE | FALSE |
| LOC107067244 | ETS-related transcription factor Elf-5-like | 0.03496 | 34.08475 | FALSE | FALSE |
| LOC107069559 | non-histone protein 10 | 0.03536 | 34.09193 | FALSE | FALSE |
| LOC107068103 | ly6/PLAUR domain-containing protein 6B-lik | 0.03553 | 34.10355 | FALSE | FALSE |
| LOC107067217 | contactin-2 | 0.03433 | 34.11827 | FALSE | FALSE |
| LOC107068099 | uncharacterized protein LOC107068099 | 0.03621 | 34.13831 | FALSE | FALSE |
| LOC107074572 | protein henna | 0.03528 | 34.1389 | FALSE | FALSE |
| LOC107067963 | splicing factor U2AF 50 kDa subunit | 0.03556 | 34.14374 | FALSE | FALSE |
| LOC107066714 | Bardet-Biedl syndrome 7 protein homolog | 0.03549 | 34.14903 | FALSE | FALSE |
| LOC107069818 | putative oxidoreductase GLYR1 homolog | 0.03557 | 34.17892 | FALSE | FALSE |
| LOC107067367 | acyl-CoA synthetase short-chain family men | 0.03668 | 34.18755 | FALSE | FALSE |
| LOC107066377 | ankyrin repeat and death domain-containin | 0.03468 | 34.18897 | FALSE | FALSE |
| LOC107069499 | uncharacterized protein LOC107069499 | 0.03455 | 34.20449 | FALSE | FALSE |
| LOC107073876 | probable chitinase 3 | 0.03525 | 34.22607 | FALSE | FALSE |
| LOC107067318 | alpha-mannosidase 2 | 0.03445 | 34.22619 | FALSE | FALSE |
| LOC107072230 | 60S ribosomal protein L36 | 0.03554 | 34.23366 | FALSE | FALSE |
| LOC107072021 | uncharacterized protein LOC107072021 | 0.03652 | 34.27474 | FALSE | FALSE |
| LOC107074229 | muscle segmentation homeobox-like | 0.03499 | 34.28125 | FALSE | FALSE |
| LOC107065597 | nucleolin 2-like | 0.03451 | 34.30448 | FALSE | FALSE |
| LOC107067168 | partitioning defective protein 6 | 0.03448 | 34.31015 | FALSE | FALSE |
| LOC107071118 | ero1-like protein | 0.03592 | 34.31527 | FALSE | FALSE |
| LOC107070762 | glutamyl-tRNA(Gln) amidotransferase subur | 0.03477 | 34.31592 | FALSE | FALSE |

| | | | | | |
|---|---|---|---|---|---|
| LOC107068177 | protein AAR2 homolog | 0.03427 | 34.324 | FALSE | FALSE |
| LOC107072946 | uncharacterized protein LOC107072946 | 0.03459 | 34.32887 | FALSE | FALSE |
| LOC107064673 | EKC/KEOPS complex subunit LAGE3 | 0.03507 | 34.33327 | FALSE | FALSE |
| LOC107064060 | general transcription factor IIF subunit 2 | 0.03596 | 34.34159 | FALSE | FALSE |
| LOC107072795 | F-box only protein 44-like | 0.0348 | 34.36074 | FALSE | FALSE |
| LOC107066119 | NADH-quinone oxidoreductase subunit B 2- | 0.03334 | 34.36929 | FALSE | FALSE |
| LOC107064876 | protein LTV1 homolog | 0.03573 | 34.37933 | FALSE | FALSE |
| LOC107070010 | zinc transporter ZIP3 | 0.03627 | 34.38896 | FALSE | FALSE |
| LOC107065074 | methyltransferase-like protein 6 | 0.03559 | 34.39182 | FALSE | FALSE |
| LOC107069426 | gamma-aminobutyric acid receptor subunit | 0.03594 | 34.39667 | FALSE | FALSE |
| LOC107064746 | 39S ribosomal protein L39, mitochondrial | 0.0357 | 34.39869 | FALSE | FALSE |
| LOC107069578 | uncharacterized protein LOC107069578 | 0.03506 | 34.40956 | FALSE | FALSE |
| LOC107070915 | protein N-terminal asparagine amidohydrol | 0.03441 | 34.41797 | FALSE | FALSE |
| LOC107065436 | cilia- and flagella-associated protein 20-like | 0.03551 | 34.41947 | FALSE | FALSE |
| LOC107074405 | uncharacterized protein LOC107074405 | 0.03528 | 34.42586 | FALSE | FALSE |
| LOC107072760 | probable ATP-dependent RNA helicase ddx2 | 0.03533 | 34.43002 | FALSE | FALSE |
| LOC107072835 | uncharacterized protein LOC107072835 | 0.03536 | 34.45046 | FALSE | FALSE |
| LOC107067816 | actin, muscle | 0.03592 | 34.45316 | FALSE | FALSE |
| LOC107071205 | uncharacterized protein LOC107071205 | 0.0354 | 34.45547 | FALSE | FALSE |
| LOC107073668 | uncharacterized protein LOC107073668 | 0.03507 | 34.47104 | FALSE | FALSE |
| LOC107070572 | N-acylneuraminate-9-phosphatase | 0.03544 | 34.4865 | FALSE | FALSE |
| LOC107065372 | phosphatidylinositol 4-phosphate 3-kinase ( | 0.0356 | 34.49029 | FALSE | FALSE |
| LOC107074174 | transmembrane and TPR repeat-containing | 0.03489 | 34.50417 | FALSE | FALSE |
| LOC107063917 | uncharacterized protein LOC107063917 | 0.03571 | 34.51424 | FALSE | FALSE |
| LOC107066049 | 4-coumarate--CoA ligase 1-like | 0.0346 | 34.51998 | FALSE | FALSE |
| LOC107064136 | collagen alpha-1(XVIII) chain | 0.03467 | 34.52526 | FALSE | FALSE |
| LOC107067972 | RNA-binding protein 26 | 0.03501 | 34.5329 | FALSE | FALSE |
| LOC107066419 | C-Myc-binding protein | 0.03449 | 34.5361 | FALSE | FALSE |
| LOC107063772 | T-complex protein 1 subunit gamma | 0.03597 | 34.53757 | FALSE | FALSE |
| LOC107070447 | translation initiation factor eIF-2B subunit b | 0.03436 | 34.55296 | FALSE | FALSE |
| LOC107064023 | homeobox protein prophet of Pit-1 | 0.03552 | 34.55883 | FALSE | FALSE |
| LOC107070144 | probable serine/threonine-protein kinase D | 0.03533 | 34.55973 | FALSE | FALSE |
| LOC107065918 | uncharacterized protein KIAA0556-like | 0.03531 | 34.57732 | FALSE | FALSE |
| LOC107072128 | uncharacterized protein LOC107072128 | 0.03431 | 34.5798 | FALSE | FALSE |
| LOC107068404 | 2-amino-3-ketobutyrate coenzyme A ligase, | 0.03441 | 34.5818 | FALSE | FALSE |
| LOC107068465 | DET1 homolog | 0.03436 | 34.59842 | FALSE | FALSE |
| LOC107071948 | lysine--tRNA ligase | 0.03369 | 34.60019 | FALSE | FALSE |
| LOC107067943 | cytochrome P450 4C1-like | 0.03539 | 34.60142 | FALSE | FALSE |
| LOC107073060 | uncharacterized protein LOC107073060 | 0.03449 | 34.60778 | FALSE | FALSE |
| LOC107068504 | nitric oxide-associated protein 1 | 0.03547 | 34.62235 | FALSE | FALSE |
| LOC107064769 | uncharacterized protein LOC107064769 | 0.03468 | 34.62676 | FALSE | FALSE |
| LOC107068192 | heat shock factor 2-binding protein-like | 0.03437 | 34.62937 | FALSE | FALSE |
| LOC107074159 | myosin regulatory light chain LC-2, mantle n | 0.03385 | 34.63033 | FALSE | FALSE |
| LOC107071026 | glycosylated lysosomal membrane protein A | 0.03477 | 34.63855 | FALSE | FALSE |
| LOC107063908 | mannose-1-phosphate guanyltransferase be | 0.03504 | 34.6399 | FALSE | FALSE |
| LOC107066232 | tyrosine-protein kinase-like otk | 0.03473 | 34.64218 | FALSE | FALSE |
| LOC107070788 | uncharacterized protein LOC107070788 | 0.03484 | 34.65625 | FALSE | FALSE |
| LOC107068826 | uncharacterized protein LOC107068826 | 0.03445 | 34.65678 | FALSE | FALSE |
| LOC107073473 | cytochrome P450 4C1-like | 0.03407 | 34.65997 | FALSE | FALSE |
| LOC107064825 | protein DPCD | 0.03428 | 34.66175 | FALSE | FALSE |

| | | | | | |
|---|---|---|---|---|---|
| LOC107069656 | transmembrane protein 179 | 0.03426 | 34.66449 | FALSE | FALSE |
| LOC107073152 | uncharacterized protein LOC107073152 | 0.03516 | 34.67981 | FALSE | FALSE |
| LOC107065985 | mitochondrial inner membrane protease su | 0.03647 | 34.68445 | FALSE | FALSE |
| LOC107066142 | sodium-dependent nutrient amino acid tran | 0.03469 | 34.68966 | FALSE | FALSE |
| LOC107070199 | INO80 complex subunit E | 0.03529 | 34.6908 | FALSE | FALSE |
| LOC107072407 | uncharacterized protein LOC107072407 | 0.03428 | 34.70036 | FALSE | FALSE |
| LOC107072393 | sister chromatid cohesion protein PDS5 hon | 0.03411 | 34.73583 | FALSE | FALSE |
| LOC107073631 | uncharacterized protein LOC107073631 | 0.03434 | 34.76211 | FALSE | FALSE |
| LOC107073338 | ADP-ribosylation factor-like protein 4C | 0.03543 | 34.76348 | FALSE | FALSE |
| LOC107071541 | uncharacterized protein LOC107071541 | 0.03451 | 34.7855 | FALSE | FALSE |
| LOC107067041 | uncharacterized protein LOC107067041 | 0.03499 | 34.79631 | FALSE | FALSE |
| LOC107063987 | DNA polymerase alpha subunit B | 0.0345 | 34.80931 | FALSE | FALSE |
| LOC107064672 | phosphatidylinositol N-acetylglucosaminyltr | 0.03435 | 34.82355 | FALSE | FALSE |
| LOC107074515 | thioredoxin domain-containing protein 17-li | 0.03499 | 34.83143 | FALSE | FALSE |
| LOC107074686 | pancreatic triacylglycerol lipase-like | 0.03492 | 34.84324 | FALSE | FALSE |
| LOC107074503 | triokinase/FMN cyclase-like | 0.03621 | 34.86052 | FALSE | FALSE |
| LOC107071471 | uncharacterized protein LOC107071471 | 0.03538 | 34.86665 | FALSE | FALSE |
| LOC107065644 | long-chain fatty acid transport protein 4-like | 0.03517 | 34.87723 | FALSE | FALSE |
| LOC107072769 | uncharacterized protein LOC107072769 | 0.03492 | 34.89096 | FALSE | FALSE |
| LOC107072596 | facilitated trehalose transporter Tret1-like | 0.03505 | 34.91125 | FALSE | FALSE |
| LOC107067304 | uncharacterized protein LOC107067304 | 0.03441 | 34.91555 | FALSE | FALSE |
| LOC107063832 | enhancer of split mgamma protein-like | 0.03454 | 34.92191 | FALSE | FALSE |
| LOC107064472 | PX domain-containing protein kinase-like pr | 0.03429 | 34.93037 | FALSE | FALSE |
| LOC107071062 | proteasome assembly chaperone 4-like | 0.03444 | 34.94189 | FALSE | FALSE |
| LOC107071086 | HD domain-containing protein 2 | 0.0349 | 34.9495 | FALSE | FALSE |
| LOC107065495 | probable nuclear transport factor 2 | 0.03492 | 34.95002 | FALSE | FALSE |
| LOC107067495 | GATA zinc finger domain-containing protein | 0.03636 | 34.95864 | FALSE | FALSE |
| LOC107063997 | nephrin-like | 0.03484 | 34.96496 | FALSE | FALSE |
| LOC107068824 | anoctamin-4 | 0.03359 | 34.97035 | FALSE | FALSE |
| LOC107072621 | GPI mannosyltransferase 4 | 0.03542 | 34.9868 | FALSE | FALSE |
| LOC107073800 | hemicentin-2-like | 0.03446 | 34.98818 | FALSE | FALSE |
| LOC107064516 | chitobiosyldiphosphodolichol beta-mannosy | 0.03423 | 34.99491 | FALSE | FALSE |
| LOC107071498 | serine-rich adhesin for platelets | 0.03539 | 34.99618 | FALSE | FALSE |
| LOC107074368 | fibrillin-2-like | 0.03524 | 35.00537 | FALSE | FALSE |
| LOC107065166 | uncharacterized protein LOC107065166 | 0.03435 | 35.0125 | FALSE | FALSE |
| LOC107070522 | SH2B adapter protein 1-like | 0.03448 | 35.01519 | FALSE | FALSE |
| LOC107066559 | alpha-(1,6)-fucosyltransferase | 0.0344 | 35.02591 | FALSE | FALSE |
| LOC107072373 | putative uncharacterized protein DDB_G027 | 0.03481 | 35.02937 | FALSE | FALSE |
| LOC107066270 | transcriptional adapter 3-A | 0.03454 | 35.04144 | FALSE | FALSE |
| LOC107067350 | ATP-binding cassette sub-family G member | 0.03389 | 35.04335 | FALSE | FALSE |
| LOC107069007 | kynurenine formamidase | 0.03402 | 35.05562 | FALSE | FALSE |
| LOC107069984 | UPF0483 protein CG5412 | 0.03595 | 35.06007 | FALSE | FALSE |
| LOC107064239 | transmembrane protein 208 | 0.03406 | 35.06072 | FALSE | FALSE |
| LOC107064927 | uncharacterized serine-rich protein C215.13 | 0.0337 | 35.06624 | FALSE | FALSE |
| LOC107065605 | serine/threonine-protein kinase pakD-like | 0.03435 | 35.07647 | FALSE | FALSE |
| LOC107066597 | X-ray repair cross-complementing protein 5 | 0.03463 | 35.10635 | FALSE | FALSE |
| LOC107065752 | mitogen-activated protein kinase 1 | 0.03414 | 35.12777 | FALSE | FALSE |
| LOC107074259 | uncharacterized protein LOC107074259 | 0.03488 | 35.12797 | FALSE | FALSE |
| LOC107074292 | uncharacterized protein LOC107074292 | 0.03418 | 35.13259 | FALSE | FALSE |
| LOC107065422 | sideroflexin-2 | 0.03317 | 35.15854 | FALSE | FALSE |

| | | | | | |
|---|---|---|---|---|---|
| LOC107066978 | leukotriene A-4 hydrolase | 0.03433 | 35.16546 | FALSE | FALSE |
| LOC107071956 | long-chain-fatty-acid--CoA ligase 5 | 0.0338 | 35.16609 | FALSE | FALSE |
| LOC107064789 | CD109 antigen | 0.03635 | 35.17317 | FALSE | FALSE |
| LOC107065405 | kinesin heavy chain | 0.03456 | 35.18163 | FALSE | FALSE |
| LOC107070086 | ras-related C3 botulinum toxin substrate 1 | 0.03524 | 35.18696 | FALSE | FALSE |
| LOC107073477 | phenylalanine--tRNA ligase beta subunit | 0.03454 | 35.19729 | FALSE | FALSE |
| LOC107069784 | small subunit processome component 20 ho | 0.03378 | 35.19981 | FALSE | FALSE |
| LOC107070098 | rho GTPase-activating protein 1 | 0.0336 | 35.20404 | FALSE | FALSE |
| LOC107067432 | nuclear receptor-binding protein homolog | 0.03596 | 35.21394 | FALSE | FALSE |
| LOC107072826 | uncharacterized protein LOC107072826 | 0.0355 | 35.22735 | FALSE | FALSE |
| LOC107066030 | methylthioribose-1-phosphate isomerase | 0.03514 | 35.22864 | FALSE | FALSE |
| LOC107069332 | uncharacterized protein LOC107069332 | 0.03438 | 35.23935 | FALSE | FALSE |
| LOC107065411 | conserved oligomeric Golgi complex subuni | 0.03343 | 35.24483 | FALSE | FALSE |
| LOC107067795 | proto-oncogene c-Fos | 0.03568 | 35.2482 | FALSE | FALSE |
| LOC107068073 | putative ammonium transporter 3 | 0.03526 | 35.25374 | FALSE | FALSE |
| LOC107067811 | retinal guanylyl cyclase 2-like | 0.035 | 35.25777 | FALSE | FALSE |
| LOC107073034 | LIRP-like | 0.03542 | 35.28766 | FALSE | FALSE |
| LOC107065784 | LOW QUALITY PROTEIN: insulin-like growth | 0.0339 | 35.30129 | FALSE | FALSE |
| LOC107071638 | uncharacterized protein LOC107071638 | 0.03512 | 35.3102 | FALSE | FALSE |
| LOC107063670 | inositol hexakisphosphate kinase 1 | 0.03475 | 35.32325 | FALSE | FALSE |
| LOC107068608 | 60S acidic ribosomal protein P1 | 0.03453 | 35.32384 | FALSE | FALSE |
| LOC107069013 | vesicle-associated membrane protein 7 | 0.03502 | 35.32432 | FALSE | FALSE |
| LOC107066808 | disintegrin and metalloproteinase domain-c | 0.0348 | 35.32966 | FALSE | FALSE |
| LOC107068017 | protein msta, isoform B | 0.03365 | 35.33568 | FALSE | FALSE |
| LOC107068789 | endoplasmic reticulum lectin 1 | 0.03446 | 35.34139 | FALSE | FALSE |
| LOC107069902 | high affinity choline transporter 1-like | 0.03443 | 35.3478 | FALSE | FALSE |
| LOC107064942 | uncharacterized protein LOC107064942 | 0.03444 | 35.37659 | FALSE | FALSE |
| LOC107064315 | organic cation transporter protein-like | 0.03523 | 35.38271 | FALSE | FALSE |
| LOC107070557 | hyccin | 0.03507 | 35.3828 | FALSE | FALSE |
| LOC107068629 | F-box only protein 42 | 0.03472 | 35.38638 | FALSE | FALSE |
| LOC107063910 | cullin-3 | 0.03504 | 35.38834 | FALSE | FALSE |
| LOC107070664 | JNK-interacting protein 3 | 0.03544 | 35.39487 | FALSE | FALSE |
| LOC107066266 | probable cytochrome P450 12a5, mitochon | 0.03424 | 35.40156 | FALSE | FALSE |
| LOC107064135 | THO complex subunit 1 | 0.03404 | 35.41085 | FALSE | FALSE |
| LOC107069489 | alkyldihydroxyacetonephosphate synthase | 0.03387 | 35.41966 | FALSE | FALSE |
| LOC107066197 | 60S ribosomal protein L39 | 0.03374 | 35.42852 | FALSE | FALSE |
| LOC107074272 | uncharacterized protein LOC107074272 | 0.03472 | 35.4322 | FALSE | FALSE |
| LOC107070550 | uncharacterized protein LOC107070550 | 0.03379 | 35.43913 | FALSE | FALSE |
| LOC107071549 | neuronal PAS domain-containing protein 4 | 0.03423 | 35.46101 | FALSE | FALSE |
| LOC107068600 | reticulon-4 | 0.03333 | 35.46121 | FALSE | FALSE |
| LOC107071679 | endoplasmic reticulum aminopeptidase 2-lil | 0.0352 | 35.47313 | FALSE | FALSE |
| LOC107067234 | myb-like protein D | 0.03414 | 35.495 | FALSE | FALSE |
| LOC107070633 | dynein light chain 4, axonemal-like | 0.03465 | 35.4972 | FALSE | FALSE |
| LOC107071389 | 40S ribosomal protein S16-like | 0.03394 | 35.50854 | FALSE | FALSE |
| LOC107064144 | DE-cadherin | 0.03517 | 35.52292 | FALSE | FALSE |
| LOC107063951 | protein gooseberry-neuro-like | 0.03615 | 35.5259 | FALSE | FALSE |
| LOC107067751 | aspartate--tRNA ligase, mitochondrial | 0.03411 | 35.52623 | FALSE | FALSE |
| LOC107073341 | uncharacterized protein LOC107073341 | 0.03477 | 35.52727 | FALSE | FALSE |
| LOC107066999 | HEAT repeat-containing protein 3 | 0.03559 | 35.53084 | FALSE | FALSE |
| LOC107070630 | lysocardiolipin acyltransferase 1-like | 0.0347 | 35.54214 | FALSE | FALSE |

| | | | | | |
|---|---|---|---|---|---|
| LOC107067002 | C-factor | 0.0343 | 35.55978 | FALSE | FALSE |
| LOC107069159 | AF4/FMR2 family member 4 | 0.03393 | 35.56121 | FALSE | FALSE |
| LOC107073758 | uncharacterized protein LOC107073758 | 0.0341 | 35.56535 | FALSE | FALSE |
| LOC107073220 | segmentation protein cap'n'collar | 0.03459 | 35.56712 | FALSE | FALSE |
| LOC107072983 | uncharacterized protein LOC107072983 | 0.0353 | 35.57021 | FALSE | FALSE |
| LOC107069779 | eyes absent homolog 4 | 0.0333 | 35.57511 | FALSE | FALSE |
| LOC107074692 | glucosamine-6-phosphate isomerase | 0.03553 | 35.5763 | FALSE | FALSE |
| LOC107063848 | uncharacterized protein LOC107063848 | 0.03524 | 35.57854 | FALSE | FALSE |
| LOC107066086 | SUN domain-containing protein 3-like | 0.033 | 35.59032 | FALSE | FALSE |
| LOC107069372 | farnesol dehydrogenase-like | 0.03503 | 35.59358 | FALSE | FALSE |
| LOC107072754 | uncharacterized protein LOC107072754 | 0.03444 | 35.6012 | FALSE | FALSE |
| LOC107071688 | 39S ribosomal protein L35, mitochondrial | 0.03533 | 35.61897 | FALSE | FALSE |
| LOC107069248 | nucleoside diphosphate-linked moiety X mo | 0.0354 | 35.62533 | FALSE | FALSE |
| LOC107064346 | serine/threonine-protein phosphatase 6 reg | 0.03401 | 35.62794 | FALSE | FALSE |
| LOC107069341 | mucin-19-like | 0.03233 | 35.66076 | FALSE | FALSE |
| LOC107071717 | piggyBac transposable element-derived pro | 0.03471 | 35.67169 | FALSE | FALSE |
| LOC107063737 | tropomyosin-2-like | 0.03404 | 35.67494 | FALSE | FALSE |
| LOC107070541 | arf-GAP with SH3 domain, ANK repeat and F | 0.03463 | 35.68026 | FALSE | FALSE |
| LOC107069714 | anaphase-promoting complex subunit 15 | 0.03401 | 35.70709 | FALSE | FALSE |
| LOC107073919 | dedicator of cytokinesis protein 7 | 0.03318 | 35.71271 | FALSE | FALSE |
| LOC107067193 | exosome complex component RRP40 | 0.03574 | 35.73044 | FALSE | FALSE |
| LOC107071000 | uncharacterized protein LOC107071000 | 0.03495 | 35.74574 | FALSE | FALSE |
| LOC107071515 | THAP domain-containing protein 5-like | 0.03503 | 35.76988 | FALSE | FALSE |
| LOC107071468 | uncharacterized protein LOC107071468 | 0.03511 | 35.78365 | FALSE | FALSE |
| LOC107064998 | dyslexia-associated protein KIAA0319 | 0.03526 | 35.8126 | FALSE | FALSE |
| LOC107070091 | protein TBRG4 | 0.03393 | 35.81512 | FALSE | FALSE |
| LOC107067961 | uncharacterized protein LOC107067961 | 0.03382 | 35.82477 | FALSE | FALSE |
| LOC107073468 | NADH dehydrogenase [ubiquinone | 0.03466 | 35.83241 | FALSE | FALSE |
| LOC107067094 | uncharacterized protein LOC107067094 | 0.03468 | 35.8374 | FALSE | FALSE |
| LOC107073912 | kinesin-related protein 4-like | 0.03386 | 35.86727 | FALSE | FALSE |
| LOC107070977 | balbiani ring protein 3 | 0.03319 | 35.87457 | FALSE | FALSE |
| LOC107074586 | mitochondrial inner membrane protein COX | 0.03563 | 35.87539 | FALSE | FALSE |
| LOC107069948 | transient receptor potential channel pyrexia | 0.03475 | 35.8993 | FALSE | FALSE |
| LOC107072928 | uncharacterized protein LOC107072928 | 0.03462 | 35.90243 | FALSE | FALSE |
| LOC107071039 | helix-loop-helix protein 1-like | 0.03388 | 35.90482 | FALSE | FALSE |
| LOC107068559 | odorant receptor 67c-like | 0.03429 | 35.91143 | FALSE | FALSE |
| LOC107074416 | LOW QUALITY PROTEIN: cubilin-like | 0.03402 | 35.95984 | FALSE | FALSE |
| LOC107068671 | homeotic protein proboscipedia | 0.03327 | 35.9736 | FALSE | FALSE |
| LOC107065828 | UPF0746 protein DDB_G0281095-like | 0.03306 | 35.99641 | FALSE | FALSE |
| LOC107072808 | serine/threonine-protein kinase 32A | 0.03378 | 36.00649 | FALSE | FALSE |
| LOC107070174 | uncharacterized protein LOC107070174 | 0.03472 | 36.02654 | FALSE | FALSE |
| LOC107073965 | calcium/calmodulin-dependent protein kina | 0.03306 | 36.03314 | FALSE | FALSE |
| LOC107068034 | voltage-dependent anion-selective channel- | 0.033 | 36.04454 | FALSE | FALSE |
| LOC107070560 | RNA polymerase-associated protein RTF1 hc | 0.03512 | 36.04783 | FALSE | FALSE |
| LOC107073271 | b(0,+)-type amino acid transporter 1-like | 0.03555 | 36.05601 | FALSE | FALSE |
| LOC107073834 | NADPH--cytochrome P450 reductase | 0.03541 | 36.05958 | FALSE | FALSE |
| LOC107067628 | putative ATPase N2B | 0.03394 | 36.07684 | FALSE | FALSE |
| LOC107067488 | LIX1-like protein | 0.03546 | 36.09041 | FALSE | FALSE |
| LOC107064406 | tyrosine-protein kinase PR2-like | 0.03365 | 36.09993 | FALSE | FALSE |
| LOC107066344 | neuroendocrine convertase 1-like | 0.03304 | 36.10494 | FALSE | FALSE |

| | | | | | |
|---|---|---|---|---|---|
| LOC107072845 | lipoma HMGIC fusion partner | 0.03288 | 36.11238 | FALSE | FALSE |
| LOC107065839 | serine/threonine-protein phosphatase 2A re | 0.03502 | 36.11256 | FALSE | FALSE |
| LOC107067433 | TBC1 domain family member 20 | 0.0342 | 36.11342 | FALSE | FALSE |
| LOC107072617 | intraflagellar transport protein 57 homolog | 0.0354 | 36.1158 | FALSE | FALSE |
| LOC107066152 | uncharacterized protein LOC107066152 | 0.03517 | 36.11876 | FALSE | FALSE |
| LOC107073134 | failed axon connections | 0.03629 | 36.13084 | FALSE | FALSE |
| LOC107065503 | probable 39S ribosomal protein L45, mitoch | 0.03487 | 36.13166 | FALSE | FALSE |
| LOC107071390 | uncharacterized protein PFB0145c-like | 0.03382 | 36.13751 | FALSE | FALSE |
| LOC107068023 | transcription initiation protein SPT3 homolc | 0.03446 | 36.1401 | FALSE | FALSE |
| LOC107070425 | rab GDP dissociation inhibitor beta | 0.03397 | 36.14207 | FALSE | FALSE |
| LOC107066468 | UPF0609 protein C4orf27 | 0.0339 | 36.16592 | FALSE | FALSE |
| LOC107072923 | uncharacterized protein LOC107072923 | 0.03346 | 36.1778 | FALSE | FALSE |
| LOC107072468 | histone demethylase UTY | 0.03339 | 36.21099 | FALSE | FALSE |
| LOC107068866 | SET and MYND domain-containing protein 4 | 0.0343 | 36.21469 | FALSE | FALSE |
| LOC107066168 | putative serine protease K12H4.7 | 0.0331 | 36.21867 | FALSE | FALSE |
| LOC107064335 | bone morphogenetic protein receptor type- | 0.03426 | 36.22381 | FALSE | FALSE |
| LOC107067140 | ninjurin-1-like | 0.03413 | 36.23188 | FALSE | FALSE |
| LOC107071343 | arf-GAP with coiled-coil, ANK repeat and PH | 0.03423 | 36.23565 | FALSE | FALSE |
| LOC107071414 | arrestin domain-containing protein 3 | 0.03429 | 36.25644 | FALSE | FALSE |
| LOC107071078 | uncharacterized protein LOC107071078 | 0.03422 | 36.25862 | FALSE | FALSE |
| LOC107071222 | coatomer subunit beta' | 0.03438 | 36.2903 | FALSE | FALSE |
| LOC107068348 | ATP-binding cassette sub-family G member | 0.03455 | 36.32486 | FALSE | FALSE |
| LOC107070701 | uncharacterized protein LOC107070701 | 0.03402 | 36.34673 | FALSE | FALSE |
| LOC107065944 | 60S ribosomal protein L5 | 0.03525 | 36.35879 | FALSE | FALSE |
| LOC107067553 | uncharacterized protein LOC107067553 | 0.0337 | 36.37364 | FALSE | FALSE |
| LOC107071333 | uncharacterized protein LOC107071333 | 0.03486 | 36.38231 | FALSE | FALSE |
| LOC107065443 | cAMP-dependent protein kinase type I regu | 0.03429 | 36.38369 | FALSE | FALSE |
| LOC107067970 | NEDD8-activating enzyme E1 catalytic subu | 0.03456 | 36.39406 | FALSE | FALSE |
| LOC107064216 | transcriptional regulator Myc-like | 0.03309 | 36.42368 | FALSE | FALSE |
| LOC107064057 | renin receptor | 0.03478 | 36.43678 | FALSE | FALSE |
| LOC107070952 | 39S ribosomal protein L9, mitochondrial | 0.03388 | 36.44425 | FALSE | FALSE |
| LOC107070607 | cuticle protein 7 | 0.03512 | 36.4553 | FALSE | FALSE |
| LOC107065407 | uncharacterized protein LOC107065407 | 0.03413 | 36.45959 | FALSE | FALSE |
| LOC107070443 | ribonuclease P protein subunit p14-like | 0.03253 | 36.47244 | FALSE | FALSE |
| LOC107072954 | uncharacterized protein LOC107072954 | 0.03348 | 36.48048 | FALSE | FALSE |
| LOC107068530 | transcription factor stalky-like | 0.03462 | 36.49211 | FALSE | FALSE |
| LOC107073403 | ubiquitin carboxyl-terminal hydrolase 36-lik | 0.03426 | 36.50043 | FALSE | FALSE |
| LOC107064601 | DNA helicase MCM8-like | 0.0333 | 36.50195 | FALSE | FALSE |
| LOC107064589 | tRNA (cytosine(34)-C(5))-methyltransferase | 0.03474 | 36.51346 | FALSE | FALSE |
| LOC107071937 | carnitine O-palmitoyltransferase 1, liver isof | 0.03411 | 36.53808 | FALSE | FALSE |
| LOC107071797 | U6 snRNA-associated Sm-like protein LSm6 | 0.03438 | 36.55217 | FALSE | FALSE |
| LOC107070989 | diphosphomevalonate decarboxylase | 0.0332 | 36.55727 | FALSE | FALSE |
| LOC107067459 | lethal(3)malignant brain tumor-like protein | 0.0341 | 36.56265 | FALSE | FALSE |
| LOC107068585 | mitochondrial thiamine pyrophosphate carr | 0.03489 | 36.57799 | FALSE | FALSE |
| LOC107066125 | LETM1 and EF-hand domain-containing prot | 0.03277 | 36.58899 | FALSE | FALSE |
| LOC107064715 | RING finger protein 11 | 0.03492 | 36.59136 | FALSE | FALSE |
| LOC107074397 | metabotropic glutamate receptor | 0.0334 | 36.59241 | FALSE | FALSE |
| LOC107064790 | uncharacterized protein LOC107064790 | 0.03311 | 36.60749 | FALSE | FALSE |
| LOC107073882 | transmembrane protein 184C | 0.03401 | 36.60889 | FALSE | FALSE |
| LOC107072510 | GATA zinc finger domain-containing protein | 0.03458 | 36.61028 | FALSE | FALSE |

| | | | | | |
|---|---|---|---|---|---|
| LOC107070937 | unconventional myosin-X | 0.03463 | 36.61506 | FALSE | FALSE |
| LOC107068499 | ejaculatory bulb-specific protein 3-like | 0.03406 | 36.63864 | FALSE | FALSE |
| LOC107067755 | uncharacterized protein LOC107067755 | 0.03358 | 36.65839 | FALSE | FALSE |
| LOC107071895 | probable histone-lysine N-methyltransfera | 0.03378 | 36.66138 | FALSE | FALSE |
| LOC107064217 | coiled-coil domain-containing protein 170 | 0.03376 | 36.66603 | FALSE | FALSE |
| LOC107066606 | ribose-5-phosphate isomerase | 0.03355 | 36.68244 | FALSE | FALSE |
| LOC107071801 | FAD-linked sulfhydryl oxidase ALR | 0.0335 | 36.68336 | FALSE | FALSE |
| LOC107066475 | nucleolar protein 10 | 0.03363 | 36.69743 | FALSE | FALSE |
| LOC107068475 | small integral membrane protein 8 | 0.03505 | 36.70269 | FALSE | FALSE |
| LOC107064394 | uncharacterized protein LOC107064394 | 0.03437 | 36.70767 | FALSE | FALSE |
| LOC107064146 | neurogenic locus Notch protein | 0.03386 | 36.70904 | FALSE | FALSE |
| LOC107074056 | adrenodoxin-like protein, mitochondrial | 0.03464 | 36.72229 | FALSE | FALSE |
| LOC107067897 | cytochrome P450 4C1-like | 0.03416 | 36.72599 | FALSE | FALSE |
| LOC107068762 | uncharacterized protein LOC107068762 | 0.03485 | 36.75212 | FALSE | FALSE |
| LOC107068248 | protein Daple | 0.03371 | 36.75929 | FALSE | FALSE |
| LOC107072549 | uncharacterized protein LOC107072549 | 0.03446 | 36.76524 | FALSE | FALSE |
| LOC107066257 | uncharacterized protein LOC107066257 | 0.03331 | 36.78234 | FALSE | FALSE |
| LOC107067642 | mediator of RNA polymerase II transcriptior | 0.0332 | 36.79398 | FALSE | FALSE |
| LOC107069043 | peroxidasin homolog | 0.03322 | 36.79732 | FALSE | FALSE |
| LOC107065040 | uncharacterized protein LOC107065040 | 0.03417 | 36.80002 | FALSE | FALSE |
| LOC107071660 | tryptophan--tRNA ligase, cytoplasmic | 0.03439 | 36.80456 | FALSE | FALSE |
| LOC107068212 | ankyrin repeat domain-containing protein 1 | 0.03239 | 36.8192 | FALSE | FALSE |
| LOC107066406 | ubiquitin-like protein 5 | 0.03352 | 36.8295 | FALSE | FALSE |
| LOC107066335 | 2-methoxy-6-polyprenyl-1,4-benzoquinol m | 0.03362 | 36.83157 | FALSE | FALSE |
| LOC107065492 | zinc finger protein 91 | 0.03328 | 36.84354 | FALSE | FALSE |
| LOC107068333 | eukaryotic translation initiation factor 3 sub | 0.03396 | 36.84379 | FALSE | FALSE |
| LOC107066014 | spermine oxidase-like | 0.03478 | 36.85243 | FALSE | FALSE |
| LOC107067929 | tubulin alpha-1 chain | 0.03353 | 36.85615 | FALSE | FALSE |
| LOC107067663 | F-box/LRR-repeat protein 14-like | 0.03458 | 36.86632 | FALSE | FALSE |
| LOC107064958 | GTP-binding protein Rit1-like | 0.03384 | 36.86918 | FALSE | FALSE |
| LOC107066212 | poly(ADP-ribose) glycohydrolase | 0.03388 | 36.86929 | FALSE | FALSE |
| LOC107073174 | laccase-2-like | 0.0341 | 36.87386 | FALSE | FALSE |
| LOC107064653 | YY1-associated factor 2 | 0.03385 | 36.88601 | FALSE | FALSE |
| LOC107067060 | uncharacterized protein At4g17910 | 0.03391 | 36.88755 | FALSE | FALSE |
| LOC107066785 | vacuolar protein sorting-associated protein | 0.0333 | 36.88828 | FALSE | FALSE |
| LOC107066402 | protein transport protein Sec31A | 0.03361 | 36.89275 | FALSE | FALSE |
| LOC107068434 | putative uncharacterized protein DDB_G02 | 0.03407 | 36.89629 | FALSE | FALSE |
| LOC107072086 | phospholipase A1 2-like | 0.03262 | 36.91694 | FALSE | FALSE |
| LOC107064627 | putative serine/threonine-protein kinase C | 0.03324 | 36.92366 | FALSE | FALSE |
| LOC107072656 | uncharacterized protein LOC107072656 | 0.0345 | 36.93711 | FALSE | FALSE |
| LOC107073203 | sodium-coupled monocarboxylate transpor | 0.03207 | 36.94719 | FALSE | FALSE |
| LOC107064971 | uncharacterized protein LOC107064971 | 0.03427 | 36.94761 | FALSE | FALSE |
| LOC107071798 | septin-2 | 0.03391 | 36.96188 | FALSE | FALSE |
| LOC107070621 | zinc metalloproteinase nas-4-like | 0.03457 | 36.9676 | FALSE | FALSE |
| LOC107072982 | uncharacterized protein LOC107072982 | 0.0336 | 36.96865 | FALSE | FALSE |
| LOC107074223 | general transcription factor IIH subunit 4 | 0.03259 | 36.9734 | FALSE | FALSE |
| LOC107072454 | tudor domain-containing protein 3 | 0.03395 | 36.97581 | FALSE | FALSE |
| LOC107074631 | prickle-like protein 3 | 0.03351 | 36.98018 | FALSE | FALSE |
| LOC107070534 | putative tyramine receptor 2 | 0.03332 | 36.98567 | FALSE | FALSE |
| LOC107070130 | ras-specific guanine nucleotide-releasing fa | 0.03379 | 36.98572 | FALSE | FALSE |

| | | | | | |
|---|---|---|---|---|---|
| LOC107067054 | uncharacterized protein LOC107067054 | 0.03318 | 36.99689 | FALSE | FALSE |
| LOC107072261 | uncharacterized protein LOC107072261 | 0.03466 | 37.01054 | FALSE | FALSE |
| LOC107063993 | golgin subfamily A member 2 | 0.03508 | 37.05308 | FALSE | FALSE |
| LOC107068873 | uncharacterized protein LOC107068873 | 0.03491 | 37.07457 | FALSE | FALSE |
| LOC107070793 | gem-associated protein 8-like | 0.03442 | 37.07487 | FALSE | FALSE |
| LOC107069582 | sugar transporter SWEET1 | 0.03435 | 37.08363 | FALSE | FALSE |
| LOC107066763 | nuclear pore complex protein Nup88 | 0.03367 | 37.09541 | FALSE | FALSE |
| LOC107067879 | nodal modulator 1 | 0.03394 | 37.10001 | FALSE | FALSE |
| LOC107066264 | cytochrome b-c1 complex subunit 2, mitoch | 0.0347 | 37.11413 | FALSE | FALSE |
| LOC107067839 | delta-like protein 1 | 0.03358 | 37.12796 | FALSE | FALSE |
| LOC107068260 | eukaryotic translation initiation factor 3 sub | 0.03314 | 37.13009 | FALSE | FALSE |
| LOC107073429 | uncharacterized protein LOC107073429 | 0.03382 | 37.14397 | FALSE | FALSE |
| LOC107065380 | RING finger protein 121 | 0.03289 | 37.158 | FALSE | FALSE |
| LOC107064444 | 3-hydroxyacyl-CoA dehydrogenase type-2-li | 0.03324 | 37.15845 | FALSE | FALSE |
| LOC107068222 | solute carrier family 12 member 7 | 0.0352 | 37.199 | FALSE | FALSE |
| LOC107070101 | uncharacterized protein LOC107070101 | 0.03364 | 37.21938 | FALSE | FALSE |
| LOC107065382 | MAGUK p55 subfamily member 6 | 0.03374 | 37.22199 | FALSE | FALSE |
| LOC107069515 | ras-related protein Rab-8A | 0.03427 | 37.22466 | FALSE | FALSE |
| LOC107072410 | striatin-interacting protein 1 homolog | 0.03461 | 37.22498 | FALSE | FALSE |
| LOC107071464 | autophagy-related protein 13 homolog | 0.03342 | 37.23167 | FALSE | FALSE |
| LOC107071849 | charged multivesicular body protein 2a | 0.03272 | 37.24311 | FALSE | FALSE |
| LOC107064158 | elongation factor 1-alpha | 0.03384 | 37.25582 | FALSE | FALSE |
| LOC107071083 | uncharacterized protein LOC107071083 | 0.03358 | 37.28158 | FALSE | FALSE |
| LOC107072278 | uncharacterized protein LOC107072278 | 0.03377 | 37.28767 | FALSE | FALSE |
| LOC107066367 | uncharacterized protein LOC107066367 | 0.03338 | 37.30332 | FALSE | FALSE |
| LOC107071153 | epidermal growth factor receptor | 0.033 | 37.31118 | FALSE | FALSE |
| LOC107069919 | segmentation protein even-skipped-like | 0.03328 | 37.31697 | FALSE | FALSE |
| LOC107066184 | serine/threonine-protein kinase tousled-like | 0.0334 | 37.31733 | FALSE | FALSE |
| LOC107070028 | uncharacterized protein LOC107070028 | 0.03283 | 37.31753 | FALSE | FALSE |
| LOC107068437 | vacuolar protein-sorting-associated protein | 0.03459 | 37.32282 | FALSE | FALSE |
| LOC107065145 | putative DNA helicase Ino80 | 0.03389 | 37.32283 | FALSE | FALSE |
| LOC107064965 | probable ribosome production factor 1 | 0.0344 | 37.32615 | FALSE | FALSE |
| LOC107066092 | uncharacterized protein LOC107066092 | 0.03414 | 37.33169 | FALSE | FALSE |
| LOC107064858 | protein turtle-like | 0.0319 | 37.3497 | FALSE | FALSE |
| LOC107073009 | phosphoribosyl pyrophosphate synthase-as | 0.03319 | 37.35127 | FALSE | FALSE |
| LOC107064670 | glutamate receptor ionotropic, kainate 2-lik | 0.03257 | 37.35339 | FALSE | FALSE |
| LOC107066659 | EP300-interacting inhibitor of differentiatio | 0.03372 | 37.39547 | FALSE | FALSE |
| LOC107071223 | DNA polymerase iota-like | 0.0341 | 37.39847 | FALSE | FALSE |
| LOC107067817 | peroxisomal membrane protein PEX16 | 0.034 | 37.405 | FALSE | FALSE |
| LOC107071104 | NADH dehydrogenase [ubiquinone | 0.03396 | 37.43411 | FALSE | FALSE |
| LOC107072306 | uncharacterized protein LOC107072306 | 0.03582 | 37.44446 | FALSE | FALSE |
| LOC107069949 | cysteine-rich protein 1-like | 0.03422 | 37.447 | FALSE | FALSE |
| LOC107069406 | uncharacterized protein LOC107069406 | 0.03235 | 37.45106 | FALSE | FALSE |
| LOC107068818 | uncharacterized protein LOC107068818 | 0.03484 | 37.46304 | FALSE | FALSE |
| Trnam-cau | Trnam-cau | 0.0332 | 37.46473 | FALSE | FALSE |
| LOC107067222 | uncharacterized protein LOC107067222 | 0.03364 | 37.48459 | FALSE | FALSE |
| LOC107063671 | peroxiredoxin-6 | 0.0326 | 37.50295 | FALSE | FALSE |
| LOC107073589 | putative fatty acyl-CoA reductase CG5065 | 0.03536 | 37.50444 | FALSE | FALSE |
| LOC107069879 | neogenin | 0.03331 | 37.55101 | FALSE | FALSE |
| LOC107068140 | ankyrin repeat domain-containing protein 4 | 0.03315 | 37.56422 | FALSE | FALSE |

| | | | | | |
|---|---|---|---|---|---|
| LOC107064654 | protein lethal(2)essential for life-like | 0.03334 | 37.56616 | FALSE | FALSE |
| LOC107071709 | 5'-nucleotidase domain-containing protein 3 | 0.03287 | 37.56774 | FALSE | FALSE |
| LOC107072248 | uncharacterized protein LOC107072248 | 0.03347 | 37.56799 | FALSE | FALSE |
| LOC107068142 | alpha/beta hydrolase domain-containing pro | 0.03403 | 37.58004 | FALSE | FALSE |
| LOC107067856 | putative uncharacterized protein DDB_G028 | 0.03448 | 37.58525 | FALSE | FALSE |
| LOC107065220 | uncharacterized protein LOC107065220 | 0.03421 | 37.59918 | FALSE | FALSE |
| LOC107065158 | uncharacterized protein LOC107065158 | 0.03333 | 37.60026 | FALSE | FALSE |
| LOC107068731 | uncharacterized protein LOC107068731 | 0.03287 | 37.61093 | FALSE | FALSE |
| LOC107072084 | mitochondrial import inner membrane trans | 0.03425 | 37.61797 | FALSE | FALSE |
| LOC107065130 | nucleoporin NUP188 homolog | 0.03343 | 37.62255 | FALSE | FALSE |
| LOC107068966 | DNA-directed RNA polymerase I subunit RPA | 0.03218 | 37.62511 | FALSE | FALSE |
| LOC107068338 | probable elongation factor 1-delta | 0.03359 | 37.63171 | FALSE | FALSE |
| LOC107069230 | MAP kinase-activating death domain protein | 0.03314 | 37.6341 | FALSE | FALSE |
| LOC107063793 | uncharacterized protein LOC107063793 | 0.03436 | 37.66863 | FALSE | FALSE |
| LOC107068781 | hemicentin-2-like | 0.03321 | 37.67563 | FALSE | FALSE |
| LOC107069740 | NADH dehydrogenase [ubiquinone | 0.03331 | 37.68332 | FALSE | FALSE |
| LOC107064633 | baculoviral IAP repeat-containing protein 5 | 0.03479 | 37.69743 | FALSE | FALSE |
| LOC107067450 | sphingomyelin synthase-related 1 | 0.03352 | 37.69845 | FALSE | FALSE |
| LOC107072974 | transcriptional coactivator YAP1 | 0.03353 | 37.70775 | FALSE | FALSE |
| LOC107067034 | peroxisomal membrane protein PMP34 | 0.03406 | 37.72038 | FALSE | FALSE |
| LOC107070795 | gamma-glutamyltranspeptidase 1-like | 0.03346 | 37.73081 | FALSE | FALSE |
| LOC107067393 | troponin I | 0.03348 | 37.73604 | FALSE | FALSE |
| LOC107066007 | beclin-1-like protein | 0.03279 | 37.73674 | FALSE | FALSE |
| LOC107074020 | endocuticle structural glycoprotein SgAbd-2 | 0.0332 | 37.76375 | FALSE | FALSE |
| LOC107063876 | mitochondrial import inner membrane trans | 0.03323 | 37.76964 | FALSE | FALSE |
| LOC107072302 | ATP synthase subunit epsilon, mitochondria | 0.03361 | 37.78244 | FALSE | FALSE |
| LOC107072837 | uncharacterized protein LOC107072837 | 0.03312 | 37.78353 | FALSE | FALSE |
| LOC107072546 | uncharacterized protein LOC107072546 | 0.03381 | 37.79533 | FALSE | FALSE |
| LOC107072214 | uncharacterized protein LOC107072214 | 0.03409 | 37.79692 | FALSE | FALSE |
| LOC107069558 | hydroxyacylglutathione hydrolase, mitochon | 0.03489 | 37.81254 | FALSE | FALSE |
| LOC107066304 | uncharacterized protein LOC107066304 | 0.03391 | 37.81948 | FALSE | FALSE |
| LOC107065056 | desumoylating isopeptidase 2 | 0.03367 | 37.834 | FALSE | FALSE |
| LOC107074168 | uncharacterized protein LOC107074168 | 0.03511 | 37.83614 | FALSE | FALSE |
| LOC107068774 | uncharacterized protein LOC107068774 | 0.03262 | 37.83841 | FALSE | FALSE |
| LOC107067195 | NEDD8 | 0.03451 | 37.84368 | FALSE | FALSE |
| LOC107072463 | protein canopy-1 | 0.03266 | 37.84972 | FALSE | FALSE |
| LOC107067370 | chitotriosidase-1-like | 0.03293 | 37.85812 | FALSE | FALSE |
| LOC107074627 | transmembrane and coiled-coil domains pro | 0.03267 | 37.86787 | FALSE | FALSE |
| LOC107068609 | uncharacterized protein LOC107068609 | 0.03329 | 37.88816 | FALSE | FALSE |
| LOC107069233 | serine-rich adhesin for platelets | 0.0341 | 37.90027 | FALSE | FALSE |
| LOC107069641 | uncharacterized protein LOC107069641 | 0.03442 | 37.9016 | FALSE | FALSE |
| LOC107066116 | zinc finger protein basonuclin-2-like | 0.03474 | 37.90742 | FALSE | FALSE |
| LOC107064260 | acyl-CoA-binding domain-containing protein | 0.0333 | 37.91111 | FALSE | FALSE |
| LOC107070031 | uncharacterized protein LOC107070031 | 0.0342 | 37.92205 | FALSE | FALSE |
| LOC107068256 | NGFI-A-binding protein homolog | 0.03391 | 37.93619 | FALSE | FALSE |
| LOC107071063 | uncharacterized protein LOC107071063 | 0.03237 | 37.94439 | FALSE | FALSE |
| LOC107074516 | ubiquitin carboxyl-terminal hydrolase 35-lik | 0.03268 | 37.94804 | FALSE | FALSE |
| LOC107065144 | protein yellow | 0.03259 | 37.94971 | FALSE | FALSE |
| LOC107066138 | troponin C, isoform 2-like | 0.03446 | 37.95237 | FALSE | FALSE |
| LOC107070405 | fatty acyl-CoA reductase 1 | 0.0337 | 37.9572 | FALSE | FALSE |

| | | | | | |
|---|---|---|---|---|---|
| LOC107072487 | ADP-ribosylation factor-like protein 6-intera | 0.03403 | 37.95926 | FALSE | FALSE |
| LOC107067729 | synaptic vesicle glycoprotein 2C-like | 0.03276 | 37.9629 | FALSE | FALSE |
| LOC107074476 | uncharacterized protein F21D5.5 | 0.03327 | 37.97442 | FALSE | FALSE |
| LOC107072900 | uncharacterized protein LOC107072900 | 0.03332 | 37.98022 | FALSE | FALSE |
| LOC107068989 | uncharacterized protein LOC107068989 | 0.03283 | 37.98566 | FALSE | FALSE |
| LOC107072813 | leucine-rich repeat-containing protein 15 | 0.03307 | 37.99096 | FALSE | FALSE |
| LOC107069428 | MAGUK p55 subfamily member 5-A | 0.03223 | 37.99982 | FALSE | FALSE |
| LOC107069960 | coatomer subunit zeta-1 | 0.03364 | 38.0024 | FALSE | FALSE |
| LOC107070090 | ras-related protein Rab-21 | 0.03232 | 38.00868 | FALSE | FALSE |
| LOC107065143 | protein yellow-like | 0.03398 | 38.036 | FALSE | FALSE |
| LOC107070061 | uncharacterized protein PF11_0213-like | 0.03304 | 38.03603 | FALSE | FALSE |
| LOC107073114 | ankyrin-3-like | 0.03341 | 38.05854 | FALSE | FALSE |
| LOC107070471 | flavin-containing monooxygenase FMO GS-( | 0.03442 | 38.06254 | FALSE | FALSE |
| LOC107069290 | protein claret segregational | 0.03313 | 38.07101 | FALSE | FALSE |
| LOC107074385 | N-terminal Xaa-Pro-Lys N-methyltransferase | 0.03303 | 38.07105 | FALSE | FALSE |
| LOC107067619 | uncharacterized protein LOC107067619 | 0.03386 | 38.0714 | FALSE | FALSE |
| LOC107068021 | zinc finger protein DZIP1L | 0.03292 | 38.07148 | FALSE | FALSE |
| LOC107067629 | alanine--tRNA ligase, cytoplasmic | 0.03285 | 38.07314 | FALSE | FALSE |
| LOC107073547 | uncharacterized protein LOC107073547 | 0.03257 | 38.07332 | FALSE | FALSE |
| LOC107063777 | coiled-coil domain-containing protein 124 | 0.03199 | 38.07985 | FALSE | FALSE |
| LOC107066546 | protein SREK1IP1-like | 0.03179 | 38.08333 | FALSE | FALSE |
| LOC107063853 | RNA-binding protein NOB1 | 0.03439 | 38.08646 | FALSE | FALSE |
| LOC107066247 | uncharacterized protein LOC107066247 | 0.0337 | 38.09719 | FALSE | FALSE |
| LOC107069562 | storkhead-box protein 2 | 0.03359 | 38.10245 | FALSE | FALSE |
| LOC107070209 | MATH and LRR domain-containing protein F | 0.03228 | 38.10321 | FALSE | FALSE |
| LOC107064671 | F-box only protein 22-like | 0.03398 | 38.11509 | FALSE | FALSE |
| LOC107072158 | double-stranded RNA-specific editase B2-lik | 0.03281 | 38.11566 | FALSE | FALSE |
| LOC107071383 | cold shock domain-containing protein E1 | 0.0336 | 38.12169 | FALSE | FALSE |
| LOC107063809 | dual serine/threonine and tyrosine protein l | 0.03338 | 38.12247 | FALSE | FALSE |
| LOC107071678 | MICOS complex subunit Mic60 | 0.03325 | 38.13137 | FALSE | FALSE |
| LOC107071341 | uncharacterized protein LOC107071341 | 0.0335 | 38.13307 | FALSE | FALSE |
| LOC107070635 | josephin-2 | 0.03348 | 38.13366 | FALSE | FALSE |
| LOC107065135 | protein LSM14 homolog B | 0.03505 | 38.13689 | FALSE | FALSE |
| LOC107067719 | myb-like protein X | 0.03217 | 38.13898 | FALSE | FALSE |
| LOC107066492 | uncharacterized protein LOC107066492 | 0.03346 | 38.16147 | FALSE | FALSE |
| LOC107073054 | serine-rich adhesin for platelets-like | 0.03337 | 38.16313 | FALSE | FALSE |
| LOC107072307 | uncharacterized protein LOC107072307 | 0.03257 | 38.17362 | FALSE | FALSE |
| LOC107066793 | probable ATP-dependent RNA helicase pitch | 0.03307 | 38.17389 | FALSE | FALSE |
| LOC107073596 | fatty acid synthase-like | 0.03285 | 38.17456 | FALSE | FALSE |
| LOC107074358 | methyltransferase-like protein 16 homolog | 0.03327 | 38.17709 | FALSE | FALSE |
| LOC107074407 | regulator of G-protein signaling 7 | 0.03304 | 38.19024 | FALSE | FALSE |
| LOC107071186 | nipped-B-like protein A | 0.03299 | 38.20367 | FALSE | FALSE |
| LOC107069056 | uncharacterized protein LOC107069056 | 0.03312 | 38.20574 | FALSE | FALSE |
| LOC107069209 | uncharacterized protein LOC107069209 | 0.03259 | 38.20976 | FALSE | FALSE |
| LOC107065988 | F-box/LRR-repeat protein 14-like | 0.03235 | 38.22039 | FALSE | FALSE |
| LOC107073400 | 1,4-alpha-glucan-branching enzyme | 0.03308 | 38.22195 | FALSE | FALSE |
| LOC107069189 | ribulose-phosphate 3-epimerase | 0.03403 | 38.22314 | FALSE | FALSE |
| LOC107063971 | HEAT repeat-containing protein 1 | 0.03279 | 38.22653 | FALSE | FALSE |
| LOC107073509 | uncharacterized protein LOC107073509 | 0.03323 | 38.2293 | FALSE | FALSE |
| LOC107066784 | dehydrodolichyl diphosphate syntase compl | 0.03418 | 38.2305 | FALSE | FALSE |

| | | | | | |
|---|---|---|---|---|---|
| LOC107071992 | arrestin domain-containing protein 3-like | 0.03345 | 38.23053 | FALSE | FALSE |
| LOC107070923 | uncharacterized protein LOC107070923 | 0.03344 | 38.26859 | FALSE | FALSE |
| LOC107071598 | uncharacterized protein LOC107071598 | 0.03264 | 38.26989 | FALSE | FALSE |
| LOC107070618 | uncharacterized protein LOC107070618 | 0.03339 | 38.27073 | FALSE | FALSE |
| LOC107067389 | uncharacterized protein LOC107067389 | 0.03382 | 38.27522 | FALSE | FALSE |
| LOC107074021 | phosphatidylinositol glycan anchor biosynth | 0.03458 | 38.27582 | FALSE | FALSE |
| LOC107070993 | PHD finger protein rhinoceros | 0.03292 | 38.27985 | FALSE | FALSE |
| LOC107068539 | uncharacterized protein LOC107068539 | 0.03297 | 38.28467 | FALSE | FALSE |
| LOC107069162 | nuclear RNA export factor 1 | 0.03305 | 38.28836 | FALSE | FALSE |
| LOC107064142 | probable phosphorylase b kinase regulatory | 0.03316 | 38.29345 | FALSE | FALSE |
| LOC107072310 | 26S proteasome non-ATPase regulatory sub | 0.03363 | 38.29597 | FALSE | FALSE |
| LOC107072493 | probable splicing factor 3B subunit 5 | 0.03239 | 38.29767 | FALSE | FALSE |
| LOC107063791 | uncharacterized protein LOC107063791 | 0.03301 | 38.30263 | FALSE | FALSE |
| LOC107074290 | uncharacterized protein LOC107074290 | 0.03386 | 38.3052 | FALSE | FALSE |
| LOC107069023 | uncharacterized protein LOC107069023 | 0.03321 | 38.31625 | FALSE | FALSE |
| LOC107064189 | 5'-3' exoribonuclease 2 homolog | 0.03299 | 38.32015 | FALSE | FALSE |
| LOC107065213 | myb-binding protein 1A-like protein | 0.0329 | 38.32021 | FALSE | FALSE |
| LOC107073373 | inhibitor of growth protein 5 | 0.03329 | 38.32462 | FALSE | FALSE |
| LOC107072149 | uncharacterized protein LOC107072149 | 0.03322 | 38.32645 | FALSE | FALSE |
| LOC107066932 | protein maelstrom 1-like | 0.0335 | 38.35774 | FALSE | FALSE |
| LOC107070948 | uncharacterized protein LOC107070948 | 0.03274 | 38.38094 | FALSE | FALSE |
| LOC107074303 | uncharacterized protein LOC107074303 | 0.0326 | 38.39854 | FALSE | FALSE |
| LOC107070956 | uncharacterized protein LOC107070956 | 0.03347 | 38.40031 | FALSE | FALSE |
| LOC107066820 | MAU2 chromatid cohesion factor homolog | 0.03352 | 38.41389 | FALSE | FALSE |
| LOC107066541 | mevalonate kinase | 0.0323 | 38.41558 | FALSE | FALSE |
| LOC107069054 | trypsin-7-like | 0.03371 | 38.42624 | FALSE | FALSE |
| LOC107074078 | homeobox protein prospero | 0.03282 | 38.42812 | FALSE | FALSE |
| LOC107065555 | slit homolog 2 protein-like | 0.03394 | 38.44661 | FALSE | FALSE |
| LOC107074509 | estradiol 17-beta-dehydrogenase 11 | 0.03411 | 38.45249 | FALSE | FALSE |
| LOC107067570 | vacuolar ATPase assembly integral membra | 0.03332 | 38.45826 | FALSE | FALSE |
| LOC107071251 | tubulin alpha chain-like | 0.03391 | 38.46378 | FALSE | FALSE |
| LOC107073790 | piggyBac transposable element-derived pro | 0.03087 | 38.46726 | FALSE | FALSE |
| LOC107070561 | HMG box transcription factor BBX | 0.03268 | 38.47619 | FALSE | FALSE |
| LOC107073641 | uncharacterized protein LOC107073641 | 0.03251 | 38.47936 | FALSE | FALSE |
| LOC107065565 | SH3 domain-binding protein 5 homolog | 0.03278 | 38.48708 | FALSE | FALSE |
| LOC107074648 | phosphofurin acidic cluster sorting protein 2 | 0.0338 | 38.49253 | FALSE | FALSE |
| LOC107073916 | GRAM domain-containing protein 3-like | 0.03295 | 38.50127 | FALSE | FALSE |
| LOC107065261 | uncharacterized protein LOC107065261 | 0.0333 | 38.52204 | FALSE | FALSE |
| LOC107072866 | histone-lysine N-methyltransferase SETMAR | 0.03388 | 38.52447 | FALSE | FALSE |
| LOC107070772 | uncharacterized protein LOC107070772 | 0.03354 | 38.52952 | FALSE | FALSE |
| LOC107070366 | uncharacterized protein LOC107070366 | 0.03234 | 38.5309 | FALSE | FALSE |
| LOC107067818 | carbonic anhydrase 2-like | 0.0323 | 38.53698 | FALSE | FALSE |
| LOC107072730 | solute carrier family 52, riboflavin transport | 0.03451 | 38.54878 | FALSE | FALSE |
| LOC107074323 | kinesin-related protein 4-like | 0.03253 | 38.56327 | FALSE | FALSE |
| LOC107067066 | 101 kDa malaria antigen-like | 0.03127 | 38.56788 | FALSE | FALSE |
| LOC107071584 | signal recognition particle subunit SRP68 | 0.03341 | 38.58422 | FALSE | FALSE |
| LOC107073498 | uncharacterized protein LOC107073498 | 0.03263 | 38.59317 | FALSE | FALSE |
| LOC107065112 | Krueppel-like factor 6 | 0.03347 | 38.59457 | FALSE | FALSE |
| LOC107066307 | putative sodium-dependent multivitamin tr | 0.03428 | 38.59957 | FALSE | FALSE |
| LOC107069228 | cholinephosphotransferase 1 | 0.03346 | 38.60861 | FALSE | FALSE |

| | | | | | |
|---|---|---|---|---|---|
| LOC107069681 | uncharacterized protein LOC107069681 | 0.03381 | 38.61264 | FALSE | FALSE |
| LOC107069973 | uncharacterized protein LOC107069973 | 0.03245 | 38.61589 | FALSE | FALSE |
| LOC107064936 | uncharacterized protein LOC107064936 | 0.03332 | 38.61852 | FALSE | FALSE |
| LOC107065760 | glyoxylate reductase/hydroxypyruvate redu | 0.0322 | 38.62199 | FALSE | FALSE |
| LOC107068031 | TGF-beta-activated kinase 1 and MAP3K7-bi | 0.03287 | 38.63049 | FALSE | FALSE |
| LOC107070898 | uncharacterized protein LOC107070898 | 0.03324 | 38.65295 | FALSE | FALSE |
| LOC107064781 | uncharacterized protein LOC107064781 | 0.03374 | 38.66023 | FALSE | FALSE |
| LOC107069297 | eukaryotic translation initiation factor 2 sub | 0.03382 | 38.66614 | FALSE | FALSE |
| LOC107071824 | uncharacterized protein LOC107071824 | 0.03291 | 38.66685 | FALSE | FALSE |
| LOC107066593 | programmed cell death protein 10 | 0.0333 | 38.66853 | FALSE | FALSE |
| LOC107068603 | putative pre-mRNA-splicing factor ATP-depe | 0.03266 | 38.69254 | FALSE | FALSE |
| LOC107072338 | uncharacterized protein LOC107072338 | 0.03362 | 38.69543 | FALSE | FALSE |
| LOC107065500 | cytochrome c oxidase assembly factor 7 hor | 0.03293 | 38.69551 | FALSE | FALSE |
| LOC107063963 | beta-parvin | 0.03219 | 38.70303 | FALSE | FALSE |
| LOC107069700 | WD repeat-containing protein 46 | 0.03356 | 38.7163 | FALSE | FALSE |
| LOC107068960 | malate dehydrogenase, mitochondrial-like | 0.03299 | 38.71796 | FALSE | FALSE |
| LOC107073187 | dnaJ homolog subfamily C member 21 | 0.03233 | 38.71886 | FALSE | FALSE |
| LOC107071753 | transcription factor SPT20 homolog | 0.03353 | 38.72035 | FALSE | FALSE |
| LOC107069431 | integrin alpha-PS1 | 0.03168 | 38.77282 | FALSE | FALSE |
| LOC107072576 | cadherin-related tumor suppressor | 0.03254 | 38.77775 | FALSE | FALSE |
| LOC107073776 | uncharacterized protein LOC107073776 | 0.03244 | 38.78537 | FALSE | FALSE |
| LOC107065805 | SUMO-activating enzyme subunit 2-like | 0.03288 | 38.79959 | FALSE | FALSE |
| LOC107067384 | homeobox protein six1-like | 0.03456 | 38.80348 | FALSE | FALSE |
| LOC107064040 | CKLF-like MARVEL transmembrane domain- | 0.03474 | 38.80411 | FALSE | FALSE |
| LOC107068265 | aldose 1-epimerase-like | 0.03391 | 38.81697 | FALSE | FALSE |
| LOC107068283 | folliculin-interacting protein 1 | 0.03248 | 38.83488 | FALSE | FALSE |
| LOC107073858 | uncharacterized protein LOC107073858 | 0.03296 | 38.84453 | FALSE | FALSE |
| LOC107074430 | inositol monophosphatase 1-like | 0.03311 | 38.85095 | FALSE | FALSE |
| LOC107065284 | FACT complex subunit Ssrp1 | 0.03465 | 38.85751 | FALSE | FALSE |
| LOC107068994 | acetylcholine receptor subunit beta-like 2 | 0.03307 | 38.8657 | FALSE | FALSE |
| LOC107073864 | myb-like protein I | 0.03312 | 38.86618 | FALSE | FALSE |
| LOC107064141 | uncharacterized protein LOC107064141 | 0.03196 | 38.87529 | FALSE | FALSE |
| LOC107068938 | probable cytochrome P450 6a14 | 0.03333 | 38.87683 | FALSE | FALSE |
| LOC107066529 | arginine-glutamic acid dipeptide repeats pro | 0.03398 | 38.88284 | FALSE | FALSE |
| LOC107065974 | acetylcholinesterase | 0.03289 | 38.89167 | FALSE | FALSE |
| LOC107074458 | fibrillin-2-like | 0.03281 | 38.89808 | FALSE | FALSE |
| LOC107073691 | alpha-protein kinase 1-like | 0.03241 | 38.89979 | FALSE | FALSE |
| LOC107066807 | FYVE, RhoGEF and PH domain-containing pr | 0.03397 | 38.90241 | FALSE | FALSE |
| LOC107072254 | solute carrier family 35 member E3-like | 0.03139 | 38.90668 | FALSE | FALSE |
| LOC107072155 | DNA primase small subunit | 0.03276 | 38.91467 | FALSE | FALSE |
| LOC107070057 | wee1-like protein kinase | 0.03321 | 38.91606 | FALSE | FALSE |
| LOC107074381 | ral guanine nucleotide dissociation stimulat | 0.0339 | 38.92378 | FALSE | FALSE |
| LOC107072268 | chondroadherin-like protein | 0.03246 | 38.92491 | FALSE | FALSE |
| LOC107071925 | ninein | 0.03204 | 38.93024 | FALSE | FALSE |
| LOC107068460 | uncharacterized protein LOC107068460 | 0.03318 | 38.94239 | FALSE | FALSE |
| LOC107071693 | ATP-dependent RNA helicase bel-like | 0.03306 | 38.94597 | FALSE | FALSE |
| LOC107069433 | GPN-loop GTPase 1 | 0.03253 | 38.94929 | FALSE | FALSE |
| LOC107068229 | uncharacterized protein LOC107068229 | 0.03284 | 38.95891 | FALSE | FALSE |
| LOC107070960 | putative leucine-rich repeat-containing prot | 0.03193 | 38.97806 | FALSE | FALSE |
| LOC107072862 | OTU domain-containing protein 5-B | 0.03235 | 38.98802 | FALSE | FALSE |

| | | | | | |
|---|---|---|---|---|---|
| LOC107074335 | E3 ubiquitin-protein ligase HECW2-like | 0.03293 | 38.99811 | FALSE | FALSE |
| LOC107065921 | aminoacyl tRNA synthase complex-interacti | 0.03117 | 39.00199 | FALSE | FALSE |
| LOC107065292 | zinc finger and SCAN domain-containing pro | 0.03134 | 39.00506 | FALSE | FALSE |
| LOC107074203 | sprouty-related, EVH1 domain-containing pr | 0.03383 | 39.01588 | FALSE | FALSE |
| LOC107072524 | LIM/homeobox protein Awh-like | 0.03312 | 39.01848 | FALSE | FALSE |
| LOC107073565 | fatty acid synthase-like | 0.03312 | 39.02738 | FALSE | FALSE |
| LOC107073337 | formin-like protein CG32138 | 0.03298 | 39.03218 | FALSE | FALSE |
| LOC107069806 | uncharacterized protein LOC107069806 | 0.03142 | 39.03354 | FALSE | FALSE |
| LOC107063705 | xanthine dehydrogenase-like | 0.03475 | 39.03416 | FALSE | FALSE |
| LOC107064915 | beta-galactosidase-1-like protein | 0.03204 | 39.03759 | FALSE | FALSE |
| LOC107073757 | piggyBac transposable element-derived pro | 0.03332 | 39.06466 | FALSE | FALSE |
| LOC107074702 | uncharacterized protein LOC107074702 | 0.03282 | 39.06576 | FALSE | FALSE |
| LOC107071666 | fumarylacetoacetate hydrolase domain-con | 0.03461 | 39.08125 | FALSE | FALSE |
| LOC107068625 | 60S ribosomal protein L28 | 0.03194 | 39.09139 | FALSE | FALSE |
| LOC107070367 | protein Fe65 homolog | 0.03333 | 39.09345 | FALSE | FALSE |
| LOC107071421 | uncharacterized protein LOC107071421 | 0.03311 | 39.11015 | FALSE | FALSE |
| LOC107069993 | gamma-secretase subunit Aph-1 | 0.03314 | 39.11611 | FALSE | FALSE |
| LOC107070273 | uncharacterized protein LOC107070273 | 0.03234 | 39.11669 | FALSE | FALSE |
| LOC107069557 | transmembrane protein 104 homolog | 0.03294 | 39.12006 | FALSE | FALSE |
| LOC107066476 | uncharacterized protein LOC107066476 | 0.03388 | 39.12384 | FALSE | FALSE |
| LOC107068541 | 60S ribosomal protein L7 | 0.03217 | 39.12704 | FALSE | FALSE |
| LOC107065887 | E3 ubiquitin-protein ligase UBR3 | 0.03076 | 39.13116 | FALSE | FALSE |
| LOC107070553 | uncharacterized protein LOC107070553 | 0.03293 | 39.13905 | FALSE | FALSE |
| LOC107074079 | uncharacterized protein LOC107074079 | 0.03364 | 39.14057 | FALSE | FALSE |
| LOC107067469 | ATP-binding cassette sub-family A member | 0.03244 | 39.14135 | FALSE | FALSE |
| LOC107069754 | tropomyosin | 0.03317 | 39.15225 | FALSE | FALSE |
| LOC107065663 | putative inorganic phosphate cotransporter | 0.03237 | 39.15762 | FALSE | FALSE |
| LOC107070563 | UDP-glucuronosyltransferase 1-8 | 0.03325 | 39.16044 | FALSE | FALSE |
| LOC107068164 | ubiquitin carboxyl-terminal hydrolase 48-lik | 0.03271 | 39.16123 | FALSE | FALSE |
| LOC107066185 | uncharacterized protein LOC107066185 | 0.03308 | 39.17661 | FALSE | FALSE |
| LOC107064485 | dynein heavy chain 10, axonemal | 0.0324 | 39.17687 | FALSE | FALSE |
| LOC107064706 | dachshund homolog 2 | 0.03338 | 39.19574 | FALSE | FALSE |
| LOC107072969 | loricrin-like | 0.03322 | 39.20892 | FALSE | FALSE |
| LOC107074609 | probable C-mannosyltransferase DPY19L1 | 0.0313 | 39.21083 | FALSE | FALSE |
| LOC107070935 | uncharacterized protein LOC107070935 | 0.03213 | 39.21168 | FALSE | FALSE |
| LOC107070451 | protein PBDC1 | 0.03196 | 39.21462 | FALSE | FALSE |
| LOC107066786 | probable dimethyladenosine transferase | 0.03249 | 39.262 | FALSE | FALSE |
| LOC107068985 | ankyrin repeat domain-containing protein 2 | 0.03383 | 39.28021 | FALSE | FALSE |
| LOC107070063 | uncharacterized protein LOC107070063 | 0.03292 | 39.28475 | FALSE | FALSE |
| LOC107068897 | semaphorin-5A | 0.03267 | 39.30029 | FALSE | FALSE |
| LOC107064761 | protein mab-21-like | 0.03316 | 39.30215 | FALSE | FALSE |
| LOC107074398 | cysteine protease ATG4D | 0.03263 | 39.30905 | FALSE | FALSE |
| LOC107065550 | mannose-1-phosphate guanyltransferase al | 0.03293 | 39.3141 | FALSE | FALSE |
| LOC107074050 | sodium channel protein Nach-like | 0.03314 | 39.32311 | FALSE | FALSE |
| LOC107073627 | uncharacterized protein LOC107073627 | 0.03407 | 39.32876 | FALSE | FALSE |
| LOC107069795 | uncharacterized protein LOC107069795 | 0.03354 | 39.33798 | FALSE | FALSE |
| LOC107064502 | histidine--tRNA ligase, cytoplasmic | 0.03293 | 39.3396 | FALSE | FALSE |
| LOC107069421 | E3 ubiquitin-protein ligase MARCH2-like | 0.03364 | 39.35064 | FALSE | FALSE |
| LOC107066244 | mpv17-like protein | 0.0319 | 39.36058 | FALSE | FALSE |
| LOC107073766 | uncharacterized protein LOC107073766 | 0.03199 | 39.37661 | FALSE | FALSE |

| | | | | | |
|---|---|---|---|---|---|
| LOC107066797 | uncharacterized protein LOC107066797 | 0.03298 | 39.38378 | FALSE | FALSE |
| LOC107070260 | glycine N-methyltransferase | 0.0335 | 39.38455 | FALSE | FALSE |
| LOC107066581 | 60S ribosomal protein L24 | 0.03341 | 39.40856 | FALSE | FALSE |
| LOC107063786 | uncharacterized protein LOC107063786 | 0.03256 | 39.41532 | FALSE | FALSE |
| LOC107069884 | uncharacterized protein LOC107069884 | 0.03241 | 39.42501 | FALSE | FALSE |
| LOC107064455 | uncharacterized protein LOC107064455 | 0.03283 | 39.42609 | FALSE | FALSE |
| LOC107072762 | neprilysin-21-like | 0.03242 | 39.43842 | FALSE | FALSE |
| LOC107071984 | cytochrome P450 6k1-like | 0.03187 | 39.44365 | FALSE | FALSE |
| LOC107063619 | ATP-dependent RNA helicase bel-like | 0.03183 | 39.44525 | FALSE | FALSE |
| LOC107074151 | arfaptin-2 | 0.03185 | 39.47064 | FALSE | FALSE |
| LOC107065445 | uncharacterized protein LOC107065445 | 0.03212 | 39.48663 | FALSE | FALSE |
| LOC107067385 | glycine-rich cell wall structural protein 2 | 0.03389 | 39.49271 | FALSE | FALSE |
| LOC107066262 | uncharacterized protein LOC107066262 | 0.03259 | 39.49726 | FALSE | FALSE |
| LOC107068694 | uncharacterized protein LOC107068694 | 0.03358 | 39.50303 | FALSE | FALSE |
| LOC107073452 | enhancer of mRNA-decapping protein 4 | 0.03383 | 39.50511 | FALSE | FALSE |
| LOC107069585 | uncharacterized protein LOC107069585 | 0.03267 | 39.51093 | FALSE | FALSE |
| LOC107064314 | phosphatidylinositide phosphatase SAC1 | 0.03168 | 39.51444 | FALSE | FALSE |
| LOC107066781 | cilia- and flagella-associated protein 61-like | 0.03236 | 39.52124 | FALSE | FALSE |
| LOC107067289 | 40S ribosomal protein S29 | 0.03189 | 39.52511 | FALSE | FALSE |
| LOC107072960 | uncharacterized protein LOC107072960 | 0.03268 | 39.54685 | FALSE | FALSE |
| LOC107069712 | ecdysone-induced protein 74EF isoform A | 0.03233 | 39.55024 | FALSE | FALSE |
| LOC107072516 | ubiquinone biosynthesis O-methyltransfera: | 0.03296 | 39.55125 | FALSE | FALSE |
| LOC107064526 | uncharacterized protein LOC107064526 | 0.03263 | 39.55175 | FALSE | FALSE |
| LOC107071366 | uncharacterized protein LOC107071366 | 0.03386 | 39.55771 | FALSE | FALSE |
| LOC107072534 | arginine kinase | 0.03242 | 39.56888 | FALSE | FALSE |
| LOC107070415 | golgin subfamily A member 6-like protein 1 | 0.03441 | 39.57466 | FALSE | FALSE |
| LOC107067759 | meiosis-specific with OB domain-containing | 0.0336 | 39.58178 | FALSE | FALSE |
| LOC107067530 | longitudinals lacking protein, isoforms A/B/I | 0.03254 | 39.58981 | FALSE | FALSE |
| LOC107069969 | polypeptide N-acetylgalactosaminyltransfer | 0.03234 | 39.59433 | FALSE | FALSE |
| LOC107067770 | organic solute transporter subunit alpha-lik( | 0.03406 | 39.60036 | FALSE | FALSE |
| LOC107069028 | ADAMTS-like protein 4 | 0.03296 | 39.6049 | FALSE | FALSE |
| LOC107070115 | uncharacterized protein LOC107070115 | 0.03283 | 39.60815 | FALSE | FALSE |
| LOC107074528 | SCY1-like protein 2 | 0.03194 | 39.61042 | FALSE | FALSE |
| LOC107074089 | RNA polymerase I-specific transcription initi | 0.03232 | 39.63575 | FALSE | FALSE |
| LOC107068155 | probable pyruvate dehydrogenase E1 comp | 0.03251 | 39.64432 | FALSE | FALSE |
| LOC107070598 | 40S ribosomal protein S21 | 0.0316 | 39.68212 | FALSE | FALSE |
| LOC107071179 | serine/threonine-protein phosphatase 6 cat | 0.03317 | 39.68328 | FALSE | FALSE |
| LOC107071652 | thyroid receptor-interacting protein 6 | 0.03319 | 39.70291 | FALSE | FALSE |
| LOC107069106 | transport and Golgi organization protein 6 h | 0.03345 | 39.71618 | FALSE | FALSE |
| LOC107068276 | venom acid phosphatase Acph-1-like | 0.03372 | 39.717 | FALSE | FALSE |
| LOC107072865 | leucine-rich repeat-containing protein 16A | 0.03307 | 39.74516 | FALSE | FALSE |
| LOC107069523 | uncharacterized protein LOC107069523 | 0.03197 | 39.77191 | FALSE | FALSE |
| LOC107069029 | photoreceptor-specific nuclear receptor-lik( | 0.03274 | 39.78234 | FALSE | FALSE |
| LOC107064004 | ribonuclease P protein subunit p40-like | 0.03376 | 39.78656 | FALSE | FALSE |
| LOC107066012 | uncharacterized protein LOC107066012 | 0.03307 | 39.79003 | FALSE | FALSE |
| LOC107070692 | uncharacterized protein LOC107070692 | 0.03319 | 39.79514 | FALSE | FALSE |
| LOC107065880 | ribonuclease H2 subunit B | 0.03145 | 39.81909 | FALSE | FALSE |
| LOC107068301 | cytochrome P450 4C1-like | 0.03129 | 39.82087 | FALSE | FALSE |
| LOC107065270 | transcription factor A, mitochondrial | 0.03227 | 39.83234 | FALSE | FALSE |
| LOC107066379 | heterogeneous nuclear ribonucleoprotein 2 | 0.03278 | 39.8352 | FALSE | FALSE |

| | | | | | |
|---|---|---|---|---|---|
| LOC107071976 | battenin | 0.03229 | 39.83648 | FALSE | FALSE |
| LOC107064276 | organic cation transporter protein | 0.03263 | 39.84602 | FALSE | FALSE |
| LOC107067337 | putative uncharacterized protein DDB_G028 | 0.03238 | 39.85088 | FALSE | FALSE |
| LOC107074472 | glutamate dehydrogenase, mitochondrial | 0.03377 | 39.85837 | FALSE | FALSE |
| LOC107066675 | putative uncharacterized protein DDB_G027 | 0.03228 | 39.88012 | FALSE | FALSE |
| LOC107069935 | uncharacterized protein LOC107069935 | 0.03227 | 39.88165 | FALSE | FALSE |
| LOC107074215 | uncharacterized protein LOC107074215 | 0.03238 | 39.88451 | FALSE | FALSE |
| LOC107070774 | midnolin-like | 0.03223 | 39.88583 | FALSE | FALSE |
| LOC107073376 | vacuolar protein sorting-associated protein | 0.03169 | 39.89557 | FALSE | FALSE |
| LOC107067404 | 43 kDa receptor-associated protein of the sy | 0.0323 | 39.90093 | FALSE | FALSE |
| LOC107067276 | stabilizer of axonemal microtubules 2-like | 0.03328 | 39.9093 | FALSE | FALSE |
| LOC107067087 | uncharacterized protein LOC107067087 | 0.03331 | 39.9148 | FALSE | FALSE |
| LOC107068281 | uncharacterized protein LOC107068281 | 0.034 | 39.91594 | FALSE | FALSE |
| LOC107064422 | dipeptidase 1-like | 0.03282 | 39.92911 | FALSE | FALSE |
| LOC107069472 | odorant receptor 30a-like | 0.03343 | 39.94573 | FALSE | FALSE |
| LOC107072977 | bladder cancer-associated protein | 0.03353 | 39.94677 | FALSE | FALSE |
| LOC107064443 | protein cueball | 0.0331 | 39.9815 | FALSE | FALSE |
| LOC107067373 | cytochrome c oxidase subunit 6b-1-like | 0.03216 | 39.98346 | FALSE | FALSE |
| LOC107069832 | protein phosphatase PTC7 homolog | 0.03197 | 39.98445 | FALSE | FALSE |
| LOC107074188 | zinc finger protein 419-like | 0.03173 | 39.99198 | FALSE | FALSE |
| LOC107069990 | uncharacterized protein LOC107069990 | 0.03336 | 39.99321 | FALSE | FALSE |
| LOC107069897 | conserved oligomeric Golgi complex subunit | 0.03281 | 40.01053 | FALSE | FALSE |
| LOC107066101 | cholinesterase 1-like | 0.03325 | 40.01546 | FALSE | FALSE |
| LOC107069500 | gustatory receptor 5a for trehalose-like | 0.03283 | 40.0253 | FALSE | FALSE |
| LOC107070894 | ras and EF-hand domain-containing protein | 0.03316 | 40.02651 | FALSE | FALSE |
| LOC107074366 | uncharacterized protein LOC107074366 | 0.03222 | 40.02878 | FALSE | FALSE |
| LOC107069669 | ubiquitin carboxyl-terminal hydrolase 30 ho | 0.0318 | 40.0344 | FALSE | FALSE |
| LOC107072448 | uncharacterized protein LOC107072448 | 0.03181 | 40.05523 | FALSE | FALSE |
| LOC107074064 | uncharacterized protein LOC107074064 | 0.03294 | 40.05572 | FALSE | FALSE |
| LOC107071707 | uncharacterized protein LOC107071707 | 0.03216 | 40.06412 | FALSE | FALSE |
| LOC107074265 | mucin-19-like | 0.03195 | 40.07445 | FALSE | FALSE |
| LOC107065264 | ecdysone 20-monooxygenase | 0.03146 | 40.09914 | FALSE | FALSE |
| LOC107068615 | uncharacterized protein LOC107068615 | 0.03176 | 40.10382 | FALSE | FALSE |
| LOC107066440 | uncharacterized protein LOC107066440 | 0.03353 | 40.11232 | FALSE | FALSE |
| LOC107073227 | paraplegin | 0.03234 | 40.1152 | FALSE | FALSE |
| LOC107068961 | adenylate kinase isoenzyme 1-like | 0.03094 | 40.11852 | FALSE | FALSE |
| LOC107070832 | uncharacterized protein LOC107070832 | 0.03274 | 40.12949 | FALSE | FALSE |
| LOC107073680 | glutamic acid-rich protein-like | 0.03246 | 40.13025 | FALSE | FALSE |
| LOC107065891 | 40S ribosomal protein S15Aa | 0.03114 | 40.1367 | FALSE | FALSE |
| LOC107066180 | dystrophin-like | 0.03201 | 40.14414 | FALSE | FALSE |
| LOC107069481 | GPI mannosyltransferase 2 | 0.03212 | 40.15103 | FALSE | FALSE |
| LOC107070312 | protein HGH1 homolog | 0.032 | 40.1923 | FALSE | FALSE |
| LOC107067797 | E3 ubiquitin-protein ligase listerin | 0.03265 | 40.19306 | FALSE | FALSE |
| LOC107071228 | transmembrane protein 222 | 0.03195 | 40.19381 | FALSE | FALSE |
| LOC107068564 | protein still life, isoform SIF type 1 | 0.03317 | 40.1952 | FALSE | FALSE |
| LOC107072851 | transmembrane protein 135-like | 0.03149 | 40.20214 | FALSE | FALSE |
| LOC107064565 | protein O-GlcNAcase | 0.03223 | 40.21318 | FALSE | FALSE |
| LOC107064304 | 60S ribosomal protein L18 | 0.03207 | 40.219 | FALSE | FALSE |
| LOC107071555 | uncharacterized protein LOC107071555 | 0.03306 | 40.22528 | FALSE | FALSE |
| LOC107072006 | leucine-rich repeat protein 1 | 0.03218 | 40.24216 | FALSE | FALSE |

| | | | | | |
|---|---|---|---|---|---|
| LOC107067377 | glyceraldehyde-3-phosphate dehydrogenas | 0.03159 | 40.24994 | FALSE | FALSE |
| LOC107066513 | segment polarity protein dishevelled homol | 0.03236 | 40.25507 | FALSE | FALSE |
| LOC107070301 | glutathione S-transferase-like | 0.03322 | 40.27272 | FALSE | FALSE |
| LOC107064883 | TD and POZ domain-containing protein 1-lik | 0.03179 | 40.27688 | FALSE | FALSE |
| LOC107065019 | aftiphilin | 0.03234 | 40.31146 | FALSE | FALSE |
| LOC107073362 | gamma-tubulin complex component 3 hom | 0.03218 | 40.32122 | FALSE | FALSE |
| LOC107070705 | uncharacterized protein LOC107070705 | 0.03173 | 40.33955 | FALSE | FALSE |
| LOC107069423 | serine/threonine-protein kinase D3 | 0.03268 | 40.34712 | FALSE | FALSE |
| LOC107069196 | molybdenum cofactor sulfurase 1 | 0.03162 | 40.37056 | FALSE | FALSE |
| LOC107073087 | importin-13 | 0.03242 | 40.37088 | FALSE | FALSE |
| LOC107071321 | uncharacterized protein LOC107071321 | 0.03055 | 40.37669 | FALSE | FALSE |
| LOC107066922 | leupaxin | 0.03051 | 40.39744 | FALSE | FALSE |
| LOC107063651 | ankyrin repeat domain-containing protein 1 | 0.03316 | 40.43015 | FALSE | FALSE |
| LOC107064007 | eukaryotic translation initiation factor 4E ty | 0.03324 | 40.44793 | FALSE | FALSE |
| LOC107070566 | mitochondrial 2-oxodicarboxylate carrier | 0.03251 | 40.45069 | FALSE | FALSE |
| LOC107073147 | receptor-mediated endocytosis protein 6 hc | 0.03221 | 40.46853 | FALSE | FALSE |
| LOC107065893 | transcription factor AP-2-epsilon | 0.03267 | 40.48159 | FALSE | FALSE |
| LOC107072432 | uncharacterized protein LOC107072432 | 0.03146 | 40.50073 | FALSE | FALSE |
| LOC107064140 | telomerase-binding protein EST1A | 0.03206 | 40.51467 | FALSE | FALSE |
| LOC107072356 | LOW QUALITY PROTEIN: protein polybromo | 0.03184 | 40.51755 | FALSE | FALSE |
| LOC107064372 | E3 ubiquitin-protein ligase TRIM71-like | 0.03175 | 40.52399 | FALSE | FALSE |
| LOC107074688 | nuclear distribution protein nudE-like 1-B | 0.03265 | 40.53753 | FALSE | FALSE |
| LOC107068423 | PDZ domain-containing protein GIPC3 | 0.03176 | 40.54342 | FALSE | FALSE |
| LOC107065559 | charged multivesicular body protein 7 | 0.03119 | 40.54475 | FALSE | FALSE |
| LOC107070939 | serine/threonine-protein kinase 10 | 0.03294 | 40.54931 | FALSE | FALSE |
| LOC107069012 | voltage-gated potassium channel subunit be | 0.03209 | 40.55666 | FALSE | FALSE |
| LOC107065552 | uncharacterized protein PF11_0213-like | 0.03282 | 40.55766 | FALSE | FALSE |
| LOC107065905 | enkurin | 0.03271 | 40.59947 | FALSE | FALSE |
| LOC107065629 | uncharacterized protein LOC107065629 | 0.03272 | 40.60115 | FALSE | FALSE |
| LOC107069860 | glucose dehydrogenase [FAD, quinone | 0.03305 | 40.60634 | FALSE | FALSE |
| Trnad-guc | Trnad-guc | 0.03262 | 40.61602 | FALSE | FALSE |
| LOC107070880 | tetraspanin-31-A | 0.0319 | 40.62793 | FALSE | FALSE |
| LOC107072880 | uncharacterized protein LOC107072880 | 0.03249 | 40.67247 | FALSE | FALSE |
| LOC107064030 | FMRFamide-related peptides-like | 0.03253 | 40.68156 | FALSE | FALSE |
| LOC107063879 | threonylcarbamoyladenosine tRNA methyltl | 0.03017 | 40.69971 | FALSE | FALSE |
| LOC107065569 | transcription elongation factor SPT4 | 0.0327 | 40.70229 | FALSE | FALSE |
| LOC107073005 | uncharacterized protein LOC107073005 | 0.03345 | 40.71552 | FALSE | FALSE |
| LOC107073061 | repressor of RNA polymerase III transcriptio | 0.03211 | 40.7159 | FALSE | FALSE |
| LOC107073401 | pentatricopeptide repeat-containing proteir | 0.03264 | 40.71867 | FALSE | FALSE |
| LOC107069842 | 15 kDa selenoprotein | 0.03225 | 40.73385 | FALSE | FALSE |
| LOC107073171 | reactive oxygen species modulator 1 | 0.03251 | 40.7619 | FALSE | FALSE |
| LOC107071726 | uncharacterized protein LOC107071726 | 0.03097 | 40.76504 | FALSE | FALSE |
| LOC107067869 | peroxisomal acyl-coenzyme A oxidase 3 | 0.03188 | 40.76739 | FALSE | FALSE |
| LOC107070912 | WD repeat-containing protein 36 | 0.03167 | 40.76966 | FALSE | FALSE |
| LOC107070473 | uncharacterized protein LOC107070473 | 0.03314 | 40.7786 | FALSE | FALSE |
| LOC107070539 | uncharacterized protein LOC107070539 | 0.03197 | 40.78068 | FALSE | FALSE |
| LOC107073937 | WD repeat-containing protein 6 | 0.03235 | 40.7852 | FALSE | FALSE |
| LOC107071484 | protein PF14_0175-like | 0.03301 | 40.78877 | FALSE | FALSE |
| LOC107074676 | zinc finger protein 2-like | 0.03208 | 40.78927 | FALSE | FALSE |
| LOC107067908 | odorant receptor 43a-like | 0.03055 | 40.7948 | FALSE | FALSE |

| | | | | | |
|---|---|---|---|---|---|
| LOC107065722 | zinc finger BED domain-containing protein 1 | 0.03254 | 40.79636 | FALSE | FALSE |
| LOC107066986 | monocarboxylate transporter 10 | 0.03171 | 40.79673 | FALSE | FALSE |
| LOC107073818 | LIM/homeobox protein Lhx3 | 0.03202 | 40.80649 | FALSE | FALSE |
| LOC107073617 | putative fatty acyl-CoA reductase CG5065 | 0.03239 | 40.8106 | FALSE | FALSE |
| LOC107064721 | suppressor APC domain-containing protein 1 | 0.03215 | 40.81723 | FALSE | FALSE |
| LOC107066220 | uncharacterized protein LOC107066220 | 0.03168 | 40.81726 | FALSE | FALSE |
| LOC107066219 | excitatory amino acid transporter | 0.03184 | 40.8244 | FALSE | FALSE |
| LOC107068458 | phosphatidylserine synthase 1 | 0.03201 | 40.83938 | FALSE | FALSE |
| LOC107072938 | RNA polymerase II subunit A C-terminal don | 0.0321 | 40.85797 | FALSE | FALSE |
| LOC107070623 | uncharacterized protein LOC107070623 | 0.03172 | 40.87109 | FALSE | FALSE |
| LOC107065831 | E3 ubiquitin-protein ligase MSL2 | 0.03215 | 40.90239 | FALSE | FALSE |
| LOC107074052 | uncharacterized protein LOC107074052 | 0.03063 | 40.91545 | FALSE | FALSE |
| LOC107073936 | nuclear pore membrane glycoprotein 210 | 0.03163 | 40.9271 | FALSE | FALSE |
| LOC107073236 | vacuolar protein sorting-associated protein | 0.03193 | 40.92969 | FALSE | FALSE |
| LOC107064101 | calcium-independent phospholipase A2-gan | 0.03194 | 40.95317 | FALSE | FALSE |
| LOC107068327 | cystathionine beta-synthase | 0.03236 | 40.95667 | FALSE | FALSE |
| LOC107072791 | uncharacterized protein LOC107072791 | 0.03241 | 40.96092 | FALSE | FALSE |
| LOC107069118 | putative ATP-dependent RNA helicase TDRD | 0.03195 | 40.98292 | FALSE | FALSE |
| LOC107070970 | cytochrome b561 domain-containing protei | 0.03283 | 40.99711 | FALSE | FALSE |
| LOC107074447 | arrestin homolog | 0.03184 | 41.00015 | FALSE | FALSE |
| LOC107067637 | uncharacterized protein LOC107067637 | 0.03131 | 41.00156 | FALSE | FALSE |
| LOC107067962 | ribosomal L1 domain-containing protein CG | 0.0315 | 41.01356 | FALSE | FALSE |
| LOC107073844 | 40S ribosomal protein S27-like | 0.03157 | 41.03565 | FALSE | FALSE |
| LOC107064109 | transferrin | 0.03015 | 41.05214 | FALSE | FALSE |
| LOC107068644 | uncharacterized protein LOC107068644 | 0.03273 | 41.06577 | FALSE | FALSE |
| LOC107067006 | kinase suppressor of Ras 2 | 0.03314 | 41.08312 | FALSE | FALSE |
| LOC107068252 | scaffold protein salvador-like | 0.03145 | 41.09084 | FALSE | FALSE |
| LOC107064681 | eukaryotic translation initiation factor 3 sub | 0.03134 | 41.09092 | FALSE | FALSE |
| LOC107073553 | cytochrome P450 4C1-like | 0.03149 | 41.09591 | FALSE | FALSE |
| LOC107065920 | nucleosome-remodeling factor subunit NUR | 0.03215 | 41.09814 | FALSE | FALSE |
| LOC107071972 | endochitinase EP3-like | 0.03219 | 41.10351 | FALSE | FALSE |
| LOC107070504 | uncharacterized protein LOC107070504 | 0.03124 | 41.12762 | FALSE | FALSE |
| LOC107068712 | nose resistant to fluoxetine protein 6-like | 0.03141 | 41.12993 | FALSE | FALSE |
| LOC107070499 | cilia- and flagella-associated protein 69-like | 0.03139 | 41.13379 | FALSE | FALSE |
| LOC107070419 | membrane-associated progesterone recepto | 0.03257 | 41.13684 | FALSE | FALSE |
| LOC107066682 | uncharacterized protein LOC107066682 | 0.03211 | 41.13758 | FALSE | FALSE |
| LOC107068519 | uncharacterized protein LOC107068519 | 0.0318 | 41.15433 | FALSE | FALSE |
| LOC107067605 | tetratricopeptide repeat protein 36 homolo | 0.03206 | 41.1554 | FALSE | FALSE |
| LOC107071332 | uncharacterized protein LOC107071332 | 0.03229 | 41.17108 | FALSE | FALSE |
| LOC107070422 | eukaryotic translation initiation factor 3 sub | 0.0313 | 41.17657 | FALSE | FALSE |
| LOC107069486 | uncharacterized protein LOC107069486 | 0.03199 | 41.18006 | FALSE | FALSE |
| LOC107066102 | regulating synaptic membrane exocytosis pr | 0.03131 | 41.19169 | FALSE | FALSE |
| LOC107066320 | lysyl oxidase homolog 2B | 0.03188 | 41.19181 | FALSE | FALSE |
| LOC107068226 | apoptosis-inducing factor 1, mitochondrial | 0.03295 | 41.20448 | FALSE | FALSE |
| LOC107068980 | mitochondrial inner membrane protease AT | 0.03292 | 41.21061 | FALSE | FALSE |
| LOC107069128 | trypsin-1-like | 0.03102 | 41.22199 | FALSE | FALSE |
| LOC107063685 | pachytene checkpoint protein 2 homolog | 0.03178 | 41.238 | FALSE | FALSE |
| LOC107071115 | mitogen-activated protein kinase kinase kin | 0.03195 | 41.23948 | FALSE | FALSE |
| LOC107073911 | uncharacterized protein LOC107073911 | 0.03056 | 41.24308 | FALSE | FALSE |
| LOC107067819 | PRA1 family protein 3 | 0.03139 | 41.24504 | FALSE | FALSE |

| | | | | | |
|---|---|---|---|---|---|
| LOC107070393 | bestrophin-4 | 0.0322 | 41.26075 | FALSE | FALSE |
| LOC107068376 | acylphosphatase-2 | 0.03224 | 41.26758 | FALSE | FALSE |
| LOC107072790 | 5-hydroxytryptamine receptor 1 | 0.03181 | 41.27248 | FALSE | FALSE |
| LOC107065005 | uncharacterized protein LOC107065005 | 0.03244 | 41.30619 | FALSE | FALSE |
| LOC107071748 | dynein regulatory complex protein 1 homol | 0.03287 | 41.31312 | FALSE | FALSE |
| LOC107064254 | hsp70-binding protein 1-like | 0.03159 | 41.32748 | FALSE | FALSE |
| LOC107065808 | protein yippee-like 1 | 0.03231 | 41.34193 | FALSE | FALSE |
| LOC107064288 | cytosolic carboxypeptidase-like protein 5 | 0.03266 | 41.34414 | FALSE | FALSE |
| LOC107069651 | autophagy-related protein 2 homolog B | 0.0326 | 41.36392 | FALSE | FALSE |
| LOC107064423 | toll-interacting protein A-like | 0.03135 | 41.36648 | FALSE | FALSE |
| LOC107072117 | extensin | 0.03194 | 41.37801 | FALSE | FALSE |
| LOC107068324 | cytochrome c oxidase assembly factor 6 hor | 0.033 | 41.38892 | FALSE | FALSE |
| LOC107064645 | uncharacterized protein LOC107064645 | 0.03143 | 41.40361 | FALSE | FALSE |
| LOC107068996 | uncharacterized protein LOC107068996 | 0.03137 | 41.42409 | FALSE | FALSE |
| LOC107072234 | calcineurin-binding protein cabin-1-like | 0.03174 | 41.42835 | FALSE | FALSE |
| LOC107071644 | fatty-acid amide hydrolase 2-B-like | 0.03311 | 41.45443 | FALSE | FALSE |
| LOC107072746 | uncharacterized protein LOC107072746 | 0.03189 | 41.47728 | FALSE | FALSE |
| LOC107066020 | uncharacterized protein MAL13P1.304-like | 0.03153 | 41.48164 | FALSE | FALSE |
| LOC107065670 | ubiquitin carboxyl-terminal hydrolase CYLD | 0.03245 | 41.4918 | FALSE | FALSE |
| LOC107072381 | juvenile hormone epoxide hydrolase 1-like | 0.03212 | 41.49297 | FALSE | FALSE |
| LOC107063697 | transient receptor potential channel pyrexia | 0.03364 | 41.52088 | FALSE | FALSE |
| LOC107069719 | ADP-ribosylation factor-like protein 2-bindir | 0.03187 | 41.52434 | FALSE | FALSE |
| LOC107065188 | protein yellow | 0.03093 | 41.53022 | FALSE | FALSE |
| LOC107063613 | uncharacterized protein LOC107063613 | 0.03085 | 41.53379 | FALSE | FALSE |
| LOC107064559 | potassium/sodium hyperpolarization-activa | 0.03273 | 41.53673 | FALSE | FALSE |
| LOC107068482 | growth arrest-specific protein 2-like | 0.03274 | 41.53943 | FALSE | FALSE |
| LOC107064150 | WD repeat-containing protein 48 | 0.03118 | 41.55751 | FALSE | FALSE |
| LOC107072129 | E3 ubiquitin-protein ligase TRAIP | 0.03111 | 41.55913 | FALSE | FALSE |
| LOC107069261 | inner centromere protein A-like | 0.03074 | 41.57165 | FALSE | FALSE |
| LOC107069294 | serrate RNA effector molecule homolog | 0.03209 | 41.57452 | FALSE | FALSE |
| LOC107064152 | 4-coumarate--CoA ligase 1-like | 0.032 | 41.57625 | FALSE | FALSE |
| LOC107065442 | leukocyte receptor cluster member 8 homo | 0.03292 | 41.59211 | FALSE | FALSE |
| LOC107067693 | uncharacterized protein LOC107067693 | 0.03223 | 41.60054 | FALSE | FALSE |
| LOC107070547 | beta-amyloid-like protein | 0.03203 | 41.60192 | FALSE | FALSE |
| LOC107070676 | guanine nucleotide-binding protein subunit | 0.0307 | 41.60406 | FALSE | FALSE |
| LOC107069335 | SCY1-like protein 2 | 0.03107 | 41.6124 | FALSE | FALSE |
| LOC107070758 | nucleolar protein 12 | 0.03136 | 41.64106 | FALSE | FALSE |
| LOC107065169 | 40S ribosomal protein S12, mitochondrial | 0.03153 | 41.64625 | FALSE | FALSE |
| LOC107067258 | NADH dehydrogenase [ubiquinone | 0.03163 | 41.67895 | FALSE | FALSE |
| LOC107068624 | cysteine dioxygenase type 1 | 0.03297 | 41.68735 | FALSE | FALSE |
| LOC107068219 | TLD domain-containing protein 1 | 0.03264 | 41.6879 | FALSE | FALSE |
| LOC107065052 | igLON family member 5-like | 0.0321 | 41.69287 | FALSE | FALSE |
| LOC107067894 | uncharacterized protein LOC107067894 | 0.03182 | 41.71222 | FALSE | FALSE |
| LOC107068143 | neuronal calcium sensor 2 | 0.03195 | 41.73174 | FALSE | FALSE |
| LOC107073040 | 2,3-dehydroadipyl-CoA hydratase-like | 0.03188 | 41.76313 | FALSE | FALSE |
| LOC107064810 | heat shock protein 75 kDa, mitochondrial | 0.03148 | 41.77743 | FALSE | FALSE |
| LOC107070020 | homeobox protein 5-like | 0.03177 | 41.77831 | FALSE | FALSE |
| LOC107065208 | regulator of nonsense transcripts 3A | 0.0312 | 41.78806 | FALSE | FALSE |
| LOC107067803 | probable small nuclear ribonucleoprotein G | 0.03178 | 41.7987 | FALSE | FALSE |
| LOC107066544 | uncharacterized protein LOC107066544 | 0.03194 | 41.81684 | FALSE | FALSE |

| | | | | | |
|---|---|---|---|---|---|
| LOC107069663 | phosphopantothenoylcysteine decarboxylas | 0.03113 | 41.82397 | FALSE | FALSE |
| LOC107068059 | ATP-binding cassette sub-family B member | 0.03122 | 41.82992 | FALSE | FALSE |
| LOC107069488 | elongation factor 1-alpha-like | 0.03207 | 41.83111 | FALSE | FALSE |
| LOC107067219 | octopamine receptor beta-3R-like | 0.03067 | 41.84358 | FALSE | FALSE |
| LOC107063841 | uncharacterized protein LOC107063841 | 0.03121 | 41.85757 | FALSE | FALSE |
| LOC107066510 | exosome complex exonuclease RRP44 | 0.03176 | 41.8675 | FALSE | FALSE |
| LOC107072547 | short-chain dehydrogenase/reductase 3-like | 0.02993 | 41.87555 | FALSE | FALSE |
| LOC107066361 | equilibrative nucleoside transporter 3 | 0.03143 | 41.89737 | FALSE | FALSE |
| LOC107064340 | UPF0415 protein C7orf25 homolog | 0.03177 | 41.9027 | FALSE | FALSE |
| LOC107070300 | geranylgeranyl transferase type-2 subunit a | 0.03162 | 41.91033 | FALSE | FALSE |
| LOC107070973 | sodium- and chloride-dependent glycine tra | 0.03217 | 41.93757 | FALSE | FALSE |
| LOC107074262 | transcription factor SOX-2-like | 0.02969 | 41.93981 | FALSE | FALSE |
| LOC107065892 | isoleucine--tRNA ligase, cytoplasmic | 0.03104 | 41.95828 | FALSE | FALSE |
| LOC107067974 | uncharacterized protein LOC107067974 | 0.03208 | 41.96074 | FALSE | FALSE |
| LOC107069845 | uncharacterized protein LOC107069845 | 0.03096 | 41.96407 | FALSE | FALSE |
| LOC107063929 | TFIIH basal transcription factor complex heli | 0.03259 | 41.98 | FALSE | FALSE |
| LOC107068705 | endocuticle structural glycoprotein SgAbd-2 | 0.0324 | 41.98637 | FALSE | FALSE |
| LOC107073734 | uncharacterized protein LOC107073734 | 0.03391 | 42.00118 | FALSE | FALSE |
| LOC107068033 | 4-coumarate--CoA ligase 1-like | 0.0328 | 42.00957 | FALSE | FALSE |
| LOC107073702 | protein toll-like | 0.03183 | 42.01816 | FALSE | FALSE |
| LOC107071052 | protein FAM32A | 0.03152 | 42.02426 | FALSE | FALSE |
| LOC107063647 | C2 domain-containing protein 5 | 0.0322 | 42.03423 | FALSE | FALSE |
| LOC107074080 | uncharacterized protein LOC107074080 | 0.03052 | 42.03754 | FALSE | FALSE |
| LOC107063922 | uncharacterized protein LOC107063922 | 0.03249 | 42.04902 | FALSE | FALSE |
| LOC107068486 | DDB1- and CUL4-associated factor 6-like | 0.03084 | 42.05253 | FALSE | FALSE |
| LOC107067434 | group XV phospholipase A2-like | 0.03167 | 42.05547 | FALSE | FALSE |
| LOC107072303 | probable 28S ribosomal protein S26, mitoch | 0.03195 | 42.05768 | FALSE | FALSE |
| LOC107065966 | hepatic leukemia factor | 0.03114 | 42.07626 | FALSE | FALSE |
| LOC107064712 | zinc finger protein ZIC 4-like | 0.03157 | 42.07655 | FALSE | FALSE |
| LOC107063961 | enhancer of split mbeta protein-like | 0.0308 | 42.08868 | FALSE | FALSE |
| LOC107069202 | uncharacterized protein LOC107069202 | 0.03193 | 42.08869 | FALSE | FALSE |
| LOC107072069 | uncharacterized protein C7orf50 homolog | 0.03112 | 42.09726 | FALSE | FALSE |
| LOC107071551 | uncharacterized protein LOC107071551 | 0.03063 | 42.10666 | FALSE | FALSE |
| LOC107065806 | uncharacterized protein LOC107065806 | 0.03234 | 42.1106 | FALSE | FALSE |
| LOC107069988 | phosphatidylcholine:ceramide cholinephosp | 0.03239 | 42.1151 | FALSE | FALSE |
| LOC107072570 | cilia- and flagella-associated protein 57 | 0.03069 | 42.12136 | FALSE | FALSE |
| LOC107065927 | transitional endoplasmic reticulum ATPase | 0.03278 | 42.12891 | FALSE | FALSE |
| LOC107071031 | receptor-type tyrosine-protein phosphatase | 0.03181 | 42.13944 | FALSE | FALSE |
| LOC107072995 | F-box/LRR-repeat protein 16 | 0.03072 | 42.14194 | FALSE | FALSE |
| LOC107068172 | proteasome subunit beta type-4 | 0.03136 | 42.14507 | FALSE | FALSE |
| LOC107071853 | UMP-CMP kinase | 0.03331 | 42.15562 | FALSE | FALSE |
| LOC107064419 | regulatory protein MIG1-like | 0.03141 | 42.15959 | FALSE | FALSE |
| LOC107067062 | uncharacterized protein LOC107067062 | 0.03112 | 42.16761 | FALSE | FALSE |
| LOC107068669 | trifunctional enzyme subunit beta, mitochor | 0.03219 | 42.17368 | FALSE | FALSE |
| LOC107074144 | transient receptor potential cation channel | 0.03267 | 42.18279 | FALSE | FALSE |
| LOC107072732 | zinc finger and BTB domain-containing prote | 0.03286 | 42.19681 | FALSE | FALSE |
| LOC107067144 | uncharacterized protein LOC107067144 | 0.03283 | 42.19735 | FALSE | FALSE |
| LOC107070176 | peptidyl-prolyl cis-trans isomerase E-like | 0.03429 | 42.19953 | FALSE | FALSE |
| LOC107073740 | microfibrillar-associated protein 1 | 0.03197 | 42.2031 | FALSE | FALSE |
| LOC107070695 | uncharacterized protein LOC107070695 | 0.03221 | 42.22039 | FALSE | FALSE |

| | | | | | |
|---|---|---|---|---|---|
| LOC107063764 | bud site selection protein 4-like | 0.03115 | 42.24045 | FALSE | FALSE |
| LOC107070261 | serine/threonine-protein phosphatase 4 reg | 0.03106 | 42.25595 | FALSE | FALSE |
| LOC107066560 | venom carboxylesterase-6-like | 0.03323 | 42.26608 | FALSE | FALSE |
| LOC107069303 | UPF0528 protein CG10038 | 0.03166 | 42.26716 | FALSE | FALSE |
| LOC107070142 | NADH dehydrogenase [ubiquinone | 0.03212 | 42.27164 | FALSE | FALSE |
| LOC107064124 | probable phospholipid hydroperoxide gluta | 0.03108 | 42.2755 | FALSE | FALSE |
| LOC107070436 | WD repeat-containing protein 34-like | 0.03149 | 42.27989 | FALSE | FALSE |
| LOC107064788 | asparagine--tRNA ligase, cytoplasmic | 0.03338 | 42.29808 | FALSE | FALSE |
| LOC107065254 | septin-1 | 0.03184 | 42.35108 | FALSE | FALSE |
| LOC107067250 | leucine-rich repeat-containing protein 49 | 0.0319 | 42.35224 | FALSE | FALSE |
| LOC107073953 | probable dolichyl pyrophosphate Glc1Man9 | 0.0319 | 42.36029 | FALSE | FALSE |
| LOC107074591 | acylglycerol kinase, mitochondrial | 0.03155 | 42.36284 | FALSE | FALSE |
| LOC107072040 | armadillo repeat-containing protein 1-like | 0.03317 | 42.37008 | FALSE | FALSE |
| LOC107072688 | carboxypeptidase N subunit 2-like | 0.03218 | 42.37712 | FALSE | FALSE |
| LOC107064602 | transcription initiation factor TFIID subunit | 0.03084 | 42.39486 | FALSE | FALSE |
| LOC107065420 | inhibitor of growth protein 3 | 0.03136 | 42.39759 | FALSE | FALSE |
| LOC107072335 | 1-phosphatidylinositol 4,5-bisphosphate ph | 0.03112 | 42.40212 | FALSE | FALSE |
| LOC107070976 | INO80 complex subunit D | 0.03093 | 42.40792 | FALSE | FALSE |
| LOC107066852 | protein spire | 0.03149 | 42.41106 | FALSE | FALSE |
| LOC107069299 | GMP reductase 1-like | 0.02995 | 42.41157 | FALSE | FALSE |
| LOC107069401 | semaphorin-2A | 0.03126 | 42.41392 | FALSE | FALSE |
| LOC107074190 | protein diaphanous | 0.03191 | 42.41744 | FALSE | FALSE |
| LOC107067108 | uncharacterized protein LOC107067108 | 0.03067 | 42.44705 | FALSE | FALSE |
| LOC107064107 | uncharacterized protein LOC107064107 | 0.03129 | 42.45701 | FALSE | FALSE |
| LOC107074045 | myb-like protein K | 0.032 | 42.46866 | FALSE | FALSE |
| LOC107067638 | synaptic vesicle glycoprotein 2B-like | 0.03129 | 42.47107 | FALSE | FALSE |
| LOC107067994 | AN1-type zinc finger protein 1-like | 0.03223 | 42.4757 | FALSE | FALSE |
| LOC107074141 | nucleolar protein 6 | 0.03177 | 42.4766 | FALSE | FALSE |
| LOC107070796 | uncharacterized protein LOC107070796 | 0.02994 | 42.48287 | FALSE | FALSE |
| LOC107066278 | MATH and LRR domain-containing protein P | 0.03109 | 42.48925 | FALSE | FALSE |
| LOC107064374 | larval cuticle protein A2B-like | 0.03289 | 42.4922 | FALSE | FALSE |
| LOC107065696 | uncharacterized protein LOC107065696 | 0.03131 | 42.49441 | FALSE | FALSE |
| LOC107064835 | X-linked retinitis pigmentosa GTPase regula | 0.03016 | 42.51565 | FALSE | FALSE |
| LOC107074227 | uncharacterized protein LOC107074227 | 0.03052 | 42.51852 | FALSE | FALSE |
| LOC107072102 | N-acetylglucosamine-1-phosphotransferase | 0.0316 | 42.5286 | FALSE | FALSE |
| LOC107066960 | elongation of very long chain fatty acids pro | 0.03122 | 42.5443 | FALSE | FALSE |
| LOC107066637 | dnaJ homolog subfamily C member 4-like | 0.03095 | 42.54765 | FALSE | FALSE |
| LOC107073778 | uncharacterized protein LOC107073778 | 0.03236 | 42.54819 | FALSE | FALSE |
| LOC107064110 | uncharacterized protein LOC107064110 | 0.03215 | 42.54882 | FALSE | FALSE |
| LOC107069704 | uncharacterized protein LOC107069704 | 0.03196 | 42.55316 | FALSE | FALSE |
| LOC107072685 | uncharacterized protein LOC107072685 | 0.0304 | 42.55849 | FALSE | FALSE |
| LOC107070700 | uncharacterized protein LOC107070700 | 0.03218 | 42.56836 | FALSE | FALSE |
| LOC107073210 | uncharacterized protein LOC107073210 | 0.02995 | 42.5718 | FALSE | FALSE |
| LOC107066825 | protein transport protein Sec16A | 0.03142 | 42.57852 | FALSE | FALSE |
| LOC107072197 | uncharacterized protein LOC107072197 | 0.03167 | 42.5874 | FALSE | FALSE |
| LOC107065148 | alpha-tocopherol transfer protein-like | 0.03116 | 42.59218 | FALSE | FALSE |
| LOC107074197 | uncharacterized protein LOC107074197 | 0.03055 | 42.61412 | FALSE | FALSE |
| LOC107067218 | discoidin domain-containing receptor 2 | 0.0307 | 42.61834 | FALSE | FALSE |
| LOC107074511 | band 7 protein AGAP004871 | 0.03034 | 42.61989 | FALSE | FALSE |
| LOC107072324 | uncharacterized protein LOC107072324 | 0.03101 | 42.62732 | FALSE | FALSE |

| | | | | | |
|---|---|---|---|---|---|
| LOC107068593 | uncharacterized protein LOC107068593 | 0.03277 | 42.63473 | FALSE | FALSE |
| LOC107066698 | partitioning defective 3 homolog | 0.03196 | 42.64211 | FALSE | FALSE |
| LOC107066386 | cytoplasmic tRNA 2-thiolation protein 2-A | 0.03086 | 42.64358 | FALSE | FALSE |
| LOC107065964 | multidrug resistance-associated protein 4-lil | 0.03038 | 42.68023 | FALSE | FALSE |
| LOC107068057 | putative leucine-rich repeat-containing prot | 0.03021 | 42.70814 | FALSE | FALSE |
| LOC107073090 | 39S ribosomal protein L52, mitochondrial | 0.03153 | 42.71348 | FALSE | FALSE |
| LOC107066974 | LYR motif-containing protein 5 | 0.03224 | 42.71759 | FALSE | FALSE |
| LOC107066695 | formin-2-like | 0.03034 | 42.72385 | FALSE | FALSE |
| LOC107068998 | nucleolar protein 14 homolog | 0.03024 | 42.7471 | FALSE | FALSE |
| LOC107065465 | protein krasavietz | 0.03142 | 42.76202 | FALSE | FALSE |
| LOC107065903 | inositol polyphosphate multikinase | 0.03138 | 42.76531 | FALSE | FALSE |
| LOC107069147 | uncharacterized protein LOC107069147 | 0.03155 | 42.78338 | FALSE | FALSE |
| LOC107066201 | ER membrane protein complex subunit 8/9 | 0.03198 | 42.7962 | FALSE | FALSE |
| LOC107072665 | putative Dol-P-Glc:Glc(2)Man(9)GlcNAc(2)-F | 0.03353 | 42.80329 | FALSE | FALSE |
| LOC107071067 | elongation of very long chain fatty acids pro | 0.03248 | 42.81213 | FALSE | FALSE |
| LOC107071946 | uncharacterized protein LOC107071946 | 0.03089 | 42.81429 | FALSE | FALSE |
| LOC107070899 | uncharacterized protein LOC107070899 | 0.03175 | 42.82912 | FALSE | FALSE |
| LOC107072215 | uncharacterized protein LOC107072215 | 0.03243 | 42.84259 | FALSE | FALSE |
| LOC107066089 | F-box protein SKIP2-like | 0.03058 | 42.86799 | FALSE | FALSE |
| LOC107072138 | aquaporin AQPAn.G | 0.03233 | 42.87027 | FALSE | FALSE |
| LOC107069224 | zinc finger protein 593 homolog | 0.03148 | 42.89757 | FALSE | FALSE |
| LOC107065925 | protein sidekick | 0.03148 | 42.92968 | FALSE | FALSE |
| LOC107074360 | clustered mitochondria protein homolog | 0.03037 | 42.93242 | FALSE | FALSE |
| LOC107072274 | uncharacterized protein LOC107072274 | 0.032 | 42.93394 | FALSE | FALSE |
| LOC107072337 | uncharacterized protein LOC107072337 | 0.03149 | 42.95509 | FALSE | FALSE |
| LOC107072819 | protein piccolo | 0.03145 | 42.9556 | FALSE | FALSE |
| LOC107069383 | uncharacterized protein LOC107069383 | 0.03101 | 42.96614 | FALSE | FALSE |
| LOC107072327 | transient receptor potential channel pyrexia | 0.03071 | 42.97542 | FALSE | FALSE |
| LOC107065990 | S-adenosylmethionine mitochondrial carrier | 0.03143 | 42.97646 | FALSE | FALSE |
| LOC107068439 | myb-like protein X | 0.03163 | 42.98218 | FALSE | FALSE |
| LOC107071704 | MOXD1 homolog 2 | 0.03099 | 42.99058 | FALSE | FALSE |
| LOC107070004 | glucose-induced degradation protein 4 hom | 0.02997 | 43.00687 | FALSE | FALSE |
| LOC107066549 | uncharacterized protein LOC107066549 | 0.03167 | 43.00754 | FALSE | FALSE |
| LOC107066564 | putative uncharacterized protein DDB_G028 | 0.03199 | 43.01343 | FALSE | FALSE |
| LOC107073160 | putative uncharacterized protein DDB_G028 | 0.03027 | 43.01418 | FALSE | FALSE |
| LOC107064847 | opsin, blue-sensitive | 0.03155 | 43.02374 | FALSE | FALSE |
| LOC107063662 | neprilysin-2-like | 0.03122 | 43.02555 | FALSE | FALSE |
| LOC107064492 | RPA-interacting protein-like | 0.03134 | 43.02612 | FALSE | FALSE |
| LOC107069666 | uncharacterized protein LOC107069666 | 0.03234 | 43.03314 | FALSE | FALSE |
| LOC107073196 | probable Golgi SNAP receptor complex men | 0.03148 | 43.03837 | FALSE | FALSE |
| LOC107068911 | odorant receptor 13a-like | 0.03162 | 43.03931 | FALSE | FALSE |
| LOC107064205 | CDP-diacylglycerol--glycerol-3-phosphate 3- | 0.03264 | 43.04378 | FALSE | FALSE |
| LOC107064294 | protein 4.1 homolog | 0.0305 | 43.04579 | FALSE | FALSE |
| LOC107064944 | uncharacterized protein LOC107064944 | 0.03118 | 43.0465 | FALSE | FALSE |
| LOC107073548 | uncharacterized protein LOC107073548 | 0.03053 | 43.08525 | FALSE | FALSE |
| LOC107073768 | uncharacterized protein LOC107073768 | 0.031 | 43.08931 | FALSE | FALSE |
| LOC107069336 | protein KRI1 homolog | 0.03103 | 43.0906 | FALSE | FALSE |
| LOC107074231 | uncharacterized protein LOC107074231 | 0.03267 | 43.12128 | FALSE | FALSE |
| LOC107066791 | cell division cycle 7-related protein kinase-li | 0.03109 | 43.13065 | FALSE | FALSE |
| LOC107070792 | uncharacterized protein LOC107070792 | 0.03247 | 43.13266 | FALSE | FALSE |

| | | | | | |
|---|---|---|---|---|---|
| LOC107068180 | mitotic spindle assembly checkpoint proteir | 0.0314 | 43.14312 | FALSE | FALSE |
| LOC107071755 | zinc finger protein 391-like | 0.03133 | 43.14538 | FALSE | FALSE |
| LOC107064431 | cytoskeleton-associated protein 5 | 0.02952 | 43.15215 | FALSE | FALSE |
| LOC107072233 | origin recognition complex subunit 1 | 0.03123 | 43.17156 | FALSE | FALSE |
| LOC107064631 | putative inorganic phosphate cotransporter | 0.02984 | 43.17889 | FALSE | FALSE |
| LOC107066240 | probable G-protein coupled receptor B0563 | 0.03144 | 43.17889 | FALSE | FALSE |
| LOC107072518 | dnaJ homolog subfamily B member 12 | 0.03108 | 43.18323 | FALSE | FALSE |
| LOC107073798 | putative fatty acyl-CoA reductase CG5065 | 0.03129 | 43.18678 | FALSE | FALSE |
| LOC107064116 | transcription elongation factor B polypeptid | 0.03104 | 43.18904 | FALSE | FALSE |
| LOC107072781 | cilia- and flagella-associated protein 58-like | 0.02963 | 43.20041 | FALSE | FALSE |
| LOC107067506 | uncharacterized protein LOC107067506 | 0.03041 | 43.20077 | FALSE | FALSE |
| LOC107063657 | myotubularin-related protein 14 | 0.03058 | 43.20129 | FALSE | FALSE |
| LOC107070805 | ESF1 homolog | 0.03203 | 43.20589 | FALSE | FALSE |
| LOC107070032 | potassium channel subfamily T member 2 | 0.03123 | 43.20967 | FALSE | FALSE |
| LOC107064530 | E3 ubiquitin-protein ligase MARCH6 | 0.03137 | 43.21714 | FALSE | FALSE |
| LOC107068213 | lysophospholipase-like protein 1 | 0.03119 | 43.22126 | FALSE | FALSE |
| LOC107069665 | beta-glucuronidase-like | 0.0315 | 43.234 | FALSE | FALSE |
| LOC107065832 | protein roadkill-like | 0.03037 | 43.23696 | FALSE | FALSE |
| LOC107064322 | uncharacterized protein LOC107064322 | 0.03093 | 43.24273 | FALSE | FALSE |
| LOC107069668 | zinc finger HIT domain-containing protein 3 | 0.03163 | 43.25507 | FALSE | FALSE |
| LOC107071459 | uncharacterized protein LOC107071459 | 0.03086 | 43.26246 | FALSE | FALSE |
| LOC107071024 | uncharacterized protein LOC107071024 | 0.032 | 43.29371 | FALSE | FALSE |
| LOC107065473 | cytosol aminopeptidase | 0.03271 | 43.29937 | FALSE | FALSE |
| LOC107066936 | uncharacterized protein LOC107066936 | 0.03129 | 43.30851 | FALSE | FALSE |
| LOC107066596 | protein bicaudal D | 0.0315 | 43.31436 | FALSE | FALSE |
| LOC107073374 | phosphomannomutase 2 | 0.03073 | 43.32093 | FALSE | FALSE |
| LOC107067068 | uncharacterized protein LOC107067068 | 0.03024 | 43.33441 | FALSE | FALSE |
| LOC107069284 | protein Spindly-like | 0.03229 | 43.33771 | FALSE | FALSE |
| LOC107066812 | uncharacterized protein LOC107066812 | 0.0314 | 43.35457 | FALSE | FALSE |
| LOC107069584 | vacuolar protein sorting-associated protein | 0.03075 | 43.36649 | FALSE | FALSE |
| LOC107071458 | armadillo repeat-containing protein 5 | 0.03054 | 43.37045 | FALSE | FALSE |
| LOC107067847 | 39S ribosomal protein L30, mitochondrial | 0.03183 | 43.37071 | FALSE | FALSE |
| LOC107071502 | putative U5 small nuclear ribonucleoproteir | 0.03143 | 43.37688 | FALSE | FALSE |
| LOC107069660 | bolA-like protein DDB_G0274169 | 0.03139 | 43.37937 | FALSE | FALSE |
| LOC107071764 | uncharacterized protein LOC107071764 | 0.03109 | 43.40169 | FALSE | FALSE |
| LOC107071388 | uncharacterized protein LOC107071388 | 0.03137 | 43.42855 | FALSE | FALSE |
| LOC107069164 | uncharacterized protein LOC107069164 | 0.03143 | 43.43786 | FALSE | FALSE |
| LOC107071933 | uncharacterized protein LOC107071933 | 0.03168 | 43.46013 | FALSE | FALSE |
| LOC107074506 | FGFR1 oncogene partner 2 homolog | 0.03122 | 43.46485 | FALSE | FALSE |
| LOC107064593 | ribosome biogenesis protein NSA2 homolog | 0.03163 | 43.46495 | FALSE | FALSE |
| LOC107069693 | activating transcription factor 7-interacting | 0.03065 | 43.47005 | FALSE | FALSE |
| LOC107070024 | maltase 1-like | 0.03123 | 43.47372 | FALSE | FALSE |
| LOC107069858 | phosphatidylinositol 3-kinase catalytic subu | 0.03051 | 43.47618 | FALSE | FALSE |
| LOC107064127 | phosphoglycolate phosphatase-like | 0.03123 | 43.49456 | FALSE | FALSE |
| LOC107064548 | uncharacterized protein LOC107064548 | 0.03097 | 43.50495 | FALSE | FALSE |
| LOC107074504 | fatty acyl-CoA reductase 2-like | 0.03081 | 43.50994 | FALSE | FALSE |
| LOC107073104 | putative uncharacterized protein FLJ37770 | 0.03135 | 43.5191 | FALSE | FALSE |
| LOC107072755 | uncharacterized protein LOC107072755 | 0.03081 | 43.52141 | FALSE | FALSE |
| LOC107070117 | protein unc-50 homolog | 0.03039 | 43.52487 | FALSE | FALSE |
| LOC107073448 | uncharacterized protein LOC107073448 | 0.03166 | 43.53048 | FALSE | FALSE |

| | | | | | |
|---|---|---|---|---|---|
| LOC107072325 | succinyl-CoA ligase [ADP-forming | 0.03001 | 43.53087 | FALSE | FALSE |
| LOC107072981 | uncharacterized protein LOC107072981 | 0.03205 | 43.5581 | FALSE | FALSE |
| LOC107070274 | replication factor C subunit 1 | 0.03037 | 43.5782 | FALSE | FALSE |
| LOC107067785 | double-stranded RNA-specific editase Adar | 0.03046 | 43.58373 | FALSE | FALSE |
| LOC107073129 | ribosome biogenesis regulatory protein hon | 0.03055 | 43.61072 | FALSE | FALSE |
| LOC107073408 | uncharacterized protein LOC107073408 | 0.03107 | 43.62189 | FALSE | FALSE |
| LOC107068664 | THO complex subunit 7 homolog | 0.0316 | 43.62491 | FALSE | FALSE |
| LOC107065472 | uncharacterized protein LOC107065472 | 0.03076 | 43.62813 | FALSE | FALSE |
| LOC107072348 | RNA-binding protein 47-like | 0.03178 | 43.63534 | FALSE | FALSE |
| LOC107071966 | uncharacterized protein LOC107071966 | 0.03162 | 43.637 | FALSE | FALSE |
| LOC107070857 | uncharacterized protein LOC107070857 | 0.02988 | 43.65407 | FALSE | FALSE |
| LOC107073832 | perilipin-2-like | 0.03139 | 43.663 | FALSE | FALSE |
| LOC107071799 | protein SCAI | 0.03134 | 43.66593 | FALSE | FALSE |
| LOC107070682 | lachesin-like | 0.02978 | 43.67686 | FALSE | FALSE |
| LOC107064560 | eukaryotic translation initiation factor 5B | 0.03043 | 43.68261 | FALSE | FALSE |
| LOC107070251 | probable cytosolic oligopeptidase A | 0.03117 | 43.68383 | FALSE | FALSE |
| LOC107071837 | uncharacterized protein LOC107071837 | 0.03087 | 43.68554 | FALSE | FALSE |
| LOC107072444 | uncharacterized protein LOC107072444 | 0.02986 | 43.6948 | FALSE | FALSE |
| LOC107064417 | uncharacterized protein LOC107064417 | 0.03038 | 43.6963 | FALSE | FALSE |
| LOC107068275 | venom acid phosphatase Acph-1-like | 0.02992 | 43.69632 | FALSE | FALSE |
| LOC107069579 | bifunctional heparan sulfate N-deacetylase/ | 0.03023 | 43.70529 | FALSE | FALSE |
| LOC107068232 | elongation factor 1-gamma | 0.03076 | 43.75504 | FALSE | FALSE |
| LOC107065560 | MIP18 family protein CG30152 | 0.03119 | 43.76664 | FALSE | FALSE |
| LOC107066530 | angiopoietin-2 | 0.02986 | 43.76736 | FALSE | FALSE |
| LOC107066734 | uncharacterized protein LOC107066734 | 0.03053 | 43.76969 | FALSE | FALSE |
| LOC107068649 | sex-determining region Y protein-like | 0.03155 | 43.78141 | FALSE | FALSE |
| LOC107070021 | Na(+)/H(+) exchange regulatory cofactor NH | 0.03106 | 43.79275 | FALSE | FALSE |
| LOC107071159 | potassium channel subfamily K member 9-li | 0.02972 | 43.80843 | FALSE | FALSE |
| LOC107069738 | cytochrome c oxidase subunit NDUFA4 | 0.03109 | 43.82913 | FALSE | FALSE |
| LOC107071639 | prostatic acid phosphatase-like | 0.03114 | 43.8401 | FALSE | FALSE |
| LOC107066956 | WD repeat-containing protein 66-like | 0.03112 | 43.84167 | FALSE | FALSE |
| LOC107073853 | EKC/KEOPS complex subunit TPRKB-like | 0.03114 | 43.87431 | FALSE | FALSE |
| LOC107073957 | actin-related protein 2 | 0.03013 | 43.87618 | FALSE | FALSE |
| LOC107071300 | DNA ligase 1-like | 0.03084 | 43.88466 | FALSE | FALSE |
| LOC107068385 | deoxyhypusine hydroxylase | 0.03183 | 43.88673 | FALSE | FALSE |
| LOC107069270 | uncharacterized protein LOC107069270 | 0.03118 | 43.88924 | FALSE | FALSE |
| LOC107068402 | cleft lip and palate transmembrane protein | 0.03018 | 43.91362 | FALSE | FALSE |
| LOC107068393 | integrator complex subunit 1 | 0.03083 | 43.92038 | FALSE | FALSE |
| LOC107063782 | tachykinins | 0.0297 | 43.93293 | FALSE | FALSE |
| LOC107065416 | UNC93-like protein | 0.03075 | 43.94125 | FALSE | FALSE |
| LOC107067695 | dynein intermediate chain 2, ciliary | 0.02986 | 43.96549 | FALSE | FALSE |
| LOC107066203 | dual specificity protein phosphatase 22-like | 0.03101 | 43.98614 | FALSE | FALSE |
| LOC107073454 | kaptin-like | 0.02939 | 43.99461 | FALSE | FALSE |
| LOC107068309 | ran-binding protein 3 | 0.03104 | 44.00265 | FALSE | FALSE |
| LOC107068987 | uncharacterized protein LOC107068987 | 0.03187 | 44.01259 | FALSE | FALSE |
| LOC107066458 | E3 ubiquitin-protein ligase RNF34 | 0.02982 | 44.01624 | FALSE | FALSE |
| LOC107074415 | PI-PLC X domain-containing protein 3 | 0.03139 | 44.03505 | FALSE | FALSE |
| LOC107070089 | xylulose kinase | 0.03088 | 44.04075 | FALSE | FALSE |
| LOC107069925 | uncharacterized protein LOC107069925 | 0.03016 | 44.05866 | FALSE | FALSE |
| LOC107064199 | cell wall protein IFF7 | 0.03271 | 44.05898 | FALSE | FALSE |

| | | | | | |
|---|---|---|---|---|---|
| LOC107066091 | uncharacterized protein LOC107066091 | 0.03223 | 44.06603 | FALSE | FALSE |
| LOC107069639 | uncharacterized protein LOC107069639 | 0.02899 | 44.07336 | FALSE | FALSE |
| LOC107070910 | GTP-binding protein SAR1a | 0.03104 | 44.07377 | FALSE | FALSE |
| LOC107068461 | uncharacterized protein LOC107068461 | 0.033 | 44.07873 | FALSE | FALSE |
| LOC107073211 | uncharacterized protein LOC107073211 | 0.03068 | 44.07933 | FALSE | FALSE |
| LOC107068011 | uncharacterized protein LOC107068011 | 0.02949 | 44.08559 | FALSE | FALSE |
| LOC107072457 | probable asparagine--tRNA ligase, mitochor | 0.03014 | 44.09586 | FALSE | FALSE |
| LOC107068210 | mitogen-activated protein kinase kinase kin | 0.03014 | 44.10162 | FALSE | FALSE |
| LOC107066002 | gamma-glutamyltranspeptidase 1-like | 0.03039 | 44.12157 | FALSE | FALSE |
| LOC107071595 | TBC1 domain family member 5 homolog A-l | 0.03073 | 44.13189 | FALSE | FALSE |
| LOC107069901 | protein tramtrack, alpha isoform-like | 0.0304 | 44.15676 | FALSE | FALSE |
| LOC107071454 | acetylcholine receptor subunit beta-like 1 | 0.03082 | 44.16318 | FALSE | FALSE |
| LOC107066336 | testis-specific serine/threonine-protein kina | 0.03139 | 44.1765 | FALSE | FALSE |
| LOC107073928 | 3-hydroxy-3-methylglutaryl-coenzyme A rec | 0.02864 | 44.18181 | FALSE | FALSE |
| LOC107065529 | 39S ribosomal protein L48, mitochondrial | 0.03171 | 44.1915 | FALSE | FALSE |
| LOC107074100 | very low-density lipoprotein receptor-like | 0.03119 | 44.19362 | FALSE | FALSE |
| LOC107071142 | FK506-binding protein 2 | 0.03066 | 44.19639 | FALSE | FALSE |
| LOC107068374 | 60S acidic ribosomal protein P0 | 0.03051 | 44.2273 | FALSE | FALSE |
| LOC107072898 | dual specificity protein kinase splB-like | 0.03089 | 44.23136 | FALSE | FALSE |
| LOC107067083 | structural maintenance of chromosomes pro | 0.03022 | 44.24335 | FALSE | FALSE |
| LOC107066761 | uncharacterized protein LOC107066761 | 0.03081 | 44.25636 | FALSE | FALSE |
| LOC107074663 | UHRF1-binding protein 1 | 0.02993 | 44.25824 | FALSE | FALSE |
| LOC107065378 | uncharacterized protein LOC107065378 | 0.02936 | 44.26024 | FALSE | FALSE |
| LOC107065534 | carbohydrate sulfotransferase 4-like | 0.03034 | 44.2658 | FALSE | FALSE |
| LOC107073157 | 28S ribosomal protein S2, mitochondrial | 0.03221 | 44.28239 | FALSE | FALSE |
| LOC107071690 | uncharacterized protein LOC107071690 | 0.03033 | 44.28389 | FALSE | FALSE |
| LOC107073994 | alkaline ceramidase | 0.03121 | 44.30053 | FALSE | FALSE |
| LOC107068060 | TBC1 domain family member 31 | 0.03003 | 44.30536 | FALSE | FALSE |
| LOC107067033 | protein disconnected-like | 0.03022 | 44.31237 | FALSE | FALSE |
| LOC107065070 | cytochrome P450 4g15-like | 0.03058 | 44.31568 | FALSE | FALSE |
| LOC107071235 | uncharacterized protein LOC107071235 | 0.03195 | 44.32063 | FALSE | FALSE |
| LOC107070309 | vacuolar protein-sorting-associated protein | 0.0297 | 44.32548 | FALSE | FALSE |
| LOC107072189 | keratin, type I cytoskeletal 9-like | 0.03049 | 44.3336 | FALSE | FALSE |
| LOC107063735 | ferrochelatase, mitochondrial | 0.03123 | 44.35147 | FALSE | FALSE |
| LOC107072846 | sialin | 0.03028 | 44.36426 | FALSE | FALSE |
| LOC107072615 | protein croquemort-like | 0.0302 | 44.36553 | FALSE | FALSE |
| LOC107073279 | protein rhomboid | 0.03081 | 44.37658 | FALSE | FALSE |
| LOC107071713 | transcription factor Ken 2-like | 0.03113 | 44.38177 | FALSE | FALSE |
| LOC107066754 | tyrosine decarboxylase-like | 0.02976 | 44.38464 | FALSE | FALSE |
| LOC107074619 | paired box protein Pax-6-like | 0.03096 | 44.38921 | FALSE | FALSE |
| LOC107074408 | serine/threonine-protein kinase mTOR | 0.02997 | 44.38927 | FALSE | FALSE |
| LOC107065795 | uncharacterized protein LOC107065795 | 0.03036 | 44.39473 | FALSE | FALSE |
| LOC107067257 | membrane-bound transcription factor site-1 | 0.02993 | 44.41001 | FALSE | FALSE |
| LOC107069904 | phosphoenolpyruvate carboxykinase [GTP | 0.03126 | 44.41299 | FALSE | FALSE |
| LOC107072825 | RNA-binding protein 24-B-like | 0.03092 | 44.41837 | FALSE | FALSE |
| LOC107063750 | N-acetylgalactosaminyltransferase 6 | 0.03161 | 44.41932 | FALSE | FALSE |
| LOC107065277 | polypyrimidine tract-binding protein 1 | 0.02907 | 44.42339 | FALSE | FALSE |
| LOC107074041 | uncharacterized protein LOC107074041 | 0.03193 | 44.42624 | FALSE | FALSE |
| LOC107071178 | probable serine/threonine-protein kinase cl | 0.03066 | 44.42712 | FALSE | FALSE |
| LOC107072942 | LOW QUALITY PROTEIN: putative uncharact | 0.03051 | 44.43164 | FALSE | FALSE |

| | | | | | |
|---|---|---|---|---|---|
| LOC107067075 | pyruvate dehydrogenase [acetyl-transferrin | 0.03123 | 44.47391 | FALSE | FALSE |
| LOC107069781 | caspase-8 | 0.02961 | 44.47786 | FALSE | FALSE |
| LOC107065002 | tripartite motif-containing protein 45-like | 0.03079 | 44.48138 | FALSE | FALSE |
| LOC107071054 | elongation of very long chain fatty acids pro | 0.03086 | 44.48802 | FALSE | FALSE |
| LOC107074636 | polyadenylate-binding protein-interacting p | 0.03054 | 44.48952 | FALSE | FALSE |
| LOC107074284 | transcription factor SOX-14-like | 0.0299 | 44.49702 | FALSE | FALSE |
| LOC107071735 | serine/threonine-protein kinase grp | 0.0318 | 44.5143 | FALSE | FALSE |
| LOC107071158 | hybrid signal transduction histidine kinase E | 0.0304 | 44.51515 | FALSE | FALSE |
| LOC107070820 | general transcriptional corepressor trfA-like | 0.0303 | 44.51907 | FALSE | FALSE |
| LOC107070000 | uncharacterized protein LOC107070000 | 0.03011 | 44.5193 | FALSE | FALSE |
| LOC107069882 | serine/threonine-protein kinase ATR | 0.02957 | 44.52245 | FALSE | FALSE |
| LOC107073391 | uncharacterized protein LOC107073391 | 0.03152 | 44.52481 | FALSE | FALSE |
| LOC107069946 | alanine aminotransferase 1 | 0.03148 | 44.52643 | FALSE | FALSE |
| LOC107067596 | histidine triad nucleotide-binding protein 3- | 0.02934 | 44.53062 | FALSE | FALSE |
| LOC107071823 | neither inactivation nor afterpotential prote | 0.03049 | 44.53452 | FALSE | FALSE |
| LOC107069807 | uncharacterized protein LOC107069807 | 0.03051 | 44.55447 | FALSE | FALSE |
| LOC107064782 | fat-like cadherin-related tumor suppressor h | 0.0316 | 44.55904 | FALSE | FALSE |
| LOC107071775 | NEDD8-conjugating enzyme Ubc12 | 0.03111 | 44.56597 | FALSE | FALSE |
| LOC107074434 | NADH dehydrogenase [ubiquinone | 0.03057 | 44.5925 | FALSE | FALSE |
| LOC107069273 | sarcoplasmic calcium-binding protein 1 | 0.02996 | 44.59958 | FALSE | FALSE |
| LOC107072525 | uncharacterized protein LOC107072525 | 0.03065 | 44.61645 | FALSE | FALSE |
| LOC107067905 | uncharacterized protein LOC107067905 | 0.03041 | 44.62081 | FALSE | FALSE |
| LOC107065138 | DNA repair and recombination protein RAD! | 0.03036 | 44.62872 | FALSE | FALSE |
| LOC107064991 | probable phosphatase phospho2 | 0.03059 | 44.63151 | FALSE | FALSE |
| LOC107065662 | T-cell immunomodulatory protein | 0.03038 | 44.68551 | FALSE | FALSE |
| LOC107073055 | leucine-rich repeat-containing protein C10o | 0.03018 | 44.68764 | FALSE | FALSE |
| LOC107071004 | D(2) dopamine receptor-like | 0.02996 | 44.69599 | FALSE | FALSE |
| LOC107066833 | ATP-binding cassette sub-family F member 1 | 0.03157 | 44.70602 | FALSE | FALSE |
| LOC107074346 | ubiquitin conjugation factor E4 B | 0.0304 | 44.70772 | FALSE | FALSE |
| LOC107065912 | membrane-bound transcription factor site-2 | 0.03125 | 44.71417 | FALSE | FALSE |
| LOC107072972 | caveolin-3-like | 0.03061 | 44.71564 | FALSE | FALSE |
| LOC107073193 | dolichyl-diphosphooligosaccharide--protein | 0.03136 | 44.74916 | FALSE | FALSE |
| LOC107074201 | uncharacterized protein LOC107074201 | 0.03068 | 44.76194 | FALSE | FALSE |
| LOC107071242 | L-galactose dehydrogenase-like | 0.03078 | 44.78497 | FALSE | FALSE |
| LOC107064692 | vacuolar protein sorting-associated protein | 0.0295 | 44.78921 | FALSE | FALSE |
| LOC107064240 | uncharacterized protein LOC107064240 | 0.03061 | 44.79019 | FALSE | FALSE |
| LOC107070538 | uncharacterized protein LOC107070538 | 0.0298 | 44.80113 | FALSE | FALSE |
| LOC107072046 | protein yellow-like | 0.02999 | 44.80964 | FALSE | FALSE |
| LOC107064454 | uncharacterized protein LOC107064454 | 0.03 | 44.81248 | FALSE | FALSE |
| LOC107072059 | T-box transcription factor TBX1 | 0.02876 | 44.8169 | FALSE | FALSE |
| LOC107067272 | tubulointerstitial nephritis antigen-like | 0.03045 | 44.83845 | FALSE | FALSE |
| LOC107065100 | UPF0518 protein AAEL005291-like | 0.03092 | 44.84029 | FALSE | FALSE |
| LOC107074031 | caspase-like | 0.03043 | 44.84951 | FALSE | FALSE |
| LOC107066893 | probable sulfite oxidase, mitochondrial | 0.03017 | 44.86341 | FALSE | FALSE |
| LOC107071340 | AT-rich interactive domain-containing prote | 0.03044 | 44.8786 | FALSE | FALSE |
| LOC107067262 | transducin beta-like protein 2 | 0.03027 | 44.90933 | FALSE | FALSE |
| LOC107067152 | progestin and adipoQ receptor family meml | 0.03029 | 44.92402 | FALSE | FALSE |
| LOC107066039 | cytoplasmic aconitate hydratase-like | 0.02936 | 44.92749 | FALSE | FALSE |
| LOC107068849 | probable WRKY transcription factor protein | 0.03026 | 44.95966 | FALSE | FALSE |
| LOC107066526 | solute carrier family 25 member 46 | 0.03109 | 44.96918 | FALSE | FALSE |

| | | | | | |
|---|---|---|---|---|---|
| LOC107073052 | dnaJ homolog subfamily C member 8 | 0.03018 | 44.98309 | FALSE | FALSE |
| LOC107069342 | coiled-coil domain-containing protein 102A | 0.0289 | 44.99589 | FALSE | FALSE |
| LOC107070665 | uncharacterized protein LOC107070665 | 0.02967 | 45.00405 | FALSE | FALSE |
| LOC107074192 | DNA/RNA-binding protein KIN17 | 0.02976 | 45.0316 | FALSE | FALSE |
| LOC107065484 | 7SK snRNA methylphosphate capping enzym | 0.03037 | 45.03272 | FALSE | FALSE |
| LOC107063652 | uncharacterized protein LOC107063652 | 0.03027 | 45.0431 | FALSE | FALSE |
| LOC107066395 | uncharacterized protein LOC107066395 | 0.0309 | 45.05849 | FALSE | FALSE |
| LOC107065487 | uncharacterized protein LOC107065487 | 0.03057 | 45.06338 | FALSE | FALSE |
| LOC107065404 | serine-rich adhesin for platelets-like | 0.02977 | 45.06455 | FALSE | FALSE |
| LOC107071467 | RNA-binding protein 28-like | 0.03022 | 45.07956 | FALSE | FALSE |
| LOC107073232 | ubiquitin-like protein 4A | 0.0312 | 45.08857 | FALSE | FALSE |
| LOC107065614 | triosephosphate isomerase | 0.03186 | 45.08983 | FALSE | FALSE |
| LOC107065683 | protein FAM45A-like | 0.02988 | 45.09267 | FALSE | FALSE |
| LOC107069055 | uncharacterized protein LOC107069055 | 0.03023 | 45.0953 | FALSE | FALSE |
| LOC107066128 | RING finger protein 37 | 0.02968 | 45.10012 | FALSE | FALSE |
| LOC107069733 | activating transcription factor of chaperone | 0.03006 | 45.10338 | FALSE | FALSE |
| LOC107070500 | nose resistant to fluoxetine protein 6-like | 0.02873 | 45.11499 | FALSE | FALSE |
| LOC107066105 | protein NDUFAF4 homolog | 0.0308 | 45.12841 | FALSE | FALSE |
| LOC107068173 | coiled-coil domain-containing protein 85C | 0.03004 | 45.13451 | FALSE | FALSE |
| LOC107065841 | zinc finger MYND domain-containing protei | 0.03094 | 45.15073 | FALSE | FALSE |
| LOC107074246 | protein asteroid | 0.0304 | 45.15123 | FALSE | FALSE |
| LOC107071624 | general vesicular transport factor p115 | 0.03055 | 45.16447 | FALSE | FALSE |
| LOC107066463 | putative uncharacterized protein DDB_G02 | 0.02899 | 45.16677 | FALSE | FALSE |
| LOC107073019 | uncharacterized protein LOC107073019 | 0.03001 | 45.16993 | FALSE | FALSE |
| LOC107068242 | uncharacterized protein LOC107068242 | 0.02978 | 45.1727 | FALSE | FALSE |
| LOC107068139 | chromosome-associated kinesin KIF4 | 0.03021 | 45.18254 | FALSE | FALSE |
| LOC107066703 | uncharacterized protein LOC107066703 | 0.0304 | 45.18588 | FALSE | FALSE |
| LOC107072651 | uncharacterized protein LOC107072651 | 0.03118 | 45.19821 | FALSE | FALSE |
| LOC107070006 | uncharacterized protein LOC107070006 | 0.03154 | 45.20206 | FALSE | FALSE |
| LOC107073254 | uncharacterized protein LOC107073254 | 0.02997 | 45.20394 | FALSE | FALSE |
| LOC107067325 | ras-related GTP-binding protein C | 0.0304 | 45.20718 | FALSE | FALSE |
| LOC107073138 | E3 ubiquitin-protein ligase MARCH5-like | 0.02961 | 45.22034 | FALSE | FALSE |
| LOC107064183 | eukaryotic translation initiation factor 3 sub | 0.02987 | 45.22373 | FALSE | FALSE |
| LOC107071221 | uncharacterized protein LOC107071221 | 0.02935 | 45.24019 | FALSE | FALSE |
| LOC107066308 | uncharacterized protein LOC107066308 | 0.03175 | 45.2517 | FALSE | FALSE |
| LOC107065582 | ufm1-specific protease 2 | 0.03083 | 45.26493 | FALSE | FALSE |
| LOC107066619 | esterase E4-like | 0.03178 | 45.26558 | FALSE | FALSE |
| LOC107065153 | kanadaptin | 0.02964 | 45.28145 | FALSE | FALSE |
| LOC107068758 | uncharacterized protein LOC107068758 | 0.03011 | 45.30204 | FALSE | FALSE |
| LOC107069905 | AP-3 complex subunit delta-1 | 0.03099 | 45.30604 | FALSE | FALSE |
| LOC107063846 | IQ and ubiquitin-like domain-containing pro | 0.03087 | 45.38883 | FALSE | FALSE |
| LOC107070236 | ras-related protein Rab-32-like | 0.03002 | 45.40351 | FALSE | FALSE |
| LOC107072637 | cytochrome P450 4C1-like | 0.03067 | 45.40732 | FALSE | FALSE |
| LOC107065186 | tyramine receptor 1-like | 0.02978 | 45.41482 | FALSE | FALSE |
| LOC107066206 | transmembrane protein 42-like | 0.02989 | 45.42041 | FALSE | FALSE |
| LOC107067243 | multiple RNA-binding domain-containing pr | 0.03014 | 45.44683 | FALSE | FALSE |
| LOC107066303 | uncharacterized protein LOC107066303 | 0.0296 | 45.47109 | FALSE | FALSE |
| LOC107067944 | protein gar2-like | 0.02894 | 45.47885 | FALSE | FALSE |
| LOC107074389 | oxysterol-binding protein-related protein 8 | 0.03043 | 45.48241 | FALSE | FALSE |
| LOC107068955 | uncharacterized protein LOC107068955 | 0.03237 | 45.49029 | FALSE | FALSE |

| | | | | | |
|---|---|---|---|---|---|
| LOC107068750 | uncharacterized protein LOC107068750 | 0.02897 | 45.50465 | FALSE | FALSE |
| LOC107066643 | RUN domain-containing protein 1 | 0.03068 | 45.54765 | FALSE | FALSE |
| LOC107064280 | uncharacterized protein LOC107064280 | 0.02998 | 45.55486 | FALSE | FALSE |
| LOC107068858 | uncharacterized threonine-rich GPI-anchore | 0.03187 | 45.57765 | FALSE | FALSE |
| LOC107068745 | acid sphingomyelinase-like phosphodiestera | 0.02929 | 45.58163 | FALSE | FALSE |
| LOC107066701 | uncharacterized protein LOC107066701 | 0.02959 | 45.60002 | FALSE | FALSE |
| LOC107069144 | 60S ribosomal protein L21 | 0.02997 | 45.6021 | FALSE | FALSE |
| LOC107068784 | uncharacterized protein LOC107068784 | 0.02856 | 45.60231 | FALSE | FALSE |
| LOC107070003 | uncharacterized protein LOC107070003 | 0.03171 | 45.60558 | FALSE | FALSE |
| LOC107064817 | uncharacterized protein LOC107064817 | 0.03002 | 45.60756 | FALSE | FALSE |
| LOC107069800 | cytochrome P450 4C1-like | 0.03005 | 45.61321 | FALSE | FALSE |
| LOC107069788 | vesicular glutamate transporter 1 | 0.02936 | 45.61786 | FALSE | FALSE |
| LOC107070610 | RING finger and transmembrane domain-co | 0.03138 | 45.62154 | FALSE | FALSE |
| LOC107069416 | homeobox protein 2-like | 0.0303 | 45.63434 | FALSE | FALSE |
| LOC107072839 | farnesol dehydrogenase-like | 0.0314 | 45.64994 | FALSE | FALSE |
| LOC107068565 | uncharacterized protein LOC107068565 | 0.03072 | 45.65687 | FALSE | FALSE |
| LOC107065124 | ubiquitin carboxyl-terminal hydrolase 31-lik | 0.02948 | 45.66243 | FALSE | FALSE |
| LOC107074671 | protein groucho-1-like | 0.03042 | 45.6628 | FALSE | FALSE |
| LOC107073325 | uncharacterized protein LOC107073325 | 0.02951 | 45.67165 | FALSE | FALSE |
| LOC107069723 | probable tRNA pseudouridine synthase 2 | 0.03041 | 45.67614 | FALSE | FALSE |
| LOC107063621 | uncharacterized protein LOC107063621 | 0.02992 | 45.68173 | FALSE | FALSE |
| LOC107073345 | uncharacterized protein LOC107073345 | 0.03032 | 45.68582 | FALSE | FALSE |
| LOC107065185 | oxysterol-binding protein-related protein 2 | 0.02861 | 45.68806 | FALSE | FALSE |
| LOC107071135 | cilia- and flagella-associated protein 44 | 0.03147 | 45.68997 | FALSE | FALSE |
| LOC107063710 | pterin-4-alpha-carbinolamine dehydratase | 0.0301 | 45.69628 | FALSE | FALSE |
| LOC107067815 | actin, clone 205-like | 0.02894 | 45.69816 | FALSE | FALSE |
| LOC107066817 | uncharacterized protein LOC107066817 | 0.03075 | 45.70696 | FALSE | FALSE |
| LOC107069085 | kielin/chordin-like protein | 0.03108 | 45.71899 | FALSE | FALSE |
| LOC107064910 | uncharacterized protein LOC107064910 | 0.03011 | 45.72364 | FALSE | FALSE |
| LOC107064022 | sodium channel protein Nach-like | 0.02946 | 45.75118 | FALSE | FALSE |
| LOC107070224 | sphingosine-1-phosphate phosphatase 2-lik | 0.02941 | 45.75354 | FALSE | FALSE |
| LOC107064569 | uncharacterized protein C11orf70 homolog | 0.02988 | 45.77122 | FALSE | FALSE |
| LOC107067331 | ubiquitin-protein ligase E3A | 0.0297 | 45.77449 | FALSE | FALSE |
| LOC107067366 | muscle-specific protein 20 | 0.02863 | 45.77952 | FALSE | FALSE |
| LOC107073310 | prolactin-releasing peptide receptor-like | 0.03071 | 45.78242 | FALSE | FALSE |
| LOC107069778 | digestive cysteine proteinase 1 | 0.02989 | 45.78309 | FALSE | FALSE |
| LOC107068545 | methyltransferase-like protein 17, mitochor | 0.03007 | 45.7849 | FALSE | FALSE |
| LOC107070634 | dynein light chain 4, axonemal-like | 0.03093 | 45.8158 | FALSE | FALSE |
| LOC107066926 | arginine--tRNA ligase, cytoplasmic | 0.03016 | 45.81851 | FALSE | FALSE |
| LOC107068468 | protein PFC0760c-like | 0.02954 | 45.84586 | FALSE | FALSE |
| LOC107070794 | DNA repair protein complementing XP-G ce | 0.02962 | 45.86873 | FALSE | FALSE |
| LOC107068993 | SWI/SNF-related matrix-associated actin-de | 0.0296 | 45.87179 | FALSE | FALSE |
| LOC107069447 | UPF0193 protein EVG1 homolog | 0.02996 | 45.88633 | FALSE | FALSE |
| LOC107069698 | tRNA-splicing ligase RtcB homolog | 0.02994 | 45.89196 | FALSE | FALSE |
| LOC107064585 | uncharacterized protein LOC107064585 | 0.02975 | 45.89686 | FALSE | FALSE |
| LOC107073295 | ERI1 exoribonuclease 3 | 0.02997 | 45.91026 | FALSE | FALSE |
| LOC107066796 | GPI transamidase component PIG-S | 0.02977 | 45.92935 | FALSE | FALSE |
| LOC107072426 | cytochrome c oxidase assembly factor 4 hor | 0.03077 | 45.93366 | FALSE | FALSE |
| LOC107071526 | GRIP and coiled-coil domain-containing prot | 0.02956 | 45.93723 | FALSE | FALSE |
| LOC107068028 | RING-box protein 1A | 0.02976 | 45.94782 | FALSE | FALSE |

| | | | | | |
|---|---|---|---|---|---|
| LOC107067294 | probable G-protein coupled receptor Mth-li | 0.03085 | 45.95089 | FALSE | FALSE |
| LOC107066943 | leucine-rich PPR motif-containing protein, m | 0.02967 | 45.95737 | FALSE | FALSE |
| LOC107069048 | alpha-taxilin | 0.03065 | 45.95808 | FALSE | FALSE |
| LOC107066273 | zinc finger protein 205-like | 0.02934 | 45.95889 | FALSE | FALSE |
| LOC107065626 | uncharacterized protein LOC107065626 | 0.02965 | 45.97544 | FALSE | FALSE |
| LOC107068913 | uncharacterized protein LOC107068913 | 0.02984 | 45.98058 | FALSE | FALSE |
| LOC107065615 | prostaglandin E synthase 3 | 0.02924 | 45.98185 | FALSE | FALSE |
| LOC107070299 | receptor-type guanylate cyclase daf-11 | 0.03026 | 45.99857 | FALSE | FALSE |
| LOC107074127 | uncharacterized protein LOC107074127 | 0.02912 | 46.00573 | FALSE | FALSE |
| LOC107070153 | cytoplasmic dynein 1 intermediate chain-lik | 0.03229 | 46.01464 | FALSE | FALSE |
| LOC107063883 | uncharacterized protein LOC107063883 | 0.02891 | 46.02317 | FALSE | FALSE |
| LOC107066160 | kinesin-like protein KIF19 | 0.0298 | 46.02506 | FALSE | FALSE |
| LOC107068665 | alpha-ketoglutarate-dependent dioxygenase | 0.02961 | 46.04389 | FALSE | FALSE |
| LOC107072708 | leucine-rich repeat-containing protein 47-lik | 0.03008 | 46.04404 | FALSE | FALSE |
| LOC107064066 | solute carrier family 26 member 10-like | 0.03159 | 46.05609 | FALSE | FALSE |
| LOC107073749 | cytochrome P450 4C1-like | 0.03067 | 46.07236 | FALSE | FALSE |
| LOC107070323 | hypoxia-inducible factor 1-alpha | 0.03013 | 46.07926 | FALSE | FALSE |
| LOC107068648 | facilitated trehalose transporter Tret1-like | 0.03025 | 46.0813 | FALSE | FALSE |
| LOC107068149 | protein unc-119 homolog B | 0.02976 | 46.08431 | FALSE | FALSE |
| LOC107068914 | V-type proton ATPase subunit e 2-like | 0.02977 | 46.10092 | FALSE | FALSE |
| LOC107067668 | HBS1-like protein | 0.03026 | 46.1383 | FALSE | FALSE |
| LOC107074473 | uncharacterized protein KIAA1841 | 0.03002 | 46.15236 | FALSE | FALSE |
| LOC107063770 | death-associated protein kinase 1-like | 0.03019 | 46.15704 | FALSE | FALSE |
| LOC107067251 | uncharacterized protein LOC107067251 | 0.03013 | 46.16072 | FALSE | FALSE |
| LOC107069253 | calcium and integrin-binding protein 1-like | 0.03052 | 46.18668 | FALSE | FALSE |
| LOC107068662 | chymotrypsin-2-like | 0.02892 | 46.19167 | FALSE | FALSE |
| LOC107071361 | methylglutaconyl-CoA hydratase, mitochon | 0.02883 | 46.2015 | FALSE | FALSE |
| LOC107069324 | uncharacterized protein LOC107069324 | 0.02973 | 46.21053 | FALSE | FALSE |
| LOC107067052 | uncharacterized protein LOC107067052 | 0.02974 | 46.22537 | FALSE | FALSE |
| LOC107064696 | protein shuttle craft | 0.02943 | 46.22747 | FALSE | FALSE |
| LOC107067807 | epsilon-sarcoglycan | 0.02889 | 46.23517 | FALSE | FALSE |
| LOC107072184 | mitochondrial import receptor subunit TOM | 0.02978 | 46.23947 | FALSE | FALSE |
| LOC107065957 | 28S ribosomal protein S28, mitochondrial | 0.03094 | 46.25142 | FALSE | FALSE |
| LOC107066097 | cytochrome c oxidase assembly factor 3, mi | 0.03018 | 46.26475 | FALSE | FALSE |
| LOC107071495 | zinc finger RNA-binding protein | 0.03077 | 46.30785 | FALSE | FALSE |
| LOC107071033 | uncharacterized protein LOC107071033 | 0.02826 | 46.32478 | FALSE | FALSE |
| LOC107068746 | glutathione S-transferase D5-like | 0.03089 | 46.33364 | FALSE | FALSE |
| LOC107065888 | uncharacterized protein LOC107065888 | 0.0306 | 46.341 | FALSE | FALSE |
| LOC107064953 | KN motif and ankyrin repeat domain-contai | 0.0296 | 46.3454 | FALSE | FALSE |
| LOC107068835 | uncharacterized protein LOC107068835 | 0.03118 | 46.34989 | FALSE | FALSE |
| LOC107069590 | sodium-dependent phosphate transport pro | 0.02941 | 46.35409 | FALSE | FALSE |
| LOC107072887 | zinc finger protein on ecdysone puffs | 0.02946 | 46.36691 | FALSE | FALSE |
| LOC107068268 | prostaglandin reductase 1-like | 0.02963 | 46.38533 | FALSE | FALSE |
| LOC107064550 | protein hook homolog | 0.03005 | 46.38557 | FALSE | FALSE |
| LOC107065547 | angiotensin-converting enzyme | 0.02912 | 46.38995 | FALSE | FALSE |
| LOC107069125 | RNA-binding protein squid | 0.03013 | 46.39431 | FALSE | FALSE |
| LOC107071671 | uncharacterized protein LOC107071671 | 0.02943 | 46.41164 | FALSE | FALSE |
| LOC107071037 | activin receptor type-1-like | 0.02973 | 46.41193 | FALSE | FALSE |
| LOC107066826 | vesicle-trafficking protein SEC22b | 0.02917 | 46.4604 | FALSE | FALSE |
| LOC107065741 | SOSS complex subunit B homolog | 0.02965 | 46.46184 | FALSE | FALSE |

| | | | | | |
|---|---|---|---|---|---|
| LOC107073633 | uncharacterized protein LOC107073633 | 0.03084 | 46.46859 | FALSE | FALSE |
| LOC107068220 | uncharacterized protein LOC107068220 | 0.02999 | 46.49093 | FALSE | FALSE |
| LOC107064888 | protein turtle | 0.0288 | 46.4981 | FALSE | FALSE |
| LOC107068560 | uncharacterized protein LOC107068560 | 0.03056 | 46.49948 | FALSE | FALSE |
| LOC107069154 | inositol polyphosphate 1-phosphatase | 0.0283 | 46.50954 | FALSE | FALSE |
| LOC107071492 | facilitated trehalose transporter Tret1-like | 0.0294 | 46.51715 | FALSE | FALSE |
| LOC107071887 | uncharacterized protein LOC107071887 | 0.03008 | 46.5324 | FALSE | FALSE |
| LOC107070146 | uncharacterized protein LOC107070146 | 0.03039 | 46.54648 | FALSE | FALSE |
| LOC107066384 | uncharacterized protein LOC107066384 | 0.02967 | 46.54833 | FALSE | FALSE |
| LOC107064016 | 60S ribosomal protein L35 | 0.02904 | 46.55405 | FALSE | FALSE |
| LOC107068120 | nurim homolog | 0.02924 | 46.57206 | FALSE | FALSE |
| LOC107071400 | bestrophin-4-like | 0.02911 | 46.58594 | FALSE | FALSE |
| LOC107070148 | tyrosine-protein phosphatase 10D | 0.03029 | 46.59891 | FALSE | FALSE |
| LOC107071042 | probable inactive tRNA-specific adenosine d | 0.03152 | 46.60011 | FALSE | FALSE |
| LOC107071430 | uncharacterized protein LOC107071430 | 0.02962 | 46.61494 | FALSE | FALSE |
| LOC107069741 | tyrosine-protein kinase Fps85D | 0.02949 | 46.61948 | FALSE | FALSE |
| LOC107074432 | inositol monophosphatase 2-like | 0.03177 | 46.62364 | FALSE | FALSE |
| Trnas-aga | Trnas-aga | 0.03032 | 46.62718 | FALSE | FALSE |
| LOC107072472 | metallo-beta-lactamase domain-containing | 0.02845 | 46.65308 | FALSE | FALSE |
| LOC107071773 | uncharacterized protein LOC107071773 | 0.03017 | 46.66569 | FALSE | FALSE |
| LOC107067408 | bone morphogenetic protein receptor type- | 0.03006 | 46.66891 | FALSE | FALSE |
| LOC107069099 | zinc finger protein 543-like | 0.02923 | 46.66995 | FALSE | FALSE |
| LOC107071610 | nascent polypeptide-associated complex su | 0.0288 | 46.67057 | FALSE | FALSE |
| LOC107073077 | E3 ubiquitin-protein ligase UBR2 | 0.02947 | 46.67825 | FALSE | FALSE |
| LOC107067591 | conserved oligomeric Golgi complex subuni | 0.02974 | 46.6876 | FALSE | FALSE |
| LOC107067872 | uncharacterized protein LOC107067872 | 0.03026 | 46.69545 | FALSE | FALSE |
| LOC107074322 | integrin beta-PS | 0.02864 | 46.69799 | FALSE | FALSE |
| LOC107069105 | contactin | 0.02997 | 46.70954 | FALSE | FALSE |
| LOC107070673 | uronyl 2-sulfotransferase-like | 0.03017 | 46.71606 | FALSE | FALSE |
| LOC107066920 | X-box-binding protein 1-like | 0.03037 | 46.74068 | FALSE | FALSE |
| LOC107064351 | uncharacterized protein LOC107064351 | 0.02879 | 46.74543 | FALSE | FALSE |
| LOC107074475 | ribosomal protein S6 kinase beta-1-like | 0.03147 | 46.80204 | FALSE | FALSE |
| LOC107068538 | protein lin-54 homolog | 0.03128 | 46.8026 | FALSE | FALSE |
| LOC107067106 | organic cation transporter protein | 0.02946 | 46.81512 | FALSE | FALSE |
| LOC107065238 | DNA-directed RNA polymerase II subunit RP | 0.02821 | 46.8382 | FALSE | FALSE |
| LOC107065194 | inner centromere protein A-like | 0.02985 | 46.84569 | FALSE | FALSE |
| LOC107073143 | protein transport protein Sec61 subunit bet | 0.02984 | 46.84822 | FALSE | FALSE |
| LOC107064719 | tetratricopeptide repeat protein 39B-like | 0.02948 | 46.84973 | FALSE | FALSE |
| LOC107063820 | CTL-like protein 1 | 0.02914 | 46.8682 | FALSE | FALSE |
| LOC107063650 | transmembrane protein 70 homolog, mitoc | 0.02999 | 46.87623 | FALSE | FALSE |
| LOC107066258 | low-density lipoprotein receptor-related pro | 0.03012 | 46.88053 | FALSE | FALSE |
| LOC107064754 | leishmanolysin-like peptidase | 0.03081 | 46.91018 | FALSE | FALSE |
| LOC107072499 | U3 small nucleolar ribonucleoprotein protei | 0.02847 | 46.91699 | FALSE | FALSE |
| LOC107067921 | Fanconi anemia group I protein homolog | 0.02954 | 46.91815 | FALSE | FALSE |
| LOC107067623 | DNA ligase 1-like | 0.02941 | 46.94188 | FALSE | FALSE |
| LOC107065439 | uncharacterized protein LOC107065439 | 0.03059 | 46.94375 | FALSE | FALSE |
| LOC107069291 | sorbitol dehydrogenase-like | 0.02949 | 46.94598 | FALSE | FALSE |
| LOC107073992 | alanine--glyoxylate aminotransferase 2-like | 0.02943 | 46.95173 | FALSE | FALSE |
| LOC107063976 | leucine-rich repeat-containing protein 59 | 0.0303 | 46.95263 | FALSE | FALSE |
| LOC107067479 | uncharacterized protein LOC107067479 | 0.0289 | 46.96429 | FALSE | FALSE |

| | | | | | |
|---|---|---|---|---|---|
| LOC107066519 | sonic hedgehog protein A | 0.03153 | 46.96488 | FALSE | FALSE |
| LOC107074375 | small conductance calcium-activated potass | 0.02957 | 46.96715 | FALSE | FALSE |
| LOC107063739 | broad-complex core protein isoforms 1/2/3, | 0.02954 | 47.00623 | FALSE | FALSE |
| LOC107072812 | gem-associated protein 7-like | 0.02937 | 47.00675 | FALSE | FALSE |
| LOC107069078 | mortality factor 4-like protein 1 | 0.03008 | 47.0325 | FALSE | FALSE |
| LOC107069097 | single-stranded DNA-binding protein, mitoc | 0.02968 | 47.04845 | FALSE | FALSE |
| LOC107073022 | general transcription factor 3C polypeptide | 0.02943 | 47.0561 | FALSE | FALSE |
| LOC107065177 | uncharacterized protein LOC107065177 | 0.03046 | 47.05725 | FALSE | FALSE |
| LOC107067146 | ribosomal protein S6 kinase beta-1-like | 0.02963 | 47.06696 | FALSE | FALSE |
| LOC107070656 | deoxynucleoside kinase | 0.02915 | 47.09334 | FALSE | FALSE |
| LOC107073918 | chromodomain-helicase-DNA-binding prote | 0.02989 | 47.095 | FALSE | FALSE |
| LOC107073302 | uncharacterized protein LOC107073302 | 0.03015 | 47.10426 | FALSE | FALSE |
| LOC107068074 | origin recognition complex subunit 4 | 0.02944 | 47.11643 | FALSE | FALSE |
| LOC107064000 | probable ATP-dependent RNA helicase spin | 0.02873 | 47.11742 | FALSE | FALSE |
| LOC107071659 | DNA replication factor Cdt1 | 0.02824 | 47.12249 | FALSE | FALSE |
| LOC107066862 | succinate dehydrogenase cytochrome b560 | 0.02935 | 47.14173 | FALSE | FALSE |
| LOC107065501 | limulus clotting factor C-like | 0.02897 | 47.16237 | FALSE | FALSE |
| LOC107070140 | uncharacterized protein LOC107070140 | 0.02986 | 47.17996 | FALSE | FALSE |
| LOC107070803 | thioredoxin domain-containing protein 15 | 0.02834 | 47.18255 | FALSE | FALSE |
| LOC107072509 | uncharacterized protein LOC107072509 | 0.02956 | 47.18346 | FALSE | FALSE |
| LOC107068876 | uncharacterized protein LOC107068876 | 0.02963 | 47.19598 | FALSE | FALSE |
| LOC107066437 | PITH domain-containing protein GA19395 | 0.02872 | 47.19637 | FALSE | FALSE |
| LOC107066023 | phosphoglycerate mutase 2 | 0.03021 | 47.20731 | FALSE | FALSE |
| Trnad-guc | Trnad-guc | 0.02923 | 47.21898 | FALSE | FALSE |
| LOC107071897 | serine-protein kinase ATM | 0.02902 | 47.24419 | FALSE | FALSE |
| LOC107064661 | neuroblastoma-amplified sequence-like | 0.02773 | 47.24478 | FALSE | FALSE |
| LOC107071975 | actin-like protein 6B | 0.03064 | 47.25383 | FALSE | FALSE |
| LOC107064583 | uncharacterized protein LOC107064583 | 0.02805 | 47.25889 | FALSE | FALSE |
| LOC107070354 | furin-like protease 1, isoform 1-CRR | 0.02995 | 47.27708 | FALSE | FALSE |
| LOC107069547 | uncharacterized protein LOC107069547 | 0.02973 | 47.27937 | FALSE | FALSE |
| LOC107069187 | protoporphyrinogen oxidase | 0.0302 | 47.28326 | FALSE | FALSE |
| LOC107074304 | serine/arginine repetitive matrix protein 1 | 0.02843 | 47.33401 | FALSE | FALSE |
| LOC107072609 | uncharacterized protein LOC107072609 | 0.02908 | 47.33835 | FALSE | FALSE |
| LOC107071496 | uncharacterized protein LOC107071496 | 0.02906 | 47.35225 | FALSE | FALSE |
| LOC107069203 | probable cytochrome P450 304a1 | 0.02922 | 47.35639 | FALSE | FALSE |
| LOC107064039 | condensin complex subunit 1 | 0.02935 | 47.3578 | FALSE | FALSE |
| LOC107067690 | vesicle-associated membrane protein-assoc | 0.03034 | 47.36362 | FALSE | FALSE |
| LOC107073275 | putative RNA polymerase II subunit B1 CTD | 0.02969 | 47.36685 | FALSE | FALSE |
| LOC107068204 | succinate dehydrogenase [ubiquinone | 0.02939 | 47.36945 | FALSE | FALSE |
| LOC107067871 | transcription factor kayak | 0.02882 | 47.37673 | FALSE | FALSE |
| LOC107065946 | charged multivesicular body protein 6-A | 0.03079 | 47.37722 | FALSE | FALSE |
| LOC107074493 | polypeptide N-acetylgalactosaminyltransfer | 0.02995 | 47.39583 | FALSE | FALSE |
| LOC107065502 | tRNA pseudouridine synthase A, mitochond | 0.03066 | 47.40144 | FALSE | FALSE |
| LOC107065456 | neuropeptide FF receptor 2-like | 0.0306 | 47.41607 | FALSE | FALSE |
| LOC107070962 | structural maintenance of chromosomes pr | 0.02837 | 47.42043 | FALSE | FALSE |
| LOC107072921 | geminin-like | 0.02921 | 47.42816 | FALSE | FALSE |
| LOC107073205 | period circadian protein | 0.02922 | 47.44994 | FALSE | FALSE |
| LOC107072934 | syntenin-1-like | 0.02949 | 47.4596 | FALSE | FALSE |
| LOC107066662 | uncharacterized protein LOC107066662 | 0.02916 | 47.46164 | FALSE | FALSE |
| LOC107067293 | intron-binding protein aquarius | 0.0291 | 47.49318 | FALSE | FALSE |

| | | | | | |
|---|---|---|---|---|---|
| LOC107064042 | tubulin beta chain-like | 0.02977 | 47.50669 | FALSE | FALSE |
| LOC107071983 | FYVE, RhoGEF and PH domain-containing pr | 0.02927 | 47.50856 | FALSE | FALSE |
| LOC107074417 | sialin | 0.02926 | 47.52272 | FALSE | FALSE |
| LOC107066306 | tensin-3 | 0.02921 | 47.54499 | FALSE | FALSE |
| LOC107065414 | translation initiation factor eIF-2B subunit e | 0.02907 | 47.54657 | FALSE | FALSE |
| LOC107068594 | uncharacterized protein LOC107068594 | 0.03123 | 47.5482 | FALSE | FALSE |
| LOC107073106 | uncharacterized protein LOC107073106 | 0.0302 | 47.5642 | FALSE | FALSE |
| LOC107073598 | uncharacterized protein LOC107073598 | 0.02865 | 47.5992 | FALSE | FALSE |
| LOC107067975 | septin-2-like | 0.02986 | 47.61095 | FALSE | FALSE |
| LOC107064308 | zinc finger CCHC domain-containing protein | 0.02849 | 47.6237 | FALSE | FALSE |
| LOC107065315 | cuticle protein 18.7-like | 0.02952 | 47.62797 | FALSE | FALSE |
| LOC107065352 | uncharacterized protein LOC107065352 | 0.02948 | 47.63872 | FALSE | FALSE |
| LOC107068162 | pyruvate dehydrogenase (acetyl-transferrin | 0.0295 | 47.64161 | FALSE | FALSE |
| LOC107067125 | tRNA-dihydrouridine(16/17) synthase [NAD | 0.02987 | 47.64174 | FALSE | FALSE |
| LOC107063957 | zinc finger MYM-type protein 3 | 0.02728 | 47.65299 | FALSE | FALSE |
| LOC107071847 | motile sperm domain-containing protein 2-l | 0.02866 | 47.66198 | FALSE | FALSE |
| LOC107071180 | 1-acyl-sn-glycerol-3-phosphate acyltransfera | 0.0282 | 47.67406 | FALSE | FALSE |
| LOC107066997 | upstream stimulatory factor-like | 0.0302 | 47.67728 | FALSE | FALSE |
| LOC107063977 | circadian clock-controlled protein-like | 0.03042 | 47.68757 | FALSE | FALSE |
| LOC107066472 | protein crooked neck | 0.02903 | 47.69322 | FALSE | FALSE |
| LOC107074026 | uncharacterized protein LOC107074026 | 0.02931 | 47.70886 | FALSE | FALSE |
| LOC107067196 | nuclear cap-binding protein subunit 1 | 0.03072 | 47.71157 | FALSE | FALSE |
| LOC107069971 | venom acid phosphatase Acph-1-like | 0.02845 | 47.71726 | FALSE | FALSE |
| LOC107066498 | uncharacterized protein LOC107066498 | 0.02945 | 47.72438 | FALSE | FALSE |
| LOC107064521 | EF-hand calcium-binding domain-containing | 0.03107 | 47.76556 | FALSE | FALSE |
| LOC107067814 | actin, clone 205-like | 0.02854 | 47.76672 | FALSE | FALSE |
| LOC107073396 | serine/threonine-protein kinase PINK1, mit | 0.02917 | 47.78155 | FALSE | FALSE |
| LOC107064053 | uncharacterized protein LOC107064053 | 0.02924 | 47.79212 | FALSE | FALSE |
| LOC107072210 | general transcription factor IIF subunit 1 | 0.02935 | 47.80029 | FALSE | FALSE |
| LOC107073803 | high affinity copper uptake protein 1-like | 0.02966 | 47.81574 | FALSE | FALSE |
| LOC107067920 | DNA polymerase zeta catalytic subunit | 0.02867 | 47.82527 | FALSE | FALSE |
| LOC107072001 | splicing factor, suppressor of white-apricot | 0.02832 | 47.84775 | FALSE | FALSE |
| LOC107073489 | heart- and neural crest derivatives-expresse | 0.02925 | 47.8529 | FALSE | FALSE |
| LOC107072270 | uncharacterized protein LOC107072270 | 0.02961 | 47.86663 | FALSE | FALSE |
| LOC107074139 | casein kinase I isoform gamma-3 | 0.02925 | 47.88583 | FALSE | FALSE |
| LOC107067571 | uncharacterized protein LOC107067571 | 0.02914 | 47.9055 | FALSE | FALSE |
| LOC107071833 | uncharacterized protein C4orf29 | 0.02858 | 47.9086 | FALSE | FALSE |
| LOC107066696 | NF-kappa-B-repressing factor | 0.02925 | 47.92077 | FALSE | FALSE |
| LOC107069114 | dnaJ homolog subfamily C member 5-like | 0.02962 | 47.92388 | FALSE | FALSE |
| LOC107068193 | monocarboxylate transporter 12-B-like | 0.02874 | 47.92559 | FALSE | FALSE |
| LOC107070444 | protein unc-13 homolog D-like | 0.02902 | 47.9263 | FALSE | FALSE |
| LOC107065868 | 60S ribosomal protein L5-like | 0.02894 | 47.93063 | FALSE | FALSE |
| LOC107066684 | uncharacterized protein LOC107066684 | 0.03091 | 47.94582 | FALSE | FALSE |
| LOC107070514 | uncharacterized protein LOC107070514 | 0.02829 | 47.96843 | FALSE | FALSE |
| LOC107065003 | androgen-induced gene 1 protein-like | 0.02917 | 47.97906 | FALSE | FALSE |
| LOC107068944 | uncharacterized protein LOC107068944 | 0.02851 | 47.98392 | FALSE | FALSE |
| LOC107067802 | neprilysin-11 | 0.02976 | 48.01998 | FALSE | FALSE |
| LOC107067382 | uncharacterized protein LOC107067382 | 0.02929 | 48.04032 | FALSE | FALSE |
| LOC107069062 | fatty-acid amide hydrolase 2-like | 0.03016 | 48.05286 | FALSE | FALSE |
| LOC107071219 | uncharacterized protein LOC107071219 | 0.02975 | 48.05567 | FALSE | FALSE |

| | | | | | |
|---|---|---|---|---|---|
| LOC107066962 | facilitated trehalose transporter Tret1-like | 0.02919 | 48.05796 | FALSE | FALSE |
| LOC107065494 | uncharacterized protein LOC107065494 | 0.02971 | 48.06093 | FALSE | FALSE |
| LOC107064237 | uncharacterized protein LOC107064237 | 0.02845 | 48.07124 | FALSE | FALSE |
| LOC107068936 | putative nuclease HARBI1 | 0.0294 | 48.07983 | FALSE | FALSE |
| LOC107063881 | proteasomal ubiquitin receptor ADRM1 | 0.02883 | 48.08379 | FALSE | FALSE |
| LOC107063805 | probable rRNA-processing protein EBP2 hor | 0.02923 | 48.08605 | FALSE | FALSE |
| LOC107069340 | DNA repair protein RAD50 | 0.02817 | 48.0906 | FALSE | FALSE |
| LOC107070408 | 40S ribosomal protein S3a | 0.02891 | 48.09811 | FALSE | FALSE |
| LOC107072738 | zinc finger protein 267 | 0.02929 | 48.10459 | FALSE | FALSE |
| LOC107071505 | geranylgeranyl transferase type-1 subunit b | 0.02905 | 48.11655 | FALSE | FALSE |
| LOC107072511 | saccharopine dehydrogenase-like oxidoredu | 0.02915 | 48.1367 | FALSE | FALSE |
| LOC107065535 | cyclin-G1 | 0.02907 | 48.13793 | FALSE | FALSE |
| LOC107064305 | inhibin beta chain | 0.02911 | 48.14343 | FALSE | FALSE |
| LOC107065517 | glutamate--cysteine ligase catalytic subunit | 0.02874 | 48.15464 | FALSE | FALSE |
| LOC107071589 | uncharacterized protein LOC107071589 | 0.02958 | 48.16923 | FALSE | FALSE |
| LOC107068720 | exportin-2 | 0.03032 | 48.17499 | FALSE | FALSE |
| LOC107067684 | delta-1-pyrroline-5-carboxylate dehydrogen | 0.02938 | 48.17642 | FALSE | FALSE |
| LOC107074548 | uncharacterized protein LOC107074548 | 0.02976 | 48.18129 | FALSE | FALSE |
| LOC107068368 | DNA excision repair protein haywire | 0.02969 | 48.18635 | FALSE | FALSE |
| LOC107065470 | ABC transporter F family member 4 | 0.03009 | 48.19342 | FALSE | FALSE |
| LOC107067345 | insulin-like growth factor-binding protein cc | 0.03001 | 48.19426 | FALSE | FALSE |
| LOC107071258 | actin-related protein 6 | 0.02844 | 48.19969 | FALSE | FALSE |
| LOC107073582 | uncharacterized protein LOC107073582 | 0.03018 | 48.22931 | FALSE | FALSE |
| LOC107070622 | alpha-tocopherol transfer protein-like | 0.02955 | 48.23278 | FALSE | FALSE |
| LOC107070481 | uncharacterized protein LOC107070481 | 0.0291 | 48.25516 | FALSE | FALSE |
| LOC107065076 | uncharacterized protein LOC107065076 | 0.02869 | 48.26481 | FALSE | FALSE |
| LOC107066515 | sodium/potassium-transporting ATPase sub | 0.02942 | 48.2698 | FALSE | FALSE |
| LOC107072183 | UDP-N-acetylglucosamine--dolichyl-phospha | 0.02935 | 48.27544 | FALSE | FALSE |
| LOC107073355 | presequence protease, mitochondrial | 0.02832 | 48.27973 | FALSE | FALSE |
| LOC107073081 | phenoloxidase 2 | 0.0291 | 48.30611 | FALSE | FALSE |
| LOC107066905 | STAM-binding protein-like A | 0.03 | 48.31865 | FALSE | FALSE |
| LOC107069014 | peptidyl-prolyl cis-trans isomerase | 0.02995 | 48.32571 | FALSE | FALSE |
| LOC107068892 | uncharacterized protein LOC107068892 | 0.02783 | 48.32575 | FALSE | FALSE |
| LOC107071756 | actin-related protein 5 | 0.03001 | 48.32622 | FALSE | FALSE |
| LOC107066043 | ADP-ribosylation factor-like protein 2 | 0.02836 | 48.33412 | FALSE | FALSE |
| LOC107071288 | uncharacterized protein LOC107071288 | 0.02845 | 48.36313 | FALSE | FALSE |
| LOC107072042 | intracellular protein transport protein USO1 | 0.0293 | 48.36936 | FALSE | FALSE |
| LOC107067893 | LOW QUALITY PROTEIN: diphthamide biosyı | 0.02835 | 48.43182 | FALSE | FALSE |
| LOC107073672 | cytochrome P450 4g15-like | 0.02913 | 48.43265 | FALSE | FALSE |
| LOC107067692 | protein scabrous-like | 0.029 | 48.43832 | FALSE | FALSE |
| LOC107065304 | leukocyte elastase inhibitor-like | 0.02819 | 48.44694 | FALSE | FALSE |
| LOC107064309 | group XIIA secretory phospholipase A2 | 0.02987 | 48.44837 | FALSE | FALSE |
| LOC107064181 | fatty acid-binding protein, adipocyte-like | 0.02879 | 48.46735 | FALSE | FALSE |
| LOC107071742 | cartilage oligomeric matrix protein | 0.02881 | 48.46758 | FALSE | FALSE |
| LOC107071817 | uncharacterized protein LOC107071817 | 0.02951 | 48.46853 | FALSE | FALSE |
| LOC107068378 | ras-related protein Rab-37-like | 0.02797 | 48.47771 | FALSE | FALSE |
| LOC107067749 | pleckstrin homology domain-containing fam | 0.02896 | 48.49017 | FALSE | FALSE |
| LOC107073071 | saccharopine dehydrogenase-like oxidoredu | 0.02895 | 48.50552 | FALSE | FALSE |
| LOC107067424 | uncharacterized protein LOC107067424 | 0.02912 | 48.5076 | FALSE | FALSE |
| LOC107072186 | uncharacterized protein LOC107072186 | 0.02946 | 48.51145 | FALSE | FALSE |

| | | | | | |
|---|---|---|---|---|---|
| LOC107072736 | cytochrome c oxidase subunit 5A, mitochon | 0.02926 | 48.51363 | FALSE | FALSE |
| LOC107072869 | TP53-regulating kinase | 0.02763 | 48.51688 | FALSE | FALSE |
| LOC107068532 | voltage-dependent calcium channel subunit | 0.02932 | 48.52406 | FALSE | FALSE |
| LOC107070648 | MATH and LRR domain-containing protein P | 0.0291 | 48.52649 | FALSE | FALSE |
| LOC107066210 | CUE domain-containing protein 1 | 0.0288 | 48.53233 | FALSE | FALSE |
| LOC107068274 | uncharacterized protein LOC107068274 | 0.02847 | 48.53641 | FALSE | FALSE |
| LOC107073679 | uncharacterized protein LOC107073679 | 0.02836 | 48.57181 | FALSE | FALSE |
| LOC107068370 | uncharacterized protein LOC107068370 | 0.02821 | 48.57682 | FALSE | FALSE |
| LOC107072471 | integrator complex subunit 9 | 0.02977 | 48.58847 | FALSE | FALSE |
| LOC107064618 | arrestin domain-containing protein 2-like | 0.02917 | 48.62553 | FALSE | FALSE |
| LOC107066193 | tyrosine-protein phosphatase 69D | 0.02968 | 48.62853 | FALSE | FALSE |
| LOC107066100 | uncharacterized protein LOC107066100 | 0.0287 | 48.62923 | FALSE | FALSE |
| LOC107066294 | tubulin glycylase 3A-like | 0.02863 | 48.64391 | FALSE | FALSE |
| LOC107064699 | Krueppel-like factor 3 | 0.02838 | 48.64632 | FALSE | FALSE |
| LOC107071470 | uncharacterized protein LOC107071470 | 0.02842 | 48.65051 | FALSE | FALSE |
| LOC107065837 | venom carboxylesterase-6-like | 0.029 | 48.65677 | FALSE | FALSE |
| LOC107066715 | uncharacterized protein LOC107066715 | 0.02751 | 48.66825 | FALSE | FALSE |
| LOC107073945 | lachesin-like | 0.02957 | 48.69415 | FALSE | FALSE |
| LOC107074068 | uncharacterized protein LOC107074068 | 0.02805 | 48.70091 | FALSE | FALSE |
| LOC107070365 | glucosidase 2 subunit beta | 0.02824 | 48.70284 | FALSE | FALSE |
| LOC107067901 | aldose reductase-like | 0.02968 | 48.71604 | FALSE | FALSE |
| LOC107065004 | uncharacterized protein LOC107065004 | 0.02855 | 48.74534 | FALSE | FALSE |
| LOC107069632 | serine/arginine repetitive matrix protein 1-l | 0.0282 | 48.76095 | FALSE | FALSE |
| LOC107066933 | 39S ribosomal protein L12, mitochondrial | 0.02998 | 48.76193 | FALSE | FALSE |
| LOC107069451 | 39S ribosomal protein L3, mitochondrial | 0.02923 | 48.76618 | FALSE | FALSE |
| LOC107067850 | transcription factor IIIB 90 kDa subunit | 0.02789 | 48.81298 | FALSE | FALSE |
| LOC107073663 | uncharacterized protein LOC107073663 | 0.02808 | 48.82376 | FALSE | FALSE |
| LOC107071094 | odorant receptor 85b-like | 0.0289 | 48.82959 | FALSE | FALSE |
| LOC107067150 | nuclear receptor subfamily 4 group A memb | 0.02927 | 48.83536 | FALSE | FALSE |
| LOC107069975 | odorant receptor coreceptor-like | 0.02949 | 48.84333 | FALSE | FALSE |
| LOC107069505 | uncharacterized protein LOC107069505 | 0.02878 | 48.85725 | FALSE | FALSE |
| LOC107067750 | serine/threonine-protein kinase DDB_G028 | 0.02842 | 48.85972 | FALSE | FALSE |
| LOC107065855 | uncharacterized protein LOC107065855 | 0.02877 | 48.86646 | FALSE | FALSE |
| LOC107069070 | uncharacterized protein LOC107069070 | 0.02789 | 48.86688 | FALSE | FALSE |
| LOC107064573 | nidogen-2 | 0.02777 | 48.87103 | FALSE | FALSE |
| LOC107069435 | replication protein A 32 kDa subunit | 0.02987 | 48.89957 | FALSE | FALSE |
| LOC107069709 | uncharacterized protein LOC107069709 | 0.02874 | 48.90229 | FALSE | FALSE |
| LOC107074489 | uncharacterized protein LOC107074489 | 0.0275 | 48.90578 | FALSE | FALSE |
| LOC107069293 | putative glutathione-specific gamma-glutam | 0.0283 | 48.91694 | FALSE | FALSE |
| LOC107066236 | uncharacterized protein LOC107066236 | 0.02832 | 48.92824 | FALSE | FALSE |
| LOC107065340 | adenosine 3'-phospho 5'-phosphosulfate tra | 0.02766 | 48.95674 | FALSE | FALSE |
| LOC107071558 | uncharacterized protein LOC107071558 | 0.0293 | 48.98176 | FALSE | FALSE |
| LOC107073066 | ubiquilin-1 | 0.0285 | 48.98964 | FALSE | FALSE |
| LOC107069360 | actin-related protein 2/3 complex subunit 1 | 0.03034 | 48.99134 | FALSE | FALSE |
| LOC107070711 | doublesex- and mab-3-related transcription | 0.0285 | 49.00615 | FALSE | FALSE |
| LOC107065763 | uncharacterized protein LOC107065763 | 0.02884 | 49.01127 | FALSE | FALSE |
| LOC107066864 | membralin | 0.02919 | 49.03415 | FALSE | FALSE |
| LOC107070825 | protein misato | 0.0294 | 49.05844 | FALSE | FALSE |
| LOC107070852 | uncharacterized protein LOC107070852 | 0.03013 | 49.06065 | FALSE | FALSE |
| LOC107074069 | uncharacterized protein LOC107074069 | 0.02933 | 49.06364 | FALSE | FALSE |

| | | | | | |
|---|---|---|---|---|---|
| LOC107069568 | sperm-tail PG-rich repeat-containing protein | 0.02864 | 49.06497 | FALSE | FALSE |
| LOC107071974 | polyubiquitin-B | 0.0291 | 49.07263 | FALSE | FALSE |
| LOC107071921 | speckle targeted PIP5K1A-regulated poly(A) | 0.02878 | 49.07529 | FALSE | FALSE |
| LOC107072080 | uncharacterized protein LOC107072080 | 0.02783 | 49.07611 | FALSE | FALSE |
| LOC107074257 | dirigent protein 10-like | 0.02947 | 49.11315 | FALSE | FALSE |
| LOC107066080 | ring canal kelch homolog | 0.028 | 49.12436 | FALSE | FALSE |
| LOC107074241 | androglobin-like | 0.02891 | 49.15884 | FALSE | FALSE |
| LOC107074308 | BAI1-associated protein 3 | 0.02843 | 49.1918 | FALSE | FALSE |
| LOC107064083 | sister chromatid cohesion protein DCC1 | 0.02954 | 49.20662 | FALSE | FALSE |
| LOC107069821 | uncharacterized protein LOC107069821 | 0.02763 | 49.21685 | FALSE | FALSE |
| LOC107074637 | NAD-dependent protein deacetylase sirtuin- | 0.02863 | 49.222 | FALSE | FALSE |
| LOC107069307 | uncharacterized protein LOC107069307 | 0.0282 | 49.22911 | FALSE | FALSE |
| LOC107068476 | uncharacterized protein LOC107068476 | 0.02914 | 49.24746 | FALSE | FALSE |
| LOC107065401 | uncharacterized protein LOC107065401 | 0.02856 | 49.26196 | FALSE | FALSE |
| LOC107073235 | cell cycle checkpoint control protein RAD9A | 0.02857 | 49.26692 | FALSE | FALSE |
| LOC107066735 | putative DNA helicase INO80 | 0.02818 | 49.28919 | FALSE | FALSE |
| LOC107066231 | uncharacterized protein LOC107066231 | 0.02872 | 49.29424 | FALSE | FALSE |
| LOC107074324 | POU domain protein 2, isoform B-like | 0.02883 | 49.2982 | FALSE | FALSE |
| LOC107064666 | uncharacterized protein LOC107064666 | 0.02854 | 49.33564 | FALSE | FALSE |
| LOC107065319 | beta-1,4-N-acetylgalactosaminyltransferase | 0.02883 | 49.33598 | FALSE | FALSE |
| LOC107068295 | protein alan shepard-like | 0.02833 | 49.3384 | FALSE | FALSE |
| LOC107070170 | uncharacterized protein LOC107070170 | 0.02861 | 49.34235 | FALSE | FALSE |
| LOC107069870 | protein tramtrack, alpha isoform-like | 0.02874 | 49.35027 | FALSE | FALSE |
| LOC107073410 | uncharacterized protein LOC107073410 | 0.02945 | 49.35315 | FALSE | FALSE |
| LOC107073525 | uncharacterized protein LOC107073525 | 0.02935 | 49.39593 | FALSE | FALSE |
| LOC107074539 | lipoamide acyltransferase component of bra | 0.02928 | 49.40936 | FALSE | FALSE |
| LOC107071461 | putative fatty acyl-CoA reductase CG8306 | 0.02765 | 49.4429 | FALSE | FALSE |
| LOC107063695 | cell wall integrity and stress response comp | 0.02936 | 49.44696 | FALSE | FALSE |
| LOC107070742 | myosin-VIIa | 0.0284 | 49.44735 | FALSE | FALSE |
| LOC107071029 | uncharacterized protein LOC107071029 | 0.0284 | 49.46286 | FALSE | FALSE |
| LOC107073137 | 60S ribosomal protein L14 | 0.02763 | 49.46861 | FALSE | FALSE |
| LOC107064125 | putative epidermal cell surface receptor | 0.02907 | 49.47035 | FALSE | FALSE |
| LOC107070111 | uncharacterized protein LOC107070111 | 0.0279 | 49.47231 | FALSE | FALSE |
| LOC107066717 | homeotic protein distal-less | 0.02942 | 49.49573 | FALSE | FALSE |
| LOC107070834 | selenium-binding protein 1 | 0.02727 | 49.50102 | FALSE | FALSE |
| LOC107072939 | uncharacterized protein LOC107072939 | 0.02853 | 49.51024 | FALSE | FALSE |
| LOC107071778 | 14-3-3 protein zeta | 0.02838 | 49.52457 | FALSE | FALSE |
| LOC107072714 | paired mesoderm homeobox protein 2-like | 0.02851 | 49.54229 | FALSE | FALSE |
| LOC107066548 | histamine H2 receptor-like | 0.02871 | 49.55547 | FALSE | FALSE |
| LOC107063886 | uncharacterized protein LOC107063886 | 0.02909 | 49.56144 | FALSE | FALSE |
| LOC107067964 | ankyrin repeat domain-containing protein 4 | 0.0287 | 49.57826 | FALSE | FALSE |
| LOC107072677 | histidine decarboxylase | 0.02841 | 49.57948 | FALSE | FALSE |
| LOC107067531 | zinc finger X-chromosomal protein-like | 0.02943 | 49.60333 | FALSE | FALSE |
| LOC107072193 | T-complex-associated testis-expressed prot | 0.02736 | 49.62056 | FALSE | FALSE |
| LOC107070833 | ceramide glucosyltransferase | 0.02917 | 49.62872 | FALSE | FALSE |
| LOC107068607 | serine/threonine-protein kinase mig-15 | 0.02971 | 49.62914 | FALSE | FALSE |
| LOC107074066 | homeobox protein Nkx-2.5-like | 0.02848 | 49.63087 | FALSE | FALSE |
| LOC107071245 | 40S ribosomal protein S9 | 0.02797 | 49.64757 | FALSE | FALSE |
| LOC107068297 | ribosome biogenesis protein WDR12 homol | 0.02911 | 49.68163 | FALSE | FALSE |
| LOC107073683 | fatty acyl-CoA reductase 1-like | 0.02932 | 49.68708 | FALSE | FALSE |

| | | | | | |
|---|---|---|---|---|---|
| LOC107066572 | caltractin | 0.02964 | 49.70318 | FALSE | FALSE |
| LOC107072391 | flap endonuclease 1 | 0.02845 | 49.70932 | FALSE | FALSE |
| LOC107066186 | thioredoxin domain-containing protein 5 | 0.02887 | 49.71655 | FALSE | FALSE |
| LOC107068823 | tyrosine-protein phosphatase non-receptor | 0.02885 | 49.72184 | FALSE | FALSE |
| LOC107073234 | endonuclease G, mitochondrial | 0.02883 | 49.72585 | FALSE | FALSE |
| LOC107072548 | uncharacterized protein LOC107072548 | 0.03037 | 49.74858 | FALSE | FALSE |
| LOC107065265 | cyclin-dependent kinase 8 | 0.02797 | 49.75759 | FALSE | FALSE |
| LOC107068259 | actin-related protein 10 | 0.02887 | 49.75781 | FALSE | FALSE |
| LOC107065262 | uncharacterized protein LOC107065262 | 0.02737 | 49.75909 | FALSE | FALSE |
| LOC107070666 | uncharacterized protein LOC107070666 | 0.02823 | 49.77356 | FALSE | FALSE |
| LOC107073416 | uncharacterized protein LOC107073416 | 0.02891 | 49.77401 | FALSE | FALSE |
| LOC107074517 | WD repeat-containing protein 24 | 0.02873 | 49.77572 | FALSE | FALSE |
| LOC107067336 | lysophospholipid acyltransferase 1 | 0.02902 | 49.77716 | FALSE | FALSE |
| LOC107066342 | glutamate receptor ionotropic, delta-2 | 0.02833 | 49.7797 | FALSE | FALSE |
| LOC107066504 | ER membrane protein complex subunit 1 | 0.02931 | 49.78253 | FALSE | FALSE |
| LOC107066457 | rap1 GTPase-GDP dissociation stimulator 1- | 0.0287 | 49.79725 | FALSE | FALSE |
| LOC107066450 | tubulin--tyrosine ligase-like protein 12 | 0.03056 | 49.79994 | FALSE | FALSE |
| LOC107063642 | SUMO-activating enzyme subunit 2 | 0.02871 | 49.82518 | FALSE | FALSE |
| LOC107066351 | zinc finger protein 665-like | 0.0284 | 49.8496 | FALSE | FALSE |
| LOC107072699 | serine/threonine-protein kinase PAK 3 | 0.02853 | 49.8519 | FALSE | FALSE |
| LOC107067956 | general odorant-binding protein 56d-like | 0.02933 | 49.85397 | FALSE | FALSE |
| LOC107066723 | iroquois-class homeodomain protein IRX-6-l | 0.02872 | 49.85486 | FALSE | FALSE |
| LOC107073636 | putative nuclease HARBI1 | 0.02869 | 49.88464 | FALSE | FALSE |
| LOC107072384 | glycogen synthase kinase-3 beta-like | 0.02951 | 49.8877 | FALSE | FALSE |
| LOC107066310 | protein l(2)37Cc | 0.02986 | 49.88836 | FALSE | FALSE |
| LOC107064882 | frizzled-2 | 0.02809 | 49.88842 | FALSE | FALSE |
| LOC107068637 | uncharacterized protein LOC107068637 | 0.02947 | 49.89018 | FALSE | FALSE |
| LOC107068583 | zinc finger CCHC domain-containing protein | 0.02989 | 49.89353 | FALSE | FALSE |
| LOC107072876 | liprin-beta-1 | 0.02961 | 49.89963 | FALSE | FALSE |
| LOC107068803 | solute carrier family 26 member 10 | 0.02915 | 49.90046 | FALSE | FALSE |
| LOC107071367 | protein son of sevenless | 0.02902 | 49.90298 | FALSE | FALSE |
| LOC107067853 | phosphoacetylglucosamine mutase | 0.02773 | 49.90861 | FALSE | FALSE |
| LOC107070075 | serine palmitoyltransferase 1 | 0.02884 | 49.91549 | FALSE | FALSE |
| LOC107069047 | innexin inx2-like | 0.02801 | 49.91661 | FALSE | FALSE |
| LOC107064464 | probable phospholipid-transporting ATPase | 0.02744 | 49.92323 | FALSE | FALSE |
| LOC107068420 | leucine carboxyl methyltransferase 1 | 0.02805 | 49.94234 | FALSE | FALSE |
| LOC107064830 | uncharacterized protein LOC107064830 | 0.02785 | 49.94913 | FALSE | FALSE |
| LOC107070548 | ectonucleoside triphosphate diphosphohydı | 0.02882 | 49.95147 | FALSE | FALSE |
| LOC107064299 | Down syndrome cell adhesion molecule-like | 0.0287 | 49.95846 | FALSE | FALSE |
| LOC107070383 | dipeptidase 1-like | 0.02909 | 49.97163 | FALSE | FALSE |
| LOC107073775 | uncharacterized protein LOC107073775 | 0.02766 | 49.99461 | FALSE | FALSE |
| LOC107066595 | vacuolar protein sorting-associated protein | 0.02903 | 50.01032 | FALSE | FALSE |
| LOC107064051 | uncharacterized protein LOC107064051 | 0.02851 | 50.02219 | FALSE | FALSE |
| LOC107066413 | ribosome production factor 2 homolog | 0.02892 | 50.03465 | FALSE | FALSE |
| LOC107074134 | methylthioribulose-1-phosphate dehydratas | 0.02886 | 50.03774 | FALSE | FALSE |
| LOC107066390 | insulin receptor substrate 1 | 0.02866 | 50.04958 | FALSE | FALSE |
| LOC107066968 | phospholipase A1 member A | 0.02787 | 50.06051 | FALSE | FALSE |
| LOC107073535 | uncharacterized protein LOC107073535 | 0.02882 | 50.06336 | FALSE | FALSE |
| LOC107067928 | cytochrome P450 4C1-like | 0.0278 | 50.0861 | FALSE | FALSE |
| LOC107066769 | peptidoglycan recognition protein-like | 0.02889 | 50.10898 | FALSE | FALSE |

| | | | | | |
|---|---|---|---|---|---|
| LOC107071378 | myosin-M heavy chain | 0.02816 | 50.14026 | FALSE | FALSE |
| LOC107071861 | proteasome subunit alpha type-5 | 0.02915 | 50.15933 | FALSE | FALSE |
| LOC107067061 | ATPase WRNIP1-like | 0.02857 | 50.16707 | FALSE | FALSE |
| LOC107074466 | alpha-actinin, sarcomeric | 0.02908 | 50.18761 | FALSE | FALSE |
| LOC107066478 | very-long-chain 3-oxoacyl-CoA reductase-lik | 0.02958 | 50.18839 | FALSE | FALSE |
| LOC107072713 | tetratricopeptide repeat protein 25-like | 0.02735 | 50.19404 | FALSE | FALSE |
| LOC107064446 | UV excision repair protein RAD23 homolog I | 0.0285 | 50.19674 | FALSE | FALSE |
| LOC107071047 | cyclin-dependent kinase 14 | 0.02699 | 50.20957 | FALSE | FALSE |
| LOC107066074 | uncharacterized protein LOC107066074 | 0.02868 | 50.22841 | FALSE | FALSE |
| LOC107073118 | uncharacterized protein LOC107073118 | 0.02802 | 50.24341 | FALSE | FALSE |
| Trnar-acg | Trnar-acg | 0.02897 | 50.26777 | FALSE | FALSE |
| LOC107071444 | coiled-coil domain-containing protein 93 | 0.02803 | 50.27414 | FALSE | FALSE |
| LOC107071525 | alpha/beta hydrolase domain-containing pr | 0.0294 | 50.27889 | FALSE | FALSE |
| LOC107064641 | lysine-specific histone demethylase 1A | 0.03035 | 50.27971 | FALSE | FALSE |
| LOC107069566 | zinc finger protein 436-like | 0.02704 | 50.29137 | FALSE | FALSE |
| LOC107065248 | venom protease-like | 0.02859 | 50.29247 | FALSE | FALSE |
| LOC107070271 | calcineurin subunit B type 2 | 0.02833 | 50.30408 | FALSE | FALSE |
| LOC107071605 | protein lunapark-B | 0.02901 | 50.30804 | FALSE | FALSE |
| LOC107067883 | uncharacterized protein LOC107067883 | 0.028 | 50.30869 | FALSE | FALSE |
| LOC107072667 | collagen alpha chain CG42342 | 0.02845 | 50.31165 | FALSE | FALSE |
| LOC107066229 | sodium-coupled monocarboxylate transport | 0.02988 | 50.31463 | FALSE | FALSE |
| LOC107073173 | sodium- and chloride-dependent GABA tran | 0.02885 | 50.31471 | FALSE | FALSE |
| LOC107069545 | uncharacterized protein LOC107069545 | 0.02864 | 50.31552 | FALSE | FALSE |
| LOC107065583 | dynein beta chain, ciliary | 0.02797 | 50.31784 | FALSE | FALSE |
| LOC107068563 | pickpocket protein 19-like | 0.02823 | 50.35389 | FALSE | FALSE |
| LOC107063845 | pyruvate dehydrogenase E1 component sub | 0.02839 | 50.3655 | FALSE | FALSE |
| LOC107067381 | uncharacterized protein LOC107067381 | 0.02874 | 50.36597 | FALSE | FALSE |
| LOC107071904 | gonadotropin-releasing hormone II receptor | 0.02859 | 50.37011 | FALSE | FALSE |
| LOC107069850 | protein C12orf4 homolog | 0.02724 | 50.37624 | FALSE | FALSE |
| LOC107073883 | titin-like | 0.02959 | 50.38057 | FALSE | FALSE |
| LOC107066692 | E3 ubiquitin-protein ligase RNF13-like | 0.02836 | 50.39541 | FALSE | FALSE |
| LOC107069962 | tonsoku-like protein | 0.02757 | 50.40751 | FALSE | FALSE |
| LOC107071953 | uncharacterized protein LOC107071953 | 0.02971 | 50.45245 | FALSE | FALSE |
| LOC107063702 | uncharacterized protein LOC107063702 | 0.02841 | 50.46578 | FALSE | FALSE |
| LOC107070510 | uncharacterized protein LOC107070510 | 0.02875 | 50.48097 | FALSE | FALSE |
| LOC107073895 | tubulin-specific chaperone A | 0.02892 | 50.48435 | FALSE | FALSE |
| LOC107068490 | organic cation transporter protein-like | 0.02892 | 50.50296 | FALSE | FALSE |
| LOC107069346 | uncharacterized protein LOC107069346 | 0.02965 | 50.50373 | FALSE | FALSE |
| LOC107073785 | uncharacterized protein LOC107073785 | 0.02777 | 50.50693 | FALSE | FALSE |
| LOC107072840 | uncharacterized protein LOC107072840 | 0.02749 | 50.54956 | FALSE | FALSE |
| LOC107069022 | uncharacterized protein LOC107069022 | 0.02891 | 50.60448 | FALSE | FALSE |
| LOC107071090 | protocadherin Fat 4 | 0.02897 | 50.61854 | FALSE | FALSE |
| LOC107068653 | uncharacterized protein LOC107068653 | 0.02819 | 50.62327 | FALSE | FALSE |
| LOC107066491 | hexokinase type 2 | 0.02799 | 50.6358 | FALSE | FALSE |
| LOC107069713 | high affinity copper uptake protein 1 | 0.02887 | 50.63661 | FALSE | FALSE |
| LOC107069141 | 39S ribosomal protein L15, mitochondrial | 0.02723 | 50.63822 | FALSE | FALSE |
| LOC107067505 | uncharacterized protein LOC107067505 | 0.02826 | 50.6412 | FALSE | FALSE |
| LOC107065122 | GDP-fucose transporter 1 | 0.02781 | 50.65904 | FALSE | FALSE |
| LOC107067202 | uncharacterized protein LOC107067202 | 0.02738 | 50.67655 | FALSE | FALSE |
| LOC107064604 | melanotransferrin | 0.02897 | 50.68508 | FALSE | FALSE |

| | | | | | |
|---|---|---|---|---|---|
| LOC107068954 | putative odorant receptor 85e | 0.02959 | 50.68594 | FALSE | FALSE |
| LOC107066462 | uncharacterized protein LOC107066462 | 0.0275 | 50.73169 | FALSE | FALSE |
| LOC107073096 | ARL14 effector protein | 0.03033 | 50.75096 | FALSE | FALSE |
| LOC107064474 | decaprenyl-diphosphate synthase subunit 2 | 0.02828 | 50.75732 | FALSE | FALSE |
| LOC107070173 | uncharacterized protein LOC107070173 | 0.02775 | 50.76484 | FALSE | FALSE |
| LOC107064977 | lariat debranching enzyme | 0.02933 | 50.79608 | FALSE | FALSE |
| LOC107073088 | uncharacterized protein C1orf50 homolog | 0.02675 | 50.79635 | FALSE | FALSE |
| LOC107072985 | nuclear transcription factor Y subunit B-4 | 0.02803 | 50.79925 | FALSE | FALSE |
| LOC107064277 | alpha-1,3-mannosyl-glycoprotein 4-beta-N-a | 0.028 | 50.83112 | FALSE | FALSE |
| LOC107064892 | heterogeneous nuclear ribonucleoprotein U | 0.02765 | 50.84561 | FALSE | FALSE |
| LOC107068639 | uncharacterized protein LOC107068639 | 0.02662 | 50.87411 | FALSE | FALSE |
| LOC107068730 | sodium-dependent proline transporter-like | 0.0279 | 50.87922 | FALSE | FALSE |
| LOC107063940 | acyl-CoA Delta(11) desaturase-like | 0.02808 | 50.88373 | FALSE | FALSE |
| LOC107071995 | alpha-amylase-like | 0.02764 | 50.89079 | FALSE | FALSE |
| LOC107063756 | charged multivesicular body protein 5 | 0.03023 | 50.89196 | FALSE | FALSE |
| LOC107073188 | ras guanine nucleotide exchange factor B | 0.03006 | 50.90765 | FALSE | FALSE |
| LOC107065638 | stress-activated protein kinase JNK | 0.02744 | 50.91322 | FALSE | FALSE |
| LOC107071364 | TM2 domain-containing protein CG10795 | 0.02877 | 50.92641 | FALSE | FALSE |
| LOC107066985 | histone-arginine methyltransferase CARMEI | 0.02883 | 50.9555 | FALSE | FALSE |
| LOC107072000 | beta-1,4-glucuronyltransferase 1 | 0.02819 | 50.9796 | FALSE | FALSE |
| LOC107064899 | 85/88 kDa calcium-independent phospholip | 0.02876 | 50.98216 | FALSE | FALSE |
| LOC107064867 | protein croquemort-like | 0.02754 | 50.9948 | FALSE | FALSE |
| LOC107072573 | mediator of RNA polymerase II transcriptior | 0.02831 | 50.9954 | FALSE | FALSE |
| LOC107064678 | tripeptidyl-peptidase 2 | 0.02899 | 51.00769 | FALSE | FALSE |
| LOC107065318 | ubiquitin carboxyl-terminal hydrolase 1 | 0.02703 | 51.01214 | FALSE | FALSE |
| LOC107072841 | uncharacterized protein LOC107072841 | 0.0284 | 51.02376 | FALSE | FALSE |
| LOC107073450 | uncharacterized protein LOC107073450 | 0.02905 | 51.04141 | FALSE | FALSE |
| LOC107068015 | uncharacterized protein LOC107068015 | 0.02765 | 51.06235 | FALSE | FALSE |
| LOC107063769 | uncharacterized protein LOC107063769 | 0.0279 | 51.08116 | FALSE | FALSE |
| LOC107072085 | exonuclease 3'-5' domain-containing protei | 0.02812 | 51.08609 | FALSE | FALSE |
| LOC107071207 | elongation factor Tu, mitochondrial-like | 0.02812 | 51.10194 | FALSE | FALSE |
| LOC107066616 | free fatty acid receptor 4-like | 0.02727 | 51.10771 | FALSE | FALSE |
| LOC107064296 | protein THEM6-like | 0.02702 | 51.12956 | FALSE | FALSE |
| LOC107066794 | protein unc-45 homolog B | 0.02763 | 51.138 | FALSE | FALSE |
| LOC107069717 | probable E3 ubiquitin-protein ligase RNF14 | 0.02774 | 51.16807 | FALSE | FALSE |
| LOC107067484 | axonemal 84 kDa protein-like | 0.02663 | 51.1738 | FALSE | FALSE |
| LOC107073390 | uncharacterized protein LOC107073390 | 0.02863 | 51.17585 | FALSE | FALSE |
| LOC107072014 | slit homolog 2 protein-like | 0.02753 | 51.17738 | FALSE | FALSE |
| LOC107064131 | homeobox protein Nkx-2.4 | 0.02834 | 51.17781 | FALSE | FALSE |
| LOC107068382 | piggyBac transposable element-derived pro | 0.02997 | 51.19547 | FALSE | FALSE |
| LOC107071907 | uncharacterized protein LOC107071907 | 0.02819 | 51.21034 | FALSE | FALSE |
| LOC107068990 | kelch-like protein diablo | 0.02756 | 51.22221 | FALSE | FALSE |
| LOC107068421 | tRNA pseudouridine(38/39) synthase | 0.02824 | 51.25236 | FALSE | FALSE |
| LOC107070488 | stress response protein NST1-like | 0.02824 | 51.25425 | FALSE | FALSE |
| LOC107072723 | uncharacterized protein LOC107072723 | 0.02866 | 51.25503 | FALSE | FALSE |
| LOC107072019 | probable serine/threonine-protein kinase D | 0.02858 | 51.2618 | FALSE | FALSE |
| LOC107069764 | fibrillin-2-like | 0.02872 | 51.27167 | FALSE | FALSE |
| LOC107065192 | glycine-rich cell wall structural protein 2-like | 0.02755 | 51.27355 | FALSE | FALSE |
| LOC107069603 | putative nuclease HARBI1 | 0.02749 | 51.27356 | FALSE | FALSE |
| LOC107065593 | LOW QUALITY PROTEIN: uncharacterized th | 0.02631 | 51.28789 | FALSE | FALSE |

| | | | | | |
|---|---|---|---|---|---|
| LOC107071022 | major facilitator superfamily domain-contai | 0.03037 | 51.28949 | FALSE | FALSE |
| LOC107069913 | metaxin-2-like | 0.02855 | 51.2935 | FALSE | FALSE |
| LOC107064209 | uncharacterized protein LOC107064209 | 0.02733 | 51.29572 | FALSE | FALSE |
| LOC107066706 | glutamate decarboxylase | 0.02806 | 51.30419 | FALSE | FALSE |
| LOC107070782 | uncharacterized protein LOC107070782 | 0.02823 | 51.30859 | FALSE | FALSE |
| LOC107065542 | sulfotransferase family cytosolic 1B membe | 0.02849 | 51.3099 | FALSE | FALSE |
| LOC107067134 | proton-coupled amino acid transporter 1 | 0.02772 | 51.31364 | FALSE | FALSE |
| LOC107069231 | dehydrogenase/reductase SDR family meml | 0.02893 | 51.3144 | FALSE | FALSE |
| LOC107069989 | cysteine and histidine-rich domain-containi | 0.02838 | 51.32444 | FALSE | FALSE |
| LOC107068930 | venom protease-like | 0.02843 | 51.32496 | FALSE | FALSE |
| LOC107063962 | probable serine/threonine-protein kinase D | 0.02701 | 51.33475 | FALSE | FALSE |
| LOC107073383 | LOW QUALITY PROTEIN: polyglutamine-repe | 0.02847 | 51.34922 | FALSE | FALSE |
| LOC107071892 | 40S ribosomal protein S16 | 0.02711 | 51.36806 | FALSE | FALSE |
| LOC107067086 | piezo-type mechanosensitive ion channel co | 0.02739 | 51.37115 | FALSE | FALSE |
| LOC107074571 | circadian locomoter output cycles protein k | 0.02729 | 51.37506 | FALSE | FALSE |
| LOC107073031 | translation factor GUF1 homolog, mitochon | 0.02737 | 51.38827 | FALSE | FALSE |
| LOC107069588 | uncharacterized protein LOC107069588 | 0.02913 | 51.38922 | FALSE | FALSE |
| LOC107070814 | membrane magnesium transporter 1 | 0.02734 | 51.39114 | FALSE | FALSE |
| LOC107064362 | growth/differentiation factor 11 | 0.02825 | 51.40326 | FALSE | FALSE |
| LOC107065288 | neurofibromin | 0.02871 | 51.42608 | FALSE | FALSE |
| LOC107067702 | oxysterol-binding protein-related protein 9 | 0.02799 | 51.43561 | FALSE | FALSE |
| LOC107066042 | protein RTF2 homolog | 0.02773 | 51.43622 | FALSE | FALSE |
| LOC107063757 | protein tincar | 0.02868 | 51.43982 | FALSE | FALSE |
| LOC107071524 | homer protein homolog 2 | 0.02772 | 51.4477 | FALSE | FALSE |
| LOC107069594 | nuclear migration protein nudC | 0.02921 | 51.45177 | FALSE | FALSE |
| LOC107064076 | leucine-rich repeat-containing protein 15-lik | 0.02917 | 51.45457 | FALSE | FALSE |
| LOC107063958 | probable E3 ubiquitin-protein ligase makori | 0.02798 | 51.46849 | FALSE | FALSE |
| LOC107064764 | alpha-2A adrenergic receptor | 0.02834 | 51.4798 | FALSE | FALSE |
| LOC107073487 | uncharacterized protein LOC107073487 | 0.0285 | 51.48569 | FALSE | FALSE |
| LOC107071109 | vascular endothelial growth factor C | 0.02737 | 51.50945 | FALSE | FALSE |
| LOC107071845 | uncharacterized protein LOC107071845 | 0.02675 | 51.52453 | FALSE | FALSE |
| LOC107072883 | ADP-ribosylation factor 2 | 0.02753 | 51.53664 | FALSE | FALSE |
| LOC107068752 | beta-1-syntrophin | 0.02891 | 51.53945 | FALSE | FALSE |
| LOC107069773 | guanine nucleotide-binding protein-like 3 ho | 0.02876 | 51.55354 | FALSE | FALSE |
| LOC107067899 | cytochrome P450 4C1-like | 0.02869 | 51.55495 | FALSE | FALSE |
| LOC107070238 | sortilin-related receptor-like | 0.02796 | 51.56384 | FALSE | FALSE |
| LOC107069528 | arylsulfatase B-like | 0.02724 | 51.5703 | FALSE | FALSE |
| LOC107066776 | MATH and LRR domain-containing protein P | 0.02883 | 51.57774 | FALSE | FALSE |
| LOC107068191 | uncharacterized protein LOC107068191 | 0.0275 | 51.59666 | FALSE | FALSE |
| LOC107070838 | RAD50-interacting protein 1 | 0.02924 | 51.60008 | FALSE | FALSE |
| LOC107066490 | protein FAM160B1-like | 0.02866 | 51.60163 | FALSE | FALSE |
| LOC107070908 | protocadherin-15 | 0.02743 | 51.60924 | FALSE | FALSE |
| LOC107073901 | alaserpin-like | 0.02801 | 51.62955 | FALSE | FALSE |
| LOC107071305 | MOB kinase activator-like 3 | 0.0283 | 51.63922 | FALSE | FALSE |
| LOC107064393 | protein PFC0760c | 0.02841 | 51.64607 | FALSE | FALSE |
| LOC107064432 | integrator complex subunit 12-like | 0.02649 | 51.6558 | FALSE | FALSE |
| LOC107066578 | signal peptidase complex subunit 3 | 0.02929 | 51.66725 | FALSE | FALSE |
| LOC107069667 | dystrophin-related protein 2-like | 0.02853 | 51.6779 | FALSE | FALSE |
| LOC107070284 | apolipoprotein D-like | 0.02723 | 51.70545 | FALSE | FALSE |
| LOC107067137 | superoxide dismutase [Cu-Zn | 0.02808 | 51.71444 | FALSE | FALSE |

| | | | | | |
|---|---|---|---|---|---|
| LOC107064968 | BMP-2-inducible protein kinase | 0.02695 | 51.72323 | FALSE | FALSE |
| LOC107068787 | orexin receptor type 2-like | 0.02861 | 51.74998 | FALSE | FALSE |
| LOC107068974 | innexin inx7-like | 0.02663 | 51.76442 | FALSE | FALSE |
| LOC107065007 | cytochrome b5 reductase 4 | 0.0273 | 51.79219 | FALSE | FALSE |
| LOC107065748 | uncharacterized protein LOC107065748 | 0.02863 | 51.7949 | FALSE | FALSE |
| LOC107072951 | cytochrome c1-2, heme protein, mitochond | 0.02759 | 51.80416 | FALSE | FALSE |
| LOC107071866 | myosin heavy chain, muscle | 0.02647 | 51.80922 | FALSE | FALSE |
| LOC107067746 | kyphoscoliosis peptidase | 0.02856 | 51.82384 | FALSE | FALSE |
| LOC107074302 | uncharacterized protein LOC107074302 | 0.02881 | 51.83274 | FALSE | FALSE |
| LOC107068847 | CAD protein | 0.02785 | 51.84269 | FALSE | FALSE |
| LOC107069791 | uncharacterized protein LOC107069791 | 0.02965 | 51.85035 | FALSE | FALSE |
| LOC107065191 | altered inheritance of mitochondria protein | 0.02824 | 51.85517 | FALSE | FALSE |
| LOC107070674 | uncharacterized protein LOC107070674 | 0.02787 | 51.86455 | FALSE | FALSE |
| LOC107070226 | protein CLEC16A | 0.02885 | 51.86844 | FALSE | FALSE |
| LOC107069985 | transcription elongation factor B polypeptid | 0.02879 | 51.87777 | FALSE | FALSE |
| LOC107074696 | peroxisomal membrane protein PEX13 | 0.02844 | 51.87995 | FALSE | FALSE |
| LOC107069544 | uncharacterized protein LOC107069544 | 0.02683 | 51.88178 | FALSE | FALSE |
| LOC107068497 | viral IAP-associated factor homolog | 0.0274 | 51.89075 | FALSE | FALSE |
| LOC107071122 | band 4.1-like protein 4 | 0.02704 | 51.89135 | FALSE | FALSE |
| LOC107071868 | palmitoyltransferase ZDHHC17 | 0.02813 | 51.89348 | FALSE | FALSE |
| LOC107066897 | uncharacterized protein LOC107066897 | 0.02765 | 51.90826 | FALSE | FALSE |
| LOC107073010 | homeobox protein prospero | 0.02793 | 51.9467 | FALSE | FALSE |
| LOC107074525 | adenomatous polyposis coli homolog | 0.02903 | 51.9473 | FALSE | FALSE |
| LOC107072415 | coiled-coil-helix-coiled-coil-helix domain-co | 0.0277 | 51.95084 | FALSE | FALSE |
| LOC107065677 | dynein heavy chain 6, axonemal | 0.02952 | 52.02243 | FALSE | FALSE |
| LOC107068233 | probable 39S ribosomal protein L49, mitoch | 0.02838 | 52.02542 | FALSE | FALSE |
| LOC107066691 | uncharacterized protein LOC107066691 | 0.02864 | 52.05562 | FALSE | FALSE |
| LOC107063888 | transcription factor 25 | 0.02649 | 52.07788 | FALSE | FALSE |
| LOC107068857 | uncharacterized protein LOC107068857 | 0.02844 | 52.09263 | FALSE | FALSE |
| LOC107068509 | uncharacterized protein LOC107068509 | 0.02781 | 52.09362 | FALSE | FALSE |
| LOC107070204 | probable citrate synthase 2, mitochondrial | 0.02888 | 52.09618 | FALSE | FALSE |
| LOC107072148 | uncharacterized protein LOC107072148 | 0.02805 | 52.11386 | FALSE | FALSE |
| LOC107065833 | DNA repair protein XRCC1 | 0.02712 | 52.1285 | FALSE | FALSE |
| LOC107066044 | uncharacterized protein LOC107066044 | 0.02772 | 52.13105 | FALSE | FALSE |
| LOC107069145 | phospholipase B1, membrane-associated-lik | 0.0286 | 52.16253 | FALSE | FALSE |
| LOC107067844 | dual specificity protein phosphatase 12-like | 0.02876 | 52.1643 | FALSE | FALSE |
| LOC107065379 | protein kinase C and casein kinase substrate | 0.02802 | 52.1748 | FALSE | FALSE |
| LOC107068681 | dynein light chain roadblock-type 2 | 0.02786 | 52.18318 | FALSE | FALSE |
| LOC107071494 | nucleolar protein 56 | 0.02766 | 52.19753 | FALSE | FALSE |
| LOC107074595 | uncharacterized protein LOC107074595 | 0.02841 | 52.21992 | FALSE | FALSE |
| LOC107066175 | micronuclear linker histone polyprotein-like | 0.0295 | 52.2222 | FALSE | FALSE |
| LOC107065043 | dopamine D2-like receptor | 0.02704 | 52.25046 | FALSE | FALSE |
| LOC107069042 | trypsin-7-like | 0.02676 | 52.25398 | FALSE | FALSE |
| LOC107065189 | kelch-like protein 10 | 0.02764 | 52.26269 | FALSE | FALSE |
| LOC107069900 | uncharacterized protein LOC107069900 | 0.0272 | 52.27632 | FALSE | FALSE |
| LOC107064227 | uncharacterized protein LOC107064227 | 0.02842 | 52.33801 | FALSE | FALSE |
| LOC107067541 | uncharacterized protein LOC107067541 | 0.02833 | 52.3488 | FALSE | FALSE |
| LOC107070775 | ATP-binding cassette sub-family D member | 0.02871 | 52.35027 | FALSE | FALSE |
| LOC107063688 | ferritin light chain, oocyte isoform | 0.0273 | 52.35177 | FALSE | FALSE |
| LOC107067169 | SHC-transforming protein 1 | 0.02678 | 52.35352 | FALSE | FALSE |

| | | | | | |
|---|---|---|---|---|---|
| LOC107064908 | ATP-binding cassette sub-family E member 1 | 0.02782 | 52.37825 | FALSE | FALSE |
| LOC107073978 | cadherin EGF LAG seven-pass G-type recept | 0.02726 | 52.3809 | FALSE | FALSE |
| LOC107071117 | uncharacterized protein LOC107071117 | 0.02682 | 52.41156 | FALSE | FALSE |
| LOC107065742 | cationic amino acid transporter 2 | 0.02859 | 52.45663 | FALSE | FALSE |
| LOC107074433 | la-related protein 6 | 0.02803 | 52.46986 | FALSE | FALSE |
| LOC107070516 | uncharacterized protein LOC107070516 | 0.02793 | 52.47614 | FALSE | FALSE |
| LOC107063618 | protein toll-like | 0.02761 | 52.47882 | FALSE | FALSE |
| LOC107064639 | ATP-dependent DNA helicase Q5-like | 0.02606 | 52.50541 | FALSE | FALSE |
| LOC107071630 | probable serine/threonine-protein kinase ts | 0.02757 | 52.50703 | FALSE | FALSE |
| LOC107069318 | protein starmaker-like | 0.02756 | 52.50744 | FALSE | FALSE |
| LOC107072121 | cullin-5 | 0.02832 | 52.53632 | FALSE | FALSE |
| LOC107069509 | uncharacterized protein LOC107069509 | 0.02899 | 52.53986 | FALSE | FALSE |
| LOC107065987 | dedicator of cytokinesis protein 1 | 0.02665 | 52.54432 | FALSE | FALSE |
| LOC107067694 | uncharacterized protein LOC107067694 | 0.02684 | 52.54478 | FALSE | FALSE |
| LOC107071420 | squamous cell carcinoma antigen recognize | 0.0261 | 52.54945 | FALSE | FALSE |
| LOC107073093 | protein PFC0760c | 0.02698 | 52.57937 | FALSE | FALSE |
| LOC107069037 | midasin | 0.027 | 52.61337 | FALSE | FALSE |
| LOC107067004 | T-complex protein 1 subunit alpha | 0.02825 | 52.61922 | FALSE | FALSE |
| LOC107071278 | mitochondrial GTPase 1 | 0.02711 | 52.62079 | FALSE | FALSE |
| LOC107066742 | uncharacterized protein LOC107066742 | 0.02804 | 52.67649 | FALSE | FALSE |
| LOC107066809 | acetylcholine receptor subunit beta-type ac | 0.02851 | 52.68739 | FALSE | FALSE |
| LOC107069551 | uncharacterized protein LOC107069551 | 0.02754 | 52.75199 | FALSE | FALSE |
| LOC107068372 | leucine-rich repeat serine/threonine-protei | 0.02596 | 52.76574 | FALSE | FALSE |
| LOC107067630 | general transcription factor IIE subunit 1 | 0.02818 | 52.76912 | FALSE | FALSE |
| LOC107065603 | prothoracicotropic hormone-like | 0.02767 | 52.77207 | FALSE | FALSE |
| LOC107070328 | uncharacterized protein LOC107070328 | 0.02869 | 52.78174 | FALSE | FALSE |
| LOC107073038 | centrosomal protein of 290 kDa | 0.02869 | 52.8183 | FALSE | FALSE |
| LOC107064021 | SAGA-associated factor 29 homolog | 0.02774 | 52.82441 | FALSE | FALSE |
| LOC107072771 | UPF0489 protein C5orf22 homolog | 0.02786 | 52.82676 | FALSE | FALSE |
| LOC107069138 | transmembrane protein 231-like | 0.0283 | 52.8353 | FALSE | FALSE |
| LOC107065781 | radial spoke head 1 homolog | 0.02777 | 52.83812 | FALSE | FALSE |
| LOC107069407 | BTB/POZ domain-containing protein 1-like | 0.02765 | 52.83869 | FALSE | FALSE |
| LOC107072106 | lysosomal thioesterase PPT2 homolog | 0.02829 | 52.84138 | FALSE | FALSE |
| LOC107072988 | N-acetyl-D-glucosamine kinase | 0.02785 | 52.85725 | FALSE | FALSE |
| LOC107068199 | uncharacterized protein LOC107068199 | 0.02887 | 52.86301 | FALSE | FALSE |
| LOC107071774 | importin-7 | 0.02748 | 52.88226 | FALSE | FALSE |
| LOC107074035 | phospholipase A1-like | 0.02821 | 52.89382 | FALSE | FALSE |
| LOC107072485 | digestive organ expansion factor homolog | 0.02755 | 52.91104 | FALSE | FALSE |
| LOC107071970 | uncharacterized protein LOC107071970 | 0.02783 | 52.91693 | FALSE | FALSE |
| LOC107067019 | lys-63-specific deubiquitinase BRCC36-like | 0.02697 | 52.92387 | FALSE | FALSE |
| LOC107067687 | lysosomal aspartic protease | 0.02825 | 52.94435 | FALSE | FALSE |
| LOC107066341 | uncharacterized protein LOC107066341 | 0.02855 | 52.9556 | FALSE | FALSE |
| LOC107071673 | tachykinin-like peptides receptor 86C | 0.02714 | 52.96845 | FALSE | FALSE |
| LOC107068748 | WD repeat-containing protein 44 | 0.02865 | 52.98377 | FALSE | FALSE |
| LOC107065008 | fatty-acid amide hydrolase 2 | 0.02724 | 52.99732 | FALSE | FALSE |
| LOC107070120 | uncharacterized protein LOC107070120 | 0.02843 | 53.00186 | FALSE | FALSE |
| LOC107074233 | uncharacterized protein LOC107074233 | 0.0275 | 53.00791 | FALSE | FALSE |
| LOC107072434 | zinc transporter 7 | 0.02763 | 53.01343 | FALSE | FALSE |
| LOC107066610 | actin-related protein 2/3 complex subunit 5 | 0.02666 | 53.02005 | FALSE | FALSE |
| LOC107069025 | GTPase-activating Rap/Ran-GAP domain-like | 0.02792 | 53.02057 | FALSE | FALSE |

| | | | | | |
|---|---|---|---|---|---|
| LOC107065229 | uncharacterized protein LOC107065229 | 0.02874 | 53.03994 | FALSE | FALSE |
| LOC107066668 | max-like protein X | 0.02683 | 53.05791 | FALSE | FALSE |
| LOC107069131 | S-adenosylmethionine synthase | 0.0267 | 53.05791 | FALSE | FALSE |
| LOC107072057 | uncharacterized protein LOC107072057 | 0.02784 | 53.05862 | FALSE | FALSE |
| LOC107067601 | uncharacterized protein LOC107067601 | 0.02791 | 53.06143 | FALSE | FALSE |
| LOC107070378 | LIM/homeobox protein Lhx5 | 0.02739 | 53.06696 | FALSE | FALSE |
| LOC107072007 | chromatin complexes subunit BAP18 | 0.02731 | 53.07502 | FALSE | FALSE |
| LOC107073607 | uncharacterized protein LOC107073607 | 0.02756 | 53.08263 | FALSE | FALSE |
| LOC107069501 | gustatory receptor 5a for trehalose-like | 0.02747 | 53.08658 | FALSE | FALSE |
| LOC107071894 | lipoma HMGIC fusion partner-like 3 protein | 0.02798 | 53.12184 | FALSE | FALSE |
| LOC107068018 | uncharacterized protein LOC107068018 | 0.02959 | 53.1343 | FALSE | FALSE |
| LOC107069531 | visual system homeobox 2-like | 0.02772 | 53.13606 | FALSE | FALSE |
| LOC107071787 | serine/threonine-protein kinase mos | 0.0277 | 53.14085 | FALSE | FALSE |
| LOC107065157 | 2-hydroxyacyl-CoA lyase 1 | 0.02721 | 53.14434 | FALSE | FALSE |
| LOC107069526 | probable cytochrome P450 6a14 | 0.02818 | 53.14462 | FALSE | FALSE |
| LOC107070308 | uncharacterized protein LOC107070308 | 0.02691 | 53.17042 | FALSE | FALSE |
| LOC107066939 | uncharacterized protein LOC107066939 | 0.0281 | 53.17069 | FALSE | FALSE |
| LOC107066431 | lysozyme c-1-like | 0.02711 | 53.19004 | FALSE | FALSE |
| LOC107073821 | uncharacterized protein LOC107073821 | 0.02678 | 53.22498 | FALSE | FALSE |
| LOC107071819 | protein tramtrack, alpha isoform | 0.02727 | 53.22764 | FALSE | FALSE |
| LOC107071814 | probable ribosome biogenesis protein RLP2 | 0.02714 | 53.23131 | FALSE | FALSE |
| LOC107063866 | ras-related protein Rab-11A | 0.02794 | 53.23266 | FALSE | FALSE |
| LOC107069046 | innexin shaking-B-like | 0.0272 | 53.23691 | FALSE | FALSE |
| LOC107065902 | symplekin | 0.0275 | 53.25219 | FALSE | FALSE |
| LOC107064916 | regulation of nuclear pre-mRNA domain-cor | 0.0291 | 53.26636 | FALSE | FALSE |
| LOC107068770 | signal recognition particle 19 kDa protein | 0.0271 | 53.27236 | FALSE | FALSE |
| LOC107067162 | protein escargot-like | 0.02764 | 53.27506 | FALSE | FALSE |
| LOC107065525 | uncharacterized protein LOC107065525 | 0.02715 | 53.29715 | FALSE | FALSE |
| LOC107071597 | calcineurin B homologous protein 1 | 0.02817 | 53.30703 | FALSE | FALSE |
| LOC107074158 | 39S ribosomal protein L32, mitochondrial | 0.02688 | 53.3142 | FALSE | FALSE |
| LOC107072863 | uncharacterized protein LOC107072863 | 0.02744 | 53.34714 | FALSE | FALSE |
| LOC107064190 | 40S ribosomal protein S7 | 0.02883 | 53.35075 | FALSE | FALSE |
| LOC107072334 | B9 domain-containing protein 2-like | 0.02657 | 53.35544 | FALSE | FALSE |
| LOC107069943 | brain protein I3 | 0.02546 | 53.35663 | FALSE | FALSE |
| LOC107070661 | adenylate kinase 9-like | 0.027 | 53.36938 | FALSE | FALSE |
| LOC107070280 | Fanconi anemia group J protein homolog | 0.02865 | 53.37643 | FALSE | FALSE |
| LOC107066079 | uncharacterized protein LOC107066079 | 0.02773 | 53.37659 | FALSE | FALSE |
| LOC107064134 | tubulin beta-1 chain | 0.02804 | 53.37702 | FALSE | FALSE |
| LOC107066880 | adenylate kinase isoenzyme 1 | 0.027 | 53.40422 | FALSE | FALSE |
| LOC107072482 | actin-binding Rho-activating protein-like | 0.02743 | 53.42784 | FALSE | FALSE |
| LOC107071883 | guanine nucleotide-releasing factor 2 | 0.02852 | 53.44584 | FALSE | FALSE |
| LOC107070616 | structure-specific endonuclease subunit slx1 | 0.02723 | 53.44763 | FALSE | FALSE |
| LOC107066162 | TATA element modulatory factor-like | 0.02656 | 53.4558 | FALSE | FALSE |
| LOC107074420 | claspin-like | 0.02618 | 53.48663 | FALSE | FALSE |
| LOC107065924 | TATA box-binding protein-like protein 1 | 0.02731 | 53.49712 | FALSE | FALSE |
| LOC107069601 | netrin receptor UNC5B-like | 0.02766 | 53.51037 | FALSE | FALSE |
| LOC107074295 | coiled-coil domain-containing protein 137 | 0.02798 | 53.5149 | FALSE | FALSE |
| LOC107068138 | palmitoyltransferase ZDHHC23 | 0.02754 | 53.51511 | FALSE | FALSE |
| LOC107065667 | uncharacterized protein LOC107065667 | 0.02726 | 53.58226 | FALSE | FALSE |
| LOC107068796 | mediator of RNA polymerase II transcriptior | 0.02746 | 53.60344 | FALSE | FALSE |

| | | | | | |
|---|---|---|---|---|---|
| LOC107073167 | alpha-protein kinase 1 | 0.02761 | 53.60553 | FALSE | FALSE |
| LOC107071213 | uncharacterized protein LOC107071213 | 0.02781 | 53.62187 | FALSE | FALSE |
| LOC107073515 | uncharacterized protein LOC107073515 | 0.02724 | 53.62855 | FALSE | FALSE |
| LOC107074561 | uncharacterized protein LOC107074561 | 0.02708 | 53.62888 | FALSE | FALSE |
| LOC107066755 | tyrosine decarboxylase-like | 0.0277 | 53.62971 | FALSE | FALSE |
| LOC107064571 | annulin | 0.02634 | 53.63402 | FALSE | FALSE |
| LOC107070100 | uncharacterized protein LOC107070100 | 0.02936 | 53.64024 | FALSE | FALSE |
| LOC107074531 | differentially expressed in FDCP 6 homolog | 0.02784 | 53.64535 | FALSE | FALSE |
| LOC107065775 | uncharacterized protein LOC107065775 | 0.02712 | 53.66126 | FALSE | FALSE |
| LOC107071864 | putative inorganic phosphate cotransporter | 0.02742 | 53.68587 | FALSE | FALSE |
| LOC107071405 | putative neutral sphingomyelinase | 0.02698 | 53.68817 | FALSE | FALSE |
| LOC107067107 | uncharacterized protein LOC107067107 | 0.02738 | 53.69893 | FALSE | FALSE |
| LOC107073959 | putative sodium-coupled neutral amino acid | 0.02676 | 53.70322 | FALSE | FALSE |
| LOC107067241 | SUMO-conjugating enzyme UBC9-B | 0.02751 | 53.70333 | FALSE | FALSE |
| LOC107063691 | GATA zinc finger domain-containing protein | 0.02612 | 53.7078 | FALSE | FALSE |
| LOC107063615 | titin | 0.02723 | 53.73698 | FALSE | FALSE |
| LOC107069937 | pinopsin-like | 0.02737 | 53.76305 | FALSE | FALSE |
| LOC107068765 | protein LAP2-like | 0.02789 | 53.7706 | FALSE | FALSE |
| LOC107069810 | uncharacterized protein LOC107069810 | 0.02753 | 53.77282 | FALSE | FALSE |
| LOC107074418 | WD repeat-containing protein 13-like | 0.02728 | 53.77334 | FALSE | FALSE |
| LOC107064162 | vacuolar protein sorting-associated protein | 0.02631 | 53.79569 | FALSE | FALSE |
| LOC107070231 | sperm-associated antigen 7 homolog | 0.0269 | 53.79628 | FALSE | FALSE |
| LOC107071168 | coiled-coil domain-containing protein 112-li | 0.02748 | 53.80718 | FALSE | FALSE |
| LOC107074391 | nucleoredoxin-like | 0.02894 | 53.9041 | FALSE | FALSE |
| LOC107070841 | 3-phosphoinositide-dependent protein kina | 0.02814 | 53.91618 | FALSE | FALSE |
| LOC107070581 | GATA zinc finger domain-containing protein | 0.03046 | 53.93418 | FALSE | FALSE |
| LOC107069076 | serine-rich adhesin for platelets | 0.028 | 53.94106 | FALSE | FALSE |
| LOC107068638 | uncharacterized protein LOC107068638 | 0.02743 | 53.96104 | FALSE | FALSE |
| LOC107073018 | uncharacterized protein LOC107073018 | 0.0284 | 53.96498 | FALSE | FALSE |
| LOC107067543 | myosin light chain alkali | 0.02646 | 53.96824 | FALSE | FALSE |
| LOC107073432 | uncharacterized protein LOC107073432 | 0.02699 | 54.00984 | FALSE | FALSE |
| LOC107072216 | uncharacterized protein LOC107072216 | 0.02833 | 54.01174 | FALSE | FALSE |
| LOC107065717 | transcription factor AP-1-like | 0.02622 | 54.01462 | FALSE | FALSE |
| LOC107066768 | uncharacterized protein LOC107066768 | 0.02749 | 54.049 | FALSE | FALSE |
| LOC107068512 | protein UBASH3A homolog | 0.02761 | 54.05066 | FALSE | FALSE |
| LOC107063915 | (11Z)-hexadec-11-enoyl-CoA conjugase-like | 0.02707 | 54.05631 | FALSE | FALSE |
| LOC107070652 | DNA damage-binding protein 1 | 0.02842 | 54.09007 | FALSE | FALSE |
| LOC107069361 | ras-related protein Rab-3 | 0.02656 | 54.12433 | FALSE | FALSE |
| LOC107072201 | myosin-2 essential light chain | 0.02738 | 54.15402 | FALSE | FALSE |
| LOC107065959 | titin | 0.02693 | 54.15562 | FALSE | FALSE |
| LOC107069311 | uncharacterized protein LOC107069311 | 0.02669 | 54.16086 | FALSE | FALSE |
| LOC107066561 | transport and Golgi organization protein 11 | 0.02761 | 54.17895 | FALSE | FALSE |
| LOC107064385 | uncharacterized protein LOC107064385 | 0.02663 | 54.18969 | FALSE | FALSE |
| LOC107073733 | transcription factor Sp8-like | 0.02708 | 54.19809 | FALSE | FALSE |
| LOC107073190 | TBC1 domain family member 25 | 0.02675 | 54.21468 | FALSE | FALSE |
| LOC107066120 | uncharacterized protein LOC107066120 | 0.02656 | 54.22604 | FALSE | FALSE |
| LOC107073877 | traB domain-containing protein | 0.02792 | 54.2624 | FALSE | FALSE |
| LOC107073186 | WD repeat-containing protein 7 | 0.02761 | 54.27263 | FALSE | FALSE |
| LOC107066625 | fructose-1,6-bisphosphatase 1 | 0.02844 | 54.28176 | FALSE | FALSE |
| LOC107071432 | cytochrome P450 4c3-like | 0.02672 | 54.28867 | FALSE | FALSE |

| | | | | | |
|---|---|---|---|---|---|
| LOC107067475 | maternal protein tudor | 0.02628 | 54.29384 | FALSE | FALSE |
| LOC107070267 | sugar transporter SWEET1 | 0.02764 | 54.29489 | FALSE | FALSE |
| LOC107069612 | uncharacterized protein LOC107069612 | 0.02835 | 54.30287 | FALSE | FALSE |
| LOC107068040 | polycomb group RING finger protein 3 | 0.02778 | 54.30844 | FALSE | FALSE |
| LOC107068964 | uncharacterized protein LOC107068964 | 0.02716 | 54.31131 | FALSE | FALSE |
| LOC107065300 | MAGUK p55 subfamily member 7 | 0.02718 | 54.34741 | FALSE | FALSE |
| LOC107066081 | translocon-associated protein subunit gamn | 0.02762 | 54.3482 | FALSE | FALSE |
| LOC107069892 | R3H domain-containing protein 1 | 0.02731 | 54.35151 | FALSE | FALSE |
| LOC107074022 | ribosome biogenesis protein TSR3 homolog | 0.02777 | 54.35247 | FALSE | FALSE |
| LOC107072445 | probable phospholipid-transporting ATPase | 0.02736 | 54.35249 | FALSE | FALSE |
| LOC107072289 | probable oligoribonuclease | 0.0275 | 54.38034 | FALSE | FALSE |
| LOC107069533 | uncharacterized protein LOC107069533 | 0.02672 | 54.39111 | FALSE | FALSE |
| LOC107063978 | hybrid signal transduction histidine kinase L | 0.02662 | 54.40701 | FALSE | FALSE |
| LOC107074076 | dynein light chain 1, axonemal-like | 0.02594 | 54.41588 | FALSE | FALSE |
| LOC107071141 | UDP-glucose 6-dehydrogenase | 0.02714 | 54.41949 | FALSE | FALSE |
| LOC107070484 | DNA polymerase delta catalytic subunit | 0.02744 | 54.42088 | FALSE | FALSE |
| LOC107067735 | slit homolog 3 protein-like | 0.02749 | 54.42878 | FALSE | FALSE |
| LOC107068848 | uncharacterized protein LOC107068848 | 0.0276 | 54.43502 | FALSE | FALSE |
| LOC107071706 | uncharacterized protein LOC107071706 | 0.02784 | 54.44441 | FALSE | FALSE |
| LOC107073866 | uncharacterized protein KIAA1109 | 0.02572 | 54.45429 | FALSE | FALSE |
| LOC107067113 | insulin-like growth factor III | 0.02744 | 54.45733 | FALSE | FALSE |
| LOC107072056 | ataxin-2 homolog | 0.02744 | 54.46338 | FALSE | FALSE |
| LOC107068506 | uncharacterized protein LOC107068506 | 0.02761 | 54.46395 | FALSE | FALSE |
| LOC107067362 | coiled-coil-helix-coiled-coil-helix domain-co | 0.02648 | 54.47624 | FALSE | FALSE |
| LOC107073543 | putative ribosome-binding factor A, mitocho | 0.02699 | 54.48257 | FALSE | FALSE |
| LOC107070530 | guanine nucleotide-binding protein subunit | 0.02872 | 54.48645 | FALSE | FALSE |
| LOC107063982 | multiple coagulation factor deficiency prote | 0.02729 | 54.48705 | FALSE | FALSE |
| LOC107068088 | NADH-ubiquinone oxidoreductase subunit 8 | 0.0268 | 54.50298 | FALSE | FALSE |
| LOC107069092 | mediator of RNA polymerase II transcription | 0.02766 | 54.50694 | FALSE | FALSE |
| LOC107065777 | nuclear RNA export factor 1-like | 0.02831 | 54.51895 | FALSE | FALSE |
| LOC107071121 | uncharacterized protein LOC107071121 | 0.02767 | 54.52037 | FALSE | FALSE |
| LOC107069598 | uncharacterized protein LOC107069598 | 0.02826 | 54.52125 | FALSE | FALSE |
| LOC107068474 | anaphase-promoting complex subunit 2 | 0.02703 | 54.55147 | FALSE | FALSE |
| LOC107067752 | UPF0389 protein CG9231 | 0.02769 | 54.56081 | FALSE | FALSE |
| LOC107072083 | beta-1,4-galactosyltransferase 7 | 0.02707 | 54.56085 | FALSE | FALSE |
| LOC107064384 | larval cuticle protein A3A-like | 0.02718 | 54.56754 | FALSE | FALSE |
| LOC107064214 | uncharacterized protein LOC107064214 | 0.02621 | 54.57518 | FALSE | FALSE |
| LOC107065908 | syntaxin-6 | 0.02759 | 54.57755 | FALSE | FALSE |
| LOC107068331 | myosin heavy chain, non-muscle | 0.02686 | 54.58174 | FALSE | FALSE |
| LOC107066973 | uncharacterized protein LOC107066973 | 0.02764 | 54.58714 | FALSE | FALSE |
| LOC107067312 | nucleolar complex protein 2 homolog | 0.02678 | 54.59015 | FALSE | FALSE |
| LOC107074556 | transmembrane emp24 domain-containing | 0.02778 | 54.60242 | FALSE | FALSE |
| LOC107067701 | homeotic protein deformed | 0.02766 | 54.61956 | FALSE | FALSE |
| LOC107071201 | uncharacterized protein LOC107071201 | 0.02737 | 54.62331 | FALSE | FALSE |
| LOC107067610 | coiled-coil domain-containing protein 96-lik | 0.027 | 54.63701 | FALSE | FALSE |
| LOC107067657 | SRSF protein kinase 3 | 0.02801 | 54.63959 | FALSE | FALSE |
| LOC107067013 | protein aveugle | 0.0272 | 54.63992 | FALSE | FALSE |
| LOC107065266 | protein SMG7-like | 0.02726 | 54.65154 | FALSE | FALSE |
| LOC107068065 | carotenoid isomerooxygenase | 0.02732 | 54.68215 | FALSE | FALSE |
| LOC107063854 | retinol dehydrogenase 11-like | 0.02774 | 54.68354 | FALSE | FALSE |

| | | | | | |
|---|---|---|---|---|---|
| LOC107071050 | elongation of very long chain fatty acids pro | 0.02742 | 54.69246 | FALSE | FALSE |
| LOC107072893 | uncharacterized protein LOC107072893 | 0.02582 | 54.69497 | FALSE | FALSE |
| LOC107069323 | probable G-protein coupled receptor No9 | 0.02679 | 54.69654 | FALSE | FALSE |
| LOC107069397 | serine/arginine repetitive matrix protein 2-l | 0.02629 | 54.71167 | FALSE | FALSE |
| LOC107070893 | conserved oligomeric Golgi complex subunit | 0.02687 | 54.71644 | FALSE | FALSE |
| LOC107071766 | FIT family protein CG10671 | 0.02714 | 54.72512 | FALSE | FALSE |
| LOC107064535 | 26S proteasome non-ATPase regulatory sub | 0.02663 | 54.73427 | FALSE | FALSE |
| LOC107070485 | trypsin-1-like | 0.02744 | 54.74916 | FALSE | FALSE |
| LOC107064439 | uncharacterized protein LOC107064439 | 0.02683 | 54.75869 | FALSE | FALSE |
| LOC107069654 | probable ATP-dependent RNA helicase DDX | 0.02828 | 54.77255 | FALSE | FALSE |
| LOC107067468 | interleukin enhancer-binding factor 2 | 0.02711 | 54.77682 | FALSE | FALSE |
| LOC107067082 | receptor-type guanylate cyclase gcy-4-like | 0.02656 | 54.77936 | FALSE | FALSE |
| LOC107071486 | putative fatty acyl-CoA reductase CG5065 | 0.02834 | 54.78179 | FALSE | FALSE |
| LOC107068467 | beta-lactamase-like protein 2 | 0.02671 | 54.78213 | FALSE | FALSE |
| LOC107074700 | tumor suppressor candidate 3 | 0.02806 | 54.7838 | FALSE | FALSE |
| LOC107072966 | uncharacterized protein LOC107072966 | 0.02607 | 54.78549 | FALSE | FALSE |
| LOC107065977 | putative uncharacterized protein DDB_G029 | 0.02702 | 54.80196 | FALSE | FALSE |
| LOC107067561 | ubiquitin domain-containing protein UBFD1 | 0.02711 | 54.8056 | FALSE | FALSE |
| LOC107064582 | uncharacterized protein LOC107064582 | 0.02656 | 54.82056 | FALSE | FALSE |
| LOC107066804 | protein held out wings | 0.02592 | 54.82247 | FALSE | FALSE |
| LOC107072314 | nucleolysin TIAR | 0.02694 | 54.82306 | FALSE | FALSE |
| LOC107064119 | ubiquitin-conjugating enzyme E2 C | 0.02696 | 54.8344 | FALSE | FALSE |
| LOC107069508 | protein NRDE2 homolog | 0.02399 | 54.84386 | FALSE | FALSE |
| LOC107069812 | uncharacterized protein LOC107069812 | 0.02693 | 54.84526 | FALSE | FALSE |
| LOC107068535 | vacuolar protein sorting-associated protein | 0.02509 | 54.84706 | FALSE | FALSE |
| LOC107063708 | uncharacterized protein LOC107063708 | 0.02773 | 54.85257 | FALSE | FALSE |
| LOC107064210 | torsin-1A-like | 0.02845 | 54.86094 | FALSE | FALSE |
| LOC107065757 | golgin subfamily A member 1-like | 0.02636 | 54.87219 | FALSE | FALSE |
| LOC107067123 | probable malonyl-CoA-acyl carrier protein tr | 0.02793 | 54.88257 | FALSE | FALSE |
| LOC107063942 | USP6 N-terminal-like protein | 0.02594 | 54.8958 | FALSE | FALSE |
| LOC107066282 | carboxypeptidase B-like | 0.02689 | 54.90444 | FALSE | FALSE |
| LOC107067491 | methionyl-tRNA formyltransferase, mitocho | 0.0266 | 54.91135 | FALSE | FALSE |
| LOC107069749 | polyamine-modulated factor 1-binding prot | 0.02663 | 54.93094 | FALSE | FALSE |
| LOC107066378 | gastrula zinc finger protein XlCGF26.1-like | 0.02712 | 54.94794 | FALSE | FALSE |
| LOC107072565 | histone-lysine N-methyltransferase, H3 lysir | 0.02608 | 54.94823 | FALSE | FALSE |
| LOC107070696 | lipase 3-like | 0.02778 | 54.96023 | FALSE | FALSE |
| LOC107064730 | uncharacterized protein LOC107064730 | 0.02744 | 54.96146 | FALSE | FALSE |
| LOC107069478 | low molecular weight phosphotyrosine prot | 0.02681 | 54.96268 | FALSE | FALSE |
| LOC107065099 | nuclear factor related to kappa-B-binding pr | 0.02921 | 54.96453 | FALSE | FALSE |
| LOC107066188 | BRCA1-associated protein | 0.0257 | 54.96897 | FALSE | FALSE |
| LOC107074331 | cyclin-dependent kinase 20-like | 0.02651 | 54.97393 | FALSE | FALSE |
| LOC107066111 | synaptic vesicle glycoprotein 2B | 0.0259 | 54.97899 | FALSE | FALSE |
| LOC107068753 | uncharacterized protein LOC107068753 | 0.02699 | 54.98027 | FALSE | FALSE |
| LOC107064886 | ubiquitin-conjugating enzyme E2 T-like | 0.02731 | 55.01606 | FALSE | FALSE |
| LOC107064695 | solute carrier family 35 member F2-like | 0.02736 | 55.02477 | FALSE | FALSE |
| LOC107071293 | GTP-binding protein RAD-like | 0.02723 | 55.03305 | FALSE | FALSE |
| LOC107068521 | alpha-ketoglutarate-dependent dioxygenase | 0.02642 | 55.0664 | FALSE | FALSE |
| LOC107067032 | ankyrin repeat and MYND domain-containir | 0.02732 | 55.07464 | FALSE | FALSE |
| LOC107066811 | uncharacterized protein LOC107066811 | 0.02776 | 55.07592 | FALSE | FALSE |
| LOC107071001 | peroxidasin | 0.02699 | 55.08845 | FALSE | FALSE |

| | | | | | |
|---|---|---|---|---|---|
| LOC107065737 | uncharacterized protein LOC107065737 | 0.02677 | 55.0936 | FALSE | FALSE |
| LOC107073585 | nose resistant to fluoxetine protein 6-like | 0.02638 | 55.11864 | FALSE | FALSE |
| LOC107069775 | DNA-directed RNA polymerase I subunit RP/ | 0.0265 | 55.11973 | FALSE | FALSE |
| LOC107070853 | J domain-containing protein | 0.02677 | 55.12064 | FALSE | FALSE |
| LOC107068239 | HEAT repeat-containing protein 5B | 0.02601 | 55.12205 | FALSE | FALSE |
| LOC107063837 | uncharacterized protein LOC107063837 | 0.02769 | 55.12567 | FALSE | FALSE |
| LOC107071177 | NADPH:adrenodoxin oxidoreductase, mitoc | 0.02646 | 55.12834 | FALSE | FALSE |
| LOC107069871 | odorant receptor 49b-like | 0.02763 | 55.13305 | FALSE | FALSE |
| LOC107068592 | probable serine/threonine-protein kinase ir | 0.02613 | 55.17854 | FALSE | FALSE |
| LOC107072686 | vesicle transport protein SEC20 | 0.02748 | 55.18232 | FALSE | FALSE |
| LOC107071220 | ubiquitin-conjugating enzyme E2-17 kDa | 0.02867 | 55.18457 | FALSE | FALSE |
| LOC107067980 | odorant receptor 2a-like | 0.02707 | 55.19125 | FALSE | FALSE |
| LOC107073082 | Kv channel-interacting protein 1-like | 0.02665 | 55.21074 | FALSE | FALSE |
| LOC107064293 | protein phosphatase 1 regulatory subunit 4: | 0.0275 | 55.22654 | FALSE | FALSE |
| LOC107073906 | uncharacterized protein LOC107073906 | 0.0262 | 55.25049 | FALSE | FALSE |
| LOC107069680 | uncharacterized protein LOC107069680 | 0.02713 | 55.26864 | FALSE | FALSE |
| LOC107072978 | uncharacterized protein LOC107072978 | 0.02645 | 55.28054 | FALSE | FALSE |
| LOC107063913 | glutamyl aminopeptidase-like | 0.02788 | 55.29491 | FALSE | FALSE |
| LOC107064358 | minor histocompatibility antigen H13 | 0.02704 | 55.3002 | FALSE | FALSE |
| LOC107072752 | uncharacterized protein LOC107072752 | 0.02741 | 55.30217 | FALSE | FALSE |
| LOC107067731 | synaptic vesicle glycoprotein 2B-like | 0.02655 | 55.32978 | FALSE | FALSE |
| LOC107069363 | uncharacterized protein LOC107069363 | 0.02703 | 55.34466 | FALSE | FALSE |
| LOC107063802 | uncharacterized protein LOC107063802 | 0.02709 | 55.35779 | FALSE | FALSE |
| LOC107063863 | probable salivary secreted peptide | 0.02567 | 55.3908 | FALSE | FALSE |
| LOC107070929 | uncharacterized protein LOC107070929 | 0.02642 | 55.3949 | FALSE | FALSE |
| LOC107069422 | adenylate cyclase type 9 | 0.027 | 55.39515 | FALSE | FALSE |
| LOC107067255 | general transcription factor 3C polypeptide | 0.02606 | 55.39531 | FALSE | FALSE |
| LOC107068574 | protein SYS1 homolog | 0.02588 | 55.39596 | FALSE | FALSE |
| LOC107066040 | F-box/WD repeat-containing protein 4-like | 0.02708 | 55.42228 | FALSE | FALSE |
| LOC107074014 | 40S ribosomal protein S6-like | 0.02755 | 55.42372 | FALSE | FALSE |
| LOC107074268 | glutamate receptor ionotropic, kainate 2-lik | 0.02725 | 55.42615 | FALSE | FALSE |
| LOC107067976 | multifunctional methyltransferase subunit T | 0.02711 | 55.42666 | FALSE | FALSE |
| LOC107074563 | uncharacterized protein LOC107074563 | 0.02778 | 55.45241 | FALSE | FALSE |
| LOC107069997 | oligosaccharyltransferase complex subunit ( | 0.028 | 55.45303 | FALSE | FALSE |
| LOC107071399 | PDZ domain-containing protein 8 | 0.02574 | 55.45974 | FALSE | FALSE |
| LOC107064807 | tyrosine-protein kinase CSK-like | 0.02474 | 55.47821 | FALSE | FALSE |
| LOC107071959 | nephrin | 0.02851 | 55.47978 | FALSE | FALSE |
| LOC107073126 | F-box/SPRY domain-containing protein 1 | 0.02689 | 55.48528 | FALSE | FALSE |
| LOC107070045 | platelet binding protein GspB-like | 0.02746 | 55.4875 | FALSE | FALSE |
| LOC107073662 | fatty acid synthase-like | 0.02757 | 55.49194 | FALSE | FALSE |
| LOC107071606 | DNA replication licensing factor Mcm3 | 0.0268 | 55.49364 | FALSE | FALSE |
| LOC107073110 | cytoplasmic dynein 2 light intermediate cha | 0.02648 | 55.51577 | FALSE | FALSE |
| LOC107072297 | mothers against decapentaplegic homolog 4 | 0.02706 | 55.52252 | FALSE | FALSE |
| LOC107069262 | uncharacterized aarF domain-containing pro | 0.02555 | 55.53162 | FALSE | FALSE |
| LOC107063794 | vascular endothelial growth factor A-A-like | 0.02829 | 55.542 | FALSE | FALSE |
| LOC107064008 | ubiquinone biosynthesis protein COQ4 hom | 0.02747 | 55.55829 | FALSE | FALSE |
| LOC107069701 | integrator complex subunit 4 | 0.02743 | 55.55901 | FALSE | FALSE |
| LOC107069918 | cytochrome P450 307a1-like | 0.02813 | 55.57513 | FALSE | FALSE |
| LOC107065050 | transient receptor potential-gamma protein | 0.02562 | 55.58488 | FALSE | FALSE |
| LOC107073837 | pre-mRNA-splicing factor ATP-dependent RI | 0.02683 | 55.58921 | FALSE | FALSE |

| | | | | | |
|---|---|---|---|---|---|
| LOC107071429 | calcium uptake protein 3, mitochondrial | 0.02626 | 55.5984 | FALSE | FALSE |
| LOC107069439 | alsin | 0.02586 | 55.60523 | FALSE | FALSE |
| LOC107063928 | G-protein coupled receptor Mth2-like | 0.02813 | 55.6135 | FALSE | FALSE |
| LOC107064103 | uncharacterized protein LOC107064103 | 0.02549 | 55.63486 | FALSE | FALSE |
| LOC107068245 | glutaredoxin-related protein 5, mitochondri | 0.02615 | 55.6672 | FALSE | FALSE |
| LOC107068747 | glutathione S-transferase 1-1-like | 0.02741 | 55.68507 | FALSE | FALSE |
| LOC107064587 | centrosomal protein of 104 kDa | 0.02627 | 55.68525 | FALSE | FALSE |
| LOC107071513 | probable nucleoporin Nup54 | 0.0271 | 55.68881 | FALSE | FALSE |
| LOC107070388 | homologous-pairing protein 2 homolog | 0.02627 | 55.70083 | FALSE | FALSE |
| LOC107063687 | protein mahjong | 0.02556 | 55.70468 | FALSE | FALSE |
| LOC107073908 | phosphoglucomutase-2 | 0.02726 | 55.70914 | FALSE | FALSE |
| LOC107072593 | transmembrane protein C9orf91 homolog | 0.02578 | 55.72121 | FALSE | FALSE |
| LOC107072479 | uncharacterized protein LOC107072479 | 0.02702 | 55.73416 | FALSE | FALSE |
| LOC107065632 | protein bicaudal C | 0.0251 | 55.74852 | FALSE | FALSE |
| LOC107065437 | chymotrypsin inhibitor | 0.02642 | 55.7488 | FALSE | FALSE |
| LOC107070416 | uncharacterized protein LOC107070416 | 0.02608 | 55.76056 | FALSE | FALSE |
| LOC107071569 | uncharacterized protein LOC107071569 | 0.02679 | 55.77314 | FALSE | FALSE |
| LOC107070495 | splicing factor U2af 38 kDa subunit-like | 0.02521 | 55.77393 | FALSE | FALSE |
| LOC107071386 | uncharacterized protein LOC107071386 | 0.02667 | 55.78413 | FALSE | FALSE |
| LOC107065720 | uncharacterized protein LOC107065720 | 0.0261 | 55.78855 | FALSE | FALSE |
| LOC107072763 | uncharacterized protein LOC107072763 | 0.02537 | 55.81993 | FALSE | FALSE |
| LOC107067987 | 4-hydroxybutyrate coenzyme A transferase- | 0.02711 | 55.82584 | FALSE | FALSE |
| LOC107068701 | endocuticle structural glycoprotein ABD-4-li | 0.02691 | 55.82751 | FALSE | FALSE |
| LOC107070891 | protein scarlet-like | 0.02596 | 55.84305 | FALSE | FALSE |
| LOC107068407 | very long-chain specific acyl-CoA dehydroge | 0.02807 | 55.84923 | FALSE | FALSE |
| LOC107064235 | probable actin-related protein 2/3 complex | 0.02644 | 55.84925 | FALSE | FALSE |
| LOC107070710 | DNA polymerase epsilon subunit 3 | 0.02699 | 55.84958 | FALSE | FALSE |
| LOC107066010 | uncharacterized protein LOC107066010 | 0.02746 | 55.89026 | FALSE | FALSE |
| LOC107072772 | mediator of RNA polymerase II transcriptior | 0.02576 | 55.91773 | FALSE | FALSE |
| LOC107065184 | hepatic leukemia factor | 0.02612 | 55.94059 | FALSE | FALSE |
| LOC107066849 | protein TRC8 homolog | 0.02761 | 55.94575 | FALSE | FALSE |
| LOC107065594 | exosome component 10 | 0.02515 | 55.95179 | FALSE | FALSE |
| LOC107072413 | uncharacterized protein LOC107072413 | 0.02507 | 55.95903 | FALSE | FALSE |
| LOC107073637 | uncharacterized protein LOC107073637 | 0.02639 | 55.9653 | FALSE | FALSE |
| LOC107071212 | nudC domain-containing protein 3 | 0.02885 | 55.96664 | FALSE | FALSE |
| LOC107063707 | glutathione S-transferase-like | 0.02695 | 55.98639 | FALSE | FALSE |
| LOC107072181 | uncharacterized protein LOC107072181 | 0.02697 | 55.99759 | FALSE | FALSE |
| LOC107074191 | neural cell adhesion molecule 2-like | 0.02671 | 55.999 | FALSE | FALSE |
| LOC107064450 | suppressor of Mek1-like | 0.02547 | 56.00915 | FALSE | FALSE |
| LOC107072859 | nuclear RNA export factor 1-like | 0.02633 | 56.01836 | FALSE | FALSE |
| LOC107068424 | 60S ribosomal protein L23a-like | 0.0267 | 56.04819 | FALSE | FALSE |
| LOC107071752 | 60S ribosomal protein L11 | 0.02622 | 56.07325 | FALSE | FALSE |
| LOC107070694 | muscarinic acetylcholine receptor M5 | 0.02697 | 56.07531 | FALSE | FALSE |
| LOC107073063 | adrenodoxin | 0.02791 | 56.08365 | FALSE | FALSE |
| LOC107070883 | uncharacterized protein LOC107070883 | 0.02535 | 56.08721 | FALSE | FALSE |
| LOC107064349 | uncharacterized protein LOC107064349 | 0.02785 | 56.08856 | FALSE | FALSE |
| LOC107073491 | la-related protein 1-like | 0.02551 | 56.09631 | FALSE | FALSE |
| LOC107070205 | transmembrane protein 50A | 0.0261 | 56.09722 | FALSE | FALSE |
| LOC107073099 | BTB/POZ domain-containing protein 6-A | 0.02763 | 56.14398 | FALSE | FALSE |
| LOC107067347 | putative homeodomain transcription factor | 0.02708 | 56.148 | FALSE | FALSE |

| | | | | | |
|---|---|---|---|---|---|
| LOC107064897 | laccase-5 | 0.02787 | 56.15065 | FALSE | FALSE |
| LOC107072698 | ras-related GTP-binding protein A | 0.02545 | 56.17332 | FALSE | FALSE |
| LOC107071800 | transmembrane protein 145-like | 0.02709 | 56.19116 | FALSE | FALSE |
| LOC107069633 | major facilitator superfamily domain-contai | 0.0252 | 56.23153 | FALSE | FALSE |
| LOC107064049 | uncharacterized protein LOC107064049 | 0.02605 | 56.24217 | FALSE | FALSE |
| LOC107072788 | protein ELYS | 0.02588 | 56.26394 | FALSE | FALSE |
| LOC107072187 | uncharacterized protein DDB_G0287625-lik | 0.02783 | 56.29176 | FALSE | FALSE |
| LOC107064169 | hairy/enhancer-of-split related with YRPW r | 0.02515 | 56.29223 | FALSE | FALSE |
| LOC107063873 | NADH dehydrogenase [ubiquinone | 0.027 | 56.30782 | FALSE | FALSE |
| LOC107067833 | calcium-binding mitochondrial carrier prote | 0.0257 | 56.30877 | FALSE | FALSE |
| LOC107066900 | peptidoglycan-recognition protein 2-like | 0.02642 | 56.3097 | FALSE | FALSE |
| LOC107071148 | uncharacterized protein LOC107071148 | 0.02589 | 56.31777 | FALSE | FALSE |
| LOC107073420 | uncharacterized protein LOC107073420 | 0.0267 | 56.32148 | FALSE | FALSE |
| LOC107071731 | protein Wnt-7b | 0.0268 | 56.33149 | FALSE | FALSE |
| LOC107072147 | malignant T-cell-amplified sequence 1 hom | 0.02704 | 56.34063 | FALSE | FALSE |
| LOC107068313 | solute carrier family 35 member E1 homolo | 0.02629 | 56.35328 | FALSE | FALSE |
| LOC107066507 | AP-1 complex subunit sigma-2 | 0.02659 | 56.36644 | FALSE | FALSE |
| LOC107071241 | luc7-like protein 3 | 0.02553 | 56.36823 | FALSE | FALSE |
| LOC107073347 | uncharacterized protein LOC107073347 | 0.02691 | 56.38267 | FALSE | FALSE |
| LOC107068554 | uncharacterized protein LOC107068554 | 0.02698 | 56.38355 | FALSE | FALSE |
| LOC107067470 | pupal cuticle protein 20-like | 0.02624 | 56.39399 | FALSE | FALSE |
| LOC107070163 | methyl farnesoate epoxidase-like | 0.02753 | 56.39508 | FALSE | FALSE |
| LOC107072276 | AP-2 complex subunit mu | 0.02642 | 56.40239 | FALSE | FALSE |
| LOC107064859 | heparan sulfate glucosamine 3-O-sulfotrans | 0.02671 | 56.40495 | FALSE | FALSE |
| LOC107072911 | uncharacterized protein LOC107072911 | 0.02695 | 56.42108 | FALSE | FALSE |
| LOC107072067 | phosphatidylinositol N-acetylglucosaminyltr | 0.02704 | 56.42605 | FALSE | FALSE |
| LOC107074165 | dolichyl-diphosphooligosaccharide--protein | 0.02718 | 56.43769 | FALSE | FALSE |
| LOC107069916 | UPF0183 protein CG7083 | 0.02744 | 56.43842 | FALSE | FALSE |
| LOC107074171 | uncharacterized protein LOC107074171 | 0.02807 | 56.44283 | FALSE | FALSE |
| LOC107068459 | flotillin-2 | 0.02643 | 56.44327 | FALSE | FALSE |
| LOC107071447 | G-box-binding factor-like | 0.02629 | 56.45142 | FALSE | FALSE |
| LOC107072948 | centrosomal protein of 97 kDa | 0.02525 | 56.46604 | FALSE | FALSE |
| LOC107066525 | uncharacterized protein LOC107066525 | 0.02656 | 56.47819 | FALSE | FALSE |
| LOC107073348 | uncharacterized protein LOC107073348 | 0.03221 | 56.50086 | FALSE | FALSE |
| LOC107066248 | 39S ribosomal protein L23, mitochondrial | 0.02555 | 56.51772 | FALSE | FALSE |
| LOC107063774 | sorting nexin-25-like | 0.02641 | 56.51915 | FALSE | FALSE |
| LOC107070250 | protein bunched, class 2/F/G isoform-like | 0.02659 | 56.51974 | FALSE | FALSE |
| LOC107064889 | HIV Tat-specific factor 1 homolog | 0.02783 | 56.52726 | FALSE | FALSE |
| LOC107065674 | DNA primase large subunit-like | 0.0272 | 56.53037 | FALSE | FALSE |
| LOC107072073 | twinkle protein, mitochondrial | 0.02612 | 56.53039 | FALSE | FALSE |
| LOC107068606 | tropomyosin-1 | 0.02639 | 56.56031 | FALSE | FALSE |
| LOC107071520 | protein PAT1 homolog 1-like | 0.02627 | 56.56575 | FALSE | FALSE |
| LOC107071169 | uncharacterized protein LOC107071169 | 0.02706 | 56.57727 | FALSE | FALSE |
| LOC107069094 | WD repeat-containing protein 75 | 0.02701 | 56.58007 | FALSE | FALSE |
| LOC107065846 | N-glycosylase/DNA lyase | 0.0263 | 56.58234 | FALSE | FALSE |
| LOC107068379 | VWFA and cache domain-containing protei | 0.02633 | 56.62334 | FALSE | FALSE |
| LOC107066383 | uncharacterized protein LOC107066383 | 0.02507 | 56.62746 | FALSE | FALSE |
| LOC107064238 | NADH dehydrogenase [ubiquinone | 0.026 | 56.64687 | FALSE | FALSE |
| LOC107073206 | uncharacterized protein LOC107073206 | 0.0251 | 56.649 | FALSE | FALSE |
| LOC107068861 | 116 kDa U5 small nuclear ribonucleoprotein | 0.0268 | 56.67662 | FALSE | FALSE |

| | | | | | |
|---|---|---|---|---|---|
| LOC107072947 | uncharacterized protein LOC107072947 | 0.02665 | 56.69583 | FALSE | FALSE |
| LOC107063930 | tubulin-specific chaperone E | 0.02606 | 56.70456 | FALSE | FALSE |
| LOC107073012 | allatostatin-A receptor | 0.02488 | 56.70674 | FALSE | FALSE |
| LOC107067180 | uncharacterized protein LOC107067180 | 0.02598 | 56.71981 | FALSE | FALSE |
| LOC107070757 | endonuclease III-like protein 1 | 0.02702 | 56.72383 | FALSE | FALSE |
| LOC107065268 | uncharacterized protein LOC107065268 | 0.0268 | 56.74811 | FALSE | FALSE |
| LOC107067634 | uncharacterized protein LOC107067634 | 0.02555 | 56.75554 | FALSE | FALSE |
| LOC107073032 | very-long-chain (3R)-3-hydroxyacyl-CoA deh | 0.02668 | 56.75577 | FALSE | FALSE |
| LOC107072797 | patched domain-containing protein 3-like | 0.02584 | 56.75757 | FALSE | FALSE |
| LOC107073013 | uncharacterized protein LOC107073013 | 0.02662 | 56.77729 | FALSE | FALSE |
| LOC107072154 | uncharacterized protein LOC107072154 | 0.02505 | 56.78968 | FALSE | FALSE |
| LOC107072987 | attractin-like protein 1 | 0.02591 | 56.79162 | FALSE | FALSE |
| LOC107065015 | DNA-directed RNA polymerase II 16 kDa pol | 0.0263 | 56.83644 | FALSE | FALSE |
| LOC107070332 | protein BCCIP homolog | 0.02745 | 56.83776 | FALSE | FALSE |
| LOC107068997 | 1-acyl-sn-glycerol-3-phosphate acyltransfera | 0.02544 | 56.85938 | FALSE | FALSE |
| LOC107065252 | mitochondrial import inner membrane trans | 0.02594 | 56.87758 | FALSE | FALSE |
| LOC107069520 | protein dispatched | 0.02588 | 56.91359 | FALSE | FALSE |
| LOC107072474 | sodium/hydrogen exchanger 7 | 0.02678 | 56.9214 | FALSE | FALSE |
| LOC107072098 | E3 ubiquitin-protein ligase RING1 | 0.02641 | 56.92174 | FALSE | FALSE |
| LOC107066887 | chitinase-like protein Idgf4 | 0.02588 | 56.93991 | FALSE | FALSE |
| LOC107068525 | transcriptional regulator ATRX-like | 0.02509 | 56.94289 | FALSE | FALSE |
| LOC107066667 | uncharacterized protein LOC107066667 | 0.02636 | 56.94586 | FALSE | FALSE |
| LOC107072770 | uncharacterized protein LOC107072770 | 0.02656 | 56.94729 | FALSE | FALSE |
| LOC107072165 | heparan-alpha-glucosaminide N-acetyltrans | 0.02628 | 56.94782 | FALSE | FALSE |
| Trnaa-cgc | Trnaa-cgc | 0.02757 | 56.96522 | FALSE | FALSE |
| LOC107069016 | uncharacterized protein LOC107069016 | 0.02628 | 56.98663 | FALSE | FALSE |
| LOC107071806 | negative elongation factor E | 0.02693 | 57.00597 | FALSE | FALSE |
| LOC107063900 | nephrin | 0.0267 | 57.01022 | FALSE | FALSE |
| LOC107074274 | uncharacterized protein LOC107074274 | 0.02644 | 57.02042 | FALSE | FALSE |
| LOC107066187 | uncharacterized protein LOC107066187 | 0.02845 | 57.02322 | FALSE | FALSE |
| LOC107065103 | calcium-transporting ATPase sarcoplasmic/e | 0.02574 | 57.02634 | FALSE | FALSE |
| LOC107074474 | lymphocyte-specific helicase-like | 0.02921 | 57.03981 | FALSE | FALSE |
| LOC107063736 | tropomyosin-2-like | 0.0262 | 57.04151 | FALSE | FALSE |
| LOC107070537 | RNA pseudouridylate synthase domain-cont | 0.02502 | 57.05062 | FALSE | FALSE |
| LOC107069674 | eukaryotic initiation factor 4A-III | 0.02686 | 57.05351 | FALSE | FALSE |
| LOC107068225 | rootletin | 0.02508 | 57.05571 | FALSE | FALSE |
| LOC107065929 | transmembrane protein 164 | 0.02718 | 57.0674 | FALSE | FALSE |
| LOC107074152 | uncharacterized protein LOC107074152 | 0.02685 | 57.07788 | FALSE | FALSE |
| LOC107071613 | protein deglycase DJ-1zDJ-1-like | 0.0264 | 57.08188 | FALSE | FALSE |
| LOC107064500 | uncharacterized protein LOC107064500 | 0.0267 | 57.08665 | FALSE | FALSE |
| LOC107066474 | sodium/hydrogen exchanger 9B1-like | 0.02617 | 57.10253 | FALSE | FALSE |
| LOC107070256 | sodium-independent sulfate anion transpor | 0.0247 | 57.13646 | FALSE | FALSE |
| LOC107067806 | phosphatidylinositol-glycan biosynthesis cla | 0.02615 | 57.14345 | FALSE | FALSE |
| LOC107072495 | armadillo repeat-containing protein 4 | 0.02682 | 57.14546 | FALSE | FALSE |
| LOC107066493 | uncharacterized protein LOC107066493 | 0.02698 | 57.15251 | FALSE | FALSE |
| LOC107068775 | nuclear export mediator factor NEMF homo | 0.02545 | 57.17548 | FALSE | FALSE |
| LOC107068195 | uncharacterized protein LOC107068195 | 0.02703 | 57.19073 | FALSE | FALSE |
| LOC107067297 | serine-rich adhesin for platelets | 0.0252 | 57.22106 | FALSE | FALSE |
| LOC107069074 | UDP-glucuronosyltransferase 2C1-like | 0.02667 | 57.25891 | FALSE | FALSE |
| LOC107068111 | serine/threonine-protein kinase 25 | 0.02743 | 57.25945 | FALSE | FALSE |

| | | | | | |
|---|---|---|---|---|---|
| LOC107064775 | ankyrin repeat domain-containing protein 7 | 0.02663 | 57.25969 | FALSE | FALSE |
| LOC107070715 | allatostatin-A receptor | 0.02576 | 57.2696 | FALSE | FALSE |
| LOC107072692 | putative uncharacterized protein DDB_G027 | 0.0258 | 57.2777 | FALSE | FALSE |
| LOC107066326 | thioredoxin-related transmembrane proteir | 0.02609 | 57.2958 | FALSE | FALSE |
| LOC107067427 | IQ domain-containing protein G | 0.02594 | 57.30446 | FALSE | FALSE |
| LOC107069661 | nucleoporin GLE1 | 0.02712 | 57.35724 | FALSE | FALSE |
| LOC107071587 | serine/threonine-protein kinase PAK 2 | 0.02542 | 57.35815 | FALSE | FALSE |
| LOC107069792 | intraflagellar transport protein 52 homolog | 0.02651 | 57.36405 | FALSE | FALSE |
| LOC107066495 | uncharacterized protein LOC107066495 | 0.02497 | 57.36429 | FALSE | FALSE |
| LOC107066741 | 26S proteasome non-ATPase regulatory sub | 0.02682 | 57.37408 | FALSE | FALSE |
| LOC107071165 | TATA-box-binding protein | 0.02752 | 57.37866 | FALSE | FALSE |
| LOC107073445 | putative fatty acyl-CoA reductase CG8306 | 0.02703 | 57.39713 | FALSE | FALSE |
| LOC107068524 | trithorax group protein osa | 0.02645 | 57.40289 | FALSE | FALSE |
| LOC107073760 | uncharacterized protein LOC107073760 | 0.02708 | 57.41439 | FALSE | FALSE |
| LOC107072674 | glutamic acid-rich protein-like | 0.02557 | 57.4156 | FALSE | FALSE |
| LOC107073529 | uncharacterized protein LOC107073529 | 0.02572 | 57.41763 | FALSE | FALSE |
| LOC107072918 | probable phosphoserine aminotransferase | 0.02558 | 57.453 | FALSE | FALSE |
| LOC107074707 | runt-related transcription factor 3-like | 0.0253 | 57.45938 | FALSE | FALSE |
| LOC107064009 | cleavage and polyadenylation specificity fac | 0.02541 | 57.48438 | FALSE | FALSE |
| LOC107073447 | (11Z)-hexadec-11-enoyl-CoA conjugase-like | 0.02602 | 57.4884 | FALSE | FALSE |
| LOC107069476 | guanine deaminase | 0.02587 | 57.49302 | FALSE | FALSE |
| LOC107063995 | IQ and ubiquitin-like domain-containing pro | 0.02604 | 57.50803 | FALSE | FALSE |
| LOC107067043 | uncharacterized protein LOC107067043 | 0.02543 | 57.50839 | FALSE | FALSE |
| LOC107074038 | uncharacterized protein LOC107074038 | 0.02444 | 57.51176 | FALSE | FALSE |
| LOC107067141 | argininosuccinate lyase | 0.0266 | 57.51273 | FALSE | FALSE |
| LOC107073357 | methyltransferase-like protein 25 | 0.02618 | 57.53018 | FALSE | FALSE |
| LOC107069373 | antichymotrypsin-2-like | 0.02651 | 57.53571 | FALSE | FALSE |
| LOC107070920 | uncharacterized protein LOC107070920 | 0.0267 | 57.5447 | FALSE | FALSE |
| LOC107064341 | tetratricopeptide repeat protein 12-like | 0.02586 | 57.55324 | FALSE | FALSE |
| LOC107070316 | optomotor-blind protein-like | 0.02558 | 57.5796 | FALSE | FALSE |
| LOC107067204 | discoidin domain-containing receptor 2-like | 0.02594 | 57.5816 | FALSE | FALSE |
| LOC107072052 | phospholipase A1-like | 0.02582 | 57.60175 | FALSE | FALSE |
| LOC107070377 | histone-lysine N-methyltransferase, H3 lysir | 0.02745 | 57.6235 | FALSE | FALSE |
| LOC107068544 | uncharacterized protein LOC107068544 | 0.02417 | 57.63962 | FALSE | FALSE |
| LOC107074011 | protein lethal(2)essential for life-like | 0.02514 | 57.65125 | FALSE | FALSE |
| LOC107069197 | myosin-G heavy chain-like | 0.02566 | 57.65331 | FALSE | FALSE |
| LOC107067700 | uncharacterized protein LOC107067700 | 0.02687 | 57.66462 | FALSE | FALSE |
| LOC107072140 | protein sarah | 0.02687 | 57.70268 | FALSE | FALSE |
| LOC107069743 | uncharacterized protein LOC107069743 | 0.02602 | 57.70701 | FALSE | FALSE |
| LOC107071472 | uncharacterized protein LOC107071472 | 0.02488 | 57.71686 | FALSE | FALSE |
| LOC107072588 | cathepsin J-like | 0.02648 | 57.72602 | FALSE | FALSE |
| LOC107071620 | putative leucine-rich repeat-containing prot | 0.02601 | 57.74618 | FALSE | FALSE |
| LOC107066411 | alpha-2-macroglobulin receptor-associated | 0.02725 | 57.76236 | FALSE | FALSE |
| LOC107072063 | uncharacterized protein LOC107072063 | 0.02502 | 57.76511 | FALSE | FALSE |
| LOC107071568 | uncharacterized protein LOC107071568 | 0.0246 | 57.80396 | FALSE | FALSE |
| LOC107066327 | LOW QUALITY PROTEIN: protein Red-like | 0.02597 | 57.80651 | FALSE | FALSE |
| LOC107069659 | set1/Ash2 histone methyltransferase compl | 0.02545 | 57.81409 | FALSE | FALSE |
| LOC107066397 | 28S ribosomal protein S33, mitochondrial | 0.0265 | 57.82626 | FALSE | FALSE |
| LOC107072497 | protein FAM173B | 0.02611 | 57.83774 | FALSE | FALSE |
| LOC107068416 | uncharacterized protein LOC107068416 | 0.02786 | 57.85628 | FALSE | FALSE |

| | | | | | |
|---|---|---|---|---|---|
| LOC107069770 | uncharacterized protein LOC107069770 | 0.0259 | 57.88289 | FALSE | FALSE |
| LOC107067737 | protein drumstick | 0.02549 | 57.89249 | FALSE | FALSE |
| LOC107069075 | WD repeat-containing protein 74 | 0.02611 | 57.89369 | FALSE | FALSE |
| LOC107068533 | interference hedgehog-like | 0.02655 | 57.89663 | FALSE | FALSE |
| LOC107067745 | uncharacterized protein LOC107067745 | 0.02588 | 57.93816 | FALSE | FALSE |
| LOC107068272 | uncharacterized protein LOC107068272 | 0.02655 | 57.95707 | FALSE | FALSE |
| LOC107073789 | proline-rich protein 2-like | 0.02665 | 57.96102 | FALSE | FALSE |
| LOC107073943 | plexin-A4 | 0.02487 | 57.96202 | FALSE | FALSE |
| LOC107068426 | ATP-binding cassette sub-family G member | 0.02628 | 57.97064 | FALSE | FALSE |
| LOC107065483 | echinoderm microtubule-associated proteir | 0.02571 | 57.97353 | FALSE | FALSE |
| LOC107074470 | 60S ribosomal protein L13a | 0.02492 | 57.98561 | FALSE | FALSE |
| LOC107064531 | alpha- and gamma-adaptin-binding protein | 0.02509 | 57.99413 | FALSE | FALSE |
| LOC107069804 | cardioacceleratory peptide receptor-like | 0.02585 | 58.01281 | FALSE | FALSE |
| LOC107066330 | coiled-coil domain-containing protein 61-lik | 0.02735 | 58.01596 | FALSE | FALSE |
| LOC107070990 | uncharacterized protein LOC107070990 | 0.02715 | 58.01632 | FALSE | FALSE |
| LOC107070474 | uncharacterized protein LOC107070474 | 0.02665 | 58.02826 | FALSE | FALSE |
| LOC107073272 | sorting nexin-16 | 0.02634 | 58.03658 | FALSE | FALSE |
| LOC107072581 | DNA repair protein RAD52 homolog | 0.0272 | 58.03754 | FALSE | FALSE |
| LOC107066098 | rho-related BTB domain-containing protein | 0.02678 | 58.0379 | FALSE | FALSE |
| LOC107067604 | uncharacterized protein LOC107067604 | 0.02434 | 58.05885 | FALSE | FALSE |
| LOC107070645 | LOW QUALITY PROTEIN: uncharacterized pr | 0.02547 | 58.06565 | FALSE | FALSE |
| LOC107072888 | uncharacterized protein LOC107072888 | 0.02599 | 58.07285 | FALSE | FALSE |
| LOC107072369 | V-type proton ATPase 16 kDa proteolipid su | 0.02553 | 58.07356 | FALSE | FALSE |
| LOC107071079 | uncharacterized protein LOC107071079 | 0.02682 | 58.07825 | FALSE | FALSE |
| LOC107068924 | uncharacterized protein LOC107068924 | 0.02595 | 58.09162 | FALSE | FALSE |
| LOC107074142 | sodium channel protein Nach-like | 0.02543 | 58.09404 | FALSE | FALSE |
| LOC107073743 | uncharacterized protein LOC107073743 | 0.02552 | 58.09487 | FALSE | FALSE |
| LOC107072610 | growth arrest-specific protein 8 | 0.02644 | 58.09911 | FALSE | FALSE |
| LOC107070420 | uncharacterized protein LOC107070420 | 0.02559 | 58.11257 | FALSE | FALSE |
| LOC107065799 | zinc finger protein 250-like | 0.02647 | 58.11577 | FALSE | FALSE |
| LOC107066095 | uncharacterized protein LOC107066095 | 0.02752 | 58.11581 | FALSE | FALSE |
| LOC107071152 | protein NDRG3 | 0.02635 | 58.12032 | FALSE | FALSE |
| LOC107067085 | peptidyl-prolyl cis-trans isomerase 6-like | 0.02609 | 58.12942 | FALSE | FALSE |
| LOC107070826 | calcium and integrin-binding family membe | 0.02615 | 58.1469 | FALSE | FALSE |
| LOC107068384 | E3 ubiquitin-protein ligase NRDP1 | 0.02551 | 58.1567 | FALSE | FALSE |
| LOC107068038 | uncharacterized protein LOC107068038 | 0.02715 | 58.15722 | FALSE | FALSE |
| LOC107067359 | uncharacterized protein LOC107067359 | 0.02598 | 58.15781 | FALSE | FALSE |
| LOC107069744 | uncharacterized protein LOC107069744 | 0.02621 | 58.1584 | FALSE | FALSE |
| LOC107073624 | lipase 3-like | 0.02542 | 58.16224 | FALSE | FALSE |
| LOC107068660 | uncharacterized protein LOC107068660 | 0.02515 | 58.16819 | FALSE | FALSE |
| LOC107070855 | uncharacterized protein LOC107070855 | 0.0248 | 58.18219 | FALSE | FALSE |
| LOC107063826 | frizzled-2 | 0.02593 | 58.1912 | FALSE | FALSE |
| LOC107069015 | mucin-12-like | 0.02567 | 58.19166 | FALSE | FALSE |
| LOC107070417 | SH3 and multiple ankyrin repeat domains pr | 0.0268 | 58.20654 | FALSE | FALSE |
| LOC107073946 | trafficking protein particle complex subunit | 0.02549 | 58.20682 | FALSE | FALSE |
| LOC107069214 | mutS protein homolog 4-like | 0.02555 | 58.2075 | FALSE | FALSE |
| LOC107069650 | uncharacterized protein LOC107069650 | 0.02627 | 58.22156 | FALSE | FALSE |
| LOC107063790 | tubulin polyglutamylase TTLL5 | 0.02561 | 58.22635 | FALSE | FALSE |
| LOC107071100 | uncharacterized protein LOC107071100 | 0.0263 | 58.22857 | FALSE | FALSE |
| LOC107073085 | dnaJ homolog subfamily C member 11 | 0.0261 | 58.23252 | FALSE | FALSE |

| | | | | | |
|---|---|---|---|---|---|
| LOC107071924 | syntaxin-18 | 0.02785 | 58.24167 | FALSE | FALSE |
| LOC107071694 | ATP-dependent RNA helicase bel-like | 0.02697 | 58.24307 | FALSE | FALSE |
| LOC107070567 | odorant receptor 85c-like | 0.02712 | 58.26475 | FALSE | FALSE |
| LOC107067641 | proton-associated sugar transporter A | 0.02554 | 58.26988 | FALSE | FALSE |
| LOC107065669 | uncharacterized protein LOC107065669 | 0.0257 | 58.27225 | FALSE | FALSE |
| LOC107065198 | uncharacterized protein LOC107065198 | 0.02638 | 58.27344 | FALSE | FALSE |
| LOC107065232 | chaoptin | 0.02669 | 58.27822 | FALSE | FALSE |
| LOC107067111 | ras-related protein Rab-6.2 | 0.02609 | 58.27945 | FALSE | FALSE |
| LOC107068813 | ADP-ribosylation factor 1 | 0.02548 | 58.28142 | FALSE | FALSE |
| LOC107070007 | uncharacterized protein LOC107070007 | 0.02606 | 58.28347 | FALSE | FALSE |
| LOC107067516 | zinc finger protein interacting with ribonucl | 0.02636 | 58.29889 | FALSE | FALSE |
| LOC107072382 | leucine-rich repeat-containing protein 58 | 0.0257 | 58.30191 | FALSE | FALSE |
| LOC107069829 | integrator complex subunit 3 | 0.02627 | 58.33709 | FALSE | FALSE |
| LOC107069205 | TELO2-interacting protein 2-like | 0.02647 | 58.34106 | FALSE | FALSE |
| LOC107066173 | NADH dehydrogenase [ubiquinone | 0.02615 | 58.37338 | FALSE | FALSE |
| LOC107073772 | uncharacterized protein LOC107073772 | 0.0259 | 58.3796 | FALSE | FALSE |
| LOC107067632 | rab-like protein 3 | 0.02693 | 58.38082 | FALSE | FALSE |
| LOC107069082 | uncharacterized protein LOC107069082 | 0.02542 | 58.42418 | FALSE | FALSE |
| LOC107072564 | bifunctional arginine demethylase and lysyl- | 0.02605 | 58.42692 | FALSE | FALSE |
| LOC107068798 | BRISC and BRCA1-A complex member 1-like | 0.02533 | 58.42788 | FALSE | FALSE |
| LOC107065196 | sperm flagellar protein 2-like | 0.02705 | 58.4497 | FALSE | FALSE |
| LOC107074623 | eukaryotic translation initiation factor 5B-lik | 0.02407 | 58.45585 | FALSE | FALSE |
| LOC107066565 | sorting nexin-13-like | 0.02438 | 58.46887 | FALSE | FALSE |
| LOC107066949 | sex peptide receptor-like | 0.02541 | 58.491 | FALSE | FALSE |
| LOC107074211 | pleiotropic regulator 1 | 0.02602 | 58.49667 | FALSE | FALSE |
| LOC107066362 | F-box only protein 7-like | 0.0251 | 58.49889 | FALSE | FALSE |
| LOC107069232 | farnesol dehydrogenase-like | 0.0261 | 58.50025 | FALSE | FALSE |
| LOC107074644 | uncharacterized protein LOC107074644 | 0.02776 | 58.50079 | FALSE | FALSE |
| LOC107066065 | uncharacterized protein LOC107066065 | 0.02418 | 58.51562 | FALSE | FALSE |
| LOC107067708 | islet cell autoantigen 1 | 0.02541 | 58.52266 | FALSE | FALSE |
| LOC107072238 | chaoptin | 0.02496 | 58.54957 | FALSE | FALSE |
| LOC107068571 | uncharacterized protein LOC107068571 | 0.02561 | 58.55002 | FALSE | FALSE |
| LOC107072726 | voltage-dependent calcium channel subunit | 0.02584 | 58.55908 | FALSE | FALSE |
| LOC107066503 | diacylglycerol kinase theta | 0.02415 | 58.59624 | FALSE | FALSE |
| LOC107073140 | GATA zinc finger domain-containing protein | 0.02619 | 58.62229 | FALSE | FALSE |
| LOC107065607 | uncharacterized protein LOC107065607 | 0.02574 | 58.63903 | FALSE | FALSE |
| LOC107070396 | uncharacterized protein LOC107070396 | 0.02698 | 58.65105 | FALSE | FALSE |
| LOC107067554 | uncharacterized protein LOC107067554 | 0.02686 | 58.65536 | FALSE | FALSE |
| LOC107071655 | agrin-like | 0.02607 | 58.66372 | FALSE | FALSE |
| LOC107072956 | uncharacterized protein LOC107072956 | 0.02653 | 58.70223 | FALSE | FALSE |
| LOC107072715 | zinc finger protein 567-like | 0.02632 | 58.70655 | FALSE | FALSE |
| LOC107069396 | uncharacterized protein LOC107069396 | 0.02676 | 58.71326 | FALSE | FALSE |
| LOC107063751 | hydroxyacid oxidase 1 | 0.02457 | 58.72535 | FALSE | FALSE |
| LOC107070172 | achaete-scute complex protein T8-like | 0.02503 | 58.726 | FALSE | FALSE |
| LOC107068933 | transient receptor potential protein | 0.02561 | 58.76344 | FALSE | FALSE |
| LOC107068104 | cytosolic carboxypeptidase NnaD | 0.02615 | 58.76395 | FALSE | FALSE |
| LOC107064900 | putative uncharacterized protein DDB_G02 | 0.02464 | 58.76463 | FALSE | FALSE |
| LOC107067877 | VIP36-like protein | 0.02547 | 58.7727 | FALSE | FALSE |
| LOC107071087 | uncharacterized protein LOC107071087 | 0.02499 | 58.77712 | FALSE | FALSE |
| LOC107074112 | Krueppel homolog 2 | 0.02588 | 58.8308 | FALSE | FALSE |

| | | | | | |
|---|---|---|---|---|---|
| LOC107070257 | putative uncharacterized protein DDB_G028 | 0.02613 | 58.83572 | FALSE | FALSE |
| LOC107071609 | arginine-glutamic acid dipeptide repeats pro | 0.02755 | 58.85752 | FALSE | FALSE |
| LOC107067742 | uncharacterized protein LOC107067742 | 0.02406 | 58.86208 | FALSE | FALSE |
| LOC107073807 | uncharacterized protein LOC107073807 | 0.02634 | 58.87152 | FALSE | FALSE |
| LOC107069957 | endophilin-B1 | 0.02625 | 58.87729 | FALSE | FALSE |
| LOC107073156 | dehydrogenase/reductase SDR family meml | 0.02692 | 58.90795 | FALSE | FALSE |
| LOC107074306 | uncharacterized protein LOC107074306 | 0.02578 | 58.91419 | FALSE | FALSE |
| LOC107072105 | venom serine carboxypeptidase | 0.0257 | 58.92988 | FALSE | FALSE |
| LOC107072734 | uncharacterized protein LOC107072734 | 0.02629 | 58.93356 | FALSE | FALSE |
| LOC107068202 | ovarian-specific serine/threonine-protein ki | 0.02612 | 58.93482 | FALSE | FALSE |
| LOC107069484 | gonadal protein gdl | 0.02594 | 58.94403 | FALSE | FALSE |
| LOC107063722 | uncharacterized protein LOC107063722 | 0.02559 | 58.9907 | FALSE | FALSE |
| LOC107067047 | sensory neuron membrane protein 2-like | 0.02644 | 59.0163 | FALSE | FALSE |
| LOC107073176 | uncharacterized protein LOC107073176 | 0.02609 | 59.0172 | FALSE | FALSE |
| LOC107073569 | LOW QUALITY PROTEIN: cleavage stimulatic | 0.02546 | 59.02109 | FALSE | FALSE |
| LOC107069444 | UPF0396 protein CG6066 | 0.02519 | 59.07388 | FALSE | FALSE |
| LOC107064045 | cell death-inducing p53-target protein 1 | 0.02708 | 59.07851 | FALSE | FALSE |
| LOC107068284 | dnaJ homolog subfamily B member 13-like | 0.02577 | 59.08467 | FALSE | FALSE |
| LOC107072239 | jerky protein homolog-like | 0.02467 | 59.09494 | FALSE | FALSE |
| LOC107069129 | high-affinity choline transporter 1 | 0.02598 | 59.10826 | FALSE | FALSE |
| LOC107064117 | valacyclovir hydrolase | 0.02487 | 59.14904 | FALSE | FALSE |
| LOC107069961 | uncharacterized protein C19orf43 homolog | 0.02497 | 59.15533 | FALSE | FALSE |
| LOC107071730 | uncharacterized protein LOC107071730 | 0.0264 | 59.16011 | FALSE | FALSE |
| LOC107067215 | rho GTPase-activating protein 18-like | 0.02519 | 59.1607 | FALSE | FALSE |
| LOC107065870 | uncharacterized protein LOC107065870 | 0.02576 | 59.17019 | FALSE | FALSE |
| LOC107065014 | eukaryotic translation initiation factor 3 sub | 0.02576 | 59.19717 | FALSE | FALSE |
| LOC107067182 | thyroid receptor-interacting protein 6 | 0.02386 | 59.19846 | FALSE | FALSE |
| LOC107073510 | uncharacterized protein LOC107073510 | 0.02484 | 59.20727 | FALSE | FALSE |
| LOC107065539 | rhythmically expressed gene 5 protein | 0.0254 | 59.22543 | FALSE | FALSE |
| LOC107069009 | 40S ribosomal protein S23 | 0.02591 | 59.23647 | FALSE | FALSE |
| LOC107074117 | myosin regulatory light chain 2 | 0.02519 | 59.24266 | FALSE | FALSE |
| LOC107070606 | alpha-tocopherol transfer protein-like | 0.02643 | 59.25919 | FALSE | FALSE |
| LOC107074288 | uncharacterized protein LOC107074288 | 0.02722 | 59.26459 | FALSE | FALSE |
| LOC107064377 | protein pellino | 0.02691 | 59.28394 | FALSE | FALSE |
| LOC107067050 | corepressor interacting with RBPJ 1 | 0.02422 | 59.30443 | FALSE | FALSE |
| LOC107065762 | cell adhesion molecule 3-like | 0.02713 | 59.3135 | FALSE | FALSE |
| LOC107071543 | calcyphosin-like protein | 0.02628 | 59.31454 | FALSE | FALSE |
| LOC107071273 | alanine--tRNA ligase, mitochondrial | 0.02582 | 59.31742 | FALSE | FALSE |
| LOC107071537 | ATP-dependent RNA helicase SUV3 homolog | 0.02594 | 59.32818 | FALSE | FALSE |
| LOC107063679 | alcohol dehydrogenase [NADP(+) | 0.02553 | 59.34878 | FALSE | FALSE |
| LOC107072821 | uncharacterized protein LOC107072821 | 0.02584 | 59.3582 | FALSE | FALSE |
| LOC107068675 | electron transfer flavoprotein subunit alpha | 0.0264 | 59.37106 | FALSE | FALSE |
| LOC107064812 | CCR4-NOT transcription complex subunit 7 | 0.02498 | 59.37151 | FALSE | FALSE |
| LOC107065355 | uncharacterized protein LOC107065355 | 0.02549 | 59.39355 | FALSE | FALSE |
| LOC107064303 | uncharacterized protein LOC107064303 | 0.02475 | 59.40311 | FALSE | FALSE |
| LOC107072745 | nose resistant to fluoxetine protein 6-like | 0.02492 | 59.41194 | FALSE | FALSE |
| LOC107069010 | uncharacterized protein LOC107069010 | 0.02582 | 59.4248 | FALSE | FALSE |
| LOC107068346 | transmembrane protein 114 | 0.02503 | 59.4335 | FALSE | FALSE |
| LOC107074240 | uncharacterized protein LOC107074240 | 0.02599 | 59.44846 | FALSE | FALSE |
| LOC107071886 | MOB kinase activator-like 2 | 0.02574 | 59.45037 | FALSE | FALSE |

| | | | | | |
|---|---|---|---|---|---|
| LOC107073100 | succinyl-CoA ligase [ADP-forming | 0.02591 | 59.45686 | FALSE | FALSE |
| LOC107065066 | L-lactate dehydrogenase-like | 0.02546 | 59.4595 | FALSE | FALSE |
| LOC107072412 | glucose-6-phosphate 1-dehydrogenase | 0.02559 | 59.46172 | FALSE | FALSE |
| LOC107073026 | uncharacterized protein LOC107073026 | 0.02549 | 59.51381 | FALSE | FALSE |
| LOC107069110 | eukaryotic translation initiation factor 2-alp | 0.02496 | 59.51385 | FALSE | FALSE |
| LOC107073428 | uncharacterized protein LOC107073428 | 0.02534 | 59.53484 | FALSE | FALSE |
| LOC107067015 | uncharacterized protein LOC107067015 | 0.02562 | 59.53616 | FALSE | FALSE |
| LOC107073318 | histone-lysine N-methyltransferase E(z) | 0.02576 | 59.55585 | FALSE | FALSE |
| LOC107073146 | uncharacterized protein LOC107073146 | 0.02675 | 59.58078 | FALSE | FALSE |
| LOC107072741 | uncharacterized protein LOC107072741 | 0.0254 | 59.5827 | FALSE | FALSE |
| LOC107066859 | myosin-IIIb-like | 0.02437 | 59.58511 | FALSE | FALSE |
| LOC107065172 | uncharacterized protein LOC107065172 | 0.0252 | 59.59 | FALSE | FALSE |
| LOC107071508 | zinc finger protein ZIC 3-like | 0.02568 | 59.59134 | FALSE | FALSE |
| LOC107071426 | uncharacterized protein LOC107071426 | 0.02612 | 59.59237 | FALSE | FALSE |
| LOC107064353 | DNA replication licensing factor MCM4 | 0.02673 | 59.61708 | FALSE | FALSE |
| LOC107073252 | U1 small nuclear ribonucleoprotein C | 0.02562 | 59.63477 | FALSE | FALSE |
| LOC107072235 | nuclear pore complex protein Nup160 hom | 0.02716 | 59.64683 | FALSE | FALSE |
| LOC107065634 | methyltransferase-like protein 14 homolog | 0.02685 | 59.65249 | FALSE | FALSE |
| LOC107066225 | WD repeat-containing protein 87-like | 0.02517 | 59.66173 | FALSE | FALSE |
| LOC107064387 | trypsin-1-like | 0.026 | 59.66529 | FALSE | FALSE |
| LOC107071439 | 40S ribosomal protein S15 | 0.02703 | 59.71401 | FALSE | FALSE |
| LOC107064605 | protein spinster | 0.02573 | 59.72485 | FALSE | FALSE |
| LOC107068369 | TAF5-like RNA polymerase II p300/CBP-asso | 0.02593 | 59.73777 | FALSE | FALSE |
| LOC107067214 | cell growth regulator with RING finger dom | 0.02684 | 59.73842 | FALSE | FALSE |
| LOC107066802 | protein Dr1 | 0.02582 | 59.74865 | FALSE | FALSE |
| LOC107064360 | UDP-glucuronosyltransferase 2A3 | 0.02601 | 59.75177 | FALSE | FALSE |
| LOC107070897 | flavin-containing monooxygenase FMO GS- | 0.02737 | 59.7589 | FALSE | FALSE |
| LOC107065479 | uncharacterized protein LOC107065479 | 0.02508 | 59.7657 | FALSE | FALSE |
| LOC107070363 | TNF receptor-associated factor family prote | 0.02591 | 59.77518 | FALSE | FALSE |
| LOC107068860 | cholesterol 7-desaturase | 0.0253 | 59.78112 | FALSE | FALSE |
| LOC107069375 | calcium uptake protein 1 homolog, mitocho | 0.02376 | 59.78637 | FALSE | FALSE |
| LOC107064786 | bursicon | 0.02614 | 59.7883 | FALSE | FALSE |
| LOC107072643 | uncharacterized protein LOC107072643 | 0.02563 | 59.79396 | FALSE | FALSE |
| LOC107063899 | cAMP-dependent protein kinase catalytic su | 0.02563 | 59.79999 | FALSE | FALSE |
| LOC107069959 | 40S ribosomal protein S18 | 0.02349 | 59.81551 | FALSE | FALSE |
| LOC107071030 | tail-anchored protein insertion receptor WR | 0.02595 | 59.82411 | FALSE | FALSE |
| LOC107069278 | katanin p80 WD40 repeat-containing subun | 0.02551 | 59.85189 | FALSE | FALSE |
| LOC107066834 | uncharacterized protein LOC107066834 | 0.02549 | 59.85991 | FALSE | FALSE |
| LOC107066371 | cytoplasmic polyadenylation element-bindi | 0.02448 | 59.8731 | FALSE | FALSE |
| LOC107067586 | uncharacterized protein LOC107067586 | 0.02635 | 59.88112 | FALSE | FALSE |
| LOC107067248 | uncharacterized protein LOC107067248 | 0.02554 | 59.88172 | FALSE | FALSE |
| LOC107063653 | mitochondrial fission regulator 1-like | 0.0272 | 59.89317 | FALSE | FALSE |
| LOC107071183 | uncharacterized protein LOC107071183 | 0.02451 | 59.89774 | FALSE | FALSE |
| LOC107064804 | uncharacterized protein LOC107064804 | 0.02589 | 59.89906 | FALSE | FALSE |
| LOC107073534 | fatty acid synthase-like | 0.02408 | 59.92218 | FALSE | FALSE |
| LOC107071576 | LOW QUALITY PROTEIN: actin-binding prote | 0.02633 | 59.92345 | FALSE | FALSE |
| LOC107071240 | guanine nucleotide-binding protein G(I)/G(S | 0.02546 | 59.92434 | FALSE | FALSE |
| LOC107072355 | facilitated trehalose transporter Tret1-like | 0.02501 | 59.93102 | FALSE | FALSE |
| LOC107069438 | RAC-beta serine/threonine-protein kinase B | 0.02475 | 59.95488 | FALSE | FALSE |
| LOC107065332 | ubiquitin-fold modifier-conjugating enzyme | 0.02638 | 59.96716 | FALSE | FALSE |

| | | | | | |
|---|---|---|---|---|---|
| LOC107074104 | uncharacterized protein LOC107074104 | 0.0252 | 59.97124 | FALSE | FALSE |
| LOC107071968 | uncharacterized protein LOC107071968 | 0.02477 | 59.97982 | FALSE | FALSE |
| LOC107068045 | putative uncharacterized protein DDB_G028 | 0.02598 | 59.98246 | FALSE | FALSE |
| LOC107072768 | uncharacterized protein LOC107072768 | 0.02507 | 59.98961 | FALSE | FALSE |
| LOC107067616 | protein kinase C-binding protein 1-like | 0.02414 | 60.00041 | FALSE | FALSE |
| LOC107070357 | UV excision repair protein RAD23 homolog I | 0.02446 | 60.00129 | FALSE | FALSE |
| LOC107069694 | uncharacterized protein LOC107069694 | 0.02583 | 60.01572 | FALSE | FALSE |
| LOC107069803 | pleckstrin homology domain-containing fam | 0.02634 | 60.0197 | FALSE | FALSE |
| LOC107065464 | serine--tRNA ligase, mitochondrial | 0.02514 | 60.02684 | FALSE | FALSE |
| LOC107071453 | proton-coupled amino acid transporter 4-lik | 0.02748 | 60.03182 | FALSE | FALSE |
| LOC107073047 | uncharacterized protein LOC107073047 | 0.02487 | 60.03596 | FALSE | FALSE |
| LOC107068053 | ATP-dependent RNA helicase Ddx1 | 0.02573 | 60.07581 | FALSE | FALSE |
| LOC107072731 | solute carrier family 52, riboflavin transport | 0.02504 | 60.07615 | FALSE | FALSE |
| LOC107072392 | probable ATP-dependent RNA helicase DDX | 0.02658 | 60.09074 | FALSE | FALSE |
| LOC107063934 | probable cytochrome P450 12b2, mitochon | 0.02497 | 60.09762 | FALSE | FALSE |
| LOC107067758 | uncharacterized protein LOC107067758 | 0.02467 | 60.09775 | FALSE | FALSE |
| LOC107072469 | uncharacterized abhydrolase domain-contai | 0.02409 | 60.10412 | FALSE | FALSE |
| LOC107067574 | TBC1 domain family member 19 | 0.02587 | 60.10592 | FALSE | FALSE |
| LOC107067439 | protein msta, isoform B-like | 0.02451 | 60.11572 | FALSE | FALSE |
| LOC107072275 | transmembrane emp24 domain-containing | 0.02497 | 60.12174 | FALSE | FALSE |
| LOC107074426 | ATP-dependent helicase brm | 0.02413 | 60.12254 | FALSE | FALSE |
| LOC107072884 | uncharacterized protein LOC107072884 | 0.02606 | 60.13273 | FALSE | FALSE |
| LOC107066373 | zinc finger protein OZF-like | 0.02545 | 60.17917 | FALSE | FALSE |
| LOC107064072 | protein-L-isoaspartate(D-aspartate) O-meth | 0.02553 | 60.19371 | FALSE | FALSE |
| LOC107064943 | DNA polymerase subunit gamma-2, mitocho | 0.02389 | 60.19721 | FALSE | FALSE |
| LOC107074354 | zinc finger protein rotund | 0.02523 | 60.22091 | FALSE | FALSE |
| LOC107064813 | uncharacterized protein LOC107064813 | 0.02455 | 60.22746 | FALSE | FALSE |
| LOC107069933 | ets DNA-binding protein pokkuri | 0.02431 | 60.23957 | FALSE | FALSE |
| LOC107067256 | DNA ligase 4-like | 0.02549 | 60.24272 | FALSE | FALSE |
| LOC107073961 | NADH dehydrogenase [ubiquinone | 0.02521 | 60.25028 | FALSE | FALSE |
| LOC107068339 | uncharacterized protein LOC107068339 | 0.02574 | 60.28103 | FALSE | FALSE |
| LOC107070190 | PDF receptor-like | 0.02564 | 60.28386 | FALSE | FALSE |
| LOC107065743 | protein zyg-11 homolog B-like | 0.02486 | 60.2879 | FALSE | FALSE |
| LOC107073653 | uncharacterized protein LOC107073653 | 0.02547 | 60.29399 | FALSE | FALSE |
| LOC107065299 | fasciclin-2 | 0.02638 | 60.30483 | FALSE | FALSE |
| LOC107070693 | uncharacterized protein LOC107070693 | 0.02529 | 60.33279 | FALSE | FALSE |
| LOC107072502 | uncharacterized protein LOC107072502 | 0.02538 | 60.33561 | FALSE | FALSE |
| LOC107071724 | SLIT-ROBO Rho GTPase-activating protein 1- | 0.02592 | 60.338 | FALSE | FALSE |
| LOC107073930 | uncharacterized protein LOC107073930 | 0.02621 | 60.33836 | FALSE | FALSE |
| LOC107074650 | vasopressin V1a receptor-like | 0.0265 | 60.34144 | FALSE | FALSE |
| LOC107064213 | inhibitory POU protein | 0.02441 | 60.3521 | FALSE | FALSE |
| LOC107071838 | tumor necrosis factor receptor superfamily | 0.02417 | 60.35391 | FALSE | FALSE |
| LOC107069466 | adenylate kinase isoenzyme 5 | 0.02554 | 60.37501 | FALSE | FALSE |
| LOC107065574 | uncharacterized protein LOC107065574 | 0.02597 | 60.37977 | FALSE | FALSE |
| LOC107072505 | putative uncharacterized protein DDB_G028 | 0.02613 | 60.38903 | FALSE | FALSE |
| LOC107072322 | uncharacterized protein LOC107072322 | 0.02492 | 60.41151 | FALSE | FALSE |
| LOC107069507 | uncharacterized protein LOC107069507 | 0.02615 | 60.41784 | FALSE | FALSE |
| LOC107069895 | high mobility group protein 20A | 0.02563 | 60.42138 | FALSE | FALSE |
| LOC107070490 | sodium-dependent neutral amino acid trans | 0.02517 | 60.42205 | FALSE | FALSE |
| LOC107069940 | DNA-directed RNA polymerases I, II, and III s | 0.0258 | 60.44714 | FALSE | FALSE |

| | | | | | |
|---|---|---|---|---|---|
| LOC107073465 | uncharacterized protein LOC107073465 | 0.025 | 60.45264 | FALSE | FALSE |
| LOC107067211 | GATA-binding factor A-like | 0.02499 | 60.46711 | FALSE | FALSE |
| LOC107074043 | uncharacterized protein LOC107074043 | 0.025 | 60.46968 | FALSE | FALSE |
| LOC107071393 | cytoplasmic FMR1-interacting protein | 0.02501 | 60.47336 | FALSE | FALSE |
| LOC107069420 | uncharacterized protein LOC107069420 | 0.02664 | 60.47537 | FALSE | FALSE |
| LOC107070779 | DDB1- and CUL4-associated factor 12 | 0.02521 | 60.49329 | FALSE | FALSE |
| LOC107066810 | calmodulin-like protein 4 | 0.02421 | 60.4943 | FALSE | FALSE |
| LOC107073421 | uncharacterized protein LOC107073421 | 0.02607 | 60.49901 | FALSE | FALSE |
| LOC107070611 | odorant receptor coreceptor | 0.02609 | 60.50996 | FALSE | FALSE |
| LOC107067130 | kinetochore protein NDC80 homolog | 0.02636 | 60.52593 | FALSE | FALSE |
| LOC107071835 | uncharacterized protein LOC107071835 | 0.02617 | 60.52779 | FALSE | FALSE |
| LOC107070029 | muscle calcium channel subunit alpha-1 | 0.02539 | 60.52847 | FALSE | FALSE |
| LOC107074645 | cyclin-dependent kinase-like 4 | 0.02517 | 60.53836 | FALSE | FALSE |
| LOC107072594 | UPF0449 protein C19orf25-like | 0.02594 | 60.54342 | FALSE | FALSE |
| LOC107067110 | probable cysteine desulfurase, mitochondri | 0.02513 | 60.54446 | FALSE | FALSE |
| LOC107073963 | ceramide synthase 6 | 0.02556 | 60.54904 | FALSE | FALSE |
| LOC107064956 | ER lumen protein-retaining receptor | 0.02604 | 60.55015 | FALSE | FALSE |
| LOC107073480 | leucine-rich repeat-containing protein 15-lik | 0.02471 | 60.55922 | FALSE | FALSE |
| LOC107070716 | protein couch potato | 0.02502 | 60.56282 | FALSE | FALSE |
| LOC107074638 | titin | 0.02414 | 60.57682 | FALSE | FALSE |
| LOC107070169 | opioid-binding protein/cell adhesion molecu | 0.02585 | 60.59826 | FALSE | FALSE |
| LOC107070979 | radial spoke head protein 6 homolog A | 0.0257 | 60.62204 | FALSE | FALSE |
| LOC107070200 | E3 ubiquitin-protein ligase MARCH3-like | 0.02591 | 60.62692 | FALSE | FALSE |
| LOC107070489 | probable serine/threonine-protein kinase d | 0.02632 | 60.63159 | FALSE | FALSE |
| LOC107069911 | NAD-dependent protein deacetylase Sirt6 | 0.02603 | 60.66476 | FALSE | FALSE |
| LOC107074542 | N-sulphoglucosamine sulphohydrolase | 0.02542 | 60.66542 | FALSE | FALSE |
| LOC107069942 | uncharacterized protein LOC107069942 | 0.02463 | 60.6913 | FALSE | FALSE |
| LOC107069163 | DNA-directed RNA polymerase III subunit RI | 0.02462 | 60.69387 | FALSE | FALSE |
| LOC107066835 | cytochrome P450 9e2-like | 0.02619 | 60.71233 | FALSE | FALSE |
| LOC107072179 | uncharacterized protein LOC107072179 | 0.02524 | 60.72077 | FALSE | FALSE |
| LOC107073512 | uncharacterized protein LOC107073512 | 0.02482 | 60.74655 | FALSE | FALSE |
| LOC107067714 | congested-like trachea protein | 0.02489 | 60.74676 | FALSE | FALSE |
| LOC107072612 | MORN repeat-containing protein 4 | 0.02527 | 60.74741 | FALSE | FALSE |
| LOC107063857 | delta(3,5)-Delta(2,4)-dienoyl-CoA isomerase | 0.02936 | 60.74901 | FALSE | FALSE |
| LOC107069702 | uncharacterized protein LOC107069702 | 0.02526 | 60.77916 | FALSE | FALSE |
| LOC107064323 | endothelin-converting enzyme 1 | 0.02419 | 60.78318 | FALSE | FALSE |
| LOC107067914 | odorant receptor 4-like | 0.02435 | 60.80429 | FALSE | FALSE |
| LOC107070325 | replication protein A 70 kDa DNA-binding su | 0.0263 | 60.8056 | FALSE | FALSE |
| LOC107065940 | probable trans-2-enoyl-CoA reductase, mitc | 0.02475 | 60.80855 | FALSE | FALSE |
| LOC107071128 | uncharacterized protein LOC107071128 | 0.02445 | 60.82046 | FALSE | FALSE |
| LOC107074317 | RNA-binding protein 39 | 0.02534 | 60.84931 | FALSE | FALSE |
| LOC107071351 | crossover junction endonuclease MUS81 | 0.02377 | 60.85247 | FALSE | FALSE |
| LOC107064784 | zinc finger CCCH domain-containing protein | 0.02401 | 60.87067 | FALSE | FALSE |
| LOC107072789 | proton-coupled folate transporter | 0.02524 | 60.87388 | FALSE | FALSE |
| LOC107069730 | uncharacterized protein LOC107069730 | 0.02544 | 60.88836 | FALSE | FALSE |
| LOC107067836 | kinesin-like protein KIF6 | 0.02516 | 60.89543 | FALSE | FALSE |
| LOC107070442 | hypoxia-inducible factor 1-alpha | 0.02531 | 60.89812 | FALSE | FALSE |
| LOC107069404 | extensin-like | 0.02644 | 60.89833 | FALSE | FALSE |
| LOC107068366 | 40S ribosomal protein S25-like | 0.02463 | 60.90095 | FALSE | FALSE |
| LOC107071641 | arf-GAP domain and FG repeat-containing p | 0.02543 | 60.9199 | FALSE | FALSE |

| | | | | | |
|---|---|---|---|---|---|
| LOC107066947 | nuclear receptor-binding factor 2-like | 0.02469 | 60.93294 | FALSE | FALSE |
| LOC107070322 | uncharacterized protein LOC107070322 | 0.02598 | 60.93342 | FALSE | FALSE |
| LOC107069731 | zinc finger protein 622 | 0.02432 | 60.95014 | FALSE | FALSE |
| LOC107064676 | glutamate receptor ionotropic, kainate 2-lik | 0.02605 | 60.9712 | FALSE | FALSE |
| LOC107064098 | uncharacterized protein LOC107064098 | 0.02603 | 60.98062 | FALSE | FALSE |
| LOC107068713 | immunoglobulin-binding protein 1b | 0.0252 | 60.99293 | FALSE | FALSE |
| LOC107071402 | leucine-rich repeat-containing protein 20 | 0.02395 | 60.9935 | FALSE | FALSE |
| LOC107072456 | CD151 antigen | 0.02377 | 60.99663 | FALSE | FALSE |
| LOC107067509 | uncharacterized protein LOC107067509 | 0.02529 | 61.00221 | FALSE | FALSE |
| LOC107068548 | uncharacterized protein LOC107068548 | 0.02471 | 61.02114 | FALSE | FALSE |
| LOC107069761 | iron-sulfur cluster assembly 2 homolog, mit | 0.02662 | 61.02851 | FALSE | FALSE |
| LOC107074620 | probable basic-leucine zipper transcription f | 0.02649 | 61.03615 | FALSE | FALSE |
| LOC107073742 | uncharacterized protein LOC107073742 | 0.02577 | 61.05088 | FALSE | FALSE |
| LOC107073122 | WAP four-disulfide core domain protein 18 | 0.02461 | 61.05957 | FALSE | FALSE |
| LOC107071740 | monoacylglycerol lipase ABHD12-like | 0.02627 | 61.06086 | FALSE | FALSE |
| LOC107072877 | zinc finger protein Pegasus | 0.02586 | 61.06191 | FALSE | FALSE |
| LOC107064945 | uncharacterized protein LOC107064945 | 0.02612 | 61.06956 | FALSE | FALSE |
| LOC107069587 | tumor protein p63-regulated gene 1-like pro | 0.02552 | 61.09062 | FALSE | FALSE |
| LOC107074055 | glutamate receptor ionotropic, delta-1-like | 0.02485 | 61.09247 | FALSE | FALSE |
| LOC107064621 | protocadherin-like wing polarity protein sta | 0.02326 | 61.09739 | FALSE | FALSE |
| LOC107072996 | ubiquitin-conjugating enzyme E2 W | 0.02383 | 61.10099 | FALSE | FALSE |
| LOC107070808 | polynucleotide 5'-hydroxyl-kinase NOL9-like | 0.02505 | 61.12037 | FALSE | FALSE |
| LOC107073438 | uncharacterized protein LOC107073438 | 0.02404 | 61.19706 | FALSE | FALSE |
| LOC107073964 | protein amalgam-like | 0.0259 | 61.19797 | FALSE | FALSE |
| LOC107069645 | PAX-interacting protein 1 | 0.02601 | 61.20135 | FALSE | FALSE |
| LOC107065226 | tubulin-specific chaperone C | 0.02485 | 61.20199 | FALSE | FALSE |
| LOC107072980 | probable ATP-dependent RNA helicase DDX | 0.02392 | 61.22304 | FALSE | FALSE |
| LOC107063973 | isocitrate dehydrogenase [NADP | 0.02461 | 61.22792 | FALSE | FALSE |
| LOC107069703 | kelch domain-containing protein 10 homolo | 0.02364 | 61.24242 | FALSE | FALSE |
| LOC107067608 | uncharacterized protein LOC107067608 | 0.02581 | 61.25919 | FALSE | FALSE |
| LOC107068479 | mediator of RNA polymerase II transcription | 0.02575 | 61.26368 | FALSE | FALSE |
| LOC107069607 | uncharacterized protein LOC107069607 | 0.02596 | 61.27018 | FALSE | FALSE |
| LOC107065151 | transcription initiation factor TFIID subunit 8 | 0.02562 | 61.27809 | FALSE | FALSE |
| LOC107071362 | small nuclear ribonucleoprotein Sm D3 | 0.02482 | 61.28271 | FALSE | FALSE |
| LOC107064088 | probable cardiolipin synthase (CMP-forming | 0.02519 | 61.29132 | FALSE | FALSE |
| LOC107067437 | protein NDNF | 0.02394 | 61.29277 | FALSE | FALSE |
| LOC107068168 | fanconi-associated nuclease 1-like | 0.0253 | 61.31442 | FALSE | FALSE |
| LOC107074441 | carboxypeptidase D | 0.02583 | 61.34204 | FALSE | FALSE |
| LOC107067924 | DEAD-box ATP-dependent RNA helicase 20- | 0.0251 | 61.35133 | FALSE | FALSE |
| LOC107066805 | uncharacterized protein LOC107066805 | 0.02501 | 61.36249 | FALSE | FALSE |
| LOC107067096 | U3 small nucleolar RNA-associated protein 6 | 0.02408 | 61.37486 | FALSE | FALSE |
| LOC107066216 | histone acetyltransferase KAT8-like | 0.02555 | 61.43247 | FALSE | FALSE |
| LOC107068285 | homeotic protein antennapedia-like | 0.02636 | 61.43269 | FALSE | FALSE |
| LOC107071226 | X-ray repair cross-complementing protein 6 | 0.02479 | 61.43922 | FALSE | FALSE |
| LOC107071147 | uncharacterized protein LOC107071147 | 0.02341 | 61.44547 | FALSE | FALSE |
| LOC107065740 | tRNA modification GTPase MnmE | 0.02595 | 61.44563 | FALSE | FALSE |
| LOC107067511 | dynein intermediate chain 2, axonemal | 0.02527 | 61.48083 | FALSE | FALSE |
| LOC107071842 | uncharacterized protein LOC107071842 | 0.02487 | 61.4856 | FALSE | FALSE |
| LOC107070386 | A disintegrin and metalloproteinase with th | 0.02552 | 61.48671 | FALSE | FALSE |
| LOC107072998 | peroxiredoxin-like | 0.02444 | 61.49961 | FALSE | FALSE |

| | | | | | |
|---|---|---|---|---|---|
| LOC107069899 | uncharacterized protein LOC107069899 | 0.02477 | 61.519 | FALSE | FALSE |
| LOC107065933 | uncharacterized protein LOC107065933 | 0.02571 | 61.52235 | FALSE | FALSE |
| LOC107067753 | coiled-coil domain-containing protein 58 | 0.02513 | 61.52518 | FALSE | FALSE |
| LOC107066918 | CD63 antigen-like | 0.02428 | 61.54805 | FALSE | FALSE |
| LOC107067658 | sodium/hydrogen exchanger 8 | 0.0257 | 61.55001 | FALSE | FALSE |
| LOC107068106 | uncharacterized protein LOC107068106 | 0.02451 | 61.555 | FALSE | FALSE |
| LOC107073395 | rab3 GTPase-activating protein non-catalyti | 0.02363 | 61.56164 | FALSE | FALSE |
| LOC107073839 | SEC14 domain and spectrin repeat-containi | 0.02509 | 61.57998 | FALSE | FALSE |
| LOC107064106 | uncharacterized protein LOC107064106 | 0.02621 | 61.58369 | FALSE | FALSE |
| LOC107074465 | retinoic acid receptor RXR | 0.02492 | 61.60392 | FALSE | FALSE |
| LOC107070660 | uncharacterized protein C1orf43 homolog | 0.02533 | 61.61434 | FALSE | FALSE |
| LOC107065980 | G8 domain-containing protein DDB_G0286 | 0.02562 | 61.62322 | FALSE | FALSE |
| LOC107065035 | mucolipin-3-like | 0.02652 | 61.62455 | FALSE | FALSE |
| LOC107067379 | secretion-regulating guanine nucleotide exc | 0.02518 | 61.63009 | FALSE | FALSE |
| LOC107069736 | vacuolar protein sorting-associated protein | 0.02451 | 61.63468 | FALSE | FALSE |
| LOC107064669 | uncharacterized protein LOC107064669 | 0.02488 | 61.63882 | FALSE | FALSE |
| LOC107070670 | A disintegrin and metalloproteinase with th | 0.02553 | 61.64603 | FALSE | FALSE |
| LOC107072139 | chondroitin proteoglycan-2-like | 0.02532 | 61.64949 | FALSE | FALSE |
| LOC107074454 | beta-1,4-mannosyltransferase egh | 0.02554 | 61.6569 | FALSE | FALSE |
| LOC107069851 | leucine-rich repeat neuronal protein 1-like | 0.0254 | 61.6694 | FALSE | FALSE |
| LOC107069243 | protein scarlet | 0.02491 | 61.68083 | FALSE | FALSE |
| LOC107071523 | sorting nexin-24-like | 0.02565 | 61.69087 | FALSE | FALSE |
| LOC107064851 | guanine nucleotide-binding protein G(q) sub | 0.02406 | 61.69224 | FALSE | FALSE |
| LOC107067955 | uncharacterized protein LOC107067955 | 0.02479 | 61.69872 | FALSE | FALSE |
| LOC107067030 | probable histone-binding protein Caf1 | 0.02441 | 61.7026 | FALSE | FALSE |
| LOC107064121 | tyrosine-protein kinase receptor torso-like | 0.02453 | 61.70584 | FALSE | FALSE |
| LOC107073924 | cyclin-dependent kinase 11B | 0.0252 | 61.71309 | FALSE | FALSE |
| LOC107068165 | zinc finger protein rotund-like | 0.02457 | 61.72732 | FALSE | FALSE |
| LOC107074279 | 46 kDa FK506-binding nuclear protein | 0.02573 | 61.74351 | FALSE | FALSE |
| LOC107074482 | uncharacterized protein LOC107074482 | 0.02456 | 61.77965 | FALSE | FALSE |
| LOC107071114 | uncharacterized protein LOC107071114 | 0.02453 | 61.78215 | FALSE | FALSE |
| LOC107064529 | coiled-coil domain-containing protein 115-li | 0.02569 | 61.79765 | FALSE | FALSE |
| LOC107074449 | thioredoxin domain-containing protein 9 | 0.02602 | 61.79952 | FALSE | FALSE |
| LOC107065498 | uncharacterized protein LOC107065498 | 0.02423 | 61.80712 | FALSE | FALSE |
| LOC107065388 | proline-rich protein 4-like | 0.02676 | 61.84363 | FALSE | FALSE |
| LOC107070255 | DDB1- and CUL4-associated factor 5 | 0.0245 | 61.84993 | FALSE | FALSE |
| LOC107069453 | uncharacterized protein LOC107069453 | 0.02478 | 61.85998 | FALSE | FALSE |
| LOC107065604 | RNA-binding protein 33-like | 0.02337 | 61.86899 | FALSE | FALSE |
| LOC107072363 | uncharacterized protein LOC107072363 | 0.02616 | 61.87092 | FALSE | FALSE |
| LOC107068956 | odorant receptor 13a-like | 0.02461 | 61.87747 | FALSE | FALSE |
| LOC107074522 | leucine-rich repeat-containing protein 70-lik | 0.02444 | 61.89148 | FALSE | FALSE |
| LOC107064251 | phosphoglycerate kinase | 0.02542 | 61.92278 | FALSE | FALSE |
| LOC107067823 | dopamine receptor 2 | 0.02341 | 61.93127 | FALSE | FALSE |
| LOC107068417 | uncharacterized protein LOC107068417 | 0.02387 | 61.93746 | FALSE | FALSE |
| LOC107066982 | ankyrin repeat and LEM domain-containing | 0.02436 | 61.94116 | FALSE | FALSE |
| LOC107069017 | uncharacterized protein LOC107069017 | 0.02494 | 61.94215 | FALSE | FALSE |
| LOC107069534 | isopentenyl-diphosphate Delta-isomerase 1 | 0.02572 | 61.95507 | FALSE | FALSE |
| LOC107063776 | neo-calmodulin-like | 0.02453 | 61.95727 | FALSE | FALSE |
| LOC107066938 | alpha-protein kinase 1 | 0.02479 | 61.9601 | FALSE | FALSE |
| LOC107064077 | prolyl endopeptidase | 0.02524 | 61.97451 | FALSE | FALSE |

| | | | | | |
|---|---|---|---|---|---|
| LOC107070065 | uncharacterized protein LOC107070065 | 0.02479 | 61.98518 | FALSE | FALSE |
| LOC107065807 | uncharacterized protein LOC107065807 | 0.0255 | 62.00376 | FALSE | FALSE |
| LOC107071790 | phosphatidate phosphatase LPIN3 | 0.02465 | 62.04488 | FALSE | FALSE |
| LOC107068387 | probable trafficking protein particle comple | 0.02534 | 62.06071 | FALSE | FALSE |
| LOC107070946 | uncharacterized protein LOC107070946 | 0.02511 | 62.06369 | FALSE | FALSE |
| LOC107066542 | two pore calcium channel protein 1-like | 0.02551 | 62.06883 | FALSE | FALSE |
| LOC107070472 | innexin inx3 | 0.02557 | 62.07214 | FALSE | FALSE |
| LOC107066633 | protein LMBR1L | 0.02455 | 62.08302 | FALSE | FALSE |
| LOC107072775 | uncharacterized protein LOC107072775 | 0.02378 | 62.08398 | FALSE | FALSE |
| LOC107070449 | probable ATP-dependent RNA helicase DDX | 0.02488 | 62.08686 | FALSE | FALSE |
| LOC107066052 | 5-formyltetrahydrofolate cyclo-ligase | 0.02432 | 62.12058 | FALSE | FALSE |
| LOC107067886 | 60 kDa SS-A/Ro ribonucleoprotein-like | 0.02486 | 62.12626 | FALSE | FALSE |
| LOC107065658 | uncharacterized protein LOC107065658 | 0.02491 | 62.13243 | FALSE | FALSE |
| LOC107065055 | 1-phosphatidylinositol 4,5-bisphosphate ph | 0.02594 | 62.15755 | FALSE | FALSE |
| LOC107064259 | histone H3.3 | 0.02577 | 62.16182 | FALSE | FALSE |
| LOC107064890 | uncharacterized protein LOC107064890 | 0.02509 | 62.1823 | FALSE | FALSE |
| LOC107074318 | G patch domain-containing protein 1 homol | 0.02279 | 62.19189 | FALSE | FALSE |
| LOC107065630 | uncharacterized protein LOC107065630 | 0.02428 | 62.22453 | FALSE | FALSE |
| LOC107065951 | uncharacterized protein LOC107065951 | 0.0261 | 62.2352 | FALSE | FALSE |
| LOC107065901 | transmembrane and coiled-coil domain-con | 0.02468 | 62.24252 | FALSE | FALSE |
| LOC107066683 | uncharacterized protein LOC107066683 | 0.02511 | 62.2516 | FALSE | FALSE |
| LOC107072267 | uncharacterized protein LOC107072267 | 0.0255 | 62.27289 | FALSE | FALSE |
| LOC107066365 | uncharacterized protein LOC107066365 | 0.02442 | 62.28007 | FALSE | FALSE |
| LOC107073773 | uncharacterized protein LOC107073773 | 0.02564 | 62.28128 | FALSE | FALSE |
| LOC107068693 | uncharacterized protein LOC107068693 | 0.02471 | 62.28233 | FALSE | FALSE |
| LOC107071870 | proline-rich protein PRCC-like | 0.0232 | 62.28248 | FALSE | FALSE |
| LOC107072824 | mediator of RNA polymerase II transcriptior | 0.02455 | 62.29155 | FALSE | FALSE |
| LOC107070164 | vesicular glutamate transporter 1 | 0.02577 | 62.29348 | FALSE | FALSE |
| LOC107072403 | ejaculatory bulb-specific protein 3-like | 0.02626 | 62.29951 | FALSE | FALSE |
| LOC107071743 | uncharacterized protein LOC107071743 | 0.02498 | 62.3789 | FALSE | FALSE |
| LOC107065131 | carnitine O-palmitoyltransferase 2, mitocho | 0.02491 | 62.3925 | FALSE | FALSE |
| LOC107068620 | uncharacterized protein MAL13P1.304-like | 0.0241 | 62.40553 | FALSE | FALSE |
| LOC107069418 | post-GPI attachment to proteins factor 3 | 0.02503 | 62.41015 | FALSE | FALSE |
| LOC107068206 | ras-related protein Rab-8A | 0.02411 | 62.41286 | FALSE | FALSE |
| LOC107067603 | uncharacterized protein LOC107067603 | 0.02444 | 62.42977 | FALSE | FALSE |
| LOC107069553 | 28S ribosomal protein S24, mitochondrial | 0.0252 | 62.46143 | FALSE | FALSE |
| LOC107065873 | uncharacterized protein LOC107065873 | 0.02404 | 62.46416 | FALSE | FALSE |
| LOC107070533 | uncharacterized protein LOC107070533 | 0.02679 | 62.46788 | FALSE | FALSE |
| LOC107068613 | splicing factor 3B subunit 4 | 0.02482 | 62.47129 | FALSE | FALSE |
| LOC107066449 | uncharacterized protein LOC107066449 | 0.02525 | 62.48031 | FALSE | FALSE |
| LOC107068851 | protein tiptop | 0.02341 | 62.49061 | FALSE | FALSE |
| LOC107065895 | division abnormally delayed protein | 0.02388 | 62.51963 | FALSE | FALSE |
| LOC107066372 | protogenin | 0.02464 | 62.54586 | FALSE | FALSE |
| LOC107067538 | carboxypeptidase Q-like | 0.0246 | 62.55395 | FALSE | FALSE |
| LOC107073440 | uncharacterized protein LOC107073440 | 0.02432 | 62.56574 | FALSE | FALSE |
| LOC107070885 | phospholipid scramblase 2-like | 0.02632 | 62.57253 | FALSE | FALSE |
| LOC107068224 | TBC domain-containing protein kinase-like p | 0.02371 | 62.62891 | FALSE | FALSE |
| LOC107064440 | putative uncharacterized protein DDB_G028 | 0.02385 | 62.62921 | FALSE | FALSE |
| LOC107071654 | zinc transporter ZIP1-like | 0.024 | 62.63624 | FALSE | FALSE |
| LOC107063880 | uncharacterized protein LOC107063880 | 0.02414 | 62.64171 | FALSE | FALSE |

| | | | | | |
|---|---|---|---|---|---|
| LOC107071957 | bone morphogenetic protein 2-like | 0.02462 | 62.65453 | FALSE | FALSE |
| LOC107063676 | serine protease easter-like | 0.02461 | 62.66485 | FALSE | FALSE |
| LOC107069327 | uncharacterized protein LOC107069327 | 0.02389 | 62.68829 | FALSE | FALSE |
| LOC107068651 | uncharacterized protein LOC107068651 | 0.02549 | 62.69268 | FALSE | FALSE |
| LOC107065825 | ankyrin-1-like | 0.02353 | 62.69676 | FALSE | FALSE |
| LOC107063783 | endoplasmic reticulum resident protein 44 | 0.02514 | 62.7104 | FALSE | FALSE |
| LOC107072793 | uncharacterized protein LOC107072793 | 0.02602 | 62.73499 | FALSE | FALSE |
| LOC107065475 | putative pre-mRNA-splicing factor ATP-depe | 0.02533 | 62.74627 | FALSE | FALSE |
| LOC107070938 | eye-specific diacylglycerol kinase | 0.02452 | 62.76887 | FALSE | FALSE |
| LOC107069300 | cGMP-dependent 3',5'-cyclic phosphodieste | 0.02399 | 62.77961 | FALSE | FALSE |
| LOC107068618 | uncharacterized protein LOC107068618 | 0.02443 | 62.78825 | FALSE | FALSE |
| LOC107071227 | uncharacterized protein LOC107071227 | 0.02518 | 62.79213 | FALSE | FALSE |
| LOC107065995 | MAD2L1-binding protein | 0.02422 | 62.82823 | FALSE | FALSE |
| LOC107065904 | epidermal retinol dehydrogenase 2-like | 0.02467 | 62.83854 | FALSE | FALSE |
| LOC107074369 | uncharacterized protein LOC107074369 | 0.02424 | 62.84534 | FALSE | FALSE |
| LOC107067333 | inner nuclear membrane protein Man1 | 0.02335 | 62.85188 | FALSE | FALSE |
| LOC107065028 | ATP-dependent RNA helicase DHX36 | 0.02695 | 62.86272 | FALSE | FALSE |
| LOC107069027 | alpha-protein kinase 1-like | 0.02426 | 62.87744 | FALSE | FALSE |
| LOC107067799 | serine/threonine/tyrosine-interacting prote | 0.0251 | 62.91047 | FALSE | FALSE |
| LOC107063921 | uncharacterized protein LOC107063921 | 0.02463 | 62.91944 | FALSE | FALSE |
| LOC107070878 | cytosolic Fe-S cluster assembly factor NUBP | 0.02436 | 62.92065 | FALSE | FALSE |
| LOC107064376 | probable transaldolase | 0.02466 | 62.92207 | FALSE | FALSE |
| LOC107072785 | protein SCO1 homolog, mitochondrial | 0.02421 | 62.92875 | FALSE | FALSE |
| LOC107071619 | succinate dehydrogenase [ubiquinone | 0.02456 | 62.93125 | FALSE | FALSE |
| LOC107067197 | mitochondrial glutamate carrier 1-like | 0.02503 | 62.93624 | FALSE | FALSE |
| LOC107063743 | vacuolar protein sorting-associated protein | 0.02468 | 62.97778 | FALSE | FALSE |
| LOC107067940 | DDRGK domain-containing protein 1 | 0.02559 | 63.00037 | FALSE | FALSE |
| LOC107070213 | uncharacterized protein LOC107070213 | 0.0254 | 63.00521 | FALSE | FALSE |
| LOC107073985 | uncharacterized protein LOC107073985 | 0.02466 | 63.00667 | FALSE | FALSE |
| LOC107071951 | uncharacterized protein LOC107071951 | 0.02594 | 63.01408 | FALSE | FALSE |
| LOC107072453 | DNA repair protein complementing XP-A cel | 0.02527 | 63.01803 | FALSE | FALSE |
| LOC107065859 | carboxy-terminal domain RNA polymerase I | 0.02467 | 63.02432 | FALSE | FALSE |
| LOC107063813 | dnaJ homolog dnj-5 | 0.02315 | 63.04842 | FALSE | FALSE |
| LOC107063655 | uncharacterized protein LOC107063655 | 0.02412 | 63.08188 | FALSE | FALSE |
| LOC107067486 | proteasome subunit beta type-7 | 0.02371 | 63.0867 | FALSE | FALSE |
| LOC107063872 | uncharacterized protein LOC107063872 | 0.02504 | 63.16605 | FALSE | FALSE |
| LOC107066664 | uncharacterized protein LOC107066664 | 0.02412 | 63.16839 | FALSE | FALSE |
| LOC107072494 | sodium-coupled neutral amino acid transpo | 0.02408 | 63.18555 | FALSE | FALSE |
| LOC107064792 | dnaJ homolog subfamily C member 17 | 0.02543 | 63.19056 | FALSE | FALSE |
| LOC107071373 | uncharacterized protein LOC107071373 | 0.02515 | 63.21235 | FALSE | FALSE |
| LOC107066148 | uncharacterized protein LOC107066148 | 0.02309 | 63.26052 | FALSE | FALSE |
| LOC107067958 | uncharacterized protein LOC107067958 | 0.02483 | 63.26602 | FALSE | FALSE |
| LOC107072051 | uncharacterized protein LOC107072051 | 0.025 | 63.28759 | FALSE | FALSE |
| LOC107064964 | THUMP domain-containing protein 3-like | 0.02397 | 63.29263 | FALSE | FALSE |
| LOC107066925 | vascular endothelial growth factor receptor | 0.02549 | 63.29623 | FALSE | FALSE |
| LOC107074051 | homeobox protein aristaless-like | 0.0247 | 63.2987 | FALSE | FALSE |
| LOC107068710 | polypeptide N-acetylgalactosaminyltransfer | 0.02454 | 63.30257 | FALSE | FALSE |
| LOC107064250 | tyrosine-protein kinase hopscotch | 0.02338 | 63.30285 | FALSE | FALSE |
| LOC107064541 | uncharacterized protein LOC107064541 | 0.02558 | 63.3156 | FALSE | FALSE |
| LOC107067971 | proliferation-associated protein 2G4 | 0.0248 | 63.3373 | FALSE | FALSE |

| | | | | | |
|---|---|---|---|---|---|
| LOC107065680 | transcription elongation regulator 1 | 0.02374 | 63.33732 | FALSE | FALSE |
| LOC107068742 | E3 ubiquitin-protein ligase RNF19A-like | 0.02376 | 63.33879 | FALSE | FALSE |
| LOC107069934 | MICOS complex subunit MIC27 | 0.02433 | 63.34814 | FALSE | FALSE |
| LOC107064984 | protein lines | 0.02461 | 63.35014 | FALSE | FALSE |
| LOC107072178 | ras-related protein RABF2b | 0.02385 | 63.35891 | FALSE | FALSE |
| LOC107065860 | ER membrane protein complex subunit 7 | 0.02333 | 63.38861 | FALSE | FALSE |
| LOC107067699 | glutamate receptor ionotropic, NMDA 2B | 0.02351 | 63.40265 | FALSE | FALSE |
| LOC107071938 | lisH domain and HEAT repeat-containing pro | 0.02363 | 63.40699 | FALSE | FALSE |
| LOC107066444 | thiamin pyrophosphokinase 1 | 0.0263 | 63.40903 | FALSE | FALSE |
| LOC107064759 | G patch domain and KOW motifs-containing | 0.02378 | 63.40965 | FALSE | FALSE |
| LOC107069517 | band 4.1-like protein 5 | 0.02379 | 63.41063 | FALSE | FALSE |
| LOC107072807 | 26S proteasome non-ATPase regulatory sub | 0.02589 | 63.44165 | FALSE | FALSE |
| LOC107066393 | 2-oxoisovalerate dehydrogenase subunit alp | 0.02521 | 63.44863 | FALSE | FALSE |
| LOC107070593 | sodium-independent sulfate anion transpor | 0.02428 | 63.45345 | FALSE | FALSE |
| LOC107070619 | glucose dehydrogenase [FAD, quinone | 0.02481 | 63.46672 | FALSE | FALSE |
| LOC107066818 | uncharacterized protein LOC107066818 | 0.02412 | 63.4668 | FALSE | FALSE |
| LOC107071352 | splicing factor 3B subunit 2 | 0.02547 | 63.51803 | FALSE | FALSE |
| LOC107068495 | glutamate receptor-like | 0.02405 | 63.52056 | FALSE | FALSE |
| LOC107074494 | putative phosphatidate phosphatase | 0.02474 | 63.52853 | FALSE | FALSE |
| LOC107068186 | proteasome assembly chaperone 1 | 0.02541 | 63.53565 | FALSE | FALSE |
| LOC107066566 | uncharacterized protein LOC107066566 | 0.02566 | 63.54623 | FALSE | FALSE |
| LOC107069218 | centrosomal protein of 135 kDa | 0.02318 | 63.5474 | FALSE | FALSE |
| LOC107069995 | protein bark beetle | 0.02565 | 63.54998 | FALSE | FALSE |
| LOC107069967 | uncharacterized protein LOC107069967 | 0.02431 | 63.55467 | FALSE | FALSE |
| LOC107064905 | elongation of very long chain fatty acids pro | 0.024 | 63.60592 | FALSE | FALSE |
| LOC107067212 | nuclear receptor subfamily 2 group E memb | 0.02341 | 63.62392 | FALSE | FALSE |
| LOC107073980 | uncharacterized protein LOC107073980 | 0.02558 | 63.63495 | FALSE | FALSE |
| LOC107066277 | uncharacterized protein LOC107066277 | 0.02396 | 63.64377 | FALSE | FALSE |
| LOC107066369 | zinc finger protein 664-like | 0.02467 | 63.65049 | FALSE | FALSE |
| LOC107063719 | 39S ribosomal protein L11, mitochondrial | 0.02466 | 63.6517 | FALSE | FALSE |
| LOC107072291 | putative lipoyltransferase 2, mitochondrial | 0.02583 | 63.66515 | FALSE | FALSE |
| LOC107073414 | protein tipE | 0.02503 | 63.69235 | FALSE | FALSE |
| LOC107068306 | RPII140-upstream gene protein | 0.02349 | 63.69312 | FALSE | FALSE |
| LOC107064203 | signal peptidase complex subunit 1 | 0.02536 | 63.69338 | FALSE | FALSE |
| LOC107067435 | ubiquitin-fold modifier 1 | 0.02585 | 63.69867 | FALSE | FALSE |
| LOC107067145 | uncharacterized protein LOC107067145 | 0.02359 | 63.71504 | FALSE | FALSE |
| LOC107073076 | transcription initiation factor TFIID subunit 3 | 0.0234 | 63.72054 | FALSE | FALSE |
| LOC107073516 | uncharacterized protein LOC107073516 | 0.02379 | 63.72577 | FALSE | FALSE |
| LOC107070650 | uncharacterized protein LOC107070650 | 0.02513 | 63.73236 | FALSE | FALSE |
| LOC107074210 | serine--tRNA ligase, cytoplasmic | 0.02487 | 63.74273 | FALSE | FALSE |
| LOC107071055 | probable U2 small nuclear ribonucleoprotei | 0.02509 | 63.75264 | FALSE | FALSE |
| LOC107071215 | roundabout homolog 2-like | 0.02408 | 63.75727 | FALSE | FALSE |
| LOC107067426 | annexin B9-like | 0.02335 | 63.76749 | FALSE | FALSE |
| LOC107068828 | pyridoxine/pyridoxamine 5'-phosphate oxid | 0.02482 | 63.7677 | FALSE | FALSE |
| LOC107066644 | protein unzipped | 0.02389 | 63.77462 | FALSE | FALSE |
| LOC107066697 | UBX domain-containing protein 4 | 0.02395 | 63.78914 | FALSE | FALSE |
| LOC107065962 | uncharacterized protein LOC107065962 | 0.02444 | 63.79741 | FALSE | FALSE |
| LOC107070437 | proline-rich nuclear receptor coactivator 2 | 0.02346 | 63.7998 | FALSE | FALSE |
| LOC107070080 | protein tyrosine phosphatase type IVA 1 | 0.02439 | 63.79994 | FALSE | FALSE |
| LOC107071803 | uncharacterized protein LOC107071803 | 0.02532 | 63.82325 | FALSE | FALSE |

| | | | | | |
|---|---|---|---|---|---|
| LOC107067101 | engulfment and cell motility protein 1 | 0.02306 | 63.83678 | FALSE | FALSE |
| LOC107074008 | protein BUD31 homolog | 0.02389 | 63.83881 | FALSE | FALSE |
| Trnas-gcu | Trnas-gcu | 0.02503 | 63.84342 | FALSE | FALSE |
| LOC107064192 | RNA 3'-terminal phosphate cyclase-like prot | 0.02482 | 63.84726 | FALSE | FALSE |
| LOC107070123 | hormonally up-regulated neu tumor-associa | 0.02542 | 63.85739 | FALSE | FALSE |
| LOC107074414 | coiled-coil domain-containing protein 47 | 0.02451 | 63.86317 | FALSE | FALSE |
| LOC107067357 | hydroxysteroid dehydrogenase-like protein | 0.02466 | 63.88403 | FALSE | FALSE |
| LOC107066313 | lipid droplet-associated hydrolase | 0.02428 | 63.92119 | FALSE | FALSE |
| LOC107066750 | MOB kinase activator-like 4 | 0.02259 | 63.92122 | FALSE | FALSE |
| LOC107065538 | uncharacterized protein LOC107065538 | 0.02709 | 63.92921 | FALSE | FALSE |
| LOC107064716 | ell-associated factor Eaf | 0.02299 | 63.93156 | FALSE | FALSE |
| LOC107070337 | gamma-aminobutyric acid receptor subunit | 0.02434 | 63.93565 | FALSE | FALSE |
| LOC107072171 | uncharacterized protein LOC107072171 | 0.02462 | 63.93604 | FALSE | FALSE |
| LOC107072853 | von Hippel-Lindau tumor suppressor homol | 0.02412 | 63.94531 | FALSE | FALSE |
| LOC107073101 | UPF0769 protein C21orf59 homolog | 0.02425 | 63.96961 | FALSE | FALSE |
| LOC107071192 | uncharacterized protein LOC107071192 | 0.02452 | 63.971 | FALSE | FALSE |
| LOC107066442 | homeobox protein B-H1 | 0.02453 | 63.97262 | FALSE | FALSE |
| LOC107070240 | acetylcholinesterase-like | 0.02422 | 63.97718 | FALSE | FALSE |
| LOC107071377 | uncharacterized protein LOC107071377 | 0.02455 | 63.98668 | FALSE | FALSE |
| LOC107068743 | transmembrane emp24 domain-containing | 0.02303 | 63.99224 | FALSE | FALSE |
| LOC107072604 | rRNA-processing protein FCF1 homolog | 0.02344 | 64.02302 | FALSE | FALSE |
| LOC107067410 | electrogenic sodium bicarbonate cotranspo | 0.0245 | 64.03798 | FALSE | FALSE |
| LOC107070737 | 40S ribosomal protein S28-like | 0.02637 | 64.0702 | FALSE | FALSE |
| LOC107065256 | uncharacterized protein LOC107065256 | 0.02517 | 64.07212 | FALSE | FALSE |
| LOC107070638 | mediator of RNA polymerase II transcriptior | 0.02435 | 64.08848 | FALSE | FALSE |
| LOC107069621 | uncharacterized protein LOC107069621 | 0.02282 | 64.10702 | FALSE | FALSE |
| LOC107067739 | citron Rho-interacting kinase-like | 0.02394 | 64.12897 | FALSE | FALSE |
| LOC107071658 | MOB kinase activator-like 1 | 0.02414 | 64.13374 | FALSE | FALSE |
| LOC107064426 | uncharacterized protein PFB0145c | 0.02616 | 64.14248 | FALSE | FALSE |
| LOC107071255 | mediator of RNA polymerase II transcriptior | 0.02402 | 64.15456 | FALSE | FALSE |
| LOC107070055 | hemicentin-2-like | 0.02477 | 64.17577 | FALSE | FALSE |
| LOC107068722 | uncharacterized protein LOC107068722 | 0.02341 | 64.18392 | FALSE | FALSE |
| LOC107067413 | neither inactivation nor afterpotential prote | 0.02313 | 64.20468 | FALSE | FALSE |
| LOC107063906 | NF-X1-type zinc finger protein NFXL1 | 0.02462 | 64.20552 | FALSE | FALSE |
| LOC107066269 | ESF1 homolog | 0.02435 | 64.20567 | FALSE | FALSE |
| LOC107067688 | staphylococcal nuclease domain-containing | 0.02417 | 64.20663 | FALSE | FALSE |
| LOC107068522 | protein ovo-like | 0.0247 | 64.20989 | FALSE | FALSE |
| LOC107063838 | LOW QUALITY PROTEIN: intracellular protei | 0.02314 | 64.24689 | FALSE | FALSE |
| LOC107069387 | uncharacterized protein LOC107069387 | 0.02486 | 64.25074 | FALSE | FALSE |
| LOC107064841 | probable tyrosine-protein phosphatase F54 | 0.02318 | 64.25337 | FALSE | FALSE |
| LOC107073949 | drebrin-like protein B | 0.02495 | 64.28309 | FALSE | FALSE |
| LOC107071354 | paired amphipathic helix protein Sin3a | 0.02351 | 64.28517 | FALSE | FALSE |
| LOC107067496 | G1/S-specific cyclin-E1 | 0.02574 | 64.29 | FALSE | FALSE |
| LOC107071538 | uncharacterized protein LOC107071538 | 0.02446 | 64.29065 | FALSE | FALSE |
| LOC107072891 | zinc finger and SCAN domain-containing pro | 0.02283 | 64.3216 | FALSE | FALSE |
| LOC107070072 | uncharacterized protein LOC107070072 | 0.02518 | 64.32749 | FALSE | FALSE |
| LOC107065360 | uncharacterized protein LOC107065360 | 0.02324 | 64.33486 | FALSE | FALSE |
| LOC107067557 | polycomb protein suz12-B | 0.02413 | 64.33508 | FALSE | FALSE |
| LOC107071911 | multidrug resistance-associated protein 4-lil | 0.02362 | 64.34413 | FALSE | FALSE |
| LOC107073893 | cytochrome c-type heme lyase | 0.02358 | 64.39575 | FALSE | FALSE |

| | | | | | |
|---|---|---|---|---|---|
| LOC107065491 | ras-like GTP-binding protein RhoL | 0.02263 | 64.40281 | FALSE | FALSE |
| LOC107072936 | putative defense protein 3 | 0.0247 | 64.40704 | FALSE | FALSE |
| LOC107070093 | uncharacterized protein LOC107070093 | 0.02439 | 64.43275 | FALSE | FALSE |
| LOC107072443 | exodeoxyribonuclease | 0.02391 | 64.48172 | FALSE | FALSE |
| LOC107069576 | uncharacterized protein LOC107069576 | 0.02615 | 64.50782 | FALSE | FALSE |
| LOC107070132 | UNC93-like protein | 0.02449 | 64.52654 | FALSE | FALSE |
| LOC107066299 | uncharacterized protein DDB_G0290587-lik | 0.02431 | 64.57815 | FALSE | FALSE |
| LOC107065182 | uncharacterized protein LOC107065182 | 0.02545 | 64.60147 | FALSE | FALSE |
| LOC107065856 | GTPase-activating protein | 0.02524 | 64.60572 | FALSE | FALSE |
| LOC107064460 | serine/arginine repetitive matrix protein 1 | 0.02303 | 64.60997 | FALSE | FALSE |
| LOC107070755 | uncharacterized protein LOC107070755 | 0.02365 | 64.63011 | FALSE | FALSE |
| LOC107069626 | uncharacterized protein LOC107069626 | 0.02359 | 64.63169 | FALSE | FALSE |
| LOC107064402 | rhoGEF domain-containing protein gxcJ | 0.02317 | 64.66229 | FALSE | FALSE |
| LOC107065415 | uncharacterized protein LOC107065415 | 0.02409 | 64.66553 | FALSE | FALSE |
| LOC107064418 | probable peptide chain release factor C12or | 0.0246 | 64.67251 | FALSE | FALSE |
| LOC107066253 | mitochondrial import receptor subunit TOM | 0.02266 | 64.68091 | FALSE | FALSE |
| LOC107070968 | zinc finger protein 510-like | 0.02433 | 64.68367 | FALSE | FALSE |
| LOC107064326 | POU domain, class 4, transcription factor 1 | 0.0249 | 64.70724 | FALSE | FALSE |
| LOC107068121 | uncharacterized protein LOC107068121 | 0.02423 | 64.73026 | FALSE | FALSE |
| LOC107074502 | eukaryotic peptide chain release factor GTP | 0.02415 | 64.73236 | FALSE | FALSE |
| LOC107068632 | uncharacterized protein LOC107068632 | 0.02455 | 64.7428 | FALSE | FALSE |
| LOC107067296 | G-protein coupled receptor Mth2-like | 0.02439 | 64.74936 | FALSE | FALSE |
| LOC107072878 | uncharacterized protein LOC107072878 | 0.02497 | 64.75799 | FALSE | FALSE |
| LOC107065201 | mesencephalic astrocyte-derived neurotrop | 0.02428 | 64.78838 | FALSE | FALSE |
| LOC107070467 | uncharacterized protein LOC107070467 | 0.02443 | 64.80133 | FALSE | FALSE |
| LOC107066037 | enolase-phosphatase E1-like | 0.02566 | 64.80322 | FALSE | FALSE |
| LOC107067931 | tubulin alpha-1 chain-like | 0.02408 | 64.8052 | FALSE | FALSE |
| LOC107068262 | E3 ubiquitin-protein ligase SMURF1 | 0.02448 | 64.80819 | FALSE | FALSE |
| LOC107066467 | uncharacterized protein LOC107066467 | 0.02461 | 64.82017 | FALSE | FALSE |
| Trnap-cgg | Trnap-cgg | 0.02391 | 64.8219 | FALSE | FALSE |
| LOC107068976 | condensin-2 complex subunit D3-like | 0.02421 | 64.82464 | FALSE | FALSE |
| LOC107067609 | protein argonaute-2-like | 0.02477 | 64.8311 | FALSE | FALSE |
| LOC107070918 | LOW QUALITY PROTEIN: eukaryotic initiatio | 0.02444 | 64.85558 | FALSE | FALSE |
| LOC107072915 | uncharacterized protein LOC107072915 | 0.02315 | 64.85942 | FALSE | FALSE |
| LOC107068037 | uncharacterized protein LOC107068037 | 0.02384 | 64.8792 | FALSE | FALSE |
| LOC107065234 | glycine cleavage system H protein, mitochor | 0.02345 | 64.88843 | FALSE | FALSE |
| LOC107065774 | uncharacterized protein LOC107065774 | 0.02514 | 64.88881 | FALSE | FALSE |
| LOC107063898 | glutamic acid-rich protein | 0.02411 | 64.89061 | FALSE | FALSE |
| LOC107063788 | serine/threonine-protein kinase BRSK2 | 0.02488 | 64.89725 | FALSE | FALSE |
| LOC107066245 | transcription initiation factor TFIID subunit 2 | 0.02306 | 64.90865 | FALSE | FALSE |
| LOC107071296 | m7GpppX diphosphatase | 0.02366 | 64.91053 | FALSE | FALSE |
| LOC107063998 | ER membrane protein complex subunit 4 | 0.02379 | 64.92133 | FALSE | FALSE |
| LOC107071681 | aminopeptidase N-like | 0.02545 | 64.92579 | FALSE | FALSE |
| LOC107070809 | E3 ubiquitin-protein ligase TRIM23-like | 0.02474 | 64.92621 | FALSE | FALSE |
| LOC107065697 | ras-related protein Rab-10 | 0.02599 | 64.93128 | FALSE | FALSE |
| LOC107072701 | DNA ligase 3 | 0.02455 | 64.93141 | FALSE | FALSE |
| LOC107072145 | mediator of RNA polymerase II transcriptior | 0.02444 | 64.93318 | FALSE | FALSE |
| LOC107073931 | U6 snRNA-associated Sm-like protein LSm2 | 0.02432 | 64.94906 | FALSE | FALSE |
| LOC107070764 | uncharacterized protein LOC107070764 | 0.02402 | 64.95771 | FALSE | FALSE |
| LOC107068772 | protein charlatan | 0.02459 | 64.96429 | FALSE | FALSE |

| | | | | | |
|---|---|---|---|---|---|
| LOC107071139 | paramyosin, long form-like | 0.02294 | 64.96727 | FALSE | FALSE |
| LOC107064147 | protein Jumonji | 0.02294 | 64.96844 | FALSE | FALSE |
| LOC107066122 | putative elongator complex protein 1 | 0.02273 | 64.97354 | FALSE | FALSE |
| LOC107065726 | E3 ubiquitin-protein ligase RNF185-like | 0.02301 | 64.97392 | FALSE | FALSE |
| LOC107069264 | tumor protein p53-inducible nuclear protei | 0.02273 | 64.98853 | FALSE | FALSE |
| LOC107070228 | PIH1 domain-containing protein 1 | 0.02508 | 64.98941 | FALSE | FALSE |
| LOC107066912 | uncharacterized protein LOC107066912 | 0.02409 | 64.99034 | FALSE | FALSE |
| LOC107071243 | solute carrier family 25 member 38-like | 0.02434 | 65.003 | FALSE | FALSE |
| LOC107066563 | uncharacterized protein LOC107066563 | 0.02277 | 65.01685 | FALSE | FALSE |
| LOC107069192 | neurexin-4 | 0.02311 | 65.03971 | FALSE | FALSE |
| LOC107066567 | serine/arginine-rich splicing factor 7-like | 0.02493 | 65.06039 | FALSE | FALSE |
| LOC107073852 | uncharacterized protein LOC107073852 | 0.0226 | 65.08079 | FALSE | FALSE |
| LOC107070186 | tether containing UBX domain for GLUT4 | 0.02461 | 65.12242 | FALSE | FALSE |
| LOC107070175 | aryl hydrocarbon receptor protein 1 | 0.0246 | 65.12477 | FALSE | FALSE |
| LOC107065030 | B-cell lymphoma/leukemia 11B | 0.02523 | 65.13266 | FALSE | FALSE |
| LOC107067529 | zinc finger X-chromosomal protein-like | 0.02371 | 65.13759 | FALSE | FALSE |
| LOC107070722 | ATP-dependent RNA helicase dbp2-like | 0.02326 | 65.13982 | FALSE | FALSE |
| LOC107070360 | cell wall protein IFF6-like | 0.02408 | 65.14879 | FALSE | FALSE |
| LOC107069543 | zinc finger and BTB domain-containing prot | 0.02406 | 65.15376 | FALSE | FALSE |
| LOC107063734 | uncharacterized protein LOC107063734 | 0.02404 | 65.15518 | FALSE | FALSE |
| LOC107066338 | cellular tumor antigen p53 | 0.02419 | 65.15937 | FALSE | FALSE |
| LOC107069487 | uncharacterized protein LOC107069487 | 0.02405 | 65.18207 | FALSE | FALSE |
| LOC107067172 | putative RNA-binding protein Luc7-like 1 | 0.02445 | 65.18811 | FALSE | FALSE |
| LOC107067423 | tankyrase | 0.02395 | 65.20376 | FALSE | FALSE |
| LOC107067661 | uncharacterized protein LOC107067661 | 0.02442 | 65.21667 | FALSE | FALSE |
| LOC107069448 | tubulin-specific chaperone cofactor E-like p | 0.02349 | 65.23904 | FALSE | FALSE |
| LOC107064630 | N-alpha-acetyltransferase 25, NatB auxiliary | 0.02365 | 65.28466 | FALSE | FALSE |
| LOC107065311 | uridine phosphorylase 1 | 0.02425 | 65.29252 | FALSE | FALSE |
| LOC107067242 | G2/mitotic-specific cyclin-A | 0.02386 | 65.30674 | FALSE | FALSE |
| LOC107068063 | aarF domain-containing protein kinase 4 | 0.0243 | 65.30709 | FALSE | FALSE |
| LOC107074456 | peroxisome biogenesis factor 1 | 0.02431 | 65.3126 | FALSE | FALSE |
| LOC107064961 | glutamyl-tRNA(Gln) amidotransferase subu | 0.02352 | 65.32151 | FALSE | FALSE |
| LOC107073597 | uncharacterized protein LOC107073597 | 0.02311 | 65.33947 | FALSE | FALSE |
| LOC107063948 | putative mediator of RNA polymerase II tra | 0.02344 | 65.37681 | FALSE | FALSE |
| LOC107066917 | CD63 antigen-like | 0.02373 | 65.37937 | FALSE | FALSE |
| LOC107069676 | uncharacterized protein LOC107069676 | 0.02494 | 65.40928 | FALSE | FALSE |
| LOC107072710 | uncharacterized protein LOC107072710 | 0.02447 | 65.41001 | FALSE | FALSE |
| LOC107071796 | intraflagellar transport protein 81 homolog | 0.02375 | 65.42525 | FALSE | FALSE |
| LOC107072151 | cleavage stimulation factor subunit 2 tau va | 0.02402 | 65.43239 | FALSE | FALSE |
| LOC107073105 | vacuolar protein sorting-associated protein | 0.0239 | 65.43588 | FALSE | FALSE |
| LOC107071082 | uncharacterized protein LOC107071082 | 0.02418 | 65.45315 | FALSE | FALSE |
| LOC107074334 | uncharacterized protein LOC107074334 | 0.02384 | 65.45649 | FALSE | FALSE |
| LOC107069312 | uncharacterized protein LOC107069312 | 0.02346 | 65.45937 | FALSE | FALSE |
| LOC107073835 | ras-related protein Rab-34 | 0.02456 | 65.46266 | FALSE | FALSE |
| LOC107072822 | transcription initiation factor TFIID subunit | 0.02381 | 65.4686 | FALSE | FALSE |
| LOC107064416 | uncharacterized protein LOC107064416 | 0.02659 | 65.47559 | FALSE | FALSE |
| LOC107066967 | melanoma-associated antigen G1-like | 0.02342 | 65.47658 | FALSE | FALSE |
| LOC107066944 | intraflagellar transport protein 74 homolog | 0.0237 | 65.48108 | FALSE | FALSE |
| LOC107067669 | uncharacterized protein LOC107067669 | 0.02468 | 65.48408 | FALSE | FALSE |
| LOC107064694 | TBC1 domain family member 22B | 0.02233 | 65.49226 | FALSE | FALSE |

| | | | | | |
|---|---|---|---|---|---|
| LOC107065338 | thiamine transporter 1-like | 0.02493 | 65.4948 | FALSE | FALSE |
| LOC107064954 | uncharacterized protein LOC107064954 | 0.0253 | 65.49671 | FALSE | FALSE |
| LOC107066420 | protein flightless-1 | 0.02441 | 65.515 | FALSE | FALSE |
| LOC107073644 | histone-lysine N-methyltransferase SETMAF | 0.02275 | 65.53137 | FALSE | FALSE |
| LOC107072223 | uncharacterized protein LOC107072223 | 0.02483 | 65.59995 | FALSE | FALSE |
| LOC107068236 | tyrosine-protein phosphatase non-receptor | 0.02421 | 65.624 | FALSE | FALSE |
| LOC107072257 | UPF0488 protein CG14286 | 0.02284 | 65.63558 | FALSE | FALSE |
| LOC107070582 | uncharacterized protein LOC107070582 | 0.02468 | 65.65838 | FALSE | FALSE |
| LOC107068991 | E3 SUMO-protein ligase PIAS3 | 0.02319 | 65.66466 | FALSE | FALSE |
| LOC107064985 | dihydropyrimidinase | 0.02376 | 65.67712 | FALSE | FALSE |
| LOC107068645 | LOW QUALITY PROTEIN: putative tRNA pseu | 0.02417 | 65.67833 | FALSE | FALSE |
| LOC107066455 | UPF0160 protein | 0.02386 | 65.6808 | FALSE | FALSE |
| LOC107064884 | uncharacterized protein LOC107064884 | 0.02457 | 65.68339 | FALSE | FALSE |
| LOC107073353 | mediator of RNA polymerase II transcriptior | 0.02436 | 65.68656 | FALSE | FALSE |
| LOC107067155 | 4-coumarate--CoA ligase 1-like | 0.02371 | 65.69432 | FALSE | FALSE |
| LOC107066157 | peptide methionine sulfoxide reductase | 0.02503 | 65.70248 | FALSE | FALSE |
| LOC107069221 | serine--tRNA ligase, mitochondrial | 0.02283 | 65.71659 | FALSE | FALSE |
| LOC107070752 | radial spoke head protein 9 homolog | 0.02338 | 65.77357 | FALSE | FALSE |
| LOC107074628 | GSK3-beta interaction protein | 0.02244 | 65.78641 | FALSE | FALSE |
| LOC107065514 | sulfotransferase 4A1-like | 0.02396 | 65.81756 | FALSE | FALSE |
| LOC107064677 | fasciclin-3 | 0.02433 | 65.82461 | FALSE | FALSE |
| LOC107064120 | dnaJ homolog subfamily C member 13 | 0.02388 | 65.82697 | FALSE | FALSE |
| LOC107073469 | uncharacterized protein LOC107073469 | 0.02493 | 65.83433 | FALSE | FALSE |
| LOC107068636 | CTD nuclear envelope phosphatase 1 homol | 0.02391 | 65.83675 | FALSE | FALSE |
| LOC107070875 | uncharacterized protein LOC107070875 | 0.02429 | 65.85702 | FALSE | FALSE |
| LOC107064129 | uncharacterized protein LOC107064129 | 0.02426 | 65.90555 | FALSE | FALSE |
| LOC107065227 | heterogeneous nuclear ribonucleoprotein R | 0.02355 | 65.90958 | FALSE | FALSE |
| LOC107064064 | sorting nexin-27 | 0.02413 | 65.91264 | FALSE | FALSE |
| LOC107069760 | transmembrane emp24 domain-containing | 0.02373 | 65.91422 | FALSE | FALSE |
| LOC107070207 | anther-specific proline-rich protein APG-like | 0.02351 | 65.91466 | FALSE | FALSE |
| LOC107069729 | uncharacterized protein LOC107069729 | 0.02403 | 65.93124 | FALSE | FALSE |
| LOC107073062 | NIF3-like protein 1 | 0.02346 | 65.94592 | FALSE | FALSE |
| LOC107063812 | synaptogyrin-1 | 0.02396 | 65.95037 | FALSE | FALSE |
| LOC107070195 | prenylcysteine oxidase-like | 0.02363 | 65.95898 | FALSE | FALSE |
| LOC107072560 | active regulator of SIRT1-like | 0.0252 | 65.9591 | FALSE | FALSE |
| LOC107072748 | uncharacterized protein LOC107072748 | 0.02333 | 65.96352 | FALSE | FALSE |
| LOC107067800 | protein NPC2 homolog | 0.02295 | 65.96383 | FALSE | FALSE |
| LOC107071554 | nicotinate phosphoribosyltransferase | 0.02462 | 65.96656 | FALSE | FALSE |
| LOC107069234 | 60S ribosomal protein L31 | 0.02488 | 65.98471 | FALSE | FALSE |
| LOC107074446 | spindle assembly abnormal protein 6 homol | 0.02284 | 65.98574 | FALSE | FALSE |
| LOC107063816 | 60S ribosomal protein L8 | 0.02311 | 65.99798 | FALSE | FALSE |
| LOC107063867 | transcription initiation factor TFIID subunit 1 | 0.02249 | 66.0228 | FALSE | FALSE |
| LOC107069430 | uncharacterized protein LOC107069430 | 0.02386 | 66.02727 | FALSE | FALSE |
| LOC107067074 | calpain-7-like | 0.02493 | 66.02866 | FALSE | FALSE |
| LOC107065042 | NSFL1 cofactor p47 | 0.02382 | 66.0421 | FALSE | FALSE |
| LOC107065657 | E3 ubiquitin-protein ligase RNF25 | 0.02363 | 66.10258 | FALSE | FALSE |
| LOC107071239 | uncharacterized protein LOC107071239 | 0.02344 | 66.11523 | FALSE | FALSE |
| LOC107072595 | striatin-3 | 0.0221 | 66.12731 | FALSE | FALSE |
| LOC107067129 | abnormal long morphology protein 1-like | 0.02641 | 66.13322 | FALSE | FALSE |
| LOC107071194 | zinc finger protein 236-like | 0.02266 | 66.1863 | FALSE | FALSE |

| | | | | | |
|---|---|---|---|---|---|
| LOC107067773 | ubiquitin | 0.0236 | 66.1941 | FALSE | FALSE |
| LOC107073467 | RING finger protein unkempt | 0.02318 | 66.19561 | FALSE | FALSE |
| LOC107071322 | CDK5 and ABL1 enzyme substrate 1 | 0.02281 | 66.23982 | FALSE | FALSE |
| LOC107073719 | protein toll-like | 0.02427 | 66.24221 | FALSE | FALSE |
| LOC107070564 | prostatic spermine-binding protein-like | 0.02453 | 66.25577 | FALSE | FALSE |
| LOC107069170 | cation-independent mannose-6-phosphate | 0.02463 | 66.27575 | FALSE | FALSE |
| LOC107072227 | U3 small nucleolar ribonucleoprotein protei | 0.02306 | 66.27642 | FALSE | FALSE |
| LOC107070784 | MATH and LRR domain-containing protein P | 0.02406 | 66.27793 | FALSE | FALSE |
| LOC107074647 | protein YIF1B | 0.02453 | 66.29527 | FALSE | FALSE |
| LOC107066034 | isoaspartyl peptidase/L-asparaginase-like | 0.0228 | 66.29915 | FALSE | FALSE |
| LOC107073847 | coiled-coil domain-containing protein 130 h | 0.02389 | 66.30066 | FALSE | FALSE |
| LOC107066364 | uncharacterized protein LOC107066364 | 0.02294 | 66.31585 | FALSE | FALSE |
| LOC107067344 | chromatin-remodeling complex ATPase chai | 0.02262 | 66.32174 | FALSE | FALSE |
| LOC107069258 | protein YIPF6 | 0.02284 | 66.33509 | FALSE | FALSE |
| LOC107074122 | mothers against decapentaplegic homolog 3 | 0.02433 | 66.34019 | FALSE | FALSE |
| LOC107068674 | putative protein FAM10A4 | 0.02332 | 66.37816 | FALSE | FALSE |
| LOC107069783 | thioredoxin-2 | 0.02199 | 66.38689 | FALSE | FALSE |
| LOC107074004 | bifunctional transcriptional activator/DNA r | 0.02341 | 66.40534 | FALSE | FALSE |
| LOC107068656 | tRNA (adenine(58)-N(1))-methyltransferase | 0.02331 | 66.40785 | FALSE | FALSE |
| LOC107064252 | peroxisomal biogenesis factor 19 | 0.02374 | 66.44436 | FALSE | FALSE |
| LOC107074016 | protein MEF2BNB homolog | 0.02294 | 66.44467 | FALSE | FALSE |
| LOC107074705 | THAP domain-containing protein 2-like | 0.02405 | 66.44875 | FALSE | FALSE |
| LOC107067494 | protein phosphatase 1 regulatory subunit 3I | 0.02387 | 66.45083 | FALSE | FALSE |
| LOC107068621 | leucine-rich repeat-containing protein 24-lik | 0.02425 | 66.46673 | FALSE | FALSE |
| LOC107070521 | putative tricarboxylate transport protein, m | 0.02328 | 66.46816 | FALSE | FALSE |
| LOC107064122 | uncharacterized protein LOC107064122 | 0.02373 | 66.47045 | FALSE | FALSE |
| LOC107069160 | uncharacterized protein LOC107069160 | 0.02374 | 66.47488 | FALSE | FALSE |
| LOC107070221 | N-alpha-acetyltransferase 38-B, NatC auxilia | 0.02401 | 66.4788 | FALSE | FALSE |
| LOC107066539 | uncharacterized protein LOC107066539 | 0.02363 | 66.49583 | FALSE | FALSE |
| LOC107070745 | cap-specific mRNA (nucleoside-2'-O-)-methy | 0.02372 | 66.4995 | FALSE | FALSE |
| LOC107067069 | uncharacterized protein LOC107067069 | 0.02334 | 66.50195 | FALSE | FALSE |
| LOC107069968 | uncharacterized protein LOC107069968 | 0.0225 | 66.50723 | FALSE | FALSE |
| LOC107069050 | uncharacterized protein LOC107069050 | 0.02407 | 66.50964 | FALSE | FALSE |
| LOC107065911 | chymotrypsin-2-like | 0.02392 | 66.51289 | FALSE | FALSE |
| LOC107065693 | cirhin | 0.02301 | 66.56829 | FALSE | FALSE |
| LOC107070609 | uncharacterized protein LOC107070609 | 0.02243 | 66.58604 | FALSE | FALSE |
| LOC107072823 | mediator of RNA polymerase II transcriptior | 0.02281 | 66.59813 | FALSE | FALSE |
| LOC107073443 | probable 2-oxoglutarate dehydrogenase E1 | 0.02361 | 66.59971 | FALSE | FALSE |
| LOC107065969 | RING finger protein 44-like | 0.02453 | 66.60396 | FALSE | FALSE |
| LOC107067923 | WD repeat-containing protein 59 | 0.02278 | 66.6101 | FALSE | FALSE |
| LOC107073464 | nose resistant to fluoxetine protein 6-like | 0.02379 | 66.62224 | FALSE | FALSE |
| LOC107068296 | splicing factor U2af 38 kDa subunit-like | 0.02479 | 66.62709 | FALSE | FALSE |
| LOC107065334 | probable Xaa-Pro aminopeptidase 3 | 0.02468 | 66.64166 | FALSE | FALSE |
| LOC107068711 | transcription factor hamlet-like | 0.02296 | 66.64705 | FALSE | FALSE |
| LOC107068269 | alpha-tocopherol transfer protein-like | 0.02278 | 66.64732 | FALSE | FALSE |
| LOC107065796 | uncharacterized protein LOC107065796 | 0.02449 | 66.64888 | FALSE | FALSE |
| LOC107064574 | uncharacterized protein LOC107064574 | 0.02487 | 66.66539 | FALSE | FALSE |
| LOC107071298 | ATP-binding cassette sub-family B member | 0.02319 | 66.67973 | FALSE | FALSE |
| LOC107070497 | fatty acid 2-hydroxylase | 0.02455 | 66.71012 | FALSE | FALSE |
| LOC107065874 | myotubularin-related protein 10-B | 0.02474 | 66.72656 | FALSE | FALSE |

| | | | | | |
|---|---|---|---|---|---|
| LOC107073418 | piggyBac transposable element-derived pro | 0.0229 | 66.72693 | FALSE | FALSE |
| LOC107065692 | uncharacterized protein LOC107065692 | 0.02363 | 66.7496 | FALSE | FALSE |
| LOC107068575 | NADH dehydrogenase [ubiquinone | 0.02488 | 66.78229 | FALSE | FALSE |
| LOC107072628 | galactoside 2-alpha-L-fucosyltransferase 3-li | 0.02446 | 66.81686 | FALSE | FALSE |
| LOC107072360 | uncharacterized protein LOC107072360 | 0.0228 | 66.82679 | FALSE | FALSE |
| LOC107068334 | probable low-specificity L-threonine aldolas | 0.0239 | 66.83018 | FALSE | FALSE |
| LOC107065712 | splicing factor 3B subunit 6-like protein | 0.02396 | 66.83321 | FALSE | FALSE |
| LOC107065973 | uncharacterized protein LOC107065973 | 0.02332 | 66.84579 | FALSE | FALSE |
| LOC107068556 | odorant receptor Or1-like | 0.02327 | 66.84959 | FALSE | FALSE |
| LOC107071556 | endochitinase | 0.02495 | 66.85523 | FALSE | FALSE |
| LOC107067597 | sphingosine-1-phosphate lyase | 0.02372 | 66.85741 | FALSE | FALSE |
| LOC107072246 | phenylalanine--tRNA ligase alpha subunit | 0.02274 | 66.8775 | FALSE | FALSE |
| LOC107072037 | calpain-A | 0.02404 | 66.89218 | FALSE | FALSE |
| LOC107067368 | uncharacterized protein LOC107067368 | 0.02405 | 66.89566 | FALSE | FALSE |
| LOC107067900 | cytochrome P450 4C1-like | 0.0234 | 66.91279 | FALSE | FALSE |
| LOC107073705 | uncharacterized protein LOC107073705 | 0.02441 | 66.92916 | FALSE | FALSE |
| LOC107071734 | splicing factor 3A subunit 3 | 0.02342 | 66.95718 | FALSE | FALSE |
| LOC107071264 | uncharacterized protein LOC107071264 | 0.02211 | 66.96018 | FALSE | FALSE |
| LOC107064263 | girdin-like | 0.02371 | 66.96942 | FALSE | FALSE |
| LOC107064947 | uncharacterized protein LOC107064947 | 0.02389 | 67.02061 | FALSE | FALSE |
| LOC107074562 | unconventional myosin-IXa-like | 0.02448 | 67.02381 | FALSE | FALSE |
| LOC107072625 | polycomb protein Sfmbt-like | 0.02444 | 67.02819 | FALSE | FALSE |
| LOC107067462 | SAFB-like transcription modulator | 0.02218 | 67.04343 | FALSE | FALSE |
| LOC107064873 | uncharacterized protein LOC107064873 | 0.02356 | 67.05094 | FALSE | FALSE |
| LOC107074143 | uncharacterized protein C9orf85 homolog | 0.02387 | 67.06293 | FALSE | FALSE |
| LOC107066359 | DNA-directed RNA polymerase II subunit RP | 0.02316 | 67.06539 | FALSE | FALSE |
| LOC107066200 | protein Hook homolog 3 | 0.02245 | 67.07142 | FALSE | FALSE |
| LOC107071596 | eukaryotic translation initiation factor 4B | 0.02231 | 67.07449 | FALSE | FALSE |
| LOC107065432 | small ubiquitin-related modifier 3 | 0.02307 | 67.07733 | FALSE | FALSE |
| LOC107063668 | uncharacterized protein LOC107063668 | 0.0247 | 67.09227 | FALSE | FALSE |
| LOC107068716 | pre-mRNA-processing factor 17 | 0.02366 | 67.12223 | FALSE | FALSE |
| LOC107071580 | uncharacterized protein LOC107071580 | 0.02415 | 67.17601 | FALSE | FALSE |
| LOC107069436 | tyrosine-protein kinase transmembrane rec | 0.02305 | 67.19044 | FALSE | FALSE |
| LOC107072424 | uncharacterized protein LOC107072424 | 0.02427 | 67.19073 | FALSE | FALSE |
| LOC107069320 | WD repeat-containing protein 47 | 0.02401 | 67.19782 | FALSE | FALSE |
| LOC107065719 | uncharacterized protein LOC107065719 | 0.02315 | 67.19868 | FALSE | FALSE |
| LOC107069836 | mitochondrial import receptor subunit TOM | 0.02228 | 67.21733 | FALSE | FALSE |
| LOC107069167 | protein toll | 0.02356 | 67.22346 | FALSE | FALSE |
| LOC107068345 | uncharacterized protein LOC107068345 | 0.0249 | 67.23677 | FALSE | FALSE |
| LOC107074583 | uncharacterized protein LOC107074583 | 0.02634 | 67.2603 | FALSE | FALSE |
| LOC107066084 | abscission/NoCut checkpoint regulator | 0.02492 | 67.26264 | FALSE | FALSE |
| LOC107064983 | tetratricopeptide repeat protein 37 | 0.02327 | 67.27196 | FALSE | FALSE |
| LOC107072329 | KRAB-A domain-containing protein 2-like | 0.02457 | 67.27593 | FALSE | FALSE |
| LOC107066773 | zinc finger protein 585A | 0.02154 | 67.30162 | FALSE | FALSE |
| LOC107073466 | meiosis arrest female protein 1 homolog | 0.02322 | 67.30933 | FALSE | FALSE |
| LOC107071418 | phospholipase DDHD1-like | 0.02342 | 67.31678 | FALSE | FALSE |
| LOC107066068 | RNA polymerase II subunit A C-terminal don | 0.02363 | 67.34784 | FALSE | FALSE |
| LOC107071460 | fatty acid synthase-like | 0.02312 | 67.3533 | FALSE | FALSE |
| LOC107064744 | uncharacterized protein LOC107064744 | 0.0238 | 67.35685 | FALSE | FALSE |
| LOC107072922 | succinate dehydrogenase assembly factor 4, | 0.02429 | 67.37403 | FALSE | FALSE |

| | | | | | |
|---|---|---|---|---|---|
| LOC107066608 | caspase-1-like | 0.02383 | 67.37667 | FALSE | FALSE |
| LOC107063851 | uncharacterized protein LOC107063851 | 0.02293 | 67.37962 | FALSE | FALSE |
| LOC107067039 | uncharacterized protein LOC107067039 | 0.02421 | 67.38377 | FALSE | FALSE |
| LOC107064682 | DCN1-like protein 1 | 0.02385 | 67.38593 | FALSE | FALSE |
| LOC107065258 | serine/threonine-protein phosphatase 2A ca | 0.0232 | 67.38905 | FALSE | FALSE |
| LOC107068240 | protein Peter pan | 0.02307 | 67.39557 | FALSE | FALSE |
| LOC107072331 | probable arginine--tRNA ligase, mitochondri | 0.02368 | 67.41482 | FALSE | FALSE |
| LOC107071646 | uncharacterized protein LOC107071646 | 0.02456 | 67.41493 | FALSE | FALSE |
| LOC107067239 | proteasome subunit alpha type-4 | 0.02475 | 67.42607 | FALSE | FALSE |
| LOC107072282 | heterogeneous nuclear ribonucleoprotein A | 0.02288 | 67.43292 | FALSE | FALSE |
| LOC107072545 | 1-phosphatidylinositol 3-phosphate 5-kinase | 0.02244 | 67.46137 | FALSE | FALSE |
| LOC107069314 | isochorismatase domain-containing protein | 0.02419 | 67.47429 | FALSE | FALSE |
| LOC107070129 | guanine nucleotide-binding protein G(o) sub | 0.02412 | 67.49778 | FALSE | FALSE |
| LOC107066345 | uncharacterized protein LOC107066345 | 0.02239 | 67.51521 | FALSE | FALSE |
| LOC107064229 | rhoGEF domain-containing protein gxcJ-like | 0.02195 | 67.52253 | FALSE | FALSE |
| LOC107072958 | short coiled-coil protein B | 0.02301 | 67.53968 | FALSE | FALSE |
| LOC107072786 | cyclin N-terminal domain-containing protein | 0.02312 | 67.54735 | FALSE | FALSE |
| LOC107073492 | relaxin receptor 1 | 0.02473 | 67.55622 | FALSE | FALSE |
| LOC107072541 | glutaminase liver isoform, mitochondrial | 0.02362 | 67.57129 | FALSE | FALSE |
| LOC107065281 | protein BTG2-like | 0.02352 | 67.58929 | FALSE | FALSE |
| LOC107068767 | progestin and adipoQ receptor family meml | 0.02296 | 67.60997 | FALSE | FALSE |
| LOC107074440 | uncharacterized protein LOC107074440 | 0.02314 | 67.6192 | FALSE | FALSE |
| LOC107071683 | acetylcholine receptor subunit alpha-like | 0.02255 | 67.62365 | FALSE | FALSE |
| LOC107071703 | 60S ribosomal protein L8 | 0.02278 | 67.62819 | FALSE | FALSE |
| LOC107073456 | rRNA-processing protein UTP23 homolog | 0.02274 | 67.62899 | FALSE | FALSE |
| LOC107072194 | Kv channel-interacting protein 1 | 0.025 | 67.64422 | FALSE | FALSE |
| LOC107073356 | ATP-dependent zinc metalloprotease YME1 | 0.02195 | 67.66268 | FALSE | FALSE |
| LOC107065706 | deoxynucleotidyltransferase terminal-intera | 0.02296 | 67.68308 | FALSE | FALSE |
| LOC107068586 | protein Fer3 | 0.02411 | 67.72178 | FALSE | FALSE |
| LOC107069266 | phosphatidylserine decarboxylase proenzym | 0.02373 | 67.74982 | FALSE | FALSE |
| LOC107074332 | tRNA methyltransferase 10 homolog A | 0.02363 | 67.75952 | FALSE | FALSE |
| LOC107072607 | serine hydrolase-like protein | 0.02427 | 67.75989 | FALSE | FALSE |
| LOC107066199 | leucine-rich repeat-containing protein 40-lik | 0.02252 | 67.76098 | FALSE | FALSE |
| LOC107066594 | uncharacterized protein LOC107066594 | 0.02382 | 67.76452 | FALSE | FALSE |
| LOC107071733 | uridine 5'-monophosphate synthase | 0.02265 | 67.76605 | FALSE | FALSE |
| LOC107073158 | succinate dehydrogenase assembly factor 2, | 0.02408 | 67.7716 | FALSE | FALSE |
| LOC107064398 | uncharacterized protein LOC107064398 | 0.02337 | 67.78735 | FALSE | FALSE |
| LOC107065377 | uncharacterized protein LOC107065377 | 0.02185 | 67.79089 | FALSE | FALSE |
| LOC107067466 | serine protease snake-like | 0.02373 | 67.80703 | FALSE | FALSE |
| LOC107063882 | exocyst complex component 4 | 0.02326 | 67.80761 | FALSE | FALSE |
| LOC107074621 | insulin-like growth factor-binding protein cc | 0.02341 | 67.81779 | FALSE | FALSE |
| LOC107066400 | angiotensin-converting enzyme-like | 0.02367 | 67.82262 | FALSE | FALSE |
| LOC107064928 | uncharacterized protein LOC107064928 | 0.02406 | 67.83248 | FALSE | FALSE |
| LOC107065012 | calcium homeostasis endoplasmic reticulum | 0.02362 | 67.83753 | FALSE | FALSE |
| LOC107070439 | astakine-like | 0.0237 | 67.85786 | FALSE | FALSE |
| LOC107064996 | CAAX prenyl protease 2 | 0.02449 | 67.86224 | FALSE | FALSE |
| LOC107064512 | uncharacterized protein LOC107064512 | 0.02553 | 67.88327 | FALSE | FALSE |
| LOC107071016 | CKLF-like MARVEL transmembrane domain- | 0.02478 | 67.88682 | FALSE | FALSE |
| LOC107073028 | uncharacterized protein LOC107073028 | 0.0226 | 67.9045 | FALSE | FALSE |
| LOC107065081 | uncharacterized protein LOC107065081 | 0.02229 | 67.90866 | FALSE | FALSE |

| | | | | | |
|---|---|---|---|---|---|
| LOC107072799 | cell division cycle protein 20 homolog | 0.02233 | 67.92929 | FALSE | FALSE |
| LOC107073819 | pyrroline-5-carboxylate reductase 1, mitoch | 0.02371 | 67.93815 | FALSE | FALSE |
| LOC107063716 | oxidation resistance protein 1 | 0.02379 | 67.93952 | FALSE | FALSE |
| LOC107073249 | alpha-soluble NSF attachment protein | 0.02398 | 67.94347 | FALSE | FALSE |
| LOC107067209 | facilitated trehalose transporter Tret1-like | 0.0228 | 67.96204 | FALSE | FALSE |
| LOC107073248 | glomulin-like | 0.02327 | 67.98663 | FALSE | FALSE |
| LOC107066487 | uncharacterized protein LOC107066487 | 0.02259 | 67.99767 | FALSE | FALSE |
| LOC107067646 | histone-lysine N-methyltransferase Suv4-20 | 0.02254 | 68.00231 | FALSE | FALSE |
| LOC107065861 | RNA-binding protein 42 | 0.02329 | 68.00517 | FALSE | FALSE |
| LOC107073549 | phosducin-like protein | 0.0217 | 68.01851 | FALSE | FALSE |
| LOC107069322 | vesicle transport protein GOT1B | 0.02349 | 68.02046 | FALSE | FALSE |
| LOC107069429 | gephyrin | 0.02533 | 68.02151 | FALSE | FALSE |
| LOC107063665 | uncharacterized protein LOC107063665 | 0.02386 | 68.03162 | FALSE | FALSE |
| LOC107072767 | putative RNA polymerase II subunit B1 CTD | 0.02209 | 68.03219 | FALSE | FALSE |
| LOC107074267 | zinc finger protein 714-like | 0.02376 | 68.05471 | FALSE | FALSE |
| LOC107066038 | serine/threonine-protein kinase S6KL-like | 0.02253 | 68.05559 | FALSE | FALSE |
| LOC107064071 | facilitated trehalose transporter Tret1-like | 0.02412 | 68.05758 | FALSE | FALSE |
| LOC107066505 | aladin | 0.02322 | 68.05939 | FALSE | FALSE |
| LOC107072460 | uncharacterized protein LOC107072460 | 0.02295 | 68.08824 | FALSE | FALSE |
| LOC107067528 | zinc finger protein 263-like | 0.02367 | 68.10803 | FALSE | FALSE |
| LOC107071544 | endoplasmin | 0.02264 | 68.11653 | FALSE | FALSE |
| LOC107071286 | DNA repair protein XRCC3-like | 0.02198 | 68.11872 | FALSE | FALSE |
| LOC107066663 | pre-rRNA-processing protein TSR1 homolog | 0.02236 | 68.1473 | FALSE | FALSE |
| LOC107067343 | homeobox protein SIX2-like | 0.02497 | 68.17293 | FALSE | FALSE |
| LOC107070013 | dynein heavy chain 5, axonemal | 0.02512 | 68.18513 | FALSE | FALSE |
| LOC107068052 | guanine nucleotide-binding protein G(s) sub | 0.02236 | 68.19415 | FALSE | FALSE |
| LOC107070506 | C2 domain-containing protein 3-like | 0.02213 | 68.19519 | FALSE | FALSE |
| LOC107067607 | glutamic acid-rich protein-like | 0.02371 | 68.20312 | FALSE | FALSE |
| LOC107064010 | probable methylmalonate-semialdehyde de | 0.02262 | 68.20977 | FALSE | FALSE |
| LOC107070253 | syntaxin-8 | 0.02301 | 68.22124 | FALSE | FALSE |
| LOC107071872 | borealin-like | 0.02341 | 68.23106 | FALSE | FALSE |
| LOC107067761 | protein odd-skipped | 0.02429 | 68.23689 | FALSE | FALSE |
| LOC107063834 | meiosis-specific nuclear structural protein 1 | 0.02365 | 68.23861 | FALSE | FALSE |
| LOC107070702 | histone-lysine N-methyltransferase SETMAF | 0.02575 | 68.24908 | FALSE | FALSE |
| LOC107072277 | calmodulin-binding transcription activator 2 | 0.02457 | 68.2611 | FALSE | FALSE |
| LOC107070259 | zinc finger protein 706-like | 0.02241 | 68.26175 | FALSE | FALSE |
| LOC107072478 | peroxisomal membrane protein 11B | 0.02287 | 68.26738 | FALSE | FALSE |
| LOC107067119 | meckelin-like | 0.02557 | 68.27442 | FALSE | FALSE |
| LOC107067838 | uncharacterized protein LOC107067838 | 0.023 | 68.27983 | FALSE | FALSE |
| LOC107070776 | uncharacterized protein LOC107070776 | 0.02331 | 68.29883 | FALSE | FALSE |
| LOC107067968 | coronin-2B-like | 0.02358 | 68.32006 | FALSE | FALSE |
| LOC107066060 | large subunit GTPase 1 homolog | 0.02415 | 68.32345 | FALSE | FALSE |
| LOC107067548 | transmembrane protein 199 | 0.02222 | 68.33348 | FALSE | FALSE |
| LOC107069088 | laminin subunit alpha-1 | 0.02265 | 68.34787 | FALSE | FALSE |
| LOC107068832 | 39S ribosomal protein L16, mitochondrial | 0.02339 | 68.35256 | FALSE | FALSE |
| LOC107070535 | glutamate receptor ionotropic, kainate 2 | 0.02406 | 68.35352 | FALSE | FALSE |
| LOC107074428 | THO complex subunit 5 homolog | 0.02301 | 68.37881 | FALSE | FALSE |
| LOC107065804 | uncharacterized protein LOC107065804 | 0.02417 | 68.45091 | FALSE | FALSE |
| LOC107065633 | phosphatidylinositol 3,4,5-trisphosphate 3-p | 0.02252 | 68.46334 | FALSE | FALSE |
| LOC107074436 | actin-related protein 2/3 complex subunit 4 | 0.02257 | 68.46821 | FALSE | FALSE |

| | | | | | |
|---|---|---|---|---|---|
| LOC107071540 | trafficking protein particle complex subunit | 0.02486 | 68.48082 | FALSE | FALSE |
| LOC107063811 | Golgi phosphoprotein 3 homolog sauron | 0.02348 | 68.50513 | FALSE | FALSE |
| LOC107067502 | uncharacterized protein LOC107067502 | 0.02304 | 68.52402 | FALSE | FALSE |
| LOC107065392 | protein max | 0.02258 | 68.54172 | FALSE | FALSE |
| LOC107070914 | tetraspanin-11-like | 0.02331 | 68.54445 | FALSE | FALSE |
| LOC107070276 | odorant receptor Or2-like | 0.02281 | 68.55313 | FALSE | FALSE |
| LOC107067902 | cytochrome P450 4C1-like | 0.02279 | 68.56525 | FALSE | FALSE |
| LOC107073086 | 5'-3' exoribonuclease 1 | 0.02294 | 68.57843 | FALSE | FALSE |
| LOC107066724 | ATP-binding cassette sub-family F member 2 | 0.0235 | 68.60052 | FALSE | FALSE |
| LOC107069263 | uncharacterized protein F13E9.13, mitochon | 0.02215 | 68.66076 | FALSE | FALSE |
| LOC107067014 | sialin | 0.02369 | 68.67184 | FALSE | FALSE |
| LOC107065664 | cyclin-dependent-like kinase 5 | 0.02336 | 68.67499 | FALSE | FALSE |
| LOC107070270 | histone deacetylase complex subunit SAP30 | 0.02424 | 68.70267 | FALSE | FALSE |
| LOC107070304 | putative uncharacterized protein DDB_G028 | 0.02302 | 68.70438 | FALSE | FALSE |
| LOC107072041 | uncharacterized protein LOC107072041 | 0.02381 | 68.7133 | FALSE | FALSE |
| LOC107074691 | uncharacterized protein LOC107074691 | 0.02357 | 68.717 | FALSE | FALSE |
| LOC107073277 | uncharacterized protein LOC107073277 | 0.02319 | 68.73588 | FALSE | FALSE |
| LOC107073836 | uncharacterized protein LOC107073836 | 0.02274 | 68.76445 | FALSE | FALSE |
| LOC107070258 | valine--tRNA ligase | 0.02269 | 68.77072 | FALSE | FALSE |
| LOC107065250 | uncharacterized protein LOC107065250 | 0.02185 | 68.78772 | FALSE | FALSE |
| LOC107071669 | uncharacterized protein LOC107071669 | 0.02387 | 68.79661 | FALSE | FALSE |
| LOC107071712 | tyrosine-protein kinase Drl | 0.02412 | 68.79884 | FALSE | FALSE |
| LOC107064164 | WD repeat-containing protein mio | 0.02353 | 68.79917 | FALSE | FALSE |
| LOC107063787 | UPF0235 protein C15orf40 homolog | 0.02322 | 68.80997 | FALSE | FALSE |
| LOC107066587 | uncharacterized protein LOC107066587 | 0.0232 | 68.81502 | FALSE | FALSE |
| LOC107064044 | uncharacterized protein LOC107064044 | 0.02406 | 68.8164 | FALSE | FALSE |
| LOC107072816 | tyrosine-protein kinase Drl-like | 0.0225 | 68.82196 | FALSE | FALSE |
| LOC107067419 | protein Wnt-6-like | 0.02273 | 68.84038 | FALSE | FALSE |
| LOC107069008 | 40S ribosomal protein S23 | 0.02142 | 68.85503 | FALSE | FALSE |
| LOC107073850 | uncharacterized protein LOC107073850 | 0.02395 | 68.87073 | FALSE | FALSE |
| LOC107070999 | acyl-coenzyme A thioesterase 13-like | 0.02326 | 68.87188 | FALSE | FALSE |
| LOC107073321 | caskin-1 | 0.02261 | 68.87558 | FALSE | FALSE |
| LOC107065637 | uncharacterized protein LOC107065637 | 0.0226 | 68.92169 | FALSE | FALSE |
| LOC107064348 | methylcrotonoyl-CoA carboxylase beta chai | 0.02327 | 68.92907 | FALSE | FALSE |
| LOC107074101 | WD repeat domain-containing protein 83 | 0.02432 | 68.93705 | FALSE | FALSE |
| LOC107064153 | 3'(2'),5'-bisphosphate nucleotidase 1 | 0.02254 | 68.94898 | FALSE | FALSE |
| LOC107066626 | oxytocin-neurophysin 1-like | 0.0234 | 68.95218 | FALSE | FALSE |
| LOC107070246 | AP-2 complex subunit sigma | 0.0241 | 68.98033 | FALSE | FALSE |
| LOC107067292 | tRNA-specific adenosine deaminase 2 | 0.02278 | 68.98034 | FALSE | FALSE |
| LOC107071095 | odorant receptor 4-like | 0.0239 | 68.98643 | FALSE | FALSE |
| LOC107068392 | lipid phosphate phosphohydrolase 1-like | 0.02301 | 68.98902 | FALSE | FALSE |
| LOC107066469 | smad nuclear interacting protein 1 | 0.02285 | 68.9916 | FALSE | FALSE |
| LOC107065941 | general transcription factor IIH subunit 5 | 0.02255 | 68.99654 | FALSE | FALSE |
| LOC107072828 | protein windpipe | 0.02409 | 69.01908 | FALSE | FALSE |
| LOC107069567 | uncharacterized protein LOC107069567 | 0.02242 | 69.02164 | FALSE | FALSE |
| Trnaq-cug | Trnaq-cug | 0.02271 | 69.03971 | FALSE | FALSE |
| LOC107073591 | eukaryotic translation initiation factor 3 sub | 0.02413 | 69.04051 | FALSE | FALSE |
| LOC107069173 | protein farnesyltransferase subunit beta | 0.02221 | 69.05868 | FALSE | FALSE |
| LOC107070001 | cyclin-T-like | 0.0235 | 69.07873 | FALSE | FALSE |
| LOC107066577 | exosome complex component RRP43-like | 0.02278 | 69.0904 | FALSE | FALSE |

| | | | | | |
|---|---|---|---|---|---|
| LOC107065703 | ribosome maturation protein SBDS | 0.02369 | 69.09661 | FALSE | FALSE |
| LOC107072062 | uncharacterized protein LOC107072062 | 0.02341 | 69.10016 | FALSE | FALSE |
| LOC107072229 | xanthine dehydrogenase | 0.02268 | 69.10119 | FALSE | FALSE |
| LOC107067453 | synaptotagmin-7 | 0.02425 | 69.10216 | FALSE | FALSE |
| LOC107069331 | serine protease HTRA2, mitochondrial | 0.02385 | 69.10374 | FALSE | FALSE |
| LOC107065785 | collagen alpha-1(V) chain-like | 0.02375 | 69.10489 | FALSE | FALSE |
| LOC107065298 | uncharacterized protein LOC107065298 | 0.02277 | 69.10875 | FALSE | FALSE |
| LOC107065167 | uncharacterized protein LOC107065167 | 0.02217 | 69.12838 | FALSE | FALSE |
| LOC107067744 | golgin-84-like | 0.02258 | 69.13827 | FALSE | FALSE |
| LOC107073056 | prolyl 4-hydroxylase subunit alpha-1 | 0.02296 | 69.14715 | FALSE | FALSE |
| LOC107069843 | uncharacterized protein LOC107069843 | 0.02343 | 69.1625 | FALSE | FALSE |
| LOC107070036 | venom acid phosphatase Acph-1-like | 0.02335 | 69.16631 | FALSE | FALSE |
| LOC107064650 | excitatory amino acid transporter-like | 0.0238 | 69.21551 | FALSE | FALSE |
| LOC107066635 | DNA-directed RNA polymerase III subunit RI | 0.02343 | 69.21981 | FALSE | FALSE |
| LOC107072114 | uncharacterized protein LOC107072114 | 0.02419 | 69.23175 | FALSE | FALSE |
| LOC107072743 | transcription factor TBF1-like | 0.02417 | 69.23264 | FALSE | FALSE |
| LOC107071032 | uncharacterized protein LOC107071032 | 0.02335 | 69.23927 | FALSE | FALSE |
| LOC107066748 | HEAT repeat-containing protein 6 | 0.02172 | 69.23938 | FALSE | FALSE |
| LOC107064017 | putative uncharacterized protein DDB_G028 | 0.02217 | 69.2436 | FALSE | FALSE |
| LOC107067545 | protein FAM76B-like | 0.02354 | 69.24597 | FALSE | FALSE |
| LOC107066435 | uncharacterized protein LOC107066435 | 0.0223 | 69.26475 | FALSE | FALSE |
| LOC107066995 | venom acid phosphatase Acph-1-like | 0.02387 | 69.26765 | FALSE | FALSE |
| LOC107064710 | slit homolog 1 protein | 0.02366 | 69.27246 | FALSE | FALSE |
| LOC107072535 | uncharacterized protein LOC107072535 | 0.02373 | 69.27743 | FALSE | FALSE |
| LOC107073419 | uncharacterized protein LOC107073419 | 0.02279 | 69.27923 | FALSE | FALSE |
| LOC107070592 | Golgi pH regulator | 0.02454 | 69.30551 | FALSE | FALSE |
| LOC107070103 | decaprenyl-diphosphate synthase subunit 1 | 0.02359 | 69.3105 | FALSE | FALSE |
| LOC107068050 | uncharacterized protein LOC107068050 | 0.02339 | 69.31357 | FALSE | FALSE |
| LOC107066252 | centromere-associated protein E | 0.02356 | 69.33953 | FALSE | FALSE |
| LOC107073726 | uncharacterized protein LOC107073726 | 0.02317 | 69.34644 | FALSE | FALSE |
| LOC107064093 | slit homolog 1 protein-like | 0.02204 | 69.35219 | FALSE | FALSE |
| LOC107066501 | ABC transporter G family member 23-like | 0.02342 | 69.36539 | FALSE | FALSE |
| LOC107069437 | muscle M-line assembly protein unc-89 | 0.02305 | 69.3904 | FALSE | FALSE |
| LOC107070229 | Golgi reassembly-stacking protein 2 | 0.02312 | 69.398 | FALSE | FALSE |
| LOC107071552 | serendipity locus protein H-1-like | 0.02446 | 69.40912 | FALSE | FALSE |
| LOC107068442 | feline leukemia virus subgroup C receptor-re | 0.02307 | 69.42924 | FALSE | FALSE |
| LOC107067089 | zinc finger homeobox protein 4 | 0.02211 | 69.46301 | FALSE | FALSE |
| LOC107071182 | probable Ras GTPase-activating protein | 0.02314 | 69.47983 | FALSE | FALSE |
| LOC107073765 | uncharacterized protein LOC107073765 | 0.02249 | 69.48349 | FALSE | FALSE |
| LOC107067951 | uncharacterized protein LOC107067951 | 0.02346 | 69.48791 | FALSE | FALSE |
| LOC107069774 | uncharacterized protein LOC107069774 | 0.02506 | 69.49048 | FALSE | FALSE |
| LOC107069524 | uncharacterized protein LOC107069524 | 0.02495 | 69.52208 | FALSE | FALSE |
| LOC107071161 | uncharacterized protein LOC107071161 | 0.02235 | 69.52483 | FALSE | FALSE |
| LOC107071171 | CAPA peptides-like | 0.02334 | 69.52988 | FALSE | FALSE |
| LOC107065139 | protein suppressor of sable | 0.02361 | 69.53075 | FALSE | FALSE |
| LOC107073878 | solute carrier family 2, facilitated glucose tra | 0.02285 | 69.53743 | FALSE | FALSE |
| LOC107070352 | T-complex protein 1 subunit epsilon | 0.02189 | 69.55527 | FALSE | FALSE |
| LOC107068347 | uncharacterized protein LOC107068347 | 0.02282 | 69.55686 | FALSE | FALSE |
| LOC107068146 | nucleolar protein 9 | 0.02269 | 69.56492 | FALSE | FALSE |
| LOC107070084 | uncharacterized protein LOC107070084 | 0.02376 | 69.57312 | FALSE | FALSE |

| | | | | | |
|---|---|---|---|---|---|
| LOC107074622 | adenylosuccinate synthetase | 0.02287 | 69.57548 | FALSE | FALSE |
| LOC107074060 | neuronal acetylcholine receptor subunit alp | 0.02188 | 69.58297 | FALSE | FALSE |
| LOC107068083 | homeobox protein ARX-like | 0.0227 | 69.60192 | FALSE | FALSE |
| LOC107071804 | ATM interactor | 0.02237 | 69.60525 | FALSE | FALSE |
| LOC107065673 | glutamyl-tRNA(Gln) amidotransferase subur | 0.02333 | 69.61207 | FALSE | FALSE |
| LOC107066170 | uncharacterized protein LOC107066170 | 0.02135 | 69.6289 | FALSE | FALSE |
| LOC107073601 | protein sickie | 0.02263 | 69.67586 | FALSE | FALSE |
| LOC107068446 | uncharacterized protein LOC107068446 | 0.02259 | 69.6888 | FALSE | FALSE |
| LOC107074429 | coiled-coil domain-containing protein 65 | 0.02208 | 69.69906 | FALSE | FALSE |
| LOC107070049 | probable G-protein coupled receptor 158 | 0.02374 | 69.70204 | FALSE | FALSE |
| LOC107069718 | SHC SH2 domain-binding protein 1 | 0.02283 | 69.74402 | FALSE | FALSE |
| LOC107071577 | pikachurin | 0.02344 | 69.74556 | FALSE | FALSE |
| LOC107072406 | 40S ribosomal protein S20 | 0.02212 | 69.77039 | FALSE | FALSE |
| LOC107069996 | centromere/kinetochore protein zw10 hom | 0.02223 | 69.7862 | FALSE | FALSE |
| LOC107067175 | ran-binding protein 9 | 0.02305 | 69.78802 | FALSE | FALSE |
| LOC107066650 | uncharacterized protein LOC107066650 | 0.02243 | 69.79599 | FALSE | FALSE |
| LOC107067100 | CD63 antigen-like | 0.02406 | 69.81172 | FALSE | FALSE |
| LOC107065218 | putative sodium-dependent multivitamin tr | 0.02275 | 69.83358 | FALSE | FALSE |
| LOC107068952 | uncharacterized protein LOC107068952 | 0.02292 | 69.84325 | FALSE | FALSE |
| LOC107065307 | BTB/POZ domain-containing protein KCTD1 | 0.02382 | 69.85768 | FALSE | FALSE |
| LOC107064015 | 39S ribosomal protein L19, mitochondrial | 0.02252 | 69.91846 | FALSE | FALSE |
| LOC107066354 | thioredoxin domain-containing protein 11 | 0.02217 | 69.93593 | FALSE | FALSE |
| LOC107071600 | nucleolar MIF4G domain-containing protein | 0.02257 | 69.94789 | FALSE | FALSE |
| LOC107065423 | caspase-1-like | 0.02293 | 69.95087 | FALSE | FALSE |
| LOC107063669 | uncharacterized protein LOC107063669 | 0.02272 | 69.96835 | FALSE | FALSE |
| LOC107068923 | regulator of G-protein signaling loco | 0.0223 | 69.99148 | FALSE | FALSE |
| LOC107070430 | uncharacterized protein LOC107070430 | 0.02247 | 69.99346 | FALSE | FALSE |
| LOC107069339 | protein escargot | 0.0239 | 70.01919 | FALSE | FALSE |
| LOC107073165 | sodium/potassium-transporting ATPase sub | 0.02348 | 70.02103 | FALSE | FALSE |
| LOC107063700 | ubiquinone biosynthesis monooxygenase C( | 0.02364 | 70.02831 | FALSE | FALSE |
| LOC107072368 | COUP transcription factor 2 | 0.02231 | 70.04822 | FALSE | FALSE |
| LOC107068676 | probable G-protein coupled receptor Mth-li | 0.0233 | 70.07449 | FALSE | FALSE |
| LOC107072336 | uncharacterized protein LOC107072336 | 0.02245 | 70.08795 | FALSE | FALSE |
| LOC107065982 | AMP deaminase 2 | 0.02369 | 70.08841 | FALSE | FALSE |
| LOC107064100 | baculoviral IAP repeat-containing protein 6 | 0.02357 | 70.11855 | FALSE | FALSE |
| LOC107069589 | sphingolipid delta(4)-desaturase DES1 | 0.02273 | 70.14098 | FALSE | FALSE |
| LOC107073536 | uncharacterized protein LOC107073536 | 0.02181 | 70.17052 | FALSE | FALSE |
| LOC107066412 | magnesium transporter NIPA2 | 0.02289 | 70.17778 | FALSE | FALSE |
| LOC107071247 | LOW QUALITY PROTEIN: NACHT and WD rep | 0.02418 | 70.18071 | FALSE | FALSE |
| LOC107068237 | serine/threonine-protein kinase pelle | 0.02259 | 70.18193 | FALSE | FALSE |
| LOC107069758 | forkhead box protein C1-B-like | 0.02205 | 70.18678 | FALSE | FALSE |
| LOC107065909 | uncharacterized protein LOC107065909 | 0.02291 | 70.18725 | FALSE | FALSE |
| LOC107067102 | probable protein S-acyltransferase 23 | 0.02283 | 70.19832 | FALSE | FALSE |
| LOC107066884 | dual oxidase maturation factor 1-like | 0.02315 | 70.21269 | FALSE | FALSE |
| LOC107068864 | ras-related protein M-Ras-like | 0.02101 | 70.2235 | FALSE | FALSE |
| LOC107067977 | uncharacterized protein LOC107067977 | 0.02295 | 70.23272 | FALSE | FALSE |
| LOC107073541 | nose resistant to fluoxetine protein 6-like | 0.02296 | 70.24285 | FALSE | FALSE |
| LOC107066934 | claspin | 0.02153 | 70.2514 | FALSE | FALSE |
| LOC107074184 | TBC1 domain family member 7 | 0.0224 | 70.2598 | FALSE | FALSE |
| LOC107072773 | lamin Dm0-like | 0.02393 | 70.26058 | FALSE | FALSE |

| | | | | | |
|---|---|---|---|---|---|
| LOC107073125 | WAP four-disulfide core domain protein 3-li | 0.02291 | 70.26456 | FALSE | FALSE |
| LOC107069750 | cilia- and flagella-associated protein 69-like | 0.02324 | 70.26591 | FALSE | FALSE |
| LOC107073282 | helicase SKI2W | 0.02097 | 70.26844 | FALSE | FALSE |
| LOC107067995 | anaphase-promoting complex subunit 13 | 0.02285 | 70.27294 | FALSE | FALSE |
| LOC107072015 | cell wall integrity and stress response comp | 0.02304 | 70.27525 | FALSE | FALSE |
| LOC107069928 | gamma-tubulin complex component 6 | 0.02268 | 70.2799 | FALSE | FALSE |
| LOC107065877 | CD81 protein | 0.02237 | 70.28725 | FALSE | FALSE |
| LOC107070594 | galactokinase | 0.02293 | 70.29066 | FALSE | FALSE |
| LOC107063927 | pre-mRNA-processing-splicing factor 8 | 0.021 | 70.29096 | FALSE | FALSE |
| LOC107065197 | cyclic AMP-dependent transcription factor / | 0.02164 | 70.308 | FALSE | FALSE |
| LOC107074611 | protein PFC0760c-like | 0.02332 | 70.31403 | FALSE | FALSE |
| LOC107074220 | uncharacterized protein LOC107074220 | 0.02188 | 70.36897 | FALSE | FALSE |
| LOC107066652 | carbonic anhydrase 2-like | 0.02363 | 70.38324 | FALSE | FALSE |
| LOC107072836 | LOW QUALITY PROTEIN: serine/threonine-p | 0.02122 | 70.38588 | FALSE | FALSE |
| LOC107074535 | probable RNA-binding protein 19 | 0.02415 | 70.39189 | FALSE | FALSE |
| LOC107073471 | cytochrome P450 4c3-like | 0.02275 | 70.40298 | FALSE | FALSE |
| LOC107072975 | serine/threonine-protein kinase 3 | 0.02322 | 70.40846 | FALSE | FALSE |
| LOC107068136 | PR domain zinc finger protein 10-like | 0.0221 | 70.41213 | FALSE | FALSE |
| LOC107071637 | retinol dehydrogenase 13-like | 0.02362 | 70.4487 | FALSE | FALSE |
| LOC107064628 | spliceosome-associated protein CWC15 hon | 0.02411 | 70.4513 | FALSE | FALSE |
| LOC107072290 | Werner Syndrome-like exonuclease | 0.02258 | 70.45793 | FALSE | FALSE |
| LOC107068082 | protein rogdi | 0.02187 | 70.47082 | FALSE | FALSE |
| LOC107070183 | S1 RNA-binding domain-containing protein : | 0.02261 | 70.48511 | FALSE | FALSE |
| LOC107070194 | uncharacterized protein LOC107070194 | 0.02349 | 70.50334 | FALSE | FALSE |
| LOC107068972 | uncharacterized protein LOC107068972 | 0.02278 | 70.51524 | FALSE | FALSE |
| LOC107065591 | APOPT family protein CG14806, mitochondı | 0.02394 | 70.51543 | FALSE | FALSE |
| LOC107070597 | uncharacterized protein LOC107070597 | 0.02379 | 70.52462 | FALSE | FALSE |
| LOC107067220 | uncharacterized protein LOC107067220 | 0.02296 | 70.53022 | FALSE | FALSE |
| LOC107074570 | uncharacterized protein LOC107074570 | 0.02148 | 70.56261 | FALSE | FALSE |
| LOC107069408 | probable ATP-dependent RNA helicase YTHI | 0.02137 | 70.60102 | FALSE | FALSE |
| LOC107068599 | ras-related protein Ral-a | 0.02175 | 70.63433 | FALSE | FALSE |
| LOC107067525 | GATA zinc finger domain-containing protein | 0.02202 | 70.65588 | FALSE | FALSE |
| LOC107073739 | toll-like receptor 4 | 0.02293 | 70.65807 | FALSE | FALSE |
| LOC107066870 | programmed cell death protein 2-like | 0.02359 | 70.66517 | FALSE | FALSE |
| LOC107068456 | glucose dehydrogenase [FAD, quinone | 0.02334 | 70.71246 | FALSE | FALSE |
| LOC107064469 | putative uncharacterized protein DDB_G02 | 0.02243 | 70.71843 | FALSE | FALSE |
| LOC107067375 | skin secretory protein xP2-like | 0.02217 | 70.72509 | FALSE | FALSE |
| LOC107063954 | uncharacterized protein LOC107063954 | 0.02347 | 70.73123 | FALSE | FALSE |
| LOC107070398 | uncharacterized protein KIAA1143 homolog | 0.02206 | 70.76823 | FALSE | FALSE |
| LOC107066883 | probable splicing factor, arginine/serine-rich | 0.02261 | 70.77464 | FALSE | FALSE |
| LOC107068114 | UPF0430 protein CG31712 | 0.02299 | 70.80924 | FALSE | FALSE |
| LOC107070215 | dopamine receptor 1-like | 0.02205 | 70.81612 | FALSE | FALSE |
| LOC107074549 | uncharacterized protein DDB_G0287625-lik | 0.02324 | 70.8531 | FALSE | FALSE |
| LOC107066821 | transcription factor Sp4-like | 0.02247 | 70.85866 | FALSE | FALSE |
| LOC107074010 | NAD-dependent protein deacetylase sirtuin- | 0.02334 | 70.87134 | FALSE | FALSE |
| LOC107074630 | protein 5NUC-like | 0.02284 | 70.88025 | FALSE | FALSE |
| LOC107065075 | transcription initiation factor TFIID subunit : | 0.02343 | 70.8888 | FALSE | FALSE |
| LOC107071779 | uncharacterized protein LOC107071779 | 0.0229 | 70.91693 | FALSE | FALSE |
| LOC107066538 | Golgi to ER traffic protein 4 homolog | 0.02231 | 70.92171 | FALSE | FALSE |
| LOC107072815 | pro-resilin-like | 0.02271 | 70.93902 | FALSE | FALSE |

| | | | | | |
|---|---|---|---|---|---|
| LOC107066831 | uncharacterized protein LOC107066831 | 0.02197 | 70.94911 | FALSE | FALSE |
| LOC107065510 | serine/arginine repetitive matrix protein 2 | 0.02216 | 70.95272 | FALSE | FALSE |
| LOC107068289 | uncharacterized protein LOC107068289 | 0.02307 | 70.95314 | FALSE | FALSE |
| LOC107064757 | synaptotagmin-10-like | 0.02345 | 70.95467 | FALSE | FALSE |
| LOC107064473 | nuclear pore complex protein Nup50 | 0.02137 | 70.96045 | FALSE | FALSE |
| LOC107068161 | glyoxylate/hydroxypyruvate reductase A-lik | 0.02265 | 70.96503 | FALSE | FALSE |
| LOC107068209 | pecanex-like protein 1 | 0.02317 | 70.97007 | FALSE | FALSE |
| LOC107069321 | ADP,ATP carrier protein | 0.0231 | 70.97126 | FALSE | FALSE |
| LOC107065522 | peptidyl-alpha-hydroxyglycine alpha-amidat | 0.02339 | 70.9938 | FALSE | FALSE |
| LOC107067499 | probable hydroxyacid-oxoacid transhydroge | 0.02254 | 71.02535 | FALSE | FALSE |
| LOC107068412 | protein HID1 | 0.02239 | 71.03895 | FALSE | FALSE |
| LOC107070768 | eukaryotic translation initiation factor 3 sub | 0.02218 | 71.04585 | FALSE | FALSE |
| LOC107073898 | endophilin-A | 0.02236 | 71.08064 | FALSE | FALSE |
| LOC107073493 | odorant receptor 82a-like | 0.02317 | 71.10174 | FALSE | FALSE |
| LOC107069831 | basic-leucine zipper transcription factor A | 0.02351 | 71.13228 | FALSE | FALSE |
| LOC107068489 | neurexin-1 | 0.02278 | 71.14153 | FALSE | FALSE |
| LOC107063966 | DNA repair protein RAD51 homolog A | 0.02232 | 71.14285 | FALSE | FALSE |
| LOC107070402 | presqualene diphosphate phosphatase | 0.02353 | 71.15515 | FALSE | FALSE |
| LOC107064665 | uncharacterized protein LOC107064665 | 0.02272 | 71.15738 | FALSE | FALSE |
| LOC107073223 | gem-associated protein 2 | 0.02287 | 71.15861 | FALSE | FALSE |
| LOC107073285 | protein toll-like | 0.02211 | 71.17082 | FALSE | FALSE |
| LOC107069413 | uncharacterized protein DDB_G0284459 | 0.02231 | 71.17568 | FALSE | FALSE |
| LOC107065375 | potassium voltage-gated channel subfamily | 0.02253 | 71.18884 | FALSE | FALSE |
| LOC107066772 | protein dead ringer-like | 0.02343 | 71.19535 | FALSE | FALSE |
| LOC107073812 | chondroitin sulfate glucuronyltransferase | 0.02108 | 71.20325 | FALSE | FALSE |
| LOC107070646 | E3 ubiquitin-protein ligase FANCL | 0.02209 | 71.20761 | FALSE | FALSE |
| LOC107074685 | dipeptidase 1 | 0.02224 | 71.21272 | FALSE | FALSE |
| LOC107070997 | uncharacterized protein LOC107070997 | 0.0226 | 71.2238 | FALSE | FALSE |
| LOC107069238 | malate dehydrogenase-like | 0.02182 | 71.22955 | FALSE | FALSE |
| LOC107067606 | uncharacterized protein LOC107067606 | 0.02142 | 71.23346 | FALSE | FALSE |
| LOC107065032 | MKRN2 opposite strand protein | 0.02254 | 71.25578 | FALSE | FALSE |
| LOC107064620 | death-associated protein 1 | 0.02172 | 71.29031 | FALSE | FALSE |
| LOC107066688 | uncharacterized protein LOC107066688 | 0.02071 | 71.29119 | FALSE | FALSE |
| LOC107063713 | uncharacterized protein LOC107063713 | 0.02215 | 71.29189 | FALSE | FALSE |
| LOC107064844 | uncharacterized protein LOC107064844 | 0.02205 | 71.32253 | FALSE | FALSE |
| LOC107069249 | uncharacterized protein LOC107069249 | 0.0233 | 71.32303 | FALSE | FALSE |
| LOC107067431 | macro domain-containing protein CT2219-li | 0.0223 | 71.32585 | FALSE | FALSE |
| LOC107070931 | ubiquitin-like protein 3 | 0.02179 | 71.34618 | FALSE | FALSE |
| LOC107068595 | gem-associated protein 6-like | 0.02132 | 71.36281 | FALSE | FALSE |
| LOC107071059 | N-alpha-acetyltransferase 11 | 0.02238 | 71.38583 | FALSE | FALSE |
| LOC107067456 | tektin-1 | 0.02145 | 71.39315 | FALSE | FALSE |
| LOC107065418 | glutathione synthetase-like | 0.02251 | 71.40238 | FALSE | FALSE |
| LOC107066757 | uncharacterized protein LOC107066757 | 0.02334 | 71.4152 | FALSE | FALSE |
| LOC107067986 | V-type proton ATPase subunit B | 0.02148 | 71.41894 | FALSE | FALSE |
| LOC107072693 | ran GTPase-activating protein 1 | 0.02369 | 71.43833 | FALSE | FALSE |
| LOC107074217 | rhophilin-2 | 0.02219 | 71.43976 | FALSE | FALSE |
| LOC107074057 | putative tRNA (cytidine(32)/guanosine(34)-2 | 0.02292 | 71.45 | FALSE | FALSE |
| LOC107063773 | uncharacterized protein LOC107063773 | 0.02191 | 71.45057 | FALSE | FALSE |
| LOC107070096 | speckle-type POZ protein | 0.02196 | 71.46285 | FALSE | FALSE |
| LOC107066751 | uncharacterized protein DDB_G0286591-lik | 0.02101 | 71.46524 | FALSE | FALSE |

| | | | | | |
|---|---|---|---|---|---|
| LOC107068208 | dnaJ homolog subfamily C member 28 | 0.02275 | 71.46553 | FALSE | FALSE |
| LOC107065078 | signaling mucin HKR1-like | 0.02188 | 71.47244 | FALSE | FALSE |
| LOC107067985 | uncharacterized protein LOC107067985 | 0.0224 | 71.47415 | FALSE | FALSE |
| LOC107071615 | tripartite motif-containing protein 2-like | 0.02196 | 71.50127 | FALSE | FALSE |
| LOC107073948 | secretory carrier-associated membrane pro | 0.02283 | 71.50828 | FALSE | FALSE |
| LOC107073385 | G-protein coupled receptor Mth2 | 0.02201 | 71.51128 | FALSE | FALSE |
| LOC107071485 | uncharacterized protein LOC107071485 | 0.02375 | 71.5179 | FALSE | FALSE |
| LOC107064561 | uncharacterized protein LOC107064561 | 0.02298 | 71.52814 | FALSE | FALSE |
| LOC107064005 | SPRY domain-containing protein 7 | 0.02164 | 71.54832 | FALSE | FALSE |
| LOC107073381 | uncharacterized protein LOC107073381 | 0.02266 | 71.54856 | FALSE | FALSE |
| LOC107073947 | chaoptin-like | 0.02349 | 71.553 | FALSE | FALSE |
| LOC107067189 | arylphorin subunit alpha-like | 0.02302 | 71.55436 | FALSE | TRUE |
| LOC107071202 | G-protein coupled receptor Mth2-like | 0.02336 | 71.56583 | FALSE | FALSE |
| LOC107065496 | transmembrane protein 209 | 0.02305 | 71.58543 | FALSE | FALSE |
| LOC107065952 | transmembrane protein 165 | 0.02276 | 71.59699 | FALSE | FALSE |
| LOC107070738 | mediator of RNA polymerase II transcriptior | 0.0229 | 71.59817 | FALSE | FALSE |
| LOC107074371 | enoyl-CoA delta isomerase 1, mitochondrial | 0.02215 | 71.60471 | FALSE | FALSE |
| LOC107066032 | transferrin | 0.02446 | 71.60653 | FALSE | FALSE |
| LOC107074132 | tryptophan 2,3-dioxygenase | 0.02338 | 71.63445 | FALSE | FALSE |
| LOC107064870 | sorting nexin-12 | 0.02243 | 71.63477 | FALSE | FALSE |
| LOC107069229 | uncharacterized protein LOC107069229 | 0.02175 | 71.64023 | FALSE | FALSE |
| LOC107065916 | WD repeat domain phosphoinositide-intera | 0.02236 | 71.64541 | FALSE | FALSE |
| LOC107071345 | multiple epidermal growth factor-like doma | 0.02116 | 71.66783 | FALSE | FALSE |
| LOC107067513 | transcriptional regulator of yeast form adhe | 0.02296 | 71.67126 | FALSE | FALSE |
| LOC107073894 | U11/U12 small nuclear ribonucleoprotein 5! | 0.02152 | 71.67675 | FALSE | FALSE |
| LOC107064401 | uncharacterized protein LOC107064401 | 0.02394 | 71.67914 | FALSE | FALSE |
| LOC107073434 | uncharacterized protein LOC107073434 | 0.02296 | 71.6867 | FALSE | FALSE |
| LOC107068868 | uncharacterized protein LOC107068868 | 0.02291 | 71.69124 | FALSE | FALSE |
| LOC107069330 | histidine triad nucleotide-binding protein 1 | 0.02216 | 71.70088 | FALSE | FALSE |
| LOC107064941 | cytochrome b-c1 complex subunit 8 | 0.0223 | 71.71451 | FALSE | FALSE |
| LOC107067881 | serine/threonine-protein kinase 11-interact | 0.02329 | 71.74427 | FALSE | FALSE |
| LOC107073397 | neurogenic locus notch homolog protein 1 | 0.02133 | 71.74697 | FALSE | FALSE |
| LOC107066280 | odorant receptor 46a, isoform A-like | 0.02275 | 71.77368 | FALSE | FALSE |
| LOC107068526 | neurotactin | 0.02191 | 71.78264 | FALSE | FALSE |
| LOC107072885 | IQ motif and SEC7 domain-containing prote | 0.02209 | 71.83516 | FALSE | FALSE |
| LOC107069394 | lon protease homolog, mitochondrial | 0.02204 | 71.84099 | FALSE | FALSE |
| LOC107066268 | 39S ribosomal protein L14, mitochondrial | 0.02328 | 71.84249 | FALSE | FALSE |
| LOC107072342 | saccharopine dehydrogenase-like oxidoredu | 0.02376 | 71.84985 | FALSE | FALSE |
| LOC107071808 | UPF0669 protein C6orf120 homolog | 0.02221 | 71.85112 | FALSE | FALSE |
| LOC107067440 | protein jim lovell | 0.02173 | 71.85687 | FALSE | FALSE |
| LOC107072300 | annexin B9-like | 0.02146 | 71.87856 | FALSE | FALSE |
| LOC107064697 | uncharacterized protein LOC107064697 | 0.02207 | 71.89495 | FALSE | FALSE |
| LOC107066239 | semaphorin-1A | 0.02236 | 71.9015 | FALSE | FALSE |
| LOC107068505 | uncharacterized protein LOC107068505 | 0.0216 | 71.91192 | FALSE | FALSE |
| LOC107072543 | uncharacterized protein C18orf8 | 0.02293 | 71.9447 | FALSE | FALSE |
| LOC107072705 | transcription initiation factor TFIID subunit : | 0.02295 | 71.9499 | FALSE | FALSE |
| LOC107072033 | hemicentin-2-like | 0.02153 | 71.97807 | FALSE | FALSE |
| LOC107067346 | mitochondrial Rho GTPase | 0.02181 | 71.99637 | FALSE | FALSE |
| LOC107070493 | venom serine protease-like | 0.02206 | 72.03825 | FALSE | FALSE |
| LOC107074341 | dihydropteridine reductase | 0.02238 | 72.04822 | FALSE | FALSE |

| | | | | | |
|---|---|---|---|---|---|
| LOC107068278 | transmembrane protease serine 9 | 0.02184 | 72.05702 | FALSE | FALSE |
| LOC107070608 | solute carrier family 22 member 8-like | 0.0217 | 72.06539 | FALSE | FALSE |
| LOC107069208 | uncharacterized protein LOC107069208 | 0.0232 | 72.07452 | FALSE | FALSE |
| LOC107072967 | uncharacterized protein LOC107072967 | 0.02296 | 72.10332 | FALSE | FALSE |
| LOC107067691 | phosphatidylinositol 4-kinase beta | 0.02392 | 72.10602 | FALSE | FALSE |
| LOC107069790 | lipase member H-like | 0.02282 | 72.11194 | FALSE | FALSE |
| LOC107067709 | type I inositol 3,4-bisphosphate 4-phosphat | 0.02264 | 72.14557 | FALSE | FALSE |
| LOC107067673 | transketolase-like protein 2 | 0.02325 | 72.16399 | FALSE | FALSE |
| LOC107068806 | protein tramtrack, beta isoform-like | 0.02269 | 72.18825 | FALSE | FALSE |
| LOC107070574 | vacuolar protein sorting-associated protein | 0.02185 | 72.20389 | FALSE | FALSE |
| LOC107072099 | protein ROP | 0.0222 | 72.20395 | FALSE | FALSE |
| LOC107069844 | zinc finger protein 1 | 0.02386 | 72.21599 | FALSE | FALSE |
| LOC107067878 | protein FAM122A | 0.02271 | 72.22443 | FALSE | FALSE |
| LOC107074693 | cAMP-responsive element-binding protein-l | 0.02344 | 72.22653 | FALSE | FALSE |
| LOC107068573 | probable N-acetyltransferase san | 0.02245 | 72.23207 | FALSE | FALSE |
| LOC107064957 | uncharacterized protein LOC107064957 | 0.02294 | 72.24047 | FALSE | FALSE |
| LOC107073712 | cyclin-T1-4-like | 0.02289 | 72.24418 | FALSE | FALSE |
| LOC107069664 | cytochrome c oxidase assembly protein CO | 0.02284 | 72.26768 | FALSE | FALSE |
| LOC107073564 | fatty acid synthase-like | 0.02223 | 72.27076 | FALSE | FALSE |
| LOC107069537 | protein Wnt-11b-1-like | 0.02197 | 72.27815 | FALSE | FALSE |
| LOC107073144 | LETM1 domain-containing protein 1 | 0.02276 | 72.28639 | FALSE | FALSE |
| LOC107068451 | glucose dehydrogenase [FAD, quinone | 0.02266 | 72.32805 | FALSE | FALSE |
| LOC107074258 | notchless protein homolog 1 | 0.02189 | 72.32833 | FALSE | FALSE |
| LOC107070368 | cytochrome P450 6a2-like | 0.02171 | 72.32999 | FALSE | FALSE |
| LOC107071279 | phosphatidylglycerophosphatase and protei | 0.02318 | 72.33046 | FALSE | FALSE |
| LOC107071519 | uncharacterized protein LOC107071519 | 0.02115 | 72.33477 | FALSE | FALSE |
| LOC107066844 | zinc finger protein 236-like | 0.02173 | 72.33798 | FALSE | FALSE |
| LOC107064046 | JNK1/MAPK8-associated membrane protein | 0.02307 | 72.338 | FALSE | FALSE |
| LOC107064312 | glutaredoxin-C4-like | 0.02294 | 72.3482 | FALSE | FALSE |
| LOC107063690 | 1-phosphatidylinositol 4,5-bisphosphate ph | 0.0224 | 72.36188 | FALSE | FALSE |
| LOC107072778 | WW domain-binding protein 11 | 0.02258 | 72.36846 | FALSE | FALSE |
| LOC107072167 | uncharacterized protein LOC107072167 | 0.02154 | 72.39174 | FALSE | FALSE |
| LOC107066027 | origin recognition complex subunit 3 | 0.02268 | 72.40087 | FALSE | FALSE |
| LOC107072380 | uncharacterized protein LOC107072380 | 0.02171 | 72.41301 | FALSE | FALSE |
| LOC107074653 | uncharacterized protein LOC107074653 | 0.0219 | 72.42175 | FALSE | FALSE |
| LOC107073198 | ras-related protein Rab-39B | 0.02281 | 72.43967 | FALSE | FALSE |
| LOC107070708 | allatostatins | 0.02219 | 72.47058 | FALSE | FALSE |
| LOC107074670 | uncharacterized protein LOC107074670 | 0.02308 | 72.47305 | FALSE | FALSE |
| LOC107071133 | protein YIPF1 | 0.02298 | 72.48138 | FALSE | FALSE |
| LOC107063919 | uncharacterized protein LOC107063919 | 0.02327 | 72.48718 | FALSE | FALSE |
| LOC107064489 | RNA binding protein fox-1 homolog 2 | 0.0223 | 72.48943 | FALSE | FALSE |
| LOC107069887 | uncharacterized protein LOC107069887 | 0.02324 | 72.48961 | FALSE | FALSE |
| LOC107064447 | uncharacterized protein LOC107064447 | 0.0218 | 72.50326 | FALSE | FALSE |
| LOC107068771 | regulator of G-protein signaling 17 | 0.02144 | 72.505 | FALSE | FALSE |
| LOC107065133 | mesoderm induction early response protein | 0.02206 | 72.50969 | FALSE | FALSE |
| LOC107074455 | run domain Beclin-1 interacting and cystein | 0.02319 | 72.51253 | FALSE | FALSE |
| LOC107064701 | WASH complex subunit FAM21 | 0.02081 | 72.52421 | FALSE | FALSE |
| LOC107066899 | potassium/sodium hyperpolarization-activa | 0.02228 | 72.53366 | FALSE | FALSE |
| LOC107072372 | glycine receptor subunit alpha-2 | 0.02131 | 72.54801 | FALSE | FALSE |
| LOC107068981 | lymphokine-activated killer T-cell-originatec | 0.02245 | 72.57711 | FALSE | FALSE |

| | | | | | |
|---|---|---|---|---|---|
| LOC107070640 | NADH-cytochrome b5 reductase-like | 0.02184 | 72.58126 | FALSE | FALSE |
| LOC107066254 | glycerophosphodiester phosphodiesterase ( | 0.02287 | 72.59093 | FALSE | FALSE |
| LOC107066575 | lambda-crystallin homolog | 0.02243 | 72.59408 | FALSE | FALSE |
| LOC107068085 | uncharacterized protein C4orf22 homolog | 0.02177 | 72.5943 | FALSE | FALSE |
| LOC107066195 | ER membrane protein complex subunit 3 | 0.02184 | 72.59773 | FALSE | FALSE |
| LOC107071021 | cleavage and polyadenylation specificity fac | 0.02403 | 72.62648 | FALSE | FALSE |
| LOC107067192 | broad-complex core protein isoforms 1/2/3, | 0.02177 | 72.63037 | FALSE | FALSE |
| LOC107068863 | cyclin-L1 | 0.02225 | 72.64804 | FALSE | FALSE |
| LOC107065640 | uncharacterized protein LOC107065640 | 0.02066 | 72.66198 | FALSE | FALSE |
| LOC107072578 | uncharacterized protein LOC107072578 | 0.02129 | 72.6725 | FALSE | FALSE |
| LOC107069947 | odorant receptor 13a-like | 0.02303 | 72.67965 | FALSE | FALSE |
| LOC107063975 | uncharacterized protein LOC107063975 | 0.02263 | 72.70884 | FALSE | FALSE |
| LOC107067804 | uncharacterized protein LOC107067804 | 0.02119 | 72.72652 | FALSE | FALSE |
| LOC107071475 | bifunctional glutamate/proline--tRNA ligase | 0.02193 | 72.72672 | FALSE | FALSE |
| LOC107071256 | kinectin | 0.02204 | 72.73183 | FALSE | FALSE |
| LOC107068320 | glucoside xylosyltransferase 1 | 0.02161 | 72.73287 | FALSE | FALSE |
| LOC107065149 | vinculin | 0.02406 | 72.73959 | FALSE | FALSE |
| LOC107073500 | uncharacterized protein LOC107073500 | 0.02257 | 72.74089 | FALSE | FALSE |
| LOC107073059 | uncharacterized protein LOC107073059 | 0.02269 | 72.80873 | FALSE | FALSE |
| LOC107064896 | intraflagellar transport protein 20 homolog | 0.02224 | 72.83748 | FALSE | FALSE |
| LOC107067174 | putative fatty acyl-CoA reductase CG8306 | 0.02166 | 72.84052 | FALSE | FALSE |
| LOC107067500 | E3 ubiquitin-protein ligase RNF146-A | 0.02234 | 72.8487 | FALSE | FALSE |
| LOC107063742 | protein-lysine N-methyltransferase mettl10 | 0.02227 | 72.89157 | FALSE | FALSE |
| LOC107070263 | serine/threonine-protein phosphatase 5 | 0.02206 | 72.90011 | FALSE | FALSE |
| LOC107073442 | mediator of RNA polymerase II transcriptior | 0.02244 | 72.9087 | FALSE | FALSE |
| LOC107074553 | protein MAK16 homolog | 0.02219 | 72.92727 | FALSE | FALSE |
| LOC107070347 | facilitated trehalose transporter Tret1-2 hor | 0.02253 | 72.94163 | FALSE | FALSE |
| LOC107074082 | probable DNA-directed RNA polymerases I a | 0.02204 | 72.97917 | FALSE | FALSE |
| LOC107069994 | RNA-binding protein Rsf1-like | 0.0219 | 73.02956 | FALSE | FALSE |
| LOC107067969 | omega-scoloptoxin-Ssm1a | 0.02208 | 73.05044 | FALSE | FALSE |
| LOC107071604 | sodium leak channel non-selective protein | 0.02194 | 73.07102 | FALSE | FALSE |
| LOC107067576 | ras-related protein Rap-2a | 0.02354 | 73.08662 | FALSE | FALSE |
| LOC107068691 | uncharacterized protein LOC107068691 | 0.02193 | 73.09697 | FALSE | FALSE |
| LOC107071497 | mediator of RNA polymerase II transcriptior | 0.02285 | 73.10122 | FALSE | FALSE |
| LOC107074654 | DNA repair protein xrcc4-like | 0.02253 | 73.10186 | FALSE | FALSE |
| LOC107064822 | protein mesh | 0.02356 | 73.10712 | FALSE | FALSE |
| LOC107073103 | uncharacterized protein LOC107073103 | 0.02339 | 73.13015 | FALSE | FALSE |
| LOC107066573 | bleomycin hydrolase | 0.0223 | 73.13151 | FALSE | FALSE |
| LOC107065645 | uncharacterized protein LOC107065645 | 0.02225 | 73.14466 | FALSE | FALSE |
| LOC107074115 | choline-phosphate cytidylyltransferase A-lik | 0.0223 | 73.14808 | FALSE | FALSE |
| LOC107067460 | fringe glycosyltransferase | 0.02355 | 73.14915 | FALSE | FALSE |
| LOC107065570 | uncharacterized protein LOC107065570 | 0.0218 | 73.16895 | FALSE | FALSE |
| LOC107072146 | RNA-binding protein 1-like | 0.02271 | 73.17843 | FALSE | FALSE |
| LOC107073865 | dysbindin protein homolog | 0.02179 | 73.1823 | FALSE | FALSE |
| LOC107069180 | uncharacterized protein LOC107069180 | 0.0218 | 73.18507 | FALSE | FALSE |
| LOC107070317 | glutathione S-transferase 1-1-like | 0.02285 | 73.195 | FALSE | FALSE |
| LOC107071989 | solute carrier family 41 member 3-like | 0.02261 | 73.19592 | FALSE | FALSE |
| LOC107064197 | B-cell receptor-associated protein 31 | 0.02172 | 73.20221 | FALSE | FALSE |
| LOC107068971 | cuticle protein 8-like | 0.02291 | 73.20401 | FALSE | FALSE |
| LOC107069767 | uncharacterized protein LOC107069767 | 0.02305 | 73.20719 | FALSE | FALSE |

| | | | | | |
|---|---|---|---|---|---|
| LOC107070376 | DNA ligase 1-like | 0.02179 | 73.20982 | FALSE | FALSE |
| LOC107071920 | uncharacterized protein LOC107071920 | 0.0231 | 73.22271 | FALSE | FALSE |
| LOC107073779 | RNA-binding protein 12B-like | 0.02088 | 73.24311 | FALSE | FALSE |
| LOC107072963 | protein MAATS1-like | 0.02354 | 73.25417 | FALSE | FALSE |
| LOC107072055 | probable ATP-dependent RNA helicase DDX | 0.02181 | 73.25795 | FALSE | FALSE |
| LOC107071209 | uncharacterized protein LOC107071209 | 0.02353 | 73.26237 | FALSE | FALSE |
| LOC107066054 | lachesin-like | 0.02156 | 73.27839 | FALSE | FALSE |
| LOC107063709 | succinate dehydrogenase [ubiquinone | 0.02219 | 73.29947 | FALSE | FALSE |
| LOC107073239 | adenylate cyclase type 2-like | 0.02259 | 73.30038 | FALSE | FALSE |
| LOC107073043 | Golgi SNAP receptor complex member 1 | 0.02207 | 73.30294 | FALSE | FALSE |
| LOC107067124 | palmitoyltransferase ZDHHC5 | 0.02201 | 73.31432 | FALSE | FALSE |
| LOC107069127 | moesin/ezrin/radixin homolog 1 | 0.02251 | 73.31824 | FALSE | FALSE |
| LOC107070766 | ubiquitin-60S ribosomal protein L40 | 0.0219 | 73.32482 | FALSE | FALSE |
| LOC107068336 | uncharacterized protein LOC107068336 | 0.02143 | 73.34343 | FALSE | FALSE |
| LOC107064168 | organic cation transporter protein-like | 0.02215 | 73.38195 | FALSE | FALSE |
| LOC107069885 | translational activator GCN1 | 0.0212 | 73.41052 | FALSE | FALSE |
| LOC107065031 | protein MTO1 homolog, mitochondrial | 0.02225 | 73.41205 | FALSE | FALSE |
| LOC107071919 | POU domain, class 6, transcription factor 1 | 0.02314 | 73.4382 | FALSE | FALSE |
| LOC107069672 | protein deltex | 0.02213 | 73.44828 | FALSE | FALSE |
| LOC107074280 | uncharacterized protein LOC107074280 | 0.02224 | 73.4634 | FALSE | FALSE |
| LOC107065181 | U1 small nuclear ribonucleoprotein A | 0.02239 | 73.50201 | FALSE | FALSE |
| LOC107067712 | WAS protein family homolog 1 | 0.02358 | 73.50582 | FALSE | FALSE |
| LOC107068419 | venom acid phosphatase Acph-1-like | 0.02128 | 73.51027 | FALSE | FALSE |
| LOC107072409 | dual specificity tyrosine-phosphorylation-re | 0.02153 | 73.51569 | FALSE | FALSE |
| LOC107068943 | transcriptional adapter 1-like | 0.02138 | 73.54166 | FALSE | FALSE |
| LOC107063933 | transcription factor E2F1 | 0.02255 | 73.55768 | FALSE | FALSE |
| LOC107071187 | uncharacterized protein LOC107071187 | 0.02246 | 73.56556 | FALSE | FALSE |
| LOC107064833 | uncharacterized protein LOC107064833 | 0.02335 | 73.56938 | FALSE | FALSE |
| LOC107071648 | uncharacterized protein LOC107071648 | 0.02234 | 73.5729 | FALSE | FALSE |
| LOC107073011 | P protein | 0.02266 | 73.59268 | FALSE | FALSE |
| LOC107068025 | myosin heavy chain, non-muscle-like | 0.02273 | 73.63266 | FALSE | FALSE |
| LOC107073941 | tensin-1-like | 0.02238 | 73.64592 | FALSE | FALSE |
| LOC107074157 | cell division cycle and apoptosis regulator pı | 0.02215 | 73.6607 | FALSE | FALSE |
| LOC107068433 | muscle M-line assembly protein unc-89 | 0.02119 | 73.67207 | FALSE | FALSE |
| LOC107069921 | uncharacterized protein LOC107069921 | 0.02135 | 73.6739 | FALSE | FALSE |
| LOC107071649 | protein PFC0760c-like | 0.02176 | 73.68829 | FALSE | FALSE |
| LOC107069222 | toll-like receptor 5 | 0.02271 | 73.68969 | FALSE | FALSE |
| LOC107074501 | LIM/homeobox protein Lhx2-like | 0.02197 | 73.70187 | FALSE | FALSE |
| LOC107070995 | sodium channel protein para | 0.02181 | 73.71003 | FALSE | FALSE |
| LOC107069976 | 60S ribosomal protein L23 | 0.02065 | 73.71164 | FALSE | FALSE |
| LOC107070249 | uncharacterized protein LOC107070249 | 0.02262 | 73.72218 | FALSE | FALSE |
| LOC107069072 | uncharacterized protein C18orf63-like | 0.02268 | 73.72431 | FALSE | FALSE |
| LOC107067664 | dipeptidase 1-like | 0.02073 | 73.7287 | FALSE | FALSE |
| LOC107074518 | pre-mRNA-processing factor 6 | 0.02312 | 73.78075 | FALSE | FALSE |
| LOC107065221 | uncharacterized protein LOC107065221 | 0.02384 | 73.78451 | FALSE | FALSE |
| LOC107068808 | alpha-L-fucosidase | 0.02249 | 73.80728 | FALSE | FALSE |
| LOC107073620 | cytochrome P450 4C1-like | 0.02184 | 73.85064 | FALSE | FALSE |
| LOC107072599 | uncharacterized protein LOC107072599 | 0.02135 | 73.87204 | FALSE | FALSE |
| LOC107072552 | uncharacterized protein LOC107072552 | 0.02161 | 73.89108 | FALSE | FALSE |
| LOC107065945 | transmembrane and ubiquitin-like domain-c | 0.0226 | 73.90398 | FALSE | FALSE |

| | | | | | |
|---|---|---|---|---|---|
| LOC107067728 | calexcitin-2 | 0.02181 | 73.90542 | FALSE | FALSE |
| LOC107064986 | ribonuclease H2 subunit C | 0.02203 | 73.91737 | FALSE | FALSE |
| LOC107072034 | speckle targeted PIP5K1A-regulated poly(A) | 0.02186 | 73.92997 | FALSE | FALSE |
| LOC107067636 | uncharacterized protein LOC107067636 | 0.02362 | 73.94857 | FALSE | FALSE |
| LOC107070678 | peripheral plasma membrane protein CASK- | 0.02135 | 73.95903 | FALSE | FALSE |
| LOC107072421 | uncharacterized protein LOC107072421 | 0.02139 | 73.99739 | FALSE | FALSE |
| LOC107064173 | fatty acid hydroxylase domain-containing pr | 0.02162 | 74.00031 | FALSE | FALSE |
| LOC107068867 | uncharacterized protein LOC107068867 | 0.02182 | 74.03343 | FALSE | FALSE |
| LOC107065127 | enolase-like | 0.02147 | 74.05424 | FALSE | FALSE |
| LOC107069237 | acetyl-CoA carboxylase | 0.02237 | 74.06818 | FALSE | FALSE |
| LOC107068870 | tropomyosin alpha-1 chain-like | 0.02273 | 74.10136 | FALSE | FALSE |
| LOC107074204 | hydrocephalus-inducing protein homolog | 0.02242 | 74.10241 | FALSE | FALSE |
| LOC107066156 | elongator complex protein 6 | 0.02135 | 74.10798 | FALSE | FALSE |
| LOC107066783 | adenylate kinase 8-like | 0.02196 | 74.11151 | FALSE | FALSE |
| LOC107066001 | uncharacterized protein LOC107066001 | 0.02146 | 74.11775 | FALSE | FALSE |
| LOC107065872 | DNA repair protein complementing XP-C cel | 0.02333 | 74.12736 | FALSE | FALSE |
| LOC107068700 | endocuticle structural glycoprotein SgAbd-4 | 0.02218 | 74.15829 | FALSE | FALSE |
| LOC107070315 | nuclear pore complex protein Nup107 | 0.02303 | 74.16639 | FALSE | FALSE |
| LOC107066471 | conserved oligomeric Golgi complex subunit | 0.02236 | 74.19037 | FALSE | FALSE |
| LOC107073070 | uncharacterized protein LOC107073070 | 0.022 | 74.1933 | FALSE | FALSE |
| LOC107074642 | protein groucho-1-like | 0.0213 | 74.209 | FALSE | FALSE |
| LOC107070519 | tetratricopeptide repeat protein 28 | 0.0205 | 74.21822 | FALSE | FALSE |
| LOC107068995 | sterol O-acyltransferase 1 | 0.02184 | 74.2526 | FALSE | FALSE |
| LOC107064803 | 4-coumarate--CoA ligase 1-like | 0.02223 | 74.26469 | FALSE | FALSE |
| LOC107065699 | serine-rich adhesin for platelets | 0.02301 | 74.28025 | FALSE | FALSE |
| LOC107071602 | chymotrypsin-1-like | 0.02236 | 74.2868 | FALSE | FALSE |
| LOC107067323 | transmembrane protein 131 | 0.02171 | 74.30256 | FALSE | FALSE |
| LOC107069529 | arylsulfatase B-like | 0.02076 | 74.34653 | FALSE | FALSE |
| LOC107065513 | probable ribonuclease ZC3H12B | 0.02227 | 74.3504 | FALSE | FALSE |
| LOC107067724 | serologically defined colon cancer antigen 8 | 0.02266 | 74.41328 | FALSE | FALSE |
| LOC107072854 | uncharacterized protein LOC107072854 | 0.02229 | 74.42969 | FALSE | FALSE |
| LOC107070092 | synaptic vesicle glycoprotein 2C-like | 0.0211 | 74.43879 | FALSE | FALSE |
| LOC107073243 | forkhead box protein C1-B-like | 0.02238 | 74.46331 | FALSE | FALSE |
| LOC107068882 | zinc finger protein 286A-like | 0.02137 | 74.472 | FALSE | FALSE |
| LOC107064612 | protein 60A | 0.02133 | 74.49875 | FALSE | FALSE |
| LOC107072949 | histone deacetylase 4 | 0.02076 | 74.5054 | FALSE | FALSE |
| LOC107067768 | WD repeat-containing protein 82 | 0.023 | 74.51586 | FALSE | FALSE |
| LOC107071476 | protein Wnt-1-like | 0.0222 | 74.51858 | FALSE | FALSE |
| LOC107066733 | tubulin polyglutamylase TTLL13P-like | 0.02211 | 74.52505 | FALSE | FALSE |
| LOC107064265 | centrosomal protein of 162 kDa-like | 0.02157 | 74.55835 | FALSE | FALSE |
| LOC107072627 | uncharacterized protein DDB_G0271670 | 0.02195 | 74.59089 | FALSE | FALSE |
| LOC107066398 | uncharacterized protein LOC107066398 | 0.0207 | 74.59939 | FALSE | FALSE |
| LOC107064703 | uncharacterized protein LOC107064703 | 0.02187 | 74.61958 | FALSE | FALSE |
| LOC107064319 | reversion-inducing cysteine-rich protein wit | 0.02359 | 74.63723 | FALSE | FALSE |
| LOC107072976 | sorting nexin-17 | 0.02321 | 74.64378 | FALSE | FALSE |
| LOC107063658 | peroxisome biogenesis factor 10-like | 0.02404 | 74.64663 | FALSE | FALSE |
| LOC107072624 | mitochondrial intermediate peptidase | 0.0219 | 74.69624 | FALSE | FALSE |
| LOC107069351 | lethal(2) giant larvae protein homolog 1 | 0.02211 | 74.71417 | FALSE | FALSE |
| LOC107067428 | serine/arginine repetitive matrix protein 2 | 0.02084 | 74.71726 | FALSE | FALSE |
| LOC107066987 | ATP synthase subunit alpha, mitochondrial | 0.02247 | 74.72362 | FALSE | FALSE |

| | | | | | |
|---|---|---|---|---|---|
| LOC107066739 | titin | 0.02134 | 74.74871 | FALSE | FALSE |
| LOC107068029 | E3 ubiquitin-protein ligase MARCH3-like | 0.02096 | 74.74924 | FALSE | FALSE |
| LOC107064572 | mitochondrial import inner membrane trans | 0.02165 | 74.75072 | FALSE | FALSE |
| LOC107066182 | H/ACA ribonucleoprotein complex subunit 3 | 0.02265 | 74.76012 | FALSE | FALSE |
| LOC107064427 | apolipoprotein D | 0.02222 | 74.77449 | FALSE | FALSE |
| LOC107073970 | uncharacterized protein LOC107073970 | 0.02301 | 74.77696 | FALSE | FALSE |
| Trnag-gcc | Trnag-gcc | 0.02143 | 74.80985 | FALSE | FALSE |
| LOC107065758 | synaptic vesicle glycoprotein 2B-like | 0.02279 | 74.85026 | FALSE | FALSE |
| LOC107067374 | soma ferritin-like | 0.02088 | 74.88974 | FALSE | FALSE |
| LOC107065463 | uncharacterized protein LOC107065463 | 0.02165 | 74.89746 | FALSE | FALSE |
| LOC107068939 | synaptic vesicular amine transporter | 0.0226 | 74.89946 | FALSE | FALSE |
| LOC107066178 | monocarboxylate transporter 9 | 0.02187 | 74.8999 | FALSE | FALSE |
| LOC107074105 | ATP synthase subunit b, mitochondrial | 0.02201 | 74.90065 | FALSE | FALSE |
| LOC107070736 | proteasome subunit alpha type-7-1 | 0.02102 | 74.91015 | FALSE | FALSE |
| LOC107074469 | glucosylceramidase | 0.02222 | 74.9376 | FALSE | FALSE |
| LOC107068266 | uroporphyrinogen-III synthase-like | 0.02239 | 74.93846 | FALSE | FALSE |
| LOC107074248 | sodium channel protein Nach-like | 0.02181 | 74.98443 | FALSE | FALSE |
| LOC107066941 | cytochrome c oxidase assembly protein CO) | 0.02244 | 75.00142 | FALSE | FALSE |
| LOC107064766 | protein takeout-like | 0.02191 | 75.03187 | FALSE | FALSE |
| LOC107070202 | ADP-ribosylation factor-like protein 5B | 0.02138 | 75.03602 | FALSE | FALSE |
| LOC107065639 | uncharacterized protein LOC107065639 | 0.0216 | 75.03862 | FALSE | FALSE |
| LOC107068013 | toll-like receptor 13 | 0.02152 | 75.04881 | FALSE | FALSE |
| LOC107064869 | glutamate [NMDA | 0.02197 | 75.06258 | FALSE | FALSE |
| LOC107072909 | rhombotin-1-like | 0.02167 | 75.07531 | FALSE | FALSE |
| LOC107063760 | arrestin domain-containing protein 4 | 0.02166 | 75.08067 | FALSE | FALSE |
| LOC107074163 | ankyrin repeat domain-containing protein 5 | 0.02156 | 75.09035 | FALSE | FALSE |
| LOC107073317 | autophagy-related protein 101 | 0.02169 | 75.09199 | FALSE | FALSE |
| LOC107067088 | ELAV-like protein 4 | 0.02268 | 75.09362 | FALSE | FALSE |
| LOC107067278 | hydroxymethylglutaryl-CoA lyase, mitochon | 0.02207 | 75.10469 | FALSE | FALSE |
| LOC107067544 | E3 ubiquitin-protein ligase RNF181-like | 0.02105 | 75.10501 | FALSE | FALSE |
| LOC107074316 | E3 ubiquitin-protein ligase RFWD3-like | 0.0211 | 75.11512 | FALSE | FALSE |
| LOC107065537 | eukaryotic translation initiation factor 4E ty | 0.02178 | 75.12182 | FALSE | FALSE |
| LOC107072483 | DNA-directed RNA polymerase III subunit RI | 0.02087 | 75.12431 | FALSE | FALSE |
| LOC107071760 | S-phase kinase-associated protein 2-like | 0.02299 | 75.14844 | FALSE | FALSE |
| LOC107073595 | cytochrome P450 4p1-like | 0.02275 | 75.15771 | FALSE | FALSE |
| LOC107066928 | proteasome maturation protein | 0.02205 | 75.17916 | FALSE | FALSE |
| LOC107071573 | programmed cell death protein 2 | 0.02205 | 75.18469 | FALSE | FALSE |
| LOC107067461 | uncharacterized protein PFB0145c | 0.02023 | 75.18904 | FALSE | FALSE |
| LOC107067448 | uncharacterized protein LOC107067448 | 0.02264 | 75.21434 | FALSE | FALSE |
| LOC107070863 | uncharacterized protein LOC107070863 | 0.02304 | 75.24642 | FALSE | FALSE |
| LOC107071098 | location of vulva defective 1-like | 0.02113 | 75.25275 | FALSE | FALSE |
| LOC107065976 | UDP-glucuronosyltransferase 1-4-like | 0.02288 | 75.26127 | FALSE | FALSE |
| LOC107064528 | MRG/MORF4L-binding protein | 0.02205 | 75.28227 | FALSE | FALSE |
| LOC107074182 | AF4/FMR2 family member 4-like | 0.02191 | 75.29842 | FALSE | FALSE |
| LOC107073415 | uncharacterized protein LOC107073415 | 0.02235 | 75.31418 | FALSE | FALSE |
| LOC107073613 | protein cycle-like | 0.02183 | 75.31694 | FALSE | FALSE |
| LOC107074461 | ubiA prenyltransferase domain-containing p | 0.0217 | 75.36879 | FALSE | FALSE |
| LOC107070888 | uncharacterized protein LOC107070888 | 0.02324 | 75.37049 | FALSE | FALSE |
| LOC107065950 | transcriptional regulatory protein GAT1-like | 0.02281 | 75.41317 | FALSE | FALSE |
| LOC107065085 | clavesin-2 | 0.02141 | 75.41389 | FALSE | FALSE |

| | | | | | |
|---|---|---|---|---|---|
| LOC107071858 | COMM domain-containing protein 3-like | 0.02184 | 75.42963 | FALSE | FALSE |
| LOC107070351 | tyrosine 3-monooxygenase | 0.0217 | 75.43243 | FALSE | FALSE |
| LOC107065434 | uncharacterized protein LOC107065434 | 0.0219 | 75.45734 | FALSE | FALSE |
| LOC107071517 | uncharacterized protein LOC107071517 | 0.02223 | 75.46536 | FALSE | FALSE |
| LOC107069039 | ubiquitin carboxyl-terminal hydrolase 16 | 0.02073 | 75.47036 | FALSE | FALSE |
| LOC107074605 | Bardet-Biedl syndrome 4 protein homolog | 0.02153 | 75.48055 | FALSE | FALSE |
| LOC107068514 | CDC42 small effector protein homolog | 0.02162 | 75.49133 | FALSE | FALSE |
| LOC107067348 | uncharacterized protein LOC107067348 | 0.02132 | 75.52572 | FALSE | FALSE |
| LOC107069552 | tropomyosin-2 | 0.02108 | 75.52882 | FALSE | FALSE |
| LOC107074392 | copper chaperone for superoxide dismutase | 0.02156 | 75.54123 | FALSE | FALSE |
| LOC107065344 | uncharacterized protein LOC107065344 | 0.02068 | 75.54881 | FALSE | FALSE |
| LOC107068877 | transcription elongation factor SPT5 | 0.02123 | 75.55531 | FALSE | FALSE |
| LOC107073838 | zinc finger SWIM domain-containing protein | 0.02232 | 75.56388 | FALSE | FALSE |
| LOC107069378 | uncharacterized protein LOC107069378 | 0.02188 | 75.56465 | FALSE | FALSE |
| LOC107067831 | uncharacterized protein LOC107067831 | 0.02126 | 75.58341 | FALSE | FALSE |
| LOC107073886 | uncharacterized protein LOC107073886 | 0.02215 | 75.58836 | FALSE | FALSE |
| LOC107065885 | DNA replication licensing factor Mcm7 | 0.02191 | 75.62489 | FALSE | FALSE |
| LOC107073848 | nuclear protein localization protein 4 homol | 0.02098 | 75.64183 | FALSE | FALSE |
| LOC107072654 | uncharacterized protein LOC107072654 | 0.02141 | 75.64592 | FALSE | FALSE |
| LOC107070854 | sodium-coupled monocarboxylate transport | 0.02206 | 75.65705 | FALSE | FALSE |
| LOC107066752 | ribonuclease P protein subunit p25-like prot | 0.02109 | 75.69664 | FALSE | FALSE |
| LOC107068859 | nuclease-sensitive element-binding protein | 0.02119 | 75.72793 | FALSE | FALSE |
| LOC107066137 | Down syndrome critical region protein 3 ho | 0.0225 | 75.73915 | FALSE | FALSE |
| LOC107074442 | nuclear speckle splicing regulatory protein 1 | 0.02143 | 75.74266 | FALSE | FALSE |
| LOC107069366 | frizzled-4 | 0.0227 | 75.75034 | FALSE | FALSE |
| LOC107069647 | Meckel syndrome type 1 protein-like | 0.021 | 75.77066 | FALSE | FALSE |
| LOC107072575 | carbonic anhydrase 2 | 0.02113 | 75.78668 | FALSE | FALSE |
| LOC107066737 | D-amino-acid oxidase | 0.02285 | 75.79328 | FALSE | FALSE |
| LOC107071138 | FH1/FH2 domain-containing protein 3 | 0.02044 | 75.81994 | FALSE | FALSE |
| Trnat-cgu | Trnat-cgu | 0.02123 | 75.83472 | FALSE | FALSE |
| LOC107063717 | prion-like-(Q/N-rich) domain-bearing protei | 0.02098 | 75.84059 | FALSE | FALSE |
| LOC107066323 | G protein-coupled receptor kinase 2 | 0.02285 | 75.84082 | FALSE | FALSE |
| LOC107068141 | neurocalcin homolog | 0.02225 | 75.85442 | FALSE | FALSE |
| LOC107066363 | phosphatase and actin regulator 4B | 0.0214 | 75.85505 | FALSE | FALSE |
| LOC107065818 | transcription termination factor 2 | 0.02198 | 75.85583 | FALSE | FALSE |
| LOC107067349 | protein C3orf33 | 0.02079 | 75.86593 | FALSE | FALSE |
| LOC107067981 | uncharacterized protein LOC107067981 | 0.02144 | 75.8748 | FALSE | FALSE |
| LOC107064625 | organic cation transporter protein-like | 0.02184 | 75.88412 | FALSE | FALSE |
| LOC107067625 | uncharacterized protein LOC107067625 | 0.02049 | 75.90619 | FALSE | FALSE |
| LOC107064350 | uncharacterized protein LOC107064350 | 0.02204 | 75.93716 | FALSE | FALSE |
| LOC107074603 | N-acetyltransferase 6 | 0.02101 | 75.96333 | FALSE | FALSE |
| LOC107073436 | mediator of RNA polymerase II transcription | 0.02322 | 75.96852 | FALSE | FALSE |
| LOC107063849 | uncharacterized protein LOC107063849 | 0.02153 | 75.99456 | FALSE | FALSE |
| LOC107067674 | 60S ribosomal protein L27a | 0.02364 | 76.00916 | FALSE | FALSE |
| LOC107068349 | protein angel | 0.02108 | 76.02018 | FALSE | FALSE |
| LOC107070058 | prenylated Rab acceptor protein 1 | 0.02211 | 76.03064 | FALSE | FALSE |
| LOC107065688 | DNA-directed RNA polymerase II subunit RP | 0.02233 | 76.06951 | FALSE | FALSE |
| LOC107064513 | uncharacterized protein LOC107064513 | 0.0217 | 76.07388 | FALSE | FALSE |
| LOC107069802 | uncharacterized protein LOC107069802 | 0.0215 | 76.07845 | FALSE | FALSE |
| LOC107066061 | nuclear protein 1 | 0.02071 | 76.09756 | FALSE | FALSE |

| | | | | | |
|---|---|---|---|---|---|
| LOC107074435 | transmembrane protein 45B-like | 0.02151 | 76.13181 | FALSE | FALSE |
| LOC107067380 | jmjC domain-containing protein 7 | 0.02199 | 76.13485 | FALSE | FALSE |
| LOC107072589 | growth/differentiation factor 8 | 0.02118 | 76.14444 | FALSE | FALSE |
| LOC107069347 | probable V-type proton ATPase subunit D 2 | 0.02083 | 76.14824 | FALSE | FALSE |
| LOC107065750 | WD repeat-containing protein 38-like | 0.02326 | 76.15524 | FALSE | FALSE |
| LOC107068910 | tyrosine-protein kinase transmembrane rec | 0.02139 | 76.16649 | FALSE | FALSE |
| LOC107071794 | calcium-transporting ATPase type 2C memb | 0.02085 | 76.17722 | FALSE | FALSE |
| LOC107063660 | odorant receptor 9a-like | 0.02126 | 76.18229 | FALSE | FALSE |
| LOC107069867 | uncharacterized protein LOC107069867 | 0.02203 | 76.18254 | FALSE | FALSE |
| LOC107064568 | 2-oxoglutarate dehydrogenase, mitochondr | 0.023 | 76.20091 | FALSE | FALSE |
| LOC107069124 | cyclin-dependent kinase 5 activator 1 | 0.02216 | 76.21225 | FALSE | FALSE |
| LOC107070879 | cytochrome c oxidase assembly protein CO) | 0.02209 | 76.21762 | FALSE | FALSE |
| LOC107064976 | survival of motor neuron-related-splicing fa | 0.02199 | 76.24372 | FALSE | FALSE |
| LOC107065587 | WD repeat-containing protein 38-like | 0.02091 | 76.27259 | FALSE | FALSE |
| LOC107069442 | dnaJ homolog subfamily B member 11 | 0.02228 | 76.29153 | FALSE | FALSE |
| LOC107074072 | DNA polymerase epsilon subunit 2 | 0.02164 | 76.29429 | FALSE | FALSE |
| LOC107063939 | protein ariadne-2 | 0.02113 | 76.33671 | FALSE | FALSE |
| LOC107065452 | bumetanide-sensitive sodium-(potassium)-c | 0.02227 | 76.37044 | FALSE | FALSE |
| LOC107074601 | mitogen-activated protein kinase kinase kin | 0.022 | 76.41614 | FALSE | FALSE |
| LOC107069611 | kinetochore-associated protein 1 | 0.02207 | 76.43573 | FALSE | FALSE |
| LOC107073955 | cathepsin L | 0.02142 | 76.43654 | FALSE | FALSE |
| LOC107068962 | sodium/potassium/calcium exchanger 4-like | 0.02053 | 76.4728 | FALSE | FALSE |
| LOC107069281 | glutathione synthetase-like | 0.02119 | 76.48176 | FALSE | FALSE |
| LOC107067880 | UPF0587 protein C1orf123 homolog | 0.02194 | 76.50415 | FALSE | FALSE |
| LOC107068014 | fibroblast growth factor receptor homolog 1 | 0.02065 | 76.51899 | FALSE | FALSE |
| LOC107063725 | uncharacterized protein LOC107063725 | 0.02184 | 76.55335 | FALSE | FALSE |
| LOC107064818 | sideroflexin-1 | 0.02252 | 76.56316 | FALSE | FALSE |
| LOC107069637 | SWI/SNF complex subunit SMARCC2 | 0.0217 | 76.57017 | FALSE | FALSE |
| LOC107072035 | uncharacterized protein LOC107072035 | 0.02217 | 76.57829 | FALSE | FALSE |
| LOC107068904 | MIT domain-containing protein 1-like | 0.02236 | 76.59427 | FALSE | FALSE |
| LOC107071123 | signal recognition particle 9 kDa protein | 0.02131 | 76.59625 | FALSE | FALSE |
| LOC107070033 | uncharacterized protein LOC107070033 | 0.02112 | 76.61967 | FALSE | FALSE |
| LOC107067993 | tetraspanin-33-like | 0.02274 | 76.64572 | FALSE | FALSE |
| LOC107064845 | tyrosine-protein phosphatase Lar | 0.02142 | 76.64797 | FALSE | FALSE |
| LOC107064705 | 1-phosphatidylinositol 4,5-bisphosphate ph | 0.02131 | 76.64938 | FALSE | FALSE |
| LOC107072832 | uncharacterized protein LOC107072832 | 0.02122 | 76.65954 | FALSE | FALSE |
| LOC107073370 | uncharacterized protein LOC107073370 | 0.02077 | 76.66573 | FALSE | FALSE |
| LOC107072744 | hybrid signal transduction protein dokA-like | 0.02327 | 76.69147 | FALSE | FALSE |
| LOC107064626 | synaptogenesis protein syg-2-like | 0.02208 | 76.70396 | FALSE | FALSE |
| LOC107068316 | uncharacterized protein LOC107068316 | 0.0224 | 76.71431 | FALSE | FALSE |
| LOC107073612 | uncharacterized protein LOC107073612 | 0.02079 | 76.76802 | FALSE | FALSE |
| LOC107066019 | DNA-directed RNA polymerase III subunit RI | 0.02132 | 76.77466 | FALSE | FALSE |
| LOC107074524 | protein lap1 | 0.02122 | 76.79071 | FALSE | FALSE |
| LOC107070094 | origin recognition complex subunit 2 | 0.02087 | 76.80144 | FALSE | FALSE |
| LOC107070763 | uncharacterized protein LOC107070763 | 0.02085 | 76.80826 | FALSE | FALSE |
| LOC107069199 | migration and invasion enhancer 1 | 0.02147 | 76.81794 | FALSE | FALSE |
| LOC107066798 | coiled-coil domain-containing protein 149 | 0.02078 | 76.82346 | FALSE | FALSE |
| LOC107067171 | heparan-alpha-glucosaminide N-acetyltrans | 0.02153 | 76.82377 | FALSE | FALSE |
| LOC107069763 | glucose dehydrogenase [FAD, quinone | 0.02158 | 76.85153 | FALSE | FALSE |
| LOC107073935 | alpha-2-macroglobulin-like | 0.02187 | 76.86431 | FALSE | FALSE |

| | | | | | |
|---|---|---|---|---|---|
| LOC107065576 | uncharacterized protein LOC107065576 | 0.02153 | 76.87747 | FALSE | FALSE |
| LOC107072725 | ribosomal protein S6 kinase alpha-5 | 0.02162 | 76.9003 | FALSE | FALSE |
| LOC107070540 | uncharacterized protein LOC107070540 | 0.021 | 76.90693 | FALSE | FALSE |
| LOC107073622 | glycosaminoglycan xylosylkinase | 0.02143 | 76.91235 | FALSE | FALSE |
| LOC107070461 | eukaryotic translation initiation factor 4 gan | 0.02077 | 76.93604 | FALSE | FALSE |
| LOC107071890 | succinate dehydrogenase [ubiquinone | 0.02281 | 76.94481 | FALSE | FALSE |
| LOC107070286 | arginine/serine-rich coiled-coil protein 2 | 0.02141 | 76.9527 | FALSE | FALSE |
| LOC107064151 | CD2 antigen cytoplasmic tail-binding proteir | 0.02047 | 77.01099 | FALSE | FALSE |
| LOC107070465 | uncharacterized protein LOC107070465 | 0.02186 | 77.01171 | FALSE | FALSE |
| LOC107074592 | eukaryotic translation initiation factor 1A, X | 0.02193 | 77.07979 | FALSE | FALSE |
| LOC107073213 | 28S ribosomal protein S22, mitochondrial | 0.02125 | 77.08124 | FALSE | FALSE |
| LOC107064182 | guanylate cyclase 32E | 0.0215 | 77.09483 | FALSE | FALSE |
| LOC107065460 | protein translation factor SUI1 homolog | 0.02182 | 77.12708 | FALSE | FALSE |
| LOC107065679 | riboflavin kinase | 0.02197 | 77.12975 | FALSE | FALSE |
| LOC107065114 | FERM, RhoGEF and pleckstrin domain-conta | 0.02269 | 77.13642 | FALSE | FALSE |
| LOC107071634 | tRNA pseudouridine synthase-like 1 | 0.02178 | 77.16169 | FALSE | FALSE |
| LOC107067237 | putative helicase mov-10-B.1 | 0.02071 | 77.17499 | FALSE | FALSE |
| LOC107064058 | glycerol-3-phosphate dehydrogenase [NAD( | 0.02097 | 77.17927 | FALSE | FALSE |
| LOC107067501 | class E basic helix-loop-helix protein 23 | 0.02189 | 77.18377 | FALSE | FALSE |
| LOC107068353 | transmembrane protein 189 | 0.02215 | 77.18718 | FALSE | FALSE |
| LOC107069295 | SID1 transmembrane family member 1-like | 0.02154 | 77.19204 | FALSE | FALSE |
| LOC107074225 | transmembrane inner ear expressed proteir | 0.02189 | 77.20408 | FALSE | FALSE |
| LOC107069083 | uncharacterized protein LOC107069083 | 0.02205 | 77.20694 | FALSE | FALSE |
| LOC107070799 | neuferricin | 0.02195 | 77.21011 | FALSE | FALSE |
| LOC107072354 | speckle-type POZ protein-like | 0.02114 | 77.22549 | FALSE | FALSE |
| LOC107070012 | TNF receptor-associated factor family prote | 0.02184 | 77.26276 | FALSE | FALSE |
| LOC107071916 | protein PET117 homolog, mitochondrial | 0.02119 | 77.30073 | FALSE | FALSE |
| LOC107072298 | piggyBac transposable element-derived pro | 0.02184 | 77.33515 | FALSE | FALSE |
| LOC107070266 | RCC1 and BTB domain-containing protein 1- | 0.02082 | 77.35618 | FALSE | FALSE |
| LOC107074245 | sex determination protein fruitless-like | 0.0214 | 77.35919 | FALSE | FALSE |
| LOC107074651 | tafazzin homolog | 0.0214 | 77.39855 | FALSE | FALSE |
| LOC107066251 | uncharacterized protein LOC107066251 | 0.02197 | 77.39953 | FALSE | FALSE |
| LOC107071737 | splicing regulator RBM11 | 0.02157 | 77.40396 | FALSE | FALSE |
| LOC107071238 | ataxin-3-like | 0.02147 | 77.41225 | FALSE | FALSE |
| LOC107072568 | uncharacterized protein LOC107072568 | 0.02185 | 77.42653 | FALSE | FALSE |
| LOC107065187 | putative uncharacterized protein DDB_G028 | 0.02225 | 77.44744 | FALSE | FALSE |
| LOC107071045 | uncharacterized protein C9orf78 | 0.02075 | 77.45849 | FALSE | FALSE |
| LOC107073879 | solute carrier family 25 member 35-like | 0.02237 | 77.47078 | FALSE | FALSE |
| LOC107064913 | transmembrane protein 242 | 0.02142 | 77.50494 | FALSE | FALSE |
| LOC107065984 | THAP domain-containing protein 4-like | 0.02143 | 77.50772 | FALSE | FALSE |
| LOC107071297 | lebercilin-like protein | 0.02264 | 77.59989 | FALSE | FALSE |
| LOC107069519 | prefoldin subunit 2 | 0.02269 | 77.61522 | FALSE | FALSE |
| LOC107069177 | uncharacterized protein LOC107069177 | 0.02172 | 77.6218 | FALSE | FALSE |
| LOC107070081 | 60S ribosomal protein L30 | 0.02213 | 77.63205 | FALSE | FALSE |
| LOC107064969 | histone acetyltransferase Tip60-like | 0.02163 | 77.63728 | FALSE | FALSE |
| LOC107068503 | uncharacterized protein LOC107068503 | 0.02117 | 77.64534 | FALSE | FALSE |
| LOC107064750 | uncharacterized protein LOC107064750 | 0.02222 | 77.64866 | FALSE | FALSE |
| LOC107064278 | transcription initiation factor TFIID subunit 9 | 0.0224 | 77.6513 | FALSE | FALSE |
| LOC107065399 | activating signal cointegrator 1 complex sub | 0.0228 | 77.65643 | FALSE | FALSE |
| LOC107064755 | uncharacterized protein LOC107064755 | 0.02117 | 77.666 | FALSE | FALSE |

| | | | | | |
|---|---|---|---|---|---|
| LOC107067942 | protein regulator of cytokinesis 1-like | 0.02126 | 77.66612 | FALSE | FALSE |
| LOC107074390 | polycomb protein Scm | 0.02111 | 77.67432 | FALSE | FALSE |
| LOC107066803 | fibroblast growth factor receptor 2-like | 0.0224 | 77.69715 | FALSE | FALSE |
| LOC107072100 | argininosuccinate synthase | 0.02016 | 77.71863 | FALSE | FALSE |
| LOC107071025 | uncharacterized protein LOC107071025 | 0.0211 | 77.72559 | FALSE | FALSE |
| LOC107074128 | coiled-coil domain-containing protein 142 | 0.02168 | 77.7394 | FALSE | FALSE |
| LOC107064894 | serine/threonine-protein kinase SBK1 | 0.02175 | 77.74013 | FALSE | FALSE |
| LOC107072115 | A disintegrin and metalloproteinase with th | 0.02071 | 77.74258 | FALSE | FALSE |
| LOC107068110 | uncharacterized protein LOC107068110 | 0.02091 | 77.75257 | FALSE | FALSE |
| LOC107074566 | E3 ubiquitin-protein ligase XIAP-like | 0.02159 | 77.77446 | FALSE | FALSE |
| LOC107065224 | probable tRNA(His) guanylyltransferase | 0.02117 | 77.78881 | FALSE | FALSE |
| LOC107065649 | mpv17-like protein 2 | 0.02108 | 77.79812 | FALSE | FALSE |
| LOC107063810 | COMM domain-containing protein 7-like | 0.02258 | 77.80575 | FALSE | FALSE |
| LOC107064839 | REM2- and Rab-like small GTPase 1 | 0.02182 | 77.80679 | FALSE | FALSE |
| LOC107066762 | ubiquitin-conjugating enzyme E2 G1 | 0.02071 | 77.8129 | FALSE | FALSE |
| LOC107070802 | glycoprotein-N-acetylgalactosamine 3-beta- | 0.02182 | 77.83103 | FALSE | FALSE |
| LOC107073592 | uncharacterized protein LOC107073592 | 0.02097 | 77.8393 | FALSE | FALSE |
| LOC107067253 | uncharacterized protein LOC107067253 | 0.02147 | 77.84568 | FALSE | FALSE |
| LOC107064871 | uncharacterized protein LOC107064871 | 0.02202 | 77.85468 | FALSE | FALSE |
| LOC107067161 | extensin | 0.02184 | 77.86717 | FALSE | FALSE |
| LOC107069536 | EF-hand domain-containing family member | 0.02016 | 77.87876 | FALSE | FALSE |
| LOC107068754 | uncharacterized protein LOC107068754 | 0.02102 | 77.90121 | FALSE | FALSE |
| LOC107074633 | FK506-binding protein 3-like | 0.02164 | 77.90127 | FALSE | FALSE |
| LOC107071928 | protein deadpan-like | 0.02075 | 77.92241 | FALSE | FALSE |
| LOC107073777 | splicing factor U2af large subunit B-like | 0.02016 | 77.92635 | FALSE | FALSE |
| LOC107072096 | uncharacterized protein LOC107072096 | 0.02133 | 77.9317 | FALSE | FALSE |
| LOC107073179 | uncharacterized protein LOC107073179 | 0.02135 | 77.93477 | FALSE | FALSE |
| LOC107071950 | netrin receptor UNC5C-like | 0.0208 | 77.9729 | FALSE | FALSE |
| LOC107065932 | CDK5RAP1-like protein | 0.02064 | 78.00167 | FALSE | FALSE |
| LOC107070178 | hypoxia up-regulated protein 1 | 0.02122 | 78.00259 | FALSE | FALSE |
| LOC107063878 | protein halfway | 0.02219 | 78.0294 | FALSE | FALSE |
| LOC107063840 | odorant receptor 22c-like | 0.0217 | 78.03012 | FALSE | FALSE |
| LOC107074538 | 39S ribosomal protein L43, mitochondrial | 0.02149 | 78.03684 | FALSE | FALSE |
| LOC107073811 | nuclear receptor coactivator 6-like | 0.02178 | 78.0474 | FALSE | FALSE |
| LOC107070243 | transmembrane protein 64 | 0.02073 | 78.05332 | FALSE | FALSE |
| LOC107068804 | phosphate carrier protein, mitochondrial | 0.02169 | 78.06101 | FALSE | FALSE |
| LOC107069574 | uncharacterized protein LOC107069574 | 0.0216 | 78.06824 | FALSE | FALSE |
| LOC107074362 | soluble lamin-associated protein of 75 kDa-l | 0.02126 | 78.07204 | FALSE | FALSE |
| LOC107071416 | glucose-6-phosphate isomerase | 0.02063 | 78.12126 | FALSE | FALSE |
| LOC107065970 | LOW QUALITY PROTEIN: rhodopsin-like | 0.02097 | 78.13345 | FALSE | FALSE |
| LOC107074370 | protein SET | 0.0214 | 78.14535 | FALSE | FALSE |
| LOC107068643 | quinone oxidoreductase-like protein 2 | 0.02157 | 78.19408 | FALSE | FALSE |
| LOC107074410 | transport and Golgi organization protein 2 h | 0.02139 | 78.19854 | FALSE | FALSE |
| LOC107070295 | histone-lysine N-methyltransferase NSD2 | 0.02086 | 78.23041 | FALSE | FALSE |
| LOC107071291 | eukaryotic translation initiation factor 4E-bi | 0.02043 | 78.233 | FALSE | FALSE |
| LOC107068553 | serine proteinase stubble | 0.02204 | 78.24754 | FALSE | FALSE |
| LOC107067457 | glyceraldehyde-3-phosphate dehydrogenase | 0.02135 | 78.24773 | FALSE | FALSE |
| LOC107072311 | pre-mRNA-processing factor 39-like | 0.02103 | 78.25298 | FALSE | FALSE |
| LOC107068856 | LOW QUALITY PROTEIN: fibrillin-1-like | 0.02158 | 78.26399 | FALSE | FALSE |
| LOC107065104 | uncharacterized protein LOC107065104 | 0.02144 | 78.29167 | FALSE | FALSE |

| | | | | | |
|---|---|---|---|---|---|
| LOC107064484 | cytochrome b-c1 complex subunit 7 | 0.02192 | 78.31916 | FALSE | FALSE |
| LOC107065345 | protein pelota | 0.02057 | 78.32566 | FALSE | FALSE |
| LOC107071993 | vacuolar protein sorting-associated protein | 0.02135 | 78.33611 | FALSE | FALSE |
| LOC107065305 | sushi, von Willebrand factor type A, EGF and | 0.0232 | 78.36013 | FALSE | FALSE |
| LOC107070936 | uncharacterized protein LOC107070936 | 0.02196 | 78.37176 | FALSE | FALSE |
| LOC107064095 | tubulin beta chain-like | 0.02189 | 78.37903 | FALSE | FALSE |
| LOC107073425 | uncharacterized protein LOC107073425 | 0.0215 | 78.4019 | FALSE | FALSE |
| LOC107067327 | 60S ribosomal protein L9 | 0.02107 | 78.40247 | FALSE | FALSE |
| LOC107067909 | odorant receptor 4-like | 0.02197 | 78.40273 | FALSE | FALSE |
| LOC107069034 | protein Cep78 homolog | 0.02243 | 78.41 | FALSE | FALSE |
| LOC107067810 | casein kinase II subunit beta | 0.02179 | 78.45328 | FALSE | FALSE |
| LOC107066630 | V-type proton ATPase subunit E | 0.02292 | 78.47267 | FALSE | FALSE |
| LOC107071252 | tRNA selenocysteine 1-associated protein 1 | 0.02166 | 78.47718 | FALSE | FALSE |
| LOC107065134 | V-type proton ATPase 21 kDa proteolipid su | 0.02123 | 78.49289 | FALSE | FALSE |
| LOC107063829 | defensin-1-like | 0.02055 | 78.51395 | FALSE | TRUE |
| LOC107069280 | UPF0598 protein CG30010 | 0.02173 | 78.55762 | FALSE | FALSE |
| LOC107072388 | allatotropin-like | 0.02203 | 78.57811 | FALSE | FALSE |
| LOC107071710 | transmembrane and TPR repeat-containing | 0.02143 | 78.5796 | FALSE | FALSE |
| LOC107071566 | uncharacterized protein LOC107071566 | 0.02069 | 78.58077 | FALSE | FALSE |
| LOC107069201 | chaoptin-like | 0.02151 | 78.6064 | FALSE | FALSE |
| LOC107072082 | uncharacterized protein LOC107072082 | 0.02193 | 78.61384 | FALSE | FALSE |
| LOC107068733 | uncharacterized protein LOC107068733 | 0.02107 | 78.62089 | FALSE | FALSE |
| LOC107069287 | NADH dehydrogenase [ubiquinone | 0.02068 | 78.62208 | FALSE | FALSE |
| LOC107071750 | HAUS augmin-like complex subunit 3 | 0.02133 | 78.66239 | FALSE | FALSE |
| LOC107067734 | putative RNA-binding protein 15 | 0.0214 | 78.6728 | FALSE | FALSE |
| LOC107067652 | phosphatidylinositol transfer protein alpha i | 0.02218 | 78.67539 | FALSE | FALSE |
| LOC107072811 | facilitated trehalose transporter Tret1-like | 0.02014 | 78.68318 | FALSE | FALSE |
| LOC107068328 | mannose-6-phosphate isomerase | 0.0215 | 78.69999 | FALSE | FALSE |
| LOC107074098 | ATP synthase-coupling factor 6, mitochondr | 0.02065 | 78.70256 | FALSE | FALSE |
| LOC107072680 | uncharacterized protein LOC107072680 | 0.02188 | 78.71525 | FALSE | FALSE |
| LOC107071038 | uncharacterized protein LOC107071038 | 0.02144 | 78.74068 | FALSE | FALSE |
| LOC107066399 | muscle LIM protein Mlp84B-like | 0.02105 | 78.7474 | FALSE | FALSE |
| LOC107071653 | myosin heavy chain 95F | 0.02165 | 78.76463 | FALSE | FALSE |
| LOC107070685 | aminopeptidase N | 0.02124 | 78.7867 | FALSE | FALSE |
| LOC107066910 | uncharacterized protein LOC107066910 | 0.02114 | 78.80173 | FALSE | FALSE |
| LOC107064885 | manganese-transporting ATPase 13A1 | 0.02178 | 78.84576 | FALSE | FALSE |
| LOC107066584 | arginase-2, mitochondrial | 0.02107 | 78.8542 | FALSE | FALSE |
| LOC107072669 | neuropeptide FF receptor 2-like | 0.02245 | 78.87732 | FALSE | FALSE |
| LOC107069922 | protein lingerer | 0.02157 | 78.98956 | FALSE | FALSE |
| LOC107074187 | lysosomal Pro-X carboxypeptidase | 0.02222 | 79.0097 | FALSE | FALSE |
| LOC107074634 | crossover junction endonuclease EME1 | 0.0213 | 79.02287 | FALSE | FALSE |
| LOC107072697 | beta carbonic anhydrase 1 | 0.02104 | 79.06424 | FALSE | FALSE |
| LOC107073646 | uncharacterized protein LOC107073646 | 0.02077 | 79.08403 | FALSE | FALSE |
| LOC107069683 | uncharacterized protein LOC107069683 | 0.02046 | 79.09532 | FALSE | FALSE |
| LOC107068453 | glucose dehydrogenase [FAD, quinone | 0.0215 | 79.10685 | FALSE | FALSE |
| LOC107064698 | GPI ethanolamine phosphate transferase 3 | 0.02102 | 79.17945 | FALSE | FALSE |
| LOC107068473 | uncharacterized protein LOC107068473 | 0.02246 | 79.18463 | FALSE | FALSE |
| LOC107067210 | transcription initiation factor TFIID subunit 4 | 0.02065 | 79.18691 | FALSE | FALSE |
| LOC107072293 | smoothelin-like protein 1 | 0.02087 | 79.20862 | FALSE | FALSE |
| LOC107067666 | splicing factor 3A subunit 2 | 0.02122 | 79.23322 | FALSE | FALSE |

| | | | | | |
|---|---|---|---|---|---|
| LOC107064092 | zinc finger protein 395 | 0.02181 | 79.24138 | FALSE | FALSE |
| LOC107071335 | exocyst complex component 5 | 0.02039 | 79.2704 | TRUE | FALSE |
| LOC107071765 | uncharacterized protein LOC107071765 | 0.02095 | 79.2829 | FALSE | FALSE |
| LOC107072989 | frizzled-9-like | 0.0211 | 79.32631 | FALSE | FALSE |
| LOC107065106 | DNA topoisomerase I, mitochondrial | 0.02099 | 79.35007 | FALSE | FALSE |
| LOC107074500 | ceramide kinase | 0.02137 | 79.36973 | FALSE | FALSE |
| LOC107074453 | importin subunit alpha | 0.02139 | 79.37219 | FALSE | FALSE |
| LOC107069244 | PERQ amino acid-rich with GYF domain-con | 0.0215 | 79.38722 | FALSE | FALSE |
| LOC107072101 | fez family zinc finger protein 2-like | 0.02105 | 79.39901 | FALSE | FALSE |
| LOC107066015 | pre-rRNA processing protein FTSJ3 | 0.02007 | 79.45828 | FALSE | FALSE |
| LOC107070131 | transcription factor SOX-11-like | 0.02283 | 79.47747 | FALSE | FALSE |
| LOC107071751 | probable serine/threonine-protein kinase fh | 0.02114 | 79.48991 | FALSE | FALSE |
| LOC107071425 | uncharacterized protein LOC107071425 | 0.02115 | 79.49837 | FALSE | FALSE |
| LOC107070373 | coiled-coil domain-containing protein 63 | 0.02156 | 79.49872 | FALSE | FALSE |
| LOC107065454 | tyrosine--tRNA ligase, cytoplasmic | 0.02139 | 79.53129 | FALSE | FALSE |
| LOC107070526 | uncharacterized protein LOC107070526 | 0.02083 | 79.53539 | FALSE | FALSE |
| LOC107071718 | intraflagellar transport protein 56 | 0.02165 | 79.54373 | FALSE | FALSE |
| LOC107074639 | exportin-7 | 0.02266 | 79.61439 | FALSE | FALSE |
| LOC107069139 | ubiquitin-conjugating enzyme E2Q-like prot | 0.02102 | 79.61663 | FALSE | FALSE |
| LOC107073386 | uncharacterized protein LOC107073386 | 0.02103 | 79.62626 | FALSE | FALSE |
| LOC107071276 | serine/threonine-protein kinase/endoribon | 0.02171 | 79.64604 | FALSE | FALSE |
| LOC107067521 | zinc finger and BTB domain-containing prot | 0.02166 | 79.64791 | FALSE | FALSE |
| LOC107070827 | pantothenate kinase 4 | 0.02033 | 79.64917 | FALSE | FALSE |
| LOC107069107 | stress protein DDR48-like | 0.02248 | 79.65004 | FALSE | FALSE |
| LOC107069613 | serine protease gd-like | 0.02165 | 79.66571 | FALSE | FALSE |
| LOC107064130 | uncharacterized protein LOC107064130 | 0.022 | 79.67617 | FALSE | FALSE |
| LOC107069620 | lysine-specific demethylase 3B | 0.02107 | 79.68936 | FALSE | FALSE |
| LOC107073367 | NADH dehydrogenase [ubiquinone | 0.02278 | 79.72441 | FALSE | FALSE |
| LOC107067247 | facilitated trehalose transporter Tret1-like | 0.02093 | 79.73398 | FALSE | FALSE |
| LOC107066302 | transcription factor Sox-2 | 0.02096 | 79.73933 | FALSE | FALSE |
| LOC107070753 | COMM domain-containing protein 8-like | 0.02175 | 79.74393 | FALSE | FALSE |
| LOC107069564 | spermine synthase | 0.02144 | 79.74898 | FALSE | FALSE |
| LOC107072506 | cytochrome c oxidase subunit 4 isoform 1, r | 0.02099 | 79.78099 | FALSE | FALSE |
| LOC107065759 | choline transporter-like protein 1 | 0.02223 | 79.78217 | FALSE | FALSE |
| LOC107069726 | spectrin alpha chain | 0.02044 | 79.79118 | FALSE | FALSE |
| LOC107071096 | uncharacterized protein LOC107071096 | 0.02085 | 79.81143 | FALSE | FALSE |
| LOC107067365 | RNA-binding protein with serine-rich domai | 0.0206 | 79.8398 | FALSE | FALSE |
| LOC107073830 | eukaryotic translation initiation factor 4E-1/ | 0.02201 | 79.84147 | FALSE | FALSE |
| LOC107070475 | serine/threonine-protein phosphatase 6 reg | 0.02107 | 79.84419 | FALSE | FALSE |
| LOC107073155 | glycerophosphocholine phosphodiesterase | 0.02182 | 79.87143 | FALSE | FALSE |
| LOC107068078 | homocysteine-responsive endoplasmic retic | 0.02134 | 79.9329 | FALSE | FALSE |
| LOC107073366 | probable ATP-dependent RNA helicase DDX | 0.02139 | 79.93925 | FALSE | FALSE |
| LOC107066909 | C-type lectin domain family 4 member E | 0.02206 | 79.96154 | FALSE | FALSE |
| LOC107074094 | protein UXT homolog | 0.02098 | 79.96453 | FALSE | FALSE |
| LOC107072827 | solute carrier organic anion transporter fam | 0.0218 | 80.00528 | FALSE | FALSE |
| LOC107068494 | uncharacterized protein LOC107068494 | 0.02105 | 80.02595 | FALSE | FALSE |
| LOC107066766 | maspardin-like | 0.02168 | 80.03533 | FALSE | FALSE |
| LOC107071830 | uncharacterized protein LOC107071830 | 0.02222 | 80.03835 | FALSE | FALSE |
| LOC107069981 | ubiquitin thioesterase otubain-like | 0.02074 | 80.06523 | FALSE | FALSE |
| LOC107070341 | PDZ and LIM domain protein 2-like | 0.02184 | 80.06556 | FALSE | FALSE |

| | | | | | |
|---|---|---|---|---|---|
| LOC107073153 | histone-lysine N-methyltransferase 2D-like | 0.02073 | 80.07389 | FALSE | FALSE |
| LOC107066841 | ubiquitin carboxyl-terminal hydrolase 14 | 0.02137 | 80.07769 | FALSE | FALSE |
| LOC107066760 | insulin-like receptor | 0.02185 | 80.09164 | FALSE | FALSE |
| LOC107072150 | esterase FE4-like | 0.02142 | 80.10197 | FALSE | FALSE |
| LOC107067717 | ATP synthase subunit O, mitochondrial | 0.022 | 80.10427 | FALSE | FALSE |
| LOC107072361 | protein no-on-transient A-like | 0.02035 | 80.10868 | FALSE | FALSE |
| LOC107063674 | biotin--protein ligase | 0.02113 | 80.13988 | FALSE | FALSE |
| LOC107064041 | WD repeat-containing protein 63-like | 0.0217 | 80.14718 | FALSE | FALSE |
| LOC107068409 | 30S ribosomal protein S11 | 0.02203 | 80.14733 | FALSE | FALSE |
| LOC107064549 | embryonic polarity protein dorsal-like | 0.02093 | 80.15302 | FALSE | FALSE |
| LOC107069452 | homeotic protein spalt-major-like | 0.02106 | 80.15766 | FALSE | FALSE |
| LOC107067932 | adenosylhomocysteinase | 0.02077 | 80.16059 | FALSE | FALSE |
| LOC107068900 | uncharacterized protein LOC107068900 | 0.02065 | 80.16937 | FALSE | FALSE |
| LOC107068702 | mitochondrial import inner membrane trans | 0.02077 | 80.17448 | FALSE | FALSE |
| LOC107072358 | ER degradation-enhancing alpha-mannosida | 0.02154 | 80.17512 | FALSE | FALSE |
| LOC107067279 | gibberellin 20 oxidase 1-like | 0.02082 | 80.17945 | FALSE | FALSE |
| LOC107066417 | microspherule protein 1 | 0.02167 | 80.21949 | FALSE | FALSE |
| LOC107069315 | deoxyuridine 5'-triphosphate nucleotidohyc | 0.02201 | 80.22066 | FALSE | FALSE |
| LOC107071977 | neuroligin-1-like | 0.02182 | 80.24 | FALSE | FALSE |
| LOC107064020 | uncharacterized protein LOC107064020 | 0.02117 | 80.24663 | FALSE | FALSE |
| LOC107071592 | serine-threonine kinase receptor-associatec | 0.02091 | 80.25092 | FALSE | FALSE |
| LOC107065240 | proliferating cell nuclear antigen | 0.02156 | 80.27333 | FALSE | FALSE |
| LOC107064099 | golgin subfamily A member 4-like | 0.02039 | 80.33889 | FALSE | FALSE |
| LOC107068358 | putative transferase CAF17 homolog, mitoc | 0.02052 | 80.38962 | FALSE | FALSE |
| LOC107068042 | 39S ribosomal protein L18, mitochondrial | 0.0208 | 80.39435 | FALSE | FALSE |
| LOC107065471 | pre-mRNA-splicing factor Slu7 | 0.02214 | 80.44911 | FALSE | FALSE |
| LOC107066571 | proteasome subunit alpha type-1 | 0.02153 | 80.46342 | FALSE | FALSE |
| LOC107067698 | uncharacterized protein LOC107067698 | 0.02059 | 80.46394 | FALSE | FALSE |
| LOC107074071 | serine/threonine-protein phosphatase 2A 5 | 0.02036 | 80.48003 | FALSE | FALSE |
| LOC107066722 | ADP-ribosylation factor 6 | 0.02069 | 80.48054 | FALSE | FALSE |
| LOC107067487 | ATP-dependent Clp protease proteolytic sul | 0.0215 | 80.4894 | FALSE | FALSE |
| LOC107071627 | coenzyme Q-binding protein COQ10 homolc | 0.02098 | 80.49375 | FALSE | FALSE |
| LOC107071069 | probable beta-hexosaminidase fdl | 0.02034 | 80.49926 | FALSE | FALSE |
| LOC107068614 | biogenesis of lysosome-related organelles c | 0.02117 | 80.50068 | FALSE | FALSE |
| LOC107066076 | bis(5'-nucleosyl)-tetraphosphatase [asymme | 0.02192 | 80.51052 | FALSE | FALSE |
| LOC107063912 | uncharacterized protein LOC107063912 | 0.02096 | 80.51351 | FALSE | FALSE |
| LOC107073422 | uncharacterized protein LOC107073422 | 0.02169 | 80.57195 | FALSE | FALSE |
| LOC107071074 | general odorant-binding protein 71 | 0.02137 | 80.59625 | FALSE | FALSE |
| LOC107070590 | sodium/potassium/calcium exchanger Nckx | 0.02101 | 80.60318 | FALSE | FALSE |
| LOC107068690 | probable NADH dehydrogenase [ubiquinone | 0.02095 | 80.63187 | FALSE | FALSE |
| LOC107070334 | uncharacterized protein LOC107070334 | 0.02176 | 80.6707 | FALSE | FALSE |
| LOC107072695 | CXXC-type zinc finger protein 1-like | 0.02105 | 80.70604 | FALSE | FALSE |
| LOC107070943 | ornithine aminotransferase, mitochondrial-l | 0.02065 | 80.71525 | FALSE | FALSE |
| LOC107069286 | ankyrin repeat and SAM domain-containing | 0.02036 | 80.73603 | FALSE | FALSE |
| LOC107071099 | odorant receptor 4-like | 0.02093 | 80.74022 | FALSE | FALSE |
| LOC107071419 | TRAF3-interacting protein 1 | 0.02046 | 80.777 | FALSE | FALSE |
| LOC107065204 | myb/SANT-like DNA-binding domain-contai | 0.02125 | 80.8371 | FALSE | FALSE |
| LOC107070233 | G-protein coupled receptor moody-like | 0.02133 | 80.85082 | FALSE | FALSE |
| LOC107068988 | uncharacterized protein DDB_G0287625-lik | 0.02284 | 80.89772 | FALSE | FALSE |
| LOC107071905 | uncharacterized protein LOC107071905 | 0.02061 | 80.90031 | FALSE | FALSE |

| | | | | | |
|---|---|---|---|---|---|
| LOC107068728 | cubilin-like | 0.02184 | 80.90284 | FALSE | FALSE |
| LOC107069073 | ankyrin repeat and EF-hand domain-contain | 0.02145 | 80.90522 | FALSE | FALSE |
| LOC107064356 | popeye domain-containing protein 3-like | 0.02008 | 80.91286 | FALSE | FALSE |
| LOC107072416 | protein wntless | 0.02163 | 80.92479 | FALSE | FALSE |
| LOC107064002 | ubiquitin conjugation factor E4 A | 0.02165 | 80.94198 | FALSE | FALSE |
| LOC107072250 | HMG box-containing protein 4 | 0.02071 | 80.94768 | FALSE | FALSE |
| LOC107065671 | GTP-binding protein 2 | 0.02102 | 80.9809 | FALSE | FALSE |
| LOC107073577 | vascular endothelial growth factor receptor | 0.02085 | 80.99807 | FALSE | FALSE |
| LOC107068315 | 26S protease regulatory subunit 7 | 0.02009 | 80.99874 | FALSE | FALSE |
| LOC107064655 | periodic tryptophan protein 1 homolog | 0.02152 | 81.0219 | FALSE | FALSE |
| LOC107073123 | meiosis arrest female protein 1-like | 0.02046 | 81.03167 | FALSE | FALSE |
| LOC107066704 | glycoprotein-N-acetylgalactosamine 3-beta- | 0.02048 | 81.05719 | FALSE | FALSE |
| LOC107069379 | uncharacterized protein LOC107069379 | 0.02095 | 81.06085 | FALSE | FALSE |
| LOC107072058 | protein DGCR14 homolog | 0.02164 | 81.06836 | FALSE | FALSE |
| LOC107064262 | peptide-N(4)-(N-acetyl-beta-glucosaminyl)a | 0.02113 | 81.11166 | FALSE | FALSE |
| LOC107070329 | ubiquitin carboxyl-terminal hydrolase 5 | 0.02114 | 81.12307 | FALSE | FALSE |
| LOC107074326 | probable signal peptidase complex subunit : | 0.02074 | 81.12479 | FALSE | FALSE |
| LOC107071282 | uncharacterized protein LOC107071282 | 0.02168 | 81.14847 | FALSE | FALSE |
| LOC107074276 | zinc finger matrin-type protein 5 | 0.02047 | 81.14929 | FALSE | FALSE |
| LOC107063766 | DNA polymerase epsilon subunit 4-like | 0.02147 | 81.17104 | FALSE | FALSE |
| LOC107065919 | cyclin-related protein FAM58A | 0.02078 | 81.19669 | FALSE | FALSE |
| LOC107074206 | uncharacterized protein LOC107074206 | 0.02116 | 81.23517 | FALSE | FALSE |
| LOC107072309 | uncharacterized protein LOC107072309 | 0.0209 | 81.23788 | FALSE | FALSE |
| LOC107072919 | zinc finger protein 830 | 0.02081 | 81.25276 | FALSE | FALSE |
| LOC107070924 | tryptase beta-2-like | 0.02072 | 81.25751 | FALSE | FALSE |
| LOC107073735 | uncharacterized protein LOC107073735 | 0.02087 | 81.2908 | FALSE | FALSE |
| LOC107073051 | zinc finger CCCH domain-containing protein | 0.02071 | 81.30991 | FALSE | FALSE |
| LOC107065125 | protein tweety | 0.02241 | 81.32957 | FALSE | FALSE |
| LOC107072529 | uncharacterized protein LOC107072529 | 0.02068 | 81.32986 | FALSE | FALSE |
| LOC107071320 | asparagine synthetase domain-containing p | 0.02072 | 81.33737 | FALSE | FALSE |
| LOC107071889 | succinate dehydrogenase [ubiquinone | 0.02131 | 81.34155 | FALSE | FALSE |
| LOC107066033 | major facilitator superfamily domain-contai | 0.02114 | 81.36706 | FALSE | FALSE |
| LOC107068124 | dual specificity protein phosphatase 3-like | 0.02003 | 81.38021 | FALSE | FALSE |
| LOC107072283 | homeobox protein rough-like | 0.0207 | 81.38245 | FALSE | FALSE |
| LOC107063654 | piggyBac transposable element-derived pro | 0.02132 | 81.38362 | FALSE | FALSE |
| LOC107064399 | uncharacterized protein LOC107064399 | 0.01939 | 81.39358 | FALSE | FALSE |
| LOC107069392 | uncharacterized protein LOC107069392 | 0.02128 | 81.4131 | FALSE | FALSE |
| LOC107070835 | WD repeat-containing protein 81 | 0.0206 | 81.42915 | FALSE | FALSE |
| LOC107065183 | 28S ribosomal protein S18c, mitochondrial | 0.02178 | 81.43603 | FALSE | FALSE |
| LOC107073068 | uncharacterized protein LOC107073068 | 0.02096 | 81.46842 | FALSE | FALSE |
| LOC107068395 | myosin-G heavy chain-like | 0.02154 | 81.48623 | FALSE | FALSE |
| LOC107066705 | FK506-binding protein 5-like | 0.02079 | 81.54781 | FALSE | FALSE |
| LOC107067522 | uncharacterized protein LOC107067522 | 0.02198 | 81.5937 | FALSE | FALSE |
| LOC107072896 | cytochrome P450 4C1 | 0.02097 | 81.61145 | FALSE | FALSE |
| LOC107074131 | uncharacterized protein LOC107074131 | 0.02243 | 81.61352 | FALSE | FALSE |
| LOC107067148 | DNA-directed RNA polymerase, mitochondr | 0.02204 | 81.61507 | FALSE | FALSE |
| LOC107071092 | protein FAM188B2 | 0.02125 | 81.62785 | FALSE | FALSE |
| LOC107064622 | protein FAM166B-like | 0.02229 | 81.62801 | FALSE | FALSE |
| LOC107064756 | glutaredoxin domain-containing cysteine-ric | 0.02096 | 81.66878 | FALSE | FALSE |
| LOC107070732 | neuromedin-U receptor 2-like | 0.02142 | 81.69556 | FALSE | FALSE |

| | | | | | |
|---|---|---|---|---|---|
| LOC107070464 | methionine aminopeptidase 2 | 0.02112 | 81.70085 | FALSE | FALSE |
| LOC107068708 | protein SMG8 | 0.02064 | 81.71253 | FALSE | FALSE |
| LOC107069191 | transcription initiation factor TFIID subunit 1 | 0.02241 | 81.71483 | FALSE | FALSE |
| LOC107074193 | uncharacterized protein LOC107074193 | 0.02095 | 81.71586 | FALSE | FALSE |
| LOC107074372 | leucine-rich repeat-containing protein 48-lik | 0.02113 | 81.7283 | FALSE | FALSE |
| LOC107070265 | ATP-binding cassette sub-family B member | 0.02399 | 81.72908 | FALSE | FALSE |
| LOC107073037 | protein pigeon | 0.02118 | 81.73712 | TRUE | FALSE |
| LOC107065646 | long-chain fatty acid transport protein 4-like | 0.0205 | 81.73843 | TRUE | FALSE |
| LOC107070907 | cullin-associated NEDD8-dissociated protein | 0.02177 | 81.75886 | TRUE | FALSE |
| LOC107070579 | mitochondrial uncoupling protein 2-like | 0.02112 | 81.76629 | FALSE | FALSE |
| LOC107069516 | ribosome biogenesis protein BMS1 homolog | 0.02164 | 81.80768 | FALSE | FALSE |
| LOC107073042 | uncharacterized protein LOC107073042 | 0.02229 | 81.81739 | FALSE | FALSE |
| LOC107067342 | cilia- and flagella-associated protein 97-like | 0.02124 | 81.82839 | FALSE | FALSE |
| LOC107068812 | spermatogenesis-associated protein 5 | 0.02091 | 81.83733 | FALSE | FALSE |
| LOC107071010 | transcription elongation factor SPT6 | 0.02051 | 81.83764 | FALSE | FALSE |
| LOC107069549 | CCR4-NOT transcription complex subunit 6-l | 0.02056 | 81.85783 | FALSE | FALSE |
| LOC107068129 | probable leucine--tRNA ligase, mitochondria | 0.02026 | 81.86305 | FALSE | FALSE |
| LOC107069283 | phosphatidylinositol 4,5-bisphosphate 3-kin | 0.02109 | 81.87697 | FALSE | FALSE |
| LOC107068310 | serine-rich adhesin for platelets-like | 0.02107 | 81.89432 | TRUE | FALSE |
| LOC107069474 | protein O-mannosyl-transferase 2-like | 0.01981 | 81.89445 | FALSE | FALSE |
| LOC107064006 | motile sperm domain-containing protein 1-l | 0.02206 | 81.89695 | FALSE | FALSE |
| LOC107070950 | HSPB1-associated protein 1 | 0.02096 | 81.90431 | FALSE | FALSE |
| LOC107070830 | uncharacterized protein LOC107070830 | 0.02137 | 81.90601 | FALSE | FALSE |
| LOC107066789 | pro-corazonin-like | 0.02033 | 81.91208 | FALSE | FALSE |
| LOC107070026 | uncharacterized protein LOC107070026 | 0.02011 | 81.93444 | FALSE | FALSE |
| LOC107065429 | uncharacterized protein LOC107065429 | 0.02085 | 81.93748 | FALSE | FALSE |
| LOC107073806 | NADH dehydrogenase [ubiquinone | 0.02162 | 81.97522 | FALSE | FALSE |
| LOC107070812 | 40S ribosomal protein S13 | 0.02134 | 81.97584 | FALSE | FALSE |
| LOC107065107 | uncharacterized protein LOC107065107 | 0.02013 | 82.01804 | FALSE | FALSE |
| LOC107063850 | uncharacterized protein LOC107063850 | 0.02125 | 82.03181 | FALSE | FALSE |
| LOC107064567 | dehydrogenase/reductase SDR family prote | 0.02245 | 82.06454 | FALSE | FALSE |
| LOC107065175 | glycine-rich cell wall structural protein-like | 0.0219 | 82.10119 | FALSE | FALSE |
| LOC107069883 | uncharacterized protein LOC107069883 | 0.02097 | 82.10147 | FALSE | FALSE |
| LOC107069857 | lactoylglutathione lyase | 0.02218 | 82.15384 | FALSE | FALSE |
| LOC107071686 | P-selectin | 0.02026 | 82.18585 | FALSE | FALSE |
| LOC107074342 | ATP synthase subunit e, mitochondrial | 0.02121 | 82.20002 | FALSE | FALSE |
| LOC107071028 | pro-resilin-like | 0.02293 | 82.20547 | FALSE | FALSE |
| LOC107064257 | DCN1-like protein 3 | 0.0207 | 82.23958 | FALSE | FALSE |
| LOC107065798 | uncharacterized protein LOC107065798 | 0.02112 | 82.25824 | TRUE | FALSE |
| LOC107072258 | guanine nucleotide-binding protein subunit | 0.02085 | 82.35703 | TRUE | FALSE |
| LOC107070641 | 40S ribosomal protein S5 | 0.02073 | 82.3807 | FALSE | FALSE |
| LOC107072914 | nucleoporin NSP1-like | 0.022 | 82.39279 | FALSE | FALSE |
| LOC107065023 | probable glutamate--tRNA ligase, mitochon | 0.02089 | 82.39717 | FALSE | FALSE |
| LOC107071667 | proline-rich protein 2-like | 0.01994 | 82.40388 | FALSE | FALSE |
| LOC107066780 | protein I'm not dead yet-like | 0.02144 | 82.42282 | FALSE | FALSE |
| LOC107072897 | cytochrome P450 4C1-like | 0.01988 | 82.46063 | FALSE | FALSE |
| LOC107072838 | nose resistant to fluoxetine protein 6-like | 0.02141 | 82.48392 | FALSE | FALSE |
| LOC107069513 | protein CLP1 homolog | 0.02162 | 82.4979 | FALSE | FALSE |
| LOC107073570 | uncharacterized protein LOC107073570 | 0.02084 | 82.5132 | FALSE | FALSE |
| LOC107070047 | uncharacterized protein LOC107070047 | 0.0217 | 82.517 | FALSE | TRUE |

| | | | | | |
|---|---|---|---|---|---|
| LOC107074150 | uncharacterized protein LOC107074150 | 0.02165 | 82.51745 | TRUE | FALSE |
| LOC107066213 | uncharacterized protein LOC107066213 | 0.02076 | 82.51889 | FALSE | FALSE |
| LOC107068928 | atrial natriuretic peptide receptor 1-like | 0.02004 | 82.52939 | FALSE | FALSE |
| LOC107065174 | eukaryotic translation initiation factor 2 sub | 0.0205 | 82.53763 | FALSE | FALSE |
| LOC107071936 | 1-acylglycerophosphocholine O-acyltransfer | 0.02061 | 82.53814 | FALSE | FALSE |
| LOC107069267 | uncharacterized protein LOC107069267 | 0.02071 | 82.54549 | FALSE | FALSE |
| LOC107072733 | uncharacterized protein LOC107072733 | 0.02266 | 82.56647 | FALSE | FALSE |
| LOC107066155 | uncharacterized protein LOC107066155 | 0.0214 | 82.5744 | FALSE | FALSE |
| LOC107065955 | E3 UFM1-protein ligase 1 homolog | 0.0216 | 82.58058 | FALSE | FALSE |
| LOC107072644 | KRAB-A domain-containing protein 2-like | 0.02067 | 82.59097 | FALSE | FALSE |
| LOC107066140 | uncharacterized protein LOC107066140 | 0.02038 | 82.62387 | FALSE | FALSE |
| LOC107064702 | V-type proton ATPase subunit F | 0.02045 | 82.62603 | FALSE | FALSE |
| LOC107063675 | DNA replication licensing factor Mcm5 | 0.02091 | 82.64141 | FALSE | FALSE |
| LOC107066727 | UDP-N-acetylhexosamine pyrophosphorylas | 0.0226 | 82.65969 | FALSE | FALSE |
| LOC107064872 | uncharacterized protein LOC107064872 | 0.02086 | 82.65976 | FALSE | FALSE |
| LOC107070043 | suppressor protein SRP40 | 0.02198 | 82.66136 | FALSE | FALSE |
| LOC107069468 | toll-like receptor 4 | 0.02113 | 82.6614 | FALSE | FALSE |
| LOC107071020 | neuroligin-4, Y-linked | 0.0217 | 82.66667 | FALSE | FALSE |
| LOC107071056 | venom dipeptidyl peptidase 4 | 0.02059 | 82.67203 | FALSE | FALSE |
| LOC107073694 | lysoplasmalogenase-like protein TMEM86A | 0.02293 | 82.67799 | FALSE | FALSE |
| LOC107073119 | organic cation transporter protein | 0.02056 | 82.68159 | FALSE | FALSE |
| LOC107068978 | b(0,+)-type amino acid transporter 1-like | 0.0207 | 82.69347 | FALSE | FALSE |
| LOC107073658 | calcium-activated potassium channel slowp | 0.02083 | 82.7093 | FALSE | FALSE |
| LOC107073075 | CCR4-NOT transcription complex subunit 11 | 0.02044 | 82.70949 | FALSE | FALSE |
| LOC107074523 | protein FAM195A-like | 0.02134 | 82.73194 | FALSE | FALSE |
| LOC107070691 | uncharacterized protein LOC107070691 | 0.02092 | 82.74219 | FALSE | FALSE |
| LOC107067820 | uncharacterized protein LOC107067820 | 0.0214 | 82.74483 | FALSE | FALSE |
| LOC107068562 | uncharacterized protein LOC107068562 | 0.02072 | 82.74622 | FALSE | FALSE |
| LOC107069148 | loricrin | 0.02159 | 82.76894 | FALSE | FALSE |
| LOC107064779 | uncharacterized protein LOC107064779 | 0.02027 | 82.77489 | FALSE | FALSE |
| LOC107071941 | protein arginine N-methyltransferase 7 | 0.0216 | 82.78002 | FALSE | FALSE |
| LOC107066758 | homeobox protein cut-like | 0.022 | 82.79917 | TRUE | FALSE |
| LOC107069571 | uncharacterized protein LOC107069571 | 0.02074 | 82.83451 | FALSE | FALSE |
| LOC107065610 | GTP-binding protein 10 homolog | 0.02155 | 82.83521 | TRUE | FALSE |
| LOC107074491 | peroxisomal multifunctional enzyme type 2- | 0.02434 | 82.83697 | FALSE | FALSE |
| LOC107066368 | protein lap4 | 0.02075 | 82.83729 | FALSE | FALSE |
| LOC107071932 | OTU domain-containing protein 6B | 0.0207 | 82.92887 | FALSE | FALSE |
| LOC107066430 | chondroitin sulfate synthase 1 | 0.02097 | 82.93962 | FALSE | FALSE |
| LOC107073224 | uncharacterized protein LOC107073224 | 0.02097 | 82.95898 | FALSE | FALSE |
| LOC107070014 | methyltransferase-like protein 4 | 0.0203 | 82.96932 | FALSE | FALSE |
| LOC107072092 | DC-STAMP domain-containing protein 1-like | 0.02091 | 82.97807 | FALSE | FALSE |
| LOC107072571 | very-long-chain enoyl-CoA reductase | 0.02045 | 83.01155 | FALSE | FALSE |
| LOC107071482 | uncharacterized protein LOC107071482 | 0.02114 | 83.02336 | FALSE | FALSE |
| LOC107065001 | probable 28S ribosomal protein S16, mitoch | 0.02138 | 83.02434 | FALSE | FALSE |
| LOC107070902 | uncharacterized protein LOC107070902 | 0.02149 | 83.07214 | FALSE | FALSE |
| LOC107070150 | mucin-5AC | 0.02219 | 83.0734 | TRUE | FALSE |
| LOC107071536 | biorientation of chromosomes in cell divisio | 0.02032 | 83.0847 | FALSE | FALSE |
| LOC107069540 | uncharacterized protein LOC107069540 | 0.02141 | 83.10127 | FALSE | FALSE |
| LOC107069685 | uncharacterized protein LOC107069685 | 0.02147 | 83.11702 | FALSE | FALSE |
| LOC107070901 | probable peroxisomal acyl-coenzyme A oxid | 0.02039 | 83.12663 | FALSE | FALSE |

| | | | | | |
|---|---|---|---|---|---|
| LOC107068518 | potassium/sodium hyperpolarization-activa | 0.02129 | 83.15004 | FALSE | FALSE |
| LOC107074350 | DNA repair and recombination protein RAD! | 0.02174 | 83.16033 | FALSE | FALSE |
| LOC107071880 | vitellogenin receptor | 0.02063 | 83.19648 | FALSE | FALSE |
| LOC107069412 | serine/threonine-protein kinase minibrain | 0.02104 | 83.19826 | FALSE | FALSE |
| LOC107065906 | uncharacterized protein LOC107065906 | 0.022 | 83.19926 | FALSE | FALSE |
| LOC107065942 | protein OSCP1 | 0.02006 | 83.21023 | FALSE | FALSE |
| LOC107068723 | tRNA-splicing endonuclease subunit Sen34 | 0.0219 | 83.2174 | FALSE | FALSE |
| LOC107066329 | ecdysone-induced protein 78C | 0.02065 | 83.23779 | FALSE | FALSE |
| LOC107073816 | venom acid phosphatase Acph-1-like | 0.02252 | 83.28469 | FALSE | FALSE |
| LOC107073261 | uncharacterized protein LOC107073261 | 0.02069 | 83.31106 | FALSE | FALSE |
| LOC107064420 | plasminogen activator inhibitor 1 RNA-bindi | 0.02053 | 83.33358 | TRUE | FALSE |
| LOC107074508 | GDP-D-glucose phosphorylase 1 | 0.02212 | 83.34245 | TRUE | FALSE |
| LOC107072871 | uncharacterized protein LOC107072871 | 0.0215 | 83.35101 | FALSE | FALSE |
| LOC107072465 | polyadenylate-binding protein 4-like | 0.02046 | 83.38751 | FALSE | FALSE |
| LOC107066181 | probable serine/threonine-protein kinase D | 0.02045 | 83.3945 | FALSE | FALSE |
| LOC107073276 | THAP domain-containing protein 1-like | 0.02163 | 83.40685 | FALSE | FALSE |
| LOC107073216 | uncharacterized protein LOC107073216 | 0.02142 | 83.42263 | FALSE | FALSE |
| LOC107063799 | uncharacterized protein LOC107063799 | 0.02149 | 83.43937 | FALSE | FALSE |
| LOC107065963 | protein tramtrack, beta isoform-like | 0.02196 | 83.4428 | FALSE | FALSE |
| LOC107068400 | gamma-tubulin complex component 5 | 0.02056 | 83.45769 | FALSE | FALSE |
| LOC107071225 | larval cuticle protein A3A-like | 0.02089 | 83.45998 | FALSE | FALSE |
| LOC107069673 | RILP-like protein homolog | 0.02086 | 83.46203 | FALSE | FALSE |
| LOC107064935 | alpha-protein kinase 1-like | 0.02118 | 83.47267 | FALSE | FALSE |
| LOC107067567 | caseinolytic peptidase B protein homolog | 0.0217 | 83.47371 | FALSE | FALSE |
| LOC107063959 | tubulin polyglutamylase TTLL4-like | 0.02165 | 83.4945 | FALSE | FALSE |
| LOC107070450 | alpha-protein kinase 1-like | 0.01935 | 83.49704 | TRUE | FALSE |
| LOC107065242 | protein RCC2 homolog | 0.01989 | 83.49772 | FALSE | FALSE |
| LOC107071575 | sine oculis-binding protein homolog | 0.02064 | 83.50812 | TRUE | FALSE |
| LOC107069193 | muscle-specific protein 20-like | 0.02101 | 83.50878 | TRUE | FALSE |
| LOC107063842 | uncharacterized protein LOC107063842 | 0.02146 | 83.54732 | FALSE | FALSE |
| LOC107063844 | uncharacterized protein LOC107063844 | 0.0215 | 83.56505 | FALSE | FALSE |
| LOC107067640 | tetratricopeptide repeat protein 21B-like | 0.02301 | 83.56894 | FALSE | FALSE |
| LOC107067514 | gastrula zinc finger protein XlCGF8.2DB-like | 0.02109 | 83.60267 | FALSE | FALSE |
| LOC107066790 | exportin-1 | 0.02211 | 83.62493 | FALSE | FALSE |
| LOC107064993 | putative ATP synthase subunit f, mitochond | 0.02123 | 83.63311 | FALSE | FALSE |
| LOC107063804 | uncharacterized protein YJR142W | 0.02163 | 83.64036 | FALSE | FALSE |
| LOC107073884 | growth arrest and DNA damage-inducible pı | 0.02229 | 83.68717 | FALSE | FALSE |
| LOC107072159 | protein transport protein Sec24C | 0.02106 | 83.68861 | FALSE | FALSE |
| LOC107070345 | protein downstream neighbor of son homol | 0.02117 | 83.70889 | FALSE | FALSE |
| LOC107063661 | uncharacterized protein LOC107063661 | 0.02277 | 83.71381 | TRUE | FALSE |
| LOC107064504 | hemK methyltransferase family member 1 | 0.02069 | 83.7726 | FALSE | FALSE |
| LOC107072169 | uncharacterized protein LOC107072169 | 0.02063 | 83.77775 | FALSE | FALSE |
| LOC107064643 | lysine-specific histone demethylase 1A-like | 0.02226 | 83.7902 | FALSE | FALSE |
| LOC107066356 | abnormal spindle-like microcephaly-associa | 0.02141 | 83.80495 | FALSE | FALSE |
| LOC107072122 | transmembrane GTPase Marf | 0.02109 | 83.8057 | FALSE | FALSE |
| LOC107063696 | uncharacterized protein PF11_0213 | 0.0203 | 83.85168 | TRUE | FALSE |
| LOC107074661 | neurotrimin-like | 0.02082 | 83.85513 | FALSE | FALSE |
| LOC107071218 | uncharacterized protein LOC107071218 | 0.02094 | 83.85537 | FALSE | FALSE |
| LOC107069385 | NAD kinase 2, mitochondrial-like | 0.02117 | 83.89445 | FALSE | FALSE |
| LOC107069462 | rRNA methyltransferase 2, mitochondrial | 0.02197 | 83.932 | TRUE | FALSE |

| | | | | | |
|---|---|---|---|---|---|
| LOC107072875 | dual oxidase | 0.02211 | 83.94255 | FALSE | FALSE |
| LOC107064292 | integrator complex subunit 8 | 0.02068 | 83.94678 | TRUE | FALSE |
| LOC107069031 | ribosomal RNA-processing protein 8 | 0.01999 | 83.95099 | FALSE | FALSE |
| LOC107066876 | dual specificity mitogen-activated protein ki | 0.02115 | 83.95391 | TRUE | FALSE |
| LOC107069021 | uncharacterized protein LOC107069021 | 0.0215 | 83.96446 | FALSE | FALSE |
| LOC107073359 | uncharacterized protein LOC107073359 | 0.02003 | 83.97699 | FALSE | FALSE |
| LOC107073784 | RNA-binding protein 12B-like | 0.02029 | 83.97923 | TRUE | FALSE |
| LOC107074374 | G-protein-signaling modulator 2 | 0.02193 | 83.9969 | FALSE | FALSE |
| LOC107067952 | uncharacterized protein LOC107067952 | 0.02128 | 84.01256 | FALSE | FALSE |
| LOC107074552 | N-acetylglucosamine-6-sulfatase-like | 0.02073 | 84.05724 | FALSE | FALSE |
| LOC107067065 | rab proteins geranylgeranyltransferase com | 0.02061 | 84.07218 | TRUE | FALSE |
| LOC107073181 | UPF0585 protein C16orf13 homolog A | 0.02048 | 84.09742 | FALSE | FALSE |
| LOC107064034 | probable glutamate receptor | 0.01971 | 84.12143 | FALSE | FALSE |
| LOC107065477 | heat shock 70 kDa protein cognate 4-like | 0.02298 | 84.16679 | TRUE | FALSE |
| LOC107064003 | kinesin-associated protein 3 | 0.0216 | 84.1765 | FALSE | FALSE |
| LOC107066459 | palmitoyltransferase ZDHHC15 | 0.02119 | 84.1859 | FALSE | FALSE |
| LOC107074083 | ankyrin repeat and zinc finger domain-conta | 0.02113 | 84.19135 | FALSE | FALSE |
| LOC107070733 | E3 ubiquitin-protein ligase LRSAM1-like | 0.01985 | 84.2203 | TRUE | FALSE |
| LOC107070087 | splicing factor 3A subunit 1 | 0.02066 | 84.22896 | FALSE | FALSE |
| LOC107071711 | troponin C, isoform 2-like | 0.02182 | 84.23003 | TRUE | FALSE |
| LOC107065110 | trifunctional enzyme subunit alpha, mitocho | 0.02146 | 84.27649 | FALSE | FALSE |
| LOC107072857 | prolyl 3-hydroxylase 1-like | 0.0225 | 84.27941 | FALSE | FALSE |
| LOC107068841 | transmembrane protein 17B-like | 0.02093 | 84.29445 | FALSE | FALSE |
| LOC107065869 | exosome complex component MTR3 | 0.02051 | 84.33151 | FALSE | FALSE |
| LOC107069952 | coiled-coil domain-containing protein 28A | 0.02142 | 84.34844 | FALSE | FALSE |
| LOC107072605 | pickpocket protein 11-like | 0.02051 | 84.39593 | FALSE | FALSE |
| LOC107067862 | endoplasmic reticulum metallopeptidase 1- | 0.02101 | 84.40827 | FALSE | FALSE |
| LOC107070603 | gamma-secretase subunit pen-2 | 0.02016 | 84.42729 | FALSE | FALSE |
| LOC107073686 | uncharacterized protein LOC107073686 | 0.02237 | 84.43199 | FALSE | FALSE |
| LOC107073987 | diacylglycerol kinase 1 | 0.02108 | 84.46554 | FALSE | FALSE |
| LOC107069456 | uncharacterized protein LOC107069456 | 0.02155 | 84.49548 | FALSE | FALSE |
| LOC107072232 | uncharacterized protein LOC107072232 | 0.02084 | 84.49984 | FALSE | FALSE |
| LOC107070147 | uncharacterized protein LOC107070147 | 0.02213 | 84.50235 | FALSE | FALSE |
| LOC107066006 | cytochrome c oxidase subunit 6A1, mitocho | 0.02158 | 84.50695 | FALSE | FALSE |
| LOC107071061 | eukaryotic translation initiation factor 5A | 0.02056 | 84.54254 | FALSE | FALSE |
| LOC107074091 | cysteine sulfinic acid decarboxylase-like | 0.02002 | 84.54599 | FALSE | FALSE |
| LOC107070340 | NADH dehydrogenase [ubiquinone | 0.02126 | 84.57303 | FALSE | FALSE |
| LOC107067023 | uncharacterized protein LOC107067023 | 0.02077 | 84.58023 | FALSE | FALSE |
| LOC107073896 | protein prenyltransferase alpha subunit rep | 0.02025 | 84.58078 | FALSE | FALSE |
| LOC107070372 | coiled-coil domain-containing protein 42 ho | 0.02129 | 84.59037 | FALSE | FALSE |
| LOC107063814 | LMBR1 domain-containing protein 2 homolo | 0.02132 | 84.61184 | TRUE | FALSE |
| LOC107071805 | mRNA turnover protein 4 homolog | 0.02099 | 84.63919 | FALSE | FALSE |
| LOC107067306 | uncharacterized protein LOC107067306 | 0.02115 | 84.63973 | TRUE | FALSE |
| LOC107068024 | RIB43A-like with coiled-coils protein 2 | 0.0209 | 84.64829 | FALSE | FALSE |
| LOC107069828 | UPF0692 protein CG33108 | 0.02055 | 84.65462 | FALSE | FALSE |
| LOC107066840 | ras-related protein Rab-23-like | 0.02174 | 84.66949 | FALSE | FALSE |
| LOC107069586 | uncharacterized protein LOC107069586 | 0.02002 | 84.68219 | TRUE | FALSE |
| LOC107073875 | probable serine hydrolase | 0.02122 | 84.70556 | FALSE | FALSE |
| LOC107066632 | profilin | 0.02091 | 84.7315 | FALSE | FALSE |
| LOC107068478 | mediator of RNA polymerase II transcriptior | 0.02091 | 84.73804 | FALSE | FALSE |

| | | | | | |
|---|---|---|---|---|---|
| LOC107071687 | putative uncharacterized protein DDB_G028 | 0.02089 | 84.75665 | FALSE | FALSE |
| LOC107069849 | SUMO-activating enzyme subunit 1 | 0.02022 | 84.77984 | FALSE | FALSE |
| LOC107074340 | E3 ubiquitin-protein ligase KCMF1-like | 0.02218 | 84.80885 | TRUE | FALSE |
| LOC107071882 | uncharacterized protein LOC107071882 | 0.02125 | 84.85119 | FALSE | FALSE |
| LOC107068487 | protein PRRC1-like | 0.02142 | 84.90467 | FALSE | FALSE |
| LOC107070239 | E3 ubiquitin-protein ligase hyd | 0.02041 | 84.90702 | TRUE | FALSE |
| LOC107064219 | uncharacterized protein LOC107064219 | 0.02107 | 84.92387 | FALSE | FALSE |
| LOC107071518 | helicase POLQ-like | 0.02177 | 84.92736 | TRUE | FALSE |
| LOC107071327 | putative leucine-rich repeat-containing prot | 0.02062 | 84.95126 | TRUE | FALSE |
| LOC107068865 | cytochrome P450 18a1 | 0.02087 | 84.95227 | FALSE | FALSE |
| LOC107074452 | inositol polyphosphate 5-phosphatase OCRL | 0.02051 | 84.96966 | FALSE | FALSE |
| LOC107069880 | actin-like protein 6B | 0.02033 | 84.98607 | FALSE | FALSE |
| LOC107067335 | uncharacterized protein LOC107067335 | 0.02219 | 85.00722 | FALSE | TRUE |
| LOC107065739 | probable serine/threonine-protein kinase rc | 0.02181 | 85.04164 | TRUE | FALSE |
| LOC107070397 | uncharacterized protein LOC107070397 | 0.0214 | 85.05868 | FALSE | FALSE |
| LOC107072002 | nucleolar GTP-binding protein 1 | 0.02016 | 85.0597 | FALSE | FALSE |
| LOC107068887 | guanine nucleotide exchange factor DBS-like | 0.02072 | 85.11008 | TRUE | FALSE |
| LOC107067590 | protein lin-7 homolog C | 0.02022 | 85.11364 | FALSE | FALSE |
| LOC107068920 | uncharacterized protein LOC107068920 | 0.02283 | 85.11672 | TRUE | FALSE |
| LOC107067482 | 28S ribosomal protein S29, mitochondrial | 0.02219 | 85.15568 | FALSE | FALSE |
| LOC107074411 | solute carrier family 35 member G1-like | 0.0213 | 85.17218 | TRUE | FALSE |
| LOC107068355 | aryl-hydrocarbon-interacting protein-like 1 | 0.02046 | 85.20216 | TRUE | FALSE |
| LOC107063649 | cell cycle checkpoint protein RAD17 | 0.02097 | 85.21048 | FALSE | FALSE |
| LOC107065285 | major facilitator superfamily domain-contai | 0.02065 | 85.21174 | FALSE | FALSE |
| LOC107069715 | nuclear pore complex protein Nup93-like | 0.02182 | 85.21865 | TRUE | FALSE |
| LOC107065370 | uncharacterized protein LOC107065370 | 0.02074 | 85.25355 | FALSE | FALSE |
| LOC107064074 | guanylate cyclase 32E-like | 0.02048 | 85.26499 | TRUE | FALSE |
| LOC107070375 | jmjC domain-containing histone demethylat | 0.02156 | 85.29709 | FALSE | FALSE |
| LOC107068151 | uncharacterized protein LOC107068151 | 0.01991 | 85.30384 | FALSE | FALSE |
| LOC107065303 | uncharacterized protein LOC107065303 | 0.02102 | 85.32929 | FALSE | FALSE |
| LOC107065291 | uncharacterized protein LOC107065291 | 0.02193 | 85.37052 | FALSE | FALSE |
| LOC107064868 | protein FAM151B | 0.02172 | 85.3771 | TRUE | FALSE |
| LOC107071289 | uncharacterized protein LOC107071289 | 0.01993 | 85.38541 | FALSE | FALSE |
| LOC107071434 | U11/U12 small nuclear ribonucleoprotein 25 | 0.02479 | 85.38663 | FALSE | FALSE |
| LOC107071174 | lisH domain-containing protein ARMC9-like | 0.02383 | 85.40709 | TRUE | FALSE |
| LOC107070761 | probable RNA helicase armi | 0.02195 | 85.42437 | TRUE | FALSE |
| LOC107063748 | E3 ubiquitin-protein ligase synoviolin A | 0.02051 | 85.44133 | FALSE | FALSE |
| LOC107065761 | histone acetyltransferase KAT2A | 0.0212 | 85.46635 | TRUE | FALSE |
| LOC107069894 | proton-coupled amino acid transporter 1-lik | 0.02062 | 85.47772 | FALSE | FALSE |
| LOC107069491 | SPARC | 0.02169 | 85.52837 | FALSE | FALSE |
| LOC107068963 | histone-lysine N-methyltransferase SETMAR | 0.0203 | 85.53881 | FALSE | FALSE |
| LOC107064115 | biogenesis of lysosome-related organelles c | 0.02171 | 85.54399 | FALSE | FALSE |
| LOC107073389 | uncharacterized protein LOC107073389 | 0.0217 | 85.55791 | FALSE | FALSE |
| LOC107072882 | small VCP/p97-interacting protein | 0.02123 | 85.60136 | FALSE | FALSE |
| LOC107064624 | vitamin K-dependent gamma-carboxylase | 0.02063 | 85.61381 | TRUE | FALSE |
| LOC107070222 | uncharacterized protein LOC107070222 | 0.02065 | 85.66382 | FALSE | FALSE |
| LOC107069367 | uncharacterized protein LOC107069367 | 0.0213 | 85.67301 | FALSE | FALSE |
| LOC107071741 | uncharacterized protein LOC107071741 | 0.02147 | 85.70232 | FALSE | FALSE |
| LOC107072228 | partner of Y14 and mago | 0.02033 | 85.71632 | FALSE | FALSE |
| LOC107068528 | uncharacterized protein LOC107068528 | 0.02165 | 85.719 | TRUE | FALSE |

| | | | | | |
|---|---|---|---|---|---|
| LOC107073453 | uncharacterized protein LOC107073453 | 0.02034 | 85.74069 | FALSE | FALSE |
| LOC107067685 | uncharacterized protein LOC107067685 | 0.02099 | 85.76509 | FALSE | FALSE |
| LOC107065827 | NADH-ubiquinone oxidoreductase 49 kDa su | 0.02083 | 85.77389 | FALSE | FALSE |
| LOC107071985 | putative ATP-dependent RNA helicase DHX3 | 0.02097 | 85.79942 | TRUE | FALSE |
| LOC107071452 | uncharacterized protein LOC107071452 | 0.02167 | 85.80113 | TRUE | FALSE |
| LOC107066112 | mannosyl-oligosaccharide 1,2-alpha-manno | 0.02191 | 85.81463 | TRUE | FALSE |
| LOC107067830 | hemolymph lipopolysaccharide-binding prot | 0.02298 | 85.8309 | FALSE | FALSE |
| LOC107067235 | uncharacterized protein LOC107067235 | 0.02099 | 85.85748 | TRUE | FALSE |
| LOC107070109 | huntingtin-interacting protein K | 0.02001 | 85.87222 | TRUE | FALSE |
| LOC107063970 | protein gooseberry-like | 0.02183 | 85.87532 | FALSE | FALSE |
| LOC107071776 | intraflagellar transport protein 122 homolog | 0.02733 | 85.8894 | TRUE | FALSE |
| LOC107067912 | uncharacterized protein LOC107067912 | 0.02196 | 85.89693 | FALSE | FALSE |
| LOC107070278 | ras-related protein Rab-28-like | 0.0206 | 85.91787 | FALSE | FALSE |
| LOC107070754 | PRELI domain-containing protein 1, mitocho | 0.02122 | 85.93226 | FALSE | FALSE |
| LOC107071825 | uncharacterized protein LOC107071825 | 0.02199 | 85.95711 | FALSE | FALSE |
| LOC107065509 | putative malate dehydrogenase 1B | 0.01988 | 85.97884 | FALSE | FALSE |
| LOC107067861 | zinc transporter 2-like | 0.02122 | 86.01678 | FALSE | FALSE |
| LOC107064284 | uncharacterized protein LOC107064284 | 0.02124 | 86.02258 | FALSE | FALSE |
| LOC107074156 | proline synthase co-transcribed bacterial ho | 0.02025 | 86.02602 | FALSE | FALSE |
| LOC107068270 | RRP15-like protein | 0.0204 | 86.0321 | FALSE | FALSE |
| LOC107069309 | E3 ubiquitin-protein ligase MYCBP2 | 0.02043 | 86.04047 | TRUE | FALSE |
| LOC107069671 | protein NipSnap | 0.02228 | 86.05521 | FALSE | FALSE |
| LOC107064594 | ubiquitin-40S ribosomal protein S27a | 0.02065 | 86.0667 | TRUE | FALSE |
| LOC107070017 | uncharacterized protein LOC107070017 | 0.02066 | 86.07237 | FALSE | FALSE |
| LOC107066085 | uncharacterized protein LOC107066085 | 0.02076 | 86.09692 | FALSE | FALSE |
| LOC107069101 | probable E3 ubiquitin-protein ligase HERC4 | 0.02218 | 86.10945 | FALSE | FALSE |
| LOC107064832 | trypsin-7-like | 0.02203 | 86.13407 | FALSE | FALSE |
| LOC107071306 | integral membrane protein GPR155 | 0.02187 | 86.14003 | TRUE | FALSE |
| LOC107066527 | dmX-like protein 2 | 0.01979 | 86.16553 | FALSE | FALSE |
| LOC107071631 | basic helix-loop-helix transcription factor sc | 0.02071 | 86.172 | TRUE | FALSE |
| LOC107065960 | ceramide synthase 5-like | 0.02068 | 86.17209 | FALSE | FALSE |
| LOC107073771 | uncharacterized protein LOC107073771 | 0.02031 | 86.19831 | FALSE | FALSE |
| LOC107069316 | uncharacterized protein LOC107069316 | 0.02053 | 86.21838 | FALSE | FALSE |
| LOC107068769 | uncharacterized protein LOC107068769 | 0.02034 | 86.23867 | FALSE | FALSE |
| LOC107068948 | late embryogenesis abundant protein, grou | 0.02139 | 86.24609 | FALSE | FALSE |
| LOC107071986 | major facilitator superfamily domain-contai | 0.02097 | 86.25073 | FALSE | FALSE |
| LOC107065819 | matrix metalloproteinase-14 | 0.02429 | 86.27828 | TRUE | FALSE |
| LOC107074097 | pinin | 0.02071 | 86.32714 | FALSE | FALSE |
| LOC107068947 | WD repeat-containing protein 35 | 0.02179 | 86.32931 | TRUE | FALSE |
| LOC107069210 | DDB1- and CUL4-associated factor 10 | 0.02139 | 86.3521 | FALSE | FALSE |
| LOC107072538 | TBC1 domain family member 13 | 0.02234 | 86.36693 | FALSE | FALSE |
| LOC107068183 | ATPase family AAA domain-containing prote | 0.0209 | 86.38285 | FALSE | FALSE |
| LOC107064895 | BTB/POZ domain-containing adapter for CU | 0.02039 | 86.39152 | FALSE | FALSE |
| LOC107071614 | inactive dipeptidyl peptidase 10 | 0.02061 | 86.40248 | TRUE | FALSE |
| LOC107065653 | metaxin-1 | 0.0212 | 86.40391 | FALSE | FALSE |
| LOC107064386 | protein dpy-30 homolog | 0.02258 | 86.4439 | TRUE | FALSE |
| Trnag-gcc | Trnag-gcc | 0.02022 | 86.44557 | FALSE | FALSE |
| LOC107069289 | uncharacterized protein LOC107069289 | 0.01994 | 86.45272 | FALSE | FALSE |
| LOC107065518 | headcase protein | 0.02162 | 86.46597 | TRUE | FALSE |
| LOC107071365 | zinc finger protein GLI1 | 0.0213 | 86.50546 | TRUE | FALSE |

| | | | | | |
|---|---|---|---|---|---|
| LOC107072820 | uncharacterized protein LOC107072820 | 0.02042 | 86.54002 | FALSE | FALSE |
| LOC107073524 | uncharacterized protein LOC107073524 | 0.02055 | 86.55217 | TRUE | FALSE |
| LOC107067626 | 40S ribosomal protein S4 | 0.02053 | 86.56459 | FALSE | FALSE |
| LOC107072498 | uncharacterized protein LOC107072498 | 0.02193 | 86.57292 | FALSE | FALSE |
| LOC107071601 | uncharacterized protein LOC107071601 | 0.02231 | 86.58182 | FALSE | FALSE |
| LOC107074679 | KAT8 regulatory NSL complex subunit 3 | 0.02273 | 86.63767 | TRUE | FALSE |
| LOC107068667 | otopetrin-1-like | 0.0206 | 86.67076 | FALSE | FALSE |
| LOC107064586 | multidrug resistance-associated protein 7 | 0.02149 | 86.68369 | TRUE | FALSE |
| LOC107064664 | inner centromere protein-like | 0.02018 | 86.70518 | FALSE | FALSE |
| LOC107067507 | uncharacterized protein LOC107067507 | 0.02227 | 86.71068 | TRUE | FALSE |
| LOC107066486 | uncharacterized protein CG4449 | 0.02092 | 86.78366 | FALSE | FALSE |
| LOC107069977 | uncharacterized protein LOC107069977 | 0.02023 | 86.82111 | TRUE | FALSE |
| LOC107068883 | uncharacterized protein LOC107068883 | 0.02026 | 86.82978 | FALSE | FALSE |
| LOC107070790 | heat shock protein 83 | 0.02088 | 86.85014 | FALSE | FALSE |
| LOC107070856 | uncharacterized protein LOC107070856 | 0.02117 | 86.85458 | FALSE | FALSE |
| LOC107064768 | uncharacterized protein LOC107064768 | 0.0209 | 86.86263 | TRUE | FALSE |
| LOC107069271 | uncharacterized protein LOC107069271 | 0.02184 | 86.87269 | FALSE | FALSE |
| LOC107065271 | putative sulfiredoxin | 0.02019 | 86.89181 | FALSE | FALSE |
| LOC107074062 | probable serine/threonine-protein kinase ki | 0.02065 | 86.90973 | FALSE | FALSE |
| LOC107063901 | glucose dehydrogenase [FAD, quinone | 0.02107 | 86.91814 | FALSE | FALSE |
| LOC107071935 | uncharacterized protein LOC107071935 | 0.02036 | 86.92522 | TRUE | FALSE |
| LOC107068619 | U1 small nuclear ribonucleoprotein 70 kDa | 0.0227 | 86.92561 | FALSE | FALSE |
| LOC107070718 | sushi, von Willebrand factor type A, EGF anc | 0.0211 | 86.94087 | FALSE | FALSE |
| LOC107063825 | uncharacterized protein LOC107063825 | 0.02144 | 86.94649 | TRUE | FALSE |
| LOC107069251 | zinc finger protein ush | 0.02062 | 86.96136 | TRUE | FALSE |
| LOC107064054 | dehydrogenase/reductase SDR family meml | 0.02045 | 87.02941 | FALSE | FALSE |
| LOC107067387 | cytochrome b5-like | 0.02096 | 87.03389 | FALSE | FALSE |
| LOC107068166 | glycylpeptide N-tetradecanoyltransferase 1 | 0.02115 | 87.03576 | FALSE | FALSE |
| LOC107064176 | dihydrolipoyllysine-residue acetyltransferas | 0.02153 | 87.04341 | FALSE | FALSE |
| LOC107072684 | neurocalcin-delta B-like | 0.02236 | 87.05561 | TRUE | FALSE |
| LOC107071777 | uncharacterized protein LOC107071777 | 0.02033 | 87.08083 | TRUE | FALSE |
| LOC107069393 | glutamine-rich protein 2-like | 0.02043 | 87.14672 | FALSE | FALSE |
| LOC107066957 | transposable element Tc1 transposase | 0.02014 | 87.16844 | FALSE | FALSE |
| LOC107066903 | uncharacterized protein LOC107066903 | 0.02298 | 87.19093 | TRUE | FALSE |
| LOC107068312 | glutamate receptor-interacting protein 1 | 0.02103 | 87.19097 | TRUE | FALSE |
| LOC107070452 | protein neuralized | 0.02056 | 87.2471 | FALSE | FALSE |
| LOC107063990 | heparan-sulfate 6-O-sulfotransferase 2 | 0.02112 | 87.26224 | FALSE | FALSE |
| LOC107067832 | 6-phosphogluconolactonase | 0.02109 | 87.27414 | FALSE | FALSE |
| LOC107069247 | Niemann-Pick C1 protein | 0.02081 | 87.28161 | FALSE | FALSE |
| LOC107063859 | receptor expression-enhancing protein 5-lik | 0.02022 | 87.32706 | FALSE | FALSE |
| LOC107073159 | sodium/potassium-transporting ATPase sub | 0.0226 | 87.34866 | TRUE | FALSE |
| LOC107070569 | alpha-methylacyl-CoA racemase | 0.02075 | 87.36008 | FALSE | FALSE |
| LOC107072408 | uncharacterized protein LOC107072408 | 0.02004 | 87.36116 | FALSE | FALSE |
| LOC107065329 | mitochondrial pyruvate carrier 1 | 0.0221 | 87.38483 | TRUE | FALSE |
| LOC107070070 | solute carrier organic anion transporter fam | 0.02193 | 87.40835 | TRUE | FALSE |
| LOC107071771 | nucleoside diphosphate kinase | 0.02142 | 87.44929 | FALSE | FALSE |
| LOC107065714 | phosphorylated CTD-interacting factor 1 | 0.02142 | 87.46239 | FALSE | FALSE |
| LOC107072414 | microtubule-associated proteins 1A/1B light | 0.02355 | 87.46683 | TRUE | FALSE |
| LOC107070922 | uncharacterized protein LOC107070922 | 0.02342 | 87.48194 | TRUE | FALSE |
| LOC107073496 | prothrombin-like | 0.02207 | 87.48957 | FALSE | FALSE |

| | | | | | |
|---|---|---|---|---|---|
| LOC107064174 | carboxypeptidase N subunit 2-like | 0.02052 | 87.49835 | FALSE | FALSE |
| LOC107068086 | glutathione S-transferase theta-1 | 0.01999 | 87.54869 | FALSE | FALSE |
| LOC107070644 | NADH dehydrogenase [ubiquinone | 0.02082 | 87.55089 | FALSE | FALSE |
| LOC107074186 | kinesin-like protein KIF3A | 0.02553 | 87.62999 | TRUE | FALSE |
| LOC107067403 | uncharacterized protein LOC107067403 | 0.02044 | 87.6401 | FALSE | FALSE |
| LOC107072756 | uncharacterized protein LOC107072756 | 0.02042 | 87.68126 | FALSE | FALSE |
| LOC107073334 | uncharacterized protein LOC107073334 | 0.02223 | 87.68258 | TRUE | FALSE |
| LOC107073225 | uncharacterized protein LOC107073225 | 0.0196 | 87.70986 | FALSE | FALSE |
| LOC107065396 | protein starmaker | 0.02078 | 87.71297 | FALSE | FALSE |
| LOC107072423 | enhancer of mRNA-decapping protein 3 | 0.02104 | 87.71381 | FALSE | FALSE |
| LOC107065398 | uncharacterized protein LOC107065398 | 0.02071 | 87.73555 | FALSE | FALSE |
| LOC107069345 | uncharacterized protein LOC107069345 | 0.02166 | 87.74511 | TRUE | FALSE |
| LOC107070551 | putative mediator of RNA polymerase II trar | 0.0215 | 87.75031 | TRUE | FALSE |
| LOC107065879 | MAR-binding filament-like protein 1 | 0.0205 | 87.75706 | FALSE | FALSE |
| LOC107071336 | protein arginine N-methyltransferase 1 | 0.02188 | 87.77468 | FALSE | FALSE |
| LOC107071700 | serine-rich adhesin for platelets | 0.02133 | 87.77809 | TRUE | FALSE |
| LOC107066260 | protein mago nashi homolog | 0.02224 | 87.78927 | TRUE | FALSE |
| LOC107065648 | translation initiation factor eIF-2B subunit g | 0.0207 | 87.79026 | FALSE | FALSE |
| LOC107068537 | probable WRKY transcription factor protein | 0.02084 | 87.79893 | TRUE | FALSE |
| LOC107064160 | protein FAM192A | 0.02083 | 87.8224 | TRUE | FALSE |
| LOC107072703 | uncharacterized protein LOC107072703 | 0.0209 | 87.85853 | FALSE | FALSE |
| LOC107071788 | U6 snRNA-associated Sm-like protein LSm5 | 0.02119 | 87.8648 | FALSE | FALSE |
| LOC107064770 | uncharacterized protein LOC107064770 | 0.01993 | 87.87599 | TRUE | FALSE |
| LOC107068027 | vacuolar-sorting protein SNF8 | 0.02129 | 87.88028 | FALSE | FALSE |
| LOC107068137 | putative golgin subfamily A member 6-like p | 0.0211 | 87.88106 | TRUE | FALSE |
| LOC107071308 | myb-like protein X | 0.02127 | 87.88875 | TRUE | FALSE |
| LOC107071934 | sodium-dependent neutral amino acid trans | 0.02136 | 87.90285 | TRUE | FALSE |
| LOC107071850 | voltage-dependent calcium channel type A s | 0.01986 | 87.906 | FALSE | FALSE |
| LOC107066432 | uncharacterized protein LOC107066432 | 0.02034 | 87.94739 | FALSE | FALSE |
| LOC107068869 | farnesol dehydrogenase-like | 0.02123 | 87.94872 | FALSE | FALSE |
| LOC107071877 | leucine-rich repeat-containing protein 24-lik | 0.02137 | 87.96464 | FALSE | FALSE |
| LOC107064476 | GRAM domain-containing protein 1B-like | 0.0208 | 87.97738 | FALSE | FALSE |
| LOC107068249 | dnaJ homolog subfamily C member 16 | 0.02286 | 88.00895 | TRUE | FALSE |
| LOC107066394 | uncharacterized protein LOC107066394 | 0.02358 | 88.05377 | TRUE | FALSE |
| LOC107068254 | uncharacterized protein LOC107068254 | 0.0217 | 88.05811 | FALSE | FALSE |
| LOC107065886 | ubiquitin-like modifier-activating enzyme 1 | 0.02024 | 88.05944 | TRUE | FALSE |
| LOC107064407 | ileal sodium/bile acid cotransporter-like | 0.01998 | 88.09459 | FALSE | FALSE |
| LOC107070037 | uncharacterized protein LOC107070037 | 0.022 | 88.12271 | TRUE | FALSE |
| LOC107067984 | uncharacterized protein LOC107067984 | 0.02164 | 88.14719 | TRUE | FALSE |
| LOC107068342 | ubiquitin-conjugating enzyme E2 R2 | 0.02061 | 88.24289 | TRUE | FALSE |
| LOC107066426 | uncharacterized protein LOC107066426 | 0.02176 | 88.24824 | TRUE | FALSE |
| LOC107070133 | netrin-1-like | 0.02128 | 88.26869 | TRUE | FALSE |
| LOC107067104 | importin subunit beta-1 | 0.02113 | 88.27071 | FALSE | FALSE |
| LOC107072065 | endoplasmic reticulum resident protein 29 | 0.02062 | 88.28236 | TRUE | FALSE |
| LOC107071635 | dual specificity protein phosphatase 19 | 0.02017 | 88.29434 | FALSE | FALSE |
| LOC107071489 | eukaryotic translation initiation factor 3 sub | 0.02037 | 88.31419 | FALSE | FALSE |
| LOC107067794 | uncharacterized protein LOC107067794 | 0.02033 | 88.32061 | FALSE | FALSE |
| LOC107065700 | leucine-rich repeat and death domain-conta | 0.02092 | 88.35687 | TRUE | FALSE |
| LOC107065617 | ribonuclease H1 | 0.02073 | 88.36005 | FALSE | FALSE |
| LOC107069350 | syntaxin-17 | 0.02149 | 88.4147 | FALSE | FALSE |

| | | | | | |
|---|---|---|---|---|---|
| LOC107071270 | uncharacterized protein LOC107071270 | 0.02039 | 88.43079 | FALSE | FALSE |
| LOC107067149 | uncharacterized protein DDB_G0271670 | 0.02025 | 88.44393 | FALSE | FALSE |
| LOC107063923 | pyruvate carboxylase, mitochondrial | 0.02024 | 88.46573 | FALSE | FALSE |
| LOC107064966 | BET1 homolog | 0.02065 | 88.47812 | FALSE | FALSE |
| LOC107068598 | transmembrane and TPR repeat-containing | 0.02032 | 88.48598 | TRUE | FALSE |
| LOC107065875 | zinc finger protein 2 homolog | 0.02062 | 88.48832 | TRUE | FALSE |
| LOC107063703 | uncharacterized protein LOC107063703 | 0.0202 | 88.51062 | FALSE | FALSE |
| LOC107066246 | transcription factor E2F5 | 0.02547 | 88.51818 | TRUE | FALSE |
| LOC107067741 | uncharacterized protein LOC107067741 | 0.02094 | 88.52291 | TRUE | FALSE |
| LOC107070104 | uncharacterized protein LOC107070104 | 0.02069 | 88.56235 | TRUE | FALSE |
| LOC107067705 | protein orai-2-like | 0.02098 | 88.60514 | FALSE | FALSE |
| LOC107071071 | uncharacterized protein LOC107071071 | 0.02083 | 88.60881 | TRUE | FALSE |
| LOC107064598 | leucine-rich repeat transmembrane neurona | 0.02096 | 88.67056 | FALSE | FALSE |
| LOC107064434 | protein BCL9 homolog | 0.02224 | 88.68227 | TRUE | FALSE |
| LOC107073606 | uncharacterized protein LOC107073606 | 0.02069 | 88.69499 | FALSE | FALSE |
| LOC107071208 | syntaxin-1A | 0.02181 | 88.69511 | FALSE | FALSE |
| LOC107074484 | uncharacterized protein LOC107074484 | 0.02184 | 88.74886 | TRUE | FALSE |
| LOC107067645 | malate dehydrogenase, cytoplasmic-like | 0.02074 | 88.77049 | TRUE | FALSE |
| LOC107063796 | nuclear pore complex protein Nup155 | 0.02076 | 88.79762 | FALSE | FALSE |
| LOC107072690 | PHD finger protein rhinoceros | 0.02065 | 88.80921 | TRUE | FALSE |
| LOC107073932 | methyltransferase-like protein 23 | 0.02115 | 88.81618 | FALSE | FALSE |
| LOC107064223 | cytochrome b5 domain-containing protein 1 | 0.02091 | 88.85173 | FALSE | FALSE |
| LOC107071330 | protein GDAP2 homolog | 0.02287 | 88.85416 | TRUE | FALSE |
| LOC107073189 | growth hormone-inducible transmembrane | 0.0207 | 88.85429 | TRUE | FALSE |
| LOC107072075 | glycosyltransferase 25 family member-like | 0.021 | 88.85782 | TRUE | FALSE |
| LOC107066357 | immediate early response 3-interacting pro | 0.02125 | 88.85917 | FALSE | FALSE |
| LOC107066592 | interferon regulatory factor 2-binding prote | 0.02032 | 88.90464 | TRUE | FALSE |
| LOC107072850 | splicing factor 3B subunit 3 | 0.02152 | 88.91858 | FALSE | FALSE |
| LOC107074003 | uncharacterized protein LOC107074003 | 0.0207 | 88.91937 | FALSE | FALSE |
| LOC107064690 | peripheral plasma membrane protein CASK | 0.0212 | 88.92464 | TRUE | FALSE |
| LOC107064809 | uncharacterized protein LOC107064809 | 0.02099 | 88.94015 | FALSE | FALSE |
| LOC107064819 | LOW QUALITY PROTEIN: tubulin gamma-1 c | 0.02235 | 88.96645 | FALSE | FALSE |
| LOC107072787 | coiled-coil domain-containing protein 50 | 0.02157 | 88.99146 | TRUE | FALSE |
| LOC107072553 | transmembrane protein 192 | 0.02098 | 89.00366 | TRUE | FALSE |
| LOC107064177 | 39S ribosomal protein L13, mitochondrial | 0.02103 | 89.01379 | TRUE | FALSE |
| LOC107067991 | bromodomain-containing protein 8 | 0.02184 | 89.03254 | TRUE | FALSE |
| LOC107069259 | tetratricopeptide repeat protein 8 | 0.02173 | 89.03887 | TRUE | FALSE |
| LOC107065249 | inorganic pyrophosphatase | 0.02045 | 89.06317 | FALSE | FALSE |
| LOC107069869 | dual specificity protein phosphatase CDC14 | 0.02098 | 89.06371 | TRUE | FALSE |
| LOC107067979 | high affinity cAMP-specific and IBMX-insens | 0.02189 | 89.07419 | TRUE | FALSE |
| LOC107074682 | sodium- and chloride-dependent GABA tran | 0.0207 | 89.07856 | TRUE | FALSE |
| LOC107065847 | SPRY domain-containing SOCS box protein 3 | 0.0226 | 89.0825 | TRUE | FALSE |
| LOC107067788 | uncharacterized protein LOC107067788 | 0.01989 | 89.10824 | TRUE | FALSE |
| LOC107065046 | lysine-specific demethylase lid | 0.02065 | 89.11036 | TRUE | FALSE |
| LOC107065393 | sideroflexin-1-like | 0.02158 | 89.1198 | TRUE | FALSE |
| LOC107065156 | synaptic vesicle membrane protein VAT-1 h | 0.02288 | 89.12103 | TRUE | FALSE |
| LOC107064154 | MICOS complex subunit mic25-a | 0.02065 | 89.12761 | TRUE | FALSE |
| LOC107063904 | neural cell adhesion molecule 2 | 0.02062 | 89.14766 | TRUE | FALSE |
| LOC107070114 | pyroglutamyl-peptidase 1 | 0.02008 | 89.20799 | TRUE | FALSE |
| LOC107067308 | uncharacterized protein LOC107067308 | 0.02241 | 89.2254 | FALSE | FALSE |

| | | | | | |
|---|---|---|---|---|---|
| LOC107064468 | uncharacterized protein LOC107064468 | 0.02053 | 89.23499 | TRUE | FALSE |
| LOC107065628 | kinesin light chain | 0.02257 | 89.24358 | TRUE | FALSE |
| LOC107068766 | F-box only protein 9 | 0.02193 | 89.25808 | FALSE | FALSE |
| LOC107068241 | ABC transporter G family member 20 | 0.0204 | 89.25931 | FALSE | FALSE |
| LOC107072036 | uncharacterized protein LOC107072036 | 0.02063 | 89.26211 | TRUE | FALSE |
| LOC107065231 | uncharacterized protein LOC107065231 | 0.01972 | 89.26434 | TRUE | FALSE |
| LOC107074604 | solute carrier family 35 member F5 | 0.02092 | 89.2851 | TRUE | FALSE |
| LOC107066935 | cyclin-dependent kinase 10 | 0.02027 | 89.3008 | FALSE | FALSE |
| LOC107067412 | uncharacterized protein LOC107067412 | 0.0207 | 89.31135 | TRUE | FALSE |
| LOC107064860 | uncharacterized protein DDB_G0283697-lik | 0.02004 | 89.31499 | TRUE | FALSE |
| LOC107065210 | DNA repair protein RAD51 homolog 4-like | 0.02073 | 89.32444 | TRUE | FALSE |
| LOC107071271 | uncharacterized protein LOC107071271 | 0.02144 | 89.33874 | TRUE | FALSE |
| LOC107065820 | O-glucosyltransferase rumi homolog | 0.02567 | 89.35105 | TRUE | FALSE |
| LOC107069132 | BTB/POZ domain-containing protein 9 | 0.02213 | 89.3793 | TRUE | FALSE |
| LOC107066972 | THUMP domain-containing protein 1 | 0.02082 | 89.4353 | FALSE | FALSE |
| LOC107065064 | D-beta-hydroxybutyrate dehydrogenase, mi | 0.02014 | 89.45778 | FALSE | FALSE |
| LOC107066729 | uncharacterized protein LOC107066729 | 0.02159 | 89.46197 | FALSE | FALSE |
| LOC107073526 | uncharacterized protein LOC107073526 | 0.02076 | 89.4667 | TRUE | FALSE |
| LOC107066093 | ubiquitin-conjugating enzyme E2 Q2 | 0.02026 | 89.50299 | TRUE | FALSE |
| LOC107070343 | protein tramtrack, beta isoform-like | 0.02387 | 89.54138 | TRUE | FALSE |
| LOC107066506 | putative hydroxypyruvate isomerase | 0.02194 | 89.54593 | FALSE | FALSE |
| LOC107067091 | nucleolar protein 58 | 0.02125 | 89.55193 | FALSE | FALSE |
| LOC107073870 | peptidyl-prolyl cis-trans isomerase FKBP8 | 0.02097 | 89.65458 | TRUE | FALSE |
| LOC107072446 | protein MIS12 homolog | 0.02036 | 89.65777 | FALSE | FALSE |
| LOC107071401 | formin-binding protein 1-like | 0.0203 | 89.66478 | FALSE | FALSE |
| LOC107064382 | uncharacterized protein LOC107064382 | 0.02079 | 89.7008 | FALSE | FALSE |
| LOC107072030 | 3-hydroxyacyl-CoA dehydrogenase type-2-li | 0.0215 | 89.74443 | TRUE | FALSE |
| LOC107068579 | signal recognition particle receptor subunit | 0.02008 | 89.77517 | TRUE | FALSE |
| LOC107070750 | uncharacterized protein DDB_G0286299-lik | 0.02452 | 89.78979 | TRUE | FALSE |
| LOC107068576 | synaptotagmin-5 | 0.02183 | 89.82076 | FALSE | FALSE |
| LOC107063941 | acyl-CoA Delta(11) desaturase | 0.02006 | 89.83208 | TRUE | FALSE |
| LOC107072436 | bromodomain-containing protein DDB_G02 | 0.02122 | 89.841 | TRUE | FALSE |
| LOC107073804 | ras-related protein Rab-2 | 0.02101 | 89.84966 | TRUE | FALSE |
| LOC107073311 | adenylate cyclase type 2 | 0.02542 | 89.8846 | TRUE | FALSE |
| LOC107072759 | uncharacterized protein LOC107072759 | 0.02066 | 89.89653 | FALSE | FALSE |
| LOC107072241 | homeobox protein invected-like | 0.02237 | 89.90024 | TRUE | FALSE |
| LOC107069334 | uncharacterized protein LOC107069334 | 0.02051 | 89.9076 | TRUE | FALSE |
| LOC107071685 | twisted gastrulation protein homolog 1-A-lil | 0.01991 | 89.91858 | TRUE | FALSE |
| LOC107070099 | uncharacterized protein LOC107070099 | 0.02063 | 89.92994 | TRUE | FALSE |
| LOC107071829 | homeobox protein extradenticle-like | 0.02115 | 89.93709 | TRUE | FALSE |
| LOC107072504 | uncharacterized protein LOC107072504 | 0.02072 | 89.95259 | TRUE | FALSE |
| LOC107070671 | uncharacterized protein LOC107070671 | 0.02208 | 89.96197 | FALSE | FALSE |
| LOC107068777 | proclotting enzyme-like | 0.02062 | 89.96325 | FALSE | FALSE |
| LOC107068779 | lysophospholipid acyltransferase 7 | 0.0215 | 89.9871 | TRUE | FALSE |
| LOC107063778 | troponin C-like | 0.01992 | 90.00256 | TRUE | FALSE |
| LOC107074641 | NACHT domain- and WD repeat-containing | 0.02244 | 90.00712 | TRUE | FALSE |
| LOC107072461 | putative fatty acyl-CoA reductase CG5065 | 0.02249 | 90.01091 | TRUE | FALSE |
| LOC107072879 | testis-specific serine/threonine-protein kina | 0.02547 | 90.04814 | TRUE | FALSE |
| LOC107072672 | uncharacterized protein LOC107072672 | 0.02126 | 90.11224 | TRUE | FALSE |
| LOC107068425 | protein SAAL1 | 0.02192 | 90.12595 | TRUE | FALSE |

| | | | | | |
|---|---|---|---|---|---|
| LOC107070362 | GRB2-associated-binding protein 2 | 0.02106 | 90.13249 | TRUE | FALSE |
| LOC107067396 | titin-like | 0.02313 | 90.14933 | TRUE | FALSE |
| LOC107065026 | uracil phosphoribosyltransferase homolog | 0.02122 | 90.15269 | FALSE | FALSE |
| LOC107068760 | homeobox protein Hox-B1a-like | 0.02016 | 90.18732 | TRUE | FALSE |
| LOC107071431 | dynein light chain Tctex-type 1 | 0.02249 | 90.19674 | TRUE | FALSE |
| LOC107064515 | uncharacterized protein LOC107064515 | 0.02148 | 90.22139 | TRUE | FALSE |
| LOC107065652 | acylphosphatase-1-like | 0.0254 | 90.26623 | TRUE | FALSE |
| LOC107070601 | ras-like protein family member 10B | 0.02024 | 90.29897 | TRUE | FALSE |
| LOC107068790 | mitochondrial import receptor subunit TOM | 0.01991 | 90.29982 | TRUE | FALSE |
| LOC107071473 | myb-related protein B | 0.02019 | 90.30198 | TRUE | FALSE |
| LOC107066163 | serine/threonine-protein kinase SMG1 | 0.02252 | 90.36434 | TRUE | FALSE |
| LOC107070536 | alkaline phosphatase 4-like | 0.02039 | 90.42235 | FALSE | FALSE |
| LOC107068445 | fibrous sheath-interacting protein 2 | 0.0201 | 90.43606 | FALSE | FALSE |
| LOC107066800 | protein SMG9 | 0.02108 | 90.44161 | TRUE | FALSE |
| LOC107073542 | uncharacterized protein LOC107073542 | 0.02178 | 90.49376 | TRUE | FALSE |
| LOC107070371 | bicaudal D-related protein homolog | 0.02046 | 90.59216 | TRUE | FALSE |
| LOC107070512 | G-box-binding factor-like | 0.02261 | 90.59956 | TRUE | FALSE |
| LOC107064544 | perilipin-1-like | 0.02217 | 90.74269 | TRUE | FALSE |
| LOC107065572 | T-lymphocyte activation antigen CD86-like | 0.021 | 90.7499 | FALSE | FALSE |
| LOC107070466 | protein shisa-5-like | 0.02063 | 90.761 | TRUE | FALSE |
| LOC107064720 | protein suppressor of forked | 0.02064 | 90.81655 | TRUE | FALSE |
| LOC107064230 | glutamate receptor U1-like | 0.02391 | 90.84344 | TRUE | FALSE |
| LOC107067523 | POU domain, class 3, transcription factor 2-l | 0.02087 | 90.84808 | TRUE | FALSE |
| LOC107066466 | uncharacterized protein LOC107066466 | 0.02042 | 90.94699 | FALSE | FALSE |
| LOC107069419 | uncharacterized protein LOC107069419 | 0.02506 | 90.9616 | TRUE | FALSE |
| LOC107074659 | beta-TrCP | 0.02341 | 90.96969 | TRUE | FALSE |
| LOC107063692 | uncharacterized protein LOC107063692 | 0.02222 | 90.99414 | TRUE | FALSE |
| LOC107066553 | probable tRNA N6-adenosine threonylcarba | 0.02224 | 91.02579 | TRUE | FALSE |
| LOC107065419 | COP9 signalosome complex subunit 1 | 0.0223 | 91.0445 | TRUE | FALSE |
| LOC107068684 | pancreatic lipase-related protein 2-like | 0.02174 | 91.05508 | TRUE | FALSE |
| LOC107072943 | uncharacterized protein LOC107072943 | 0.02291 | 91.13759 | TRUE | FALSE |
| LOC107066183 | DNA topoisomerase 2 | 0.02144 | 91.14566 | TRUE | FALSE |
| LOC107073097 | mitochondrial pyruvate carrier 2-like | 0.02004 | 91.18297 | TRUE | FALSE |
| LOC107072103 | uncharacterized protein LOC107072103 | 0.02176 | 91.19413 | TRUE | FALSE |
| LOC107069469 | uncharacterized protein LOC107069469 | 0.02056 | 91.19523 | TRUE | FALSE |
| LOC107064960 | uncharacterized protein LOC107064960 | 0.02006 | 91.24184 | TRUE | FALSE |
| LOC107073237 | ubiquitin fusion degradation protein 1 homo | 0.02141 | 91.25602 | TRUE | FALSE |
| LOC107070974 | uncharacterized protein LOC107070974 | 0.02118 | 91.25659 | TRUE | FALSE |
| LOC107073780 | uncharacterized protein LOC107073780 | 0.0215 | 91.27351 | TRUE | FALSE |
| LOC107068292 | mitochondrial import inner membrane trans | 0.01976 | 91.31045 | TRUE | FALSE |
| LOC107072031 | uncharacterized protein LOC107072031 | 0.02204 | 91.32999 | TRUE | FALSE |
| LOC107072519 | venom allergen 5-like | 0.02142 | 91.3411 | TRUE | FALSE |
| LOC107070160 | neurotrimin-like | 0.02159 | 91.35744 | FALSE | FALSE |
| LOC107074459 | uncharacterized protein LOC107074459 | 0.03996 | 91.36198 | TRUE | FALSE |
| LOC107070244 | hepatocyte growth factor-regulated tyrosin | 0.02019 | 91.38323 | TRUE | FALSE |
| LOC107073614 | transmembrane protein 183B-like | 0.02072 | 91.39443 | TRUE | FALSE |
| LOC107070864 | carbonic anhydrase 7-like | 0.02004 | 91.407 | FALSE | FALSE |
| LOC107065536 | fatty acid synthase-like | 0.02136 | 91.42083 | TRUE | FALSE |
| LOC107069617 | glutathione S-transferase 1, isoform C | 0.02205 | 91.46611 | TRUE | FALSE |
| LOC107073530 | uncharacterized protein LOC107073530 | 0.02043 | 91.47708 | FALSE | FALSE |

| | | | | | |
|---|---|---|---|---|---|
| LOC107074520 | G8 domain-containing protein DDB_G02863 | 0.02152 | 91.4827 | TRUE | FALSE |
| LOC107068654 | enolase | 0.0203 | 91.5019 | TRUE | FALSE |
| LOC107065312 | uncharacterized protein LOC107065312 | 0.02158 | 91.50204 | FALSE | FALSE |
| LOC107066221 | serine/threonine-protein phosphatase 6 reg | 0.0229 | 91.52792 | TRUE | FALSE |
| LOC107074002 | uncharacterized protein DDB_G0285917-lik | 0.02038 | 91.53372 | TRUE | FALSE |
| LOC107067173 | replication factor C subunit 2 | 0.02111 | 91.56026 | FALSE | FALSE |
| LOC107071507 | gamma-aminobutyric acid type B receptor s | 0.02671 | 91.6017 | TRUE | FALSE |
| LOC107069434 | NFU1 iron-sulfur cluster scaffold homolog, r | 0.01989 | 91.71062 | TRUE | FALSE |
| LOC107067903 | tektin-3-like | 0.02118 | 91.7148 | TRUE | FALSE |
| LOC107068201 | lissencephaly-1 homolog | 0.02067 | 91.73008 | TRUE | FALSE |
| LOC107074386 | uncharacterized protein LOC107074386 | 0.02187 | 91.80315 | TRUE | FALSE |
| LOC107064290 | E3 ubiquitin-protein ligase RNF8-like | 0.02046 | 91.81196 | TRUE | FALSE |
| LOC107072220 | uncharacterized protein LOC107072220 | 0.02157 | 91.86091 | TRUE | FALSE |
| LOC107074330 | protein CNPPD1 | 0.02042 | 91.87974 | TRUE | FALSE |
| LOC107068367 | ubiquitin-protein ligase E3C | 0.02114 | 91.88175 | TRUE | FALSE |
| LOC107069002 | presenilins-associated rhomboid-like protei | 0.02288 | 91.88327 | TRUE | FALSE |
| LOC107069546 | uncharacterized protein LOC107069546 | 0.02054 | 91.88427 | TRUE | FALSE |
| LOC107068184 | 26S proteasome non-ATPase regulatory sub | 0.02452 | 91.88904 | TRUE | FALSE |
| LOC107072694 | abhydrolase domain-containing protein 16A | 0.02239 | 91.90899 | TRUE | FALSE |
| LOC107069354 | Krueppel-like factor 7 | 0.02108 | 91.92568 | TRUE | FALSE |
| LOC107073375 | GPI transamidase component PIG-T | 0.02083 | 91.93307 | FALSE | FALSE |
| LOC107064538 | disks large homolog 5 | 0.02147 | 91.94526 | TRUE | FALSE |
| LOC107070412 | cationic amino acid transporter 3 | 0.02111 | 91.94925 | TRUE | FALSE |
| LOC107072418 | ubiquitin-conjugating enzyme E2 H | 0.02107 | 91.9636 | TRUE | FALSE |
| LOC107067305 | chaoptin-like | 0.02241 | 91.96782 | TRUE | FALSE |
| LOC107065079 | TPR-containing protein DDB_G0280363-like | 0.02073 | 91.97476 | TRUE | FALSE |
| LOC107072299 | uncharacterized protein LOC107072299 | 0.02329 | 91.97651 | TRUE | FALSE |
| LOC107068749 | fragile X mental retardation syndrome-relat | 0.02811 | 91.99485 | TRUE | FALSE |
| LOC107070709 | protein MCM10 homolog | 0.02103 | 91.99748 | FALSE | FALSE |
| LOC107073228 | zinc metalloproteinase nas-4-like | 0.01981 | 91.99907 | FALSE | FALSE |
| LOC107070724 | gem-associated protein 5 | 0.02698 | 92.0256 | TRUE | FALSE |
| LOC107065199 | testican-2 | 0.02618 | 92.03451 | TRUE | FALSE |
| LOC107066075 | cysteine-rich venom protein 6-like | 0.0205 | 92.05315 | FALSE | FALSE |
| LOC107065403 | trypsin-2-like | 0.02175 | 92.09514 | TRUE | FALSE |
| LOC107067822 | uncharacterized protein LOC107067822 | 0.02208 | 92.11388 | TRUE | FALSE |
| LOC107072074 | glycosyltransferase 25 family member-like | 0.0214 | 92.15188 | TRUE | FALSE |
| LOC107068438 | cyclin-dependent kinases regulatory subunit | 0.0206 | 92.17426 | FALSE | FALSE |
| LOC107064962 | host cell factor 1 | 0.02128 | 92.24276 | TRUE | FALSE |
| LOC107072597 | uncharacterized protein LOC107072597 | 0.02059 | 92.24532 | TRUE | FALSE |
| LOC107066259 | succinyl-CoA ligase [ADP/GDP-forming | 0.02243 | 92.24952 | TRUE | FALSE |
| LOC107069329 | gastrulation defective protein 1 homolog | 0.02083 | 92.2804 | TRUE | FALSE |
| LOC107066464 | diphthine methyl ester synthase | 0.02518 | 92.31084 | TRUE | FALSE |
| LOC107071532 | ADAMTS-like protein 4 | 0.02176 | 92.32713 | TRUE | FALSE |
| LOC107069186 | CTP synthase | 0.02069 | 92.34526 | TRUE | FALSE |
| LOC107072366 | uncharacterized protein LOC107072366 | 0.02291 | 92.35045 | TRUE | FALSE |
| LOC107063626 | piggyBac transposable element-derived pro | 0.04456 | 92.38648 | TRUE | FALSE |
| LOC107066673 | uncharacterized protein LOC107066673 | 0.02059 | 92.38948 | TRUE | FALSE |
| LOC107072245 | coiled-coil domain-containing protein 22 ho | 0.02119 | 92.41387 | TRUE | FALSE |
| LOC107067178 | AN1-type zinc finger protein 2A-like | 0.02035 | 92.45608 | TRUE | FALSE |
| LOC107067904 | otopetrin-3-like | 0.02035 | 92.53267 | TRUE | FALSE |

| | | | | | |
|---|---|---|---|---|---|
| LOC107063974 | protoheme IX farnesyltransferase, mitochor | 0.02055 | 92.54333 | TRUE | FALSE |
| LOC107064038 | mucin-2-like | 0.02066 | 92.54876 | TRUE | FALSE |
| LOC107071435 | uncharacterized protein LOC107071435 | 0.02092 | 92.62079 | TRUE | FALSE |
| LOC107074481 | cell division control protein 6 homolog | 0.02253 | 92.63008 | TRUE | FALSE |
| LOC107065854 | uncharacterized protein LOC107065854 | 0.02117 | 92.68762 | TRUE | FALSE |
| LOC107068405 | ATPase ASNA1 homolog | 0.02017 | 92.71232 | TRUE | FALSE |
| LOC107070189 | basement membrane-specific heparan sulfa | 0.02156 | 92.77953 | TRUE | FALSE |
| LOC107064596 | apoptosis-resistant E3 ubiquitin protein liga | 0.02045 | 92.81155 | TRUE | FALSE |
| LOC107068365 | 40S ribosomal protein S25 | 0.02128 | 92.8357 | TRUE | FALSE |
| LOC107072398 | protein kish-A | 0.02055 | 92.85891 | FALSE | FALSE |
| LOC107069737 | phosphopantothenate--cysteine ligase | 0.02089 | 92.88698 | FALSE | FALSE |
| LOC107066521 | elongation factor Tu GTP-binding domain-cc | 0.02199 | 92.89508 | TRUE | FALSE |
| LOC107067599 | uncharacterized protein LOC107067599 | 0.02191 | 92.90776 | TRUE | FALSE |
| LOC107066051 | interferon-inducible double-stranded RNA-c | 0.10748 | 92.90968 | TRUE | FALSE |
| LOC107073131 | peptidyl-prolyl cis-trans isomerase NIMA-int | 0.02217 | 92.9105 | TRUE | FALSE |
| LOC107064674 | transcription elongation factor 1 homolog | 0.02151 | 93.04774 | TRUE | FALSE |
| LOC107072020 | protein jagunal | 0.02082 | 93.05382 | TRUE | FALSE |
| LOC107066226 | integrin-alpha FG-GAP repeat-containing pro | 0.02206 | 93.11716 | TRUE | FALSE |
| LOC107074558 | uncharacterized protein LOC107074558 | 0.02462 | 93.13103 | TRUE | FALSE |
| LOC107073938 | glycerol kinase | 0.02121 | 93.13768 | TRUE | FALSE |
| LOC107063683 | 2-phosphoxylose phosphatase 1 | 0.02125 | 93.16571 | TRUE | FALSE |
| LOC107070384 | antizyme inhibitor 2-like | 0.02117 | 93.17413 | TRUE | FALSE |
| LOC107071246 | transformation/transcription domain-associ | 0.02246 | 93.21496 | TRUE | FALSE |
| LOC107067786 | DNA primase large subunit-like | 0.02253 | 93.23148 | TRUE | FALSE |
| LOC107070934 | kinesin-like protein KIF16B | 0.02012 | 93.23485 | TRUE | FALSE |
| LOC107073605 | uncharacterized protein LOC107073605 | 0.02192 | 93.23944 | TRUE | FALSE |
| LOC107068290 | tubulin delta chain-like | 0.02413 | 93.25212 | TRUE | FALSE |
| LOC107072765 | iodotyrosine deiodinase 1 | 0.02449 | 93.27939 | TRUE | FALSE |
| LOC107063638 | cilia- and flagella-associated protein 36 | 0.02048 | 93.29984 | TRUE | FALSE |
| LOC107073461 | LOW QUALITY PROTEIN: craniofacial develo | 0.02224 | 93.34285 | TRUE | FALSE |
| LOC107070149 | mismatch repair endonuclease PMS2 | 0.02332 | 93.40106 | TRUE | FALSE |
| LOC107069497 | jmjC domain-containing protein 4 | 0.01966 | 93.41458 | TRUE | FALSE |
| LOC107069830 | uncharacterized protein LOC107069830 | 0.02109 | 93.41771 | TRUE | FALSE |
| LOC107068004 | uncharacterized protein LOC107068004 | 0.0239 | 93.43474 | TRUE | FALSE |
| LOC107072488 | uncharacterized protein LOC107072488 | 0.02067 | 93.43882 | TRUE | FALSE |
| LOC107070836 | MKL/myocardin-like protein 1 | 0.02808 | 93.46794 | TRUE | FALSE |
| LOC107072026 | BTB/POZ domain-containing protein 10 | 0.02256 | 93.48714 | TRUE | FALSE |
| LOC107071277 | ATP-dependent RNA helicase dbp2-like | 0.02042 | 93.53248 | TRUE | FALSE |
| LOC107069093 | integrator complex subunit 2 | 0.02448 | 93.53384 | TRUE | FALSE |
| LOC107067224 | microtubule-associated protein futsch-like | 0.02131 | 93.53848 | FALSE | FALSE |
| LOC107065257 | protein phosphatase 1 regulatory subunit 7 | 0.02187 | 93.55002 | TRUE | FALSE |
| LOC107069510 | vang-like protein 2 | 0.02141 | 93.57981 | TRUE | FALSE |
| Trnak-uuu | Trnak-uuu | 0.02067 | 93.58671 | TRUE | FALSE |
| LOC107067889 | uncharacterized protein LOC107067889 | 0.02276 | 93.6196 | TRUE | FALSE |
| LOC107065170 | NEDD4 family-interacting protein 1-like | 0.02114 | 93.62495 | TRUE | FALSE |
| LOC107073685 | nose resistant to fluoxetine protein 6-like | 0.02055 | 93.62716 | TRUE | FALSE |
| LOC107069236 | homeobox protein PKNOX2-like | 0.02481 | 93.63898 | TRUE | FALSE |
| LOC107065601 | defective pharyngeal development protein 4 | 0.0202 | 93.66771 | TRUE | FALSE |
| LOC107067395 | zinc transporter 9 | 0.02054 | 93.6809 | TRUE | FALSE |
| LOC107065931 | ankyrin repeat domain-containing protein 6 | 0.02063 | 93.69743 | TRUE | FALSE |

| | | | | | |
|---|---|---|---|---|---|
| LOC107066352 | nesprin-1 | 0.02099 | 93.69911 | TRUE | FALSE |
| LOC107074394 | uncharacterized protein LOC107074394 | 0.02122 | 93.70728 | TRUE | FALSE |
| LOC107070079 | uncharacterized protein LOC107070079 | 0.02063 | 93.73008 | TRUE | FALSE |
| LOC107064865 | uncharacterized protein LOC107064865 | 0.02085 | 93.75403 | TRUE | FALSE |
| LOC107065462 | adhesion G protein-coupled receptor A3 | 0.0225 | 93.76129 | TRUE | FALSE |
| LOC107073671 | girdin-like | 0.02223 | 93.78859 | TRUE | FALSE |
| LOC107067303 | TAR DNA-binding protein 43-like | 0.02047 | 93.79098 | TRUE | FALSE |
| LOC107073960 | insulin gene enhancer protein ISL-1 | 0.0211 | 93.81105 | TRUE | FALSE |
| LOC107067653 | short-chain specific acyl-CoA dehydrogenase | 0.02012 | 93.87633 | TRUE | FALSE |
| LOC107064791 | vacuolar fusion protein CCZ1 homolog | 0.02536 | 93.87846 | TRUE | FALSE |
| LOC107072162 | forkhead box protein P2 | 0.02641 | 93.90212 | TRUE | FALSE |
| LOC107070959 | methyltransferase-like protein 7B | 0.02155 | 93.92132 | TRUE | FALSE |
| LOC107070406 | putative fatty acyl-CoA reductase CG8306 | 0.02109 | 93.93027 | TRUE | FALSE |
| LOC107067849 | collagen alpha-5(IV) chain-like | 0.02155 | 93.94184 | TRUE | FALSE |
| LOC107072932 | ATP-dependent DNA helicase PIF1 | 0.02244 | 93.94317 | TRUE | FALSE |
| LOC107070326 | uncharacterized protein LOC107070326 | 0.02056 | 93.96116 | TRUE | FALSE |
| LOC107066115 | NCK-interacting protein with SH3 domain | 0.0207 | 93.9635 | TRUE | FALSE |
| LOC107067455 | ralBP1-associated Eps domain-containing pr | 0.02131 | 94.01195 | TRUE | FALSE |
| LOC107070992 | embryonic stem cell-specific 5-hydroxymeth | 0.02074 | 94.02489 | TRUE | FALSE |
| LOC107071334 | eukaryotic initiation factor 4A-I | 0.02184 | 94.06067 | TRUE | FALSE |
| LOC107074598 | uncharacterized protein LOC107074598 | 0.02139 | 94.06602 | TRUE | FALSE |
| LOC107071891 | vacuolar fusion protein MON1 homolog A | 0.02174 | 94.06748 | TRUE | FALSE |
| LOC107070456 | pupal cuticle protein G1A-like | 0.02265 | 94.07505 | TRUE | FALSE |
| LOC107064774 | amine sulfotransferase-like | 0.02081 | 94.07508 | TRUE | FALSE |
| LOC107072901 | uncharacterized protein LOC107072901 | 0.02039 | 94.07666 | TRUE | FALSE |
| LOC107074028 | putative leucine-rich repeat-containing prot | 0.02078 | 94.10596 | TRUE | FALSE |
| LOC107067633 | uncharacterized protein LOC107067633 | 0.02662 | 94.12296 | TRUE | FALSE |
| LOC107064575 | uncharacterized protein LOC107064575 | 0.02238 | 94.17186 | TRUE | FALSE |
| LOC107067267 | probable Bax inhibitor 1 | 0.02159 | 94.17786 | TRUE | FALSE |
| LOC107073175 | uncharacterized protein LOC107073175 | 0.02088 | 94.23274 | TRUE | FALSE |
| LOC107073336 | guanine nucleotide-binding protein subunit | 0.02045 | 94.26613 | TRUE | FALSE |
| LOC107074254 | centrosomal protein of 112 kDa-like | 0.02328 | 94.32157 | TRUE | FALSE |
| LOC107066358 | zinc finger protein OZF-like | 0.02054 | 94.34613 | TRUE | FALSE |
| LOC107069226 | AT-rich interactive domain-containing prote | 0.02257 | 94.35142 | TRUE | FALSE |
| LOC107066979 | trafficking protein particle complex subunit | 0.02158 | 94.35963 | TRUE | FALSE |
| LOC107072462 | uncharacterized protein LOC107072462 | 0.02026 | 94.38409 | TRUE | FALSE |
| LOC107069496 | odorant receptor 43a-like | 0.0251 | 94.38666 | TRUE | FALSE |
| LOC107074328 | 39S ribosomal protein L37, mitochondrial | 0.02057 | 94.41642 | TRUE | FALSE |
| LOC107069550 | protein YIPF5 | 0.02052 | 94.41792 | TRUE | FALSE |
| LOC107072377 | anaphase-promoting complex subunit cdh1- | 0.0243 | 94.43162 | TRUE | FALSE |
| LOC107070191 | sodium-independent sulfate anion transpor | 0.02136 | 94.45226 | TRUE | FALSE |
| LOC107067282 | homeobox protein 9-like | 0.02021 | 94.50899 | TRUE | FALSE |
| LOC107071381 | transport and Golgi organization protein 1 | 0.0222 | 94.55685 | TRUE | FALSE |
| LOC107069122 | transmembrane 9 superfamily member 2 | 0.0207 | 94.56075 | TRUE | FALSE |
| LOC107066916 | CD63 antigen-like | 0.02149 | 94.57408 | TRUE | FALSE |
| LOC107069068 | chromatin modification-related protein ME/ | 0.02182 | 94.5836 | TRUE | FALSE |
| LOC107065745 | PP2C-like domain-containing protein CG980 | 0.02439 | 94.58678 | TRUE | FALSE |
| LOC107066016 | serine protease snake-like | 0.01997 | 94.59177 | TRUE | FALSE |
| LOC107067740 | isovaleryl-CoA dehydrogenase, mitochondri | 0.02342 | 94.63974 | TRUE | FALSE |
| LOC107069301 | prolactin regulatory element-binding protei | 0.02118 | 94.66787 | TRUE | FALSE |

| | | | | | |
|---|---|---|---|---|---|
| LOC107068982 | syntaxin-1B-like | 0.0235 | 94.66807 | TRUE | FALSE |
| LOC107073092 | synapse-associated protein 1 | 0.02296 | 94.68208 | TRUE | FALSE |
| LOC107068319 | homeobox protein 6-like | 0.02825 | 94.68763 | TRUE | FALSE |
| LOC107070815 | calreticulin | 0.02079 | 94.70008 | TRUE | FALSE |
| LOC107071189 | uncharacterized protein LOC107071189 | 0.02114 | 94.71144 | TRUE | FALSE |
| LOC107067411 | tRNA (guanine-N(7)-)-methyltransferase | 0.02264 | 94.71749 | TRUE | FALSE |
| LOC107064493 | dedicator of cytokinesis protein 9 | 0.02287 | 94.71758 | TRUE | FALSE |
| LOC107063624 | serine protease easter-like | 0.02065 | 94.72514 | TRUE | FALSE |
| LOC107074393 | 39S ribosomal protein L33, mitochondrial | 0.02161 | 94.72778 | TRUE | FALSE |
| LOC107067132 | elongation factor 1-alpha | 0.02397 | 94.74364 | TRUE | FALSE |
| LOC107071531 | methylosome subunit pICln | 0.02336 | 94.76204 | TRUE | FALSE |
| LOC107073979 | 40S ribosomal protein S6-like | 0.02251 | 94.76594 | TRUE | FALSE |
| LOC107067718 | beta-1,3-glucan-binding protein-like | 0.02038 | 94.83139 | TRUE | FALSE |
| LOC107070479 | uncharacterized protein LOC107070479 | 0.01993 | 94.85686 | TRUE | FALSE |
| LOC107070877 | uncharacterized protein LOC107070877 | 0.02669 | 94.85854 | TRUE | FALSE |
| LOC107064363 | uncharacterized protein LOC107064363 | 0.02111 | 94.88756 | TRUE | FALSE |
| LOC107066728 | cyclin-H | 0.0199 | 94.89076 | TRUE | FALSE |
| LOC107065508 | C-terminal-binding protein | 0.02087 | 94.92107 | TRUE | FALSE |
| LOC107064333 | cGMP-dependent protein kinase, isozyme 2 | 0.02287 | 94.92521 | TRUE | FALSE |
| LOC107067221 | uncharacterized protein LOC107067221 | 0.0218 | 94.96034 | TRUE | FALSE |
| LOC107064024 | kinesin-like protein KIF18A | 0.02958 | 94.99526 | TRUE | FALSE |
| LOC107064658 | sn1-specific diacylglycerol lipase alpha | 0.0222 | 95.01105 | TRUE | FALSE |
| LOC107063884 | protein argonaute-2 | 0.02228 | 95.02847 | TRUE | FALSE |
| LOC107064816 | uncharacterized protein LOC107064816 | 0.02091 | 95.04075 | TRUE | FALSE |
| LOC107068381 | protein-lysine N-methyltransferase n6amt2 | 0.02396 | 95.07606 | TRUE | FALSE |
| LOC107068689 | ATP synthase subunit gamma, mitochondria | 0.02141 | 95.08051 | TRUE | FALSE |
| LOC107067682 | uncharacterized protein LOC107067682 | 0.02091 | 95.08141 | TRUE | FALSE |
| LOC107073393 | fidgetin-like protein 1 | 0.02037 | 95.1004 | TRUE | FALSE |
| LOC107065647 | tRNA (adenine(58)-N(1))-methyltransferase | 0.02222 | 95.13285 | TRUE | FALSE |
| LOC107066624 | heat shock 70 kDa protein cognate 5 | 0.02507 | 95.15676 | TRUE | FALSE |
| LOC107068663 | ATP-dependent RNA helicase DDX54 | 0.02156 | 95.16684 | TRUE | FALSE |
| LOC107064242 | uncharacterized protein LOC107064242 | 0.02115 | 95.17079 | TRUE | FALSE |
| LOC107069839 | uncharacterized protein LOC107069839 | 0.02116 | 95.17975 | TRUE | FALSE |
| LOC107071745 | ELMO domain-containing protein F | 0.02337 | 95.18145 | TRUE | FALSE |
| LOC107066161 | transducin beta-like protein 3 | 0.02278 | 95.1999 | TRUE | FALSE |
| LOC107068389 | T-box-containing protein TBXT | 0.02138 | 95.20808 | TRUE | FALSE |
| LOC107067846 | retinal homeobox protein Rx2-like | 0.02639 | 95.21811 | TRUE | FALSE |
| LOC107073398 | uncharacterized protein LOC107073398 | 0.02183 | 95.23894 | TRUE | FALSE |
| LOC107068695 | CDK-activating kinase assembly factor MAT1 | 0.02278 | 95.26646 | TRUE | FALSE |
| LOC107072935 | palmitoyltransferase ZDHHC6 | 0.02094 | 95.27599 | TRUE | FALSE |
| LOC107066915 | ubiquitin-conjugating enzyme E2-24 kDa | 0.02146 | 95.3368 | TRUE | FALSE |
| LOC107064814 | uncharacterized protein LOC107064814 | 0.02812 | 95.3584 | TRUE | FALSE |
| LOC107066447 | diphthine--ammonia ligase | 0.02287 | 95.40022 | TRUE | FALSE |
| LOC107070570 | uncharacterized protein LOC107070570 | 0.02328 | 95.41201 | TRUE | FALSE |
| LOC107071642 | cytoplasmic protein NCK1 | 0.02091 | 95.41208 | TRUE | FALSE |
| LOC107069954 | coiled-coil domain-containing protein 25 | 0.02157 | 95.48718 | TRUE | FALSE |
| LOC107070400 | uncharacterized protein LOC107070400 | 0.02217 | 95.50453 | TRUE | FALSE |
| LOC107071758 | putative sodium-coupled neutral amino acic | 0.02057 | 95.54353 | TRUE | FALSE |
| LOC107063637 | tetratricopeptide repeat protein 1 | 0.02293 | 95.5637 | TRUE | FALSE |
| LOC107067915 | odorant receptor 4-like | 0.02228 | 95.61268 | TRUE | FALSE |

| | | | | | |
|---|---|---|---|---|---|
| LOC107066485 | uncharacterized protein LOC107066485 | 0.02297 | 95.62681 | TRUE | FALSE |
| LOC107066823 | coronin-6 | 0.02088 | 95.63283 | TRUE | FALSE |
| LOC107065792 | transcription initiation factor TFIID subunit 1 | 0.02355 | 95.66822 | TRUE | FALSE |
| LOC107072927 | geminin-like | 0.02002 | 95.67188 | TRUE | FALSE |
| LOC107068043 | uncharacterized protein LOC107068043 | 0.02321 | 95.69092 | TRUE | FALSE |
| LOC107072735 | enolase-phosphatase E1-like | 0.02235 | 95.69584 | TRUE | FALSE |
| LOC107069541 | zinc finger protein 384-like | 0.02049 | 95.78129 | TRUE | FALSE |
| LOC107065994 | S-methyl-5'-thioadenosine phosphorylase-li | 0.02333 | 95.79098 | TRUE | FALSE |
| LOC107066671 | F-box/LRR-repeat protein 7-like | 0.02285 | 95.81574 | TRUE | FALSE |
| LOC107066710 | uncharacterized protein C18orf19 homolog | 0.02014 | 95.88965 | TRUE | FALSE |
| LOC107069024 | nuclear hormone receptor FTZ-F1 beta | 0.02331 | 95.89007 | TRUE | FALSE |
| LOC107072780 | TPPP family protein CG45057-like | 0.021 | 95.90296 | TRUE | FALSE |
| LOC107065838 | actin-binding protein anillin-like | 0.03584 | 95.90883 | TRUE | FALSE |
| LOC107072399 | protein pygopus | 0.02132 | 95.93209 | TRUE | FALSE |
| LOC107069699 | nuclear pore complex protein Nup214 | 0.02186 | 95.93468 | TRUE | FALSE |
| LOC107073854 | uncharacterized protein LOC107073854 | 0.02294 | 95.93736 | TRUE | FALSE |
| LOC107063759 | copper-transporting ATPase 1 | 0.02142 | 95.96704 | TRUE | FALSE |
| LOC107064267 | pre-mRNA-splicing factor Syf2 | 0.02098 | 95.99616 | TRUE | FALSE |
| LOC107073533 | uncharacterized protein LOC107073533 | 0.02123 | 95.99883 | TRUE | FALSE |
| LOC107074677 | U4/U6.U5 tri-snRNP-associated protein 2 | 0.02188 | 96.01352 | TRUE | FALSE |
| LOC107065749 | GATA-binding factor C-like | 0.02517 | 96.02377 | TRUE | FALSE |
| LOC107063984 | uncharacterized protein LOC107063984 | 0.02273 | 96.03226 | TRUE | FALSE |
| LOC107064501 | oxysterol-binding protein 1 | 0.02359 | 96.03293 | TRUE | FALSE |
| LOC107065020 | zinc finger protein 678 | 0.02057 | 96.09897 | TRUE | FALSE |
| LOC107064352 | uncharacterized protein LOC107064352 | 0.02244 | 96.10107 | TRUE | FALSE |
| LOC107071918 | catenin delta-2 | 0.03013 | 96.18495 | TRUE | FALSE |
| LOC107065083 | LOW QUALITY PROTEIN: alkaline phosphata | 0.02112 | 96.19215 | TRUE | FALSE |
| LOC107066634 | yrdC domain-containing protein, mitochond | 0.02293 | 96.21156 | TRUE | FALSE |
| LOC107073150 | 26S protease regulatory subunit 8 | 0.0217 | 96.2206 | TRUE | FALSE |
| LOC107073164 | intraflagellar transport protein 80 homolog | 0.02285 | 96.25052 | TRUE | FALSE |
| LOC107072097 | bromodomain-containing protein 2-like | 0.02316 | 96.29059 | TRUE | FALSE |
| LOC107069530 | sodium- and chloride-dependent GABA tran | 0.02106 | 96.2922 | TRUE | FALSE |
| LOC107073488 | hydroxylysine kinase | 0.02078 | 96.29368 | TRUE | FALSE |
| LOC107066237 | ras-like GTP-binding protein Rho1 | 0.02291 | 96.31131 | TRUE | FALSE |
| LOC107073747 | uncharacterized protein LOC107073747 | 0.02074 | 96.3245 | TRUE | FALSE |
| LOC107067780 | uncharacterized protein LOC107067780 | 0.02086 | 96.3831 | TRUE | FALSE |
| LOC107067415 | acetylcholine receptor subunit alpha-like | 0.02408 | 96.40282 | TRUE | FALSE |
| LOC107068726 | 26S proteasome non-ATPase regulatory sub | 0.02349 | 96.40824 | TRUE | FALSE |
| LOC107073046 | uncharacterized protein LOC107073046 | 0.02813 | 96.43567 | TRUE | FALSE |
| LOC107066274 | uncharacterized protein LOC107066274 | 0.02522 | 96.4485 | TRUE | FALSE |
| LOC107068070 | facilitated trehalose transporter Tret1-like | 0.02216 | 96.4492 | TRUE | FALSE |
| LOC107063896 | probable ATP-dependent RNA helicase DHX | 0.02333 | 96.45731 | TRUE | FALSE |
| LOC107071380 | syntaxin-5 | 0.02095 | 96.45932 | TRUE | FALSE |
| LOC107072961 | uncharacterized protein LOC107072961 | 0.0218 | 96.46995 | TRUE | FALSE |
| LOC107068623 | uncharacterized protein LOC107068623 | 0.02135 | 96.47394 | TRUE | FALSE |
| LOC107073786 | uncharacterized protein LOC107073786 | 0.02353 | 96.47451 | TRUE | FALSE |
| LOC107072577 | uncharacterized protein LOC107072577 | 0.04007 | 96.56285 | TRUE | FALSE |
| LOC107069769 | 39S ribosomal protein L55, mitochondrial | 0.02168 | 96.57457 | TRUE | FALSE |
| LOC107065694 | protein lifeguard 1-like | 0.02058 | 96.57611 | TRUE | FALSE |
| LOC107069793 | nuclear transcription factor Y subunit gamm | 0.02122 | 96.58781 | TRUE | FALSE |

| | | | | | |
|---|---|---|---|---|---|
| LOC107068452 | glucose dehydrogenase [FAD, quinone | 0.02172 | 96.62196 | TRUE | FALSE |
| LOC107067738 | fatty acid synthase | 0.03702 | 96.62198 | TRUE | FALSE |
| LOC107066127 | uncharacterized protein LOC107066127 | 0.02093 | 96.7137 | TRUE | FALSE |
| LOC107071217 | aquaporin AQPcic-like | 0.02173 | 96.71928 | TRUE | FALSE |
| LOC107074111 | phosphatidylinositol 3-kinase regulatory sub | 0.03833 | 96.73094 | TRUE | FALSE |
| LOC107066712 | protein slowmo | 0.02239 | 96.75575 | TRUE | FALSE |
| LOC107068580 | pyridoxine-5'-phosphate oxidase-like | 0.0214 | 96.7783 | TRUE | FALSE |
| LOC107066562 | uncharacterized protein LOC107066562 | 0.02085 | 96.77927 | TRUE | FALSE |
| LOC107067341 | unconventional myosin-XV | 0.02071 | 96.83021 | TRUE | FALSE |
| LOC107068546 | uncharacterized protein LOC107068546 | 0.0237 | 96.88219 | TRUE | FALSE |
| LOC107072196 | poly(ADP-ribose) glycohydrolase ARH3-like | 0.0237 | 96.89918 | TRUE | FALSE |
| LOC107067895 | uncharacterized protein LOC107067895 | 0.02091 | 96.9059 | TRUE | FALSE |
| LOC107066190 | NADH dehydrogenase [ubiquinone | 0.0203 | 96.91606 | TRUE | FALSE |
| LOC107074488 | putative E3 ubiquitin-protein ligase UBR7 | 0.02275 | 96.92632 | TRUE | FALSE |
| LOC107070011 | uncharacterized protein LOC107070011 | 0.02471 | 96.94987 | TRUE | FALSE |
| LOC107069415 | uncharacterized protein LOC107069415 | 0.1417 | 96.96238 | TRUE | FALSE |
| LOC107070549 | persulfide dioxygenase ETHE1, mitochondri | 0.02123 | 96.9725 | TRUE | FALSE |
| LOC107070165 | gastrin/cholecystokinin type B receptor-like | 0.02135 | 97.03883 | TRUE | FALSE |
| LOC107070637 | 60S ribosomal protein L10 | 0.02017 | 97.06653 | TRUE | FALSE |
| LOC107072312 | cadherin-86C | 0.0269 | 97.0818 | TRUE | FALSE |
| LOC107070816 | exosome complex component CSL4 | 0.0217 | 97.08329 | TRUE | FALSE |
| LOC107065661 | uncharacterized protein LOC107065661 | 0.02213 | 97.10805 | TRUE | FALSE |
| LOC107070223 | exocyst complex component 8 | 0.04059 | 97.10908 | TRUE | FALSE |
| LOC107068075 | probable glutamine-dependent NAD(+) synt | 0.02457 | 97.11758 | TRUE | FALSE |
| LOC107069958 | odorant receptor 13a-like | 0.02218 | 97.12188 | TRUE | FALSE |
| LOC107072913 | uncharacterized protein LOC107072913 | 0.02244 | 97.14973 | TRUE | FALSE |
| LOC107070525 | caprin homolog | 0.02117 | 97.15568 | TRUE | FALSE |
| LOC107073990 | probable 28S rRNA (cytosine-C(5))-methyltr | 0.02107 | 97.18227 | TRUE | FALSE |
| LOC107064132 | 28S ribosomal protein S5, mitochondrial | 0.02123 | 97.21864 | TRUE | FALSE |
| LOC107072912 | uncharacterized protein LOC107072912 | 0.02104 | 97.2312 | TRUE | FALSE |
| LOC107069950 | ubiquitin-associated domain-containing pro | 0.02324 | 97.31224 | TRUE | FALSE |
| LOC107074049 | uncharacterized protein LOC107074049 | 0.02088 | 97.33865 | TRUE | FALSE |
| LOC107065998 | AMP deaminase 2-like | 0.0204 | 97.34064 | TRUE | FALSE |
| LOC107068064 | synaptic vesicle glycoprotein 2C-like | 0.03109 | 97.35164 | TRUE | FALSE |
| LOC107074419 | uncharacterized protein LOC107074419 | 0.02392 | 97.36568 | TRUE | FALSE |
| LOC107071570 | protein transport protein Sec23A | 0.02053 | 97.39413 | TRUE | FALSE |
| LOC107069727 | mitochondrial ribonuclease P protein 3 | 0.0218 | 97.40719 | TRUE | FALSE |
| LOC107073049 | glucose-induced degradation protein 8 hom | 0.02471 | 97.41529 | TRUE | FALSE |
| LOC107066414 | protein cornichon | 0.02164 | 97.44379 | TRUE | FALSE |
| LOC107072088 | DNA excision repair protein ERCC-1 | 0.02253 | 97.44386 | TRUE | FALSE |
| LOC107071166 | sorting and assembly machinery componen | 0.02103 | 97.4713 | TRUE | FALSE |
| LOC107064519 | uncharacterized protein LOC107064519 | 0.04037 | 97.49595 | TRUE | TRUE |
| LOC107074422 | serine/threonine-protein kinase PAK mbt | 0.02896 | 97.52265 | TRUE | FALSE |
| LOC107072798 | protein inturned | 0.02111 | 97.54401 | TRUE | FALSE |
| LOC107072598 | protein catecholamines up | 0.02025 | 97.58552 | TRUE | FALSE |
| LOC107069909 | odorant receptor 13a-like | 0.0219 | 97.60193 | TRUE | FALSE |
| LOC107070137 | dnaJ homolog subfamily C member 10-like | 0.02413 | 97.61856 | TRUE | FALSE |
| LOC107067552 | beta-1,3-galactosyltransferase 1-like | 0.02267 | 97.65741 | TRUE | FALSE |
| LOC107073809 | spermatogenesis-associated protein 20 | 0.02487 | 97.66194 | TRUE | FALSE |
| LOC107067183 | uncharacterized protein LOC107067183 | 0.02336 | 97.70535 | TRUE | FALSE |

| | | | | | |
|---|---|---|---|---|---|
| LOC107072782 | uncharacterized protein LOC107072782 | 0.02349 | 97.70912 | TRUE | FALSE |
| LOC107073724 | uncharacterized protein LOC107073724 | 0.02121 | 97.73304 | TRUE | FALSE |
| LOC107066989 | disks large homolog 4-like | 0.02394 | 97.73881 | TRUE | FALSE |
| LOC107069649 | leucine-rich repeat and immunoglobulin-like | 0.02873 | 97.7636 | TRUE | FALSE |
| LOC107072475 | uncharacterized protein LOC107072475 | 0.02483 | 97.8099 | TRUE | FALSE |
| LOC107072666 | synaptic vesicle glycoprotein 2B-like | 0.02374 | 97.8958 | TRUE | FALSE |
| LOC107065474 | mediator of RNA polymerase II transcription | 0.02385 | 97.96188 | TRUE | FALSE |
| LOC107073172 | breast cancer anti-estrogen resistance prote | 0.02458 | 98.0017 | TRUE | FALSE |
| LOC107064540 | uncharacterized protein LOC107064540 | 0.0262 | 98.00788 | TRUE | FALSE |
| LOC107067756 | glycine-rich cell wall structural protein-like | 0.02185 | 98.05121 | TRUE | FALSE |
| LOC107070958 | tyramine beta-hydroxylase | 0.02358 | 98.05453 | TRUE | FALSE |
| LOC107069936 | uncharacterized protein LOC107069936 | 0.02362 | 98.08085 | TRUE | FALSE |
| LOC107074451 | uncharacterized protein LOC107074451 | 0.02776 | 98.08971 | TRUE | FALSE |
| LOC107070453 | uncharacterized protein LOC107070453 | 0.02063 | 98.09308 | TRUE | FALSE |
| LOC107069722 | arrestin homolog | 0.02529 | 98.11044 | TRUE | FALSE |
| LOC107070657 | focadhesin | 0.04274 | 98.17529 | TRUE | FALSE |
| LOC107073710 | vascular endothelial growth factor receptor | 0.02607 | 98.20411 | TRUE | FALSE |
| LOC107066088 | BAG domain-containing protein Samui | 0.02793 | 98.20877 | TRUE | FALSE |
| LOC107073166 | zinc finger FYVE domain-containing protein | 0.02307 | 98.2492 | TRUE | FALSE |
| LOC107073053 | uncharacterized protein LOC107073053 | 0.02352 | 98.27778 | TRUE | FALSE |
| LOC107064052 | uncharacterized protein LOC107064052 | 0.0205 | 98.29973 | FALSE | FALSE |
| LOC107065725 | THO complex subunit 6 homolog | 0.02142 | 98.34344 | TRUE | FALSE |
| LOC107067683 | cyclic AMP-responsive element-binding prot | 0.0204 | 98.39953 | TRUE | FALSE |
| LOC107065171 | E3 ubiquitin-protein ligase TRIM33-like | 0.02874 | 98.41497 | TRUE | FALSE |
| LOC107065230 | tolloid-like protein 1 | 0.03441 | 98.41827 | TRUE | FALSE |
| LOC107067989 | inositol polyphosphate 5-phosphatase K-like | 0.02128 | 98.44343 | TRUE | FALSE |
| LOC107071234 | protein LSM12 homolog | 0.02254 | 98.49369 | TRUE | FALSE |
| LOC107067388 | cytochrome b5-like | 0.0225 | 98.50944 | TRUE | FALSE |
| LOC107072531 | fatty acid synthase | 0.02515 | 98.52295 | TRUE | FALSE |
| LOC107072449 | uncharacterized protein LOC107072449 | 0.03058 | 98.5312 | TRUE | FALSE |
| LOC107068698 | mitochondrial fission 1 protein-like | 0.02062 | 98.53934 | TRUE | FALSE |
| LOC107064615 | NHP2-like protein 1 homolog | 0.02063 | 98.55243 | TRUE | FALSE |
| LOC107072614 | ubiquitin domain-containing protein 1 | 0.02851 | 98.55892 | TRUE | FALSE |
| LOC107063892 | RNA polymerase-associated protein CTR9 ho | 0.021 | 98.58116 | TRUE | FALSE |
| LOC107068390 | E3 ubiquitin-protein ligase parkin | 0.02141 | 98.60811 | TRUE | FALSE |
| LOC107071999 | phospholipase D2 | 0.02417 | 98.6194 | TRUE | FALSE |
| LOC107072944 | uncharacterized protein LOC107072944 | 0.02397 | 98.63009 | TRUE | FALSE |
| LOC107072663 | actin-interacting protein 1 | 0.02574 | 98.63384 | TRUE | FALSE |
| LOC107071901 | protein AATF-like | 0.02462 | 98.63799 | TRUE | FALSE |
| LOC107065279 | venom carboxylesterase-6-like | 0.03254 | 98.65136 | TRUE | FALSE |
| LOC107065357 | translationally-controlled tumor protein hor | 0.02195 | 98.69702 | TRUE | FALSE |
| LOC107070767 | filamin-A | 0.0246 | 98.7051 | TRUE | FALSE |
| LOC107068500 | ejaculatory bulb-specific protein 3-like | 0.0253 | 98.70894 | TRUE | FALSE |
| LOC107068406 | inositol-trisphosphate 3-kinase A | 0.09005 | 98.71374 | TRUE | FALSE |
| LOC107070735 | synaptic vesicle glycoprotein 2B-like | 0.02383 | 98.71528 | TRUE | FALSE |
| LOC107065917 | monocyte to macrophage differentiation fa | 0.02261 | 98.75715 | TRUE | FALSE |
| LOC107065137 | uncharacterized protein LOC107065137 | 0.02207 | 98.75726 | TRUE | FALSE |
| LOC107063714 | uncharacterized protein LOC107063714 | 0.02086 | 98.77436 | TRUE | FALSE |
| LOC107067813 | tetratricopeptide repeat protein 27 | 0.02641 | 98.77546 | TRUE | FALSE |
| LOC107072240 | probable chitinase 3 | 0.02394 | 98.77632 | TRUE | FALSE |

| | | | | | |
|---|---|---|---|---|---|
| LOC107068343 | microsomal glutathione S-transferase 1-like | 0.022 | 98.81509 | TRUE | FALSE |
| LOC107068912 | uncharacterized protein LOC107068912 | 0.02239 | 98.83249 | TRUE | FALSE |
| LOC107074613 | NAD kinase-like | 0.02246 | 98.85113 | TRUE | FALSE |
| LOC107074074 | 40S ribosomal protein S2 | 0.03093 | 98.87129 | TRUE | FALSE |
| LOC107064595 | ubiquitin-40S ribosomal protein S27a | 0.02157 | 98.87335 | TRUE | FALSE |
| LOC107068163 | protein CDV3 homolog | 0.03168 | 98.87484 | TRUE | FALSE |
| LOC107068922 | uncharacterized protein LOC107068922 | 0.02393 | 98.90009 | TRUE | FALSE |
| LOC107066827 | thymidylate kinase | 0.02389 | 98.92239 | TRUE | FALSE |
| LOC107064820 | uncharacterized protein LOC107064820 | 0.04458 | 98.99357 | TRUE | FALSE |
| LOC107074427 | uncharacterized protein LOC107074427 | 0.02658 | 99.10002 | TRUE | FALSE |
| LOC107071862 | uncharacterized protein LOC107071862 | 0.02359 | 99.1613 | TRUE | FALSE |
| LOC107072673 | cadherin-89D | 0.02253 | 99.17063 | TRUE | FALSE |
| LOC107067072 | uncharacterized protein LOC107067072 | 0.02237 | 99.17649 | TRUE | FALSE |
| LOC107071462 | uncharacterized protein LOC107071462 | 0.02237 | 99.18558 | TRUE | FALSE |
| LOC107072491 | EF-hand calcium-binding domain-containing | 0.02373 | 99.19376 | TRUE | FALSE |
| LOC107073036 | nucleoporin NUP53 | 0.02214 | 99.23658 | TRUE | FALSE |
| LOC107070005 | KAT8 regulatory NSL complex subunit 2-like | 0.02484 | 99.2588 | TRUE | FALSE |
| LOC107071127 | 26S protease regulatory subunit 10B | 0.02988 | 99.26694 | TRUE | FALSE |
| LOC107073246 | uncharacterized protein LOC107073246 | 0.02375 | 99.2764 | TRUE | FALSE |
| LOC107064745 | tryptophan 5-hydroxylase 1 | 0.02163 | 99.29015 | TRUE | FALSE |
| LOC107068706 | protein brambleberry-like | 0.0213 | 99.29822 | TRUE | FALSE |
| LOC107064711 | puff-specific protein Bx42 | 0.02076 | 99.31761 | TRUE | FALSE |
| LOC107071005 | ubiquitin-conjugating enzyme E2 J1-like | 0.0214 | 99.32699 | TRUE | FALSE |
| LOC107064487 | uncharacterized protein LOC107064487 | 0.03172 | 99.3336 | TRUE | FALSE |
| LOC107070642 | uncharacterized protein LOC107070642 | 0.02772 | 99.34228 | TRUE | FALSE |
| LOC107073977 | uncharacterized protein LOC107073977 | 0.02455 | 99.36685 | TRUE | FALSE |
| LOC107073008 | dynein assembly factor 5, axonemal | 0.02243 | 99.37022 | TRUE | FALSE |
| LOC107065685 | peroxidase | 0.02265 | 99.39022 | TRUE | FALSE |
| LOC107068773 | POU domain protein CF1A | 0.02844 | 99.44426 | TRUE | FALSE |
| LOC107072855 | uncharacterized protein LOC107072855 | 0.021 | 99.5062 | TRUE | FALSE |
| LOC107064590 | cdc42 homolog | 0.02483 | 99.52905 | TRUE | FALSE |
| LOC107066946 | uncharacterized protein LOC107066946 | 0.02127 | 99.54424 | TRUE | FALSE |
| LOC107066612 | transcription cofactor vestigial-like protein 4 | 0.02338 | 99.55104 | TRUE | FALSE |
| LOC107068805 | NF-kappa-B inhibitor cactus | 0.0262 | 99.55184 | TRUE | FALSE |
| LOC107074353 | enoyl-CoA delta isomerase 2, mitochondrial | 0.02741 | 99.64489 | TRUE | FALSE |
| LOC107065395 | uncharacterized protein LOC107065395 | 0.03384 | 99.66063 | TRUE | FALSE |
| LOC107066955 | chromosome transmission fidelity protein 1 | 0.0284 | 99.67204 | TRUE | FALSE |
| LOC107067841 | E3 ubiquitin-protein ligase MIB2 | 0.02228 | 99.70903 | TRUE | FALSE |
| LOC107069834 | epoxide hydrolase 4-like | 0.02129 | 99.77371 | TRUE | FALSE |
| LOC107068893 | uncharacterized protein LOC107068893 | 0.03017 | 99.81545 | TRUE | FALSE |
| LOC107063731 | huntingtin | 0.03609 | 99.84246 | TRUE | FALSE |
| LOC107070881 | tektin-3-like | 0.02165 | 99.85029 | TRUE | FALSE |
| LOC107068682 | rho GTPase-activating protein 100F | 0.02426 | 99.89383 | TRUE | FALSE |
| LOC107071198 | mitochondrial amidoxime-reducing compon | 0.07878 | 99.93349 | TRUE | FALSE |
| LOC107070282 | histone deacetylase complex subunit SAP13 | 0.02247 | 99.93967 | TRUE | FALSE |
| LOC107065844 | metabotropic glutamate receptor-like | 0.02638 | 99.96525 | TRUE | FALSE |
| LOC107068030 | cell wall protein DAN4 | 0.08454 | 99.9685 | TRUE | FALSE |
| LOC107067049 | RING finger protein 10 | 0.02341 | 99.98783 | TRUE | FALSE |
| LOC107072562 | uncharacterized protein LOC107072562 | 0.02187 | 99.9887 | TRUE | FALSE |
| LOC107069455 | general transcription factor 3C polypeptide | 0.02228 | 99.99539 | TRUE | FALSE |

| | | | | | |
|---|---|---|---|---|---|
| LOC107065822 | pseudouridine-5'-phosphatase | 0.02174 | 100.00525 | TRUE | FALSE |
| LOC107063744 | immunoglobulin domain-containing protein | 0.03235 | 100.0115 | TRUE | FALSE |
| LOC107074582 | uncharacterized protein LOC107074582 | 0.02285 | 100.05897 | TRUE | FALSE |
| LOC107073917 | vitellogenin-like | 0.02009 | 100.06462 | TRUE | FALSE |
| LOC107070807 | polynucleotide 5'-hydroxyl-kinase NOL9-like | 0.0285 | 100.06565 | TRUE | FALSE |
| LOC107065765 | condensin complex subunit 3-like | 0.03441 | 100.07444 | TRUE | FALSE |
| LOC107068411 | phospholipase A-2-activating protein | 0.02142 | 100.11731 | TRUE | FALSE |
| LOC107068792 | O-phosphoseryl-tRNA(Sec) selenium transfe | 0.02099 | 100.2554 | TRUE | FALSE |
| LOC107065314 | zygotic gap protein knirps-like | 0.02369 | 100.26396 | TRUE | FALSE |
| LOC107070568 | endoplasmic reticulum-Golgi intermediate c | 0.02627 | 100.27281 | TRUE | FALSE |
| LOC107067206 | uncharacterized protein LOC107067206 | 0.02156 | 100.29205 | TRUE | FALSE |
| LOC107069011 | sodium/potassium/calcium exchanger 3 | 0.02722 | 100.30931 | TRUE | FALSE |
| LOC107070804 | zinc finger protein 845-like | 0.02134 | 100.35219 | TRUE | FALSE |
| LOC107073247 | heat shock protein 70 A2-like | 0.02285 | 100.35644 | TRUE | FALSE |
| LOC107073704 | uncharacterized protein LOC107073704 | 0.02236 | 100.35755 | TRUE | FALSE |
| LOC107073274 | negative elongation factor B-like | 0.02392 | 100.37078 | TRUE | FALSE |
| LOC107063741 | procollagen-lysine,2-oxoglutarate 5-dioxyge | 0.02264 | 100.37487 | TRUE | FALSE |
| LOC107073588 | uncharacterized protein LOC107073588 | 0.03349 | 100.37786 | TRUE | FALSE |
| LOC107072620 | histone deacetylase 6 | 0.02537 | 100.39272 | TRUE | FALSE |
| LOC107073260 | uncharacterized protein LOC107073260 | 0.02271 | 100.39969 | TRUE | FALSE |
| LOC107070559 | uncharacterized protein LOC107070559 | 0.02184 | 100.41782 | TRUE | FALSE |
| LOC107068793 | uncharacterized protein LOC107068793 | 0.02484 | 100.50786 | TRUE | FALSE |
| LOC107070359 | uncharacterized protein LOC107070359 | 0.02396 | 100.58885 | TRUE | FALSE |
| LOC107068597 | inositol polyphosphate multikinase-like | 0.02966 | 100.62391 | TRUE | FALSE |
| LOC107067369 | uncharacterized protein LOC107067369 | 0.02209 | 100.64162 | TRUE | FALSE |
| LOC107065412 | xenotropic and polytropic retrovirus recept( | 0.02459 | 100.71209 | TRUE | FALSE |
| LOC107065049 | polypeptide N-acetylgalactosaminyltransfer | 0.02135 | 100.71802 | TRUE | FALSE |
| LOC107070721 | peroxidase-like | 0.02138 | 100.74788 | TRUE | FALSE |
| LOC107073797 | uncharacterized protein LOC107073797 | 0.02323 | 100.75683 | TRUE | FALSE |
| LOC107070966 | protein kinase C | 0.02439 | 100.76977 | TRUE | FALSE |
| LOC107071324 | MATH and LRR domain-containing protein P | 0.02666 | 100.77529 | TRUE | FALSE |
| LOC107067232 | N-acetyltransferase 10 | 0.02093 | 100.78156 | TRUE | FALSE |
| LOC107065027 | SH3 domain-binding glutamic acid-rich prot( | 0.02763 | 100.81646 | TRUE | FALSE |
| LOC107066165 | ATP synthase subunit d, mitochondrial-like | 0.02273 | 100.85716 | TRUE | FALSE |
| LOC107065356 | inactive dipeptidyl peptidase 10 | 0.02224 | 100.89152 | TRUE | FALSE |
| LOC107070545 | opioid-binding protein/cell adhesion molecu | 0.02481 | 100.92067 | TRUE | FALSE |
| LOC107071528 | uncharacterized threonine-rich GPI-anchore | 0.02166 | 101.00262 | TRUE | FALSE |
| LOC107069030 | sodium/potassium/calcium exchanger 3-like | 0.09977 | 101.03313 | TRUE | FALSE |
| LOC107067131 | calnexin-like | 0.0296 | 101.07258 | TRUE | FALSE |
| LOC107066153 | SUZ domain-containing protein 1 | 0.02361 | 101.07452 | TRUE | FALSE |
| LOC107070303 | AFG3-like protein 2 | 0.0226 | 101.08668 | TRUE | FALSE |
| LOC107069204 | abl interactor 2 | 0.02226 | 101.1022 | TRUE | FALSE |
| LOC107074067 | uncharacterized protein LOC107074067 | 0.0279 | 101.20705 | TRUE | FALSE |
| LOC107074253 | uncharacterized protein LOC107074253 | 0.03087 | 101.209 | TRUE | FALSE |
| LOC107073629 | seipin | 0.02334 | 101.28185 | TRUE | FALSE |
| LOC107074108 | beta-hexosaminidase subunit beta-like | 0.03521 | 101.29612 | TRUE | FALSE |
| LOC107073326 | homeobox protein extradenticle | 0.02473 | 101.30035 | TRUE | FALSE |
| LOC107066621 | uncharacterized protein LOC107066621 | 0.02152 | 101.3114 | TRUE | FALSE |
| LOC107066099 | vacuolar protein sorting-associated protein | 0.03271 | 101.31872 | TRUE | FALSE |
| LOC107071534 | DNA-binding protein D-ETS-4-like | 0.02582 | 101.32139 | TRUE | FALSE |

| | | | | | |
|---|---|---|---|---|---|
| LOC107072439 | neprilysin-like | 0.02821 | 101.35003 | TRUE | FALSE |
| LOC107069381 | uncharacterized protein LOC107069381 | 0.02478 | 101.35016 | TRUE | FALSE |
| LOC107074597 | actin-binding protein IPP | 0.02386 | 101.36128 | TRUE | FALSE |
| LOC107072292 | ectopic P granules protein 5 homolog | 0.03372 | 101.37121 | TRUE | FALSE |
| LOC107071274 | testis-expressed sequence 2 protein | 0.02522 | 101.37739 | TRUE | FALSE |
| LOC107066879 | mitochondrial coenzyme A transporter SLC2 | 0.02222 | 101.39381 | TRUE | FALSE |
| LOC107064123 | uncharacterized protein LOC107064123 | 0.03232 | 101.40266 | TRUE | FALSE |
| LOC107069772 | probable serine/threonine-protein kinase D | 0.02921 | 101.44244 | TRUE | FALSE |
| LOC107072587 | microphthalmia-associated transcription fac | 0.02042 | 101.47879 | TRUE | FALSE |
| LOC107072219 | 28S ribosomal protein S21, mitochondrial | 0.02515 | 101.48069 | TRUE | FALSE |
| LOC107065531 | sodium channel protein 60E | 0.0293 | 101.51647 | TRUE | FALSE |
| LOC107070967 | uncharacterized protein LOC107070967 | 0.02494 | 101.5254 | TRUE | FALSE |
| LOC107065935 | mucin-17 | 0.02149 | 101.57206 | TRUE | FALSE |
| LOC107068181 | uncharacterized protein LOC107068181 | 0.02952 | 101.5903 | TRUE | FALSE |
| LOC107064878 | V-type proton ATPase subunit G | 0.02624 | 101.59086 | TRUE | FALSE |
| LOC107066164 | uncharacterized protein LOC107066164 | 0.02445 | 101.6067 | TRUE | FALSE |
| LOC107071200 | WD repeat-containing protein 18 | 0.03212 | 101.66743 | TRUE | FALSE |
| LOC107068144 | protein CBFA2T2 | 0.02331 | 101.68162 | TRUE | FALSE |
| LOC107065286 | graves disease carrier protein | 0.03687 | 101.70447 | TRUE | FALSE |
| LOC107070136 | mucin-5AC-like | 0.02049 | 101.74022 | TRUE | FALSE |
| LOC107068879 | uncharacterized protein LOC107068879 | 0.02498 | 101.74194 | TRUE | FALSE |
| LOC107069103 | phosphoribosylformylglycinamidine synthas | 0.03487 | 101.8306 | TRUE | FALSE |
| LOC107073294 | uncharacterized protein LOC107073294 | 0.02294 | 101.84124 | TRUE | FALSE |
| LOC107074065 | uncharacterized protein LOC107074065 | 0.02299 | 101.85757 | TRUE | FALSE |
| LOC107068686 | spectrin beta chain, non-erythrocytic 5 | 0.02067 | 101.87202 | TRUE | FALSE |
| LOC107066439 | lysosome-associated membrane glycoprote | 0.02251 | 101.89846 | TRUE | FALSE |
| LOC107071509 | uncharacterized protein LOC107071509 | 0.13473 | 101.94104 | TRUE | FALSE |
| LOC107072722 | 5-demethoxyubiquinone hydroxylase, mitoc | 0.02254 | 102.01125 | TRUE | FALSE |
| LOC107072296 | 5-hydroxytryptamine receptor 2B-like | 0.02536 | 102.05217 | TRUE | FALSE |
| LOC107067508 | uncharacterized protein LOC107067508 | 0.03851 | 102.10693 | TRUE | FALSE |
| LOC107068176 | biogenesis of lysosome-related organelles c | 0.03316 | 102.10971 | TRUE | FALSE |
| LOC107068263 | uncharacterized protein LOC107068263 | 0.02483 | 102.16091 | TRUE | FALSE |
| LOC107067135 | uroporphyrinogen decarboxylase | 0.02366 | 102.16251 | TRUE | FALSE |
| LOC107074425 | transmembrane protein 35 | 0.11713 | 102.19718 | TRUE | FALSE |
| LOC107074310 | dynein light chain 1, axonemal-like | 0.06891 | 102.21183 | TRUE | FALSE |
| LOC107068498 | putative glycerol kinase 5 | 0.02119 | 102.22561 | TRUE | FALSE |
| LOC107067045 | zinc finger protein 497-like | 0.02516 | 102.28221 | TRUE | FALSE |
| LOC107070515 | uncharacterized protein LOC107070515 | 0.02634 | 102.28628 | TRUE | FALSE |
| LOC107069635 | uncharacterized protein LOC107069635 | 0.03105 | 102.28985 | TRUE | FALSE |
| LOC107074400 | GTP:AMP phosphotransferase AK3, mitocho | 0.02283 | 102.31657 | TRUE | FALSE |
| LOC107064743 | uncharacterized protein LOC107064743 | 0.02113 | 102.32479 | TRUE | FALSE |
| LOC107066003 | UPF0454 protein C12orf49 homolog | 0.02521 | 102.32643 | TRUE | FALSE |
| LOC107066680 | barH-like 1 homeobox protein | 0.02423 | 102.32901 | TRUE | FALSE |
| LOC107066878 | mitotic checkpoint protein BUB3 | 0.02182 | 102.34887 | TRUE | FALSE |
| LOC107064145 | retinoblastoma-like protein 1 | 0.0295 | 102.38844 | TRUE | FALSE |
| LOC107070252 | transmembrane protein 184B | 0.03094 | 102.4074 | TRUE | FALSE |
| LOC107069618 | putative thiamine transporter SLC35F3 | 0.0362 | 102.41777 | TRUE | FALSE |
| LOC107074382 | uncharacterized protein LOC107074382 | 0.04536 | 102.4899 | TRUE | FALSE |
| LOC107069616 | B-box type zinc finger protein ncl-1 | 0.02645 | 102.55615 | TRUE | FALSE |
| LOC107064389 | transmembrane protease serine 9-like | 0.02425 | 102.64656 | TRUE | FALSE |

| | | | | | |
|---|---|---|---|---|---|
| LOC107072263 | probable WRKY transcription factor protein | 0.0221 | 102.65034 | TRUE | FALSE |
| LOC107070336 | uncharacterized protein LOC107070336 | 0.02032 | 102.66191 | TRUE | FALSE |
| LOC107064534 | autophagy protein 12-like | 0.02448 | 102.67288 | TRUE | FALSE |
| LOC107066436 | ras-related protein Rab-14 | 0.02716 | 102.67906 | TRUE | FALSE |
| LOC107072843 | uncharacterized protein LOC107072843 | 0.02428 | 102.69001 | TRUE | FALSE |
| LOC107073384 | mitochondrial sodium/hydrogen exchanger | 0.02275 | 102.69455 | TRUE | FALSE |
| LOC107069577 | uncharacterized protein LOC107069577 | 0.03132 | 102.71536 | TRUE | FALSE |
| LOC107068677 | exosome complex component RRP41 | 0.02265 | 102.7301 | TRUE | FALSE |
| LOC107068185 | 40S ribosomal protein S11 | 0.02698 | 102.7773 | TRUE | FALSE |
| LOC107073731 | uncharacterized protein LOC107073731 | 0.02218 | 102.81836 | TRUE | FALSE |
| LOC107065654 | CDK5 regulatory subunit-associated protein | 0.02834 | 102.84012 | TRUE | FALSE |
| LOC107068217 | kinesin-related protein 4 | 0.02786 | 102.86285 | TRUE | FALSE |
| LOC107071088 | RING finger and SPRY domain-containing pr | 0.08668 | 102.88481 | TRUE | FALSE |
| LOC107063897 | LOW QUALITY PROTEIN: ceramide phospho | 0.02201 | 102.9671 | TRUE | FALSE |
| LOC107072437 | protein FAM50 homolog | 0.02224 | 103.02606 | TRUE | FALSE |
| LOC107071440 | cyclin-dependent kinase 7 | 0.0419 | 103.04551 | TRUE | FALSE |
| LOC107066389 | 39S ribosomal protein L17, mitochondrial | 0.02313 | 103.04852 | TRUE | FALSE |
| LOC107069100 | putative uncharacterized protein DDB_G02 | 0.03511 | 103.05331 | TRUE | FALSE |
| LOC107070785 | uncharacterized protein LOC107070785 | 0.02398 | 103.07458 | TRUE | FALSE |
| LOC107067480 | mediator of RNA polymerase II transcriptior | 0.02233 | 103.09213 | TRUE | FALSE |
| LOC107070324 | hexosaminidase D-like | 0.02423 | 103.12937 | TRUE | FALSE |
| LOC107070269 | acyl-CoA:lysophosphatidylglycerol acyltrans | 0.02224 | 103.16271 | TRUE | FALSE |
| LOC107070994 | NTF2-related export protein | 0.02047 | 103.17618 | TRUE | FALSE |
| LOC107074675 | protein zer-1 homolog | 0.0276 | 103.20893 | TRUE | FALSE |
| LOC107069370 | aromatic-L-amino-acid decarboxylase-like | 0.0208 | 103.20906 | TRUE | FALSE |
| LOC107069565 | ATP-dependent DNA helicase Q4 | 0.02764 | 103.22569 | TRUE | FALSE |
| LOC107064778 | neurogenic protein mastermind-like | 0.03995 | 103.23245 | TRUE | FALSE |
| LOC107068000 | aldose reductase-like | 0.03894 | 103.27376 | TRUE | FALSE |
| LOC107066331 | optic atrophy 3 protein homolog | 0.02507 | 103.36499 | TRUE | FALSE |
| LOC107074123 | Down syndrome cell adhesion molecule-like | 0.02509 | 103.38201 | TRUE | FALSE |
| LOC107067613 | pupal cuticle protein Edg-91-like | 0.05004 | 103.38392 | TRUE | FALSE |
| LOC107069583 | uncharacterized protein LOC107069583 | 0.02224 | 103.51373 | TRUE | FALSE |
| LOC107067422 | uncharacterized protein LOC107067422 | 0.02349 | 103.55915 | TRUE | FALSE |
| LOC107068234 | serine/threonine-protein phosphatase 2A 5 | 0.02905 | 103.56696 | TRUE | FALSE |
| LOC107072559 | nicotinamide riboside kinase 1 | 0.02125 | 103.5774 | TRUE | FALSE |
| LOC107063720 | proteasome-associated protein ECM29 hom | 0.02877 | 103.58859 | TRUE | FALSE |
| LOC107063821 | E3 ubiquitin-protein ligase Hakai | 0.02419 | 103.62755 | TRUE | FALSE |
| LOC107069161 | uncharacterized protein LOC107069161 | 0.02612 | 103.63108 | TRUE | FALSE |
| LOC107067037 | PEST proteolytic signal-containing nuclear p | 0.03445 | 103.69598 | TRUE | FALSE |
| LOC107066942 | transcription and mRNA export factor ENY2 | 0.02304 | 103.69864 | TRUE | FALSE |
| LOC107064685 | uncharacterized protein LOC107064685 | 0.03231 | 103.77444 | TRUE | FALSE |
| LOC107072784 | PI-PLC X domain-containing protein 1-like | 0.03087 | 103.80286 | TRUE | FALSE |
| LOC107070969 | uncharacterized protein LOC107070969 | 0.0235 | 103.81565 | TRUE | FALSE |
| LOC107070957 | putative phospholipase B-like lamina ancest | 0.05605 | 103.81833 | TRUE | FALSE |
| LOC107073914 | putative GTP-binding protein 6 | 0.05933 | 103.8306 | TRUE | FALSE |
| LOC107066906 | cell wall integrity and stress response comp | 0.03511 | 103.86814 | TRUE | FALSE |
| LOC107064397 | uncharacterized protein LOC107064397 | 0.06008 | 103.91808 | TRUE | FALSE |
| LOC107070018 | twitchin | 0.03575 | 103.92878 | TRUE | FALSE |
| LOC107074587 | uncharacterized protein LOC107074587 | 0.0215 | 103.93832 | TRUE | FALSE |
| LOC107072182 | beta-1,4-mannosyl-glycoprotein 4-beta-N-a | 0.02744 | 103.96515 | TRUE | FALSE |

| | | | | | |
|---|---|---|---|---|---|
| LOC107069625 | uncharacterized protein LOC107069625 | 0.02376 | 103.9758 | TRUE | FALSE |
| LOC107070187 | excitatory amino acid transporter 3-like | 0.02185 | 103.99595 | TRUE | FALSE |
| LOC107066316 | uncharacterized protein LOC107066316 | 0.02113 | 104.00709 | TRUE | FALSE |
| LOC107067578 | uncharacterized protein LOC107067578 | 0.02061 | 104.01143 | TRUE | FALSE |
| LOC107072830 | solute carrier organic anion transporter fam | 0.02568 | 104.02921 | TRUE | FALSE |
| LOC107072662 | modifier of mdg4-like | 0.02398 | 104.03654 | TRUE | FALSE |
| LOC107067834 | disheveled-associated activator of morphog | 0.02807 | 104.03702 | TRUE | FALSE |
| LOC107074363 | density-regulated protein homolog | 0.02277 | 104.09004 | TRUE | FALSE |
| LOC107068855 | growth factor receptor-bound protein 14-lik | 0.06497 | 104.12852 | TRUE | FALSE |
| LOC107073939 | protein timeless homolog | 0.05521 | 104.12993 | TRUE | FALSE |
| LOC107065704 | cob(I)yrinic acid a,c-diamide adenosyltransf | 0.02435 | 104.13451 | TRUE | FALSE |
| LOC107068935 | farnesol dehydrogenase-like | 0.024 | 104.13488 | TRUE | FALSE |
| LOC107065548 | repetitive proline-rich cell wall protein 2-like | 0.1597 | 104.15212 | TRUE | FALSE |
| LOC107070311 | uncharacterized protein LOC107070311 | 0.02171 | 104.16483 | TRUE | FALSE |
| LOC107065152 | zinc finger protein 511 | 0.02804 | 104.18037 | TRUE | FALSE |
| LOC107066202 | uncharacterized protein LOC107066202 | 0.02485 | 104.18119 | TRUE | FALSE |
| LOC107065953 | ribonuclease inhibitor | 0.04128 | 104.19386 | TRUE | FALSE |
| LOC107070138 | signal recognition particle 14 kDa protein | 0.02188 | 104.21063 | TRUE | FALSE |
| LOC107071311 | uncharacterized protein LOC107071311 | 0.02241 | 104.22823 | TRUE | FALSE |
| LOC107068012 | protein phosphatase 1 regulatory subunit p | 0.02535 | 104.24323 | TRUE | FALSE |
| LOC107072670 | uncharacterized protein LOC107072670 | 0.02846 | 104.25024 | TRUE | TRUE |
| LOC107064490 | uncharacterized protein LOC107064490 | 0.02747 | 104.25044 | TRUE | FALSE |
| LOC107067031 | sodium-dependent noradrenaline transport | 0.02273 | 104.3144 | TRUE | FALSE |
| LOC107068715 | uncharacterized protein LOC107068715 | 0.02507 | 104.32004 | TRUE | FALSE |
| LOC107072119 | putative uncharacterized protein DDB_G028 | 0.02096 | 104.32274 | TRUE | FALSE |
| LOC107071873 | uncharacterized protein LOC107071873 | 0.02887 | 104.32278 | TRUE | FALSE |
| LOC107069998 | calcium and integrin-binding protein 1-like | 0.02643 | 104.3535 | TRUE | FALSE |
| LOC107068352 | phosphatidylinositol 4-kinase alpha | 0.03472 | 104.3867 | TRUE | FALSE |
| LOC107072208 | AP-3 complex subunit sigma-2 | 0.02882 | 104.48488 | TRUE | FALSE |
| LOC107067364 | alkylglycerol monooxygenase-like | 0.02127 | 104.49194 | TRUE | FALSE |
| LOC107066058 | KH domain-containing protein akap-1 | 0.02451 | 104.50203 | TRUE | FALSE |
| LOC107072613 | MMS19 nucleotide excision repair protein h | 0.07981 | 104.51353 | TRUE | FALSE |
| LOC107072668 | putative peptidyl-tRNA hydrolase PTRHD1 | 0.06256 | 104.54255 | TRUE | FALSE |
| LOC107071190 | protein kinase C-binding protein NELL1-like | 0.02981 | 104.56098 | TRUE | FALSE |
| LOC107070813 | mediator of RNA polymerase II transcription | 0.02414 | 104.57715 | TRUE | FALSE |
| LOC107068466 | double-strand-break repair protein rad21 ho | 0.12067 | 104.58321 | TRUE | FALSE |
| LOC107073638 | cytochrome P450 4C1-like | 0.02777 | 104.64281 | TRUE | FALSE |
| LOC107064075 | uncharacterized protein LOC107064075 | 0.02485 | 104.64443 | TRUE | FALSE |
| LOC107074652 | RAB6A-GEF complex partner protein 2 | 0.03419 | 104.65391 | TRUE | FALSE |
| LOC107064667 | uncharacterized protein LOC107064667 | 0.02959 | 104.65434 | TRUE | FALSE |
| LOC107065621 | popeye domain-containing protein 3-like | 0.0244 | 104.67167 | TRUE | FALSE |
| LOC107071559 | probable DNA mismatch repair protein Msh | 0.02556 | 104.67923 | TRUE | FALSE |
| LOC107067870 | protein NPC2 homolog | 0.05436 | 104.71765 | TRUE | FALSE |
| LOC107068921 | M-phase inducer phosphatase | 0.02485 | 104.74281 | TRUE | FALSE |
| LOC107064161 | NADH dehydrogenase [ubiquinone | 0.02893 | 104.74615 | TRUE | FALSE |
| LOC107069896 | protein MMS22-like | 0.02193 | 104.75179 | TRUE | FALSE |
| LOC107070166 | cytochrome P450 4g15-like | 0.02084 | 104.77017 | TRUE | FALSE |
| LOC107070631 | regulator of microtubule dynamics protein 1 | 0.03327 | 104.78881 | TRUE | FALSE |
| LOC107068069 | uncharacterized protein LOC107068069 | 0.03252 | 104.80593 | TRUE | FALSE |
| LOC107072203 | uncharacterized protein LOC107072203 | 0.02554 | 104.84783 | TRUE | FALSE |

| | | | | | |
|---|---|---|---|---|---|
| LOC107071616 | histone-lysine N-methyltransferase PRDM9 | 0.07411 | 104.87093 | TRUE | FALSE |
| LOC107073874 | 6-phosphofructo-2-kinase/fructose-2,6-bisp | 0.09424 | 104.88017 | TRUE | FALSE |
| LOC107065848 | insulin-like growth factor 2 mRNA-binding p | 0.02912 | 104.9209 | TRUE | FALSE |
| LOC107071607 | protein dopey-1 homolog | 0.02812 | 104.922 | TRUE | FALSE |
| LOC107067319 | 3'-5' ssDNA/RNA exonuclease TatD | 0.03057 | 104.94779 | TRUE | FALSE |
| LOC107063968 | neuroglobin-like | 0.04321 | 104.98085 | TRUE | FALSE |
| LOC107067207 | BTB/POZ domain-containing protein KCTD1 | 0.0437 | 105.00285 | TRUE | FALSE |
| LOC107073231 | probable serine/threonine-protein kinase fh | 0.03344 | 105.00353 | TRUE | FALSE |
| LOC107065764 | uncharacterized protein LOC107065764 | 0.02305 | 105.0536 | TRUE | FALSE |
| LOC107067154 | Bardet-Biedl syndrome 2 protein homolog | 0.02274 | 105.06141 | TRUE | FALSE |
| LOC107068673 | 4-hydroxybenzoate polyprenyltransferase, r | 0.02517 | 105.07766 | TRUE | FALSE |
| LOC107065381 | uncharacterized protein LOC107065381 | 0.0221 | 105.12968 | TRUE | FALSE |
| LOC107068152 | protein KTI12 homolog | 0.02838 | 105.16317 | TRUE | FALSE |
| LOC107067352 | ras-related protein Rab-1A | 0.03889 | 105.17099 | TRUE | FALSE |
| LOC107072486 | ubiquitin-conjugating enzyme E2 S | 0.02335 | 105.2003 | TRUE | FALSE |
| LOC107070054 | molybdenum cofactor biosynthesis protein | 0.02662 | 105.23329 | TRUE | FALSE |
| LOC107067261 | ankyrin-3-like | 0.02558 | 105.2381 | TRUE | FALSE |
| LOC107067105 | tektin-B1 | 0.03478 | 105.2594 | TRUE | FALSE |
| LOC107068611 | ribonucleoside-diphosphate reductase large | 0.02537 | 105.27173 | TRUE | FALSE |
| LOC107069446 | UPF0472 protein C16orf72 homolog | 0.02504 | 105.30306 | TRUE | FALSE |
| LOC107065736 | DNA topoisomerase 2-binding protein 1 | 0.02406 | 105.31342 | TRUE | FALSE |
| LOC107068485 | translation elongation factor 2 | 0.02478 | 105.3444 | TRUE | FALSE |
| LOC107070600 | protein PFF0380w | 0.02881 | 105.34468 | TRUE | FALSE |
| LOC107068566 | maternal B9.10 protein-like | 0.02621 | 105.43379 | TRUE | FALSE |
| LOC107068683 | ribosomal protein S6 kinase delta-1 | 0.02133 | 105.50335 | TRUE | FALSE |
| LOC107073214 | myotubularin-related protein 9 | 0.02303 | 105.51591 | TRUE | FALSE |
| LOC107065543 | uncharacterized protein LOC107065543 | 0.0271 | 105.52377 | TRUE | FALSE |
| LOC107072387 | putative uncharacterized protein DDB_G028 | 0.02886 | 105.59558 | TRUE | FALSE |
| LOC107064470 | kinesin-like protein CG14535 | 0.03444 | 105.59675 | TRUE | FALSE |
| LOC107072496 | phosphoglucomutase | 0.02725 | 105.65784 | TRUE | FALSE |
| LOC107064317 | malate dehydrogenase, cytoplasmic | 0.02532 | 105.68061 | TRUE | FALSE |
| LOC107071675 | anosmin-1 | 0.0268 | 105.69441 | TRUE | FALSE |
| LOC107066666 | PRKCA-binding protein | 0.02715 | 105.75351 | TRUE | FALSE |
| LOC107065978 | calcitonin gene-related peptide type 1 recep | 0.02658 | 105.76124 | TRUE | FALSE |
| LOC107070185 | tRNA-dihydrouridine(47) synthase [NAD(P)( | 0.02683 | 105.77471 | TRUE | FALSE |
| LOC107074018 | uncharacterized protein C1orf198 homolog | 0.02338 | 105.77538 | TRUE | FALSE |
| LOC107065766 | proto-oncogene tyrosine-protein kinase rec | 0.02625 | 105.77947 | TRUE | FALSE |
| LOC107068890 | adenylate kinase | 0.02898 | 105.79575 | TRUE | FALSE |
| LOC107073573 | farnesol dehydrogenase-like | 0.02077 | 105.83435 | TRUE | FALSE |
| LOC107067139 | superoxide dismutase [Cu-Zn | 0.03054 | 105.84035 | TRUE | FALSE |
| LOC107070241 | exosome complex component RRP45 | 0.02415 | 105.86203 | TRUE | FALSE |
| LOC107064918 | SKI/DACH domain-containing protein 1-like | 0.02841 | 105.87035 | TRUE | FALSE |
| LOC107068294 | transcriptional repressor protein YY1-like | 0.02606 | 105.90088 | TRUE | FALSE |
| LOC107064370 | solute carrier family 35 member G1-like | 0.06263 | 105.94705 | TRUE | FALSE |
| LOC107067852 | RNA-binding protein 4-like | 0.02595 | 105.94909 | TRUE | FALSE |
| LOC107065327 | uncharacterized aarF domain-containing pro | 0.02373 | 105.98787 | TRUE | FALSE |
| LOC107066851 | alpha-catulin | 0.03898 | 106.02562 | TRUE | FALSE |
| LOC107070839 | protein phosphatase 1 regulatory subunit 1 | 0.02461 | 106.03213 | TRUE | FALSE |
| LOC107070842 | uncharacterized protein LOC107070842 | 0.02445 | 106.08182 | TRUE | FALSE |
| LOC107073474 | uncharacterized protein LOC107073474 | 0.03725 | 106.08631 | TRUE | FALSE |

| | | | | | |
|---|---|---|---|---|---|
| LOC107074657 | guanylate kinase | 0.02201 | 106.11744 | TRUE | FALSE |
| LOC107066550 | uncharacterized protein LOC107066550 | 0.02374 | 106.20145 | TRUE | FALSE |
| LOC107072252 | ATP-dependent RNA helicase DHX8 | 0.03329 | 106.25007 | TRUE | FALSE |
| LOC107072652 | excitatory amino acid transporter 1-like | 0.02837 | 106.26509 | TRUE | FALSE |
| LOC107064834 | uncharacterized protein LOC107064834 | 0.03154 | 106.27821 | TRUE | FALSE |
| LOC107071582 | nardilysin-like | 0.07669 | 106.34484 | TRUE | FALSE |
| LOC107065718 | serum response factor homolog B-like | 0.02346 | 106.351 | TRUE | FALSE |
| LOC107070580 | uncharacterized protein LOC107070580 | 0.02219 | 106.35904 | TRUE | FALSE |
| LOC107073841 | AP-1 complex subunit beta-1 | 0.03039 | 106.37119 | TRUE | FALSE |
| LOC107068072 | cilia- and flagella-associated protein 45 | 0.02291 | 106.40004 | TRUE | FALSE |
| LOC107069910 | beta-1,3-galactosyltransferase 5-like | 0.02637 | 106.43169 | TRUE | FALSE |
| LOC107065695 | uridine-cytidine kinase-like 1 | 0.04436 | 106.45752 | TRUE | FALSE |
| LOC107071232 | uncharacterized protein LOC107071232 | 0.02497 | 106.52487 | TRUE | FALSE |
| LOC107074698 | RNA-binding protein MEX3B | 0.03888 | 106.634 | TRUE | FALSE |
| LOC107065520 | uncharacterized protein LOC107065520 | 0.0307 | 106.63426 | TRUE | FALSE |
| LOC107070981 | uncharacterized protein LOC107070981 | 0.04438 | 106.72221 | TRUE | FALSE |
| LOC107068298 | GTPase Era, mitochondrial | 0.0432 | 106.73337 | TRUE | FALSE |
| LOC107067383 | uncharacterized protein LOC107067383 | 0.02702 | 106.74812 | TRUE | FALSE |
| LOC107064800 | chaoptin-like | 0.02579 | 106.75348 | TRUE | FALSE |
| LOC107072343 | cell division cycle protein 123 homolog | 0.0242 | 106.78116 | TRUE | FALSE |
| LOC107064558 | WASH complex subunit CCDC53 | 0.02307 | 106.7833 | TRUE | FALSE |
| LOC107065710 | histone-lysine N-methyltransferase Suv4-20 | 0.04639 | 106.80212 | TRUE | FALSE |
| LOC107066047 | hemicentin-1-like | 0.02977 | 106.83194 | TRUE | FALSE |
| LOC107064907 | ES1 protein homolog, mitochondrial-like | 0.02997 | 106.88577 | TRUE | FALSE |
| LOC107068738 | RWD domain-containing protein 1 | 0.03051 | 106.9477 | TRUE | FALSE |
| LOC107070358 | uncharacterized protein LOC107070358 | 0.02509 | 106.94926 | TRUE | FALSE |
| LOC107066409 | cAMP-dependent protein kinase catalytic su | 0.04718 | 106.99487 | TRUE | FALSE |
| LOC107064328 | neuronal PAS domain-containing protein 2-l | 0.02217 | 107.19216 | TRUE | FALSE |
| LOC107066952 | piggyBac transposable element-derived pro | 0.02297 | 107.2159 | TRUE | FALSE |
| LOC107065823 | CUGBP Elav-like family member 1 | 0.0531 | 107.21616 | TRUE | FALSE |
| LOC107065037 | uncharacterized protein LOC107065037 | 0.05 | 107.231 | TRUE | FALSE |
| LOC107064291 | protein mothers against dpp | 0.03275 | 107.27236 | TRUE | FALSE |
| LOC107070985 | LOW QUALITY PROTEIN: homeobox protein | 0.10957 | 107.30402 | TRUE | FALSE |
| LOC107069399 | WAS/WASL-interacting protein family meml | 0.03547 | 107.32838 | TRUE | FALSE |
| LOC107072351 | DNA primase large subunit-like | 0.03102 | 107.33501 | TRUE | FALSE |
| LOC107068836 | putative zinc metalloproteinase YIL108W | 0.03167 | 107.33734 | TRUE | FALSE |
| LOC107067451 | uncharacterized protein LOC107067451 | 0.02572 | 107.35129 | TRUE | FALSE |
| LOC107071702 | dnaJ homolog subfamily C member 3 | 0.02994 | 107.40409 | TRUE | FALSE |
| LOC107069646 | rhomboid-related protein 2 | 0.0249 | 107.41136 | TRUE | FALSE |
| LOC107066523 | serum response factor-binding protein 1-lik | 0.02409 | 107.43572 | TRUE | FALSE |
| LOC107068587 | uncharacterized protein LOC107068587 | 0.02713 | 107.45313 | TRUE | FALSE |
| LOC107073623 | peroxidase-like | 0.02221 | 107.46021 | TRUE | FALSE |
| LOC107068937 | farnesol dehydrogenase-like | 0.08021 | 107.4903 | TRUE | FALSE |
| LOC107073001 | DNA-directed RNA polymerase III subunit RI | 0.03428 | 107.49182 | TRUE | FALSE |
| LOC107069411 | prefoldin subunit 1 | 0.02807 | 107.50492 | TRUE | FALSE |
| LOC107074239 | chromosome transmission fidelity protein 8 | 0.07052 | 107.57193 | TRUE | FALSE |
| LOC107070218 | spectrin beta chain | 0.02814 | 107.61037 | TRUE | FALSE |
| LOC107064287 | sodium/bile acid cotransporter 7-like | 0.02769 | 107.64107 | TRUE | FALSE |
| LOC107065623 | DNA-directed RNA polymerases I, II, and III s | 0.0264 | 107.66762 | TRUE | FALSE |
| LOC107063937 | DNA-directed RNA polymerase III subunit RI | 0.02393 | 107.66937 | TRUE | FALSE |

| | | | | | |
|---|---|---|---|---|---|
| LOC107067053 | pituitary homeobox x | 0.03429 | 107.67585 | TRUE | FALSE |
| LOC107074635 | BCL2/adenovirus E1B 19 kDa protein-intera | 0.03037 | 107.72451 | TRUE | FALSE |
| LOC107071888 | uncharacterized protein LOC107071888 | 0.03916 | 107.75499 | TRUE | FALSE |
| LOC107065361 | cytochrome P450 4C1-like | 0.0267 | 107.79124 | TRUE | TRUE |
| LOC107065368 | sialin-like | 0.06743 | 107.79958 | TRUE | FALSE |
| LOC107074361 | serine/threonine-protein kinase ULK2 | 0.03365 | 107.80585 | TRUE | FALSE |
| LOC107068337 | transcription initiation factor TFIID subunit 7 | 0.03761 | 107.83347 | TRUE | FALSE |
| LOC107070687 | autophagy-related protein 16-1 | 0.0363 | 107.84475 | TRUE | FALSE |
| LOC107064329 | CLIP-associating protein 1 | 0.02406 | 107.88127 | TRUE | FALSE |
| LOC107072025 | SNAPIN protein homolog | 0.03173 | 107.88491 | TRUE | FALSE |
| LOC107072530 | 2-oxoglutarate and iron-dependent oxygena | 0.14707 | 107.91538 | TRUE | FALSE |
| LOC107071994 | osteoclast-stimulating factor 1-like | 0.10321 | 107.93781 | TRUE | FALSE |
| LOC107067515 | zinc finger protein 585B-like | 0.02224 | 107.95382 | TRUE | FALSE |
| LOC107074145 | polyglutamine-binding protein 1 | 0.03031 | 108.15586 | TRUE | FALSE |
| LOC107067156 | ras-interacting protein RIP3-like | 0.0282 | 108.20391 | TRUE | FALSE |
| LOC107068577 | histidine protein methyltransferase 1 homo | 0.1279 | 108.21118 | TRUE | FALSE |
| LOC107071810 | coiled-coil domain-containing protein 43 | 0.02444 | 108.24108 | TRUE | FALSE |
| LOC107073074 | serine/threonine-protein kinase VRK1-like | 0.02299 | 108.28189 | TRUE | FALSE |
| LOC107066719 | ubiquitin-conjugating enzyme E2 J2 | 0.02471 | 108.31887 | TRUE | FALSE |
| LOC107064430 | uncharacterized protein LOC107064430 | 0.06616 | 108.36921 | TRUE | FALSE |
| LOC107073567 | uncharacterized protein LOC107073567 | 0.02668 | 108.40735 | TRUE | FALSE |
| LOC107070401 | zinc finger and BTB domain-containing prote | 0.08836 | 108.43898 | TRUE | FALSE |
| LOC107067493 | uncharacterized protein LOC107067493 | 0.02347 | 108.46206 | TRUE | FALSE |
| LOC107069716 | RNA pseudouridylate synthase domain-cont | 0.06899 | 108.46491 | TRUE | FALSE |
| LOC107072716 | peptide chain release factor 1-like, mitocho | 0.03308 | 108.50017 | TRUE | FALSE |
| LOC107069255 | sepiapterin reductase | 0.08385 | 108.50225 | TRUE | FALSE |
| LOC107070675 | adenosine monophosphate-protein transfer | 0.10387 | 108.5065 | TRUE | FALSE |
| LOC107067489 | uncharacterized protein LOC107067489 | 0.03782 | 108.53042 | TRUE | FALSE |
| LOC107074402 | protein bowel-like | 0.0303 | 108.59448 | TRUE | FALSE |
| LOC107066782 | probable tRNA (uracil-O(2)-)-methyltransfer | 0.0244 | 108.67363 | TRUE | FALSE |
| LOC107063969 | protein archease-like | 0.02712 | 108.70644 | TRUE | FALSE |
| LOC107067302 | uncharacterized protein LOC107067302 | 0.02718 | 108.72098 | TRUE | FALSE |
| LOC107073065 | protein SMG5 | 0.04707 | 108.81936 | TRUE | FALSE |
| LOC107071851 | cytochrome b-c1 complex subunit 6, mitoch | 0.04465 | 108.82062 | TRUE | FALSE |
| LOC107065890 | superoxide dismutase [Cu-Zn | 0.02593 | 108.92431 | TRUE | FALSE |
| LOC107073781 | COP9 signalosome complex subunit 5-like | 0.02768 | 108.96112 | TRUE | FALSE |
| LOC107072906 | elongation of very long chain fatty acids pro | 0.04506 | 108.97746 | TRUE | TRUE |
| LOC107074179 | phospholipase D3-like | 0.09024 | 108.99167 | TRUE | FALSE |
| LOC107067356 | basic salivary proline-rich protein 2 | 0.03102 | 109.00818 | TRUE | FALSE |
| LOC107067109 | ubiquitin-conjugating enzyme E2-17 kDa | 0.0362 | 109.03299 | TRUE | FALSE |
| LOC107063758 | RNA 3'-terminal phosphate cyclase | 0.02658 | 109.1012 | TRUE | FALSE |
| LOC107065160 | adenylate cyclase type 3-like | 0.02946 | 109.1021 | TRUE | FALSE |
| LOC107074533 | cell division control protein 45 homolog | 0.02205 | 109.10457 | TRUE | FALSE |
| LOC107067271 | autism susceptibility gene 2 protein-like | 0.04208 | 109.13758 | TRUE | FALSE |
| LOC107066315 | peptidyl-prolyl cis-trans isomerase FKBP1A | 0.02812 | 109.16786 | TRUE | FALSE |
| LOC107071203 | uncharacterized protein LOC107071203 | 0.0261 | 109.1861 | TRUE | FALSE |
| LOC107072411 | piggyBac transposable element-derived pro | 0.02935 | 109.20616 | TRUE | FALSE |
| LOC107071915 | high-affinity potassium transport protein | 0.03292 | 109.29283 | TRUE | FALSE |
| LOC107071126 | uncharacterized protein LOC107071126 | 0.03857 | 109.30423 | TRUE | FALSE |
| LOC107065698 | 60S ribosomal protein L15 | 0.04406 | 109.32701 | TRUE | FALSE |

| | | | | | |
|---|---|---|---|---|---|
| LOC107065655 | E3 ubiquitin-protein ligase UBR4 | 0.0289 | 109.33569 | TRUE | FALSE |
| LOC107073192 | protein O-mannosyltransferase 1 | 0.03311 | 109.34786 | TRUE | FALSE |
| LOC107069638 | uncharacterized protein LOC107069638 | 0.07632 | 109.35814 | TRUE | FALSE |
| LOC107073148 | amyloid protein-binding protein 2 | 0.06322 | 109.37807 | TRUE | FALSE |
| LOC107064603 | 72 kDa inositol polyphosphate 5-phosphata: | 0.03397 | 109.39883 | TRUE | FALSE |
| LOC107071899 | uncharacterized protein LOC107071899 | 0.02532 | 109.40275 | TRUE | FALSE |
| LOC107068502 | GATA zinc finger domain-containing protein | 0.0242 | 109.46035 | TRUE | FALSE |
| LOC107066674 | S-adenosylmethionine mitochondrial carrier | 0.03803 | 109.47361 | TRUE | FALSE |
| LOC107065618 | INO80 complex subunit C | 0.05534 | 109.56345 | TRUE | FALSE |
| LOC107064707 | importin-9 | 0.02308 | 109.58751 | TRUE | FALSE |
| LOC107072172 | ornithine decarboxylase antizyme 1 | 0.02668 | 109.63101 | TRUE | FALSE |
| LOC107067300 | ATP-binding cassette sub-family G member | 0.02395 | 109.71083 | TRUE | FALSE |
| LOC107067157 | zinc finger protein 13 | 0.02814 | 109.75359 | TRUE | FALSE |
| LOC107068175 | GDP-fucose protein O-fucosyltransferase 2 | 0.02315 | 109.76828 | TRUE | FALSE |
| LOC107066223 | uncharacterized protein LOC107066223 | 0.02321 | 109.80049 | TRUE | FALSE |
| LOC107066631 | proline dehydrogenase 1, mitochondrial | 0.02864 | 109.83451 | TRUE | FALSE |
| LOC107068068 | uncharacterized protein CG10915 | 0.02129 | 109.84389 | TRUE | FALSE |
| LOC107065986 | RNA-binding motif protein, X-linked 2 | 0.03157 | 109.85348 | TRUE | FALSE |
| LOC107067285 | U6 snRNA-associated Sm-like protein LSm3 | 0.04806 | 109.85773 | TRUE | FALSE |
| LOC107067320 | N-alpha-acetyltransferase 40 | 0.02604 | 109.86725 | TRUE | FALSE |
| LOC107068440 | uncharacterized protein LOC107068440 | 0.02568 | 109.89426 | TRUE | FALSE |
| LOC107068907 | histone H4 transcription factor | 0.04429 | 109.93547 | TRUE | FALSE |
| LOC107072222 | uncharacterized protein LOC107072222 | 0.1103 | 109.95502 | TRUE | FALSE |
| LOC107068279 | nucleolar protein 12-like | 0.02593 | 109.97443 | TRUE | FALSE |
| LOC107066856 | uncharacterized protein LOC107066856 | 0.03048 | 110.00374 | TRUE | FALSE |
| LOC107064651 | ER membrane protein complex subunit 6 | 0.07424 | 110.03941 | TRUE | FALSE |
| LOC107070886 | 1,2-dihydroxy-3-keto-5-methylthiopentene | 0.03235 | 110.04439 | TRUE | FALSE |
| LOC107067966 | uncharacterized protein LOC107067966 | 0.0282 | 110.08271 | TRUE | FALSE |
| LOC107069104 | alpha-N-acetylgalactosaminidase | 0.02811 | 110.12425 | TRUE | FALSE |
| LOC107069623 | uncharacterized protein LOC107069623 | 0.03977 | 110.23254 | TRUE | FALSE |
| LOC107067778 | uncharacterized protein LOC107067778 | 0.0258 | 110.2489 | TRUE | FALSE |
| LOC107073696 | uncharacterized protein LOC107073696 | 0.02368 | 110.29183 | TRUE | FALSE |
| LOC107065324 | protein eyes shut | 0.02503 | 110.32792 | TRUE | FALSE |
| LOC107066272 | homeobox protein OTX2-A | 0.02457 | 110.32896 | TRUE | FALSE |
| LOC107070599 | transcription factor Sp9 | 0.02987 | 110.36168 | TRUE | FALSE |
| LOC107073815 | targeting protein for Xklp2-like | 0.02391 | 110.38332 | TRUE | FALSE |
| LOC107068801 | CUE domain-containing protein 2 | 0.02677 | 110.38722 | TRUE | FALSE |
| LOC107070039 | uncharacterized protein LOC107070039 | 0.03701 | 110.52573 | TRUE | FALSE |
| LOC107068642 | uncharacterized protein LOC107068642 | 0.0237 | 110.53191 | TRUE | FALSE |
| LOC107072894 | uncharacterized protein LOC107072894 | 0.02746 | 110.562 | TRUE | FALSE |
| LOC107072095 | UPF0605 protein CG18335-like | 0.02656 | 110.58258 | TRUE | FALSE |
| LOC107063972 | phosphatidylinositol N-acetylglucosaminyltr | 0.04716 | 110.59694 | TRUE | FALSE |
| LOC107065425 | uncharacterized protein LOC107065425 | 0.03144 | 110.6618 | TRUE | FALSE |
| LOC107065842 | scavenger receptor class B member 1 | 0.02296 | 110.67338 | TRUE | FALSE |
| LOC107065849 | putative fatty acyl-CoA reductase CG5065 | 0.02449 | 110.69759 | TRUE | FALSE |
| LOC107065225 | V-type proton ATPase subunit d | 0.03042 | 110.73044 | TRUE | FALSE |
| LOC107071064 | Golgi resident protein GCP60 | 0.06523 | 110.76273 | TRUE | FALSE |
| LOC107072806 | actin-binding LIM protein 2 | 0.06003 | 110.78272 | TRUE | FALSE |
| LOC107072127 | cytochrome P450 4C1-like | 0.14913 | 110.82256 | TRUE | FALSE |
| LOC107069923 | proton-coupled amino acid transporter 1-lik | 0.02655 | 110.84393 | TRUE | FALSE |

| | | | | | |
|---|---|---|---|---|---|
| LOC107074380 | tRNA-splicing endonuclease subunit Sen2 | 0.0335 | 110.85226 | TRUE | FALSE |
| LOC107067789 | uncharacterized protein LOC107067789 | 0.04126 | 110.86546 | TRUE | FALSE |
| LOC107064067 | cholinesterase 2-like | 0.02942 | 110.88999 | TRUE | FALSE |
| LOC107068949 | farnesol dehydrogenase-like | 0.02631 | 110.89802 | TRUE | FALSE |
| LOC107073479 | protein toll-like | 0.02736 | 110.95066 | TRUE | FALSE |
| LOC107066300 | uncharacterized protein LOC107066300 | 0.09814 | 110.9514 | TRUE | FALSE |
| LOC107066123 | uncharacterized protein LOC107066123 | 0.0268 | 110.95161 | TRUE | FALSE |
| LOC107065926 | leptin receptor gene-related protein | 0.03713 | 111.05943 | TRUE | FALSE |
| LOC107074615 | HIG1 domain family member 2A, mitochonc | 0.0328 | 111.07698 | TRUE | FALSE |
| LOC107073695 | uncharacterized protein LOC107073695 | 0.02627 | 111.16596 | TRUE | FALSE |
| LOC107066024 | histone RNA hairpin-binding protein | 0.03466 | 111.17626 | TRUE | FALSE |
| LOC107073207 | RNA-binding protein 45 | 0.05564 | 111.26976 | TRUE | FALSE |
| LOC107071674 | homeobox protein MOX-1 | 0.0283 | 111.2824 | TRUE | FALSE |
| LOC107064138 | abhydrolase domain-containing protein 4 | 0.03734 | 111.29346 | TRUE | FALSE |
| LOC107069886 | uncharacterized protein LOC107069886 | 0.02199 | 111.32579 | TRUE | FALSE |
| LOC107071626 | glycosylphosphatidylinositol anchor attachn | 0.02973 | 111.34627 | TRUE | FALSE |
| LOC107065812 | ubiquitin carboxyl-terminal hydrolase 3-like | 0.04658 | 111.35868 | TRUE | FALSE |
| LOC107073226 | forkhead box protein E1-like | 0.02714 | 111.38131 | TRUE | FALSE |
| LOC107072574 | dystroglycan-like | 0.02605 | 111.39197 | TRUE | FALSE |
| LOC107064857 | protein enabled-like | 0.06673 | 111.40543 | TRUE | FALSE |
| LOC107071699 | uncharacterized protein LOC107071699 | 0.02426 | 111.40868 | TRUE | FALSE |
| LOC107066421 | vacuole membrane protein 1-like | 0.02878 | 111.44564 | TRUE | FALSE |
| LOC107066858 | uncharacterized protein LOC107066858 | 0.03794 | 111.48818 | TRUE | FALSE |
| LOC107064799 | chymotrypsinogen 2-like | 0.0289 | 111.537 | TRUE | FALSE |
| LOC107066334 | glycerol kinase-like | 0.02275 | 111.56284 | TRUE | FALSE |
| LOC107071757 | histone acetyltransferase KAT7 | 0.03173 | 111.65008 | TRUE | FALSE |
| LOC107073412 | uncharacterized protein LOC107073412 | 0.02726 | 111.69788 | TRUE | FALSE |
| LOC107064195 | 60S ribosomal protein L27 | 0.03826 | 111.83402 | TRUE | FALSE |
| LOC107071287 | CCHC-type zinc finger protein CG3800 | 0.02909 | 111.91743 | TRUE | FALSE |
| LOC107068511 | serum response factor homolog | 0.0449 | 111.92001 | TRUE | FALSE |
| LOC107069619 | uncharacterized protein LOC107069619 | 0.07369 | 111.94997 | TRUE | FALSE |
| LOC107072166 | protein tyrosine phosphatase domain-conta | 0.06484 | 111.97461 | TRUE | FALSE |
| LOC107073335 | uncharacterized protein LOC107073335 | 0.02891 | 111.97551 | TRUE | FALSE |
| LOC107072081 | UDP-glucose 4-epimerase-like | 0.05265 | 111.9774 | TRUE | FALSE |
| LOC107067147 | ribokinase-like | 0.04331 | 111.98715 | TRUE | FALSE |
| LOC107064345 | facilitated trehalose transporter Tret1-like | 0.07476 | 112.00043 | TRUE | FALSE |
| LOC107072320 | GPN-loop GTPase 2 | 0.1262 | 112.02652 | TRUE | FALSE |
| LOC107066977 | prefoldin subunit 4 | 0.02737 | 112.04943 | TRUE | FALSE |
| LOC107067311 | probable Rho GTPase-activating protein CG! | 0.03686 | 112.0561 | TRUE | FALSE |
| LOC107071474 | UDP-glucuronosyltransferase 2B4-like | 0.12053 | 112.10973 | TRUE | FALSE |
| LOC107072941 | exocyst complex component 2 | 0.02649 | 112.12564 | TRUE | FALSE |
| LOC107066940 | xaa-Pro aminopeptidase 1-like | 0.02137 | 112.13212 | TRUE | FALSE |
| LOC107070615 | fructose-bisphosphate aldolase-like | 0.0606 | 112.13838 | TRUE | FALSE |
| LOC107071931 | tigger transposable element-derived proteir | 0.02477 | 112.17003 | TRUE | FALSE |
| LOC107067436 | heat shock factor-binding protein 1 | 0.0263 | 112.19759 | TRUE | FALSE |
| LOC107064359 | proteasome subunit beta type-2-like | 0.04611 | 112.26012 | TRUE | FALSE |
| LOC107071783 | actin-related protein 1 | 0.07623 | 112.26967 | TRUE | FALSE |
| LOC107074483 | uncharacterized protein LOC107074483 | 0.02593 | 112.31507 | TRUE | FALSE |
| LOC107064167 | follistatin | 0.04615 | 112.39167 | TRUE | FALSE |
| LOC107068304 | methyltransferase-like protein 2-A | 0.03091 | 112.43941 | TRUE | FALSE |

| | | | | | |
|---|---|---|---|---|---|
| LOC107073024 | cell wall integrity and stress response comp | 0.02595 | 112.46806 | TRUE | FALSE |
| LOC107069140 | WW domain-containing oxidoreductase | 0.08482 | 112.52069 | TRUE | FALSE |
| LOC107063815 | huntingtin-interacting protein 1-like | 0.05987 | 112.54028 | TRUE | FALSE |
| LOC107065458 | vesicle transport protein SFT2B | 0.03578 | 112.5554 | TRUE | FALSE |
| LOC107064802 | prefoldin subunit 6 | 0.02365 | 112.64289 | TRUE | FALSE |
| LOC107070095 | zinc finger protein 37-like | 0.03019 | 112.65054 | TRUE | FALSE |
| LOC107072874 | anaphase-promoting complex subunit 10 | 0.07403 | 112.66161 | TRUE | FALSE |
| LOC107064836 | uncharacterized protein LOC107064836 | 0.02405 | 112.67878 | TRUE | FALSE |
| LOC107072805 | TNF receptor-associated factor 5 | 0.03504 | 112.71357 | TRUE | FALSE |
| LOC107067126 | neural Wiskott-Aldrich syndrome protein | 0.04096 | 112.7347 | TRUE | FALSE |
| LOC107069979 | uncharacterized protein LOC107069979 | 0.02531 | 112.74723 | TRUE | FALSE |
| LOC107068601 | cytochrome P450 4g15 | 0.04332 | 112.81053 | TRUE | TRUE |
| LOC107067249 | uncharacterized protein LOC107067249 | 0.08163 | 112.81932 | TRUE | FALSE |
| LOC107069963 | U6 snRNA phosphodiesterase | 0.02456 | 112.82605 | TRUE | FALSE |
| LOC107064428 | uncharacterized protein LOC107064428 | 0.02382 | 112.93667 | TRUE | FALSE |
| LOC107067401 | uncharacterized protein LOC107067401 | 0.02382 | 112.95675 | TRUE | FALSE |
| LOC107070426 | F-box/LRR-repeat protein 3-like | 0.02297 | 112.99841 | TRUE | FALSE |
| LOC107073309 | condensin complex subunit 2-like | 0.0338 | 113.03758 | TRUE | FALSE |
| LOC107069306 | uncharacterized protein LOC107069306 | 0.03941 | 113.07358 | TRUE | FALSE |
| LOC107070786 | uncharacterized protein LOC107070786 | 0.02889 | 113.09527 | TRUE | FALSE |
| LOC107064556 | sentrin-specific protease 8-like | 0.02176 | 113.16857 | TRUE | FALSE |
| LOC107065678 | magnesium-dependent phosphatase 1-like | 0.02577 | 113.19631 | TRUE | FALSE |
| LOC107065549 | protein disulfide-isomerase A6 | 0.10957 | 113.23631 | TRUE | FALSE |
| LOC107068794 | stress-induced-phosphoprotein 1 | 0.02699 | 113.26691 | TRUE | FALSE |
| LOC107072072 | uncharacterized protein LOC107072072 | 0.03473 | 113.29717 | TRUE | FALSE |
| LOC107064536 | uncharacterized protein LOC107064536 | 0.02345 | 113.29934 | TRUE | FALSE |
| LOC107073117 | uncharacterized protein LOC107073117 | 0.03535 | 113.30657 | TRUE | FALSE |
| LOC107063663 | ecdysone receptor | 0.0601 | 113.3315 | TRUE | FALSE |
| LOC107067392 | CD151 antigen-like | 0.05837 | 113.36495 | TRUE | FALSE |
| LOC107071856 | cytosolic endo-beta-N-acetylglucosaminidas | 0.05707 | 113.38174 | TRUE | FALSE |
| LOC107065038 | uncharacterized protein LOC107065038 | 0.09831 | 113.42853 | TRUE | FALSE |
| LOC107068135 | lysosomal-trafficking regulator | 0.026 | 113.44054 | TRUE | FALSE |
| LOC107072761 | uncharacterized protein LOC107072761 | 0.04285 | 113.52352 | TRUE | FALSE |
| LOC107070975 | dolichyl-diphosphooligosaccharide--protein | 0.1496 | 113.56051 | TRUE | FALSE |
| LOC107074379 | uncharacterized protein LOC107074379 | 0.02915 | 113.57167 | TRUE | FALSE |
| LOC107067095 | nucleoside diphosphate kinase 6 | 0.02538 | 113.59667 | TRUE | FALSE |
| LOC107073871 | peptidyl-prolyl cis-trans isomerase 5 | 0.03327 | 113.68977 | TRUE | FALSE |
| LOC107072164 | putative mediator of RNA polymerase II tra | 0.04588 | 113.70632 | TRUE | TRUE |
| LOC107064875 | blood vessel epicardial substance-A-like | 0.04843 | 113.7099 | TRUE | FALSE |
| LOC107070154 | xylosyltransferase oxt | 0.0898 | 113.72366 | TRUE | FALSE |
| LOC107066954 | uncharacterized protein LOC107066954 | 0.02491 | 113.74451 | TRUE | FALSE |
| LOC107070052 | LON peptidase N-terminal domain and RINC | 0.05797 | 113.8164 | TRUE | FALSE |
| LOC107063914 | peptide transporter family 1 | 0.02313 | 113.8191 | TRUE | FALSE |
| LOC107071294 | voltage-dependent anion-selective channel- | 0.03249 | 113.8349 | TRUE | FALSE |
| LOC107074445 | pleckstrin homology domain-containing fan | 0.03317 | 113.83967 | TRUE | FALSE |
| LOC107065359 | cytochrome P450 4C1-like | 0.02898 | 113.87277 | TRUE | TRUE |
| LOC107063684 | A-kinase anchor protein 9-like | 0.06515 | 113.88839 | TRUE | FALSE |
| LOC107074599 | BTB/POZ domain-containing protein 7 | 0.05463 | 113.90223 | TRUE | FALSE |
| LOC107068108 | uncharacterized protein LOC107068108 | 0.03092 | 113.90546 | TRUE | FALSE |
| LOC107068739 | NADP-dependent malic enzyme | NA | 113.97644 | TRUE | FALSE |

| | | | | | |
|---|---|---|---|---|---|
| LOC107072064 | alpha-tocopherol transfer protein-like | 0.0958 | 113.98986 | TRUE | FALSE |
| LOC107072994 | uncharacterized protein LOC107072994 | 0.0375 | 114.00837 | TRUE | FALSE |
| LOC107072801 | dual specificity protein phosphatase CDC14I | 0.0266 | 114.01535 | TRUE | FALSE |
| LOC107068880 | uncharacterized protein LOC107068880 | 0.04429 | 114.03616 | TRUE | FALSE |
| LOC107066324 | protein PFC0760c-like | 0.05712 | 114.07355 | TRUE | FALSE |
| LOC107073558 | uncharacterized protein LOC107073558 | 0.0511 | 114.14322 | TRUE | FALSE |
| LOC107074086 | dynein heavy chain 8, axonemal | 0.04913 | 114.21752 | TRUE | FALSE |
| LOC107072175 | uncharacterized protein LOC107072175 | 0.02358 | 114.2207 | TRUE | FALSE |
| LOC107067236 | UDP-N-acetylglucosamine transferase subu | 0.0229 | 114.25257 | TRUE | FALSE |
| LOC107071719 | type 1 phosphatidylinositol 4,5-bisphosphat | 0.03222 | 114.26896 | TRUE | FALSE |
| LOC107072678 | probable nuclear hormone receptor HR3 | 0.04747 | 114.30777 | TRUE | FALSE |
| LOC107064733 | uncharacterized protein LOC107064733 | 0.02338 | 114.3399 | TRUE | FALSE |
| LOC107069840 | transmembrane protein 41B | 0.03601 | 114.35173 | TRUE | FALSE |
| LOC107065756 | homogentisate 1,2-dioxygenase | 0.02899 | 114.36964 | TRUE | FALSE |
| LOC107073725 | uncharacterized protein LOC107073725 | 0.02169 | 114.38083 | TRUE | FALSE |
| LOC107065087 | putative uncharacterized protein DDB_G02 | 0.03269 | 114.39472 | TRUE | FALSE |
| LOC107072353 | uncharacterized protein LOC107072353 | 0.02583 | 114.46845 | TRUE | FALSE |
| LOC107066613 | ADP-ribosylation factor GTPase-activating p | 0.09378 | 114.52283 | TRUE | FALSE |
| LOC107067254 | myb-like protein D | 0.03223 | 114.55723 | TRUE | FALSE |
| LOC107065981 | protein PFC0760c-like | 0.02644 | 114.59282 | TRUE | FALSE |
| LOC107068084 | uncharacterized protein LOC107068084 | 0.0273 | 114.61199 | TRUE | FALSE |
| LOC107066250 | protein dachsous | 0.05579 | 114.61204 | TRUE | FALSE |
| LOC107067188 | arylphorin subunit beta-like | 0.0283 | 114.65978 | TRUE | TRUE |
| LOC107073531 | peptidyl-alpha-hydroxyglycine alpha-amidat | 0.04065 | 114.77214 | TRUE | FALSE |
| LOC107064096 | tubulin beta chain-like | 0.10462 | 114.80018 | TRUE | FALSE |
| LOC107064951 | uncharacterized protein LOC107064951 | 0.03036 | 114.82566 | TRUE | FALSE |
| LOC107065009 | uncharacterized protein LOC107065009 | 0.03055 | 114.83864 | TRUE | FALSE |
| LOC107064211 | DNA-3-methyladenine glycosylase-like | 0.02734 | 114.8452 | TRUE | FALSE |
| LOC107063693 | uncharacterized protein KIAA0195 | 0.03574 | 114.85152 | TRUE | FALSE |
| LOC107063865 | uncharacterized protein LOC107063865 | 0.03568 | 114.85838 | TRUE | TRUE |
| LOC107068942 | 40S ribosomal protein S13-like | 0.03503 | 114.8619 | TRUE | FALSE |
| LOC107065433 | protein PET100 homolog, mitochondrial | 0.0307 | 115.02475 | TRUE | FALSE |
| LOC107067228 | uncharacterized protein LOC107067228 | 0.02332 | 115.05715 | TRUE | FALSE |
| LOC107073457 | uncharacterized protein LOC107073457 | 0.02609 | 115.08128 | TRUE | FALSE |
| LOC107067284 | putative inorganic phosphate cotransporter | 0.03186 | 115.08503 | TRUE | FALSE |
| LOC107072161 | uncharacterized protein LOC107072161 | 0.02888 | 115.09335 | TRUE | FALSE |
| LOC107070818 | leucine-rich repeats and immunoglobulin-lik | 0.03968 | 115.11158 | TRUE | FALSE |
| LOC107069241 | vesicle transport protein USE1 | 0.03253 | 115.11317 | TRUE | FALSE |
| LOC107074298 | MORN repeat-containing protein 3-like | 0.03265 | 115.12116 | TRUE | FALSE |
| LOC107065592 | H/ACA ribonucleoprotein complex non-core | 0.05152 | 115.12302 | TRUE | FALSE |
| LOC107070462 | transcription initiation factor IIB | 0.02383 | 115.15884 | TRUE | FALSE |
| LOC107074338 | RING-box protein 2 | 0.02694 | 115.17077 | TRUE | FALSE |
| LOC107067399 | dolichyl-phosphate beta-glucosyltransferase | 0.02888 | 115.21416 | TRUE | FALSE |
| LOC107069003 | calcyclin-binding protein | 0.02992 | 115.23002 | TRUE | FALSE |
| LOC107066230 | box A-binding factor-like | 0.03163 | 115.25854 | TRUE | FALSE |
| LOC107073128 | surfeit locus protein 1 | 0.09536 | 115.3082 | TRUE | FALSE |
| LOC107071136 | dentin sialophosphoprotein-like | 0.06274 | 115.31213 | TRUE | TRUE |
| LOC107072584 | trafficking protein particle complex subunit | 0.04324 | 115.3472 | TRUE | FALSE |
| LOC107066053 | protein jagged-1 | 0.06448 | 115.40256 | TRUE | FALSE |
| LOC107068569 | uncharacterized protein LOC107068569 | 0.04779 | 115.44126 | TRUE | FALSE |

| | | | | | |
|---|---|---|---|---|---|
| LOC107068341 | evolutionarily conserved signaling intermed | 0.08724 | 115.4943 | TRUE | FALSE |
| LOC107071900 | uncharacterized protein LOC107071900 | 0.03803 | 115.50375 | TRUE | FALSE |
| LOC107068875 | uncharacterized protein LOC107068875 | 0.02508 | 115.52891 | TRUE | FALSE |
| LOC107072955 | uncharacterized protein LOC107072955 | 0.1103 | 115.54392 | TRUE | FALSE |
| LOC107064085 | sodium/calcium exchanger 1 | 0.07252 | 115.58437 | TRUE | FALSE |
| LOC107066312 | dosage compensation regulator | 0.04512 | 115.6095 | TRUE | FALSE |
| LOC107065096 | DAZ-associated protein 2-like | 0.04615 | 115.668 | TRUE | FALSE |
| LOC107064684 | uncharacterized protein LOC107064684 | 0.05964 | 115.68482 | TRUE | FALSE |
| LOC107074252 | uncharacterized protein LOC107074252 | 0.02801 | 115.69224 | TRUE | FALSE |
| LOC107069799 | ankyrin repeat domain-containing protein S | 0.05168 | 115.70778 | TRUE | FALSE |
| LOC107063875 | uncharacterized protein LOC107063875 | 0.10578 | 115.75537 | TRUE | FALSE |
| LOC107074213 | glycine receptor subunit alpha-4 | 0.07798 | 115.77045 | TRUE | FALSE |
| LOC107065519 | chymotrypsin inhibitor-like | 0.06362 | 115.80421 | TRUE | FALSE |
| LOC107065351 | histone-lysine N-methyltransferase SETMAF | 0.02454 | 115.82757 | TRUE | FALSE |
| LOC107064378 | mitochondrial 2-oxoglutarate/malate carrie | 0.03142 | 115.84274 | TRUE | FALSE |
| LOC107064364 | uncharacterized protein LOC107064364 | 0.0783 | 115.95182 | TRUE | FALSE |
| LOC107070589 | chondroitin sulfate proteoglycan 4 | 0.05261 | 115.99661 | TRUE | FALSE |
| LOC107066288 | microtubule-associated protein RP/EB famil | 0.03083 | 116.01375 | TRUE | FALSE |
| LOC107072376 | uncharacterized membrane protein DDB_G( | 0.13127 | 116.01889 | TRUE | FALSE |
| LOC107068132 | probable cation-transporting ATPase 13A3 | 0.02837 | 116.07787 | TRUE | FALSE |
| LOC107066317 | DENN domain-containing protein 4C | 0.0334 | 116.09521 | TRUE | FALSE |
| LOC107073136 | zinc finger protein 28-like | 0.03912 | 116.10155 | TRUE | FALSE |
| LOC107072321 | uncharacterized protein LOC107072321 | 0.02945 | 116.11738 | TRUE | FALSE |
| LOC107064509 | protein lin-9 homolog | 0.03522 | 116.1657 | TRUE | FALSE |
| LOC107066756 | peptidyl-prolyl cis-trans isomerase H | 0.03251 | 116.1778 | TRUE | FALSE |
| LOC107073576 | uncharacterized protein LOC107073576 | 0.02628 | 116.22308 | TRUE | FALSE |
| LOC107069137 | protein gustavus | 0.03789 | 116.2645 | TRUE | FALSE |
| LOC107064693 | uncharacterized protein LOC107064693 | 0.07574 | 116.30308 | TRUE | FALSE |
| LOC107065780 | putative uncharacterized protein DDB_G028 | 0.09618 | 116.30801 | TRUE | FALSE |
| LOC107070668 | cyclin-dependent kinase 9-like | 0.03601 | 116.44191 | TRUE | FALSE |
| LOC107070122 | EH domain-containing protein 3 | 0.04746 | 116.44821 | TRUE | FALSE |
| LOC107067579 | probable elongator complex protein 3 | 0.07644 | 116.49786 | TRUE | FALSE |
| LOC107065228 | uncharacterized protein LOC107065228 | 0.02612 | 116.57221 | TRUE | FALSE |
| LOC107068371 | PHD finger protein 12 | 0.04137 | 116.58222 | TRUE | FALSE |
| LOC107071500 | uncharacterized protein LOC107071500 | 0.07946 | 116.60268 | TRUE | FALSE |
| LOC107067009 | uncharacterized protein LOC107067009 | 0.10571 | 116.74348 | TRUE | TRUE |
| LOC107070714 | uncharacterized protein LOC107070714 | 0.02987 | 116.74571 | TRUE | FALSE |
| LOC107072205 | cyclic AMP-dependent transcription factor / | 0.02703 | 116.79869 | TRUE | FALSE |
| LOC107067363 | rabenosyn-5 | 0.04031 | 116.83248 | TRUE | FALSE |
| LOC107068035 | zinc finger protein 391-like | 0.03368 | 116.86851 | TRUE | FALSE |
| LOC107070773 | uncharacterized protein LOC107070773 | 0.07555 | 116.87114 | TRUE | FALSE |
| LOC107070987 | carboxyl-terminal PDZ ligand of neuronal ni | 0.08487 | 116.8868 | TRUE | FALSE |
| LOC107065102 | arginyl-tRNA--protein transferase 1 | 0.05076 | 116.91132 | TRUE | FALSE |
| LOC107064108 | uncharacterized protein LOC107064108 | 0.02628 | 116.96314 | TRUE | FALSE |
| LOC107064204 | BRCA1-associated RING domain protein 1-lil | 0.1141 | 116.97523 | TRUE | FALSE |
| LOC107071841 | rRNA-processing protein FYV7 | 0.14457 | 116.97955 | TRUE | FALSE |
| LOC107070314 | CSC1-like protein 2 | 0.03061 | 116.99596 | TRUE | FALSE |
| LOC107071185 | uncharacterized protein LOC107071185 | 0.04658 | 116.99841 | TRUE | FALSE |
| LOC107068414 | pro-resilin-like | 0.04371 | 117.0007 | TRUE | FALSE |
| LOC107073933 | acyl-CoA-binding domain-containing proteir | 0.03254 | 117.08376 | TRUE | FALSE |

| | | | | | |
|---|---|---|---|---|---|
| LOC107069298 | myrosinase 1-like | 0.09721 | 117.08789 | TRUE | TRUE |
| LOC107074287 | ribose-phosphate pyrophosphokinase 1 | 0.02961 | 117.08922 | TRUE | FALSE |
| LOC107067665 | C-type lectin mannose-binding isoform-like | 0.06339 | 117.19649 | TRUE | FALSE |
| LOC107071150 | uncharacterized protein LOC107071150 | 0.02566 | 117.20031 | TRUE | FALSE |
| LOC107065353 | venom dipeptidyl peptidase 4 | 0.02271 | 117.2274 | TRUE | FALSE |
| LOC107071446 | phosphorylase b kinase gamma catalytic cha | 0.03448 | 117.25682 | TRUE | FALSE |
| LOC107073124 | ragulator complex protein LAMTOR2 homol | 0.02397 | 117.35566 | TRUE | FALSE |
| LOC107074706 | segmentation protein Runt-like | 0.02356 | 117.37467 | TRUE | FALSE |
| LOC107070751 | zinc finger CCCH domain-containing protein | 0.04985 | 117.46742 | TRUE | FALSE |
| LOC107064084 | serine protease inhibitor dipetalogastin | 0.04146 | 117.49415 | TRUE | FALSE |
| LOC107072537 | ataxin-7-like protein 3 | 0.02685 | 117.5133 | TRUE | FALSE |
| LOC107072077 | UPF0047 protein YjbQ-like | 0.09001 | 117.54277 | TRUE | FALSE |
| LOC107069728 | mitochondrial folate transporter/carrier | 0.1218 | 117.56796 | TRUE | FALSE |
| LOC107065173 | histone-lysine N-methyltransferase SUV39H | 0.07331 | 117.66269 | TRUE | FALSE |
| LOC107073332 | uncharacterized protein LOC107073332 | 0.02937 | 117.6822 | TRUE | FALSE |
| LOC107067378 | dynein heavy chain 7, axonemal | 0.06771 | 117.68732 | TRUE | FALSE |
| LOC107069591 | transmembrane protein 53 | 0.0347 | 117.78498 | TRUE | FALSE |
| LOC107071912 | regulator of nonsense transcripts 1 homolo | 0.02915 | 117.78784 | TRUE | FALSE |
| LOC107064662 | T-complex protein 1 subunit zeta | 0.04653 | 117.83362 | TRUE | FALSE |
| LOC107074007 | 5'-AMP-activated protein kinase subunit be | 0.02784 | 117.86019 | TRUE | FALSE |
| LOC107068330 | protein Tob1 | 0.10569 | 117.88611 | TRUE | FALSE |
| LOC107071506 | clathrin interactor 1 | 0.04278 | 117.90549 | TRUE | FALSE |
| LOC107072689 | putative uncharacterized protein DDB_G02 | 0.05735 | 117.92865 | TRUE | FALSE |
| LOC107067498 | beta-1,3-galactosyltransferase brn | 0.03819 | 117.96824 | TRUE | FALSE |
| LOC107067301 | ATP-binding cassette sub-family G member | 0.0511 | 118.03001 | TRUE | FALSE |
| LOC107070404 | S-phase kinase-associated protein 1 | 0.04635 | 118.12991 | TRUE | FALSE |
| LOC107068819 | uncharacterized protein LOC107068819 | 0.08723 | 118.15525 | TRUE | FALSE |
| LOC107067783 | E3 ubiquitin-protein ligase MIB1 | 0.0378 | 118.17594 | TRUE | FALSE |
| LOC107071363 | radial spoke head 10 homolog B-like | 0.08503 | 118.21864 | TRUE | FALSE |
| LOC107072243 | tyrocidine synthase 3 | 0.05117 | 118.23689 | TRUE | FALSE |
| LOC107073774 | uncharacterized protein LOC107073774 | 0.03001 | 118.27971 | TRUE | FALSE |
| LOC107067885 | histone-lysine N-methyltransferase 2D-like | 0.03332 | 118.28565 | TRUE | FALSE |
| LOC107074557 | RNA polymerase II elongation factor Ell-like | 0.16417 | 118.2972 | TRUE | FALSE |
| LOC107063738 | broad-complex core protein isoforms 1/2/3 | 0.03617 | 118.31765 | TRUE | FALSE |
| LOC107074543 | uncharacterized protein LOC107074543 | 0.0496 | 118.33395 | TRUE | FALSE |
| LOC107067594 | peptide deformylase, mitochondrial-like | 0.04353 | 118.38646 | TRUE | FALSE |
| LOC107069165 | synaptosomal-associated protein 25 | 0.05693 | 118.41776 | TRUE | FALSE |
| LOC107070669 | catalase | 0.05648 | 118.41792 | TRUE | FALSE |
| LOC107065223 | dihydroorotate dehydrogenase (quinone), n | 0.10886 | 118.44756 | TRUE | FALSE |
| LOC107072247 | 28S ribosomal protein S27, mitochondrial-li | 0.06619 | 118.46386 | TRUE | FALSE |
| LOC107065876 | double-strand break repair protein MRE11A | 0.05765 | 118.4729 | TRUE | FALSE |
| LOC107071331 | kynurenine 3-monooxygenase | 0.1798 | 118.48953 | TRUE | FALSE |
| LOC107070911 | membrane-associated protein Hem | 0.03264 | 118.50857 | TRUE | FALSE |
| LOC107071396 | folylpolyglutamate synthase, mitochondrial | 0.02776 | 118.52081 | TRUE | FALSE |
| LOC107070242 | casein kinase II subunit alpha | 0.11 | 118.52658 | TRUE | FALSE |
| LOC107063768 | uncharacterized protein LOC107063768 | 0.03667 | 118.54525 | TRUE | FALSE |
| LOC107073872 | GPI inositol-deacylase | 0.03691 | 118.55195 | TRUE | FALSE |
| LOC107071547 | double-stranded RNA-binding protein Stauf | 0.1771 | 118.56048 | TRUE | FALSE |
| LOC107071871 | leucine-rich repeat-containing protein 51-lik | 0.056 | 118.60712 | TRUE | FALSE |
| LOC107071727 | E3 ubiquitin-protein ligase RNF123-like | 0.04668 | 118.61635 | TRUE | FALSE |

| | | | | | |
|---|---|---|---|---|---|
| LOC107071875 | peroxisomal targeting signal 1 receptor | 0.04266 | 118.71978 | TRUE | FALSE |
| LOC107070212 | silk gland factor 1-like | 0.06169 | 118.72425 | TRUE | FALSE |
| LOC107065342 | TNF receptor-associated factor 4 | 0.02706 | 118.73079 | TRUE | FALSE |
| LOC107071292 | RNA-binding protein 40 | 0.03659 | 118.83865 | TRUE | FALSE |
| LOC107070392 | putative transcription factor SOX-14 | 0.02528 | 118.89617 | TRUE | FALSE |
| LOC107073392 | uncharacterized protein LOC107073392 | 0.035 | 119.01416 | TRUE | FALSE |
| LOC107073956 | collagen type IV alpha-3-binding protein | 0.02944 | 119.03769 | TRUE | FALSE |
| LOC107067265 | RNA-binding protein cabeza | 0.04237 | 119.04758 | TRUE | FALSE |
| LOC107074138 | folliculin | 0.03402 | 119.05115 | TRUE | FALSE |
| LOC107066547 | uncharacterized protein LOC107066547 | 0.05119 | 119.05928 | TRUE | FALSE |
| LOC107064748 | dephospho-CoA kinase | 0.18773 | 119.0633 | TRUE | FALSE |
| LOC107074437 | nuclear factor 1 X-type | 0.05801 | 119.18401 | TRUE | FALSE |
| LOC107064946 | peflin | 0.043 | 119.19266 | TRUE | FALSE |
| LOC107066517 | upstream stimulatory factor 2-like | 0.02537 | 119.24109 | TRUE | FALSE |
| LOC107071397 | probable uridine-cytidine kinase | 0.02778 | 119.31534 | TRUE | FALSE |
| LOC107072783 | secretin receptor-like | 0.03693 | 119.34259 | TRUE | FALSE |
| LOC107068480 | exosome complex exonuclease RRP42-like | 0.06986 | 119.35252 | TRUE | FALSE |
| LOC107073973 | WD and tetratricopeptide repeats protein 1 | 0.03873 | 119.35781 | TRUE | FALSE |
| LOC107069060 | stress response protein NST1-like | 0.06672 | 119.38272 | TRUE | FALSE |
| LOC107068430 | cytochrome P450 6a2-like | 0.0252 | 119.38591 | TRUE | FALSE |
| LOC107074584 | DNA-dependent protein kinase catalytic sub | 0.04685 | 119.40417 | TRUE | FALSE |
| LOC107066460 | uncharacterized protein LOC107066460 | 0.05728 | 119.43465 | TRUE | FALSE |
| LOC107064218 | bipolar kinesin KRP-130-like | 0.11817 | 119.52999 | TRUE | FALSE |
| LOC107071284 | stimulator of interferon genes protein | 0.03886 | 119.5841 | TRUE | FALSE |
| LOC107068891 | growth hormone secretagogue receptor typ | 0.12113 | 119.59087 | TRUE | FALSE |
| LOC107072279 | stress response protein NST1 | 0.06997 | 119.60279 | TRUE | TRUE |
| LOC107063935 | GTP-binding protein Rheb homolog | 0.02938 | 119.63329 | TRUE | FALSE |
| LOC107068221 | bromodomain-containing protein 7 | 0.04029 | 119.64431 | TRUE | FALSE |
| LOC107067592 | tachykinin-like peptides receptor 99D | 0.0335 | 119.64986 | TRUE | FALSE |
| LOC107068679 | protein Simiate | 0.02746 | 119.68074 | TRUE | FALSE |
| LOC107068970 | 28S ribosomal protein S15, mitochondrial | 0.03415 | 119.83812 | TRUE | FALSE |
| LOC107071623 | lysozyme-like | 0.02869 | 119.94178 | TRUE | FALSE |
| LOC107063765 | uncharacterized protein LOC107063765 | 0.1028 | 119.94584 | TRUE | FALSE |
| LOC107069846 | protein fem-1 homolog CG6966 | 0.03931 | 119.95141 | TRUE | FALSE |
| LOC107068846 | ADP,ATP carrier protein-like | 0.03697 | 119.97563 | TRUE | FALSE |
| LOC107065162 | ubiquitin-conjugating enzyme E2 variant 2 | 0.09733 | 119.97772 | TRUE | FALSE |
| LOC107074023 | DNA helicase MCM9-like | 0.0355 | 120.01463 | TRUE | FALSE |
| LOC107067842 | uncharacterized protein LOC107067842 | 0.0331 | 120.02424 | TRUE | FALSE |
| LOC107073350 | uncharacterized protein LOC107073350 | 0.04073 | 120.02789 | TRUE | FALSE |
| LOC107064494 | protein phosphatase methylesterase 1 | 0.04049 | 120.05503 | TRUE | FALSE |
| LOC107068160 | enhancer of polycomb homolog 1 | 0.04263 | 120.06122 | TRUE | FALSE |
| LOC107067644 | mitochondrial ribosome-associated GTPase | 0.02592 | 120.17046 | TRUE | FALSE |
| LOC107064623 | basic helix-loop-helix neural transcription fa | 0.0367 | 120.17809 | TRUE | FALSE |
| LOC107065011 | peptidyl-prolyl cis-trans isomerase CWC27 h | 0.02628 | 120.22462 | TRUE | FALSE |
| LOC107068950 | uncharacterized protein LOC107068950 | 0.03183 | 120.24509 | TRUE | FALSE |
| LOC107068852 | zinc finger protein 729-like | 0.02932 | 120.35278 | TRUE | FALSE |
| LOC107073360 | uncharacterized protein LOC107073360 | 0.03046 | 120.37912 | TRUE | FALSE |
| LOC107068791 | cytosolic purine 5'-nucleotidase | 0.06913 | 120.38475 | TRUE | FALSE |
| LOC107063828 | defensin-1-like | 0.02757 | 120.39271 | TRUE | FALSE |
| LOC107067317 | palmitoyltransferase ZDHHC5 | 0.03497 | 120.47607 | TRUE | FALSE |

| | | | | | |
|---|---|---|---|---|---|
| LOC107071111 | DNA-directed RNA polymerase I subunit RP/ | 0.04023 | 120.49293 | TRUE | FALSE |
| LOC107065858 | DNA-binding protein D-ETS-6-like | 0.03759 | 120.49644 | TRUE | FALSE |
| LOC107069134 | 2',5'-phosphodiesterase 12 | 0.1459 | 120.52955 | TRUE | FALSE |
| LOC107066951 | cytochrome P450 4C1-like | 0.04675 | 120.64543 | TRUE | FALSE |
| LOC107066385 | neuroparsin-A-like | 0.05697 | 120.74686 | TRUE | FALSE |
| LOC107065938 | uncharacterized protein LOC107065938 | 0.04079 | 120.77136 | TRUE | FALSE |
| LOC107066072 | G2/mitotic-specific cyclin-B | 0.13473 | 120.81421 | TRUE | FALSE |
| LOC107074438 | uncharacterized protein LOC107074438 | 0.03396 | 120.82143 | TRUE | FALSE |
| LOC107068432 | cactin | 0.06412 | 120.82187 | TRUE | FALSE |
| LOC107066689 | dystonin | 0.04787 | 120.86312 | TRUE | FALSE |
| LOC107068776 | serine/threonine-protein kinase greatwall | 0.06225 | 120.87541 | TRUE | FALSE |
| LOC107066528 | dynein heavy chain 7, axonemal-like | 0.04827 | 120.93784 | TRUE | FALSE |
| LOC107074166 | 40S ribosomal protein SA | 0.1661 | 120.96623 | TRUE | FALSE |
| LOC107068589 | potassium voltage-gated channel protein Sh | 0.06815 | 121.02069 | TRUE | FALSE |
| LOC107072507 | atypical protein kinase C | 0.04468 | 121.14315 | TRUE | FALSE |
| LOC107066660 | uncharacterized protein LOC107066660 | 0.05944 | 121.25208 | TRUE | FALSE |
| LOC107068087 | proteasome subunit beta type-3 | 0.03846 | 121.32551 | TRUE | FALSE |
| LOC107063887 | cullin-1 | 0.04142 | 121.53979 | TRUE | FALSE |
| LOC107069677 | uncharacterized protein F02A9.4b | 0.03562 | 121.58216 | TRUE | FALSE |
| LOC107074029 | uncharacterized protein PFB0765w-like | 0.05814 | 121.64177 | TRUE | FALSE |
| LOC107071973 | Down syndrome cell adhesion molecule-like | 0.03747 | 121.71029 | TRUE | FALSE |
| LOC107070040 | RNA-binding protein squid-like | 0.09356 | 121.79485 | TRUE | FALSE |
| LOC107066392 | piwi-like protein 1 | 0.18247 | 121.80117 | TRUE | FALSE |
| LOC107072060 | serine/threonine-protein phosphatase alpha | 0.03347 | 121.87807 | TRUE | FALSE |
| LOC107071665 | uncharacterized protein LOC107071665 | 0.06812 | 121.89157 | TRUE | FALSE |
| LOC107069941 | activating signal cointegrator 1 complex sub | 0.05749 | 121.89537 | TRUE | FALSE |
| LOC107064392 | leucine-rich repeat-containing protein 23-lik | 0.02805 | 121.99462 | TRUE | FALSE |
| LOC107071313 | uncharacterized protein LOC107071313 | 0.03787 | 121.99743 | TRUE | FALSE |
| LOC107067764 | odorant receptor 4-like | 0.05895 | 122.05496 | TRUE | FALSE |
| LOC107073202 | probable galactose-1-phosphate uridylyltran | 0.14147 | 122.16092 | TRUE | FALSE |
| LOC107067654 | protein FAM195A-like | 0.06128 | 122.17879 | TRUE | FALSE |
| LOC107070167 | E3 ubiquitin-protein ligase znrf1 | 0.05425 | 122.18396 | TRUE | FALSE |
| LOC107070219 | peroxiredoxin-2-like | 0.05018 | 122.22113 | TRUE | FALSE |
| LOC107063946 | uncharacterized protein LOC107063946 | 0.04551 | 122.4054 | TRUE | FALSE |
| LOC107065164 | transmembrane protein 18 | 0.15487 | 122.41368 | TRUE | FALSE |
| LOC107065212 | PHD finger-like domain-containing protein 5 | 0.02426 | 122.41896 | TRUE | FALSE |
| LOC107064717 | uncharacterized protein LOC107064717 | 0.04843 | 122.53592 | TRUE | FALSE |
| LOC107068194 | lysM and putative peptidoglycan-binding do | 0.02936 | 122.54235 | TRUE | FALSE |
| LOC107065523 | zinc finger protein Xfin-like | 0.06483 | 122.59426 | TRUE | FALSE |
| LOC107069835 | uncharacterized protein LOC107069835 | 0.02974 | 122.64862 | TRUE | FALSE |
| LOC107064609 | CCAAT/enhancer-binding protein | 0.05246 | 122.66128 | TRUE | FALSE |
| LOC107071684 | uncharacterized protein LOC107071684 | 0.1535 | 122.69402 | TRUE | FALSE |
| LOC107068081 | probable 18S rRNA (guanine-N(7))-methyltr | 0.02987 | 122.71704 | TRUE | FALSE |
| LOC107068020 | spatacsin | 0.07115 | 122.73993 | TRUE | FALSE |
| LOC107071023 | uncharacterized protein LOC107071023 | 0.07503 | 122.75771 | TRUE | FALSE |
| LOC107072540 | GTP cyclohydrolase 1 | 0.03073 | 122.78884 | TRUE | FALSE |
| LOC107072696 | CDP-diacylglycerol--inositol 3-phosphatidylt | 0.12927 | 122.8405 | TRUE | FALSE |
| LOC107066991 | uncharacterized protein LOC107066991 | 0.03382 | 122.89368 | TRUE | FALSE |
| LOC107071301 | uncharacterized protein LOC107071301 | 0.04854 | 122.96733 | TRUE | FALSE |
| LOC107068945 | AP-3 complex subunit mu-1 | 0.026 | 122.97337 | TRUE | FALSE |

| | | | | | |
|---|---|---|---|---|---|
| LOC107069670 | TELO2-interacting protein 1 homolog | 0.04331 | 123.03342 | TRUE | FALSE |
| LOC107070319 | odorant receptor 49b-like | 0.03301 | 123.03496 | TRUE | FALSE |
| LOC107071353 | GPI ethanolamine phosphate transferase 1 | 0.07393 | 123.06185 | TRUE | FALSE |
| LOC107063686 | solute carrier family 25 member 36-A | 0.04844 | 123.10015 | TRUE | FALSE |
| LOC107065643 | serine/threonine-protein kinase NLK | 0.03703 | 123.24792 | TRUE | FALSE |
| LOC107065810 | furin-like protease 2 | 0.08047 | 123.34662 | TRUE | FALSE |
| LOC107068179 | uncharacterized protein LOC107068179 | 0.06976 | 123.37663 | TRUE | FALSE |
| LOC107065499 | transcriptional activator protein Pur-beta-B | 0.07444 | 123.39954 | TRUE | FALSE |
| LOC107068829 | transmembrane protein 132E | 0.09813 | 123.5046 | TRUE | FALSE |
| LOC107069338 | calpain-C | 0.16557 | 123.60785 | TRUE | FALSE |
| LOC107063925 | uncharacterized protein LOC107063925 | 0.10323 | 123.72795 | TRUE | FALSE |
| LOC107069302 | 26S proteasome non-ATPase regulatory sub | 0.11113 | 123.73742 | TRUE | FALSE |
| LOC107065595 | proto-oncogene tyrosine-protein kinase rec | 0.07031 | 123.82937 | TRUE | FALSE |
| LOC107071987 | gamma-tubulin complex component 4 | 0.03341 | 123.91515 | TRUE | FALSE |
| LOC107065770 | polyadenylate-binding protein 2 | 0.03987 | 123.94316 | TRUE | FALSE |
| LOC107071307 | myeloid leukemia factor | 0.05124 | 123.9717 | TRUE | FALSE |
| LOC107069739 | carbohydrate sulfotransferase 11-like | 0.05747 | 123.99552 | TRUE | FALSE |
| LOC107066607 | ADP-ribosylation factor-like protein 13B | 0.06503 | 124.05394 | TRUE | FALSE |
| LOC107064827 | cuticle protein 16.5, isoform A | 0.03453 | 124.05676 | TRUE | FALSE |
| LOC107069183 | uncharacterized protein LOC107069183 | 0.06284 | 124.06596 | TRUE | FALSE |
| LOC107070575 | protein FAM50 homolog | 0.03164 | 124.11664 | TRUE | FALSE |
| LOC107069158 | serine/threonine-protein kinase tricorner | 0.05099 | 124.13921 | TRUE | FALSE |
| LOC107064269 | TGF-beta-activated kinase 1 and MAP3K7-bi | 0.06155 | 124.14843 | TRUE | FALSE |
| LOC107073027 | uncharacterized protein LOC107073027 | 0.04125 | 124.16156 | TRUE | FALSE |
| LOC107074512 | akirin-2 | 0.05159 | 124.25591 | TRUE | FALSE |
| LOC107067580 | ABC transporter G family member 14 | 0.10723 | 124.344 | TRUE | FALSE |
| LOC107068764 | dual specificity protein kinase CLK2 | 0.10703 | 124.35072 | TRUE | FALSE |
| LOC107072427 | H/ACA ribonucleoprotein complex subunit 1 | 0.16507 | 124.38305 | TRUE | FALSE |
| LOC107074286 | histone-binding protein N1/N2-like | 0.1479 | 124.41611 | TRUE | FALSE |
| LOC107066792 | myotubularin-related protein 3 | 0.07632 | 124.56957 | TRUE | FALSE |
| LOC107074300 | four and a half LIM domains protein 3 | 0.0932 | 124.5872 | TRUE | TRUE |
| LOC107071131 | uncharacterized protein LOC107071131 | 0.16457 | 124.68827 | TRUE | FALSE |
| LOC107063909 | ubiquitin-conjugating enzyme E2 L3 | 0.12147 | 124.73428 | TRUE | FALSE |
| LOC107073007 | protein singed | 0.1106 | 124.7391 | TRUE | FALSE |
| LOC107072653 | single-stranded DNA-binding protein 3 | 0.1374 | 124.89794 | TRUE | FALSE |
| LOC107066965 | late histone H1-like | 0.05593 | 124.90144 | TRUE | FALSE |
| LOC107071512 | dentin sialophosphoprotein-like | 0.13383 | 124.92541 | TRUE | FALSE |
| LOC107067777 | p53 and DNA damage-regulated protein 1 | 0.06711 | 124.93415 | TRUE | FALSE |
| LOC107069358 | uncharacterized protein LOC107069358 | 0.18163 | 124.94643 | TRUE | TRUE |
| LOC107067537 | microtubule-associated protein futsch-like | 0.02839 | 124.97457 | TRUE | FALSE |
| LOC107070556 | U4/U6 small nuclear ribonucleoprotein Prp4 | 0.10105 | 125.00908 | TRUE | FALSE |
| LOC107071110 | uncharacterized protein LOC107071110 | 0.0406 | 125.01345 | TRUE | FALSE |
| LOC107068797 | L-ascorbate oxidase-like | 0.05395 | 125.06509 | TRUE | FALSE |
| LOC107067973 | uncharacterized protein LOC107067973 | 0.03855 | 125.12443 | TRUE | FALSE |
| LOC107065444 | serine-rich adhesin for platelets | 0.05861 | 125.12963 | TRUE | FALSE |
| LOC107074329 | uncharacterized protein LOC107074329 | 0.04678 | 125.15003 | TRUE | FALSE |
| LOC107069762 | dipeptidyl peptidase 9 | 0.04309 | 125.16245 | TRUE | FALSE |
| LOC107071561 | uncharacterized protein LOC107071561 | 0.05554 | 125.19719 | TRUE | FALSE |
| LOC107065461 | insulin-degrading enzyme | 0.03997 | 125.24424 | TRUE | FALSE |
| LOC107066558 | pumilio homolog 2 | 0.10428 | 125.24429 | TRUE | FALSE |

| | | | | | |
|---|---|---|---|---|---|
| LOC107074347 | NECAP-like protein CG9132 | 0.03049 | 125.28616 | TRUE | FALSE |
| LOC107065118 | peroxidasin-like | 0.02829 | 125.29631 | TRUE | FALSE |
| LOC107070245 | translational activator of cytochrome c oxid | 0.11473 | 125.32157 | TRUE | FALSE |
| LOC107074684 | male-enhanced antigen 1 | 0.1426 | 125.36948 | TRUE | FALSE |
| LOC107064675 | scavenger receptor class B member 1 | 0.04501 | 125.37793 | TRUE | FALSE |
| LOC107065374 | potassium voltage-gated channel subfamily | 0.0451 | 125.48899 | TRUE | FALSE |
| LOC107069217 | leucine-rich repeat neuronal protein 1-like | 0.04524 | 125.50953 | TRUE | FALSE |
| LOC107069066 | larval cuticle protein A1A-like | 0.05136 | 125.53851 | TRUE | FALSE |
| LOC107068375 | uncharacterized protein LOC107068375 | 0.0734 | 125.59749 | TRUE | FALSE |
| LOC107074575 | transcriptional enhancer factor TEF-1 | 0.11007 | 125.60194 | TRUE | FALSE |
| LOC107071881 | gastrula zinc finger protein xFG20-1-like | 0.10923 | 125.61992 | TRUE | FALSE |
| LOC107064533 | cilia- and flagella-associated protein 52 | 0.07982 | 125.71915 | TRUE | FALSE |
| LOC107073958 | histone deacetylase Rpd3 | 0.04478 | 125.86228 | TRUE | FALSE |
| LOC107069328 | cyclin-dependent kinase 4-like | 0.03546 | 125.89825 | TRUE | FALSE |
| LOC107066489 | BRCA1-A complex subunit Abraxas-like | 0.02926 | 125.95096 | TRUE | FALSE |
| LOC107069019 | rab5 GDP/GTP exchange factor | 0.02531 | 126.05237 | TRUE | FALSE |
| LOC107065579 | proclotting enzyme | 0.02593 | 126.13307 | TRUE | FALSE |
| LOC107064320 | ras-related protein Rac1 | 0.04837 | 126.13492 | TRUE | FALSE |
| LOC107066104 | IgA FC receptor-like | 0.09832 | 126.25327 | TRUE | FALSE |
| LOC107074181 | leucine-rich repeat and calponin homology ( | 0.06034 | 126.30731 | TRUE | FALSE |
| LOC107064258 | histone H3.3 | 0.03655 | 126.3152 | TRUE | FALSE |
| LOC107074499 | TOM1-like protein 2 | 0.06188 | 126.32082 | TRUE | FALSE |
| LOC107067707 | homeotic protein Sex combs reduced | 0.0303 | 126.33634 | TRUE | FALSE |
| LOC107071651 | protein Malvolio | 0.03696 | 126.45126 | TRUE | FALSE |
| LOC107069848 | protein groucho | 0.05768 | 126.46424 | TRUE | FALSE |
| LOC107067252 | 39S ribosomal protein L54, mitochondrial | 0.10474 | 126.56303 | TRUE | FALSE |
| LOC107073132 | glutamine synthetase 2 cytoplasmic | NA | 126.57478 | TRUE | FALSE |
| LOC107067322 | uncharacterized protein LOC107067322 | 0.1706 | 126.5923 | TRUE | FALSE |
| LOC107067577 | TBC1 domain family member 4 | 0.05915 | 126.66775 | TRUE | FALSE |
| LOC107069349 | dopamine N-acetyltransferase-like | 0.04254 | 126.82665 | TRUE | FALSE |
| LOC107070546 | cofilin/actin-depolymerizing factor homolog | 0.10204 | 127.01982 | TRUE | FALSE |
| LOC107072490 | cytoplasmic tRNA 2-thiolation protein 1 | 0.04296 | 127.07069 | TRUE | FALSE |
| LOC107070587 | clusterin-associated protein 1 | 0.16913 | 127.07761 | TRUE | FALSE |
| LOC107069130 | lachesin-like | 0.05105 | 127.0828 | TRUE | FALSE |
| LOC107068838 | uncharacterized protein C7orf26 homolog | 0.03678 | 127.19018 | TRUE | FALSE |
| LOC107065341 | uncharacterized protein LOC107065341 | 0.12163 | 127.19755 | TRUE | FALSE |
| LOC107065438 | uncharacterized protein LOC107065438 | 0.03804 | 127.26128 | TRUE | FALSE |
| LOC107073649 | uncharacterized protein LOC107073649 | 0.04938 | 127.315 | TRUE | FALSE |
| LOC107067307 | tight junction protein ZO-2 | 0.11523 | 127.3174 | TRUE | FALSE |
| LOC107068115 | uncharacterized protein LOC107068115 | 0.05527 | 127.71038 | TRUE | FALSE |
| LOC107067059 | heat shock 70 kDa protein cognate 4 | 0.02919 | 127.74667 | TRUE | FALSE |
| LOC107064339 | zinc finger protein 346 | 0.05278 | 127.76274 | TRUE | FALSE |
| LOC107070407 | putative fatty acyl-CoA reductase CG5065 | 0.09024 | 127.76594 | TRUE | TRUE |
| LOC107066848 | LYR motif-containing protein 9-like | 0.04968 | 127.8263 | TRUE | FALSE |
| LOC107067176 | heterogeneous nuclear ribonucleoprotein H | 0.04306 | 127.87255 | TRUE | FALSE |
| LOC107069123 | apoptotic chromatin condensation inducer i | 0.07376 | 127.92135 | TRUE | FALSE |
| LOC107074607 | uncharacterized protein LOC107074607 | 0.05963 | 127.98139 | TRUE | FALSE |
| LOC107069890 | uncharacterized protein LOC107069890 | 0.11837 | 128.151 | TRUE | FALSE |
| LOC107067122 | uncharacterized protein LOC107067122 | 0.11347 | 128.21108 | TRUE | TRUE |
| LOC107070379 | N-acetylglucosaminyl-phosphatidylinositol d | 0.04612 | 128.21259 | TRUE | FALSE |

| | | | | | |
|---|---|---|---|---|---|
| LOC107069136 | uncharacterized protein LOC107069136 | NA | 128.2175 | TRUE | FALSE |
| LOC107067093 | uncharacterized protein LOC107067093 | 0.03881 | 128.22184 | TRUE | FALSE |
| LOC107073446 | fatty acyl-CoA reductase 1-like | 0.08018 | 128.30372 | TRUE | TRUE |
| LOC107067891 | tyrosine aminotransferase | 0.04752 | 128.36821 | TRUE | FALSE |
| LOC107068633 | F-box/LRR-repeat protein 2 | 0.1703 | 128.43007 | TRUE | FALSE |
| LOC107066445 | uncharacterized protein LOC107066445 | 0.03968 | 128.4568 | TRUE | FALSE |
| LOC107073423 | uncharacterized protein LOC107073423 | NA | 128.52734 | TRUE | FALSE |
| LOC107066980 | cell cycle control protein 50A | 0.06246 | 128.62257 | TRUE | FALSE |
| LOC107069169 | uncharacterized protein LOC107069169 | 0.0288 | 128.7077 | TRUE | FALSE |
| LOC107071265 | uncharacterized protein LOC107071265 | 0.08828 | 128.71501 | TRUE | FALSE |
| LOC107071369 | piggyBac transposable element-derived pro | 0.05424 | 128.7514 | TRUE | FALSE |
| LOC107070680 | uncharacterized protein LOC107070680 | 0.14237 | 128.81941 | TRUE | TRUE |
| LOC107071539 | probable serine/threonine-protein kinase ya | 0.08554 | 128.84845 | TRUE | FALSE |
| LOC107069133 | interleukin-1 receptor accessory protein-like | 0.10913 | 128.92926 | TRUE | FALSE |
| LOC107068736 | uncharacterized protein LOC107068736 | 0.10293 | 128.94098 | TRUE | FALSE |
| LOC107069630 | cleavage and polyadenylation specificity fac | 0.05724 | 128.94987 | TRUE | FALSE |
| LOC107064488 | kelch-like protein 5 | 0.05876 | 128.98998 | TRUE | FALSE |
| LOC107072950 | uncharacterized protein LOC107072950 | 0.04139 | 129.01488 | TRUE | FALSE |
| LOC107071124 | homeodomain-interacting protein kinase 2 | 0.04509 | 129.07655 | TRUE | FALSE |
| LOC107063924 | uncharacterized protein LOC107063924 | 0.06051 | 129.09533 | TRUE | FALSE |
| LOC107066824 | uncharacterized protein LOC107066824 | 0.04067 | 129.11356 | TRUE | FALSE |
| LOC107073382 | uncharacterized protein LOC107073382 | 0.15183 | 129.12161 | TRUE | FALSE |
| LOC107073783 | uncharacterized protein T26G10.4-like | 0.03527 | 129.14456 | TRUE | FALSE |
| LOC107070667 | SPARC-related modular calcium-binding pro | 0.16337 | 129.17309 | TRUE | FALSE |
| LOC107069686 | U7 snRNA-associated Sm-like protein LSm11 | 0.03597 | 129.17831 | TRUE | FALSE |
| LOC107072917 | nucleosome assembly protein 1-like 1-B | 0.04044 | 129.24144 | TRUE | FALSE |
| LOC107066509 | brahma-associated protein of 60 kDa | 0.11623 | 129.26158 | TRUE | FALSE |
| LOC107073482 | protein toll-like | 0.06221 | 129.32317 | TRUE | FALSE |
| LOC107063831 | protein aurora borealis | 0.06374 | 129.36597 | TRUE | FALSE |
| LOC107074234 | omega-conotoxin-like protein 1 | 0.0888 | 129.41317 | TRUE | FALSE |
| LOC107068001 | TPPP family protein CG45057-like | 0.06638 | 129.44782 | TRUE | FALSE |
| LOC107064796 | 4-coumarate--CoA ligase 1-like | 0.03979 | 129.47605 | TRUE | FALSE |
| LOC107071857 | uncharacterized protein LOC107071857 | 0.06489 | 129.51153 | TRUE | FALSE |
| LOC107071325 | replication factor C subunit 4 | 0.04785 | 129.57144 | TRUE | FALSE |
| LOC107071945 | cysteine--tRNA ligase, cytoplasmic | 0.05369 | 129.58043 | TRUE | FALSE |
| LOC107066638 | pre-mRNA 3'-end-processing factor FIP1 | 0.1374 | 129.66576 | TRUE | FALSE |
| LOC107064334 | uncharacterized protein LOC107064334 | 0.03235 | 129.78968 | TRUE | FALSE |
| LOC107066456 | UPF0184 protein C9orf16 homolog | 0.04019 | 129.80481 | TRUE | FALSE |
| LOC107071066 | elongation of very long chain fatty acids pro | 0.07312 | 129.86211 | TRUE | TRUE |
| LOC107067099 | uncharacterized protein LOC107067099 | 0.09273 | 129.86704 | TRUE | FALSE |
| LOC107073380 | mediator of RNA polymerase II transcription | 0.14247 | 129.87015 | TRUE | FALSE |
| LOC107071860 | uncharacterized protein LOC107071860 | 0.0313 | 129.88024 | TRUE | FALSE |
| LOC107073290 | protein HEXIM1-like | 0.0585 | 129.97869 | TRUE | FALSE |
| LOC107066284 | protein vein | 0.08315 | 130.02652 | TRUE | FALSE |
| LOC107070843 | protein APCDD1-like | 0.12743 | 130.15176 | TRUE | FALSE |
| LOC107072195 | DNA-binding protein Ets97D | 0.05786 | 130.17537 | TRUE | FALSE |
| LOC107072428 | tetra-peptide repeat homeobox protein 1-li | 0.11373 | 130.18069 | TRUE | FALSE |
| LOC107065791 | cell differentiation protein RCD1 homolog | 0.03371 | 130.18596 | TRUE | FALSE |
| LOC107063694 | acetylcholinesterase-like | 0.07189 | 130.21057 | TRUE | FALSE |
| LOC107067314 | proteasome activator complex subunit 4B-li | 0.04277 | 130.28648 | TRUE | FALSE |

| | | | | | |
|---|---|---|---|---|---|
| LOC107069305 | homeobox protein Nkx-6.1-like | 0.04538 | 130.30223 | TRUE | FALSE |
| LOC107067706 | armadillo repeat-containing protein 8-like | 0.14867 | 130.36991 | TRUE | FALSE |
| LOC107069464 | uncharacterized protein LOC107069464 | 0.1388 | 130.41679 | TRUE | FALSE |
| LOC107064198 | arylsulfatase B-like | 0.04075 | 130.43362 | TRUE | FALSE |
| LOC107066875 | protein hairy | 0.06085 | 130.51575 | TRUE | FALSE |
| LOC107068906 | T-complex protein 1 subunit beta | 0.13647 | 130.52293 | TRUE | FALSE |
| LOC107070181 | ankyrin repeat domain-containing protein 5 | 0.1123 | 130.56134 | TRUE | FALSE |
| LOC107068422 | uncharacterized protein LOC107068422 | 0.115 | 130.72579 | TRUE | FALSE |
| LOC107074169 | uncharacterized protein LOC107074169 | 0.05856 | 130.77261 | TRUE | FALSE |
| LOC107072286 | uncharacterized protein LOC107072286 | 0.09226 | 130.90823 | TRUE | FALSE |
| LOC107070391 | glutamate--cysteine ligase regulatory subun | 0.03402 | 131.0206 | TRUE | FALSE |
| LOC107068007 | probable calcium-binding protein CML22 | 0.06467 | 131.13514 | TRUE | FALSE |
| LOC107067324 | uncharacterized protein LOC107067324 | 0.07734 | 131.14572 | TRUE | FALSE |
| LOC107066744 | CREB-regulated transcription coactivator 1-l | 0.0983 | 131.14641 | TRUE | FALSE |
| LOC107070578 | uncharacterized protein LOC107070578 | 0.03165 | 131.24272 | TRUE | FALSE |
| LOC107073084 | uncharacterized protein LOC107073084 | 0.034 | 131.28722 | TRUE | FALSE |
| LOC107065010 | mitochondrial dicarboxylate carrier-like | NA | 131.32287 | TRUE | FALSE |
| LOC107064607 | serine/arginine-rich splicing factor 1B | 0.07096 | 131.39133 | TRUE | FALSE |
| LOC107069643 | dynein regulatory complex subunit 7 | 0.15193 | 131.44505 | TRUE | FALSE |
| LOC107066687 | trinucleotide repeat-containing gene 6C pro | 0.16607 | 131.52463 | TRUE | FALSE |
| LOC107074198 | tetratricopeptide repeat protein 30A | 0.0719 | 131.71395 | TRUE | FALSE |
| LOC107065557 | transmembrane protein adipocyte-associate | 0.04463 | 131.77463 | TRUE | FALSE |
| LOC107067848 | collagen alpha-1(IV) chain | 0.16 | 132.24536 | TRUE | FALSE |
| LOC107070828 | small glutamine-rich tetratricopeptide repea | 0.06219 | 132.26675 | TRUE | FALSE |
| LOC107066568 | receptor-interacting serine/threonine-prote | 0.05537 | 132.32414 | TRUE | FALSE |
| LOC107070759 | E3 SUMO-protein ligase NSE2-like | 0.04558 | 132.33937 | TRUE | FALSE |
| LOC107068170 | putative gamma-glutamylcyclotransferase C | 0.04147 | 132.38591 | TRUE | FALSE |
| LOC107072089 | transcriptional adapter 2B | 0.02901 | 132.40661 | TRUE | FALSE |
| LOC107072645 | transcription initiation factor TFIID subunit 1 | 0.11307 | 132.44283 | TRUE | FALSE |
| LOC107064466 | uncharacterized protein LOC107064466 | 0.03452 | 132.44522 | TRUE | FALSE |
| LOC107070741 | 39S ribosomal protein L50, mitochondrial | 0.04306 | 132.50891 | TRUE | FALSE |
| LOC107071112 | uncharacterized protein LOC107071112 | 0.15197 | 132.53648 | TRUE | FALSE |
| LOC107068470 | succinyl-CoA ligase [GDP-forming | 0.1751 | 132.54656 | TRUE | FALSE |
| LOC107067128 | sesquipedalian-1-like | 0.0719 | 132.68726 | TRUE | FALSE |
| LOC107066901 | uncharacterized protein LOC107066901 | 0.06329 | 132.69169 | TRUE | FALSE |
| LOC107072188 | NADH dehydrogenase [ubiquinone | 0.1322 | 132.70548 | TRUE | FALSE |
| LOC107070653 | ADP-ribosylation factor-like protein 16 | 0.08326 | 132.77098 | TRUE | FALSE |
| LOC107067821 | mushroom body large-type Kenyon cell-spe | 0.1581 | 132.84453 | TRUE | FALSE |
| LOC107065659 | chondroitin proteoglycan 2-like | 0.04712 | 132.91582 | TRUE | FALSE |
| LOC107071795 | acyl-CoA Delta(11) desaturase-like | 0.1898 | 132.91931 | TRUE | TRUE |
| LOC107071701 | leucine-rich repeat-containing protein 70-lik | 0.1132 | 132.97068 | TRUE | FALSE |
| LOC107067670 | selenoprotein K-like | 0.0485 | 133.05359 | TRUE | FALSE |
| LOC107068977 | WD repeat-containing protein 3 | 0.04903 | 133.23428 | TRUE | FALSE |
| LOC107064940 | uncharacterized protein LOC107064940 | NA | 133.38341 | TRUE | TRUE |
| LOC107066598 | uncharacterized protein LOC107066598 | 0.08484 | 133.50672 | TRUE | FALSE |
| LOC107065863 | SET and MYND domain-containing protein 4 | 0.09871 | 133.51383 | TRUE | FALSE |
| LOC107068056 | probable cation-transporting ATPase 13A3 | 0.08099 | 133.55446 | TRUE | FALSE |
| LOC107068287 | RNMT-activating mini protein | 0.08306 | 133.72381 | TRUE | FALSE |
| LOC107064649 | metallophosphoesterase domain-containing | 0.06689 | 133.73551 | TRUE | FALSE |
| LOC107072970 | homeobox protein onecut-like | 0.09993 | 133.83612 | TRUE | FALSE |

| | | | | | |
|---|---|---|---|---|---|
| LOC107063746 | CCA tRNA nucleotidyltransferase 1, mitocho | 0.05437 | 133.84992 | TRUE | FALSE |
| LOC107068616 | ejaculatory bulb-specific protein 3 | 0.07279 | 133.95974 | TRUE | TRUE |
| LOC107074348 | multidrug resistance protein homolog 49-lik | 0.07472 | 133.98418 | TRUE | FALSE |
| LOC107068477 | dynein light chain 2, cytoplasmic | 0.08392 | 134.00877 | TRUE | FALSE |
| LOC107072207 | GATA zinc finger domain-containing protein | 0.04963 | 134.24176 | TRUE | FALSE |
| LOC107068740 | E3 ubiquitin-protein ligase RNF19A-like | 0.13937 | 134.24264 | TRUE | FALSE |
| LOC107070625 | zinc finger protein 608-like | 0.1344 | 134.25317 | TRUE | FALSE |
| LOC107070771 | ribonucleoside-diphosphate reductase large | 0.03983 | 134.25959 | TRUE | FALSE |
| LOC107068119 | mitochondrial ubiquitin ligase activator of n | 0.05343 | 134.28472 | TRUE | FALSE |
| LOC107066443 | UPF0565 protein C2orf69 homolog | 0.0393 | 134.31947 | TRUE | FALSE |
| LOC107064917 | peptidylglycine alpha-hydroxylating monoox | 0.08365 | 134.47553 | TRUE | FALSE |
| LOC107074569 | uncharacterized protein LOC107074569 | 0.13337 | 134.60578 | TRUE | FALSE |
| LOC107067569 | 97 kDa heat shock protein | 0.12987 | 134.62171 | TRUE | FALSE |
| LOC107071493 | uncharacterized protein LOC107071493 | 0.14703 | 134.66758 | TRUE | TRUE |
| LOC107072470 | FUN14 domain-containing protein 1 | 0.03281 | 134.69064 | TRUE | FALSE |
| LOC107064157 | stAR-related lipid transfer protein 7, mitoch | 0.08272 | 134.72403 | TRUE | FALSE |
| LOC107070655 | peptidyl-prolyl cis-trans isomerase-like | 0.06541 | 134.79244 | TRUE | FALSE |
| LOC107072910 | uncharacterized protein LOC107072910 | 0.09408 | 134.91705 | TRUE | FALSE |
| LOC107074077 | uncharacterized protein LOC107074077 | 0.058 | 134.93066 | TRUE | FALSE |
| LOC107069052 | transcription factor BTF3 homolog 4 | NA | 134.93102 | TRUE | FALSE |
| LOC107074126 | uncharacterized protein LOC107074126 | 0.08203 | 134.98573 | TRUE | FALSE |
| LOC107074230 | probable cytochrome P450 6a13 | 0.09166 | 135.06106 | TRUE | FALSE |
| LOC107074088 | casein kinase I isoform alpha-like | 0.04221 | 135.3619 | TRUE | FALSE |
| LOC107068704 | endocuticle structural glycoprotein SgAbd-1 | 0.06792 | 135.42295 | TRUE | FALSE |
| LOC107071664 | mitochondrial distribution and morphology | 0.18563 | 135.56088 | TRUE | FALSE |
| LOC107069242 | protein white | 0.1286 | 135.56212 | TRUE | FALSE |
| LOC107065115 | circadian clock-controlled protein-like | 0.03105 | 135.68247 | TRUE | FALSE |
| LOC107070870 | PCNA-associated factor-like | 0.05369 | 135.71228 | TRUE | FALSE |
| LOC107065108 | metallophosphoesterase 1 | 0.12023 | 135.76071 | TRUE | FALSE |
| LOC107065069 | knirps-related protein-like | 0.08449 | 135.86623 | TRUE | FALSE |
| LOC107073462 | uncharacterized protein LOC107073462 | 0.06709 | 135.88925 | TRUE | FALSE |
| LOC107064383 | uncharacterized protein LOC107064383 | 0.06043 | 136.10084 | TRUE | FALSE |
| LOC107069089 | small ubiquitin-related modifier-like | 0.1328 | 136.1529 | TRUE | FALSE |
| LOC107068837 | uncharacterized protein LOC107068837 | 0.04856 | 136.18938 | TRUE | FALSE |
| LOC107070088 | uncharacterized protein LOC107070088 | 0.10319 | 136.21779 | TRUE | FALSE |
| LOC107069573 | MATH and LRR domain-containing protein P | 0.02919 | 136.29477 | TRUE | FALSE |
| LOC107068909 | M-phase phosphoprotein 6 | 0.11323 | 136.4906 | TRUE | FALSE |
| LOC107064396 | methylenetetrahydrofolate reductase | 0.04351 | 136.6203 | TRUE | FALSE |
| LOC107065051 | uncharacterized protein LOC107065051 | 0.15137 | 136.7072 | TRUE | FALSE |
| LOC107069891 | poly(A) RNA polymerase, mitochondrial-like | 0.0345 | 136.78544 | TRUE | FALSE |
| LOC107070501 | uncharacterized protein LOC107070501 | 0.0601 | 136.90648 | TRUE | FALSE |
| LOC107067003 | B-cell CLL/lymphoma 7 protein family meml | 0.04784 | 137.06717 | TRUE | FALSE |
| LOC107065283 | protein abnormal spindle | 0.08681 | 137.08427 | TRUE | FALSE |
| LOC107068377 | zinc finger protein 62-like | 0.05908 | 137.13257 | TRUE | FALSE |
| LOC107067713 | uncharacterized protein LOC107067713 | 0.10494 | 137.22825 | TRUE | FALSE |
| LOC107066427 | nucleolar protein 4-like | 0.15647 | 137.5165 | TRUE | FALSE |
| LOC107072720 | DNA-binding protein Ewg | 0.15403 | 137.6065 | TRUE | FALSE |
| LOC107071144 | uncharacterized protein LOC107071144 | 0.04673 | 137.62834 | TRUE | FALSE |
| LOC107063666 | cAMP-specific 3',5'-cyclic phosphodiesterase | 0.14827 | 137.6864 | TRUE | FALSE |
| LOC107073364 | uncharacterized protein LOC107073364 | 0.08985 | 137.78703 | TRUE | FALSE |

| | | | | | |
|---|---|---|---|---|---|
| LOC107067474 | elongation factor Ts, mitochondrial | 0.1593 | 137.80381 | TRUE | FALSE |
| LOC107065317 | uncharacterized protein LOC107065317 | 0.07194 | 137.82427 | TRUE | FALSE |
| LOC107073770 | uncharacterized protein LOC107073770 | 0.04067 | 137.86832 | TRUE | FALSE |
| LOC107068109 | uncharacterized protein LOC107068109 | 0.05223 | 137.87247 | TRUE | FALSE |
| LOC107065708 | uncharacterized protein LOC107065708 | 0.03898 | 137.89042 | TRUE | FALSE |
| LOC107069759 | protein SGT1 homolog | 0.04986 | 137.91181 | TRUE | FALSE |
| LOC107074296 | loss of heterozygosity 12 chromosomal regi | 0.09292 | 137.96025 | TRUE | FALSE |
| LOC107071125 | inactive peptidyl-prolyl cis-trans isomerase I | 0.12887 | 138.15519 | TRUE | FALSE |
| LOC107064148 | uncharacterized family 31 glucosidase KIAA: | 0.09579 | 138.17313 | TRUE | FALSE |
| LOC107065578 | toll-like receptor 8 | 0.09488 | 138.19634 | TRUE | TRUE |
| LOC107065545 | uncharacterized protein LOC107065545 | 0.07228 | 138.33458 | TRUE | FALSE |
| LOC107074095 | uncharacterized protein LOC107074095 | 0.11737 | 138.42067 | TRUE | FALSE |
| LOC107067560 | biogenesis of lysosome-related organelles c | 0.12517 | 138.42601 | TRUE | FALSE |
| LOC107074690 | kinesin-like protein KIF9 | 0.1204 | 138.46704 | TRUE | FALSE |
| LOC107064687 | 40S ribosomal protein S24 | 0.04967 | 138.46712 | TRUE | FALSE |
| LOC107066589 | histone-lysine N-methyltransferase setd3 | 0.06115 | 138.4868 | TRUE | FALSE |
| LOC107074036 | uncharacterized protein LOC107074036 | 0.06742 | 138.54006 | TRUE | FALSE |
| LOC107068735 | unconventional myosin-Va | 0.09243 | 138.7669 | TRUE | FALSE |
| LOC107071721 | out at first protein | 0.13563 | 138.81826 | TRUE | FALSE |
| LOC107065093 | nucleolysin TIAR | 0.12823 | 138.84794 | TRUE | FALSE |
| LOC107068540 | uncharacterized protein LOC107068540 | 0.08108 | 138.87676 | TRUE | FALSE |
| LOC107073435 | uncharacterized protein LOC107073435 | NA | 139.16523 | TRUE | FALSE |
| LOC107067330 | prohibitin-2 | 0.08866 | 139.25071 | TRUE | FALSE |
| LOC107065767 | Golgi apparatus protein 1 | 0.15357 | 139.3313 | TRUE | FALSE |
| LOC107065865 | etoposide-induced protein 2.4 homolog | 0.04705 | 139.35933 | TRUE | FALSE |
| LOC107068251 | uncharacterized protein LOC107068251 | 0.17773 | 139.36272 | TRUE | FALSE |
| LOC107070320 | protein Skeletor, isoforms B/C | 0.15317 | 139.38689 | TRUE | FALSE |
| LOC107073630 | retinoblastoma-binding protein 5 homolog | 0.12017 | 139.50432 | TRUE | FALSE |
| LOC107074439 | exostosin-1 | 0.05119 | 139.51123 | TRUE | FALSE |
| LOC107069149 | metabotropic glutamate receptor 6-like | 0.04938 | 139.51247 | TRUE | FALSE |
| LOC107072481 | 26S protease regulatory subunit 4 | 0.09979 | 139.51681 | TRUE | FALSE |
| LOC107066914 | poly(rC)-binding protein 4-like | 0.18687 | 139.846 | TRUE | TRUE |
| LOC107071337 | ubiquitin carboxyl-terminal hydrolase 34 | 0.148 | 139.89055 | TRUE | FALSE |
| LOC107072648 | zinc finger protein 598 | 0.06064 | 140.03694 | TRUE | FALSE |
| LOC107070729 | neutral ceramidase | 0.16353 | 140.05438 | TRUE | FALSE |
| LOC107070604 | CRAL-TRIO domain-containing protein DDB_ | 0.1581 | 140.16639 | TRUE | FALSE |
| LOC107070996 | mimitin, mitochondrial | 0.04002 | 140.29674 | TRUE | FALSE |
| LOC107073424 | aprataxin and PNK-like factor | 0.05647 | 140.29931 | TRUE | FALSE |
| LOC107067016 | splicing factor 1-like | 0.09007 | 140.36633 | TRUE | FALSE |
| LOC107069782 | putative mediator of RNA polymerase II trar | 0.16243 | 140.37125 | TRUE | TRUE |
| LOC107066130 | gonadotropin-releasing hormone receptor | 0.1272 | 140.4322 | TRUE | FALSE |
| LOC107073571 | disks large-associated protein 4-like | 0.0392 | 140.47044 | TRUE | FALSE |
| LOC107066205 | WD repeat and FYVE domain-containing prc | 0.15703 | 140.54669 | TRUE | FALSE |
| LOC107064481 | uncharacterized protein LOC107064481 | 0.1219 | 140.60313 | TRUE | FALSE |
| LOC107072986 | sodium-independent sulfate anion transpor | 0.15147 | 140.71527 | TRUE | FALSE |
| LOC107073944 | integrator complex subunit 7 | 0.09184 | 140.80325 | TRUE | FALSE |
| LOC107063843 | uncharacterized protein LOC107063843 | 0.05185 | 140.86184 | TRUE | FALSE |
| LOC107066654 | N-alpha-acetyltransferase 30-like | 0.07025 | 141.0842 | TRUE | FALSE |
| LOC107066860 | ras association domain-containing protein 8 | 0.08558 | 141.10586 | TRUE | FALSE |
| LOC107071663 | uncharacterized protein LOC107071663 | 0.12717 | 141.1077 | TRUE | FALSE |

| | | | | | |
|---|---|---|---|---|---|
| LOC107074471 | uncharacterized protein LOC107074471 | 0.04221 | 141.14758 | TRUE | FALSE |
| LOC107071370 | fatty acid synthase-like | 0.03443 | 141.29263 | TRUE | FALSE |
| LOC107068602 | serine palmitoyltransferase 2 | 0.08478 | 141.30204 | TRUE | FALSE |
| LOC107067447 | protein tumorous imaginal discs, mitochonc | 0.10027 | 141.41859 | TRUE | FALSE |
| LOC107070152 | tctex1 domain-containing protein 2-like | 0.0492 | 141.4521 | TRUE | FALSE |
| LOC107067659 | uncharacterized protein LOC107067659 | 0.1549 | 141.5569 | TRUE | FALSE |
| LOC107064065 | zinc finger protein 408-like | 0.1717 | 141.67684 | TRUE | FALSE |
| LOC107066094 | neurexin-2 | 0.17487 | 141.90676 | TRUE | FALSE |
| LOC107069881 | uncharacterized protein LOC107069881 | 0.09591 | 141.98604 | TRUE | FALSE |
| LOC107068953 | protein RMD5 homolog A | NA | 142.06802 | TRUE | FALSE |
| LOC107067181 | DNA N6-methyl adenine demethylase | 0.15673 | 142.10811 | TRUE | FALSE |
| LOC107070292 | uncharacterized protein LOC107070292 | 0.06454 | 142.13999 | TRUE | FALSE |
| LOC107074270 | uncharacterized protein LOC107074270 | 0.1914 | 142.21419 | TRUE | FALSE |
| LOC107065516 | proline-rich receptor-like protein kinase PEF | 0.05434 | 142.26828 | TRUE | FALSE |
| LOC107068244 | 4-coumarate--CoA ligase 1 | 0.1272 | 142.36212 | TRUE | FALSE |
| LOC107072364 | uncharacterized protein LOC107072364 | 0.07343 | 142.41248 | TRUE | FALSE |
| LOC107074640 | bifunctional 3'-phosphoadenosine 5'-phospl | 0.16307 | 142.56165 | TRUE | FALSE |
| LOC107071971 | mitochondrial sodium/hydrogen exchanger | 0.0495 | 142.7515 | TRUE | FALSE |
| LOC107071193 | leucine-rich repeat-containing protein 24-lik | 0.1608 | 142.79785 | TRUE | FALSE |
| LOC107065600 | uncharacterized protein LOC107065600 | 0.1219 | 142.81514 | TRUE | FALSE |
| LOC107068131 | uncharacterized protein LOC107068131 | 0.1039 | 142.88064 | TRUE | FALSE |
| LOC107064777 | uncharacterized protein LOC107064777 | 0.03889 | 142.93363 | TRUE | FALSE |
| LOC107069120 | brain mitochondrial carrier protein 1 | 0.13927 | 143.10503 | TRUE | FALSE |
| LOC107063836 | neurotrimin-like | 0.05468 | 143.33068 | TRUE | FALSE |
| LOC107064149 | putative uncharacterized protein DDB_G02{ | 0.04894 | 143.43635 | TRUE | FALSE |
| LOC107068130 | probable chitinase 2 | 0.17093 | 143.46022 | TRUE | FALSE |
| LOC107070394 | zinc finger protein jing homolog | 0.11617 | 143.46712 | TRUE | FALSE |
| LOC107065150 | uncharacterized protein LOC107065150 | 0.16183 | 143.57 | TRUE | FALSE |
| LOC107073161 | cyclin-dependent kinase 9 | 0.03316 | 143.58312 | TRUE | FALSE |
| LOC107064479 | YTH domain-containing family protein 3-like | 0.1269 | 143.63241 | TRUE | FALSE |
| LOC107072332 | coatomer subunit delta | 0.07362 | 143.9894 | TRUE | FALSE |
| LOC107071303 | zinc finger protein Elbow-like | 0.06958 | 144.10539 | TRUE | FALSE |
| LOC107070925 | myosin light chain kinase, smooth muscle-lil | 0.1724 | 144.163 | TRUE | FALSE |
| LOC107072729 | F-box/WD repeat-containing protein 9-like | 0.11917 | 144.22236 | TRUE | FALSE |
| LOC107070294 | nucleotide exchange factor SIL1 | 0.03743 | 144.2482 | TRUE | FALSE |
| LOC107070658 | uncharacterized protein LOC107070658 | 0.06969 | 144.39351 | TRUE | FALSE |
| LOC107066073 | retinaldehyde-binding protein 1-like | 0.08464 | 144.42765 | TRUE | FALSE |
| LOC107066048 | regulator of chromosome condensation-like | NA | 144.44444 | TRUE | FALSE |
| LOC107070192 | 45 kDa calcium-binding protein | 0.05139 | 144.44704 | TRUE | FALSE |
| LOC107063672 | LOW QUALITY PROTEIN: uncharacterized pr | 0.09083 | 144.47017 | TRUE | FALSE |
| LOC107067287 | sperm flagellar protein 1-like | 0.10372 | 144.47198 | TRUE | FALSE |
| LOC107071295 | UTP--glucose-1-phosphate uridylyltransfera | 0.13477 | 144.64736 | TRUE | FALSE |
| LOC107066179 | iron-sulfur cluster assembly scaffold protein | NA | 144.68195 | TRUE | FALSE |
| LOC107074687 | uncharacterized protein LOC107074687 | 0.1001 | 144.755 | TRUE | FALSE |
| LOC107066753 | uncharacterized protein LOC107066753 | 0.05666 | 144.76344 | TRUE | FALSE |
| LOC107068685 | vacuolar protein sorting-associated protein | 0.06586 | 144.78269 | TRUE | FALSE |
| LOC107064837 | mediator of RNA polymerase II transcriptior | 0.14617 | 144.93922 | TRUE | FALSE |
| LOC107073905 | UPF0704 protein C6orf165 | NA | 144.96194 | TRUE | FALSE |
| LOC107069061 | uncharacterized protein DDB_G0271670 | 0.05844 | 145.167 | TRUE | FALSE |
| LOC107066649 | uncharacterized protein LOC107066649 | 0.07874 | 145.37246 | TRUE | FALSE |

| | | | | | |
|---|---|---|---|---|---|
| LOC107063633 | chromatin modification-related protein EAF | NA | 145.45058 | TRUE | TRUE |
| LOC107064036 | 28S ribosomal protein S31, mitochondrial | 0.04513 | 145.91897 | TRUE | FALSE |
| NA | NA | 0.06509 | 145.97639 | TRUE | FALSE |
| LOC107071108 | uncharacterized protein LOC107071108 | 0.1165 | 146.02104 | TRUE | FALSE |
| LOC107064949 | adult-specific cuticular protein ACP-22-like | 0.03403 | 146.15178 | TRUE | FALSE |
| LOC107066672 | mitochondrial import inner membrane trans | 0.12487 | 146.3443 | TRUE | FALSE |
| LOC107067732 | FAS-associated factor 1 | 0.13923 | 146.53068 | TRUE | FALSE |
| LOC107070470 | dedicator of cytokinesis protein 3 | NA | 146.59289 | TRUE | FALSE |
| LOC107064191 | tRNA-specific adenosine deaminase 1 | 0.06536 | 146.59865 | TRUE | FALSE |
| LOC107064902 | pseudouridylate synthase 7 homolog | 0.09553 | 146.60442 | TRUE | FALSE |
| LOC107070800 | uncharacterized protein LOC107070800 | 0.10129 | 146.90472 | TRUE | FALSE |
| LOC107070720 | apolipophorins | 0.16317 | 146.90957 | TRUE | FALSE |
| LOC107067589 | protein-tyrosine sulfotransferase | 0.05668 | 146.97493 | TRUE | FALSE |
| LOC107064241 | uncharacterized protein LOC107064241 | 0.17807 | 147.06143 | TRUE | TRUE |
| LOC107073991 | upstream activation factor subunit spp27 | 0.04681 | 147.08052 | TRUE | FALSE |
| LOC107071943 | protein PIH1D3 | 0.03325 | 147.11 | TRUE | FALSE |
| LOC107072691 | ubiquitin-conjugating enzyme E2 N | 0.17897 | 147.16811 | TRUE | FALSE |
| LOC107071821 | guanine nucleotide exchange factor MSS4 | 0.10454 | 147.2028 | TRUE | FALSE |
| LOC107065713 | uncharacterized protein LOC107065713 | 0.06269 | 147.32056 | TRUE | FALSE |
| LOC107073455 | probable multidrug resistance-associated pr | 0.1131 | 147.95499 | TRUE | FALSE |
| LOC107065934 | uncharacterized protein LOC107065934 | 0.03681 | 148.10434 | TRUE | FALSE |
| LOC107070119 | TNF receptor-associated factor 6-like | 0.06161 | 148.11572 | TRUE | FALSE |
| LOC107073291 | uncharacterized protein LOC107073291 | 0.14493 | 148.14739 | TRUE | FALSE |
| LOC107069542 | F-box-like/WD repeat-containing protein TB | 0.13087 | 148.15321 | TRUE | FALSE |
| LOC107067748 | ribonuclease kappa | 0.18367 | 148.28967 | TRUE | FALSE |
| LOC107070177 | diphosphoinositol polyphosphate phosphoh | 0.07618 | 148.35765 | TRUE | FALSE |
| LOC107067812 | probable low affinity copper uptake protein | 0.06253 | 148.37816 | TRUE | FALSE |
| LOC107068862 | cytochrome P450 306a1 | 0.12187 | 148.46726 | TRUE | FALSE |
| LOC107071140 | TGF-beta receptor type-1 | 0.07767 | 148.57848 | TRUE | FALSE |
| LOC107070595 | protein lifeguard 3-like | 0.17363 | 148.60212 | TRUE | FALSE |
| LOC107066799 | gustatory receptor for sugar taste 43a-like | 0.04918 | 148.71007 | TRUE | FALSE |
| LOC107067205 | proton-coupled amino acid transporter 4 | 0.08293 | 148.8469 | TRUE | FALSE |
| LOC107069924 | odorant receptor 13a-like | 0.03943 | 148.92129 | TRUE | FALSE |
| LOC107066465 | protein EMRE homolog, mitochondrial | 0.08163 | 148.98678 | TRUE | FALSE |
| LOC107073230 | uncharacterized protein LOC107073230 | 0.09783 | 149.0178 | TRUE | FALSE |
| LOC107074655 | ribosomal protein S6 kinase 2 beta | 0.09509 | 149.03519 | TRUE | FALSE |
| LOC107067223 | protein singed wings 2 | 0.18953 | 149.0584 | TRUE | FALSE |
| LOC107069190 | uncharacterized protein LOC107069190 | 0.11357 | 149.06014 | TRUE | FALSE |
| LOC107066108 | peroxidasin | 0.06399 | 149.08694 | TRUE | FALSE |
| LOC107074325 | reticulocalbin-2 | 0.19173 | 149.087 | TRUE | FALSE |
| LOC107066690 | protein abrupt | 0.12433 | 149.09977 | TRUE | FALSE |
| LOC107070460 | uncharacterized protein LOC107070460 | NA | 149.32997 | TRUE | FALSE |
| LOC107064577 | 5-oxoprolinase | 0.19133 | 149.34661 | TRUE | FALSE |
| LOC107065556 | potassium/sodium hyperpolarization-activa | 0.05807 | 149.70173 | TRUE | FALSE |
| LOC107066603 | uncharacterized protein LOC107066603 | 0.064 | 149.78875 | TRUE | FALSE |
| LOC107074352 | protein FAM76A | 0.10127 | 149.8286 | TRUE | FALSE |
| LOC107065362 | nitrate reductase [NAD(P)H | 0.11187 | 149.84099 | TRUE | FALSE |
| LOC107064924 | uncharacterized protein LOC107064924 | 0.09618 | 149.86726 | TRUE | FALSE |
| LOC107066963 | cytosolic carboxypeptidase 6 | 0.0743 | 150.20325 | TRUE | FALSE |
| LOC107068360 | spondin-1-like | 0.1811 | 150.24906 | TRUE | FALSE |

| | | | | | |
|---|---|---|---|---|---|
| LOC107068888 | odorant receptor 13a-like | 0.1158 | 150.34611 | TRUE | TRUE |
| LOC107068080 | RING finger protein 113A | 0.09008 | 150.3811 | TRUE | FALSE |
| LOC107065469 | uncharacterized protein C3orf18-like | 0.07521 | 150.45682 | TRUE | FALSE |
| LOC107065851 | biogenesis of lysosome-related organelles c | 0.08153 | 150.55362 | TRUE | FALSE |
| LOC107073625 | protein toll-like | 0.03789 | 150.70024 | TRUE | FALSE |
| LOC107070778 | WD repeat-containing protein 43-like | 0.06601 | 150.92947 | TRUE | FALSE |
| LOC107065641 | sterile alpha and TIR motif-containing prote | 0.13487 | 151.14754 | TRUE | FALSE |
| LOC107067582 | CUGBP Elav-like family member 4 | 0.1812 | 151.19312 | TRUE | FALSE |
| LOC107068508 | MAP kinase-interacting serine/threonine-pr | NA | 151.32565 | TRUE | FALSE |
| LOC107069610 | probable JmjC domain-containing histone d | NA | 151.37277 | TRUE | FALSE |
| LOC107069427 | lipase 3-like | 0.0497 | 151.46287 | TRUE | FALSE |
| LOC107069142 | uncharacterized protein LOC107069142 | 0.1772 | 151.58266 | TRUE | FALSE |
| LOC107067449 | beta-1,4-glucuronyltransferase 1-like | 0.08024 | 151.7975 | TRUE | FALSE |
| LOC107073825 | DNA mismatch repair protein MutL-like | 0.1827 | 151.81131 | TRUE | FALSE |
| LOC107066134 | exostosin-2 | 0.13757 | 152.52224 | TRUE | FALSE |
| LOC107071466 | transmembrane protein 120 homolog | 0.15677 | 152.56168 | TRUE | FALSE |
| LOC107070486 | ADP-ribosylation factor-like protein 1 | 0.10525 | 152.8533 | TRUE | FALSE |
| LOC107072539 | LIM/homeobox protein Awh-like | 0.08634 | 152.87485 | TRUE | FALSE |
| LOC107068214 | thioredoxin, mitochondrial | 0.155 | 152.91033 | TRUE | FALSE |
| LOC107072650 | glycerate kinase-like | 0.06676 | 152.91445 | TRUE | FALSE |
| LOC107065882 | RNA-binding protein Musashi homolog Rbp( | 0.14697 | 153.01215 | TRUE | FALSE |
| LOC107063640 | glutamate receptor ionotropic, kainate 2 | 0.14173 | 153.07506 | TRUE | FALSE |
| LOC107064082 | focal adhesion kinase 1 | 0.1427 | 153.17697 | TRUE | FALSE |
| LOC107066291 | uncharacterized protein LOC107066291 | 0.09167 | 153.21858 | TRUE | FALSE |
| LOC107066677 | GA-binding protein subunit beta-2 | 0.10072 | 153.25601 | TRUE | FALSE |
| LOC107070237 | sensory neuron membrane protein 1 | 0.04416 | 153.27896 | TRUE | TRUE |
| LOC107072477 | uncharacterized protein LOC107072477 | 0.14327 | 153.30638 | TRUE | FALSE |
| LOC107071450 | piggyBac transposable element-derived pro | NA | 153.43427 | TRUE | FALSE |
| LOC107071344 | COP9 signalosome complex subunit 7 | 0.09409 | 153.4826 | TRUE | FALSE |
| LOC107064937 | dynein heavy chain 7, axonemal-like | 0.18933 | 153.60882 | TRUE | FALSE |
| LOC107066828 | box C/D snoRNA protein 1 | 0.05328 | 153.69647 | TRUE | FALSE |
| LOC107067353 | uncharacterized protein LOC107067353 | 0.12803 | 153.79194 | TRUE | TRUE |
| LOC107065821 | transforming growth factor beta regulator 1 | 0.13017 | 153.86431 | TRUE | FALSE |
| LOC107072168 | cilia- and flagella-associated protein 70 | NA | 153.86641 | TRUE | FALSE |
| LOC107064325 | probable tyrosyl-DNA phosphodiesterase | 0.06955 | 153.94069 | TRUE | FALSE |
| LOC107066218 | zinc finger CCHC domain-containing protein | 0.06073 | 154.06292 | TRUE | FALSE |
| LOC107069026 | uncharacterized protein LOC107069026 | 0.09695 | 154.13401 | TRUE | FALSE |
| LOC107070747 | uncharacterized protein LOC107070747 | 0.07658 | 154.18285 | TRUE | FALSE |
| LOC107068300 | ATP-dependent RNA helicase WM6 | 0.10394 | 154.28346 | TRUE | FALSE |
| LOC107069951 | probable queuine tRNA-ribosyltransferase | 0.08106 | 154.62924 | TRUE | FALSE |
| LOC107071348 | eukaryotic translation initiation factor 3 sub | 0.10963 | 154.72383 | TRUE | FALSE |
| LOC107066333 | cGMP-specific 3',5'-cyclic phosphodiesteras | 0.1741 | 154.72491 | TRUE | FALSE |
| LOC107065128 | histone H2A-like | 0.10899 | 154.83985 | TRUE | TRUE |
| LOC107071668 | cytoplasmic polyadenylation element-bindi | 0.1886 | 154.86892 | TRUE | TRUE |
| LOC107073556 | uncharacterized protein LOC107073556 | 0.04495 | 155.09453 | TRUE | FALSE |
| LOC107065021 | chromobox protein homolog 1 | 0.06198 | 155.34915 | TRUE | FALSE |
| LOC107068189 | monocarboxylate transporter 3-like | 0.1329 | 155.58061 | TRUE | FALSE |
| LOC107074625 | uncharacterized protein LOC107074625 | 0.09115 | 155.9397 | TRUE | FALSE |
| LOC107072341 | uncharacterized protein LOC107072341 | 0.10823 | 156.15286 | TRUE | FALSE |
| LOC107064086 | polyribonucleotide nucleotidyltransferase 1 | 0.07311 | 156.15448 | TRUE | FALSE |

| | | | | | |
|---|---|---|---|---|---|
| LOC107063885 | uncharacterized protein LOC107063885 | 0.12897 | 156.28173 | TRUE | FALSE |
| LOC107071368 | uncharacterized protein LOC107071368 | 0.14387 | 156.42449 | TRUE | FALSE |
| LOC107064843 | neuropeptide-like 3 | NA | 156.51897 | TRUE | FALSE |
| LOC107070739 | nudC domain-containing protein 1 | 0.10749 | 156.79609 | TRUE | FALSE |
| LOC107063754 | uncharacterized protein LOC107063754 | 0.03941 | 156.95826 | TRUE | FALSE |
| LOC107073578 | uncharacterized protein LOC107073578 | 0.05782 | 156.95856 | TRUE | FALSE |
| LOC107065233 | nuclear hormone receptor FTZ-F1 | 0.17273 | 156.99478 | TRUE | FALSE |
| LOC107070690 | uncharacterized protein LOC107070690 | 0.04396 | 157.03148 | TRUE | FALSE |
| LOC107070930 | NADH-cytochrome b5 reductase 3 | 0.15283 | 157.08006 | TRUE | FALSE |
| LOC107066640 | zinc/cadmium resistance protein | 0.16137 | 157.40241 | TRUE | FALSE |
| LOC107064783 | sarcolemmal membrane-associated protein | 0.10174 | 157.41712 | TRUE | FALSE |
| LOC107069369 | uncharacterized protein LOC107069369 | 0.1291 | 157.49343 | TRUE | FALSE |
| LOC107068718 | uncharacterized protein LOC107068718 | 0.12383 | 157.57018 | TRUE | FALSE |
| LOC107072774 | intraflagellar transport protein 22 homolog | 0.03448 | 157.72253 | TRUE | FALSE |
| LOC107073372 | beta-1,3-galactosyltransferase 6 | 0.05315 | 157.78117 | TRUE | FALSE |
| LOC107068627 | uncharacterized protein LOC107068627 | 0.1214 | 157.83173 | TRUE | FALSE |
| LOC107069482 | pyruvate dehydrogenase E1 component sub | 0.08493 | 157.98348 | TRUE | FALSE |
| LOC107065216 | protein expanded | 0.18333 | 158.02544 | TRUE | FALSE |
| LOC107063792 | dyslexia susceptibility 1 candidate gene 1 pr | NA | 158.10519 | TRUE | TRUE |
| LOC107072003 | tribbles homolog 2 | 0.05815 | 158.1115 | TRUE | FALSE |
| LOC107064316 | zinc finger C4H2 domain-containing protein | 0.10697 | 158.31417 | TRUE | FALSE |
| LOC107073017 | general transcription factor II-I repeat doma | 0.08211 | 158.34415 | TRUE | FALSE |
| LOC107068364 | uncharacterized protein LOC107068364 | 0.07891 | 158.40926 | TRUE | FALSE |
| LOC107070435 | RING finger protein 44 | NA | 158.54598 | TRUE | FALSE |
| LOC107063636 | protein unc-80 homolog | 0.1704 | 158.57045 | TRUE | FALSE |
| LOC107067743 | uncharacterized protein LOC107067743 | 0.09156 | 158.58291 | TRUE | FALSE |
| LOC107064264 | MOXD1 homolog 1 | NA | 158.66002 | TRUE | TRUE |
| LOC107071392 | ubiquitin carboxyl-terminal hydrolase isozyr | 0.07834 | 158.86717 | TRUE | FALSE |
| LOC107068843 | flotillin-1 | NA | 158.93738 | TRUE | FALSE |
| LOC107067025 | uncharacterized protein LOC107067025 | 0.1258 | 158.99141 | TRUE | TRUE |
| LOC107066293 | cytospin-B | NA | 159.12845 | TRUE | FALSE |
| LOC107064853 | uncharacterized protein LOC107064853 | 0.17787 | 159.16227 | TRUE | FALSE |
| LOC107070810 | uncharacterized protein LOC107070810 | 0.0467 | 159.23455 | TRUE | FALSE |
| LOC107066018 | nuclear hormone receptor E75 | NA | 159.29061 | TRUE | FALSE |
| LOC107069081 | homeobox protein Nkx-2.2a-like | 0.08023 | 159.30326 | TRUE | FALSE |
| LOC107065922 | uncharacterized protein LOC107065922 | NA | 159.812 | TRUE | FALSE |
| LOC107069057 | charged multivesicular body protein 1b | 0.16977 | 159.84633 | TRUE | FALSE |
| LOC107074600 | survival of motor neuron protein | 0.09267 | 160.09107 | TRUE | FALSE |
| LOC107073708 | uncharacterized protein LOC107073708 | 0.16467 | 160.22235 | TRUE | FALSE |
| LOC107069084 | uncharacterized protein LOC107069084 | NA | 160.31946 | TRUE | FALSE |
| LOC107064302 | Krueppel homolog 1-like | NA | 160.3688 | TRUE | TRUE |
| LOC107072528 | serine/threonine-protein kinase ICK | NA | 160.41449 | TRUE | FALSE |
| LOC107071188 | carbonic anhydrase-related protein 10 | 0.13767 | 160.85876 | TRUE | FALSE |
| LOC107066716 | uncharacterized protein LOC107066716 | 0.04693 | 160.87026 | TRUE | FALSE |
| LOC107074333 | DNA-directed RNA polymerase II subunit RP | 0.08971 | 160.87927 | TRUE | FALSE |
| LOC107064554 | uncharacterized protein LOC107064554 | 0.07251 | 160.91962 | TRUE | FALSE |
| LOC107063797 | uncharacterized protein LOC107063797 | 0.07172 | 161.02449 | TRUE | FALSE |
| LOC107065176 | pyridoxal-dependent decarboxylase domain | 0.14897 | 161.44532 | TRUE | FALSE |
| LOC107069389 | putative acyl-CoA-binding protein | 0.08727 | 161.53429 | TRUE | FALSE |
| LOC107065727 | transmembrane protein 138 | 0.08189 | 161.86596 | TRUE | FALSE |

| | | | | | |
|---|---|---|---|---|---|
| LOC107074073 | hypoxia-inducible factor prolyl hydroxylase | 0.14073 | 161.88126 | TRUE | FALSE |
| LOC107067158 | RNA polymerase II degradation factor 1 | NA | 161.98971 | TRUE | TRUE |
| LOC107064735 | uncharacterized protein LOC107064735 | 0.08006 | 161.99219 | TRUE | FALSE |
| LOC107067978 | zeta-sarcoglycan | 0.1035 | 162.22821 | TRUE | FALSE |
| LOC107068008 | cell cycle checkpoint protein RAD1 | 0.1312 | 162.24468 | TRUE | FALSE |
| LOC107069256 | uncharacterized protein LOC107069256 | 0.139 | 162.32579 | TRUE | FALSE |
| LOC107074314 | hexosaminidase D-like | 0.07088 | 162.41476 | TRUE | FALSE |
| LOC107071199 | uncharacterized protein LOC107071199 | 0.1459 | 162.48596 | TRUE | FALSE |
| LOC107066902 | histone-lysine N-methyltransferase 2C-like | NA | 162.881 | TRUE | FALSE |
| LOC107072959 | fibroblast growth factor 13-like | 0.15577 | 162.98794 | TRUE | FALSE |
| LOC107064222 | alanine--glyoxylate aminotransferase 2 hom | NA | 163.07339 | TRUE | TRUE |
| LOC107070438 | uncharacterized protein LOC107070438 | 0.14387 | 163.1001 | TRUE | FALSE |
| LOC107073544 | uncharacterized protein LOC107073544 | NA | 163.35462 | TRUE | FALSE |
| LOC107066843 | calcium channel flower | 0.07395 | 163.42015 | TRUE | FALSE |
| LOC107073288 | growth hormone-regulated TBC protein 1-A | 0.09082 | 163.50447 | TRUE | FALSE |
| LOC107064656 | protein Asterix | 0.06137 | 163.51798 | TRUE | FALSE |
| LOC107066167 | alpha-tocopherol transfer protein-like | NA | 163.67241 | TRUE | FALSE |
| LOC107066788 | gustatory receptor 68a-like | 0.11147 | 163.76345 | TRUE | FALSE |
| LOC107066497 | max-interacting protein 1-like | NA | 164.07789 | TRUE | TRUE |
| LOC107074200 | 26S proteasome non-ATPase regulatory sub | NA | 164.12949 | TRUE | TRUE |
| LOC107065006 | uncharacterized protein LOC107065006 | 0.06837 | 164.15091 | TRUE | FALSE |
| LOC107064234 | uncharacterized protein LOC107064234 | 0.12413 | 164.18621 | TRUE | FALSE |
| LOC107067793 | ubiquitin-like modifier-activating enzyme 5 | NA | 164.21946 | TRUE | FALSE |
| LOC107064537 | uncharacterized protein LOC107064537 | 0.19183 | 164.51768 | TRUE | FALSE |
| LOC107069697 | uncharacterized protein LOC107069697 | 0.16627 | 164.54147 | TRUE | FALSE |
| LOC107073831 | myocyte-specific enhancer factor 2 | 0.1562 | 164.61106 | TRUE | FALSE |
| LOC107067185 | arylphorin subunit beta-like | NA | 164.61296 | TRUE | TRUE |
| LOC107072308 | chaoptin | NA | 164.76198 | TRUE | TRUE |
| LOC107073568 | protein FAM46C | NA | 164.95759 | TRUE | TRUE |
| LOC107068066 | RNA polymerase II-associated factor 1 homo | NA | 165.01625 | TRUE | FALSE |
| LOC107064179 | m7GpppN-mRNA hydrolase | 0.07958 | 165.09974 | TRUE | FALSE |
| LOC107068655 | aryl hydrocarbon receptor nuclear transloca | NA | 165.26445 | TRUE | FALSE |
| LOC107073846 | protein aubergine | 0.07688 | 165.44941 | TRUE | FALSE |
| LOC107065482 | uncharacterized protein LOC107065482 | NA | 165.64104 | TRUE | FALSE |
| LOC107064660 | probable GDP-L-fucose synthase | 0.10529 | 166.35295 | TRUE | FALSE |
| LOC107071784 | S-phase kinase-associated protein 2-like | 0.1305 | 166.62719 | TRUE | FALSE |
| LOC107065063 | uncharacterized protein LOC107065063 | 0.13037 | 166.96243 | TRUE | FALSE |
| LOC107064461 | probable serine/threonine-protein kinase rc | 0.16223 | 167.17603 | TRUE | FALSE |
| LOC107073995 | proteasome subunit alpha type-6-like | 0.11963 | 167.42707 | TRUE | FALSE |
| LOC107068543 | E3 ubiquitin-protein ligase RNF126 | NA | 167.69124 | TRUE | FALSE |
| LOC107070844 | uncharacterized protein LOC107070844 | NA | 167.8245 | TRUE | FALSE |
| LOC107070689 | uncharacterized protein LOC107070689 | 0.0585 | 167.95755 | TRUE | FALSE |
| LOC107072438 | uncharacterized protein C15orf41 homolog | NA | 167.98083 | TRUE | FALSE |
| LOC107064221 | uncharacterized protein LOC107064221 | NA | 168.12404 | TRUE | TRUE |
| LOC107072753 | zinc finger protein ubi-d4 A | 0.1173 | 168.29211 | TRUE | FALSE |
| LOC107070605 | alpha-tocopherol transfer protein-like | 0.13623 | 168.47994 | TRUE | FALSE |
| LOC107066711 | fas apoptotic inhibitory molecule 1 | 0.1846 | 168.58541 | TRUE | FALSE |
| LOC107068154 | carnosine N-methyltransferase | NA | 168.74431 | TRUE | FALSE |
| LOC107071044 | uncharacterized protein LOC107071044 | 0.18593 | 168.77309 | TRUE | FALSE |
| LOC107067026 | uncharacterized protein LOC107067026 | 0.1845 | 168.90489 | TRUE | TRUE |

| | | | | | |
|---|---|---|---|---|---|
| LOC107071280 | zinc finger protein Noc | 0.18843 | 169.40702 | TRUE | FALSE |
| LOC107066966 | histone H2B-like | 0.16763 | 169.49884 | TRUE | TRUE |
| LOC107064220 | uncharacterized protein LOC107064220 | NA | 169.56771 | TRUE | TRUE |
| LOC107073413 | DNA polymerase epsilon subunit C | 0.1709 | 169.72963 | TRUE | FALSE |
| LOC107071696 | synapsin | 0.17503 | 170.76909 | TRUE | FALSE |
| LOC107065853 | flexible cuticle protein 12-like | 0.12503 | 170.85257 | TRUE | FALSE |
| LOC107070749 | ATP-binding cassette sub-family G member | 0.07465 | 171.37131 | TRUE | FALSE |
| LOC107065297 | protein bric-a-brac 2-like | NA | 171.49698 | TRUE | TRUE |
| LOC107070074 | extensin | 0.17493 | 171.63877 | TRUE | FALSE |
| LOC107073180 | homeobox protein unplugged | 0.08132 | 171.82514 | TRUE | FALSE |
| LOC107074399 | selenide, water dikinase | NA | 171.97826 | TRUE | FALSE |
| LOC107068230 | thioredoxin-like protein 4A | 0.08394 | 172.63483 | TRUE | FALSE |
| LOC107073185 | BTB/POZ domain-containing protein 6-like | 0.18043 | 172.79954 | TRUE | FALSE |
| LOC107068444 | F-actin-capping protein subunit alpha | 0.07384 | 173.69774 | TRUE | FALSE |
| LOC107067024 | uncharacterized protein LOC107067024 | 0.17903 | 175.11407 | TRUE | TRUE |
| LOC107071902 | multivesicular body subunit 12A | 0.10015 | 175.56244 | TRUE | FALSE |
| LOC107068481 | glutamine--fructose-6-phosphate aminotrar | NA | 175.60565 | TRUE | FALSE |
| LOC107068187 | copper homeostasis protein cutC homolog | 0.12993 | 175.76714 | TRUE | FALSE |
| LOC107064597 | glutathione S-transferase 1-like | 0.11683 | 175.86464 | TRUE | FALSE |
| LOC107066611 | ubiquitin-conjugating enzyme E2 G2 | 0.1567 | 176.2373 | TRUE | FALSE |
| LOC107071545 | uncharacterized protein LOC107071545 | 0.08292 | 176.41453 | TRUE | FALSE |
| LOC107072852 | pyrroline-5-carboxylate reductase | NA | 177.19842 | TRUE | FALSE |
| LOC107074235 | glutamate receptor ionotropic, delta-1-like | 0.0647 | 177.47811 | TRUE | FALSE |
| LOC107064297 | RNA-binding protein lark | NA | 179.28477 | TRUE | FALSE |
| LOC107064555 | RNA polymerase II degradation factor 1 | 0.16977 | 179.39998 | TRUE | FALSE |
| LOC107074148 | uncharacterized protein LOC107074148 | NA | 179.81426 | TRUE | TRUE |
| LOC107074530 | uncharacterized protein LOC107074530 | 0.18413 | 180.0187 | TRUE | FALSE |
| LOC107074606 | sex peptide receptor | NA | 180.13299 | TRUE | FALSE |
| LOC107073083 | nose resistant to fluoxetine protein 6 | 0.08014 | 181.30891 | TRUE | FALSE |
| LOC107073989 | ubiquitin carboxyl-terminal hydrolase isozyr | NA | 181.37085 | TRUE | FALSE |
| LOC107066480 | uncharacterized protein LOC107066480 | 0.14443 | 181.72871 | TRUE | FALSE |
| LOC107067703 | ubiquitin-like protein 7 | 0.14683 | 182.36636 | TRUE | FALSE |
| LOC107074238 | protein zwilch homolog | 0.18613 | 182.48558 | TRUE | FALSE |
| LOC107067286 | coiled-coil domain-containing protein 103 | NA | 182.67577 | TRUE | TRUE |
| LOC107073993 | cleavage stimulation factor subunit 1 | 0.1721 | 182.96037 | TRUE | FALSE |
| LOC107063916 | acyl-CoA Delta(11) desaturase-like | NA | 183.00519 | TRUE | FALSE |
| LOC107072152 | uncharacterized protein LOC107072152 | 0.08312 | 183.12628 | TRUE | FALSE |
| LOC107065267 | uncharacterized protein LOC107065267 | NA | 183.83705 | TRUE | TRUE |
| LOC107067280 | beta-mannosidase | 0.1068 | 183.85619 | TRUE | FALSE |
| LOC107070446 | TM2 domain-containing protein CG11103 | NA | 184.24729 | TRUE | FALSE |
| LOC107066629 | DCN1-like protein 5 | 0.16847 | 184.46712 | TRUE | FALSE |
| LOC107068311 | putative protein kinase C delta type homolc | NA | 184.78431 | TRUE | FALSE |
| LOC107065057 | uncharacterized protein LOC107065057 | NA | 184.97686 | TRUE | FALSE |
| LOC107064838 | histone H2B-like | NA | 186.98075 | TRUE | TRUE |
| LOC107072226 | NADPH oxidase 5 | 0.06858 | 187.44004 | TRUE | FALSE |
| LOC107073793 | extensin-like | 0.09179 | 187.56495 | TRUE | FALSE |
| LOC107070046 | serine-aspartate repeat-containing protein I | 0.09782 | 188.3454 | TRUE | FALSE |
| LOC107065449 | uncharacterized protein LOC107065449 | NA | 188.82905 | TRUE | FALSE |
| LOC107065493 | 4-hydroxyphenylpyruvate dioxygenase | 0.09032 | 189.84631 | TRUE | FALSE |
| LOC107073984 | cadherin-23 | NA | 190.43003 | TRUE | FALSE |

| | | | | | |
|---|---|---|---|---|---|
| LOC107064035 | uncharacterized protein LOC107064035 | NA | 190.4644 | TRUE | TRUE |
| LOC107074492 | G/T mismatch-specific thymine DNA glycosy | NA | 190.49265 | TRUE | FALSE |
| LOC107069780 | sodium channel protein Nach-like | NA | 190.54227 | TRUE | TRUE |
| LOC107067028 | histone H4-like | 0.13637 | 191.15477 | TRUE | TRUE |
| LOC107067446 | uncharacterized protein LOC107067446 | NA | 191.42659 | TRUE | TRUE |
| LOC107068886 | uncharacterized protein LOC107068886 | 0.08555 | 191.61696 | TRUE | FALSE |
| LOC107064343 | oocyte zinc finger protein XlCOF22 | NA | 192.36597 | TRUE | TRUE |
| LOC107064973 | protein phosphatase 1B | NA | 192.93096 | TRUE | FALSE |
| LOC107073681 | fragile X mental retardation syndrome-relat | NA | 193.00712 | TRUE | FALSE |
| LOC107067027 | histone H4 | 0.1919 | 193.26184 | TRUE | TRUE |
| LOC107074315 | E3 ubiquitin-protein ligase HUWE1 | NA | 194.03592 | TRUE | FALSE |
| LOC107074667 | cell division cycle protein 16 homolog | 0.19313 | 194.22779 | TRUE | FALSE |
| LOC107064159 | serine/threonine-protein phosphatase 4 cat | NA | 194.22991 | TRUE | FALSE |
| LOC107073660 | uncharacterized protein LOC107073660 | NA | 195.08571 | TRUE | TRUE |
| LOC107070789 | uncharacterized protein LOC107070789 | 0.15323 | 195.26083 | TRUE | TRUE |
| LOC107065276 | circadian clock-controlled protein-like | NA | 196.95077 | TRUE | FALSE |
| LOC107066998 | histone H2A | NA | 197.1903 | TRUE | TRUE |
| LOC107067555 | WW domain-binding protein 2 | 0.124 | 197.67295 | TRUE | FALSE |
| LOC107065336 | transmembrane protein 43 homolog | NA | 199.0931 | TRUE | FALSE |
| LOC107064369 | muskelin | NA | 199.76578 | TRUE | FALSE |
| LOC107073753 | piggyBac transposable element-derived pro | 0.13617 | 201.17917 | TRUE | FALSE |
| LOC107067965 | COP9 signalosome complex subunit 6 | NA | 201.36377 | TRUE | FALSE |
| LOC107064611 | forkhead box protein K1 | NA | 203.10989 | TRUE | FALSE |
| LOC107070787 | uncharacterized protein LOC107070787 | NA | 204.28355 | TRUE | TRUE |
| LOC107068501 | ejaculatory bulb-specific protein 3-like | 0.09688 | 205.13379 | TRUE | TRUE |
| LOC107064914 | nuclear envelope phosphatase-regulatory su | NA | 207.62383 | TRUE | FALSE |
| LOC107064442 | histone H3 | NA | 207.81843 | TRUE | TRUE |
| LOC107073333 | uncharacterized protein LOC107073333 | 0.11763 | 207.99112 | TRUE | FALSE |
| LOC107068628 | uncharacterized protein LOC107068628 | NA | 208.48837 | TRUE | FALSE |
| LOC107064248 | ATP-dependent RNA helicase DDX42 | 0.18817 | 208.98102 | TRUE | FALSE |
| LOC107065263 | histone H3 | NA | 209.68052 | TRUE | TRUE |
| LOC107073951 | cleavage and polyadenylation specificity fac | NA | 209.75924 | TRUE | FALSE |
| LOC107073611 | uncharacterized protein LOC107073611 | NA | 209.91375 | TRUE | TRUE |
| LOC107068157 | histone H2A | NA | 213.07651 | TRUE | TRUE |
| LOC107072323 | uncharacterized protein LOC107072323 | NA | 213.22011 | TRUE | FALSE |
| LOC107064451 | histone H4 | NA | 220.02515 | TRUE | TRUE |
| LOC107067260 | transient receptor potential cation channel | NA | 220.51601 | TRUE | TRUE |
| LOC107073714 | uncharacterized protein LOC107073714 | NA | 227.3013 | TRUE | TRUE |
| LOC107065294 | zinc carboxypeptidase-like | NA | 228.45533 | TRUE | TRUE |
| LOC107064462 | histone H4 | NA | 230.47419 | TRUE | TRUE |
| LOC107067723 | toll-like receptor 13 | NA | 243.06574 | TRUE | FALSE |
| LOC107073715 | uncharacterized protein LOC107073715 | NA | 249.54366 | TRUE | TRUE |
| LOC107073769 | uncharacterized protein LOC107073769 | NA | 249.93765 | TRUE | TRUE |
