## Supplementary Table S2 for "The molecular basis of socially-mediated phenotypic plasticity in a eusocial paper wasp"

**Table S2.** Genes remaining in the final, optimised support vector machine. **Weight:** The weight of each feature in the original, unoptimised SVM. Weights were calculated as the matrix product of the SVM's coefficients and its support vector values.

| GeneID | ProteinID | Weight |
|---|---|---|
| LOC107065294 | zinc carboxypeptidase-like | -370.0013714 |
| LOC107072323 | uncharacterized protein LOC107072323 | -354.3646309 |
| LOC107068628 | uncharacterized protein LOC107068628 | -346.7546244 |
| LOC107068157 | histone H2A | -328.9259244 |
| LOC107068501 | ejaculatory bulb-specific protein 3-like | -323.8720195 |
| LOC107065276 | circadian clock-controlled protein-like | -314.2768057 |
| LOC107065449 | uncharacterized protein LOC107065449 | -310.5148419 |
| LOC107073984 | cadherin-23 | -308.502078 |
| LOC107074315 | E3 ubiquitin-protein ligase HUWE1 | -307.4376141 |
| LOC107068187 | copper homeostasis protein cutC homolog | -304.8900247 |
| LOC107070446 | TM2 domain-containing protein CG11103 | -302.3422127 |
| LOC107072152 | uncharacterized protein LOC107072152 | -302.1907432 |
| LOC107074606 | sex peptide receptor | -300.9952454 |
| LOC107074530 | uncharacterized protein LOC107074530 | -299.3212716 |
| LOC107065057 | uncharacterized protein LOC107065057 | -297.4571349 |
| LOC107064035 | uncharacterized protein LOC107064035 | -293.1652327 |
| LOC107072438 | uncharacterized protein C15orf41 homolog | -292.1275146 |
| LOC107074238 | protein zwilch homolog | -291.4796956 |
| LOC107071545 | uncharacterized protein LOC107071545 | -291.4599548 |
| LOC107068230 | thioredoxin-like protein 4A | -290.5238349 |
| LOC107070074 | extensin | -290.3179988 |
| LOC107073083 | nose resistant to fluoxetine protein 6 | -289.9785502 |
| LOC107065267 | uncharacterized protein LOC107065267 | -289.7898739 |
| LOC107065006 | uncharacterized protein LOC107065006 | -288.9881326 |
| LOC107068444 | F-actin-capping protein subunit alpha | -288.9802941 |
| LOC107074235 | glutamate receptor ionotropic, delta-1-like | -287.8136277 |
| LOC107072226 | NADPH oxidase 5 | -287.3781288 |
| LOC107070810 | uncharacterized protein LOC107070810 | -287.2373047 |
| LOC107064597 | glutathione S-transferase 1-like | -285.2841625 |
| LOC107066716 | uncharacterized protein LOC107066716 | -280.9842217 |
| LOC107070689 | uncharacterized protein LOC107070689 | -280.4334862 |
| LOC107069190 | uncharacterized protein LOC107069190 | -279.5282455 |
| LOC107069427 | lipase 3-like | -276.6333245 |
| LOC107064554 | uncharacterized protein LOC107064554 | -276.0669186 |
| LOC107064735 | uncharacterized protein LOC107064735 | -275.2467847 |
| LOC107070749 | ATP-binding cassette sub-family G member 4-like | -273.0342525 |
| LOC107064853 | uncharacterized protein LOC107064853 | -272.0715442 |
| LOC107063636 | protein unc-80 homolog | -271.5944551 |
| LOC107066902 | histone-lysine N-methyltransferase 2C-like | -271.3548301 |
| LOC107072959 | fibroblast growth factor 13-like | -271.3412411 |
| LOC107067793 | ubiquitin-like modifier-activating enzyme 5 | -270.7140414 |
| LOC107066134 | exostosin-2 | -268.8014228 |
| LOC107073846 | protein aubergine | -268.2017294 |
| LOC107073017 | general transcription factor II-I repeat domain-containing prc | -267.9089106 |
| LOC107064316 | zinc finger C4H2 domain-containing protein | -267.8316526 |

| | | |
|---|---|---|
| LOC107069256 | uncharacterized protein LOC107069256 | -267.3554852 |
| LOC107069084 | uncharacterized protein LOC107069084 | -267.2414945 |
| LOC107070605 | alpha-tocopherol transfer protein-like | -266.9450865 |
| LOC107064234 | uncharacterized protein LOC107064234 | -266.570854 |
| LOC107063797 | uncharacterized protein LOC107063797 | -266.5349859 |
| LOC107069482 | pyruvate dehydrogenase E1 component subunit beta, mitoch | -266.279321 |
| LOC107074600 | survival of motor neuron protein | -265.3456312 |
| LOC107068364 | uncharacterized protein LOC107068364 | -264.9644888 |
| LOC107064783 | sarcolemmal membrane-associated protein | -264.3414884 |
| LOC107067158 | RNA polymerase II degradation factor 1 | -264.3037509 |
| LOC107071784 | S-phase kinase-associated protein 2-like | -264.0982017 |
| LOC107072168 | cilia- and flagella-associated protein 70 | -263.2740478 |
| LOC107065063 | uncharacterized protein LOC107065063 | -262.7268915 |
| LOC107065853 | flexible cuticle protein 12-like | -262.70609 |
| LOC107065021 | chromobox protein homolog 1 | -261.9698808 |
| LOC107070690 | uncharacterized protein LOC107070690 | -261.9407028 |
| LOC107065482 | uncharacterized protein LOC107065482 | -261.8493181 |
| LOC107065713 | uncharacterized protein LOC107065713 | -261.8374049 |
| LOC107069697 | uncharacterized protein LOC107069697 | -260.8291073 |
| LOC107074200 | 26S proteasome non-ATPase regulatory subunit 9 | -260.0756218 |
| LOC107066293 | cytospin-B | -259.776804 |
| LOC107068843 | flotillin-1 | -259.7673132 |
| NA | NA | -259.6662182 |
| LOC107067185 | arylphorin subunit beta-like | -259.4053866 |
| LOC107068214 | thioredoxin, mitochondrial | -258.9697033 |
| LOC107070119 | TNF receptor-associated factor 6-like | -258.530167 |
| LOC107066649 | uncharacterized protein LOC107066649 | -258.0811788 |
| LOC107066333 | cGMP-specific 3',5'-cyclic phosphodiesterase | -258.0299771 |
| LOC107066799 | gustatory receptor for sugar taste 43a-like | -257.86845 |
| LOC107063640 | glutamate receptor ionotropic, kainate 2 | -257.76459 |
| LOC107070739 | nudC domain-containing protein 1 | -257.7164546 |
| LOC107069389 | putative acyl-CoA-binding protein | -257.6802572 |
| LOC107072528 | serine/threonine-protein kinase ICK | -257.0656487 |
| LOC107071392 | ubiquitin carboxyl-terminal hydrolase isozyme L5 | -256.2234331 |
| LOC107066167 | alpha-tocopherol transfer protein-like | -256.1515264 |
| LOC107070595 | protein lifeguard 3-like | -256.0047642 |
| LOC107066603 | uncharacterized protein LOC107066603 | -255.292544 |
| LOC107072650 | glycerate kinase-like | -255.1625225 |
| LOC107064924 | uncharacterized protein LOC107064924 | -255.094201 |
| LOC107071348 | eukaryotic translation initiation factor 3 subunit K | -255.089314 |
| LOC107074687 | uncharacterized protein LOC107074687 | -255.0687031 |
| LOC107064843 | neuropeptide-like 3 | -254.595071 |
| LOC107064086 | polyribonucleotide nucleotidyltransferase 1, mitochondrial | -254.5561055 |
| LOC107065362 | nitrate reductase [NAD(P)H | -254.4174132 |
| LOC107066094 | neurexin-2 | -253.9026339 |
| LOC107072729 | F-box/WD repeat-containing protein 9-like | -253.6369291 |
| LOC107071344 | COP9 signalosome complex subunit 7 | -252.614383 |
| LOC107064191 | tRNA-specific adenosine deaminase 1 | -252.5542694 |
| LOC107063885 | uncharacterized protein LOC107063885 | -251.3466131 |
| LOC107069951 | probable queuine tRNA-ribosyltransferase | -250.7333895 |

| | | |
|---|---|---|
| LOC107064937 | dynein heavy chain 7, axonemal-like | -250.6851799 |
| LOC107064837 | mediator of RNA polymerase II transcription subunit 2-like | -250.225624 |
| LOC107074625 | uncharacterized protein LOC107074625 | -250.063833 |
| LOC107073708 | uncharacterized protein LOC107073708 | -249.8932022 |
| LOC107064082 | focal adhesion kinase 1 | -249.6922266 |
| LOC107072477 | uncharacterized protein LOC107072477 | -249.4612866 |
| LOC107066465 | protein EMRE homolog, mitochondrial | -249.0404148 |
| LOC107064264 | MOXD1 homolog 1 | -248.910181 |
| LOC107070237 | sensory neuron membrane protein 1 | -248.7438541 |
| LOC107065851 | biogenesis of lysosome-related organelles complex 1 subunit | -248.6848741 |
| LOC107071450 | piggyBac transposable element-derived protein 4-like | -248.3056892 |
| LOC107067205 | proton-coupled amino acid transporter 4 | -247.9656485 |
| LOC107068360 | spondin-1-like | -247.9644587 |
| LOC107073571 | disks large-associated protein 4-like | -246.8762531 |
| LOC107071368 | uncharacterized protein LOC107071368 | -246.6134751 |
| LOC107066291 | uncharacterized protein LOC107066291 | -246.4760153 |
| LOC107073625 | protein toll-like | -246.2345248 |
| LOC107071140 | TGF-beta receptor type-1 | -246.1351472 |
| LOC107066108 | peroxidasin | -245.2825588 |
| LOC107073455 | probable multidrug resistance-associated protein lethal(2)03 | -245.1828523 |
| LOC107068131 | uncharacterized protein LOC107068131 | -244.8802733 |
| LOC107064149 | putative uncharacterized protein DDB_G0282133 | -243.8862803 |
| LOC107070800 | uncharacterized protein LOC107070800 | -243.6821416 |
| LOC107074471 | uncharacterized protein LOC107074471 | -243.6226785 |
| LOC107071971 | mitochondrial sodium/hydrogen exchanger 9B2-like | -243.2802796 |
| LOC107068540 | uncharacterized protein LOC107068540 | -242.8148718 |
| LOC107069891 | poly(A) RNA polymerase, mitochondrial-like | -242.3836436 |
| LOC107064902 | pseudouridylate synthase 7 homolog | -241.8534171 |
| LOC107066672 | mitochondrial import inner membrane translocase subunit T | -239.1819356 |
| LOC107067569 | 97 kDa heat shock protein | -238.7451697 |
| LOC107071337 | ubiquitin carboxyl-terminal hydrolase 34 | -238.5575506 |
| LOC107069924 | odorant receptor 13a-like | -238.1884087 |
| LOC107074077 | uncharacterized protein LOC107074077 | -237.8060147 |
| LOC107066205 | WD repeat and FYVE domain-containing protein 3 | -237.6788106 |
| LOC107064036 | 28S ribosomal protein S31, mitochondrial | -237.4534557 |
| LOC107074640 | bifunctional 3'-phosphoadenosine 5'-phosphosulfate synthas | -236.8837737 |
| LOC107064949 | adult-specific cuticular protein ACP-22-like | -236.4016714 |
| LOC107070870 | PCNA-associated factor-like | -236.2220235 |
| LOC107064317 | malate dehydrogenase, cytoplasmic | -236.2202257 |
| LOC107071881 | gastrula zinc finger protein xFG20-1-like | -235.7007385 |
| LOC107073290 | protein HEXIM1-like | -235.5563172 |
| LOC107066753 | uncharacterized protein LOC107066753 | -235.2325438 |
| LOC107071721 | out at first protein | -235.0415559 |
| LOC107068115 | uncharacterized protein LOC107068115 | -234.8951075 |
| LOC107072645 | transcription initiation factor TFIID subunit 10-like | -234.7230463 |
| LOC107069464 | uncharacterized protein LOC107069464 | -234.6135489 |
| LOC107068735 | unconventional myosin-Va | -234.4710322 |
| LOC107070778 | WD repeat-containing protein 43-like | -234.2927192 |
| LOC107067748 | ribonuclease kappa | -233.8484586 |
| LOC107071561 | uncharacterized protein LOC107071561 | -233.5924621 |

| | | |
|---|---|---|
| LOC107070771 | ribonucleoside-diphosphate reductase large subunit | -233.4641768 |
| LOC107065283 | protein abnormal spindle | -233.2542942 |
| LOC107065108 | metallophosphoesterase 1 | -233.0851826 |
| LOC107064466 | uncharacterized protein LOC107064466 | -232.4137357 |
| LOC107064481 | uncharacterized protein LOC107064481 | -232.3980445 |
| LOC107068627 | uncharacterized protein LOC107068627 | -232.1545531 |
| LOC107071369 | piggyBac transposable element-derived protein 4-like | -232.1502911 |
| LOC107063672 | LOW QUALITY PROTEIN: uncharacterized protein LOC107063 | -232.1339157 |
| LOC107067560 | biogenesis of lysosome-related organelles complex 1 subunit | -232.1227804 |
| LOC107065767 | Golgi apparatus protein 1 | -231.7589572 |
| LOC107072648 | zinc finger protein 598 | -231.295856 |
| LOC107071945 | cysteine--tRNA ligase, cytoplasmic | -230.9438997 |
| LOC107070294 | nucleotide exchange factor SIL1 | -230.2649001 |
| LOC107068377 | zinc finger protein 62-like | -229.6059668 |
| LOC107066824 | uncharacterized protein LOC107066824 | -229.5072663 |
| LOC107069881 | uncharacterized protein LOC107069881 | -229.3020977 |
| LOC107068888 | odorant receptor 13a-like | -228.9185894 |
| LOC107069573 | MATH and LRR domain-containing protein PFE0570w | -228.8288739 |
| LOC107065317 | uncharacterized protein LOC107065317 | -228.7648667 |
| LOC107066130 | gonadotropin-releasing hormone receptor | -228.761343 |
| LOC107071987 | gamma-tubulin complex component 4 | -228.4027702 |
| LOC107067474 | elongation factor Ts, mitochondrial | -228.3056759 |
| LOC107072986 | sodium-independent sulfate anion transporter-like | -228.2965031 |
| LOC107071295 | UTP--glucose-1-phosphate uridylyltransferase | -228.2471499 |
| LOC107069066 | larval cuticle protein A1A-like | -228.1188551 |
| LOC107065600 | uncharacterized protein LOC107065600 | -227.6976209 |
| LOC107063831 | protein aurora borealis | -227.4871641 |
| LOC107070729 | neutral ceramidase | -227.1316802 |
| LOC107065865 | etoposide-induced protein 2.4 homolog | -227.1092589 |
| LOC107069061 | uncharacterized protein DDB_G0271670 | -226.5109641 |
| LOC107072089 | transcriptional adapter 2B | -225.9820409 |
| LOC107067314 | proteasome activator complex subunit 4B-like | -225.8698808 |
| LOC107068130 | probable chitinase 2 | -225.7143724 |
| LOC107067654 | protein FAM195A-like | -225.5558015 |
| LOC107068909 | M-phase phosphoprotein 6 | -225.5404013 |
| LOC107067670 | selenoprotein K-like | -225.4032033 |
| LOC107064946 | peflin | -225.0538576 |
| LOC107068740 | E3 ubiquitin-protein ligase RNF19A-like | -224.9048876 |
| LOC107069134 | 2',5'-phosphodiesterase 12 | -224.8152638 |
| LOC107066654 | N-alpha-acetyltransferase 30-like | -224.6389628 |
| LOC107065115 | circadian clock-controlled protein-like | -224.6371829 |
| LOC107064777 | uncharacterized protein LOC107064777 | -224.492792 |
| LOC107064533 | cilia- and flagella-associated protein 52 | -224.4679997 |
| LOC107068819 | uncharacterized protein LOC107068819 | -224.4025208 |
| LOC107072801 | dual specificity protein phosphatase CDC14B-like | -224.2842763 |
| LOC107065523 | zinc finger protein Xfin-like | -224.2562984 |
| LOC107070604 | CRAL-TRIO domain-containing protein DDB_G0278031-like | -224.2442369 |
| LOC107074088 | casein kinase I isoform alpha-like | -223.5388211 |
| LOC107069941 | activating signal cointegrator 1 complex subunit 2 | -223.1860014 |
| LOC107065791 | cell differentiation protein RCD1 homolog | -223.1402642 |

| | | |
|---|---|---|
| LOC107074684 | male-enhanced antigen 1 | -223.1187472 |
| LOC107067252 | 39S ribosomal protein L54, mitochondrial | -222.9368145 |
| LOC107074439 | exostosin-1 | -222.7859338 |
| LOC107063887 | cullin-1 | -222.7226469 |
| LOC107071125 | inactive peptidyl-prolyl cis-trans isomerase FKBP6 | -222.7121492 |
| LOC107064334 | uncharacterized protein LOC107064334 | -222.2881877 |
| LOC107067498 | beta-1,3-galactosyltransferase brn | -222.0699215 |
| LOC107073991 | upstream activation factor subunit spp27 | -221.8077206 |
| LOC107070911 | membrane-associated protein Hem | -221.5096359 |
| LOC107067378 | dynein heavy chain 7, axonemal | -221.367458 |
| LOC107069890 | uncharacterized protein LOC107069890 | -221.3477115 |
| LOC107066980 | cell cycle control protein 50A | -221.3167056 |
| LOC107068704 | endocuticle structural glycoprotein SgAbd-1-like | -221.2267134 |
| LOC107070658 | uncharacterized protein LOC107070658 | -221.0467063 |
| LOC107068837 | uncharacterized protein LOC107068837 | -220.8358986 |
| LOC107071512 | dentin sialophosphoprotein-like | -220.6709672 |
| LOC107064320 | ras-related protein Rac1 | -220.4991244 |
| LOC107067659 | uncharacterized protein LOC107067659 | -220.4461219 |
| LOC107073462 | uncharacterized protein LOC107073462 | -220.3374544 |
| LOC107073905 | UPF0704 protein C6orf165 | -220.3162784 |
| LOC107068679 | protein Simiate | -220.2902274 |
| LOC107065051 | uncharacterized protein LOC107065051 | -220.1509567 |
| LOC107071131 | uncharacterized protein LOC107071131 | -220.1295236 |
| LOC107069242 | protein white | -219.986872 |
| LOC107068375 | uncharacterized protein LOC107068375 | -219.933405 |
| LOC107072077 | UPF0047 protein YjbQ-like | -219.8882723 |
| LOC107072691 | ubiquitin-conjugating enzyme E2 N | -219.4053229 |
| LOC107064148 | uncharacterized family 31 glucosidase KIAA1161 | -219.3988658 |
| LOC107074169 | uncharacterized protein LOC107074169 | -218.9483164 |
| LOC107065150 | uncharacterized protein LOC107065150 | -218.8419553 |
| LOC107074347 | NECAP-like protein CG9132 | -218.7371789 |
| LOC107065863 | SET and MYND domain-containing protein 4-like | -218.701939 |
| LOC107067265 | RNA-binding protein cabeza | -218.6990249 |
| LOC107071292 | RNA-binding protein 40 | -218.6270214 |
| LOC107066456 | UPF0184 protein C9orf16 homolog | -218.6006503 |
| LOC107071701 | leucine-rich repeat-containing protein 70-like | -218.1860852 |
| LOC107065780 | putative uncharacterized protein DDB_G0282133 | -218.0044554 |
| LOC107074607 | uncharacterized protein LOC107074607 | -217.9914612 |
| LOC107070219 | peroxiredoxin-2-like | -216.9633886 |
| LOC107073136 | zinc finger protein 28-like | -216.922027 |
| LOC107066443 | UPF0565 protein C2orf69 homolog | -216.8929703 |
| LOC107073382 | uncharacterized protein LOC107073382 | -216.800763 |
| LOC107068001 | TPPP family protein CG45057-like | -216.7828765 |
| LOC107071353 | GPI ethanolamine phosphate transferase 1 | -216.649087 |
| LOC107069643 | dynein regulatory complex subunit 7 | -216.5986863 |
| LOC107068470 | succinyl-CoA ligase [GDP-forming | -215.9667986 |
| LOC107065102 | arginyl-tRNA--protein transferase 1 | -215.8302617 |
| LOC107068175 | GDP-fucose protein O-fucosyltransferase 2 | -215.6885128 |
| LOC107070818 | leucine-rich repeats and immunoglobulin-like domains prote | -215.6805517 |
| LOC107069183 | uncharacterized protein LOC107069183 | -215.2782942 |

| | | |
|---|---|---|
| LOC107067764 | odorant receptor 4-like | -214.8757719 |
| LOC107065341 | uncharacterized protein LOC107065341 | -214.6522904 |
| LOC107069136 | uncharacterized protein LOC107069136 | -214.5884251 |
| LOC107066607 | ADP-ribosylation factor-like protein 13B | -214.4871109 |
| LOC107065592 | H/ACA ribonucleoprotein complex non-core subunit NAF1-lik | -214.3326671 |
| LOC107074126 | uncharacterized protein LOC107074126 | -214.2696997 |
| LOC107073649 | uncharacterized protein LOC107073649 | -214.1746261 |
| LOC107066460 | uncharacterized protein LOC107066460 | -214.0313004 |
| LOC107064687 | 40S ribosomal protein S24 | -213.6436518 |
| LOC107073531 | peptidyl-alpha-hydroxyglycine alpha-amidating lyase 1-like | -213.4265061 |
| LOC107065010 | mitochondrial dicarboxylate carrier-like | -213.2465523 |
| LOC107071265 | uncharacterized protein LOC107071265 | -213.2276854 |
| LOC107063924 | uncharacterized protein LOC107063924 | -213.0516083 |
| LOC107064603 | 72 kDa inositol polyphosphate 5-phosphatase | -212.9156832 |
| LOC107068170 | putative gamma-glutamylcyclotransferase CG2811 | -212.8672917 |
| LOC107069241 | vesicle transport protein USE1 | -212.5122456 |
| LOC107064827 | cuticle protein 16.5, isoform A | -212.4051836 |
| LOC107067126 | neural Wiskott-Aldrich syndrome protein | -212.263724 |
| LOC107073084 | uncharacterized protein LOC107073084 | -212.1314643 |
| LOC107065087 | putative uncharacterized protein DDB_G0282133 | -211.7266623 |
| LOC107069963 | U6 snRNA phosphodiesterase | -211.6591121 |
| LOC107072894 | uncharacterized protein LOC107072894 | -211.3501571 |
| LOC107068298 | GTPase Era, mitochondrial | -211.1966581 |
| LOC107071284 | stimulator of interferon genes protein | -211.1451644 |
| LOC107065438 | uncharacterized protein LOC107065438 | -211.0491692 |
| LOC107072064 | alpha-tocopherol transfer protein-like | -210.9302694 |
| LOC107074198 | tetratricopeptide repeat protein 30A | -210.8069049 |
| LOC107070095 | zinc finger protein 37-like | -210.802278 |
| LOC107070589 | chondroitin sulfate proteoglycan 4 | -210.7302031 |
| LOC107072910 | uncharacterized protein LOC107072910 | -210.6996337 |
| LOC107073770 | uncharacterized protein LOC107073770 | -210.3746927 |
| LOC107069328 | cyclin-dependent kinase 4-like | -210.2774651 |
| LOC107068056 | probable cation-transporting ATPase 13A3 | -210.2318971 |
| LOC107066848 | LYR motif-containing protein 9-like | -210.1627415 |
| LOC107071506 | clathrin interactor 1 | -210.1254822 |
| LOC107072540 | GTP cyclohydrolase 1 | -209.9667285 |
| LOC107066547 | uncharacterized protein LOC107066547 | -209.7216385 |
| LOC107070223 | exocyst complex component 8 | -209.4584621 |
| LOC107071500 | uncharacterized protein LOC107071500 | -209.3871298 |
| LOC107074584 | DNA-dependent protein kinase catalytic subunit-like | -209.2019846 |
| LOC107068736 | uncharacterized protein LOC107068736 | -208.9821093 |
| LOC107068135 | lysosomal-trafficking regulator | -208.7623023 |
| LOC107070720 | apolipophorins | -208.5171614 |
| LOC107072428 | tetra-peptide repeat homeobox protein 1-like | -208.4944678 |
| LOC107066689 | dystonin | -208.2425396 |
| LOC107068477 | dynein light chain 2, cytoplasmic | -208.1851418 |
| LOC107074230 | probable cytochrome P450 6a13 | -208.1803161 |
| LOC107070501 | uncharacterized protein LOC107070501 | -208.1714046 |
| LOC107067363 | rabenosyn-5 | -208.1627672 |
| LOC107069169 | uncharacterized protein LOC107069169 | -208.1017393 |

| | | |
|---|---|---|
| LOC107070245 | translational activator of cytochrome c oxidase 1 | -207.8287271 |
| LOC107072584 | trafficking protein particle complex subunit 3 | -207.7652146 |
| LOC107071331 | kynurenine 3-monooxygenase | -207.7246199 |
| LOC107071023 | uncharacterized protein LOC107071023 | -207.44875 |
| LOC107073202 | probable galactose-1-phosphate uridylyltransferase | -207.3442967 |
| LOC107067966 | uncharacterized protein LOC107067966 | -206.8264077 |
| LOC107073872 | GPI inositol-deacylase | -206.810825 |
| LOC107070404 | S-phase kinase-associated protein 1 | -206.7488395 |
| LOC107073132 | glutamine synthetase 2 cytoplasmic | -206.5441441 |
| LOC107064651 | ER membrane protein complex subunit 6 | -206.5057941 |
| LOC107071860 | uncharacterized protein LOC107071860 | -206.4527968 |
| LOC107068221 | bromodomain-containing protein 7 | -206.339012 |
| LOC107066053 | protein jagged-1 | -206.2552759 |
| LOC107066072 | G2/mitotic-specific cyclin-B | -206.0917551 |
| LOC107068179 | uncharacterized protein LOC107068179 | -206.0212404 |
| LOC107070314 | CSC1-like protein 2 | -205.9972367 |
| LOC107070587 | clusterin-associated protein 1 | -205.8474431 |
| LOC107064684 | uncharacterized protein LOC107064684 | -205.7496423 |
| LOC107072994 | uncharacterized protein LOC107072994 | -205.7056026 |
| LOC107071294 | voltage-dependent anion-selective channel-like | -205.6370864 |
| LOC107074499 | TOM1-like protein 2 | -205.5681655 |
| LOC107069123 | apoptotic chromatin condensation inducer in the nucleus | -205.5126803 |
| LOC107065876 | double-strand break repair protein MRE11A | -205.3258786 |
| LOC107066445 | uncharacterized protein LOC107066445 | -205.1703246 |
| LOC107073309 | condensin complex subunit 2-like | -205.0178873 |
| LOC107069093 | integrator complex subunit 2 | -204.8832624 |
| LOC107066316 | uncharacterized protein LOC107066316 | -204.8758161 |
| LOC107068020 | spatacsin | -204.7068321 |
| LOC107069349 | dopamine N-acetyltransferase-like | -204.5017793 |
| LOC107071757 | histone acetyltransferase KAT7 | -204.4161462 |
| LOC107065164 | transmembrane protein 18 | -204.2001084 |
| LOC107066977 | prefoldin subunit 4 | -204.1717303 |
| LOC107073933 | acyl-CoA-binding domain-containing protein 6 | -204.1318163 |
| LOC107072805 | TNF receptor-associated factor 5 | -203.9723746 |
| LOC107071064 | Golgi resident protein GCP60 | -203.9695006 |
| LOC107068290 | tubulin delta chain-like | -203.8614357 |
| LOC107074234 | omega-conotoxin-like protein 1 | -203.6862001 |
| LOC107072530 | 2-oxoglutarate and iron-dependent oxygenase domain-conta | -203.6215246 |
| LOC107070578 | uncharacterized protein LOC107070578 | -203.5170721 |
| LOC107068371 | PHD finger protein 12 | -203.4244727 |
| LOC107069670 | TELO2-interacting protein 1 homolog | -202.9669847 |
| LOC107072095 | UPF0605 protein CG18335-like | -202.9022784 |
| LOC107069739 | carbohydrate sulfotransferase 11-like | -202.8775848 |
| LOC107066613 | ADP-ribosylation factor GTPase-activating protein 1 | -202.7905964 |
| LOC107067307 | tight junction protein ZO-2 | -202.7124465 |
| LOC107065890 | superoxide dismutase [Cu-Zn | -202.6102191 |
| LOC107065118 | peroxidasin-like | -202.6097604 |
| LOC107066317 | DENN domain-containing protein 4C | -202.2243491 |
| LOC107070759 | E3 SUMO-protein ligase NSE2-like | -202.154723 |
| LOC107073027 | uncharacterized protein LOC107073027 | -202.1228412 |

| | | |
|---|---|---|
| LOC107072696 | CDP-diacylglycerol--inositol 3-phosphatidyltransferase | -201.9092859 |
| LOC107072081 | UDP-glucose 4-epimerase-like | -201.8797505 |
| LOC107065444 | serine-rich adhesin for platelets | -201.8704994 |
| LOC107064291 | protein mothers against dpp | -201.8301312 |
| LOC107069677 | uncharacterized protein F02A9.4b | -201.8212657 |
| LOC107072060 | serine/threonine-protein phosphatase alpha-2 isoform | -201.735671 |
| LOC107070843 | protein APCDD1-like | -201.7293208 |
| LOC107070675 | adenosine monophosphate-protein transferase FICD homolo | -201.6868572 |
| LOC107064364 | uncharacterized protein LOC107064364 | -201.6383954 |
| LOC107065433 | protein PET100 homolog, mitochondrial | -201.130888 |
| LOC107074213 | glycine receptor subunit alpha-4 | -200.6267158 |
| LOC107063909 | ubiquitin-conjugating enzyme E2 L3 | -200.445356 |
| LOC107069840 | transmembrane protein 41B | -200.4304224 |
| LOC107074029 | uncharacterized protein PFB0765w-like | -200.2887683 |
| LOC107074138 | folliculin | -200.1124013 |
| LOC107065461 | insulin-degrading enzyme | -199.9171189 |
| LOC107067885 | histone-lysine N-methyltransferase 2D-like | -199.7301887 |
| LOC107071607 | protein dopey-1 homolog | -199.5449741 |
| LOC107066312 | dosage compensation regulator | -199.3321075 |
| LOC107073482 | protein toll-like | -199.2926068 |
| LOC107068695 | CDK-activating kinase assembly factor MAT1 | -199.2714208 |
| LOC107069060 | stress response protein NST1-like | -199.1626996 |
| LOC107074086 | dynein heavy chain 8, axonemal | -199.1359486 |
| LOC107068432 | cactin | -199.0999119 |
| LOC107066099 | vacuolar protein sorting-associated protein 13B | -199.0950023 |
| LOC107067311 | probable Rho GTPase-activating protein CG5521 | -199.0301332 |
| LOC107064623 | basic helix-loop-helix neural transcription factor TAP-like | -198.8298883 |
| LOC107071474 | UDP-glucuronosyltransferase 2B4-like | -198.783474 |
| LOC107065618 | INO80 complex subunit C | -198.7132187 |
| LOC107074483 | uncharacterized protein LOC107074483 | -198.6013766 |
| LOC107071301 | uncharacterized protein LOC107071301 | -198.5189985 |
| LOC107070886 | 1,2-dihydroxy-3-keto-5-methylthiopentene dioxygenase | -198.4660796 |
| LOC107071912 | regulator of nonsense transcripts 1 homolog | -198.2670038 |
| LOC107073159 | sodium/potassium-transporting ATPase subunit beta-2-like | -198.151751 |
| LOC107074543 | uncharacterized protein LOC107074543 | -197.9370804 |
| LOC107073128 | surfeit locus protein 1 | -197.8791293 |
| LOC107063897 | LOW QUALITY PROTEIN: ceramide phosphoethanolamine syr | -197.6777213 |
| LOC107064428 | uncharacterized protein LOC107064428 | -197.6298437 |
| LOC107064940 | uncharacterized protein LOC107064940 | -197.5391965 |
| LOC107072376 | uncharacterized membrane protein DDB_G0293934-like | -197.2497165 |
| LOC107064345 | facilitated trehalose transporter Tret1-like | -197.2341498 |
| LOC107067302 | uncharacterized protein LOC107067302 | -197.2309102 |
| LOC107070969 | uncharacterized protein LOC107070969 | -197.1469405 |
| LOC107066324 | protein PFC0760c-like | -197.117321 |
| LOC107069255 | sepiapterin reductase | -197.0617847 |
| LOC107070515 | uncharacterized protein LOC107070515 | -197.0453585 |
| LOC107064796 | 4-coumarate--CoA ligase 1-like | -197.0284511 |
| LOC107068677 | exosome complex component RRP41 | -196.8266885 |
| LOC107072783 | secretin receptor-like | -196.6779826 |
| LOC107072072 | uncharacterized protein LOC107072072 | -196.6758111 |

| | | |
|---|---|---|
| LOC107071185 | uncharacterized protein LOC107071185 | -196.6718128 |
| LOC107067848 | collagen alpha-1(IV) chain | -196.6361331 |
| LOC107069103 | phosphoribosylformylglycinamidine synthase | -196.6242313 |
| LOC107074179 | phospholipase D3-like | -196.6154659 |
| LOC107071841 | rRNA-processing protein FYV7 | -196.3438513 |
| LOC107069306 | uncharacterized protein LOC107069306 | -196.2636037 |
| LOC107072486 | ubiquitin-conjugating enzyme E2 S | -196.2186123 |
| LOC107073148 | amyloid protein-binding protein 2 | -195.978817 |
| LOC107072439 | neprilysin-like | -195.8449016 |
| LOC107067249 | uncharacterized protein LOC107067249 | -195.8121813 |
| LOC107063684 | A-kinase anchor protein 9-like | -195.7078747 |
| LOC107070967 | uncharacterized protein LOC107070967 | -195.6927821 |
| LOC107069104 | alpha-N-acetylgalactosaminidase | -195.6754555 |
| LOC107070319 | odorant receptor 49b-like | -195.5050243 |
| LOC107069003 | calcyclin-binding protein | -195.4704973 |
| LOC107071873 | uncharacterized protein LOC107071873 | -195.3273095 |
| LOC107064674 | transcription elongation factor 1 homolog | -195.3001092 |
| LOC107067009 | uncharacterized protein LOC107067009 | -195.2062457 |
| LOC107069619 | uncharacterized protein LOC107069619 | -195.166262 |
| LOC107071915 | high-affinity potassium transport protein | -194.9720336 |
| LOC107065628 | kinesin light chain | -194.7164107 |
| LOC107064490 | uncharacterized protein LOC107064490 | -194.6281332 |
| LOC107068611 | ribonucleoside-diphosphate reductase large subunit-like | -194.5889282 |
| LOC107071287 | CCHC-type zinc finger protein CG3800 | -194.5441263 |
| LOC107065655 | E3 ubiquitin-protein ligase UBR4 | -194.3347972 |
| LOC107073207 | RNA-binding protein 45 | -194.0977649 |
| LOC107068970 | 28S ribosomal protein S15, mitochondrial | -194.0678329 |
| LOC107070005 | KAT8 regulatory NSL complex subunit 2-like | -193.8979174 |
| LOC107074599 | BTB/POZ domain-containing protein 7 | -193.8267197 |
| LOC107070154 | xylosyltransferase oxt | -193.8093308 |
| LOC107071396 | folylpolyglutamate synthase, mitochondrial-like | -193.7436844 |
| LOC107070807 | polynucleotide 5'-hydroxyl-kinase NOL9-like | -193.6508083 |
| LOC107066164 | uncharacterized protein LOC107066164 | -193.6409522 |
| LOC107074328 | 39S ribosomal protein L37, mitochondrial | -193.3594927 |
| LOC107070426 | F-box/LRR-repeat protein 3-like | -193.2208765 |
| LOC107065173 | histone-lysine N-methyltransferase SUV39H2 | -193.1183775 |
| LOC107066719 | ubiquitin-conjugating enzyme E2 J2 | -192.9974147 |
| LOC107073124 | ragulator complex protein LAMTOR2 homolog | -192.7582105 |
| LOC107066414 | protein cornichon | -192.6379345 |
| LOC107074679 | KAT8 regulatory NSL complex subunit 3 | -192.5785306 |
| LOC107066385 | neuroparsin-A-like | -192.4629115 |
| LOC107066202 | uncharacterized protein LOC107066202 | -192.4209708 |
| LOC107069728 | mitochondrial folate transporter/carrier | -192.39613 |
| LOC107064370 | solute carrier family 35 member G1-like | -192.2829268 |
| LOC107073914 | putative GTP-binding protein 6 | -192.2755623 |
| LOC107068880 | uncharacterized protein LOC107068880 | -192.0003552 |
| LOC107074067 | uncharacterized protein LOC107074067 | -191.8901421 |
| LOC107068087 | proteasome subunit beta type-3 | -191.7907503 |
| LOC107073990 | probable 28S rRNA (cytosine-C(5))-methyltransferase | -191.5537077 |
| LOC107064329 | CLIP-associating protein 1 | -191.5227622 |

| | | |
|---|---|---|
| LOC107073695 | uncharacterized protein LOC107073695 | -191.5195879 |
| LOC107066392 | piwi-like protein 1 | -191.5132038 |
| LOC107068907 | histone H4 transcription factor | -191.504076 |
| LOC107068414 | pro-resilin-like | -191.4431424 |
| LOC107069979 | uncharacterized protein LOC107069979 | -191.1780588 |
| LOC107064878 | V-type proton ATPase subunit G | -191.1750351 |
| LOC107064733 | uncharacterized protein LOC107064733 | -191.1430809 |
| LOC107072020 | protein jagunal | -190.9411852 |
| LOC107069132 | BTB/POZ domain-containing protein 9 | -190.9027466 |
| LOC107068739 | NADP-dependent malic enzyme | -190.8173192 |
| LOC107073457 | uncharacterized protein LOC107073457 | -190.6232911 |
| LOC107070218 | spectrin beta chain | -190.5712661 |
| LOC107064675 | scavenger receptor class B member 1 | -190.5617491 |
| LOC107066250 | protein dachsous | -190.5240865 |
| LOC107070631 | regulator of microtubule dynamics protein 1-like | -190.4634819 |
| LOC107073815 | targeting protein for Xklp2-like | -190.4211151 |
| LOC107073696 | uncharacterized protein LOC107073696 | -190.3965657 |
| LOC107067436 | heat shock factor-binding protein 1 | -190.082562 |
| LOC107068132 | probable cation-transporting ATPase 13A3 | -189.9898035 |
| LOC107063892 | RNA polymerase-associated protein CTR9 homolog | -189.9893839 |
| LOC107073332 | uncharacterized protein LOC107073332 | -189.945553 |
| LOC107072243 | tyrocidine synthase 3 | -189.9275738 |
| LOC107070842 | uncharacterized protein LOC107070842 | -189.8040933 |
| LOC107063815 | huntingtin-interacting protein 1-like | -189.5870142 |
| LOC107063731 | huntingtin | -189.5304856 |
| LOC107069638 | uncharacterized protein LOC107069638 | -189.4685498 |
| LOC107071862 | uncharacterized protein LOC107071862 | -189.3097247 |
| LOC107067383 | uncharacterized protein LOC107067383 | -189.3015225 |
| LOC107073049 | glucose-induced degradation protein 8 homolog | -189.2825046 |
| LOC107069727 | mitochondrial ribonuclease P protein 3 | -189.2751636 |
| LOC107074338 | RING-box protein 2 | -189.0853842 |
| LOC107070122 | EH domain-containing protein 3 | -189.0297488 |
| LOC107068683 | ribosomal protein S6 kinase delta-1 | -188.7521129 |
| LOC107065792 | transcription initiation factor TFIID subunit 11 | -188.7404597 |
| LOC107068891 | growth hormone secretagogue receptor type 1-like | -188.4624912 |
| LOC107064791 | vacuolar fusion protein CCZ1 homolog | -188.4370561 |
| LOC107065038 | uncharacterized protein LOC107065038 | -188.4034395 |
| LOC107066272 | homeobox protein OTX2-A | -188.2336969 |
| LOC107066288 | microtubule-associated protein RP/EB family member 1 | -188.1939892 |
| LOC107064667 | uncharacterized protein LOC107064667 | -188.1877248 |
| LOC107064328 | neuronal PAS domain-containing protein 2-like | -188.0798876 |
| LOC107068937 | farnesol dehydrogenase-like | -187.621281 |
| LOC107063946 | uncharacterized protein LOC107063946 | -187.5305148 |
| LOC107067285 | U6 snRNA-associated Sm-like protein LSm3 | -187.4949509 |
| LOC107064515 | uncharacterized protein LOC107064515 | -187.4810609 |
| LOC107068279 | nucleolar protein 12-like | -187.4653437 |
| LOC107066906 | cell wall integrity and stress response component 2-like | -187.4336949 |
| LOC107070767 | filamin-A | -187.422904 |
| LOC107068738 | RWD domain-containing protein 1 | -187.3932151 |
| LOC107068875 | uncharacterized protein LOC107068875 | -187.3474038 |

| | | |
|---|---|---|
| LOC107074445 | pleckstrin homology domain-containing family M member 1 | -187.2626014 |
| LOC107072689 | putative uncharacterized protein DDB_G0274405 | -187.2065951 |
| LOC107074481 | cell division control protein 6 homolog | -187.11834 |
| LOC107067592 | tachykinin-like peptides receptor 99D | -187.1098436 |
| LOC107071623 | lysozyme-like | -187.0477821 |
| LOC107070714 | uncharacterized protein LOC107070714 | -186.9126692 |
| LOC107071626 | glycosylphosphatidylinositol anchor attachment 1 protein | -186.7179711 |
| LOC107072252 | ATP-dependent RNA helicase DHX8 | -186.5735355 |
| LOC107067682 | uncharacterized protein LOC107067682 | -186.5496434 |
| LOC107072387 | putative uncharacterized protein DDB_G0282133 | -186.4755015 |
| LOC107074239 | chromosome transmission fidelity protein 8 homolog | -186.4382306 |
| LOC107068181 | uncharacterized protein LOC107068181 | -186.4161811 |
| LOC107071727 | E3 ubiquitin-protein ligase RNF123-like | -186.3856149 |
| LOC107064218 | bipolar kinesin KRP-130-like | -186.2408894 |
| LOC107065353 | venom dipeptidyl peptidase 4 | -186.1384167 |
| LOC107070994 | NTF2-related export protein | -186.0507814 |
| LOC107064693 | uncharacterized protein LOC107064693 | -185.9552559 |
| LOC107074380 | tRNA-splicing endonuclease subunit Sen2 | -185.9511546 |
| LOC107071871 | leucine-rich repeat-containing protein 51-like | -185.9294821 |
| LOC107063828 | defensin-1-like | -185.9167146 |
| LOC107068846 | ADP,ATP carrier protein-like | -185.8405936 |
| LOC107073024 | cell wall integrity and stress response component 1-like | -185.7419854 |
| LOC107068069 | uncharacterized protein LOC107068069 | -185.6310195 |
| LOC107073001 | DNA-directed RNA polymerase III subunit RPC8 | -185.6189356 |
| LOC107073192 | protein O-mannosyltransferase 1 | -185.5245474 |
| LOC107071702 | dnaJ homolog subfamily C member 3 | -185.3953318 |
| LOC107070839 | protein phosphatase 1 regulatory subunit 12A | -185.3226686 |
| LOC107072411 | piggyBac transposable element-derived protein 4-like | -185.2843154 |
| LOC107070600 | protein PFF0380w | -185.2344951 |
| LOC107064907 | ES1 protein homolog, mitochondrial-like | -185.2185355 |
| LOC107073874 | 6-phosphofructo-2-kinase/fructose-2,6-bisphosphatase | -184.840003 |
| LOC107068072 | cilia- and flagella-associated protein 45 | -184.8102144 |
| LOC107071994 | osteoclast-stimulating factor 1-like | -184.7934553 |
| LOC107072320 | GPN-loop GTPase 2 | -184.6214365 |
| LOC107063968 | neuroglobin-like | -184.568061 |
| LOC107071582 | nardilysin-like | -184.5628367 |
| LOC107065286 | graves disease carrier protein | -184.5186625 |
| LOC107071005 | ubiquitin-conjugating enzyme E2 J1-like | -184.5090863 |
| LOC107065171 | E3 ubiquitin-protein ligase TRIM33-like | -184.37427 |
| LOC107068566 | maternal B9.10 protein-like | -184.3325799 |
| LOC107064378 | mitochondrial 2-oxoglutarate/malate carrier protein-like | -184.1637986 |
| LOC107067135 | uroporphyrinogen decarboxylase | -184.0624119 |
| LOC107070244 | hepatocyte growth factor-regulated tyrosine kinase substrat | -184.0250552 |
| LOC107068706 | protein brambleberry-like | -183.9824475 |
| LOC107067633 | uncharacterized protein LOC107067633 | -183.9532645 |
| LOC107068580 | pyridoxine-5'-phosphate oxidase-like | -183.9203931 |
| LOC107070358 | uncharacterized protein LOC107070358 | -183.8922499 |
| LOC107063626 | piggyBac transposable element-derived protein 4-like | -183.8791407 |
| LOC107065462 | adhesion G protein-coupled receptor A3 | -183.7280855 |
| LOC107064145 | retinoblastoma-like protein 1 | -183.7262711 |

| | | |
|---|---|---|
| LOC107067392 | CD151 antigen-like | -183.6051028 |
| LOC107072620 | histone deacetylase 6 | -183.5151249 |
| LOC107072613 | MMS19 nucleotide excision repair protein homolog | -183.5125331 |
| LOC107072944 | uncharacterized protein LOC107072944 | -183.5077816 |
| LOC107070138 | signal recognition particle 14 kDa protein | -183.4271247 |
| LOC107068440 | uncharacterized protein LOC107068440 | -183.4011675 |
| LOC107065986 | RNA-binding motif protein, X-linked 2 | -183.3809502 |
| LOC107072353 | uncharacterized protein LOC107072353 | -183.3790004 |
| LOC107068794 | stress-induced-phosphoprotein 1 | -183.3040211 |
| LOC107072247 | 28S ribosomal protein S27, mitochondrial-like | -182.8829135 |
| LOC107070966 | protein kinase C | -182.8792934 |
| LOC107074675 | protein zer-1 homolog | -182.8210099 |
| LOC107074615 | HIG1 domain family member 2A, mitochondrial | -182.7278832 |
| LOC107063720 | proteasome-associated protein ECM29 homolog | -182.6321553 |
| LOC107070655 | peptidyl-prolyl cis-trans isomerase-like | -182.5799292 |
| LOC107065654 | CDK5 regulatory subunit-associated protein 3 | -182.5740226 |
| LOC107069686 | U7 snRNA-associated Sm-like protein LSm11 | -182.4220706 |
| LOC107063637 | tetratricopeptide repeat protein 1 | -182.3204237 |
| LOC107069998 | calcium and integrin-binding protein 1-like | -182.3074293 |
| LOC107064802 | prefoldin subunit 6 | -182.079221 |
| LOC107063914 | peptide transporter family 1 | -181.9850784 |
| LOC107066942 | transcription and mRNA export factor ENY2 | -181.9401054 |
| LOC107068352 | phosphatidylinositol 4-kinase alpha | -181.8336867 |
| LOC107072874 | anaphase-promoting complex subunit 10 | -181.8198095 |
| LOC107072296 | 5-hydroxytryptamine receptor 2B-like | -181.7472282 |
| LOC107072263 | probable WRKY transcription factor protein 1 | -181.5975001 |
| LOC107072531 | fatty acid synthase | -181.5946902 |
| LOC107065736 | DNA topoisomerase 2-binding protein 1 | -181.5335144 |
| LOC107064290 | E3 ubiquitin-protein ligase RNF8-like | -181.1737813 |
| LOC107064951 | uncharacterized protein LOC107064951 | -180.8385378 |
| LOC107064748 | dephospho-CoA kinase | -180.7341594 |
| LOC107073231 | probable serine/threonine-protein kinase fhkE | -180.709865 |
| LOC107064397 | uncharacterized protein LOC107064397 | -180.6061034 |
| LOC107064711 | puff-specific protein Bx42 | -180.6055498 |
| LOC107067300 | ATP-binding cassette sub-family G member 1 | -180.4853622 |
| LOC107068137 | putative golgin subfamily A member 6-like protein 6 | -180.4319131 |
| LOC107067578 | uncharacterized protein LOC107067578 | -180.3211893 |
| LOC107064287 | sodium/bile acid cotransporter 7-like | -180.2926185 |
| LOC107071616 | histone-lysine N-methyltransferase PRDM9 | -180.2596368 |
| LOC107069886 | uncharacterized protein LOC107069886 | -180.140543 |
| LOC107069497 | jmjC domain-containing protein 4 | -180.1179816 |
| LOC107067401 | uncharacterized protein LOC107067401 | -179.9995416 |
| LOC107069496 | odorant receptor 43a-like | -179.97685 |
| LOC107074488 | putative E3 ubiquitin-protein ligase UBR7 | -179.9707575 |
| LOC107071901 | protein AATF-like | -179.8084566 |
| LOC107069030 | sodium/potassium/calcium exchanger 3-like | -179.75306 |
| LOC107065257 | protein phosphatase 1 regulatory subunit 7 | -179.6809101 |
| LOC107071473 | myb-related protein B | -179.4235144 |
| LOC107070343 | protein tramtrack, beta isoform-like | -179.3731161 |
| LOC107071888 | uncharacterized protein LOC107071888 | -179.3612736 |

| | | |
|---|---|---|
| LOC107067422 | uncharacterized protein LOC107067422 | -179.3457226 |
| LOC107065351 | histone-lysine N-methyltransferase SETMAR-like | -179.3286438 |
| LOC107072668 | putative peptidyl-tRNA hydrolase PTRHD1 | -179.3279859 |
| LOC107064918 | SKI/DACH domain-containing protein 1-like | -179.2753934 |
| LOC107064096 | tubulin beta chain-like | -179.2086725 |
| LOC107068064 | synaptic vesicle glycoprotein 2C-like | -179.1447056 |
| LOC107068498 | putative glycerol kinase 5 | -179.0999364 |
| LOC107070981 | uncharacterized protein LOC107070981 | -179.012316 |
| LOC107071851 | cytochrome b-c1 complex subunit 6, mitochondrial | -178.9984796 |
| LOC107072716 | peptide chain release factor 1-like, mitochondrial | -178.945678 |
| LOC107069618 | putative thiamine transporter SLC35F3 | -178.7833191 |
| LOC107072351 | DNA primase large subunit-like | -178.7351657 |
| LOC107074451 | uncharacterized protein LOC107074451 | -178.678098 |
| LOC107069411 | prefoldin subunit 1 | -178.6655542 |
| LOC107067834 | disheveled-associated activator of morphogenesis 1 | -178.5073891 |
| LOC107064487 | uncharacterized protein LOC107064487 | -178.4941868 |
| LOC107070324 | hexosaminidase D-like | -178.4577289 |
| LOC107068935 | farnesol dehydrogenase-like | -178.426947 |
| LOC107068425 | protein SAAL1 | -178.3048012 |
| LOC107071198 | mitochondrial amidoxime-reducing component 1-like | -178.0226663 |
| LOC107064363 | uncharacterized protein LOC107064363 | -178.009964 |
| LOC107067480 | mediator of RNA polymerase II transcription subunit 11-like | -177.945372 |
| LOC107071559 | probable DNA mismatch repair protein Msh6 | -177.8327701 |
| LOC107072735 | enolase-phosphatase E1-like | -177.8132767 |
| LOC107070241 | exosome complex component RRP45 | -177.8088703 |
| LOC107066447 | diphthine--ammonia ligase | -177.7817124 |
| LOC107068337 | transcription initiation factor TFIID subunit 7 | -177.7118053 |
| LOC107064836 | uncharacterized protein LOC107064836 | -177.6930391 |
| LOC107065152 | zinc finger protein 511 | -177.3733666 |
| LOC107065849 | putative fatty acyl-CoA reductase CG5065 | -177.3225715 |
| LOC107064544 | perilipin-1-like | -176.9609835 |
| LOC107065847 | SPRY domain-containing SOCS box protein 3-like | -176.897961 |
| LOC107064685 | uncharacterized protein LOC107064685 | -176.8551206 |
| LOC107067369 | uncharacterized protein LOC107067369 | -176.8471142 |
| LOC107073479 | protein toll-like | -176.7395428 |
| LOC107070657 | focadhesin | -176.6926423 |
| LOC107067031 | sodium-dependent noradrenaline transporter | -176.3081037 |
| LOC107072219 | 28S ribosomal protein S21, mitochondrial | -176.1534756 |
| LOC107068642 | uncharacterized protein LOC107068642 | -176.0737498 |
| LOC107065981 | protein PFC0760c-like | -175.8201122 |
| LOC107073008 | dynein assembly factor 5, axonemal | -175.818541 |
| LOC107069329 | gastrulation defective protein 1 homolog | -175.6911399 |
| LOC107069896 | protein MMS22-like | -175.6150387 |
| LOC107066300 | uncharacterized protein LOC107066300 | -175.6074682 |
| LOC107065745 | PP2C-like domain-containing protein CG9801 | -175.5624813 |
| LOC107065678 | magnesium-dependent phosphatase 1-like | -175.4758333 |
| LOC107064820 | uncharacterized protein LOC107064820 | -175.4288557 |
| LOC107064800 | chaoptin-like | -175.3878691 |
| LOC107066421 | vacuole membrane protein 1-like | -175.3491456 |
| LOC107070637 | 60S ribosomal protein L10 | -175.3262596 |

| | | |
|---|---|---|
| LOC107073781 | COP9 signalosome complex subunit 5-like | -175.002292 |
| LOC107066315 | peptidyl-prolyl cis-trans isomerase FKBP1A | -174.8976874 |
| LOC107066439 | lysosome-associated membrane glycoprotein 1-like | -174.7749254 |
| LOC107065842 | scavenger receptor class B member 1 | -174.7300573 |
| LOC107064204 | BRCA1-associated RING domain protein 1-like | -174.6104626 |
| LOC107068341 | evolutionarily conserved signaling intermediate in Toll pathw | -174.4701394 |
| LOC107067188 | arylphorin subunit beta-like | -174.4428652 |
| LOC107073294 | uncharacterized protein LOC107073294 | -174.3289268 |
| LOC107071246 | transformation/transcription domain-associated protein | -174.3094539 |
| LOC107074400 | GTP:AMP phosphotransferase AK3, mitochondrial | -174.2999867 |
| LOC107064586 | multidrug resistance-associated protein 7 | -174.2260286 |
| LOC107068890 | adenylate kinase | -174.1887153 |
| LOC107068597 | inositol polyphosphate multikinase-like | -174.1848279 |
| LOC107071380 | syntaxin-5 | -174.1239652 |
| LOC107073065 | protein SMG5 | -174.0410996 |
| LOC107073573 | farnesol dehydrogenase-like | -174.0021509 |
| LOC107067072 | uncharacterized protein LOC107067072 | -173.9404938 |
| LOC107072694 | abhydrolase domain-containing protein 16A | -173.8549111 |
| LOC107069769 | 39S ribosomal protein L55, mitochondrial | -173.750925 |
| LOC107071324 | MATH and LRR domain-containing protein PFE0570w-like | -173.7489278 |
| LOC107067105 | tektin-B1 | -173.6956993 |
| LOC107066612 | transcription cofactor vestigial-like protein 4 | -173.637784 |
| LOC107066916 | CD63 antigen-like | -173.6285579 |
| LOC107070191 | sodium-independent sulfate anion transporter | -173.5365401 |
| LOC107065037 | uncharacterized protein LOC107065037 | -173.5228695 |
| LOC107073412 | uncharacterized protein LOC107073412 | -173.5103689 |
| LOC107072203 | uncharacterized protein LOC107072203 | -173.3906917 |
| LOC107068893 | uncharacterized protein LOC107068893 | -173.3681214 |
| LOC107068000 | aldose reductase-like | -173.2286056 |
| LOC107069002 | presenilins-associated rhomboid-like protein, mitochondrial | -173.2220331 |
| LOC107065230 | tolloid-like protein 1 | -173.1233784 |
| LOC107065225 | V-type proton ATPase subunit d | -173.1063839 |
| LOC107072240 | probable chitinase 3 | -172.8759859 |
| LOC107064774 | amine sulfotransferase-like | -172.8591199 |
| LOC107074422 | serine/threonine-protein kinase PAK mbt | -172.7818863 |
| LOC107064834 | uncharacterized protein LOC107064834 | -172.7441548 |
| LOC107072292 | ectopic P granules protein 5 homolog | -172.7014187 |
| LOC107068163 | protein CDV3 homolog | -172.5935105 |
| LOC107074310 | dynein light chain 1, axonemal-like | -172.5168048 |
| LOC107070359 | uncharacterized protein LOC107070359 | -172.5010358 |
| LOC107074065 | uncharacterized protein LOC107074065 | -172.2997517 |
| LOC107068411 | phospholipase A-2-activating protein | -172.205387 |
| LOC107073117 | uncharacterized protein LOC107073117 | -172.1873847 |
| LOC107071274 | testis-expressed sequence 2 protein | -172.1483452 |
| LOC107072879 | testis-specific serine/threonine-protein kinase 3-like | -172.0970433 |
| LOC107071699 | uncharacterized protein LOC107071699 | -172.0927034 |
| LOC107065543 | uncharacterized protein LOC107065543 | -172.0865127 |
| LOC107063744 | immunoglobulin domain-containing protein oig-1-like | -171.6582958 |
| LOC107064558 | WASH complex subunit CCDC53 | -171.6467888 |
| LOC107068367 | ubiquitin-protein ligase E3C | -171.5830552 |

| | | |
|---|---|---|
| LOC107065548 | repetitive proline-rich cell wall protein 2-like | -171.5508577 |
| LOC107072366 | uncharacterized protein LOC107072366 | -171.5140741 |
| LOC107070252 | transmembrane protein 184B | -171.5075055 |
| LOC107066952 | piggyBac transposable element-derived protein 4-like | -171.3531071 |
| LOC107066621 | uncharacterized protein LOC107066621 | -171.3375781 |
| LOC107069301 | prolactin regulatory element-binding protein | -171.3009985 |
| LOC107069068 | chromatin modification-related protein MEAF6 | -171.2368902 |
| LOC107074604 | solute carrier family 35 member F5 | -171.2103609 |
| LOC107068879 | uncharacterized protein LOC107068879 | -171.1506246 |
| LOC107067232 | N-acetyltransferase 10 | -171.0768631 |
| LOC107069565 | ATP-dependent DNA helicase Q4 | -170.8462171 |
| LOC107072196 | poly(ADP-ribose) glycohydrolase ARH3-like | -170.7752114 |
| LOC107070400 | uncharacterized protein LOC107070400 | -170.7714387 |
| LOC107073046 | uncharacterized protein LOC107073046 | -170.5746435 |
| LOC107073710 | vascular endothelial growth factor receptor 2-like | -170.5453596 |
| LOC107066163 | serine/threonine-protein kinase SMG1 | -170.4689147 |
| LOC107066436 | ras-related protein Rab-14 | -170.378414 |
| LOC107070580 | uncharacterized protein LOC107070580 | -170.3774193 |
| LOC107068949 | farnesol dehydrogenase-like | -170.3704689 |
| LOC107065765 | condensin complex subunit 3-like | -170.3232488 |
| LOC107067740 | isovaleryl-CoA dehydrogenase, mitochondrial | -170.2646745 |
| LOC107066673 | uncharacterized protein LOC107066673 | -170.2634747 |
| LOC107070549 | persulfide dioxygenase ETHE1, mitochondrial | -170.233877 |
| LOC107068176 | biogenesis of lysosome-related organelles complex 1 subunit | -170.1252242 |
| LOC107069834 | epoxide hydrolase 4-like | -169.9489185 |
| LOC107074613 | NAD kinase-like | -169.8889418 |
| LOC107074253 | uncharacterized protein LOC107074253 | -169.7984318 |
| LOC107063937 | DNA-directed RNA polymerase III subunit RPC7-like | -169.7745253 |
| LOC107064540 | uncharacterized protein LOC107064540 | -169.6644465 |
| LOC107067049 | RING finger protein 10 | -169.658914 |
| LOC107072843 | uncharacterized protein LOC107072843 | -169.6210416 |
| LOC107065327 | uncharacterized aarF domain-containing protein kinase 2-like | -169.5103707 |
| LOC107071335 | exocyst complex component 5 | -169.349045 |
| LOC107064720 | protein suppressor of forked | -169.3167746 |
| LOC107067207 | BTB/POZ domain-containing protein KCTD1 | -169.2103919 |
| LOC107067508 | uncharacterized protein LOC107067508 | -169.1343665 |
| LOC107074353 | enoyl-CoA delta isomerase 2, mitochondrial | -168.9134335 |
| LOC107067320 | N-alpha-acetyltransferase 40 | -168.9050618 |
| LOC107068319 | homeobox protein 6-like | -168.785719 |
| LOC107068921 | M-phase inducer phosphatase | -168.7035793 |
| LOC107068406 | inositol-trisphosphate 3-kinase A | -168.6463748 |
| LOC107065761 | histone acetyltransferase KAT2A | -168.5852901 |
| LOC107067813 | tetratricopeptide repeat protein 27 | -168.4144915 |
| LOC107070137 | dnaJ homolog subfamily C member 10-like | -168.3401044 |
| LOC107068686 | spectrin beta chain, non-erythrocytic 5 | -168.2872538 |
| LOC107065661 | uncharacterized protein LOC107065661 | -168.1580698 |
| LOC107072377 | anaphase-promoting complex subunit cdh1-like | -168.1454446 |
| LOC107066562 | uncharacterized protein LOC107066562 | -168.1247553 |
| LOC107067493 | uncharacterized protein LOC107067493 | -168.0848129 |
| LOC107067741 | uncharacterized protein LOC107067741 | -168.0499393 |

| | | |
|---|---|---|
| LOC107067989 | inositol polyphosphate 5-phosphatase K-like | -168.0350496 |
| LOC107068920 | uncharacterized protein LOC107068920 | -168.0154079 |
| LOC107069415 | uncharacterized protein LOC107069415 | -167.88285 |
| LOC107066024 | histone RNA hairpin-binding protein | -167.6699186 |
| LOC107068793 | uncharacterized protein LOC107068793 | -167.6474144 |
| LOC107074520 | G8 domain-containing protein DDB_G0286311 | -167.6405948 |
| LOC107072798 | protein inturned | -167.5686095 |
| LOC107074186 | kinesin-like protein KIF3A | -167.4940134 |
| LOC107066352 | nesprin-1 | -167.4738616 |
| LOC107065020 | zinc finger protein 678 | -167.2196411 |
| LOC107065854 | uncharacterized protein LOC107065854 | -167.1693274 |
| LOC107066183 | DNA topoisomerase 2 | -167.1672662 |
| LOC107065917 | monocyte to macrophage differentiation factor 2 | -167.0590384 |
| LOC107064211 | DNA-3-methyladenine glycosylase-like | -166.9676965 |
| LOC107073725 | uncharacterized protein LOC107073725 | -166.8884675 |
| LOC107065820 | O-glucosyltransferase rumi homolog | -166.8802772 |
| LOC107065610 | GTP-binding protein 10 homolog | -166.8411972 |
| LOC107065798 | uncharacterized protein LOC107065798 | -166.7385266 |
| LOC107072172 | ornithine decarboxylase antizyme 1 | -166.7109508 |
| LOC107067594 | peptide deformylase, mitochondrial-like | -166.5919627 |
| LOC107067870 | protein NPC2 homolog | -166.4707801 |
| LOC107067341 | unconventional myosin-XV | -166.4558942 |
| LOC107070724 | gem-associated protein 5 | -166.4079881 |
| LOC107074425 | transmembrane protein 35 | -166.3978295 |
| LOC107070037 | uncharacterized protein LOC107070037 | -166.281548 |
| LOC107067489 | uncharacterized protein LOC107067489 | -166.2118689 |
| LOC107074597 | actin-binding protein IPP | -166.1026272 |
| LOC107064161 | NADH dehydrogenase [ubiquinone | -165.9439402 |
| LOC107073260 | uncharacterized protein LOC107073260 | -165.8528866 |
| LOC107067788 | uncharacterized protein LOC107067788 | -165.7933914 |
| LOC107068598 | transmembrane and TPR repeat-containing protein CG4050-l | -165.6722117 |
| LOC107066712 | protein slowmo | -165.669369 |
| LOC107068684 | pancreatic lipase-related protein 2-like | -165.5974723 |
| LOC107064590 | cdc42 homolog | -165.5751502 |
| LOC107072491 | EF-hand calcium-binding domain-containing protein 1 | -165.5012087 |
| LOC107070166 | cytochrome P450 4g15-like | -165.209839 |
| LOC107066955 | chromosome transmission fidelity protein 18 homolog | -165.2001501 |
| LOC107070559 | uncharacterized protein LOC107070559 | -165.0865706 |
| LOC107074411 | solute carrier family 35 member G1-like | -165.0406677 |
| LOC107072784 | PI-PLC X domain-containing protein 1-like | -164.9689863 |
| LOC107064292 | integrator complex subunit 8 | -164.8988311 |
| LOC107072299 | uncharacterized protein LOC107072299 | -164.7604466 |
| LOC107068663 | ATP-dependent RNA helicase DDX54 | -164.709919 |
| LOC107067412 | uncharacterized protein LOC107067412 | -164.6723184 |
| LOC107071431 | dynein light chain Tctex-type 1 | -164.5432592 |
| LOC107069909 | odorant receptor 13a-like | -164.5293005 |
| LOC107065838 | actin-binding protein anillin-like | -164.4376994 |
| LOC107070881 | tektin-3-like | -164.3851059 |
| LOC107072436 | bromodomain-containing protein DDB_G0270170-like | -164.3700792 |
| LOC107064538 | disks large homolog 5 | -164.3682675 |

| | | |
|---|---|---|
| LOC107066260 | protein mago nashi homolog | -164.2889986 |
| LOC107068698 | mitochondrial fission 1 protein-like | -163.8466065 |
| LOC107073384 | mitochondrial sodium/hydrogen exchanger 9B2-like | -163.8247899 |
| LOC107069722 | arrestin homolog | -163.6513407 |
| LOC107068779 | lysophospholipid acyltransferase 7 | -163.3926423 |
| LOC107064389 | transmembrane protease serine 9-like | -163.3518541 |
| LOC107070959 | methyltransferase-like protein 7B | -163.2938859 |
| LOC107074427 | uncharacterized protein LOC107074427 | -163.2520875 |
| LOC107067904 | otopetrin-3-like | -163.2042485 |
| LOC107065419 | COP9 signalosome complex subunit 1 | -163.1467831 |
| LOC107072670 | uncharacterized protein LOC107072670 | -162.978183 |
| LOC107071776 | intraflagellar transport protein 122 homolog | -162.9540905 |
| LOC107063778 | troponin C-like | -162.9218336 |
| LOC107068249 | dnaJ homolog subfamily C member 16 | -162.8089523 |
| LOC107069161 | uncharacterized protein LOC107069161 | -162.7341723 |
| LOC107072161 | uncharacterized protein LOC107072161 | -162.6421112 |
| LOC107063741 | procollagen-lysine,2-oxoglutarate 5-dioxygenase 3 | -162.5486653 |
| LOC107067065 | rab proteins geranylgeranyltransferase component A 1 | -162.513389 |
| LOC107069122 | transmembrane 9 superfamily member 2 | -162.3673213 |
| LOC107064868 | protein FAM151B | -162.3358854 |
| LOC107067206 | uncharacterized protein LOC107067206 | -162.2863723 |
| LOC107071330 | protein GDAP2 homolog | -162.0593514 |
| LOC107073246 | uncharacterized protein LOC107073246 | -161.7929343 |
| LOC107073488 | hydroxylysine kinase | -161.7415551 |
| LOC107072220 | uncharacterized protein LOC107072220 | -161.6974616 |
| LOC107074028 | putative leucine-rich repeat-containing protein DDB_G02905 | -161.6900835 |
| LOC107067991 | bromodomain-containing protein 8 | -161.6836043 |
| LOC107065935 | mucin-17 | -161.4790075 |
| LOC107065647 | tRNA (adenine(58)-N(1))-methyltransferase non-catalytic sub | -161.4339516 |
| LOC107064493 | dedicator of cytokinesis protein 9 | -161.3857073 |
| LOC107071745 | ELMO domain-containing protein F | -161.3529035 |
| LOC107070054 | molybdenum cofactor biosynthesis protein 1 | -161.0669159 |
| LOC107070934 | kinesin-like protein KIF16B | -160.9868397 |
| LOC107073671 | girdin-like | -160.8580147 |
| LOC107066047 | hemicentin-1-like | -160.8392167 |
| LOC107072652 | excitatory amino acid transporter 1-like | -160.7374164 |
| LOC107074386 | uncharacterized protein LOC107074386 | -160.6555342 |
| LOC107072519 | venom allergen 5-like | -160.5068108 |
| LOC107067552 | beta-1,3-galactosyltransferase 1-like | -160.4245568 |
| LOC107067388 | cytochrome b5-like | -160.2806521 |
| LOC107068947 | WD repeat-containing protein 35 | -160.2596866 |
| LOC107063696 | uncharacterized protein PF11_0213 | -160.0392312 |
| LOC107073841 | AP-1 complex subunit beta-1 | -159.5500308 |
| LOC107064468 | uncharacterized protein LOC107064468 | -159.453151 |
| LOC107067786 | DNA primase large subunit-like | -159.4445162 |
| LOC107070039 | uncharacterized protein LOC107070039 | -159.4019052 |
| LOC107065393 | sideroflexin-1-like | -159.389118 |
| LOC107065844 | metabotropic glutamate receptor-like | -159.2503754 |
| LOC107070187 | excitatory amino acid transporter 3-like | -159.1184059 |
| LOC107073780 | uncharacterized protein LOC107073780 | -159.0512198 |

| | | |
|---|---|---|
| LOC107071614 | inactive dipeptidyl peptidase 10 | -159.0477488 |
| LOC107065231 | uncharacterized protein LOC107065231 | -158.8797866 |
| LOC107064770 | uncharacterized protein LOC107064770 | -158.8066463 |
| LOC107064352 | uncharacterized protein LOC107064352 | -158.6907908 |
| LOC107072312 | cadherin-86C | -158.6737572 |
| LOC107066394 | uncharacterized protein LOC107066394 | -158.6543929 |
| LOC107064816 | uncharacterized protein LOC107064816 | -158.6250558 |
| LOC107063984 | uncharacterized protein LOC107063984 | -158.4574625 |
| LOC107069419 | uncharacterized protein LOC107069419 | -158.4046479 |
| LOC107073037 | protein pigeon | -158.3202779 |
| LOC107065875 | zinc finger protein 2 homolog | -158.3145912 |
| LOC107070877 | uncharacterized protein LOC107070877 | -158.2415765 |
| Trnak-uuu | Trnak-uuu | -158.233434 |
| LOC107073175 | uncharacterized protein LOC107073175 | -158.0922158 |
| LOC107067131 | calnexin-like | -158.0681949 |
| LOC107069370 | aromatic-L-amino-acid decarboxylase-like | -157.9690027 |
| LOC107071381 | transport and Golgi organization protein 1 | -157.9409791 |
| LOC107064160 | protein FAM192A | -157.9226425 |
| LOC107070371 | bicaudal D-related protein homolog | -157.8756002 |
| LOC107066190 | NADH dehydrogenase [ubiquinone | -157.8468467 |
| LOC107066221 | serine/threonine-protein phosphatase 6 regulatory subunit 3 | -157.6384644 |
| LOC107064575 | uncharacterized protein LOC107064575 | -157.5830863 |
| LOC107071507 | gamma-aminobutyric acid type B receptor subunit 1 | -157.5665996 |
| LOC107067653 | short-chain specific acyl-CoA dehydrogenase, mitochondrial | -157.5023451 |
| LOC107063661 | uncharacterized protein LOC107063661 | -157.4956259 |
| LOC107065652 | acylphosphatase-1-like | -157.4299417 |
| LOC107064615 | NHP2-like protein 1 homolog | -157.4166443 |
| LOC107070512 | G-box-binding factor-like | -157.4041478 |
| LOC107067756 | glycine-rich cell wall structural protein-like | -157.157651 |
| LOC107070136 | mucin-5AC-like | -156.9123187 |
| LOC107067780 | uncharacterized protein LOC107067780 | -156.9047828 |
| LOC107071308 | myb-like protein X | -156.676008 |
| LOC107067305 | chaoptin-like | -156.6510785 |
| LOC107070239 | E3 ubiquitin-protein ligase hyd | -156.5159838 |
| LOC107073786 | uncharacterized protein LOC107073786 | -156.5107459 |
| LOC107067979 | high affinity cAMP-specific and IBMX-insensitive 3',5'-cyclic p | -156.3616465 |
| LOC107072030 | 3-hydroxyacyl-CoA dehydrogenase type-2-like | -156.3437816 |
| LOC107074587 | uncharacterized protein LOC107074587 | -156.3208783 |
| LOC107069977 | uncharacterized protein LOC107069977 | -156.2675169 |
| LOC107068381 | protein-lysine N-methyltransferase n6amt2 | -156.2370359 |
| LOC107065739 | probable serine/threonine-protein kinase roco9 | -156.2282404 |
| LOC107063714 | uncharacterized protein LOC107063714 | -156.2085014 |
| LOC107067306 | uncharacterized protein LOC107067306 | -156.1920788 |
| LOC107068004 | uncharacterized protein LOC107068004 | -156.0139335 |
| LOC107074254 | centrosomal protein of 112 kDa-like | -155.6551584 |
| LOC107074393 | 39S ribosomal protein L33, mitochondrial | -155.6149405 |
| LOC107074659 | beta-TrCP | -155.5829815 |
| LOC107064745 | tryptophan 5-hydroxylase 1 | -155.5189045 |
| LOC107069309 | E3 ubiquitin-protein ligase MYCBP2 | -155.5019461 |
| LOC107065210 | DNA repair protein RAD51 homolog 4-like | -155.3230787 |

| | | |
|---|---|---|
| LOC107072562 | uncharacterized protein LOC107072562 | -155.2603745 |
| LOC107073731 | uncharacterized protein LOC107073731 | -155.2014628 |
| LOC107067523 | POU domain, class 3, transcription factor 2-like | -155.1967547 |
| LOC107070149 | mismatch repair endonuclease PMS2 | -154.9641894 |
| LOC107073614 | transmembrane protein 183B-like | -154.8441788 |
| LOC107066165 | ATP synthase subunit d, mitochondrial-like | -154.7736046 |
| LOC107064230 | glutamate receptor U1-like | -154.7547121 |
| LOC107070070 | solute carrier organic anion transporter family member 4A1 | -154.7188586 |
| LOC107067645 | malate dehydrogenase, cytoplasmic-like | -154.597909 |
| LOC107071518 | helicase POLQ-like | -154.4959551 |
| LOC107067267 | probable Bax inhibitor 1 | -154.4549915 |
| LOC107071532 | ADAMTS-like protein 4 | -154.3534819 |
| LOC107065700 | leucine-rich repeat and death domain-containing protein 1-li | -154.1556028 |
| LOC107068546 | uncharacterized protein LOC107068546 | -154.0841365 |
| LOC107068922 | uncharacterized protein LOC107068922 | -153.9212739 |
| LOC107068452 | glucose dehydrogenase [FAD, quinone | -153.7906791 |
| LOC107071777 | uncharacterized protein LOC107071777 | -153.6560478 |
| LOC107071685 | twisted gastrulation protein homolog 1-A-like | -153.507014 |
| LOC107073938 | glycerol kinase | -153.3167345 |
| LOC107069434 | NFU1 iron-sulfur cluster scaffold homolog, mitochondrial-like | -153.2916091 |
| LOC107072414 | microtubule-associated proteins 1A/1B light chain 3B-like | -153.2328051 |
| LOC107071071 | uncharacterized protein LOC107071071 | -153.2147655 |
| LOC107071217 | aquaporin AQPcic-like | -153.1715064 |
| LOC107073150 | 26S protease regulatory subunit 8 | -153.1541572 |
| LOC107064024 | kinesin-like protein KIF18A | -152.9883138 |
| LOC107073854 | uncharacterized protein LOC107073854 | -152.9519336 |
| LOC107070165 | gastrin/cholecystokinin type B receptor-like | -152.9261721 |
| LOC107070551 | putative mediator of RNA polymerase II transcription subunit | -152.9193137 |
| LOC107070733 | E3 ubiquitin-protein ligase LRSAM1-like | -152.9094768 |
| LOC107074508 | GDP-D-glucose phosphorylase 1 | -152.814595 |
| LOC107072103 | uncharacterized protein LOC107072103 | -152.7756908 |
| LOC107072690 | PHD finger protein rhinoceros | -152.5155254 |
| LOC107069381 | uncharacterized protein LOC107069381 | -152.4526797 |
| LOC107069259 | tetratricopeptide repeat protein 8 | -152.0794398 |
| LOC107074598 | uncharacterized protein LOC107074598 | -152.0654876 |
| LOC107065046 | lysine-specific demethylase lid | -151.9632457 |
| LOC107067221 | uncharacterized protein LOC107067221 | -151.6545377 |
| LOC107067903 | tektin-3-like | -151.5828731 |
| LOC107072961 | uncharacterized protein LOC107072961 | -151.5185441 |
| LOC107065822 | pseudouridine-5'-phosphatase | -151.4474545 |
| LOC107066464 | diphthine methyl ester synthase | -151.2985794 |
| LOC107071306 | integral membrane protein GPR155 | -151.2922553 |
| LOC107068887 | guanine nucleotide exchange factor DBS-like | -151.2383803 |
| LOC107072418 | ubiquitin-conjugating enzyme E2 H | -151.2036118 |
| LOC107064501 | oxysterol-binding protein 1 | -151.0752311 |
| LOC107070525 | caprin homolog | -150.9841856 |
| LOC107068790 | mitochondrial import receptor subunit TOM40 homolog 1-lik | -150.5580225 |
| LOC107072245 | coiled-coil domain-containing protein 22 homolog | -150.457873 |
| LOC107065279 | venom carboxylesterase-6-like | -150.0801329 |
| LOC107074394 | uncharacterized protein LOC107074394 | -149.9822922 |

| | | |
|---|---|---|
| LOC107063825 | uncharacterized protein LOC107063825 | -149.7720343 |
| LOC107068623 | uncharacterized protein LOC107068623 | -149.7190432 |
| LOC107064154 | MICOS complex subunit mic25-a | -149.7156448 |
| LOC107067889 | uncharacterized protein LOC107067889 | -149.5079594 |
| LOC107065886 | ubiquitin-like modifier-activating enzyme 1 | -149.4089467 |
| LOC107065357 | translationally-controlled tumor protein homolog | -149.3219862 |
| LOC107065646 | long-chain fatty acid transport protein 4-like | -149.0872576 |
| LOC107073797 | uncharacterized protein LOC107073797 | -149.0836179 |
| LOC107070466 | protein shisa-5-like | -148.5225566 |
| LOC107072901 | uncharacterized protein LOC107072901 | -148.4789739 |
| LOC107073804 | ras-related protein Rab-2 | -148.3162415 |
| LOC107071189 | uncharacterized protein LOC107071189 | -148.1514233 |
| LOC107063904 | neural cell adhesion molecule 2 | -148.0974943 |
| LOC107069625 | uncharacterized protein LOC107069625 | -147.9934679 |
| LOC107068982 | syntaxin-1B-like | -147.9150473 |
| LOC107072780 | TPPP family protein CG45057-like | -147.8439576 |
| LOC107070099 | uncharacterized protein LOC107070099 | -147.8398892 |
| LOC107069530 | sodium- and chloride-dependent GABA transporter 1-like | -147.6923906 |
| LOC107068689 | ATP synthase subunit gamma, mitochondrial | -146.8135499 |
| LOC107071174 | lisH domain-containing protein ARMC9-like | -146.7797946 |
| LOC107065329 | mitochondrial pyruvate carrier 1 | -146.5302399 |
| LOC107065819 | matrix metalloproteinase-14 | -145.7530669 |
| LOC107074150 | uncharacterized protein LOC107074150 | -145.6525255 |
| LOC107072932 | ATP-dependent DNA helicase PIF1 | -145.43963 |
| LOC107067235 | uncharacterized protein LOC107067235 | -144.7931856 |
| LOC107070412 | cationic amino acid transporter 3 | -144.5761145 |
| LOC107068528 | uncharacterized protein LOC107068528 | -144.4893201 |
| LOC107074641 | NACHT domain- and WD repeat-containing protein 1 | -144.1513995 |
| LOC107067599 | uncharacterized protein LOC107067599 | -143.7766812 |
| LOC107068355 | aryl-hydrocarbon-interacting protein-like 1 | -143.7120404 |
| LOC107063814 | LMBR1 domain-containing protein 2 homolog | -143.6904321 |
| LOC107074682 | sodium- and chloride-dependent GABA transporter 1-like | -143.5421798 |
| LOC107070570 | uncharacterized protein LOC107070570 | -143.3562596 |
| LOC107067849 | collagen alpha-5(IV) chain-like | -143.2108781 |
| LOC107064624 | vitamin K-dependent gamma-carboxylase | -143.2083968 |
| LOC107068579 | signal recognition particle receptor subunit beta | -143.0061406 |
| LOC107069345 | uncharacterized protein LOC107069345 | -142.9638268 |
| LOC107072258 | guanine nucleotide-binding protein subunit gamma-e | -142.9055575 |
| LOC107070907 | cullin-associated NEDD8-dissociated protein 1 | -142.753568 |
| LOC107069617 | glutathione S-transferase 1, isoform C | -142.3223165 |
| LOC107072787 | coiled-coil domain-containing protein 50 | -141.8927252 |
| LOC107069462 | rRNA methyltransferase 2, mitochondrial | -141.8015675 |
| LOC107065694 | protein lifeguard 1-like | -141.3702788 |
| LOC107069334 | uncharacterized protein LOC107069334 | -140.6689845 |
| LOC107066259 | succinyl-CoA ligase [ADP/GDP-forming | -140.1845903 |
| LOC107070104 | uncharacterized protein LOC107070104 | -139.8552171 |
| LOC107073977 | uncharacterized protein LOC107073977 | -138.9942911 |
| LOC107071435 | uncharacterized protein LOC107071435 | -138.9788244 |
| LOC107068726 | 26S proteasome non-ATPase regulatory subunit 6 | -138.201785 |
| LOC107070761 | probable RNA helicase armi | -137.8267407 |

| | | |
|---|---|---|
| LOC107067915 | odorant receptor 4-like | -135.1785456 |
| LOC107072488 | uncharacterized protein LOC107072488 | -134.9950072 |
| LOC107065170 | NEDD4 family-interacting protein 1-like | -134.7523828 |
| LOC107070815 | calreticulin | -133.9630951 |
| LOC107072065 | endoplasmic reticulum resident protein 29 | -130.5945717 |
| LOC107070992 | embryonic stem cell-specific 5-hydroxymethylcytosine-bindir | -128.8242497 |
| LOC107064768 | uncharacterized protein LOC107064768 | -126.1957905 |
| LOC107068654 | enolase | -115.9318156 |
| LOC107067059 | heat shock 70 kDa protein cognate 4 | -102.4769607 |
| LOC107066237 | ras-like GTP-binding protein Rho1 | -95.48844467 |
| LOC107067352 | ras-related protein Rab-1A | -15.3091519 |
| LOC107067109 | ubiquitin-conjugating enzyme E2-17 kDa | 19.78899762 |
| LOC107074330 | protein CNPPD1 | 79.23185044 |
| LOC107071334 | eukaryotic initiation factor 4A-I | 83.51627388 |
| LOC107068201 | lissencephaly-1 homolog | 89.65638784 |
| LOC107072036 | uncharacterized protein LOC107072036 | 93.17853118 |
| LOC107073870 | peptidyl-prolyl cis-trans isomerase FKBP8 | 94.79574927 |
| LOC107072241 | homeobox protein invected-like | 99.55977789 |
| LOC107067395 | zinc transporter 9 | 99.69031428 |
| LOC107073189 | growth hormone-inducible transmembrane protein-like | 101.471794 |
| LOC107070362 | GRB2-associated-binding protein 2 | 101.7015476 |
| LOC107065998 | AMP deaminase 2-like | 103.5563484 |
| LOC107073629 | seipin | 108.2006011 |
| LOC107074002 | uncharacterized protein DDB_G0285917-like | 109.5070618 |
| LOC107069510 | vang-like protein 2 | 109.8591461 |
| LOC107067984 | uncharacterized protein LOC107067984 | 110.6914978 |
| LOC107072553 | transmembrane protein 192 | 111.7936662 |
| LOC107070721 | peroxidase-like | 113.2269915 |
| LOC107070615 | fructose-bisphosphate aldolase-like | 113.395822 |
| LOC107068389 | T-box-containing protein TBXT | 113.868047 |
| LOC107070479 | uncharacterized protein LOC107070479 | 113.8719866 |
| LOC107067132 | elongation factor 1-alpha | 114.0432547 |
| LOC107070109 | huntingtin-interacting protein K | 114.0701572 |
| LOC107071277 | ATP-dependent RNA helicase dbp2-like | 114.6872365 |
| LOC107069699 | nuclear pore complex protein Nup214 | 114.8522058 |
| LOC107073809 | spermatogenesis-associated protein 20 | 114.8673368 |
| LOC107066876 | dual specificity mitogen-activated protein kinase kinase 7-like | 114.9437729 |
| LOC107064960 | uncharacterized protein LOC107064960 | 115.3879244 |
| LOC107074484 | uncharacterized protein LOC107074484 | 115.8532518 |
| LOC107074677 | U4/U6.U5 tri-snRNP-associated protein 2 | 116.2574629 |
| LOC107074340 | E3 ubiquitin-protein ligase KCMF1-like | 117.0136475 |
| LOC107071452 | uncharacterized protein LOC107071452 | 117.2537292 |
| LOC107070384 | antizyme inhibitor 2-like | 117.4298445 |
| LOC107065623 | DNA-directed RNA polymerases I, II, and III subunit RPABC3 | 118.0354597 |
| LOC107066112 | mannosyl-oligosaccharide 1,2-alpha-mannosidase IA | 118.3411343 |
| LOC107063683 | 2-phosphoxylose phosphatase 1 | 118.8726934 |
| LOC107070450 | alpha-protein kinase 1-like | 118.9312416 |
| LOC107069550 | protein YIPF5 | 119.5468003 |
| LOC107066115 | NCK-interacting protein with SH3 domain | 119.8433644 |
| LOC107066521 | elongation factor Tu GTP-binding domain-containing protein | 120.1769665 |

| | | |
|---|---|---|
| LOC107071365 | zinc finger protein GLI1 | 120.3467089 |
| LOC107064962 | host cell factor 1 | 121.3730537 |
| LOC107070922 | uncharacterized protein LOC107070922 | 121.3732826 |
| LOC107064658 | sn1-specific diacylglycerol lipase alpha | 121.4624664 |
| LOC107069455 | general transcription factor 3C polypeptide 5 | 122.213114 |
| LOC107067841 | E3 ubiquitin-protein ligase MIB2 | 122.2408632 |
| LOC107071891 | vacuolar fusion protein MON1 homolog A | 122.324795 |
| LOC107064434 | protein BCL9 homolog | 122.3696751 |
| LOC107067178 | AN1-type zinc finger protein 2A-like | 122.4070989 |
| LOC107065994 | S-methyl-5'-thioadenosine phosphorylase-like | 122.7049174 |
| LOC107071985 | putative ATP-dependent RNA helicase DHX33 | 122.764294 |
| LOC107071088 | RING finger and SPRY domain-containing protein 1-like | 122.8125062 |
| LOC107064038 | mucin-2-like | 122.8630843 |
| LOC107068749 | fragile X mental retardation syndrome-related protein 1 | 122.8874208 |
| LOC107063941 | acyl-CoA Delta(11) desaturase | 123.5946497 |
| LOC107069715 | nuclear pore complex protein Nup93-like | 123.7902622 |
| LOC107069830 | uncharacterized protein LOC107069830 | 124.2570837 |
| LOC107072935 | palmitoyltransferase ZDHHC6 | 124.3310922 |
| LOC107063638 | cilia- and flagella-associated protein 36 | 124.7733482 |
| LOC107070568 | endoplasmic reticulum-Golgi intermediate compartment pro | 125.0054723 |
| LOC107074582 | uncharacterized protein LOC107074582 | 125.0321487 |
| LOC107064177 | 39S ribosomal protein L13, mitochondrial | 125.0716655 |
| LOC107071127 | 26S protease regulatory subunit 10B | 125.2008451 |
| LOC107071934 | sodium-dependent neutral amino acid transporter B(0)AT3 | 125.3739192 |
| LOC107072684 | neurocalcin-delta B-like | 125.4836869 |
| LOC107071829 | homeobox protein extradenticle-like | 125.7755174 |
| LOC107067718 | beta-1,3-glucan-binding protein-like | 126.2349977 |
| LOC107072943 | uncharacterized protein LOC107072943 | 126.2937262 |
| LOC107072119 | putative uncharacterized protein DDB_G0282133 | 126.5833285 |
| LOC107071528 | uncharacterized threonine-rich GPI-anchored glycoprotein PJ | 126.6466974 |
| LOC107072598 | protein catecholamines up | 126.9195449 |
| LOC107072075 | glycosyltransferase 25 family member-like | 127.0197429 |
| LOC107067852 | RNA-binding protein 4-like | 127.4520098 |
| LOC107074419 | uncharacterized protein LOC107074419 | 127.4667728 |
| LOC107068342 | ubiquitin-conjugating enzyme E2 R2 | 127.7665628 |
| LOC107071307 | myeloid leukemia factor | 128.1035653 |
| LOC107069869 | dual specificity protein phosphatase CDC14A-like | 128.9371678 |
| LOC107066003 | UPF0454 protein C12orf49 homolog | 129.1919451 |
| LOC107073747 | uncharacterized protein LOC107073747 | 129.3675999 |
| LOC107074657 | guanylate kinase | 129.4018161 |
| LOC107063624 | serine protease easter-like | 129.4384328 |
| LOC107072927 | geminin-like | 129.7287384 |
| LOC107073393 | fidgetin-like protein 1 | 129.8676449 |
| LOC107068292 | mitochondrial import inner membrane translocase subunit T | 129.9170226 |
| LOC107063759 | copper-transporting ATPase 1 | 129.9613986 |
| LOC107073398 | uncharacterized protein LOC107073398 | 130.0324624 |
| LOC107070114 | pyroglutamyl-peptidase 1 | 130.0733028 |
| LOC107066915 | ubiquitin-conjugating enzyme E2-24 kDa | 130.1435238 |
| LOC107069586 | uncharacterized protein LOC107069586 | 130.2678825 |
| LOC107074459 | uncharacterized protein LOC107074459 | 130.2759108 |

| | | |
|---|---|---|
| LOC107071234 | protein LSM12 homolog | 130.4914666 |
| LOC107072031 | uncharacterized protein LOC107072031 | 130.5019645 |
| LOC107066728 | cyclin-H | 130.6534448 |
| LOC107066153 | SUZ domain-containing protein 1 | 130.8432427 |
| LOC107072722 | 5-demethoxyubiquinone hydroxylase, mitochondrial | 130.9341846 |
| LOC107073036 | nucleoporin NUP53 | 130.9742263 |
| LOC107073131 | peptidyl-prolyl cis-trans isomerase NIMA-interacting 1-like | 130.9920735 |
| LOC107071327 | putative leucine-rich repeat-containing protein DDB_G02905 | 131.1055744 |
| LOC107072504 | uncharacterized protein LOC107072504 | 131.1458174 |
| LOC107066093 | ubiquitin-conjugating enzyme E2 Q2 | 131.232796 |
| LOC107068310 | serine-rich adhesin for platelets-like | 131.2367707 |
| LOC107069399 | WAS/WASL-interacting protein family member 1-like | 131.324064 |
| LOC107072449 | uncharacterized protein LOC107072449 | 131.434124 |
| LOC107072074 | glycosyltransferase 25 family member-like | 131.6948055 |
| LOC107064267 | pre-mRNA-splicing factor Syf2 | 132.0237433 |
| LOC107071531 | methylosome subunit pICln | 132.2380996 |
| LOC107069546 | uncharacterized protein LOC107069546 | 132.3419823 |
| LOC107069958 | odorant receptor 13a-like | 133.1122144 |
| LOC107067282 | homeobox protein 9-like | 133.4797268 |
| LOC107068312 | glutamate receptor-interacting protein 1 | 133.7430896 |
| LOC107064074 | guanylate cyclase 32E-like | 133.7576416 |
| LOC107066553 | probable tRNA N6-adenosine threonylcarbamoyltransferase | 133.7737218 |
| LOC107068945 | AP-3 complex subunit mu-1 | 133.8807711 |
| LOC107072830 | solute carrier organic anion transporter family member 2A1- | 134.0058411 |
| LOC107064596 | apoptosis-resistant E3 ubiquitin protein ligase 1 | 134.044802 |
| LOC107066331 | optic atrophy 3 protein homolog | 134.0781169 |
| LOC107066940 | xaa-Pro aminopeptidase 1-like | 134.1097894 |
| LOC107069950 | ubiquitin-associated domain-containing protein 1 | 134.2070048 |
| LOC107068537 | probable WRKY transcription factor protein 1 | 134.3169551 |
| LOC107063821 | E3 ubiquitin-protein ligase Hakai | 134.3554666 |
| LOC107070269 | acyl-CoA:lysophosphatidylglycerol acyltransferase 1-like | 134.3669325 |
| LOC107064386 | protein dpy-30 homolog | 134.4206908 |
| LOC107069186 | CTP synthase | 135.0133804 |
| LOC107073979 | 40S ribosomal protein S6-like | 135.0538989 |
| LOC107071271 | uncharacterized protein LOC107071271 | 135.0769837 |
| LOC107073334 | uncharacterized protein LOC107073334 | 135.0949402 |
| LOC107072097 | bromodomain-containing protein 2-like | 135.2309072 |
| LOC107066592 | interferon regulatory factor 2-binding protein-like B | 135.3942482 |
| LOC107068760 | homeobox protein Hox-B1a-like | 135.6064348 |
| LOC107065518 | headcase protein | 136.0299528 |
| LOC107071700 | serine-rich adhesin for platelets | 136.0377818 |
| LOC107073533 | uncharacterized protein LOC107073533 | 136.0509244 |
| LOC107067683 | cyclic AMP-responsive element-binding protein 1-like | 136.1277598 |
| LOC107072088 | DNA excision repair protein ERCC-1 | 136.1497397 |
| LOC107070311 | uncharacterized protein LOC107070311 | 136.4087191 |
| LOC107073336 | guanine nucleotide-binding protein subunit alpha homolog | 136.5615653 |
| LOC107073461 | LOW QUALITY PROTEIN: craniofacial development protein 2- | 136.5762111 |
| LOC107071758 | putative sodium-coupled neutral amino acid transporter 7 | 136.6957222 |
| LOC107073053 | uncharacterized protein LOC107073053 | 136.7588592 |
| LOC107068405 | ATPase ASNA1 homolog | 136.8076429 |

| | | |
|---|---|---|
| LOC107068030 | cell wall protein DAN4 | 136.9438013 |
| LOC107074074 | 40S ribosomal protein S2 | 136.9804326 |
| LOC107073311 | adenylate cyclase type 2 | 137.0281239 |
| LOC107066946 | uncharacterized protein LOC107066946 | 137.0782485 |
| LOC107073164 | intraflagellar transport protein 80 homolog | 137.1144012 |
| LOC107067303 | TAR DNA-binding protein 43-like | 137.2432806 |
| LOC107065156 | synaptic vesicle membrane protein VAT-1 homolog-like | 137.2955049 |
| LOC107069236 | homeobox protein PKNOX2-like | 137.4723822 |
| LOC107065027 | SH3 domain-binding glutamic acid-rich protein homolog | 137.5070593 |
| LOC107066358 | zinc finger protein OZF-like | 137.5419117 |
| LOC107064860 | uncharacterized protein DDB_G0283697-like | 137.6638117 |
| LOC107070282 | histone deacetylase complex subunit SAP130 | 137.9141772 |
| LOC107065314 | zygotic gap protein knirps-like | 138.256303 |
| LOC107067537 | microtubule-associated protein futsch-like | 138.3438988 |
| LOC107072587 | microphthalmia-associated transcription factor | 138.4399043 |
| LOC107066634 | yrdC domain-containing protein, mitochondrial | 138.4431917 |
| LOC107073074 | serine/threonine-protein kinase VRK1-like | 138.5176515 |
| LOC107064865 | uncharacterized protein LOC107064865 | 138.6058831 |
| LOC107065079 | TPR-containing protein DDB_G0280363-like | 139.247502 |
| LOC107074111 | phosphatidylinositol 3-kinase regulatory subunit alpha-like | 139.2987867 |
| LOC107066123 | uncharacterized protein LOC107066123 | 139.299376 |
| LOC107068043 | uncharacterized protein LOC107068043 | 139.3940604 |
| LOC107065412 | xenotropic and polytropic retrovirus receptor 1 homolog | 139.5118332 |
| LOC107073784 | RNA-binding protein 12B-like | 139.5301749 |
| LOC107072597 | uncharacterized protein LOC107072597 | 139.6082749 |
| LOC107074049 | uncharacterized protein LOC107074049 | 139.7077958 |
| LOC107070406 | putative fatty acyl-CoA reductase CG8306 | 139.7081858 |
| LOC107072162 | forkhead box protein P2 | 139.7088751 |
| LOC107072399 | protein pygopus | 139.8790877 |
| LOC107071575 | sine oculis-binding protein homolog | 139.9112843 |
| LOC107069226 | AT-rich interactive domain-containing protein 2-like | 140.1159837 |
| LOC107073166 | zinc finger FYVE domain-containing protein 1-like | 140.225423 |
| LOC107064075 | uncharacterized protein LOC107064075 | 140.4136387 |
| LOC107070150 | mucin-5AC | 140.4322184 |
| LOC107066088 | BAG domain-containing protein Samui | 140.4821889 |
| LOC107067139 | superoxide dismutase [Cu-Zn | 140.8364469 |
| LOC107069204 | abl interactor 2 | 140.8624034 |
| LOC107072026 | BTB/POZ domain-containing protein 10 | 140.885171 |
| LOC107067364 | alkylglycerol monooxygenase-like | 140.8955843 |
| LOC107066758 | homeobox protein cut-like | 140.897703 |
| LOC107065137 | uncharacterized protein LOC107065137 | 141.0187611 |
| LOC107069193 | muscle-specific protein 20-like | 141.1359939 |
| LOC107072559 | nicotinamide riboside kinase 1 | 141.1719799 |
| LOC107071675 | anosmin-1 | 141.3097833 |
| LOC107066016 | serine protease snake-like | 141.3282793 |
| LOC107063692 | uncharacterized protein LOC107063692 | 141.3507067 |
| LOC107073237 | ubiquitin fusion degradation protein 1 homolog | 141.3792223 |
| LOC107073247 | heat shock protein 70 A2-like | 141.4175822 |
| LOC107069165 | synaptosomal-associated protein 25 | 141.6017179 |
| LOC107066274 | uncharacterized protein LOC107066274 | 141.6834544 |

| | | |
|---|---|---|
| LOC107064690 | peripheral plasma membrane protein CASK | 141.7166954 |
| LOC107071570 | protein transport protein Sec23A | 141.718222 |
| LOC107068184 | 26S proteasome non-ATPase regulatory subunit 8 | 141.7962068 |
| LOC107071918 | catenin delta-2 | 141.8162433 |
| LOC107070816 | exosome complex component CSL4 | 141.9444354 |
| LOC107070545 | opioid-binding protein/cell adhesion molecule-like | 142.0305287 |
| LOC107066800 | protein SMG9 | 142.1056336 |
| LOC107072437 | protein FAM50 homolog | 142.1096266 |
| LOC107072673 | cadherin-89D | 142.1211664 |
| LOC107071440 | cyclin-dependent kinase 7 | 142.2306918 |
| LOC107070601 | ras-like protein family member 10B | 142.2576998 |
| LOC107070804 | zinc finger protein 845-like | 142.2615692 |
| LOC107070011 | uncharacterized protein LOC107070011 | 142.3490742 |
| LOC107065477 | heat shock 70 kDa protein cognate 4-like | 142.3650375 |
| LOC107068764 | dual specificity protein kinase CLK2 | 142.4374991 |
| LOC107070453 | uncharacterized protein LOC107070453 | 142.4856466 |
| LOC107069469 | uncharacterized protein LOC107069469 | 142.5476245 |
| LOC107070462 | transcription initiation factor IIB | 142.8710821 |
| LOC107071631 | basic helix-loop-helix transcription factor scleraxis | 142.9007745 |
| LOC107066161 | transducin beta-like protein 3 | 143.1648862 |
| LOC107073724 | uncharacterized protein LOC107073724 | 143.2138996 |
| LOC107065083 | LOW QUALITY PROTEIN: alkaline phosphatase-like | 143.2800553 |
| LOC107067455 | ralBP1-associated Eps domain-containing protein 1 | 143.426299 |
| LOC107066903 | uncharacterized protein LOC107066903 | 143.6172302 |
| LOC107073274 | negative elongation factor B-like | 143.6329364 |
| LOC107074382 | uncharacterized protein LOC107074382 | 143.6919419 |
| LOC107067822 | uncharacterized protein LOC107067822 | 143.7658492 |
| LOC107070750 | uncharacterized protein DDB_G0286299-like | 143.7759152 |
| LOC107069954 | coiled-coil domain-containing protein 25 | 143.8068555 |
| LOC107065425 | uncharacterized protein LOC107065425 | 143.8180553 |
| LOC107069793 | nuclear transcription factor Y subunit gamma-like | 143.8646892 |
| LOC107066879 | mitochondrial coenzyme A transporter SLC25A42-like | 143.9967091 |
| LOC107070326 | uncharacterized protein LOC107070326 | 144.2434847 |
| LOC107073214 | myotubularin-related protein 9 | 144.2843933 |
| LOC107068343 | microsomal glutathione S-transferase 1-like | 144.292456 |
| LOC107072462 | uncharacterized protein LOC107072462 | 144.4311609 |
| LOC107073685 | nose resistant to fluoxetine protein 6-like | 144.649989 |
| LOC107066782 | probable tRNA (uracil-O(2)-)-methyltransferase | 144.702094 |
| LOC107071999 | phospholipase D2 | 144.7325466 |
| LOC107063974 | protoheme IX farnesyltransferase, mitochondrial | 144.7336501 |
| LOC107070133 | netrin-1-like | 144.7418646 |
| LOC107074018 | uncharacterized protein C1orf198 homolog | 144.7437068 |
| LOC107072577 | uncharacterized protein LOC107072577 | 144.81388 |
| LOC107068792 | O-phosphoseryl-tRNA(Sec) selenium transferase | 144.9330869 |
| LOC107066523 | serum response factor-binding protein 1-like | 145.0382495 |
| LOC107065601 | defective pharyngeal development protein 4-like | 145.0731237 |
| LOC107068912 | uncharacterized protein LOC107068912 | 145.1296113 |
| LOC107066680 | barH-like 1 homeobox protein | 145.194515 |
| LOC107063896 | probable ATP-dependent RNA helicase DHX35 | 145.245969 |
| LOC107065931 | ankyrin repeat domain-containing protein 6 | 145.2470573 |

| | | |
|---|---|---|
| LOC107065199 | testican-2 | 145.3165461 |
| LOC107064799 | chymotrypsinogen 2-like | 145.4815299 |
| LOC107065049 | polypeptide N-acetylgalactosaminyltransferase 5 | 145.5945258 |
| LOC107065725 | THO complex subunit 6 homolog | 145.6508205 |
| LOC107070642 | uncharacterized protein LOC107070642 | 145.7943414 |
| LOC107068068 | uncharacterized protein CG10915 | 145.8579641 |
| LOC107065621 | popeye domain-containing protein 3-like | 145.8992741 |
| LOC107066226 | integrin-alpha FG-GAP repeat-containing protein 2-like | 145.9569187 |
| LOC107073326 | homeobox protein extradenticle | 146.006915 |
| LOC107071931 | tigger transposable element-derived protein 1-like | 146.0133405 |
| LOC107065474 | mediator of RNA polymerase II transcription subunit 28-like | 146.2316779 |
| LOC107066989 | disks large homolog 4-like | 146.2426427 |
| LOC107069910 | beta-1,3-galactosyltransferase 5-like | 146.3076936 |
| LOC107074145 | polyglutamine-binding protein 1 | 146.3621529 |
| LOC107071232 | uncharacterized protein LOC107071232 | 146.4791776 |
| LOC107067507 | uncharacterized protein LOC107067507 | 146.4979874 |
| LOC107065749 | GATA-binding factor C-like | 146.6458283 |
| LOC107066979 | trafficking protein particle complex subunit 2-like protein | 146.7781673 |
| LOC107066823 | coronin-6 | 146.789656 |
| LOC107067895 | uncharacterized protein LOC107067895 | 146.7933483 |
| LOC107072663 | actin-interacting protein 1 | 146.9894716 |
| LOC107072461 | putative fatty acyl-CoA reductase CG5065 | 147.1632762 |
| LOC107067396 | titin-like | 147.1705496 |
| LOC107064123 | uncharacterized protein LOC107064123 | 147.1919285 |
| LOC107067411 | tRNA (guanine-N(7)-)-methyltransferase | 147.2101557 |
| LOC107069772 | probable serine/threonine-protein kinase DDB_G0282963 | 147.4058284 |
| LOC107071935 | uncharacterized protein LOC107071935 | 147.488806 |
| LOC107071711 | troponin C, isoform 2-like | 147.7625482 |
| LOC107065011 | peptidyl-prolyl cis-trans isomerase CWC27 homolog | 147.994088 |
| LOC107070052 | LON peptidase N-terminal domain and RING finger protein 3 | 148.0782891 |
| LOC107066954 | uncharacterized protein LOC107066954 | 148.1182294 |
| LOC107066426 | uncharacterized protein LOC107066426 | 148.2517558 |
| LOC107070974 | uncharacterized protein LOC107070974 | 148.587489 |
| LOC107066485 | uncharacterized protein LOC107066485 | 148.7380521 |
| LOC107070185 | tRNA-dihydrouridine(47) synthase [NAD(P)(+) | 148.7532557 |
| LOC107065458 | vesicle transport protein SFT2B | 148.8720419 |
| LOC107066246 | transcription factor E2F5 | 148.8759157 |
| LOC107072782 | uncharacterized protein LOC107072782 | 148.9447805 |
| LOC107072496 | phosphoglucomutase | 149.0065904 |
| LOC107068805 | NF-kappa-B inhibitor cactus | 149.1027697 |
| LOC107065953 | ribonuclease inhibitor | 149.1720926 |
| LOC107066389 | 39S ribosomal protein L17, mitochondrial | 149.2587575 |
| LOC107067451 | uncharacterized protein LOC107067451 | 149.2846662 |
| LOC107069541 | zinc finger protein 384-like | 149.3233322 |
| LOC107070456 | pupal cuticle protein G1A-like | 149.3561971 |
| LOC107068365 | 40S ribosomal protein S25 | 149.4044552 |
| LOC107072765 | iodotyrosine deiodinase 1 | 149.4764133 |
| LOC107069583 | uncharacterized protein LOC107069583 | 149.4873632 |
| LOC107064594 | ubiquitin-40S ribosomal protein S27a | 149.5675047 |
| LOC107068144 | protein CBFA2T2 | 149.5913794 |

| | | |
|---|---|---|
| LOC107065395 | uncharacterized protein LOC107065395 | 149.6090179 |
| LOC107072662 | modifier of mdg4-like | 149.6311914 |
| LOC107068390 | E3 ubiquitin-protein ligase parkin | 149.6535488 |
| LOC107066710 | uncharacterized protein C18orf19 homolog B | 149.7308407 |
| LOC107066878 | mitotic checkpoint protein BUB3 | 149.7741956 |
| LOC107070401 | zinc finger and BTB domain-containing protein 41 | 149.7806285 |
| LOC107069251 | zinc finger protein ush | 150.0939873 |
| LOC107072614 | ubiquitin domain-containing protein 1 | 150.3258294 |
| LOC107069024 | nuclear hormone receptor FTZ-F1 beta | 150.3333426 |
| LOC107073605 | uncharacterized protein LOC107073605 | 150.389355 |
| LOC107070575 | protein FAM50 homolog | 150.4654276 |
| LOC107070189 | basement membrane-specific heparan sulfate proteoglycan c | 151.0972196 |
| LOC107069446 | UPF0472 protein C16orf72 homolog | 151.2169267 |
| LOC107067156 | ras-interacting protein RIP3-like | 151.226237 |
| LOC107069923 | proton-coupled amino acid transporter 1-like | 151.3112468 |
| LOC107069623 | uncharacterized protein LOC107069623 | 151.3146893 |
| LOC107070785 | uncharacterized protein LOC107070785 | 151.4689932 |
| LOC107069616 | B-box type zinc finger protein ncl-1 | 151.4709178 |
| LOC107064707 | importin-9 | 151.5563759 |
| LOC107072208 | AP-3 complex subunit sigma-2 | 151.6992731 |
| LOC107070079 | uncharacterized protein LOC107070079 | 151.7929445 |
| LOC107074652 | RAB6A-GEF complex partner protein 2 | 151.9282388 |
| LOC107067147 | ribokinase-like | 152.0808896 |
| LOC107064242 | uncharacterized protein LOC107064242 | 152.1315282 |
| LOC107068075 | probable glutamine-dependent NAD(+) synthetase | 152.1718271 |
| LOC107068500 | ejaculatory bulb-specific protein 3-like | 152.2112033 |
| LOC107073638 | cytochrome P450 4C1-like | 152.2263462 |
| LOC107073960 | insulin gene enhancer protein ISL-1 | 152.2996727 |
| LOC107066058 | KH domain-containing protein akap-1 | 152.3667992 |
| LOC107068682 | rho GTPase-activating protein 100F | 152.4901454 |
| LOC107067037 | PEST proteolytic signal-containing nuclear protein-like | 152.5492024 |
| LOC107066827 | thymidylate kinase | 152.6681016 |
| LOC107067415 | acetylcholine receptor subunit alpha-like | 152.7193387 |
| LOC107065704 | cob(I)yrinic acid a,c-diamide adenosyltransferase, mitochond | 152.8307313 |
| LOC107068263 | uncharacterized protein LOC107068263 | 152.8588451 |
| LOC107068070 | facilitated trehalose transporter Tret1-like | 152.9163849 |
| LOC107073524 | uncharacterized protein LOC107073524 | 152.9830394 |
| LOC107064534 | autophagy protein 12-like | 152.9972974 |
| LOC107065223 | dihydroorotate dehydrogenase (quinone), mitochondrial | 153.1043622 |
| LOC107066051 | interferon-inducible double-stranded RNA-dependent protei | 153.1121815 |
| LOC107066550 | uncharacterized protein LOC107066550 | 153.1210443 |
| LOC107066334 | glycerol kinase-like | 153.1679688 |
| LOC107073526 | uncharacterized protein LOC107073526 | 153.3295296 |
| LOC107064333 | cGMP-dependent protein kinase, isozyme 2 forms cD4/T1/T3 | 153.3405159 |
| LOC107063758 | RNA 3'-terminal phosphate cyclase | 153.4417049 |
| LOC107072912 | uncharacterized protein LOC107072912 | 153.690359 |
| LOC107072913 | uncharacterized protein LOC107072913 | 153.8786536 |
| LOC107071783 | actin-related protein 1 | 153.9493641 |
| LOC107070958 | tyramine beta-hydroxylase | 153.9522656 |
| LOC107073567 | uncharacterized protein LOC107073567 | 154.1170614 |

| | | |
|---|---|---|
| LOC107064430 | uncharacterized protein LOC107064430 | 154.2448815 |
| LOC107063884 | protein argonaute-2 | 154.3085015 |
| LOC107066409 | cAMP-dependent protein kinase catalytic subunit beta-like | 154.4581259 |
| LOC107070836 | MKL/myocardin-like protein 1 | 154.4649922 |
| LOC107069635 | uncharacterized protein LOC107069635 | 154.4783167 |
| LOC107065381 | uncharacterized protein LOC107065381 | 154.8205836 |
| LOC107072855 | uncharacterized protein LOC107072855 | 154.8297108 |
| LOC107068852 | zinc finger protein 729-like | 154.9580691 |
| LOC107071642 | cytoplasmic protein NCK1 | 154.9667982 |
| LOC107068466 | double-strand-break repair protein rad21 homolog | 155.0085061 |
| LOC107069839 | uncharacterized protein LOC107069839 | 155.0107624 |
| LOC107071810 | coiled-coil domain-containing protein 43 | 155.2768553 |
| LOC107071150 | uncharacterized protein LOC107071150 | 155.2785985 |
| LOC107071203 | uncharacterized protein LOC107071203 | 155.4317507 |
| LOC107072955 | uncharacterized protein LOC107072955 | 155.5139862 |
| LOC107071900 | uncharacterized protein LOC107071900 | 155.6661914 |
| LOC107067846 | retinal homeobox protein Rx2-like | 155.751812 |
| LOC107074533 | cell division control protein 45 homolog | 155.7653837 |
| LOC107070687 | autophagy-related protein 16-1 | 155.8954447 |
| LOC107065978 | calcitonin gene-related peptide type 1 receptor-like | 155.9649377 |
| LOC107065508 | C-terminal-binding protein | 156.0031069 |
| LOC107069646 | rhomboid-related protein 2 | 156.2279001 |
| LOC107071200 | WD repeat-containing protein 18 | 156.2716455 |
| LOC107065403 | trypsin-2-like | 156.2929292 |
| LOC107065342 | TNF receptor-associated factor 4 | 156.3621822 |
| LOC107066671 | F-box/LRR-repeat protein 7-like | 156.370804 |
| LOC107066489 | BRCA1-A complex subunit Abraxas-like | 156.4988259 |
| LOC107073474 | uncharacterized protein LOC107073474 | 156.7526141 |
| LOC107073956 | collagen type IV alpha-3-binding protein | 156.8021248 |
| LOC107064814 | uncharacterized protein LOC107064814 | 156.8572647 |
| LOC107072025 | SNAPIN protein homolog | 156.8645375 |
| LOC107065536 | fatty acid synthase-like | 156.8918034 |
| LOC107074558 | uncharacterized protein LOC107074558 | 156.9366818 |
| LOC107072574 | dystroglycan-like | 156.9858387 |
| LOC107067261 | ankyrin-3-like | 157.0991925 |
| LOC107069649 | leucine-rich repeat and immunoglobulin-like domain-contain | 157.1446554 |
| LOC107067738 | fatty acid synthase | 157.2108754 |
| LOC107073092 | synapse-associated protein 1 | 157.2729247 |
| LOC107073172 | breast cancer anti-estrogen resistance protein 3 | 157.4167957 |
| LOC107064138 | abhydrolase domain-containing protein 4 | 157.43069 |
| LOC107067053 | pituitary homeobox x | 157.4388979 |
| LOC107071462 | uncharacterized protein LOC107071462 | 157.4705889 |
| LOC107074108 | beta-hexosaminidase subunit beta-like | 157.6682482 |
| LOC107067319 | 3'-5' ssDNA/RNA exonuclease TatD | 157.6956755 |
| LOC107072222 | uncharacterized protein LOC107072222 | 157.7007138 |
| LOC107067183 | uncharacterized protein LOC107067183 | 157.7398028 |
| LOC107073871 | peptidyl-prolyl cis-trans isomerase 5 | 157.7684817 |
| LOC107068152 | protein KTI12 homolog | 157.7747201 |
| LOC107072672 | uncharacterized protein LOC107072672 | 158.297488 |
| LOC107072906 | elongation of very long chain fatty acids protein AAEL008004 | 158.3146671 |

| | | |
|---|---|---|
| LOC107065718 | serum response factor homolog B-like | 158.4411676 |
| LOC107064778 | neurogenic protein mastermind-like | 158.6301029 |
| LOC107073588 | uncharacterized protein LOC107073588 | 159.0945904 |
| LOC107067613 | pupal cuticle protein Edg-91-like | 159.1616111 |
| LOC107071509 | uncharacterized protein LOC107071509 | 159.254241 |
| LOC107065356 | inactive dipeptidyl peptidase 10 | 159.325505 |
| LOC107069354 | Krueppel-like factor 7 | 159.4459933 |
| LOC107064132 | 28S ribosomal protein S5, mitochondrial | 159.4495715 |
| LOC107072182 | beta-1,4-mannosyl-glycoprotein 4-beta-N-acetylglucosaminy | 159.5013299 |
| LOC107067095 | nucleoside diphosphate kinase 6 | 159.5115493 |
| LOC107070018 | twitchin | 159.7010242 |
| LOC107068773 | POU domain protein CF1A | 159.7019659 |
| LOC107066223 | uncharacterized protein LOC107066223 | 159.8803382 |
| LOC107071126 | uncharacterized protein LOC107071126 | 159.953767 |
| LOC107066856 | uncharacterized protein LOC107066856 | 160.0010296 |
| LOC107072343 | cell division cycle protein 123 homolog | 160.2562658 |
| LOC107071397 | probable uridine-cytidine kinase | 160.3261325 |
| LOC107073917 | vitellogenin-like | 160.4178851 |
| LOC107064519 | uncharacterized protein LOC107064519 | 160.5877513 |
| LOC107064536 | uncharacterized protein LOC107064536 | 160.6050251 |
| LOC107074379 | uncharacterized protein LOC107074379 | 160.7194907 |
| LOC107065848 | insulin-like growth factor 2 mRNA-binding protein 1 | 161.2241639 |
| LOC107074123 | Down syndrome cell adhesion molecule-like protein Dscam2 | 161.3582631 |
| LOC107069011 | sodium/potassium/calcium exchanger 3 | 161.4411032 |
| LOC107065368 | sialin-like | 161.5556697 |
| LOC107066851 | alpha-catulin | 161.5842497 |
| LOC107067154 | Bardet-Biedl syndrome 2 protein homolog | 161.7009512 |
| LOC107073097 | mitochondrial pyruvate carrier 2-like | 161.7402652 |
| LOC107066674 | S-adenosylmethionine mitochondrial carrier protein-like | 161.844908 |
| LOC107065579 | proclotting enzyme | 161.9277003 |
| LOC107068836 | putative zinc metalloproteinase YIL108W | 162.0074644 |
| LOC107068012 | protein phosphatase 1 regulatory subunit pprA-like | 162.0167544 |
| LOC107072941 | exocyst complex component 2 | 162.0436397 |
| LOC107071446 | phosphorylase b kinase gamma catalytic chain, skeletal musc | 162.2894614 |
| LOC107067045 | zinc finger protein 497-like | 162.2963095 |
| LOC107068715 | uncharacterized protein LOC107068715 | 162.4072866 |
| LOC107074007 | 5'-AMP-activated protein kinase subunit beta-1-like | 162.5390201 |
| LOC107070336 | uncharacterized protein LOC107070336 | 162.7100329 |
| LOC107066127 | uncharacterized protein LOC107066127 | 162.8008862 |
| LOC107064359 | proteasome subunit beta type-2-like | 162.8498262 |
| LOC107074363 | density-regulated protein homolog | 163.1061756 |
| LOC107065549 | protein disulfide-isomerase A6 | 163.2525689 |
| LOC107065926 | leptin receptor gene-related protein | 163.2631262 |
| LOC107068485 | translation elongation factor 2 | 163.400967 |
| LOC107067644 | mitochondrial ribosome-associated GTPase 2 | 163.5117737 |
| LOC107071534 | DNA-binding protein D-ETS-4-like | 163.5613503 |
| LOC107070735 | synaptic vesicle glycoprotein 2B-like | 163.9234145 |
| LOC107065531 | sodium channel protein 60E | 164.3003987 |
| LOC107066666 | PRKCA-binding protein | 164.3081589 |
| LOC107074287 | ribose-phosphate pyrophosphokinase 1 | 164.7446168 |

| | | |
|---|---|---|
| LOC107067254 | myb-like protein D | 164.9626094 |
| LOC107072475 | uncharacterized protein LOC107072475 | 165.0147923 |
| LOC107070391 | glutamate--cysteine ligase regulatory subunit | 165.0153229 |
| LOC107066631 | proline dehydrogenase 1, mitochondrial | 165.0189876 |
| LOC107068294 | transcriptional repressor protein YY1-like | 165.0769503 |
| LOC107067778 | uncharacterized protein LOC107067778 | 165.1161588 |
| LOC107073542 | uncharacterized protein LOC107073542 | 165.1589916 |
| LOC107071311 | uncharacterized protein LOC107071311 | 165.256836 |
| LOC107070668 | cyclin-dependent kinase 9-like | 165.3459996 |
| LOC107070599 | transcription factor Sp9 | 165.4587895 |
| LOC107073226 | forkhead box protein E1-like | 165.5976768 |
| LOC107068502 | GATA zinc finger domain-containing protein 10-like | 165.6459843 |
| LOC107073704 | uncharacterized protein LOC107073704 | 165.7051321 |
| LOC107066756 | peptidyl-prolyl cis-trans isomerase H | 165.7915359 |
| LOC107071875 | peroxisomal targeting signal 1 receptor | 166.0215412 |
| LOC107065009 | uncharacterized protein LOC107065009 | 166.1074529 |
| LOC107068217 | kinesin-related protein 4 | 166.1156662 |
| LOC107065685 | peroxidase | 166.349583 |
| LOC107064470 | kinesin-like protein CG14535 | 166.5456333 |
| LOC107070957 | putative phospholipase B-like lamina ancestor | 166.6464573 |
| LOC107070303 | AFG3-like protein 2 | 166.6507396 |
| LOC107071166 | sorting and assembly machinery component 50 homolog | 166.7502703 |
| LOC107069936 | uncharacterized protein LOC107069936 | 166.804108 |
| LOC107064595 | ubiquitin-40S ribosomal protein S27a | 166.9239105 |
| LOC107064067 | cholinesterase 2-like | 166.9909477 |
| LOC107064420 | plasminogen activator inhibitor 1 RNA-binding protein | 167.0509312 |
| LOC107067515 | zinc finger protein 585B-like | 167.5446951 |
| LOC107072205 | cyclic AMP-dependent transcription factor ATF-2 | 167.588588 |
| LOC107069577 | uncharacterized protein LOC107069577 | 167.7673985 |
| LOC107068855 | growth factor receptor-bound protein 14-like | 167.7733016 |
| LOC107064857 | protein enabled-like | 168.0395624 |
| LOC107065557 | transmembrane protein adipocyte-associated 1 homolog | 168.1344842 |
| LOC107067271 | autism susceptibility gene 2 protein-like | 168.1606165 |
| LOC107064743 | uncharacterized protein LOC107064743 | 168.5782152 |
| LOC107074635 | BCL2/adenovirus E1B 19 kDa protein-interacting protein 3 | 168.6585618 |
| LOC107066624 | heat shock 70 kDa protein cognate 5 | 168.9793027 |
| LOC107064509 | protein lin-9 homolog | 169.016542 |
| LOC107068801 | CUE domain-containing protein 2 | 169.1227149 |
| LOC107071899 | uncharacterized protein LOC107071899 | 169.2769234 |
| LOC107068577 | histidine protein methyltransferase 1 homolog | 169.2829116 |
| LOC107065520 | uncharacterized protein LOC107065520 | 169.36621 |
| LOC107069835 | uncharacterized protein LOC107069835 | 169.7763916 |
| LOC107068108 | uncharacterized protein LOC107068108 | 169.8607087 |
| LOC107065695 | uridine-cytidine kinase-like 1 | 170.1199586 |
| LOC107072127 | cytochrome P450 4C1-like | 170.3977359 |
| LOC107067236 | UDP-N-acetylglucosamine transferase subunit ALG14 | 170.7178313 |
| LOC107067093 | uncharacterized protein LOC107067093 | 170.8143898 |
| LOC107067789 | uncharacterized protein LOC107067789 | 170.8383156 |
| LOC107066901 | uncharacterized protein LOC107066901 | 170.8696296 |
| LOC107073973 | WD and tetratricopeptide repeats protein 1-like | 170.8854351 |

| | | |
|---|---|---|
| LOC107074329 | uncharacterized protein LOC107074329 | 170.9915493 |
| LOC107074361 | serine/threonine-protein kinase ULK2 | 171.1700335 |
| LOC107069716 | RNA pseudouridylate synthase domain-containing protein 2-l | 171.3575827 |
| LOC107064875 | blood vessel epicardial substance-A-like | 171.3626275 |
| LOC107065324 | protein eyes shut | 171.4456789 |
| LOC107065710 | histone-lysine N-methyltransferase Suv4-20-like | 171.4609337 |
| LOC107067228 | uncharacterized protein LOC107067228 | 171.4656132 |
| LOC107068035 | zinc finger protein 391-like | 171.5936801 |
| LOC107073623 | peroxidase-like | 171.5953871 |
| LOC107074402 | protein bowel-like | 171.800961 |
| LOC107064556 | sentrin-specific protease 8-like | 172.0050921 |
| LOC107074698 | RNA-binding protein MEX3B | 172.2110236 |
| LOC107070392 | putative transcription factor SOX-14 | 172.21824 |
| LOC107064108 | uncharacterized protein LOC107064108 | 172.5927472 |
| LOC107065766 | proto-oncogene tyrosine-protein kinase receptor Ret-like | 172.6660026 |
| LOC107065228 | uncharacterized protein LOC107065228 | 172.7744626 |
| LOC107071943 | protein PIH1D3 | 172.9468298 |
| LOC107072761 | uncharacterized protein LOC107072761 | 173.0435508 |
| LOC107065361 | cytochrome P450 4C1-like | 173.0484565 |
| LOC107067579 | probable elongator complex protein 3 | 173.0717866 |
| LOC107065096 | DAZ-associated protein 2-like | 173.1687512 |
| LOC107073360 | uncharacterized protein LOC107073360 | 173.2640233 |
| LOC107071856 | cytosolic endo-beta-N-acetylglucosaminidase | 173.2954519 |
| LOC107066991 | uncharacterized protein LOC107066991 | 173.3083071 |
| LOC107073939 | protein timeless homolog | 173.4905322 |
| LOC107070813 | mediator of RNA polymerase II transcription subunit 15 | 173.496912 |
| LOC107068304 | methyltransferase-like protein 2-A | 173.5538937 |
| LOC107068673 | 4-hydroxybenzoate polyprenyltransferase, mitochondrial | 173.5828655 |
| LOC107065764 | uncharacterized protein LOC107065764 | 173.6024727 |
| LOC107068160 | enhancer of polycomb homolog 1 | 173.6762863 |
| LOC107068084 | uncharacterized protein LOC107068084 | 173.7023539 |
| LOC107069140 | WW domain-containing oxidoreductase | 173.8436118 |
| LOC107070773 | uncharacterized protein LOC107070773 | 173.8774866 |
| LOC107063969 | protein archease-like | 174.0666293 |
| LOC107070669 | catalase | 174.2630351 |
| LOC107071684 | uncharacterized protein LOC107071684 | 174.3761421 |
| LOC107072950 | uncharacterized protein LOC107072950 | 174.3984162 |
| LOC107070751 | zinc finger CCCH domain-containing protein 15 homolog | 174.4278317 |
| LOC107073958 | histone deacetylase Rpd3 | 174.5218186 |
| LOC107064258 | histone H3.3 | 174.546841 |
| LOC107063765 | uncharacterized protein LOC107063765 | 174.7692011 |
| LOC107070985 | LOW QUALITY PROTEIN: homeobox protein homothorax | 174.7751058 |
| LOC107064195 | 60S ribosomal protein L27 | 174.8103895 |
| LOC107069100 | putative uncharacterized protein DDB_G0290521 | 174.8339481 |
| LOC107067317 | palmitoyltransferase ZDHHC5 | 174.9321114 |
| LOC107067356 | basic salivary proline-rich protein 2 | 175.0462521 |
| LOC107063865 | uncharacterized protein LOC107063865 | 175.0544293 |
| LOC107070975 | dolichyl-diphosphooligosaccharide--protein glycosyltransfera | 175.1129234 |
| LOC107068234 | serine/threonine-protein phosphatase 2A 56 kDa regulatory | 175.1815359 |
| LOC107069137 | protein gustavus | 175.2496914 |

| | | |
|---|---|---|
| LOC107065812 | ubiquitin carboxyl-terminal hydrolase 3-like | 175.3048334 |
| LOC107072175 | uncharacterized protein LOC107072175 | 175.6840349 |
| LOC107063693 | uncharacterized protein KIAA0195 | 176.1255669 |
| LOC107072666 | synaptic vesicle glycoprotein 2B-like | 176.1675845 |
| LOC107072537 | ataxin-7-like protein 3 | 176.2511846 |
| LOC107066792 | myotubularin-related protein 3 | 176.4254523 |
| LOC107063935 | GTP-binding protein Rheb homolog | 176.7803564 |
| LOC107072678 | probable nuclear hormone receptor HR3 | 176.8287921 |
| LOC107068587 | uncharacterized protein LOC107068587 | 176.9447062 |
| LOC107072166 | protein tyrosine phosphatase domain-containing protein 1-lil | 177.2383986 |
| LOC107067399 | dolichyl-phosphate beta-glucosyltransferase | 177.3119361 |
| LOC107071190 | protein kinase C-binding protein NELL1-like | 177.4202244 |
| LOC107065359 | cytochrome P450 4C1-like | 177.5255355 |
| LOC107074557 | RNA polymerase II elongation factor Ell-like | 178.0625627 |
| LOC107070987 | carboxyl-terminal PDZ ligand of neuronal nitric oxide synthas | 178.1714201 |
| LOC107068480 | exosome complex exonuclease RRP42-like | 178.4313959 |
| LOC107068601 | cytochrome P450 4g15 | 178.5880473 |
| LOC107071313 | uncharacterized protein LOC107071313 | 178.6468713 |
| LOC107071547 | double-stranded RNA-binding protein Staufen homolog 2 | 178.6543889 |
| LOC107066528 | dynein heavy chain 7, axonemal-like | 178.6567332 |
| LOC107068511 | serum response factor homolog | 178.7772911 |
| LOC107068791 | cytosolic purine 5'-nucleotidase | 178.9472466 |
| LOC107064198 | arylsulfatase B-like | 179.0536219 |
| LOC107066517 | upstream stimulatory factor 2-like | 179.1137039 |
| LOC107065858 | DNA-binding protein D-ETS-6-like | 179.135795 |
| LOC107071719 | type 1 phosphatidylinositol 4,5-bisphosphate 4-phosphatase | 179.2299435 |
| LOC107069846 | protein fem-1 homolog CG6966 | 179.2583812 |
| LOC107063663 | ecdysone receptor | 179.2926164 |
| LOC107074296 | loss of heterozygosity 12 chromosomal region 1 protein hom | 179.5567945 |
| LOC107064494 | protein phosphatase methylesterase 1 | 179.5852072 |
| LOC107067176 | heterogeneous nuclear ribonucleoprotein H-like | 179.8848755 |
| LOC107067783 | E3 ubiquitin-protein ligase MIB1 | 180.2632502 |
| LOC107063746 | CCA tRNA nucleotidyltransferase 1, mitochondrial | 180.545703 |
| LOC107073350 | uncharacterized protein LOC107073350 | 180.6550114 |
| LOC107068194 | lysM and putative peptidoglycan-binding domain-containing | 180.7039673 |
| LOC107070786 | uncharacterized protein LOC107070786 | 180.8598049 |
| LOC107065519 | chymotrypsin inhibitor-like | 180.9034662 |
| LOC107064662 | T-complex protein 1 subunit zeta | 181.3237392 |
| LOC107073335 | uncharacterized protein LOC107073335 | 181.779153 |
| LOC107063875 | uncharacterized protein LOC107063875 | 181.908268 |
| LOC107068185 | 40S ribosomal protein S11 | 182.4074908 |
| LOC107070181 | ankyrin repeat domain-containing protein 50 | 182.7008498 |
| LOC107066858 | uncharacterized protein LOC107066858 | 182.8817596 |
| LOC107063925 | uncharacterized protein LOC107063925 | 182.8996678 |
| LOC107065162 | ubiquitin-conjugating enzyme E2 variant 2 | 183.0766234 |
| LOC107065823 | CUGBP Elav-like family member 1 | 183.2114441 |
| LOC107063972 | phosphatidylinositol N-acetylglucosaminyltransferase subuni | 183.4423533 |
| LOC107073558 | uncharacterized protein LOC107073558 | 183.5659222 |
| LOC107066951 | cytochrome P450 4C1-like | 183.7562162 |
| LOC107064085 | sodium/calcium exchanger 1 | 183.8835954 |

| | | |
|---|---|---|
| LOC107068081 | probable 18S rRNA (guanine-N(7))-methyltransferase | 183.9237396 |
| LOC107067301 | ATP-binding cassette sub-family G member 1-like | 183.9943653 |
| LOC107071363 | radial spoke head 10 homolog B-like | 184.2063545 |
| LOC107068829 | transmembrane protein 132E | 184.8921012 |
| LOC107063738 | broad-complex core protein isoforms 1/2/3/4/5-like | 184.9366482 |
| LOC107073380 | mediator of RNA polymerase II transcription subunit 14 | 184.9565306 |
| LOC107063694 | acetylcholinesterase-like | 185.3514369 |
| LOC107068797 | L-ascorbate oxidase-like | 185.5331177 |
| LOC107068942 | 40S ribosomal protein S13-like | 185.6767246 |
| LOC107065698 | 60S ribosomal protein L15 | 185.6799927 |
| LOC107063768 | uncharacterized protein LOC107063768 | 185.9513458 |
| LOC107064917 | peptidylglycine alpha-hydroxylating monooxygenase | 186.1017476 |
| LOC107070212 | silk gland factor 1-like | 186.1847403 |
| LOC107064167 | follistatin | 186.2345929 |
| LOC107069302 | 26S proteasome non-ATPase regulatory subunit 13 | 186.3538137 |
| LOC107066965 | late histone H1-like | 186.6089963 |
| LOC107065160 | adenylate cyclase type 3-like | 186.7033181 |
| LOC107067157 | zinc finger protein 13 | 186.7282078 |
| LOC107065212 | PHD finger-like domain-containing protein 5A | 186.7321084 |
| LOC107069799 | ankyrin repeat domain-containing protein SOWAHC | 186.8504983 |
| LOC107074706 | segmentation protein Runt-like | 186.935259 |
| LOC107065756 | homogentisate 1,2-dioxygenase | 186.9423218 |
| LOC107074166 | 40S ribosomal protein SA | 187.0074797 |
| LOC107068589 | potassium voltage-gated channel protein Shaker | 187.0225443 |
| LOC107064084 | serine protease inhibitor dipetalogastin | 187.415282 |
| LOC107064488 | kelch-like protein 5 | 187.4954157 |
| LOC107072164 | putative mediator of RNA polymerase II transcription subunit | 187.7661732 |
| LOC107073007 | protein singed | 187.8699387 |
| LOC107067665 | C-type lectin mannose-binding isoform-like | 187.9806751 |
| LOC107068602 | serine palmitoyltransferase 2 | 187.9975609 |
| LOC107065643 | serine/threonine-protein kinase NLK | 188.0059465 |
| LOC107074023 | DNA helicase MCM9-like | 188.022066 |
| LOC107071136 | dentin sialophosphoprotein-like | 188.1680789 |
| LOC107074286 | histone-binding protein N1/N2-like | 188.3456572 |
| LOC107068776 | serine/threonine-protein kinase greatwall | 188.4810266 |
| LOC107072195 | DNA-binding protein Ets97D | 188.693447 |
| LOC107067577 | TBC1 domain family member 4 | 188.7407519 |
| LOC107065069 | knirps-related protein-like | 189.0115175 |
| LOC107074252 | uncharacterized protein LOC107074252 | 189.1617718 |
| LOC107064649 | metallophosphoesterase domain-containing protein 1 | 189.2213438 |
| LOC107071539 | probable serine/threonine-protein kinase yakA | 189.2318862 |
| LOC107067122 | uncharacterized protein LOC107067122 | 189.2814308 |
| LOC107067284 | putative inorganic phosphate cotransporter | 189.4710606 |
| LOC107072490 | cytoplasmic tRNA 2-thiolation protein 1 | 190.0465027 |
| LOC107067842 | uncharacterized protein LOC107067842 | 190.0505601 |
| LOC107071674 | homeobox protein MOX-1 | 190.1416107 |
| LOC107069358 | uncharacterized protein LOC107069358 | 190.2983906 |
| LOC107072279 | stress response protein NST1 | 190.3909103 |
| LOC107073774 | uncharacterized protein LOC107073774 | 190.459256 |
| LOC107069019 | rab5 GDP/GTP exchange factor | 190.7188886 |

| | | |
|---|---|---|
| LOC107066104 | IgA FC receptor-like | 190.9919155 |
| LOC107065770 | polyadenylate-binding protein 2 | 191.0385599 |
| LOC107069298 | myrosinase 1-like | 191.1544896 |
| LOC107067777 | p53 and DNA damage-regulated protein 1 | 191.2675661 |
| LOC107074298 | MORN repeat-containing protein 3-like | 191.4668595 |
| LOC107067707 | homeotic protein Sex combs reduced | 191.5752779 |
| LOC107068430 | cytochrome P450 6a2-like | 191.5931919 |
| LOC107074512 | akirin-2 | 191.6335517 |
| LOC107068838 | uncharacterized protein C7orf26 homolog | 191.635866 |
| LOC107069338 | calpain-C | 191.6671241 |
| LOC107073392 | uncharacterized protein LOC107073392 | 191.6718692 |
| LOC107072427 | H/ACA ribonucleoprotein complex subunit 1-like | 192.5246631 |
| LOC107070680 | uncharacterized protein LOC107070680 | 192.6224595 |
| LOC107070556 | U4/U6 small nuclear ribonucleoprotein Prp4 | 192.7046137 |
| LOC107072507 | atypical protein kinase C | 192.7625719 |
| LOC107068330 | protein Tob1 | 192.868089 |
| LOC107070379 | N-acetylglucosaminyl-phosphatidylinositol de-N-acetylase | 192.9954175 |
| LOC107074181 | leucine-rich repeat and calponin homology domain-containir | 193.0566031 |
| LOC107063686 | solute carrier family 25 member 36-A | 193.1094076 |
| LOC107071110 | uncharacterized protein LOC107071110 | 193.2217722 |
| LOC107070407 | putative fatty acyl-CoA reductase CG5065 | 193.3889752 |
| LOC107072720 | DNA-binding protein Ewg | 193.4814464 |
| LOC107074095 | uncharacterized protein LOC107074095 | 193.5380475 |
| LOC107066660 | uncharacterized protein LOC107066660 | 193.7304636 |
| LOC107066230 | box A-binding factor-like | 193.7808875 |
| LOC107068953 | protein RMD5 homolog A | 194.0464384 |
| LOC107069591 | transmembrane protein 53 | 194.8197987 |
| LOC107066860 | ras association domain-containing protein 8 | 195.1026752 |
| LOC107072321 | uncharacterized protein LOC107072321 | 195.5042405 |
| LOC107068508 | MAP kinase-interacting serine/threonine-protein kinase 1 | 195.5956921 |
| LOC107073446 | fatty acyl-CoA reductase 1-like | 195.8994062 |
| LOC107066558 | pumilio homolog 2 | 196.0176252 |
| LOC107073576 | uncharacterized protein LOC107073576 | 196.0250369 |
| LOC107068950 | uncharacterized protein LOC107068950 | 196.0333687 |
| LOC107072806 | actin-binding LIM protein 2 | 196.0753971 |
| LOC107070667 | SPARC-related modular calcium-binding protein 2 | 196.4016995 |
| LOC107070242 | casein kinase II subunit alpha | 196.6281721 |
| LOC107074437 | nuclear factor 1 X-type | 196.771324 |
| LOC107065938 | uncharacterized protein LOC107065938 | 196.828172 |
| LOC107064609 | CCAAT/enhancer-binding protein | 196.8569591 |
| LOC107069762 | dipeptidyl peptidase 9 | 196.8754406 |
| LOC107073630 | retinoblastoma-binding protein 5 homolog | 196.9607569 |
| LOC107066568 | receptor-interacting serine/threonine-protein kinase 4-like | 197.3781898 |
| LOC107074438 | uncharacterized protein LOC107074438 | 197.4210903 |
| LOC107064392 | leucine-rich repeat-containing protein 23-like | 197.5032042 |
| LOC107074575 | transcriptional enhancer factor TEF-1 | 197.5599081 |
| LOC107071665 | uncharacterized protein LOC107071665 | 197.7876424 |
| LOC107068080 | RING finger protein 113A | 197.8029402 |
| LOC107071651 | protein Malvolio | 197.8890404 |
| LOC107067713 | uncharacterized protein LOC107067713 | 197.9059932 |

| | | |
|---|---|---|
| LOC107066677 | GA-binding protein subunit beta-2 | 198.1451968 |
| LOC107071111 | DNA-directed RNA polymerase I subunit RPA12 | 198.335342 |
| LOC107066589 | histone-lysine N-methyltransferase setd3 | 198.7033987 |
| LOC107064157 | stAR-related lipid transfer protein 7, mitochondrial-like | 199.4900805 |
| LOC107071066 | elongation of very long chain fatty acids protein AAEL008004 | 199.5255046 |
| LOC107068119 | mitochondrial ubiquitin ligase activator of nfkb 1-like | 199.9059356 |
| LOC107064339 | zinc finger protein 346 | 199.9774095 |
| LOC107071112 | uncharacterized protein LOC107071112 | 200.3876146 |
| LOC107065810 | furin-like protease 2 | 200.4187577 |
| LOC107068244 | 4-coumarate--CoA ligase 1 | 200.5874567 |
| LOC107068569 | uncharacterized protein LOC107068569 | 200.9032265 |
| LOC107068633 | F-box/LRR-repeat protein 2 | 200.9594982 |
| LOC107069133 | interleukin-1 receptor accessory protein-like 1-B | 201.3514928 |
| LOC107071795 | acyl-CoA Delta(11) desaturase-like | 201.451174 |
| LOC107066875 | protein hairy | 201.587872 |
| LOC107067580 | ABC transporter G family member 14 | 201.6000241 |
| LOC107072653 | single-stranded DNA-binding protein 3 | 202.5331182 |
| LOC107070040 | RNA-binding protein squid-like | 202.5522451 |
| LOC107069130 | lachesin-like | 202.6476799 |
| LOC107069158 | serine/threonine-protein kinase tricorner | 202.7362796 |
| LOC107067973 | uncharacterized protein LOC107067973 | 202.9111861 |
| LOC107074300 | four and a half LIM domains protein 3 | 203.1155095 |
| LOC107068616 | ejaculatory bulb-specific protein 3 | 203.2168324 |
| LOC107063666 | cAMP-specific 3',5'-cyclic phosphodiesterase, isoform F | 203.3570639 |
| LOC107072332 | coatomer subunit delta | 203.6103753 |
| LOC107067128 | sesquipedalian-1-like | 203.6697597 |
| LOC107071857 | uncharacterized protein LOC107071857 | 203.7471926 |
| LOC107073364 | uncharacterized protein LOC107073364 | 204.0937477 |
| LOC107066638 | pre-mRNA 3'-end-processing factor FIP1 | 204.1949753 |
| LOC107068977 | WD repeat-containing protein 3 | 204.6676376 |
| LOC107066744 | CREB-regulated transcription coactivator 1-like | 204.8196397 |
| LOC107064717 | uncharacterized protein LOC107064717 | 204.828441 |
| LOC107064607 | serine/arginine-rich splicing factor 1B | 204.9132626 |
| LOC107065374 | potassium voltage-gated channel subfamily KQT member 1 | 204.9696943 |
| LOC107066284 | protein vein | 205.2056929 |
| LOC107069120 | brain mitochondrial carrier protein 1 | 205.2909989 |
| LOC107069305 | homeobox protein Nkx-6.1-like | 205.2921147 |
| LOC107068109 | uncharacterized protein LOC107068109 | 205.6304124 |
| LOC107063843 | uncharacterized protein LOC107063843 | 205.9058435 |
| LOC107072917 | nucleosome assembly protein 1-like 1-B | 205.9649809 |
| LOC107067706 | armadillo repeat-containing protein 8-like | 205.9735157 |
| LOC107066509 | brahma-associated protein of 60 kDa | 206.1109358 |
| LOC107073944 | integrator complex subunit 7 | 206.3018958 |
| LOC107065516 | proline-rich receptor-like protein kinase PERK9 | 206.7009182 |
| LOC107068007 | probable calcium-binding protein CML22 | 206.8382554 |
| LOC107073424 | aprataxin and PNK-like factor | 206.9904946 |
| LOC107069759 | protein SGT1 homolog | 207.0027 |
| LOC107064396 | methylenetetrahydrofolate reductase | 207.2660237 |
| LOC107070167 | E3 ubiquitin-protein ligase znrf1 | 207.458656 |
| LOC107065708 | uncharacterized protein LOC107065708 | 207.4737283 |

| | | |
|---|---|---|
| LOC107071973 | Down syndrome cell adhesion molecule-like protein Dscam2 | 207.7099306 |
| LOC107073423 | uncharacterized protein LOC107073423 | 207.7831959 |
| LOC107071124 | homeodomain-interacting protein kinase 2 | 207.9681702 |
| LOC107074348 | multidrug resistance protein homolog 49-like | 208.2278205 |
| LOC107072286 | uncharacterized protein LOC107072286 | 208.3102925 |
| LOC107068251 | uncharacterized protein LOC107068251 | 208.3628923 |
| LOC107067891 | tyrosine aminotransferase | 208.5638103 |
| LOC107070088 | uncharacterized protein LOC107070088 | 208.5950841 |
| LOC107068422 | uncharacterized protein LOC107068422 | 208.8344405 |
| LOC107064479 | YTH domain-containing family protein 3-like | 208.9883756 |
| LOC107069089 | small ubiquitin-related modifier-like | 209.0200514 |
| LOC107068862 | cytochrome P450 306a1 | 209.0430394 |
| LOC107071108 | uncharacterized protein LOC107071108 | 209.1567691 |
| LOC107073783 | uncharacterized protein T26G10.4-like | 209.2341128 |
| LOC107071325 | replication factor C subunit 4 | 209.4144446 |
| LOC107072970 | homeobox protein onecut-like | 209.5154356 |
| LOC107066914 | poly(rC)-binding protein 4-like | 209.789009 |
| LOC107070741 | 39S ribosomal protein L50, mitochondrial | 209.9323916 |
| LOC107067099 | uncharacterized protein LOC107067099 | 210.2208316 |
| LOC107065499 | transcriptional activator protein Pur-beta-B | 210.3985471 |
| LOC107064269 | TGF-beta-activated kinase 1 and MAP3K7-binding protein 3 | 210.6024724 |
| LOC107069149 | metabotropic glutamate receptor 6-like | 210.6505254 |
| LOC107069630 | cleavage and polyadenylation specificity factor subunit CG71 | 210.8711941 |
| LOC107072207 | GATA zinc finger domain-containing protein 1 | 211.2846339 |
| LOC107068287 | RNMT-activating mini protein | 211.4377165 |
| LOC107065934 | uncharacterized protein LOC107065934 | 211.6518194 |
| LOC107067003 | B-cell CLL/lymphoma 7 protein family member A | 211.6658658 |
| LOC107069848 | protein groucho | 211.7026298 |
| LOC107074325 | reticulocalbin-2 | 211.7213298 |
| LOC107073161 | cyclin-dependent kinase 9 | 212.1079286 |
| LOC107072188 | NADH dehydrogenase [ubiquinone | 212.15983 |
| LOC107069057 | charged multivesicular body protein 1b | 212.3892037 |
| LOC107068300 | ATP-dependent RNA helicase WM6 | 212.6187824 |
| LOC107065595 | proto-oncogene tyrosine-protein kinase receptor Ret | 212.6617723 |
| LOC107073291 | uncharacterized protein LOC107073291 | 212.6767639 |
| LOC107065659 | chondroitin proteoglycan 2-like | 212.685187 |
| LOC107070653 | ADP-ribosylation factor-like protein 16 | 212.7695226 |
| LOC107071493 | uncharacterized protein LOC107071493 | 213.0957273 |
| LOC107071144 | uncharacterized protein LOC107071144 | 213.4664688 |
| LOC107070996 | mimitin, mitochondrial | 214.1902671 |
| LOC107065233 | nuclear hormone receptor FTZ-F1 | 214.2812795 |
| LOC107066073 | retinaldehyde-binding protein 1-like | 214.4004505 |
| LOC107073435 | uncharacterized protein LOC107073435 | 214.4915535 |
| LOC107074569 | uncharacterized protein LOC107074569 | 214.5298308 |
| LOC107067732 | FAS-associated factor 1 | 214.6574825 |
| LOC107074690 | kinesin-like protein KIF9 | 215.6321423 |
| LOC107067324 | uncharacterized protein LOC107067324 | 216.0902005 |
| LOC107067821 | mushroom body large-type Kenyon cell-specific protein 1 | 216.3198854 |
| LOC107067449 | beta-1,4-glucuronyltransferase 1-like | 216.5783139 |
| LOC107069217 | leucine-rich repeat neuronal protein 1-like | 216.7917023 |

| | | |
|---|---|---|
| LOC107072341 | uncharacterized protein LOC107072341 | 217.0227489 |
| LOC107067330 | prohibitin-2 | 217.3821874 |
| LOC107070625 | zinc finger protein 608-like | 217.663177 |
| LOC107070546 | cofilin/actin-depolymerizing factor homolog | 217.7813483 |
| LOC107066427 | nucleolar protein 4-like | 217.8176247 |
| LOC107066687 | trinucleotide repeat-containing gene 6C protein | 218.0087117 |
| LOC107064383 | uncharacterized protein LOC107064383 | 218.1449896 |
| LOC107065578 | toll-like receptor 8 | 218.2937008 |
| LOC107070152 | tctex1 domain-containing protein 2-like | 218.3274018 |
| LOC107067181 | DNA N6-methyl adenine demethylase | 218.5796841 |
| LOC107067322 | uncharacterized protein LOC107067322 | 218.6211749 |
| LOC107074036 | uncharacterized protein LOC107074036 | 218.6923803 |
| LOC107067447 | protein tumorous imaginal discs, mitochondrial | 218.9480141 |
| LOC107068906 | T-complex protein 1 subunit beta | 219.703049 |
| LOC107069052 | transcription factor BTF3 homolog 4 | 220.1140709 |
| LOC107071663 | uncharacterized protein LOC107071663 | 220.2377566 |
| LOC107070192 | 45 kDa calcium-binding protein | 221.563363 |
| LOC107071821 | guanine nucleotide exchange factor MSS4 | 221.8922007 |
| LOC107074270 | uncharacterized protein LOC107074270 | 222.2413291 |
| LOC107065128 | histone H2A-like | 222.4977553 |
| LOC107066828 | box C/D snoRNA protein 1 | 223.1739832 |
| LOC107069142 | uncharacterized protein LOC107069142 | 223.2285409 |
| LOC107065821 | transforming growth factor beta regulator 1 | 223.9334721 |
| LOC107070486 | ADP-ribosylation factor-like protein 1 | 224.4313256 |
| LOC107067016 | splicing factor 1-like | 224.653422 |
| LOC107066218 | zinc finger CCHC domain-containing protein 10-like | 224.8452702 |
| LOC107066598 | uncharacterized protein LOC107066598 | 225.2657305 |
| LOC107071303 | zinc finger protein Elbow-like | 225.703295 |
| LOC107065093 | nucleolysin TIAR | 225.7889489 |
| LOC107069542 | F-box-like/WD repeat-containing protein TBL1XR1 | 225.9438128 |
| LOC107070828 | small glutamine-rich tetratricopeptide repeat-containing pro | 226.0271242 |
| LOC107073825 | DNA mismatch repair protein MutL-like | 226.630511 |
| LOC107065176 | pyridoxal-dependent decarboxylase domain-containing prote | 226.7178569 |
| LOC107067582 | CUGBP Elav-like family member 4 | 226.7657838 |
| LOC107073230 | uncharacterized protein LOC107073230 | 226.8782225 |
| LOC107068685 | vacuolar protein sorting-associated protein 37B | 227.1518602 |
| LOC107071370 | fatty acid synthase-like | 227.3807827 |
| LOC107073372 | beta-1,3-galactosyltransferase 6 | 227.5686992 |
| LOC107070320 | protein Skeletor, isoforms B/C | 228.2200113 |
| LOC107064577 | 5-oxoprolinase | 228.4726233 |
| LOC107065469 | uncharacterized protein C3orf18-like | 228.4938759 |
| LOC107072364 | uncharacterized protein LOC107072364 | 228.5116081 |
| LOC107066048 | regulator of chromosome condensation-like | 228.5606409 |
| LOC107069782 | putative mediator of RNA polymerase II transcription subunit | 228.7236818 |
| LOC107071664 | mitochondrial distribution and morphology protein 12-like | 228.8885448 |
| LOC107067743 | uncharacterized protein LOC107067743 | 230.2090501 |
| LOC107074655 | ribosomal protein S6 kinase 2 beta | 230.3590021 |
| LOC107070394 | zinc finger protein jing homolog | 230.3934174 |
| LOC107071466 | transmembrane protein 120 homolog | 230.6468311 |
| LOC107064241 | uncharacterized protein LOC107064241 | 230.9441649 |

| | | |
|---|---|---|
| LOC107071193 | leucine-rich repeat-containing protein 24-like | 231.0085396 |
| LOC107067812 | probable low affinity copper uptake protein 2 | 231.112599 |
| LOC107064065 | zinc finger protein 408-like | 231.8656817 |
| LOC107063633 | chromatin modification-related protein EAF1-like | 232.7889461 |
| LOC107064325 | probable tyrosyl-DNA phosphodiesterase | 232.7924744 |
| LOC107066963 | cytosolic carboxypeptidase 6 | 232.8293962 |
| LOC107066018 | nuclear hormone receptor E75 | 233.6039525 |
| LOC107069081 | homeobox protein Nkx-2.2a-like | 234.0547581 |
| LOC107074352 | protein FAM76A | 234.1422083 |
| LOC107070747 | uncharacterized protein LOC107070747 | 234.4310344 |
| LOC107070925 | myosin light chain kinase, smooth muscle-like | 235.9601273 |
| LOC107067589 | protein-tyrosine sulfotransferase | 236.4870349 |
| LOC107067353 | uncharacterized protein LOC107067353 | 237.2001536 |
| LOC107072774 | intraflagellar transport protein 22 homolog | 237.204386 |
| LOC107068543 | E3 ubiquitin-protein ligase RNF126 | 237.4228027 |
| LOC107070292 | uncharacterized protein LOC107070292 | 237.4603468 |
| LOC107067223 | protein singed wings 2 | 237.6015996 |
| LOC107065545 | uncharacterized protein LOC107065545 | 237.6236317 |
| LOC107070177 | diphosphoinositol polyphosphate phosphohydrolase 1 | 238.0491804 |
| LOC107072470 | FUN14 domain-containing protein 1 | 238.1002926 |
| LOC107071199 | uncharacterized protein LOC107071199 | 238.7921346 |
| LOC107070470 | dedicator of cytokinesis protein 3 | 239.0494009 |
| LOC107063754 | uncharacterized protein LOC107063754 | 239.361095 |
| LOC107071696 | synapsin | 240.5957425 |
| LOC107066788 | gustatory receptor 68a-like | 240.9968374 |
| LOC107070435 | RING finger protein 44 | 241.1025335 |
| LOC107070460 | uncharacterized protein LOC107070460 | 241.2095407 |
| LOC107066640 | zinc/cadmium resistance protein | 241.7087017 |
| LOC107074333 | DNA-directed RNA polymerase II subunit RPB3 | 241.8753694 |
| LOC107065556 | potassium/sodium hyperpolarization-activated cyclic nucleot | 242.056483 |
| LOC107065216 | protein expanded | 242.0646246 |
| LOC107065882 | RNA-binding protein Musashi homolog Rbp6 | 242.0762963 |
| LOC107074314 | hexosaminidase D-like | 242.9728263 |
| LOC107072481 | 26S protease regulatory subunit 4 | 243.4604162 |
| LOC107073556 | uncharacterized protein LOC107073556 | 243.5122 |
| LOC107073578 | uncharacterized protein LOC107073578 | 243.6368653 |
| LOC107068189 | monocarboxylate transporter 3-like | 244.4762453 |
| LOC107074073 | hypoxia-inducible factor prolyl hydroxylase | 245.7281242 |
| LOC107064656 | protein Asterix | 246.5805721 |
| LOC107070930 | NADH-cytochrome b5 reductase 3 | 247.1108126 |
| LOC107064222 | alanine--glyoxylate aminotransferase 2 homolog 1, mitochon | 247.2035925 |
| LOC107065641 | sterile alpha and TIR motif-containing protein 1 | 248.1144221 |
| LOC107069610 | probable JmjC domain-containing histone demethylation pro | 248.2775637 |
| LOC107071668 | cytoplasmic polyadenylation element-binding protein 2 | 248.3976519 |
| LOC107066966 | histone H2B-like | 249.4492142 |
| LOC107063836 | neurotrimin-like | 249.5927643 |
| LOC107069369 | uncharacterized protein LOC107069369 | 250.6583414 |
| LOC107069026 | uncharacterized protein LOC107069026 | 250.8380529 |
| LOC107067025 | uncharacterized protein LOC107067025 | 251.136797 |
| LOC107064179 | m7GpppN-mRNA hydrolase | 251.3858521 |

| | | |
|---|---|---|
| LOC107067978 | zeta-sarcoglycan | 252.022446 |
| LOC107063792 | dyslexia susceptibility 1 candidate gene 1 protein homolog | 252.2170258 |
| LOC107067287 | sperm flagellar protein 1-like | 252.6710708 |
| LOC107072539 | LIM/homeobox protein Awh-like | 253.0215196 |
| LOC107068154 | carnosine N-methyltransferase | 253.2208301 |
| LOC107066690 | protein abrupt | 254.4726929 |
| LOC107064302 | Krueppel homolog 1-like | 255.5045401 |
| LOC107064461 | probable serine/threonine-protein kinase roco4 | 256.0173644 |
| LOC107066711 | fas apoptotic inhibitory molecule 1 | 256.682369 |
| LOC107073288 | growth hormone-regulated TBC protein 1-A | 256.9995563 |
| LOC107067026 | uncharacterized protein LOC107067026 | 257.0161337 |
| LOC107073831 | myocyte-specific enhancer factor 2 | 257.8769675 |
| LOC107065922 | uncharacterized protein LOC107065922 | 258.0385053 |
| LOC107068008 | cell cycle checkpoint protein RAD1 | 258.1997755 |
| LOC107073180 | homeobox protein unplugged | 258.6583308 |
| LOC107066497 | max-interacting protein 1-like | 259.4726077 |
| LOC107066843 | calcium channel flower | 260.1219185 |
| LOC107067280 | beta-mannosidase | 260.8235726 |
| LOC107064555 | RNA polymerase II degradation factor 1 | 261.3299704 |
| LOC107071044 | uncharacterized protein LOC107071044 | 261.6013788 |
| LOC107071188 | carbonic anhydrase-related protein 10 | 261.9603339 |
| LOC107066179 | iron-sulfur cluster assembly scaffold protein IscU | 262.139496 |
| LOC107073568 | protein FAM46C | 262.2860821 |
| LOC107064537 | uncharacterized protein LOC107064537 | 262.5448292 |
| LOC107070438 | uncharacterized protein LOC107070438 | 262.8346173 |
| LOC107064660 | probable GDP-L-fucose synthase | 262.8408564 |
| LOC107068066 | RNA polymerase II-associated factor 1 homolog | 263.0343234 |
| LOC107064221 | uncharacterized protein LOC107064221 | 265.1242839 |
| LOC107067024 | uncharacterized protein LOC107067024 | 265.7629756 |
| LOC107068655 | aryl hydrocarbon receptor nuclear translocator homolog | 265.9601585 |
| LOC107064220 | uncharacterized protein LOC107064220 | 267.3014859 |
| LOC107073185 | BTB/POZ domain-containing protein 6-like | 268.4739753 |
| LOC107073413 | DNA polymerase epsilon subunit C | 268.9466066 |
| LOC107070844 | uncharacterized protein LOC107070844 | 268.9886281 |
| LOC107072003 | tribbles homolog 2 | 269.5348635 |
| LOC107072308 | chaoptin | 269.9853985 |
| LOC107072852 | pyrroline-5-carboxylate reductase | 273.3654913 |
| LOC107064838 | histone H2B-like | 273.6126741 |
| LOC107065727 | transmembrane protein 138 | 274.8662902 |
| LOC107064297 | RNA-binding protein lark | 275.5186673 |
| LOC107068718 | uncharacterized protein LOC107068718 | 276.2321288 |
| LOC107071902 | multivesicular body subunit 12A | 276.4446477 |
| LOC107073544 | uncharacterized protein LOC107073544 | 276.6530713 |
| LOC107066611 | ubiquitin-conjugating enzyme E2 G2 | 277.2965614 |
| LOC107074399 | selenide, water dikinase | 280.3724769 |
| LOC107065297 | protein bric-a-brac 2-like | 280.4389661 |
| LOC107073995 | proteasome subunit alpha type-6-like | 280.8428815 |
| LOC107068481 | glutamine--fructose-6-phosphate aminotransferase [isomeriz | 281.0266156 |
| LOC107072753 | zinc finger protein ubi-d4 A | 283.4433483 |
| LOC107064369 | muskelin | 286.0582171 |

| | | |
|---|---|---|
| LOC107066629 | DCN1-like protein 5 | 286.3233051 |
| LOC107067028 | histone H4-like | 287.0439799 |
| LOC107070046 | serine-aspartate repeat-containing protein I-like | 288.2463326 |
| LOC107073989 | ubiquitin carboxyl-terminal hydrolase isozyme L3 | 290.8109839 |
| LOC107073793 | extensin-like | 290.9651537 |
| LOC107067703 | ubiquitin-like protein 7 | 291.1283381 |
| LOC107074148 | uncharacterized protein LOC107074148 | 291.5130186 |
| LOC107073993 | cleavage stimulation factor subunit 1 | 292.2710658 |
| LOC107069780 | sodium channel protein Nach-like | 293.3815496 |
| LOC107063916 | acyl-CoA Delta(11) desaturase-like | 293.6172668 |
| LOC107074667 | cell division cycle protein 16 homolog | 294.8543606 |
| LOC107067286 | coiled-coil domain-containing protein 103 | 295.2287126 |
| LOC107064159 | serine/threonine-protein phosphatase 4 catalytic subunit | 295.4132318 |
| LOC107064343 | oocyte zinc finger protein XlCOF22 | 296.2795188 |
| LOC107070789 | uncharacterized protein LOC107070789 | 298.4835184 |
| LOC107073333 | uncharacterized protein LOC107073333 | 301.2328325 |
| LOC107066480 | uncharacterized protein LOC107066480 | 301.4112364 |
| LOC107068886 | uncharacterized protein LOC107068886 | 301.6870867 |
| LOC107067027 | histone H4 | 301.8051475 |
| LOC107071280 | zinc finger protein Noc | 303.8295012 |
| LOC107067446 | uncharacterized protein LOC107067446 | 305.2987913 |
| LOC107068311 | putative protein kinase C delta type homolog | 305.4462319 |
| LOC107064973 | protein phosphatase 1B | 305.9102488 |
| LOC107074492 | G/T mismatch-specific thymine DNA glycosylase-like | 306.2999023 |
| LOC107066998 | histone H2A | 306.6493154 |
| LOC107065493 | 4-hydroxyphenylpyruvate dioxygenase | 308.7436853 |
| LOC107065336 | transmembrane protein 43 homolog | 308.7515963 |
| LOC107073660 | uncharacterized protein LOC107073660 | 314.0385085 |
| LOC107064442 | histone H3 | 314.2778927 |
| LOC107073681 | fragile X mental retardation syndrome-related protein 2-like | 314.8950762 |
| LOC107073753 | piggyBac transposable element-derived protein 4-like | 315.1729915 |
| LOC107065263 | histone H3 | 317.5004488 |
| LOC107070787 | uncharacterized protein LOC107070787 | 322.1509332 |
| LOC107067965 | COP9 signalosome complex subunit 6 | 323.1902651 |
| LOC107064451 | histone H4 | 336.7151952 |
| LOC107064611 | forkhead box protein K1 | 337.3023562 |
| LOC107064914 | nuclear envelope phosphatase-regulatory subunit 1 | 339.9667725 |
| LOC107067555 | WW domain-binding protein 2 | 341.1954276 |
| LOC107073951 | cleavage and polyadenylation specificity factor subunit 4 | 342.6782727 |
| LOC107073611 | uncharacterized protein LOC107073611 | 346.474736 |
| LOC107064248 | ATP-dependent RNA helicase DDX42 | 350.5700524 |
| LOC107064462 | histone H4 | 351.1580757 |
| LOC107067260 | transient receptor potential cation channel subfamily V mem | 353.3003793 |
| LOC107073714 | uncharacterized protein LOC107073714 | 371.0793583 |
| LOC107067723 | toll-like receptor 13 | 399.0640142 |
| LOC107073715 | uncharacterized protein LOC107073715 | 403.2239028 |
| LOC107073769 | uncharacterized protein LOC107073769 | 409.2356781 |
