## Supplementary Table S3 for "The molecular basis of socially-mediated phenotypic plasticity in a eusocial paper wasp"

**Table S3.** Genes identified as differentially expressed between control workers and queens by DESeq2 at adjusted p<0.05. **BaseMean**: Mean expression of the gene in the reference group (queens). **Log2LFC**: Log(2) fold change of the gene in the comparison group (workers) compared to the reference group. **LFCSE**: Standard error of the log fold-change. **Stat**: Wald test statistic for the given comparison. **pvalue**: p-value for the given test statistic. **padjusted**: Adjusted p-values generated using Benjamini-Hochberg correction with alpha = 0.05.

| GeneID | ProteinID | BaseMean | Log2FC | LFCSE | Stat | pvalue | padjusted |
|---|---|---|---|---|---|---|---|
| LOC107064940 | uncharacterize( | 374.2368 | 1.5891 | 0.2751 | 3.6505 | 0.00026174 | 0.03676 |
| LOC107067122 | uncharacterize( | 30.3036 | -2.1856 | 0.3009 | -5.3201 | 0.00000010 | 0.00003 |
| LOC107072670 | uncharacterize( | 15.4322 | 3.8287 | 0.6794 | 4.7741 | 0.00000181 | 0.00046 |
| LOC107071493 | uncharacterize( | 338.9425 | -1.4165 | 0.2239 | -3.7136 | 0.00020433 | 0.03133 |
| LOC107068601 | cytochrome P4. | 969.1238 | -6.1858 | 0.9103 | -6.1529 | 0.00000000 | 0.00000 |
| LOC107073714 | uncharacterize( | 37.5979 | -2.5317 | 0.3543 | -5.4952 | 0.00000004 | 0.00001 |
| LOC107073715 | uncharacterize( | 206.8934 | -4.9115 | 0.2028 | -21.3352 | 0.00000000 | 0.00000 |
| LOC107071795 | acyl-CoA Delta( | 7.5613 | -3.7137 | 0.8460 | -3.6985 | 0.00021689 | 0.03279 |
| LOC107071668 | cytoplasmic pol | 347.8662 | -1.2798 | 0.1721 | -4.0384 | 0.00005381 | 0.00963 |
| LOC107070237 | sensory neuron | 53.5613 | 2.6788 | 0.4487 | 4.6662 | 0.00000307 | 0.00072 |
| LOC107071136 | dentin sialopho | 705.2829 | -1.9629 | 0.3211 | -4.2907 | 0.00001781 | 0.00390 |
| LOC107066914 | poly(rC)-bindin | 16364.4075 | -1.2360 | 0.1574 | -4.1369 | 0.00003520 | 0.00672 |
| LOC107066914 | poly(rC)-bindin | 16364.4075 | -1.2360 | 0.1574 | -4.1369 | 0.00003520 | 0.00672 |
| LOC107063865 | uncharacterize( | 3.8797 | -5.2030 | 1.1629 | -3.9710 | 0.00007157 | 0.01219 |
| LOC107068616 | ejaculatory bull | 2435.2533 | -2.9785 | 0.5056 | -4.7341 | 0.00000220 | 0.00054 |
| LOC107072906 | elongation of v | 405.9568 | -2.3679 | 0.4154 | -4.2916 | 0.00001774 | 0.00390 |
| LOC107074300 | four and a half | 20206.1106 | -1.6274 | 0.2736 | -3.8101 | 0.00013893 | 0.02226 |
| LOC107074300 | four and a half | 20206.1106 | -1.6274 | 0.2736 | -3.8101 | 0.00013893 | 0.02226 |
| LOC107074300 | prickle-like prot | 20206.1106 | -1.6274 | 0.2736 | -3.8101 | 0.00013893 | 0.02226 |
| LOC107074300 | GDNF-inducible | 20206.1106 | -1.6274 | 0.2736 | -3.8101 | 0.00013893 | 0.02226 |
| LOC107074300 | zinc finger and | 20206.1106 | -1.6274 | 0.2736 | -3.8101 | 0.00013893 | 0.02226 |
| LOC107074300 | zinc finger prot | 20206.1106 | -1.6274 | 0.2736 | -3.8101 | 0.00013893 | 0.02226 |
| LOC107066966 | histone H2B-lik | 50.8434 | -2.7566 | 0.3134 | -6.9302 | 0.00000000 | 0.00000 |
| LOC107069298 | myrosinase 1-li | 21.6950 | -3.4816 | 0.5891 | -4.9167 | 0.00000088 | 0.00023 |
| LOC107067286 | coiled-coil dom | 119.6298 | -2.1036 | 0.1994 | -7.6171 | 0.00000000 | 0.00000 |
| LOC107067260 | transient recep | 600.8711 | -2.7584 | 0.2039 | -10.6612 | 0.00000000 | 0.00000 |
| LOC107074200 | 26S proteasom | 3036.3962 | 2.8011 | 0.3067 | 7.2247 | 0.00000000 | 0.00000 |
| LOC107064462 | histone H4 | 93.1582 | -3.6095 | 0.3174 | -9.5286 | 0.00000000 | 0.00000 |
| LOC107063792 | dyslexia suscep | 162.7474 | -1.4118 | 0.1925 | -4.2958 | 0.00001741 | 0.00390 |
| LOC107070407 | putative fatty a | 760.1575 | -2.3962 | 0.4382 | -4.1338 | 0.00003568 | 0.00672 |
| LOC107073611 | uncharacterize( | 2053.1830 | -1.6941 | 0.1462 | -7.5869 | 0.00000000 | 0.00000 |
| LOC107064519 | uncharacterize( | 8.4806 | -3.2945 | 0.6737 | -4.0219 | 0.00005774 | 0.01016 |
| LOC107065267 | uncharacterize( | 64.5832 | 2.9445 | 0.6441 | 3.6632 | 0.00024908 | 0.03613 |
| LOC107063633 | chromatin mod | 341.5356 | -1.3411 | 0.2083 | -3.6308 | 0.00028259 | 0.03850 |
| LOC107067353 | uncharacterize( | 206.4963 | -3.6500 | 0.4943 | -6.2001 | 0.00000000 | 0.00000 |
| LOC107065128 | histone H2A-lik | 8.8175 | -2.0794 | 0.3574 | -4.1818 | 0.00002892 | 0.00599 |
| LOC107066497 | max-interacting | 944.0240 | -1.1042 | 0.1423 | -3.6486 | 0.00026369 | 0.03676 |
| LOC107067185 | arylphorin subu | 1598.7232 | 5.1313 | 0.7878 | 5.7711 | 0.00000001 | 0.00000 |
| LOC107070789 | uncharacterize( | 31.0700 | -1.6900 | 0.2219 | -4.9798 | 0.00000064 | 0.00018 |
| LOC107074148 | uncharacterize( | 65.7700 | -2.2637 | 0.2415 | -6.9514 | 0.00000000 | 0.00000 |
| LOC107073446 | fatty acyl-CoA r | 41.7800 | -3.3629 | 0.6850 | -4.0554 | 0.00005004 | 0.00910 |
| LOC107065294 | zinc carboxype | 7.3356 | 2.7068 | 0.5452 | 3.8922 | 0.00009933 | 0.01666 |

| | | | | | | | |
|---|---|---|---|---|---|---|---|
| LOC107073660 | uncharacterized | 50.2558 | -2.5488 | 0.2242 | -8.7597 | 0.00000000 | 0.00000 |
| LOC107068157 | histone H2A | 389.3246 | 0.9992 | 0.1000 | 4.1415 | 0.00003451 | 0.00672 |
| LOC107072308 | chaoptin | 52.1370 | -1.6615 | 0.2066 | -5.2108 | 0.00000019 | 0.00005 |
| LOC107065361 | cytochrome P4 | 167.0913 | -4.0527 | 0.5720 | -6.0623 | 0.00000000 | 0.00000 |
| LOC107067446 | uncharacterized | 376.7458 | -1.1026 | 0.1368 | -3.7830 | 0.00015494 | 0.02410 |
| LOC107072279 | stress response | 12383.2209 | -1.6757 | 0.3025 | -3.6062 | 0.00031067 | 0.04168 |
| LOC107073769 | uncharacterized | 526.4029 | -1.9862 | 0.2018 | -6.9431 | 0.00000000 | 0.00000 |
| LOC107063829 | defensin-1-like | 42.6869 | -4.2235 | 0.8752 | -4.1572 | 0.00003221 | 0.00644 |
| LOC107064241 | uncharacterized | 138.6684 | -4.1137 | 0.5596 | -6.3055 | 0.00000000 | 0.00000 |
| LOC107064220 | uncharacterized | 117.9459 | -6.3221 | 0.5360 | -10.7032 | 0.00000000 | 0.00000 |
| LOC107065359 | cytochrome P4 | 138.7041 | -3.5351 | 0.5616 | -5.2534 | 0.00000015 | 0.00004 |
| LOC107071066 | elongation of v | 39.8942 | -3.2936 | 0.5060 | -5.3535 | 0.00000009 | 0.00003 |
| LOC107065297 | protein bric-a-b | 405.7567 | -1.4613 | 0.1764 | -4.9687 | 0.00000067 | 0.00018 |
| LOC107067158 | RNA polymeras | 10.5154 | 2.6992 | 0.5194 | 4.0708 | 0.00004686 | 0.00867 |
| LOC107070680 | uncharacterized | 142.1665 | -3.9859 | 0.6023 | -5.6466 | 0.00000002 | 0.00001 |
| LOC107067009 | uncharacterized | 709.0995 | 2.6239 | 0.5258 | 3.8774 | 0.00010559 | 0.01744 |
| LOC107067009 | proteoglycan 4 | 709.0995 | 2.6239 | 0.5258 | 3.8774 | 0.00010559 | 0.01744 |
| LOC107067009 | sialidase-like | 709.0995 | 2.6239 | 0.5258 | 3.8774 | 0.00010559 | 0.01744 |
| LOC107064302 | Krueppel homo | 809.4749 | -1.1795 | 0.1422 | -4.1811 | 0.00002900 | 0.00599 |
| LOC107064264 | MOXD1 homolo | 78.6771 | 1.4672 | 0.2416 | 3.6520 | 0.00026025 | 0.03676 |
| LOC107073568 | protein FAM46 | 296.2237 | -1.0600 | 0.1287 | -3.6924 | 0.00022214 | 0.03312 |
| LOC107064035 | uncharacterized | 24.7265 | 1.4823 | 0.2503 | 3.5856 | 0.00033633 | 0.04457 |
| LOC107069780 | sodium channe | 113.8775 | -2.1569 | 0.2159 | -7.2814 | 0.00000000 | 0.00000 |
| LOC107069782 | putative media | 363.4498 | -1.4742 | 0.2227 | -3.9926 | 0.00006536 | 0.01132 |
| LOC107067335 | uncharacterized | 20.3818 | -2.3476 | 0.4626 | -3.8104 | 0.00013875 | 0.02226 |
| LOC107067188 | arylphorin subu | 11.3921 | 3.5677 | 0.7043 | 4.2352 | 0.00002284 | 0.00490 |
| LOC107067189 | arylphorin subu | 8.3786 | 4.5050 | 0.8328 | 4.7068 | 0.00000252 | 0.00060 |
| LOC107070787 | uncharacterized | 582.8461 | -2.2337 | 0.2016 | -8.1762 | 0.00000000 | 0.00000 |
| LOC107069358 | uncharacterized | 33.5679 | -2.7348 | 0.3218 | -6.6810 | 0.00000000 | 0.00000 |
| LOC107068888 | odorant recept | 9.4441 | 1.5725 | 0.2607 | 3.7876 | 0.00015211 | 0.02401 |
| LOC107072164 | putative media | 3232.3273 | -1.5367 | 0.2622 | -3.6300 | 0.00028339 | 0.03850 |
| LOC107065578 | toll-like recepto | 522.2755 | -2.1924 | 0.4384 | -3.6665 | 0.00024589 | 0.03613 |
| LOC107067026 | uncharacterized | 83.9264 | -1.8983 | 0.2417 | -5.4344 | 0.00000005 | 0.00002 |
| LOC107067028 | histone H4-like | 29.3521 | -2.1186 | 0.2757 | -5.5624 | 0.00000003 | 0.00001 |
| LOC107066998 | histone H2A | 173.0184 | -3.7910 | 0.3269 | -9.8079 | 0.00000000 | 0.00000 |
| LOC107070047 | uncharacterized | 39.1535 | 1.8565 | 0.3060 | 4.1558 | 0.00003241 | 0.00644 |
| LOC107065263 | histone H3 | 35.5913 | -3.3476 | 0.3005 | -9.1925 | 0.00000000 | 0.00000 |
| LOC107064451 | histone H4 | 215.0863 | -3.8234 | 0.2966 | -10.9199 | 0.00000000 | 0.00000 |
| LOC107064442 | histone H3 | 46.5586 | -3.1472 | 0.2880 | -8.8955 | 0.00000000 | 0.00000 |
| LOC107064343 | oocyte zinc fing | 611.6337 | -5.9602 | 0.4413 | -12.1802 | 0.00000000 | 0.00000 |
| LOC107064221 | uncharacterized | 155.2180 | -4.8412 | 0.4578 | -9.2968 | 0.00000000 | 0.00000 |
| LOC107064222 | alanine--glyoxy | 49.9772 | -3.9623 | 0.4530 | -7.4556 | 0.00000000 | 0.00000 |
| LOC107067027 | histone H4 | 363.7658 | -2.3225 | 0.2523 | -6.8864 | 0.00000000 | 0.00000 |
| LOC107067024 | uncharacterized | 104.3222 | -2.4935 | 0.2744 | -6.9543 | 0.00000000 | 0.00000 |
| LOC107067025 | uncharacterized | 203.5795 | -1.8750 | 0.2502 | -5.1552 | 0.00000025 | 0.00007 |
| LOC107068501 | ejaculatory bull | 408.9823 | 4.3460 | 0.7920 | 4.7488 | 0.00000205 | 0.00051 |
| LOC107064838 | histone H2B-lik | 28.7526 | -2.3868 | 0.3214 | -5.6060 | 0.00000002 | 0.00001 |
