## Supplementary Table S4 for "The molecular basis of socially-mediated phenotypic plasticity in a eusocial paper wasp"

**Table S4.** Genes identified as differentially expressed between control workers and queen removal individuals by DESeq2 at adjusted p<0.05. **BaseMean**: Mean expression of the gene in the reference group (queens). **Log2LFC**: Log(2) fold change of the gene in the comparison group (workers) compared to the reference group. **LFCSE**: Standard error of the log fold-change. **Stat**: Wald test statistic for the given comparison. **pvalue**: p-value for the given test statistic. **padjusted**: Adjusted p-values generated using Benjamini-Hochberg correction with alpha = 0.05.

| GeneID | ProteinID | BaseMean | Log2FC | LFCSE | Stat | pvalue | padjusted |
|---|---|---|---|---|---|---|---|
| LOC107074674 | circumsporozoite protein | 1699.8702 | 6.2215 | 0.9414 | 5.9874 | 0.00000000 | 0.00002289 |
| LOC107063902 | glucose dehydrogenase [I | 2450.5809 | 6.5242 | 1.0223 | 5.8099 | 0.00000001 | 0.00003354 |
| LOC107067185 | arylphorin subunit beta-li | 1426.6354 | -3.2743 | 0.6266 | -4.2923 | 0.00001768 | 0.03796339 |
| LOC107064220 | uncharacterized protein L | 39.9903 | 4.2777 | 0.8146 | 4.5331 | 0.00000581 | 0.01559900 |
| LOC107064343 | oocyte zinc finger protein | 239.0962 | 4.1107 | 0.7227 | 4.8785 | 0.00000107 | 0.00382490 |
