## Supplementary Figure S1 for "The molecular basis of socially-mediated phenotypic plasticity in a eusocial paper wasp"

**Supplementary figure S1.** (A) Root mean squared three-fold cross-validation error of SVMs trained using an iteratively decreasing number of genes. In each iteration, the lowest-weight gene of the previous model was removed and a new model was trained with the remaining genes. For each model, three-fold cross validation error was taken as the mean of 20 calculations using randomly-selected validation bins. Figure shows a moving average with a window size of 50. Red dashed line shows the minimum validation error achieved. (B) Significantly enriched gene ontology terms at p<0.01 among 1992 caste-informative genes identified by SVM classification. Ontologies: BP = biological process; CC = cellular component; MF = molecular function. (C) Absolute feature weights of the 1992 genes in the optimised SVM. 81 genes identified by DESeq2 as differentially expressed between queens and control workers are marked in blue. (D-F) Distribution of phenotypic traits of sequenced individuals from queen removal colonies. Where possible, individuals were selected to represent as wide as possible a range of values for (D) Ovarian development; (E) Dominance; and (F) Phenotypic caste identity ('queenness'). Full methodology for the calculation of phenotypic indices is given in Taylor et al (2020). (G) First two principal components generated by PC analysis of 1992 caste-associated genes identified by SVM classification.

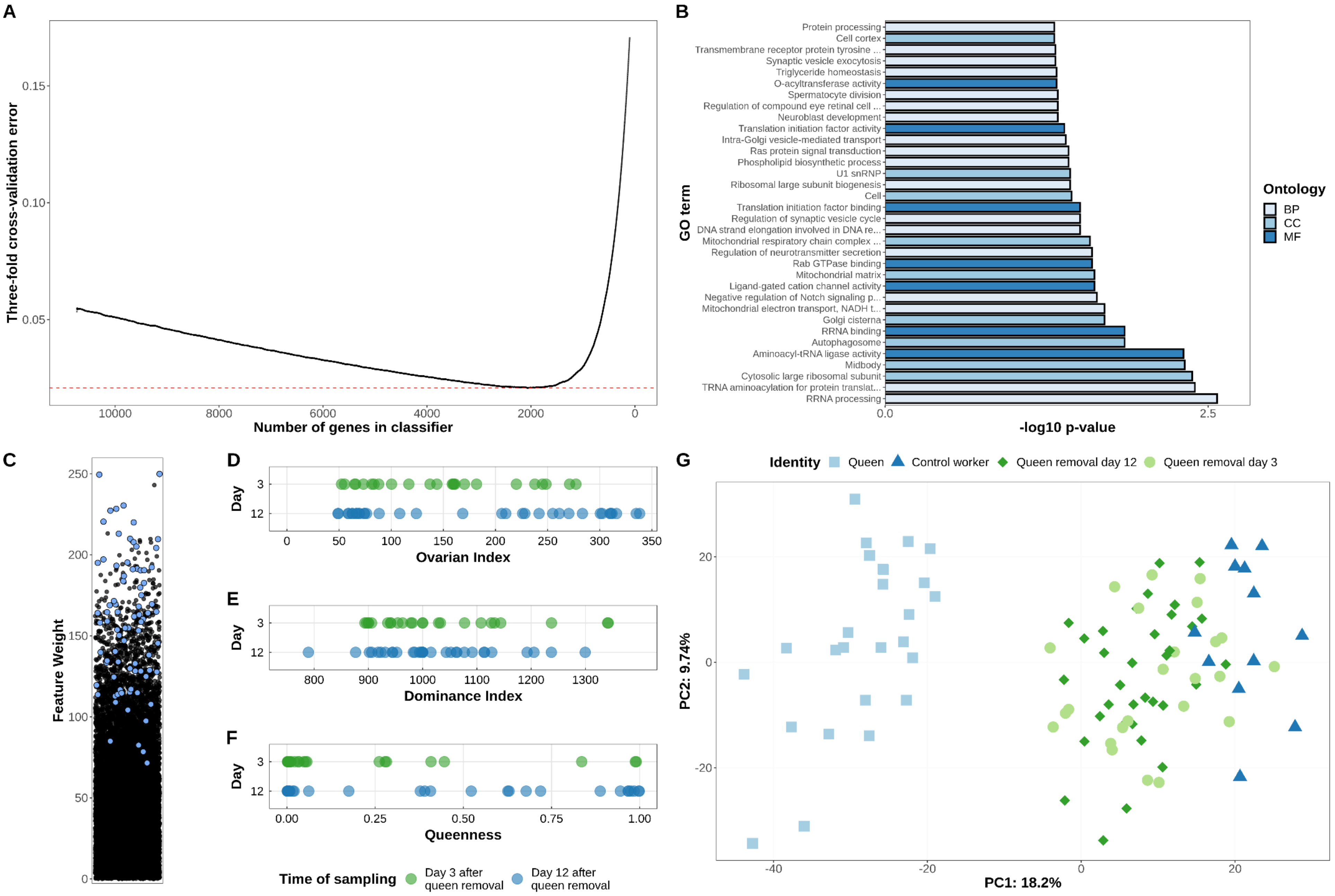
