## Supplementary Phenotypic Data for "The molecular basis of socially-mediated phenotypic plasticity in a eusocial paper wasp"

**Phenotypic data pertaining to samples analysed in the study. Treatment:** The treatment group to which the individual was assigned. C = sham removal; QR = queen removal; 3 = three-day observation following manipulation; 12 = twelve-day observation following manipulation. **Role:** Whether individual was originally a foundress queen or a worker. **Age:** Age in days at time of sampling. **Ovaries:** Ovarian index at time of sampling, measured following Cini et al (2013). **Elo:** Elo rating for the individual based on intragroup dominance interactions in the three days prior to sampling, as described in Taylor et al (2020). **Queenness:** For queen removal individuals, caste identity at the time of sampling, measured as described in Taylor et al (2020). **NestID:** Nest from which the individual was collected.

| ID | TreatmentGroup | Role | Age | Ovaries | Elo | Queenness | NestID |
|---|---|---|---|---|---|---|---|
| F01RD | C12 | queen | NA | 321.24 | 1000 | NA | 1 |
| F03BL | QR12 | queen | NA | 309.33 | 1000 | NA | 3 |
| F06BL | C12 | queen | NA | 311.4543 | 1055.546 | NA | 6 |
| F08YL | C3 | queen | NA | 258.2917 | 1215.259 | NA | 8 |
| F09BL | C3 | queen | NA | 329.9887 | 1141.21 | NA | 9 |
| F13GR | C12 | queen | NA | 275.229 | 1000 | NA | 13 |
| F14GR | QR12 | queen | NA | 315.33 | 1087.754 | NA | 14 |
| F16BL | C12 | queen | NA | 298.479 | 1121.08 | NA | 16 |
| F28GRGR | C3 | queen | NA | 338.4463 | 1226.191 | NA | 28 |
| F31GR | C3 | queen | NA | 237.1933 | 1083.965 | NA | 31 |
| F38BL | QR3 | queen | NA | 225.1998 | 1000 | NA | 38 |
| F42OR | QR12 | queen | NA | 319.78 | 1139.232 | NA | 42 |
| F45OR | QR12 | queen | NA | 337.8 | 1093.033 | NA | 45 |
| F47RD | QR3 | queen | NA | 307.5175 | 1135.236 | NA | 47 |
| F48SL | QR12 | queen | NA | 327.45 | 1224.578 | NA | 48 |
| F49RD | C3 | queen | NA | 287.5078 | 1191.317 | NA | 49 |
| F51RD | QR3 | queen | NA | 305.82 | 1135.831 | NA | 51 |
| F54GR | C3 | queen | NA | 311.8055 | 1000 | NA | 54 |
| F56YL | C3 | queen | NA | 281.8258 | 1056.417 | NA | 56 |
| F58SL | C12 | queen | NA | 313.7658 | 1050 | NA | 58 |
| F65ORWH | QR12 | queen | NA | 354.2058 | 1332.34 | NA | 65 |
| F67YLBL | QR12 | queen | NA | 351.16 | 1190.331 | NA | 67 |
| F69OR | QR3 | queen | NA | 322.7108 | 1077.252 | NA | 69 |
| F70SL | C3 | queen | NA | 332.3168 | 1127.83 | NA | 70 |
| F71BL | QR3 | queen | NA | 332.32 | 1050 | NA | 71 |
| F72RDWH | C3 | queen | NA | 323.816 | 1134.372 | NA | 72 |
| W01RDYL | C12 | worker_ctı | 20 | 195.6292 | 1050 | NA | 1 |
| W03BLGR | QR12 | worker_qr | 11 | 124.1157 | 1000 | 0.0201716 | 3 |
| W03GR | QR12 | worker_qr | 23 | 66.76183 | 923.1059 | 0.0020961 | 3 |
| W03SL | QR12 | worker_qr | 25 | 316.0947 | 1076.894 | 0.9685319 | 3 |
| W04BLOR | QR12 | worker_qr | 24 | 66.43367 | 876.4723 | 0.0016821 | 4 |
| W04GRSL | QR12 | worker_qr | 24 | 209.8983 | 931.0553 | 0.1752899 | 4 |
| W04YLWH | QR12 | worker_qr | 24 | 302.3557 | 1192.472 | 0.984416 | 4 |
| W05RDRD | QR12 | worker_qr | 23 | 270.3795 | 944.3507 | 0.6260931 | 5 |
| W08RDOR | C3 | worker_ctı | 9 | 117.0052 | 950 | NA | 8 |
| W09RDOR | C3 | worker_ctı | 16 | 52.903 | 978.77 | NA | 9 |
| W13YLSL | C12 | worker_ctı | 23 | 93.32967 | 1000 | NA | 13 |
| W14GRRD | QR12 | worker_qr | 16 | 62.876 | 994.3113 | 0.0025354 | 14 |
| W14SLBL | QR12 | worker_qr | 25 | 309.8395 | 1043.475 | 0.945193 | 14 |

| | | | | | | | |
|---|---|---|---|---|---|---|---|
| W14WHOI | QR12 | worker_qr | 23 | 69.438 | 1000 | 0.0032421 | 14 |
| W28BLOR | QR3 | worker_qr | 3 | 100.303 | 1000 | 0.0091329 | 38 |
| W28ORYL | C3 | worker_ctı | 7 | 46.88967 | 966.0638 | NA | 28 |
| W33BLYL | QR12 | worker_qr | 24 | 241.8343 | 1000 | 0.5223633 | 33 |
| W33GRSL | QR12 | worker_qr | 18 | 108.127 | 1062.003 | 0.0166572 | 33 |
| W33RDBL | QR12 | worker_qr | 13 | 168.5412 | 952.3063 | 0.0617044 | 33 |
| W33SL | QR12 | worker_qr | 28 | 68.9615 | 1127.415 | 0.0056954 | 33 |
| W38RDBL | QR3 | worker_qr | 21 | 245.001 | 954.2636 | 0.4462304 | 38 |
| W38RDWI | QR3 | worker_qr | 24 | 237.6817 | 900.004 | 0.2822045 | 38 |
| W38YLBL | QR3 | worker_qr | 33 | 277.2408 | 1342.101 | 0.9904993 | 38 |
| W40GROR | C12 | worker_ctı | 16 | 171.622 | 1093.673 | NA | 40 |
| W42GRSL | QR12 | worker_qr | 24 | 311.8067 | 1114.157 | 0.9752185 | 42 |
| W42RDWI | QR12 | worker_qr | 19 | 261.2623 | 1000 | 0.6781197 | 42 |
| W42WHBL | QR12 | worker_qr | 17 | 226.0015 | 1000 | 0.3905995 | 42 |
| W42WHRI | QR12 | worker_qr | 17 | 283.3712 | 944.7295 | 0.7189861 | 42 |
| W42WHRI | QR12 | worker_qr | 14 | 300.5748 | 1000 | 0.888133 | 42 |
| W44GRBL | QR3 | worker_qr | 9 | 73.37083 | 1107.685 | 0.00611 | 44 |
| W45RDWI | QR12 | worker_qr | 11 | 58.695 | 997.2191 | 0.0022316 | 45 |
| W45SL | QR12 | worker_qr | 16 | 334.987 | 1237.241 | 0.9972978 | 45 |
| W45WHRI | QR12 | worker_qr | 3 | 76.433 | 982.9388 | 0.0037832 | 45 |
| W45YLBL | QR12 | worker_qr | 14 | 59.21017 | 918.0188 | 0.0016135 | 45 |
| W46ORYL | C12 | worker_ctı | 1 | 84.818 | 992.3 | NA | 46 |
| W47GRBL | QR3 | worker_qr | 28 | 143.8445 | 978.401 | 0.0336721 | 47 |
| W47RDWI | QR3 | worker_qr | 21 | 158.5257 | 900.2997 | 0.0322066 | 47 |
| W47WHBL | QR3 | worker_qr | 30 | 271.0488 | 1340.866 | 0.9875367 | 47 |
| W47WHYL | QR3 | worker_qr | 26 | 170.2187 | 905.909 | 0.0475288 | 47 |
| W47YLBL | QR3 | worker_qr | 26 | 220.0398 | 1032.809 | 0.4083222 | 47 |
| W48GRBL2 | QR12 | worker_qr | 20 | 254.9603 | 901.9123 | 0.4069915 | 48 |
| W48RDWI | QR12 | worker_qr | 26 | 310.5328 | 1091.374 | 0.9671629 | 48 |
| W48YLWH | QR12 | worker_qr | 16 | 88.39633 | 1062.981 | 0.0084098 | 48 |
| W49RDYL | C3 | worker_ctı | 10 | 69.87567 | 973.6778 | NA | 49 |
| W51BL | QR3 | worker_qr | 13 | 160.0855 | 1237.459 | 0.2615344 | 51 |
| W51GRYL | QR3 | worker_qr | 11 | 83.47333 | 1029.384 | 0.0060019 | 51 |
| W51ORBL | QR3 | worker_qr | 9 | 55.30983 | 893.5864 | 0.0012866 | 51 |
| W51SLWH | QR3 | worker_qr | 9 | 161.1707 | 963.1064 | 0.0527963 | 51 |
| W52BLGR | QR3 | worker_qr | 12 | 52.3265 | 1126.494 | 0.0030758 | 52 |
| W54ORYL | C3 | worker_ctı | 3 | 61.24167 | 1000 | NA | 54 |
| W56YLBL | C3 | worker_ctı | 2 | 66.714 | 1000 | NA | 56 |
| W57BLOR | QR3 | worker_qr | 21 | 81.44583 | 1077.237 | 0.0070622 | 57 |
| W57RDYL | QR3 | worker_qr | 4 | 159.6192 | 980.9025 | 0.0564687 | 57 |
| W58BLWH | C12 | worker_ctı | 13 | 59.0565 | 1000 | NA | 58 |
| W65BLWH | QR12 | worker_qr | 12 | 338.3238 | 1299.603 | 0.9988337 | 65 |
| W65ORBL | QR12 | worker_qr | 8 | 49.022 | 1015.273 | 0.0017347 | 65 |
| W65RDGR | QR12 | worker_qr | 19 | 228.1342 | 985.476 | 0.3781139 | 65 |
| W65RDOR | QR12 | worker_qr | 21 | 74.43683 | 1052.003 | 0.0048938 | 65 |
| W67BLWH | QR12 | worker_qr | 17 | 74.00483 | 1113.127 | 0.0064127 | 67 |
| W67GROR | QR12 | worker_qr | 22 | 206.0752 | 1205.221 | 0.6297013 | 67 |
| W67RDBL | QR12 | worker_qr | 12 | 49.15117 | 789.5007 | 0.0006963 | 67 |
| W67WHBL | QR12 | worker_qr | 18 | 62.571 | 906.2553 | 0.001704 | 67 |
| W69BLOR | QR3 | worker_qr | 3 | 116.8372 | 935.7363 | 0.0109871 | 69 |

| | | | | | | | |
|---|---|---|---|---|---|---|---|
| W69BLYL | QR3 | worker_qr: | 7 | 137.1695 | 941.105 | 0.0216033 | 69 |
| W69SLBL | QR3 | worker_qr: | 3 | 65.88783 | 1000 | 0.0028771 | 69 |
| W69YLSL | QR3 | worker_qr: | 11 | 181.7857 | 1132.562 | 0.2781554 | 69 |
| W70RDYL | C3 | worker_ctr | 17 | 79.93483 | 1007.52 | NA | 70 |
| W71RDBL | QR3 | worker_qr: | 17 | 64.871 | 941.1492 | 0.0021393 | 71 |
| W71YLWH | QR3 | worker_qr: | 17 | 248.544 | 1143.861 | 0.8355728 | 71 |
| W71YLYL | QR3 | worker_qr: | 13 | 88.029 | 896.7783 | 0.0036039 | 71 |
